# The host genotype actively shapes its microbiome across generations in laboratory mice

**DOI:** 10.1101/2024.03.14.584950

**Authors:** Laurentiu Benga, Anna Rehm, Christina Gougoula, Philipp Westhoff, Thorsten Wachtmeister, W. Peter M. Benten, Eva Engelhardt, Andreas P.M. Weber, Karl Köhrer, Martin Sager, Stefan Janssen

**Author notes:** Correspondence: Laurentiu Benga Stefan Janssen. contributed equally to this work (Laurentiu Benga, Anna Rehm).

## Abstract

**Background:** The microbiome greatly affects health and wellbeing. Evolutionarily, it is doubtful that a host would rely on chance alone to pass on microbial colonization to its offspring. However, the literature currently offers only limited evidence regarding two alternative hypotheses: active microbial shaping by host genetic factors or transmission of a microbial maternal legacy.

**Results:** To further dissect the influence of host genetics and maternal inheritance, we collected 2-cell stage embryos from two representative wildtypes, C57BL6/J and BALB/c, and transferred a mixture of both genotype embryos into hybrid recipient mice to be inoculated by an identical microbiome at birth.

**Conclusions:** Observing the offspring for six generations unequivocally emphasizes the impact of host genetic factors over maternal legacy in constant environments, akin to murine laboratory experiments. Interestingly, maternal legacy solely controlled the microbiome in the first offspring generation. However, current evidence supporting maternal legacy has not extended beyond this initial generation, resolving the aforementioned debate.

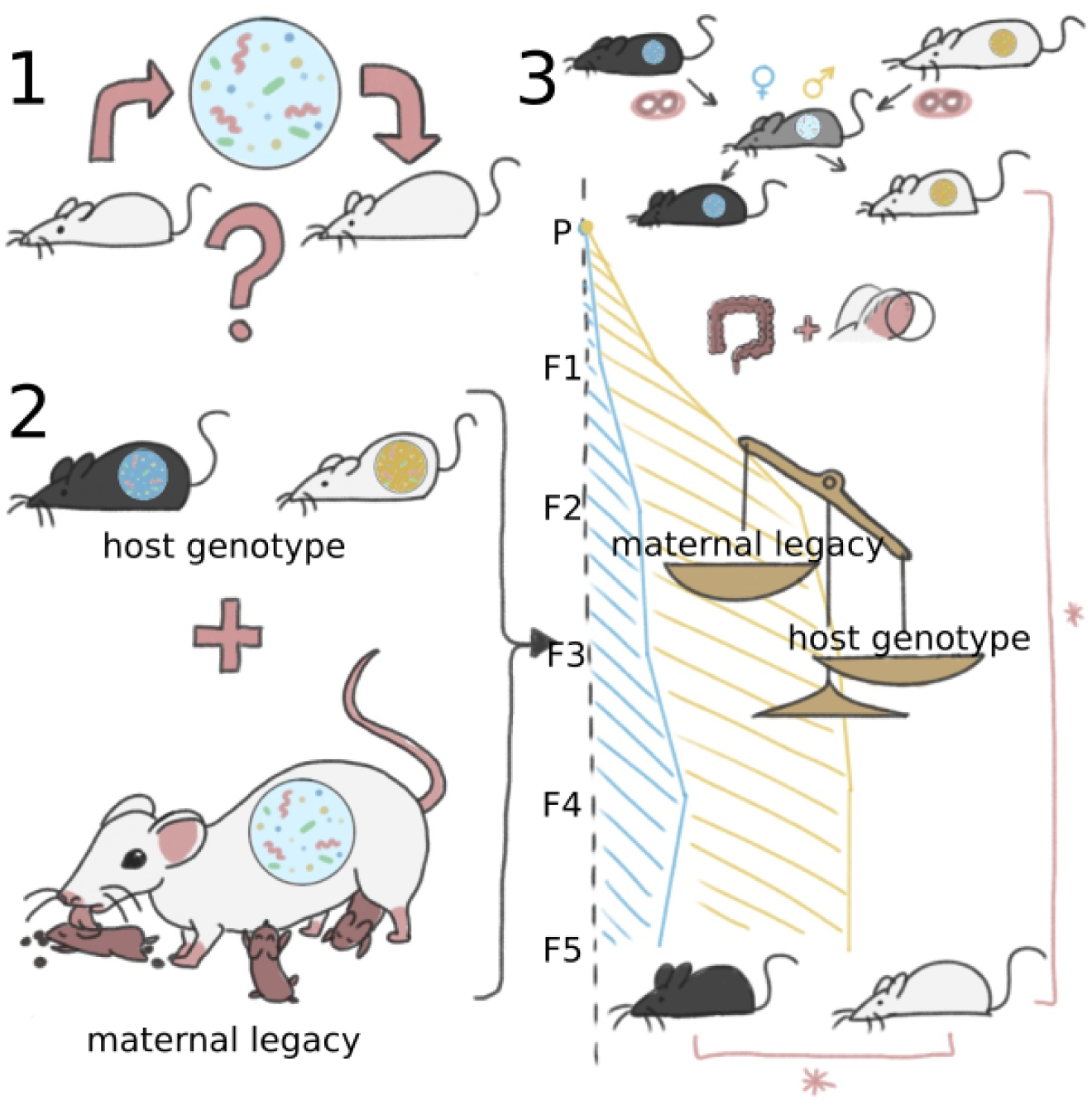
graphical abstract.

## Introduction

The human body is colonized by as many microbial cells as there are human cells [1]. Research of the last decades revealed the enormous impact of the microbiome on human health and wellbeing (see [2] for a review). Numerous factors have been identified that modulate the human colonizing microbiome like diet, exercise, animal contact and many more (see [3] for a review). It is the combination of the host’s genotype and its microbiome that together drive the host’s phenotype [4].

Understanding the mechanisms controlling the formation and function of microbial communities is essential in human biology. Standardization of endogenous and exogenous variables, such as genotype and environmental factors is hardly achievable in larger experimental animal models and impossible in humans. Therefore, mice are consecrated as the most used laboratory animals due to their advantages for experimental work. They served in deciphering fundamental physiological and pathological aspects in mammals. Available murine models range in complexity from simplified microbial communities, such as “Schaedler flora” [5], “altered Schaedler flora” [6], GM15 [7], Oligo-Mouse-Microbiota 12 [8] or humanized-microbiome models [9, 10] over specified pathogen free (SPF) laboratory mice [11], which are altered in a series of biochemical gut parameters [12], immunological [13] or anti-cancer fitness-promoting traits [14], to the more complex, wild mouse microbiota models [15].

From an evolutionary perspective, it seems unlikely that the host leaves microbial composition to chance. Extreme (genetically modified) genotypes affect the functionality of the immune system and thus contribute to changes in the composition of gut microbiota [16–20]. Multiple quantitative trait loci (QTL) from specific genomic regions seem to contribute to the host tailoring of the microbiome [21–24]. Two factors, among other undefined loci, are the major histocompatibility complex (MHC; H-2 in mice), as demonstrated by the analyses of bacteria-derived cellular fatty acids [25] and IBD susceptibility involved genes, such as caspase recruitment domain member 9 (Card9) [26]. Some studies showed genome-wide linkage with abundances of specific microbial taxa such as *Lactobacillus* [21, 27] or *Faecalibacterium prausnitzii* [28] whereas others document the influence of the “host genotype” and the environment on the whole microbiome [29–31].

Researchers aware of the importance of the microbiome in the experimental work proposed to scientific journal editors a mandatory documentation of all factors that may influence the microbiome, such as host genotype, husbandry details or experimental methods [4]. Factors like diet, bedding material, drugs, cage mates or ventilation are relatively easy to control for. The control of factors, which we subsume as maternal legacy, like passage through the birth canal, weaning, coprophagy and grooming is almost impossible or implies a significant increase in resources. However, they are known to impact the microbiome and thus most likely impact the host phenotype [19, 31, 32]. One could even speculate that maternal legacy alone is the evolutionary process to vertically transmit a defined microbiome to offspring generations.

The central question remains open, namely which of the two factors maternal legacy or host genotype contribute (more) to the active shaping of a host’s microbiome? A practical implication could be that strain differences from mice of alternative vendors would harmonize under identical environmental conditions through cross fostering if maternal legacy was to dominate microbial composition.

Existing literature is inconclusive about effect sizes of maternal legacy vs. host genotype. The vendor and genetic background, in terms of host genotype, seem to influence murine gut microbiota [33]. Nevertheless, studies using embryo transplantation and litter cross-fostering in mouse rearing and housing document that rather environmental conditions and maternal legacy exert a dominant contribution in shaping microbiota composition. A drift of the microbiota to a host genotype and facility-specific composition seem to occur under the influence of these factors [34]. Also, authors of [35] assumed that the foster mother’s gut microbiota rather than the host genotype influence gut microbiota composition in early life [36], whereas the study of Korach-Rechtman et al., 2019 indicates dominance of host genotype over the maternal inoculation by crossbreeding experiments [31]. Overall, authors of [37] account for the host genotype less than 20% of the gut microbiota variation in mice whereas the findings of [38] suggest that in humans the gut microbiome and host genotype are largely independent.

Numerous rodent studies that conclude on the influence of host genotype were performed either in immune defective phenotypes or were drawn secondarily to the main goals of the respective studies, often on highly related mice, which were purchased from commercial vendors shortly before the beginning of the respective study. In addition, no natural course of microbiota colonization and transmission over the generations was followed, rather artificial colonization with/or in association to antibiotics treatment were performed [39]. Moreover, most studies exclusively focused on the gut microbiome. Only recently, pioneering studies regarding the influence of host genotype on the microbiome of other body sites such as skin [40] and respiratory tract [41, 42] have been conducted in human and murine lung [43], while surveys for e.g. the genital tract are still missing.

To disentangle the factors host genotype and maternal legacy, we here obtained presumably microbial free 2-cell stage embryos of two representative wildtypes, namely C57BL6/J (B6J, n=42) and BALB/c (C, n=57), and transferred a mix of embryos into six SPF hybrid recipient mice (RM), which were generated from B6J dams and C sires. Therefore, offspring started from the same microbiome, acquired through maternal legacy of RM.

We continued the experiment over five generations of separated breeding, while minimizing impact of environmental factors through housing in individually ventilated cages (IVC). For reference, we also sampled six SPF-mice of each host genotype independently obtained from our mouse facility (Duesseldorf, Germany), housed in open cages instead of IVC, and bought from a commercial vendor (Janvier, France).

To further dissect maternal legacy, we implemented three cage lineages per host genotype, i.e. strict inbreed lineages that never came in contact in the following generations.

We applied 16S rRNA gene sequencing of colon content and skin of ear to obtain microbiome profiles of 334 mice in total. It was shown for immunodeficient mice that the host genotype itself alters the microbiome and leads to profound metabolome systemic and not just local effects within the gut [44]. For systemic insights, we therefore collected blood serum to obtain metabolomic data.

Our data show that under controlled environmental factors host genotype is the driving factor in microbiome composition over multiple generations in inbred laboratory mice. However, the maternal legacy effect is non-negatable, especially in earlier generations. Our analysis also documents a host genotype dependent increase of particular pathobiont microorganisms such as *Akkermansia muciniphila*, as well as host genotype specific metabolome correspondence.

## Material and methods

### Mouse strains and husbandry procedures

The mice strains C57BL/6J (B6J), BALB/c (C) and their F1 hybrid B6CF1 (RM) originated from the specified-pathogen-free (SPF) colony of the Central Unit for Animal Research and Animal Welfare Affairs (ZETT) Duesseldorf. They were free of all agents listed in Table 3 of the FELASA recommendations for health monitoring of rodents [11] and supplementary of *Staphylococcus aureus*, *Proteus spp.*, *Klebsiella spp*. *Bordetella bronchiseptica*, *Bordetella pseudohinzii, Pseudomonas aeruginosa*, *Muribacter muris* and *dermatophytes*. The access to this microbiological unit was restricted to a few animal caretakers through a sit-over barrier-system and complete change of clothes with sterile clothes consisting of suit-overall, underwear, socks, shoes, face-mask, head-cover and gloves. This unit was populated exclusively with mice strains hygienically sanitized by means of embryo-transfer. For the experiment, the mice were kept in individually ventilated cages (IVC) filled with Shepherd’s™ ALPHA-dri® bedding sheets (Shepherd Speciality Papers, Kalamazoo, USA) and had access *ad libitum* to autoclaved rodent chow (Ssniff, Soest, Germany) and acidified water. All cages were located in the same IVC rack during the whole period of the experiment and were housed under 12:12 h light/dark cycles, at a 22±2°C room temperature and 55±5% humidity. All mice cages were changed weekly with autoclaved fresh cages containing the same bedding, food and water.

### Study design and sampling

To obtain B6J and C embryos, female mice were intraperitoneally super-ovulated using 7 IU PMSG for B6J and 5 IU PMSG for C (Intergonan® 240 IE/mL, MSD Tiergesundheit, Unterschleißheim, Germany) and 7 IU hCG for B6J and 5 IU hCG for C (Predalon® 5000 IE, Essex Pharma GmbH, Waltrop, Germany) 48h later, followed by mating with males of the same strain. On day 1.5 after hCG administration, embryo donors were sacrificed, their oviducts extracted and the embryos at the 2-cell stage flushed using M2 medium (Sigma-Aldrich, Munich, Germany) according to [45]. An average number of eight 2-cell embryos of each B6J and C strains were transferred into the oviduct of each of the six pseudo-pregnant related B6CF1 recipient foster mothers (RM) used in this study as described previously [45]. The six RM were further placed in three individually ventilated cages, each containing two of the RM, where they gave birth after approximately 19 days to the mice parental (P) generation consisting of 16 B6J and 5 C mice (Figure 1). The P mice males and females were weaned at the age of approx. three weeks and placed together in two male and two female cages until the age of approx. seven weeks when they were either used for mating or placed in separate male or female cages until they reached the adult sampling age of fifteen weeks when sampling occurred. Three days before mating, dirty bedding originating from the males’ cages was transferred to the respective female’s cage in order to synchronize the ovulation. Three P generation breeding trios of one male and two females were settled for the B6J mice whereas for the C mice the only male available was mated for three days with two of the C females and then transferred to the second C female cage. For the following generations, a breeding trio was settled from each previous breeding cage, except for the F2 generation of C strain, where two breeding trios were settled from a cage (Figure 1). The exact cage location and lineage, the sex and the number of mice resulted per host genotype and generation can be depicted in Figure 1. In addition, twelve mice each (3 males and 3 females of each B6J and of C strain respectively), originating from Janvier Labs (Le Genest-Saint-Isle, France) and ZETT Duesseldorf respectively were included as controls. Duesseldorf controls were housed in open cages until sampling. Janvier controls were purchased at an age of 14 weeks and afterwards housed in IVC. The sampling occurred at the age of fifteen weeks for all mice, except for a few singular breeding mothers mice that still had to nurse for one or two further weeks and reached thus sixteen or seventeen weeks at sampling. The age of fifteen weeks was chosen since at this age the mice display a stable mature gut microbiome [46].

**Figure 1:**
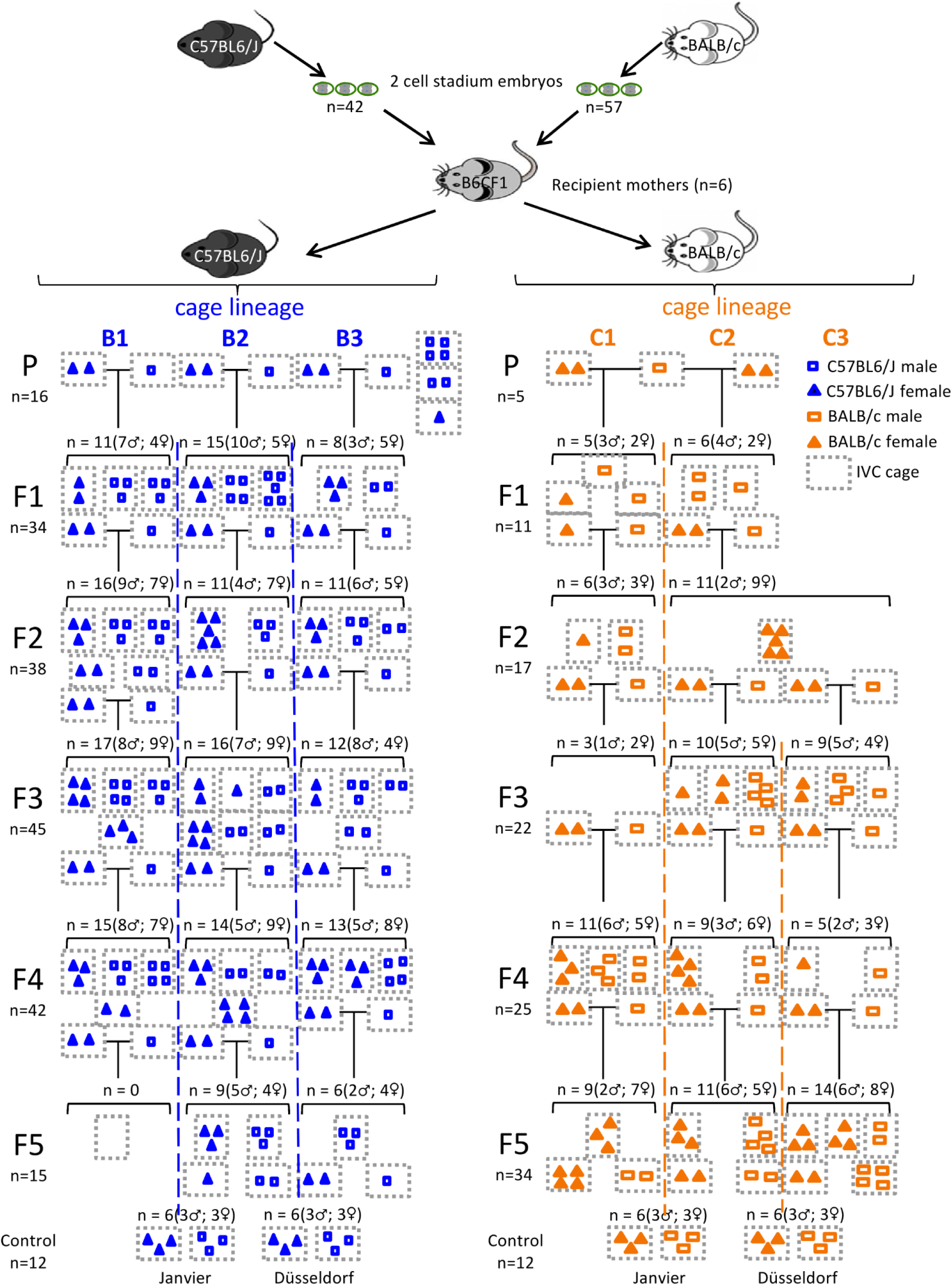
Breeding Strategy. We obtained C57BL6/J (B6J, n=42) and BALB/c (C, n=57) 2-cell stadium embryos from donor mice. A mix of both host genotype embryos was transferred into six recipient dams of a hybrid host genotype B6CF1, such that each dam gave birth to pups of both “host genotypes”. Offspring (P generation) was separated by host genotype into six cage lineages (B1-3, C1-3). Inbred for generations F1 to F5 always occurred within the same cage lineage (dashed lines). Gray dots indicate individually ventilated cages. Open squares and solid triangles indicate male and female mice, respectively, while blue icons indicate B6J and orange icons C host genotype, respectively. Last row gives numbers for control mice.

### Sample collection and DNA extraction

To harvest the samples, 15 weeks old mice were euthanized by bleeding in narcosis. The collected blood served for sera preparation. Next, approximately 2/3 of the left earlobe and the two to three most distal fecal pellets from the colon were harvested using sterile instruments and used for the analysis of the skin and gut microbiome respectively. All samples were placed into 1.5 mL sterile Eppendorf cups and immediately frozen at –80°C until further use. All samples were collected between 8:00 and 11:00 a.m. on several days. DNA extraction from colon pellets and skin was performed using the DNeasy PowerSoil and DNeasy Blood & Tissue Kit (Qiagen, Hilden, Germany) respectively using the manufacturer’s protocol. In the final step, DNA was eluted in EB buffer (Qiagen) and the yield was measured by Nanodrop One (Thermo Fisher Scientific, Waltham, USA). Extracted DNA was frozen at –20°C until further processing.

### 16S Amplicon library preparation and sequencing

Genomic DNA samples used for 16S rRNA gene sequencing were quantified by photometric measurement using NanoDrop One device (Thermo Fisher Scientific Inc.). Preparation of the 16S rRNA gene amplicon libraries for the Illumina MiSeq System was performed according to the Illumina 16S metagenomics protocol (Part #15044223 Rev. B) sequencing the V3-V4 region of the 16S rRNA gene (primers: FWD:CCTACGGGNGGCWGCAG, REV:GACTACHVGGGTATCTAATCC) with the change of the material input to 1 µL of the sample volume. Two Illumina i5 and i7 8bp barcodes were used for each sample for a 384 multiplexing schema. Final libraries were analyzed for fragment length distribution with the Fragment Analyzer (Agilent Technologies, Inc.) using the HS NGS Fragment Kit (1-6,000bp) assay (DNF-474). Concentrations were determined by fluorometric measurement using the Qubit Fluorometer and a DNA high sensitive assay (Thermo Fisher Scientific Inc.). Libraries were normalized to 2 nM, equimolar pooled and subsequently sequenced on a MiSeq system (Illumina Inc) with a read setup of 2×301bp by using a MiSeq Reagent v3 (600-cycle) Kit with three flowcells in total.

### Statistical Analysis

The base calling and simultaneous demultiplexing was done via `bcl2fastq` (v2.19.0.316) and primers were trimmed using `cutadapt` (v2.10, [47]). `Cutadapt` removes adapter sequences from high-throughput sequencing reads. Quality controlled sequence data was imported into the `Qiita` study management platform (https://qiita.ucsd.edu/; hosted at UC San Diego, [48]) under study ID 12148. Through `Qiita`, we used `QIIME` (v1.9.1 [49]) to clip reads to regions above a Phred score of 3, drop reads containing N base calls and trim reads to 150bp. The generation of feature tables was performed by *de-novo* amplicon sequence variant (ASV) determination using the `Deblur` approach (v1.1.0 [50]). Taxonomy for `Deblur` sequences was assigned via the `q2-feature-classifier` ([51]) of `QIIME2` (v2023.2 [52]) using the pre-trained Naive Bayes classifier https://data.qiime2.org/classifiers/greengenes/gg_2022_10_backbone_full_length.nb.qza, which is based on full length ribosomal sequences of `Greengenes2` [53]. As `GreenGenes2`’s taxonomy currently lacks labels for mitochondria and chloroplasts, we classified ASV sequences against the older `GreenGenes` (v13.8, [54]) database specifically ASVs assigned to ‘`c` `Chloroplast`’ or ‘`f` `mitochondria`’ as a pre-filtering. Low biomass skin samples have been controlled against “kitome” contamination [55] through `Decontam` [56] as suggested [57]. We used `Decontam` as provided through `QIIME2` version amplicon-2024.5 in “combined” mode and a threshold of 0.5.

In the following, the ASV feature table was used to determine the alpha– and beta diversities using QIIME2 as well as differential abundance analysis.

We used `q2-fragment-insertion` of `QIIME2` (v2023.5 [58]) to phylogenetically place all `Deblur` sequences into the reference `Greengenes` 13.8 99% identity tree [54] to obtain a phylogeny for downstream phylogenetic aware alpha– and beta-diversity metrics, i.e. Faith’s phylogenetic diversity index [59], weighted and unweighted UniFrac [60].

### Alpha and Beta Diversity

We chose a rarefaction depth of 1,000 reads per sample for skin samples and 6,000 for gut samples by analyzing alpha rarefaction curves for the three metrics “observed_features”, “Shannon” and “Faith’s PD” using 10 iterations for every depth. These depths were best for representing the highest taxonomic diversity while losing the least number of samples in our dataset. Alpha diversity was calculated using the plain number of observed features (richness), Shannon index, Chao1 and Faith’s phylogenetic diversity index (Faith PD). Beta diversity was calculated using the phylogenetic measure weighted and unweighted UniFrac, as well as the non-phylogenetic measure Bray-Curtis dissimilarity [61] and Jaccard-Needham dissimilarity [62]. Dissimilarity was visualized as Principal Coordinate Analysis (PCoA) in a 3D Emperor plot [63]. Significance between groups in alpha diversity was assessed by two-sided Mann-Whitney-Wilcoxon or Kruskal Wallis tests and for beta diversity group significance with PERMANOVA using 9,999 permutations, correcting via the Benjamini Hochberg approach.

### Differential Abundance Analysis

Statistically significant differentially abundant taxa were identified using analysis of composition of microbiomes (ANCOM) as a QIIME2 plugin [64].

### Joint analysis with Robertson et al. data

We obtained raw read files for [65] from NCBI’s BioProject with accession number PRJEB28381 and trimmed V4 primers 515F (Parada) and 806R (Apprill) off the reads (`cutadapt` v2.10, [47]). Further downstream processing (e.g. ASV calling, taxonomy assignments, filtering) was done identically to our dataset, see above. As both datasets target different variable regions (V4 and V3-V4 for Robertson et al. and ours, respectively) not a single ASV nucleotide sequence will be shared between both. We therefore limited alpha– and beta-diversity analysis to phylogenetic metrics, which indirectly merged the datasets by phylogenetically placing ASVs into the same `Greengenes` 13.8 99% identity tree [54]. The joint feature table was rarefied to 6,000 reads per sample.

### Metabolome analysis by GC-MS

Ten serum samples from the generations F3 and F4 belonging to each B6J and C host genotype were chosen for GC-MS based metabolic profiling, following previously established protocols [44]. Metabolite extraction was conducted with minor modifications to the methodology described by [66]. In brief, 1 mL of a –20°C cooled extraction solution composed of acetonitrile (ACN) / isopropanol (IPA) / water (H_2_O) (3:3:2, v/v/v) was mixed with 30 µL of a 25 µM internal standard (ISTD) solution (ribitol and N,N-dimethylphenylalanine). Then, 20 µL of sample was added to the extraction solution, vortexed for 10 seconds, shaken for 5 min and then centrifuged for 2 min at 14,000 rcf at 4°C. Next, two 450 µL aliquots of the supernatant were transferred to new tubes and 500 µL of an ice-cold solution of ACN / water (50:50, v/v) were added to remove any excess protein. After additional centrifugation for 2 min at 14,000 rcf, the supernatant was transferred to a pre-cooled tube and dried by vacuum centrifugation.

The dried sample was reconstituted in 150 µL of the extraction solution and dried again via vacuum centrifugation after transfer into a glass vial. The sample was derivatized with methoxyamine hydrochloride and N-methyl-N-(trimethylsilyl) trifluoroacetamide as described in [67]. After incubation for two hours at room temperature, 1 µL was injected into a GC-MS system (7890A GC and a 5977B MSD, Agilent Technologies) and chromatography was performed as described in [68]. Metabolite identification was performed on two levels. A quality control (QC) sample containing a mixture of target compounds was included as a reference to identify target compounds in the sample based on mass spectra similarity and retention time (annotation level: reference). In addition, the `AMDIS` software (http://chemdata.nist.gov/mass-spc/amdis/ v2.72, 2014) was used for deconvolution of mass spectra of target peaks before comparing spectra to the NIST14 Mass Spectral Library (https://www.nist.gov/srd/nist-standard-reference-database-1a-v14). Matches with more than 80% mass spectra similarity were assigned accordingly (annotation level: NIST match). Peaks were integrated using the software `MassHunter Quantitative` (v b08.00, Agilent Technologies). For relative quantification, metabolite peak areas were normalized to the peak area of the internal standard ribitol.

### Triglycerides quantification

Triglycerides in serum were recorded using the colorimetric Triglyceride Quantification Kit (Catalog number MAK266, Sigma-Aldrich, Darmstadt, Germany) according to manufacturer’s protocol.

## Results

### The host genotype overrides the maternal legacy effect over generations in constant environments

Conflicting reports about the dominance of host genotype or maternal legacy on microbial composition and the suggestion of Robertson and co-authors [65] to first generate F2 littermates for maximal microbial homogeneity before conducting genotype-phenotype experiments let us set out our breeding experiment in which we followed microbial composition of two commonly used wildtype strains across six generations (see Figure 1).

Our microbial gut data show a robust overriding impact of host genotype over maternal legacy for constant environments. Interestingly, this effect is not yet pronounced in the P generation (p=0.08, two-sided Mann-Whitney-Wilcoxon test), but in all that follow (Figure 2A: p < 0.01, except F5 and Figure 2B: p < 0.03, except p=0.91 for P). The unproportionally smaller number of B6J mice in generation F5 (10.9 mice on average, but only 0, 9 and 6 mice in F5 for cage lines B1, B2 and B3, respectively) led to a significantly smaller sampled microbial diversity (<u>Figure S1</u>A: p < 0.014 for B2, p < 0.004 for B3) and is primarily prohibiting detection of the above mentioned effect. Displaying the data as a PCoA of weighted UniFrac distances reveals two distinct clusters belonging to each of the “host genotypes” respectively (Figure 3). Although considered wildtypes, both “host genotypes” have different immune system responses, which probably drives the differential microbial composition in the gut, as the main host-microbiome interface in mammals. The significantly lower alpha diversity of C animals persists even when correcting for the number of founding sires in the P generation; three vs. one for B6J and C, respectively (<u>Figure S2</u>).

**Figure 2:**
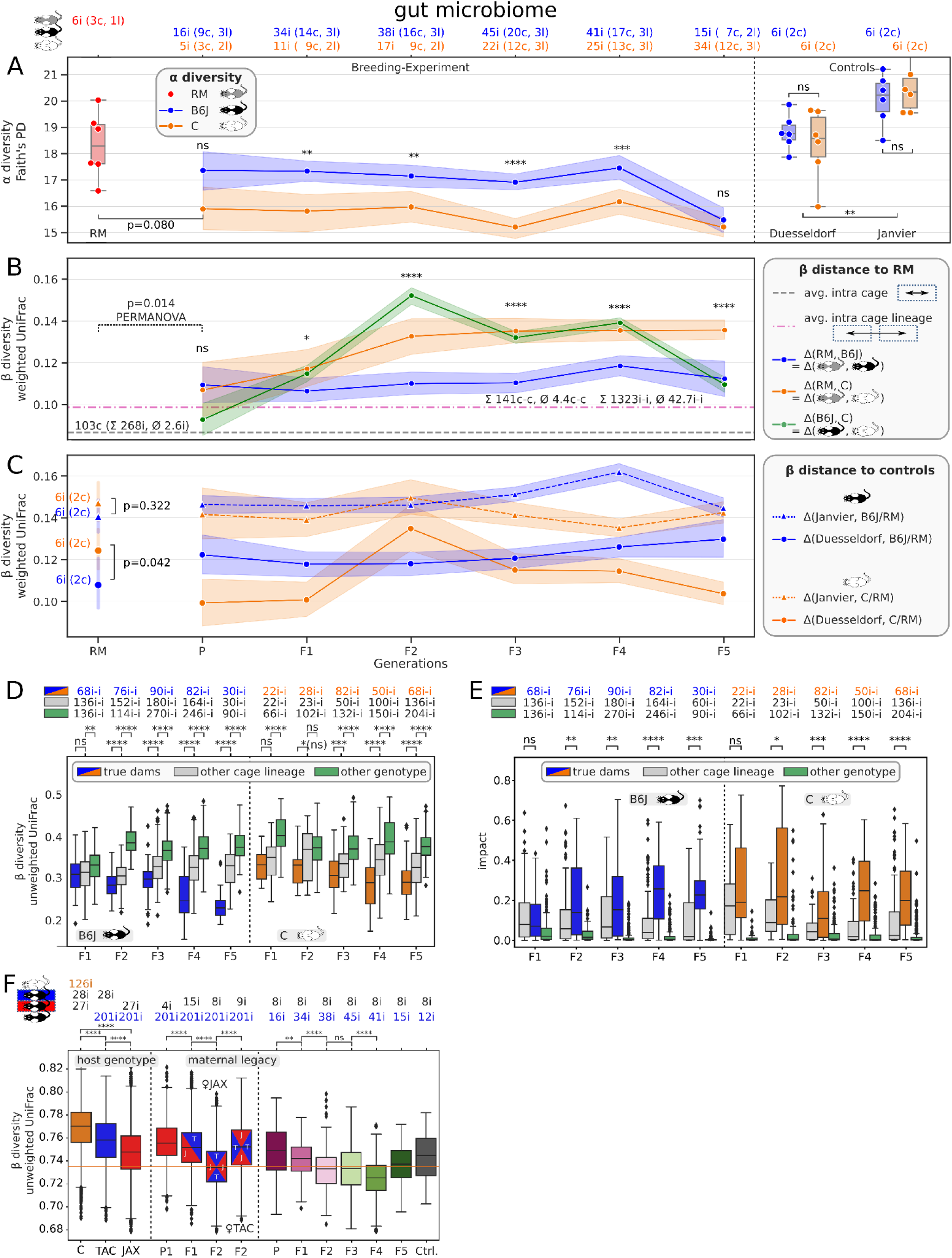
Trajectory of host genotype gut microbiome differentiation. A) Y-axis is Faith’s Phylogenetic diversity. X-axis is mouse generation or control group. Labels on top list numbers of individual mice (i), cages (c) and cage lineages (l) of which samples were aggregated by color: RM, B6J and C in red, blue and orange, respectively. B) Distances between six RM and 303 breeding experiment samples, grouped by host genotype (B6J=blue, C=orange) in terms of weighted UniFrac beta diversity. Green band indicates distance between “host genotypes”, not to RM. Gray dashed line is the mean pairwise distance between individuals housed in the same cage, i.e. between biological replicates; 103 cages with 268 individuals and 2.6 individuals per cage on average where considered. Magenta dashed line is the mean distance of individuals from two different cages of the same host genotype, same cage lineage and same generation; 1323 pairs of individuals (i-i) were considered with 42.7 i-i pairs on average per cage lineage and generation; considering housing, this are 141 different cage to cage (c-c) pairs with 4.4 c-c pairs on average per cage lineage and generation. *C)* Distances between 24 control and 303 breeding experiment plus six RM samples in terms of weighted UniFrac. We grouped control samples into host genotype and Janvier vs. Duesseldorf, such that each group consisted of 6 mice housed in 2 cages. *C)* Comparison of similarities between true dam to offspring (=true dams), dam to mice of same generation, other cage lineage (=other cage lineage) and dam to same generation different host genotype (=other genotype) in terms of unweighted UniFrac. Top label indicates the number of pairwise distances. *E)* Impact of maternal microbiome on offspring microbial composition in terms of source tracking for true dam to offspring (=true dams) and dam to mice of same generation, other cage lineage (=other cage lineage) and other host genotype (=other genotype). *F)* Joint analysis with Robertson et al. data. The y-axis is unweighted UniFrac. First three boxes summarize pairwise distances between our C and Robertson’s mice, our B6J and Robertson’s TAC and our B6J and Robertson’s JAX mice in orange, blue and red, respectively. Next four boxes relate our B6J mice with Robertson’s P1, F1 and F2 generation, where the latter is split into maternal JAX and maternal TAC mice. Subsetting Robertson’s F2 maternal JAX mice, the last seven boxes summarize distances to our six generations and B6J controls. We used Mann-Whitney-Wilcoxon for all statistical tests with Benjamini-Hochberg correction for multiple testing (ns: not significant, ****: p ≤ 0.0001, ***: p ≤ 0.001, **: p ≤ 0.01, * p ≤ 0.05)

**Figure 3:**
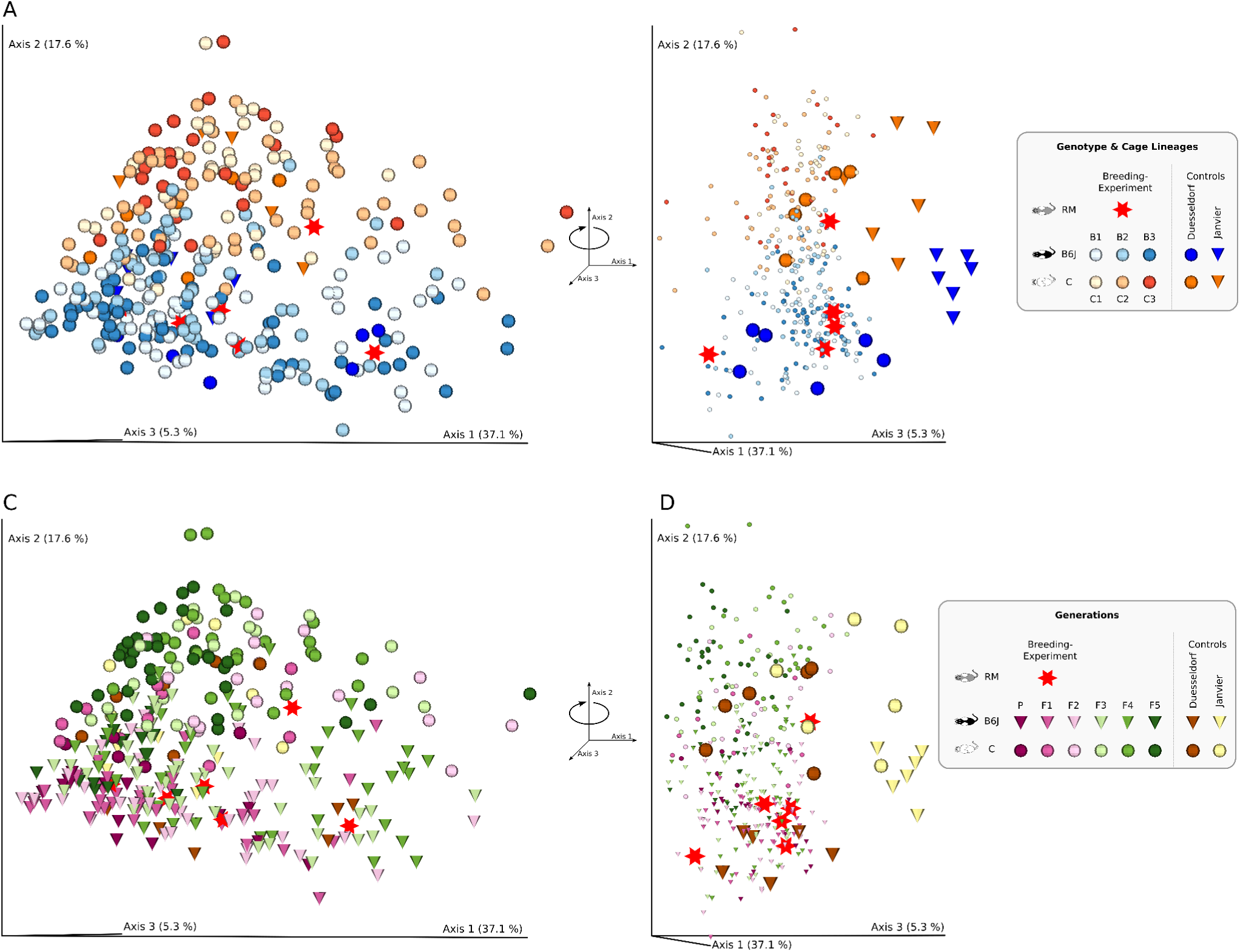
Gut Microbial Diversity. PCoA of weighted UniFrac distances for 333 colon samples. A) Colored by host genotype and cage lineage. B) Rotation of A along Axis 2. C) Same PCoA as in A, but color here indicates generation. D) Rotation of C along Axis 2.

The RM mice are generated by mating female B6J with male C mice. The high microbial similarity between “host genotypes” at the P generation (no significant difference, see above) seems to favor the maternal legacy effect; the weighted UniFrac distance between “host genotypes” (green line in Figure 2B) is lower than the average distance between any pair of mice from different cages within the same cage lineage (this comprises host genotype, magenta dashed line). In accordance, the RM microbiome is more similar to B6J Duesseldorf controls than to C Duesseldorf controls (Figure 2C, p=0.042, two-sided Mann-Whitney-Wilcoxon test). Due to open-instead of individually-ventilated-cages, control mice were exposed to a more relaxed environment and might therefore lack significant differences in alpha diversity (right part of Figure 2A: p=0.82 for Duesseldorf and Janvier controls, two-sided Mann-Whitney-Wilcoxon test). However, controls show strong separation by host genotype in beta diversity (p < 0.007 for all four tested metrics, PERMANOVA test with 9,999 permutations). The microbiome between RM and mice of the P generation is significantly different (PERMANOVA tests with 9,999 permutations on weighted UniFrac: p < 0.016), which might result from the relatively invasive embryo transplantation with preceding skin disinfection as an environmental distortion. This would explain the drop in alpha diversity, although not being significant (p=0.080, two-sided Mann-Whitney-Wilcoxon test). To explain the significant microbial dissociation in all following generations (F1 to F5) though, we must favor host genotype over maternal legacy, especially since microbiomes started quite homogeneously in the P generation, but drifted apart from F1 onwards, while we kept the environment constant. The relatively constant trajectory (blue line=B6J in Figure 2B) and the continuously increasing difference (orange line=C in Figure 2B) emphasizes that the microbiome of the hybrid RM mice is dominantly that of B6J mice and that there must be an active shaping in the C animals.

Despite the clear dominance of host genotype, environmental aspects easily exceed this effect, as can be seen by the significantly higher alpha diversity (Figure 2A, p=0.0013) of Janvier control mice, which were bought from a commercial vendor and only acclimatized for one week in our local facility prior to sampling at 15 weeks of age and the generally larger beta diversity distance of mice from our breeding experiment with Janvier controls, compared to Duesseldorf controls (Figure 2C: dashed vs. solid lines, respectively). The environment is probably furthermore a limiting factor for the degree of host genotype specific tailoring of the microbiome. Our narrow environment (autoclaved cages, autoclaved rodent chow and autoclaved bedding, acidified water, individually ventilated cages) provided a restricted set of microbes the hosts could source from, such that host genotype differences peek around F2 and probably shows intergenerational cycling thereafter [65, 69]. This would explain the flips in distance of C to control mice (orange lines in Figure 2C), whereas distances of B6J to control mice remain relatively stable as only the C host genotype actively tailors its microbiome away from the shared starting point, which is already B6J-like, in the P generation.

### Exploring family relations confirms the presence of a maternal legacy effect

The observation of C mice’s microbiome distance to RM increasing faster and stronger than between B6J mice and RM is a result of a maternal legacy effect as the mothers of the RM mice were of the B6J host genotype. In fact, RM samples are closer to B6J than to C Duesseldorf controls (p=0.042, two-sided Mann-Whitney-Wilcoxon test, Figure 2C).

As we established three cage lineages per host genotype, we could investigate maternal legacy in detail by comparing microbial distances within true family relations, i.e. individuals with their *true dams*, and non-family relations, i.e. distances to dams of *other cage lineages* but the same host genotype and last, distance to dams from the *other host genotype* (Figure 2D). Except for the F1 to F2 relation in C, we observe significant differences between the three categories with *true dams* showing the smallest distances towards their children. Using the SourceTracker [70] tool to estimate seeding capacity of alternative microbial sources to compose the offspring’s microbiome confirms that the *true dams* microbiome has the strongest impact (Figure 2E).

The maternal legacy effect might explain the dip in alpha diversity of B6J mice in generation F5 as well. Plotting the diversity by cage lineage (<u>Figure S1</u>) shows that samples in generation F4 of cage lineage B1 (blue) have exceptionally high alpha diversity, compared to B2 (orange) or B3 (green), quantified as Faith’s PD or as number of observed features. Although B3 significantly loses alpha diversity in F5, the mean across cage lineages in F5 also unproportionally suffers from a lack of B1 samples, with presumably high(er) diversity. This indicates that diversity can be crucially impacted by maternal legacy per generation.

Effect size analysis on gut data confirms dominance of host genotype. Mouse generation and maternal legacy have smaller but significant effect sizes, which has been reported previously, pointing out the importance of the mother in murine microbiome experiments (Figure 4).

**Figure 4:**
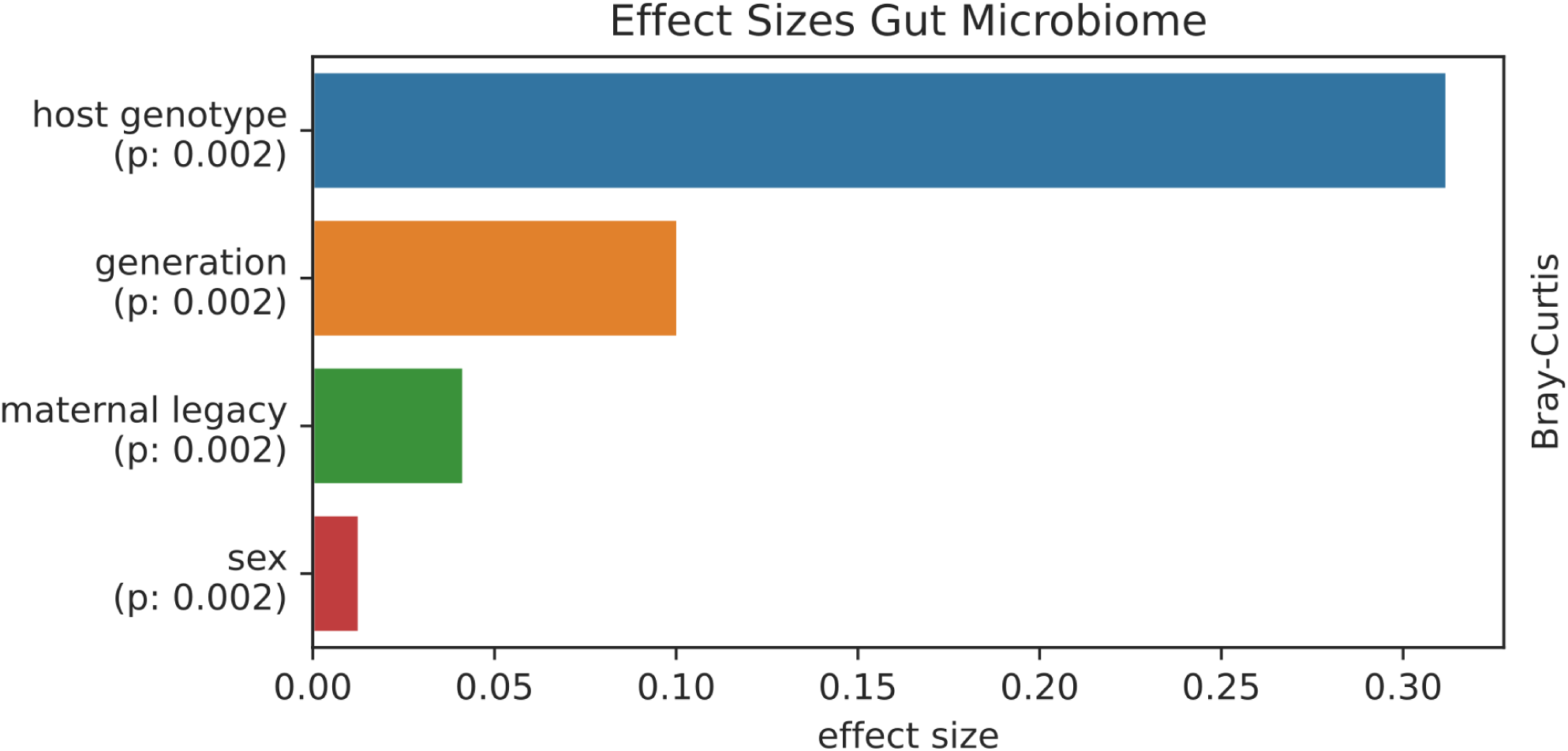
Effect Sizes Analysis. Forward step redundancy analysis with a linear model composed of host genotype, generation, sex and cage lineage (=maternal legacy) on Bray-Curtis distances.

### Joint-Analysis with independent data corroborates effects of host genotype and maternal legacy

In [65], authors investigated whether co-housing or F2 littermates, which we consider as the amalgamated effect of host genotype and maternal legacy, would lead to a more homogeneous microbiome prior to performing murine studies. Obtaining mice of very close “host genotypes”, namely the substrains C57BL6/J and C57BL6/N from two different vendors Jackson Laboratories (JAX) and Taconic Farms (TAC), respectively, they concluded that F2 littermates had a significantly higher impact on microbial standardization than co-housing. Thanks to published raw sequences from colon samples and prompt support with metadata (personal communication) we were able to perform a joint analysis.

We quantified the microbial distances between our C host genotype (n=126 mice of generations P to F5 and controls) and Robertson’s parental and littermate mice of both vendors (n=28 TAC + n=27 JAX), which are of a B6-like host genotype, as the orange box in Figure 2F. These distances are significantly larger than distances between our closely matching B6J (n=201 mice, p ≪ 10^-4^) host genotype and Robertson’s TAC (n=28, blue box, p ≪ 10^-4^) or JAX (n=27, red box, p ≪ 10^-4^) mice. This emphasizes the strong impact of host genotype on microbial composition. The decrease in distances of host genotype matching mice between different vendor dependent sub-strains (blue to red) aligns to the fact that our mice originate from a Jackson Laboratories purchase. This difference is significant and interestingly of similar magnitude as mismatching “host genotypes” (orange to blue). Practically, this could imply that not only the genotype of the utilized mice but also the substrain must be normalized for future murine microbiome experiments.

The maternal legacy effect probably complicates matters. Robertson et al. generated F2 hybrids of vendor sub-strains in two fashions: ♀JAX used JAX dams and TAC sires, while ♀TAC used TAC dams and JAX sires. Comparing distances between our n=201 B6J mice and Robertson’s n=8 F2 ♀JAX samples (sixth box in Figure 2F) or Robertson’s n=9 F2 TAC samples (seventh box in Figure 2F) illustrates that maternal legacy shapes significantly different microbiomes (p ≪ 10^-4^). As we employed a pure B6J host genotype of dam and sire for breeding, it is convincing that distances to Robertson’s F2 ♀JAX samples are significantly smaller than to F2 ♀TAC. This finding lets us specify the above recommendation to normalize or at least record female lineage for murine microbiome experiments.

Significantly decreasing distances from P1 (forth box) to F1 (fifth box) and F1 to F2 (sixth box) recapitulates Robertson’s recommendation to generate F2 mice prior to experimentation. Furthermore, stratifying Robertson’s 27 JAX mice (but not the 28 TAC mice) by generations (fourth to sixth box in Figure 2F), shows that they indeed become significantly more similar to our n=201 B6J mice over time – in accordance with our previous observations. Despite marked biological and technical differences between Robertson’s and our microbiome profiling, it is interesting to see, that our B6J samples become significantly more similar (except F3 and F5) to Robertson’s F2 JAX samples with preceding generations (seven rightmost boxes in Figure 2F). This might point to a universal host genotype specific core microbiome and warrants further investigation. Due to different variable 16S rRNA gene regions, we assume incompatible taxonomic assignments (cf. tremendous shifts in *Bacteroidota* / *Firmicutes_A* ratio in <u>Figure S</u>3) and therefore refrain from further investigations on taxonomic features.

Taken together we concur with Robertson et al. that F2 littermates should become the gold standard for microbial studies and we add that host genotype down to a level of substrain together with maternal legacy must be controlled for.

### Skin microbiome shows effects of “host genetics” but lacks maternal legacy

We sampled the skin of the left earlobe of all mice in addition to the previously discussed colon samples by processing the whole tissue in order to also capture sub-epidermis bacteria e.g. in hair follicles [71]. Lower biomass led to fewer reads and subsequently more samples were lost through quality control, invalidating application of statistical tests due to low sample numbers for some of the following comparisons.

As in the gut, host genotype shapes the skin microbial communities in a genotype dependent manner (<u>Figure S</u>4) although the results are not as decisive (Figure 5B: two-sided Mann-Whitney-Wilcoxon tests: generation P (p=0.057), F1 (p=0.201), F2 (p=0.074), F3 (p=0.007), F4 (p=0.392), F5 (p < 0.001)). The RM skin microbiome flips between being more similar to B6J and C three times throughout the P to F5 generations. Using control mice as reference instead (Figure 5C), the skin microbiome seems to be more similar to B6J for all but the P generation. Interestingly, alpha diversity (measured as Faith’s PD, Figure 5A) is never significantly different across any generation; this is also true for the alternative metrics “Shannon diversity”, “Chao1” and “observed features”, i.e. the raw number of different ASVs.

**Figure 5:**
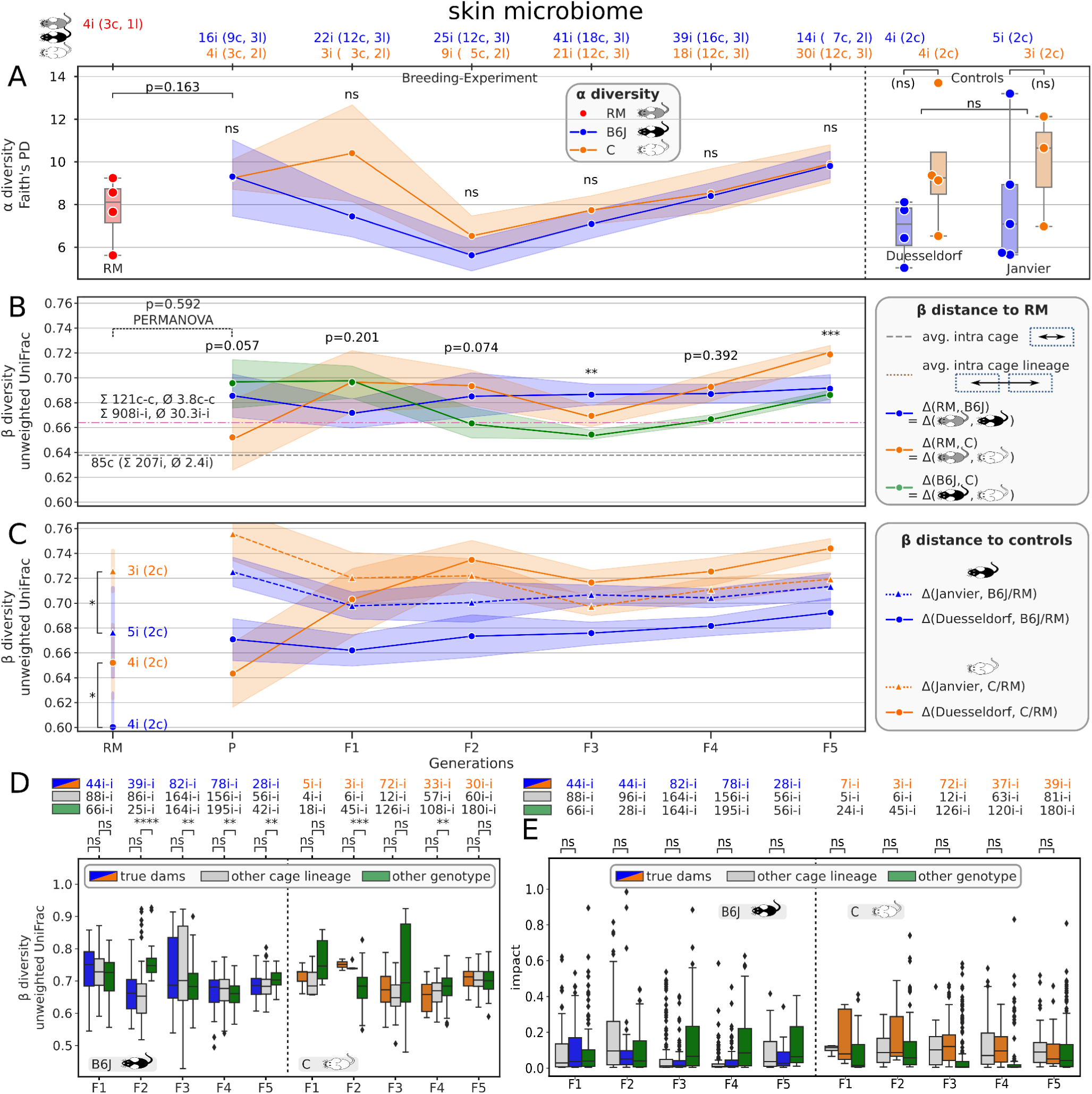
Trajectory of host genotype <u>skin</u> microbiome differentiation. Structure of this figure is identical to Figure 2 but for skin microbiome samples. Please consult the legend of Figure 2 for details. Differences are: Panel F is missing, since Robertson et al. did not collect skin samples. Panels B and C show unweighted instead of weighted UniFrac distances.

We were not able to measure a maternal legacy effect in the skin samples, neither by comparing alpha diversity (<u>Figure S</u>5), beta diversity (Figure 5D) nor by source tracking (Figure 5E). Again, low sample numbers prohibit statistical testing, but microbial alpha diversity between Janvier and Duesseldorf controls seem to be markedly different (Figure 5A), pointing to a stronger “environmental” impact on microbial composition.

Due to the cohousing of mice and their social nature, we cannot rule out the possibility of microbes from the gut transferring to the skin through factors like coprophagy and mutual grooming (cf. <u>Figure S</u>6 for gut/skin differences). Using five chow and five bedding control samples of the lots used for mice housing in addition to gut microbiome samples, stratified by host genotype and generation as “sources”, we quantified the contribution of community assembly in the skin (“sink”) via source tracking (<u>Figure S7</u>). Note that different read coverage between sinks and sources likely skews results. The source tracking analysis shows that the skin microbial community is only composed of 6% on average of microbes found in host genotype matching gut samples of the same generation. Microbes from the “opposite” host genotype gut microbiome account for negligible 1% on average. Cage bedding material (5%) and mice chow (16%) had similar or approx. three-fold stronger impact on skin microbiome assembly, whereas the huge majority of community composition remains unknown (68%), which might actually represent the “true” skin microbiome. We conclude that environmental effects dominate the skin microbiome with clear imprinting of host genotype tailoring but no detectable maternal legacy effects.

### Select taxa like *Akkermansia muciniphila* are linked to host genotype in both gut and skin microbiomes

Bacteria of the phyla Bacteroidota (75.13%) and Firmicutes_A (17.73%), dominate the baseline gut microbiota, whereas the Proteobacteria (2.28%) play a subordinate role (<u>Figure</u> <u>S</u>8A). In contrast, the phyla Firmicutes_D (45.88%) and Proteobacteria (20.53%) make up the majority of skin microbiota, whereas the Bacteroidota (12.32%) were much less abundant (<u>Figure S</u>8B). The taxonomic composition at genus level is presented in Figure 6A and <u>Figure S</u>8C for gut and skin, respectively. Both “host genotypes” shared most of the taxa in both gut and skin microbiomes over the generations and cage lineages. However, singular genera occurred preferentially only in B6J or C in the skin, as well as in combinations of generations or cage lineages in both gut and skin (<u>Figures S5</u> and <u>S6</u>). Collapsing the 946 gut ASVs to 102 named species level, ANCOM found 33 species to be significantly differentially abundant between B6J and C “host genotypes”, of which 21 had very low abundances (Figure 6B). Higher abundance of three species of the Bacteroides genus in C mice and higher abundance of six species of the *Muribaculacea* family, *Parasutterella*, *Ruminiclostridium*, *E siraeum* and *A. muciniphila* in B6J mice suggests a host genotype specific enrichment of particular taxa. Stratifying the relative abundance of *A. muciniphila* per generation (Figure 6C) shows an equally high abundance in the RM foster mothers, corroborating that the host genotype dependent shaping of the gut microbiome works through modulation of individual taxa transferred by the mother via maternal legacy.

**Figure 6:**
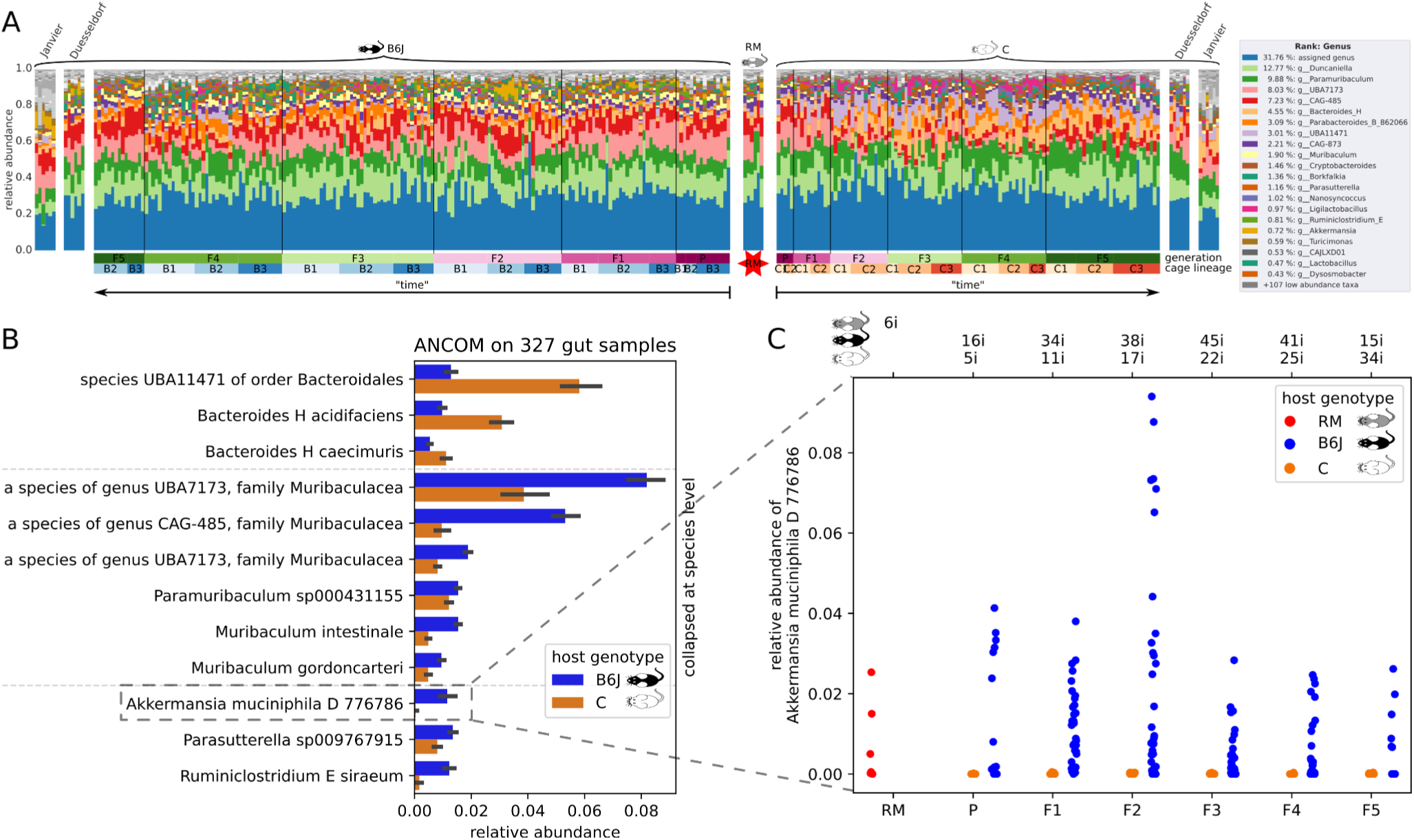
Gut Taxonomy: A) Taxonomic composition of 333 gut microbiome samples on genus level. Purple to green bar indicates generations, blueish and orange bar indicate cage lineage. B) Mean relative abundance of significantly differentially abundant species between “host genotypes”, determined via ANCOM. Further 21 species were excluded due to very low abundances. C) All relative abundances of Species Akkermansia muciniphila D 776786, stratified by generation (x-axis) and host genotype (hue).

### Blood serum metabolites correlate with host genotype and its colonizing gut microbiome

We quantified serum triglycerides in all 333 mice, to further investigate host and microbiome interaction. Interestingly, we found the same host genotype dependent correlation as with the gut microbiome, namely very similar triglycerides levels (p∼0.96, two sided Mann-Whitney-Wilcoxon test) between RM and B6J mice (Figure 7E) and significantly different (p < 0.0005) levels between RM and C mice. However, as triglycerides serum levels are already significantly different (p < 0.009) between RM and C in the first generation (P) after embryo transfer, a generation for which we could not detect microbial differences, we conclude that triglycerides levels are directly controlled by host genotype which in turn might help shaping the microbiome.

**Figure 7:**
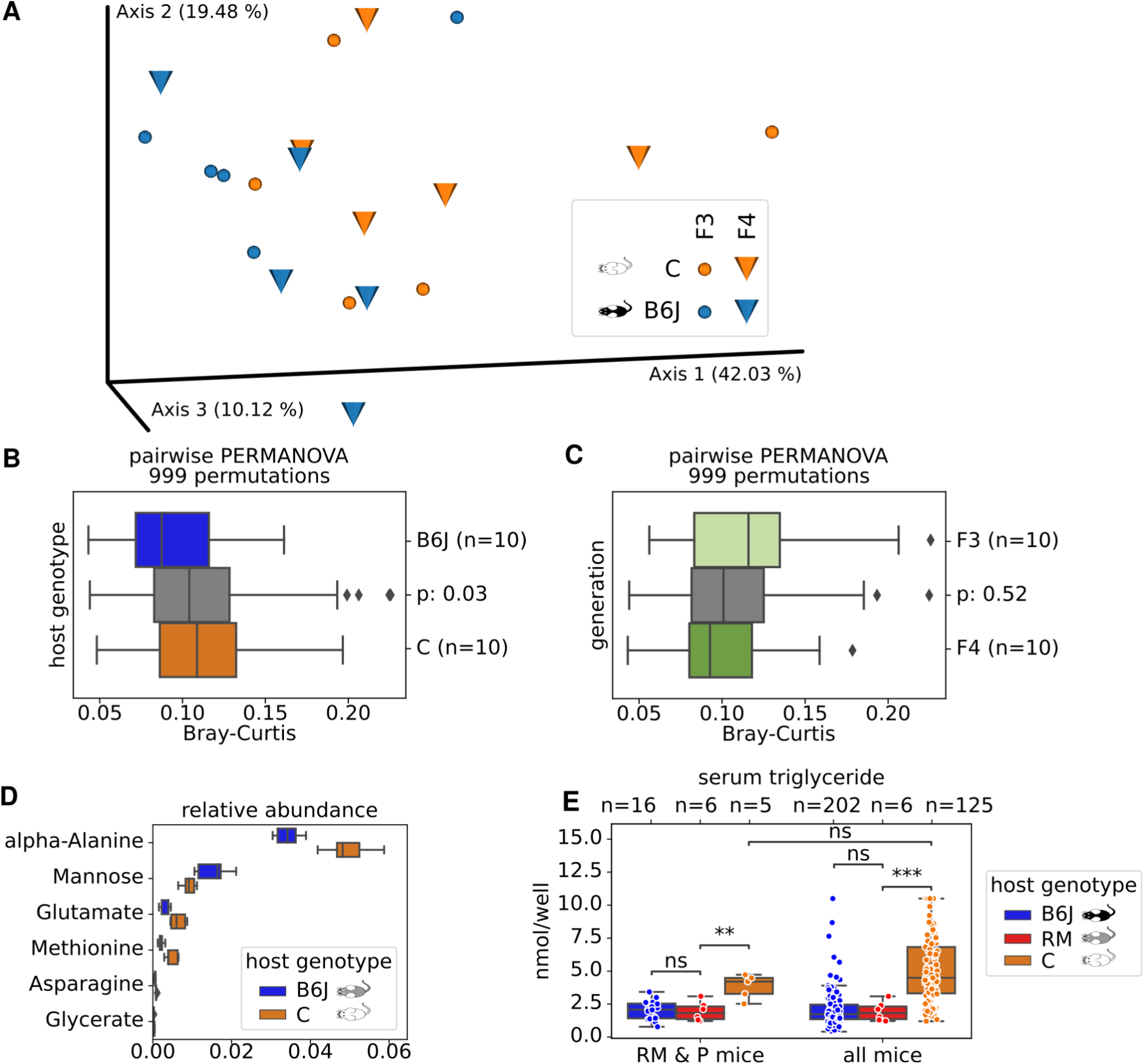
Gut Metabolite Diversity. A) PCoA of Bray-Curtis distances for 40 blood serum samples, obtained from 5 mice each in F3 and F4 of cage lineages B2, B3 and C3, C2, respectively. We quantified 41 metabolites per sample. B) Pairwise Bray-Curtis distances within “host genotypes” (B6J=blue, C=orange boxes) and between (gray) are significantly different as assessed via PERMANOVA. C) Pairwise Bray-Curtis distances within “generations” (F3=light green, F4=dark green) and between (gray) are not significantly different as assessed via PERMANOVA. D) Relative abundances of six metabolites, found to be significantly different by dsFDR between “host genotypes”. E) Serum triglyceride concentrations, stratified by host genotype (hue) for only early generations (=RM & P mice) and all generations (=all mice). Significance was assessed by Mann-Whitney-Wilcoxon with Benjamini-Hochberg correction.

It has been previously reported that the blood serum is a means to communicate gut microbial differences into the host organism. We therefore measured 41 serum metabolites via GC-MS of ten selected mice of B6J and C “host genotypes” each (Figure 7A). To capture temporal changes, we sampled mice of the generations F3 and F4 for which we found pronounced microbial differences. To exclude maternal legacy we intentionally sampled different cage lineages in both generations, i.e. B2, C3 and B3, C2 in F3 and F4, respectively. Quantified as Bray-Curtis pairwise distances, we found significant (p=0.02, PERMANOVA with 999 permutations) differences in serum metabolite profiles between “host genotypes” (Figure 7B). Closer inspection via dsFDR found six out of the 41 metabolites to be differentially abundant between “host genotypes” (Figure 7D). From our limited metabolome data, we can only speculate about directionality, but the observed differences might be a direct result of gut microbial metabolite production, which penetrates into the host’s bloodstream. We could not detect differences (p=0.52, PERMANOVA test with 999 permutations) between generations F3 and F4 (Figure 7C).

## Discussion

### The host genotype shapes its host’s microbiome

To what extent the host genotype affects the microbiome composition, and whether this effect is general or impacts only certain taxa is still subject to debate. The littermates are regarded as gold standards in microbiome standardization of experimental groups [65]. The immune system dwells with the microbial world and extreme immune altered “host genotypes” clearly influence the composition and diversity of the gut microbiome [16–18, 20]. Nevertheless, studying the impact of unmodified “host genotypes” on the microbiome is more difficult, because the effects are usually softer and cannot be directly attributed to particular engineered genes. However, the contribution of particular genes or genomic quantitative trait loci (QTL) to the microbiome tailoring [40, 72] or associations to microbial taxonomies and especially to particular genera such as *Bifidobacterium* has been demonstrated [73, 74]. The host genotype possibly acts on the microbiome by the innate and adaptive immune systems, which sequentially shape the gut microbiota, lipid metabolism and stat3 phosphorylation [75] and thus applies different evolutionary within-host selection forces to the microbial communities [76]. The two mouse strains B6J and C differ substantially in their immune responses to various infectious agents and are seen as prototypes for Th1 and Th2 immune response, respectively [77, 78], which may induce through microbiome-immune system interaction, different microbial communities [79].

Our approach studies whether differences in the microbiome can occur over generations in littermates of different “host genotypes” (B6J and C) in a constant environment, after a natural course of colonization with a common microbiome of B6CF1 recipient mothers. Moreover, the two main microbiome ecological niches, the gut and the skin are considered, since knowledge on body sites other than gut is currently sparse.

We demonstrated that the host genotype essentially contributes to the active shaping of the gut microbiome and has a powerful influence on the host’s metagenome. Once, directly through its stable genome, but in addition indirectly through tailoring of the flexibel composition of microbes that colonize the host. Despite the limitations in the profiling of the skin microbiome mentioned below, it seems that role of the host genotype is less decisive for the formation of the skin microbiome, which might be rather environmental dependent. Since the host genotype is heritable, this is an important factor in the microbiome evolution over the generations in spite of possible changes in the environment [80, 81].

Interestingly, we observed that the microbiome of our B6J mice was more similar to Robertson’s B6J than to Robertson’s B6N mice despite marked spatio-temporal (2019 vs. 2022, Canada vs. Germany), biological (sampling at 15 vs. 8 weeks of age, acidified vs. non acidified water, cages changed weekly vs. bi-monthly, commercial pelleted food vs. autoclaved chow) and technical (different technicians, V34 vs. V4 16S rRNA gene region, different sequencing centers) differences. This finding sustains that host genotype differences at substrain level (B6J vs. B6N) are still enough to produce host genotype related shaping of the gut microbiome and might point to a universal host genotype specific core microbiome, which warrants further investigation. Despite the clear dominance of host genotype, environmental aspects easily affect the microbiome tailoring, as can be seen by the significantly higher alpha diversity of Janvier control mice compared to Duesseldorf mice (Figures 3A). The environment is probably a limiting factor for the degree of host genotype specific tailoring of the microbiome in our experiment. Thus, the environment and host genotype decisively influence the composition of gut murine microbiota [33].

Whether or not the host genotype differences in the skin microbiome of the earlobe are actively tailored by the mice or is an indirect reflection of the active tailoring of the gut microbiome with subsequent spreading of these microbes to their skin cannot be unambiguously answered from our data. Fewer viable bacteria than predicted by bacterial DNA profiles colonize the skin surface [71]. It is plausible that bacterial environmental noise might impact the recording of the skin microbiome. Nevertheless, the environmental noise should partially originate from the viable skin-associated bacteria that are predominantly located in hair follicles and other cutaneous invaginations [71] and correlate taxonomically. Our analysis included the hair follicle and invaginations by including the whole ear lobe skin and not only bacteria from the upper layers of the epidermis. Moreover, since mammalian skin is a highly specialized habitat, capable of strong selection from available species pools [82], filtering thus probably occurs by host own forces and shapes the pool of bacteria that lead to this type of contamination. The influence of environmental noise on the microbiome could be partially reduced by the RNA-based profiling as a preferred screening method [82]. Nevertheless even a RNA-based profiling still records the living contaminants such as the ones acquired by coprophagy or from cage environmental sources such as bedding/food. Although we used a DNA-based profiling, the source tracking analysis shows that the impact to which the environmental contaminants drive microbial composition of the skin samples, play only a subordinate role. (<u>Figure S7</u>), implying a host genotype active tailoring also in the case of skin microbiome but to a much lower extent as for the gut. The interactions between the host immune system and skin is presumed to be much less intensive. External skin is in general more prone to environmental conditions and thus much harder to control for.

### The host genotype also shapes its metabolome

Multiple health and disease markers are correlated with the composition of the gut microbiome in humans [83]. In addition, the human gut microbiome affects the host serum metabolome and is linked to insulin resistance [84].

Using GC-MS-based metabolomics, we demonstrated differential expression and abundances of serum metabolites among selected B6J and C mice. Our findings indicate that the differences in microbiome could modulate together with the host genotype the expression of systemic markers (see Figure 7).

### The host genotype enriches specific microbial taxa

Analysis of the taxa variation between B6J and C mice in our study revealed that particular taxa were enriched by “host genotypes”. Interestingly, [85] observed 22 taxa to have a significantly higher abundance in B6J than C mice, including *Akkermansia* and *Ruminococcus*. (We assume equivalence between genera *Ruminiclostridium* and *Ruminococcus* in our data, as 99.6% of *Ruminiclostridium* reads classify as *Ruminococcus* when using Horne et al. outdated GreenGenes version.) The similar enriched abundances of *Akkermansia* and *Ruminococcus* in both studies, regardless of the experimental design, suggest that the gut environment of B6J but not of C mice is auspicious for these taxa. This may be due to increased availability of niche energy source Muc-2 in B6J, since *A. muciniphila* has the ability to degrade Muc-2 O-glycans *in vitro* [86]. *A. muciniphila* is an important pathobiont influencing numerous animal experimental phenotypes and accounts for 1-5% of the gut microbial community in healthy human adults, being a marker of a healthy microbiome and increasing the integrity of the intestinal barrier in both humans and mice [87]. There are obvious relationships between *A. muciniphila* and chronic inflammatory metabolic diseases such as type 2 diabetes, obesity, and IBD [88–90]. Interestingly, *A. muciniphila* accounted for up to 9% of the gut microbiota of the B6J but not of the C mice of our study (Figure 6C).

Overall, most of the microbial genera were shared by both “host genotypes” and inherited over all generations (Figure 6A), although differential taxa abundances occurred between “host genotypes” (Figure 6B), whereas singular genera were present only in some “host genotypes”, cage lineages and generations. Moreover, particular genera jumped over some generations, probably under the detection limit, and reappeared in a later generation (<u>Figure</u> <u>S9</u> and <u>S10</u>).

### The maternal legacy imprints the microbiome

The intergenerational changes recorded in our data are in accordance with previous studies. When inbred mouse strains were transferred into a new facility [69] and [65] reported minor changes of intestinal microbial composition and/or function across generations. Moreover, such studies suggest that even the more resilient wilding’s gut microbiota [15] are expected to change as animals are housed under laboratory conditions [91]. An expected host genotype independent finding was thus the cage lineage specificity, emphasizing the role of maternal legacy in microbiome heredity (Figures 2A, <u>3</u>D and <u>3</u>E) similar with previous studies [92]. Importantly, maternal legacy does not necessarily mean maternal microbiome if the male remains in the female cage during pup rearing favoring also the paternal horizontal transmission of microbial taxa [65]. Overall, we here documented by microbiome source tracking that the maternal legacy and the dam itself are responsible for most of the gut microbiome transmission to the offspring (Figure 2E and 4E). In our study, maternal legacy represents the second most important endogenous factor contributing to the shaping of the gut microbiome after host genotype effects.

### The host genotype is dominating maternal legacy, which both shape the host’s microbiome

Multiple studies attribute the host genotype a certain degree of influence on the gut microbiome [29, 30, 37], whereas others attribute to the host genotype a secondary role [34, 36] or no importance in microbiome shaping [93] at all. The authors of [38] examined the host genotype and microbiome data from 1,046 healthy human individuals with several distinct ancestral origins who share a relatively common environment, and found that the gut microbiome is not significantly associated with genetic ancestry, concluding that host genotype have a minor role in shaping the microbiome composition. Nevertheless, a narrow standardization of human individuals to a level similar to mice studies concerning host genotype (inbreeding) and environmental conditions is not achievable.

Previous work to disentangle the impact of host genotype and maternal legacy on the composition of offspring microbiome only sampled the first offspring generation [33, 34, 37]. Since no significant microbial difference could be detected after embryo transfer, cross-fostering or co-housing, exactly as in the microbial profiles in our P generation, authors rightfully concluded that maternal legacy dominates any “host genetics” effects, if present at all. Interestingly, a dominance of the host genotype over the maternal inoculation was also documented by cross-breeding of inbred mice [31]. However, both scenarios were based on a temporally limited observation.

The straight experimental design of our study, spanning seven generations of mice, with identical exogenous parameters regardless of the endogenous host genotype dichotomy, clearly shows that microbial differences manifest in the F1 generation and further increase over time, at least for constant environments. We therefore argue that our data is compatible with previous contradicting findings, however our longer temporal sampling suggests the opposite conclusion, namely that “host genetics” dominates maternal legacy, which for itself, but to a lesser extent, is also acting in tailoring the microbiome. According to a forward step redundancy analysis on Bray-Curtis dissimilarities, host genotype turned out to be the main driver of gut microbial diversity with an effect size of 0.312, followed by generation (0.100), “maternal legacy” (0.04205) and sex (0.013) (Figure 4). The weak signal on skin microbiome indicates that the microbiomes of different anatomical sites are driven with different power by intrinsic and extrinsic influences such as host genotype and environment. The remaining charred size effects for the microbiomes were possibly driven by the common environmental factors in this study. It is reasonably at this time point to hypothesize that the gut microbiome as an “intern” microbiome, without strong environmental contact is either prone to changes by host own factors such as host genotype, whereas microbiomes with high environmental contact such as the skin microbiome appear in this study more environmentally dependent.

### Outlook

The implication of the host genotype in shaping microbial communities of further body sites such as of the genital and respiratory mucosa should be addressed in the future. Future research may also document whether the microbiome tailoring by the host genotype can explain why some mice strains are more suitable for particular experimental models then others. For example, it would be interesting to study whether the host genotype dependent enrichment of the same particular taxa occurs independently in multiple facilities or whether the *A. muciniphila* dependent phenotypes could be recapitulated in mice strains that behave refractory to *A. muciniphila* enrichment such as C mice. Overall, the host genotype related shaping of the gut microbiome points to the existence of a universal host genotype specific core microbiome in inbreed laboratory mouse strains that warrants further investigation.

## Conclusion

Our results conclude that microbial communities at different body sites are driven by different endogen and exogenous factors. While the host genotype strongly influences the active shaping of the gut microbiome, it appears that the skin microbiome is more prone to environmental conditions. Although the microbial genes clearly outnumber genes directly encoded by the host, we propose here that the host genes, as the stable part of the holobiont, play a leading role expressing phenotypes through its microbiome shaping capacity; possibly through the establishment of universal host genotype specific core microbiota.

## Declarations

### Ethics approval and consent to participate

All procedures on animals were performed in accordance with the European guidelines for use and care of laboratory animals: European Commission Directive 2010/63/EU of the European Parliament and of the Council of 22 September 2010 on the protection of animals used for scientific purposes. Experiments were approved by the State Office for Nature, Environment and Consumer Protection (LANUV, State of North Rhine-Westphalia, Germany) under the number 81-02.04.2017.A383.

### Consent for publication

Not applicable.

### Availability of data and materials

The trimmed, demultiplexed fastq sequencing data supporting the conclusions of this article are available in the European Nucleotide Archive, PRJEB70879 https://www.ebi.ac.uk/ena/browser/view/PRJEB70879. The GC-MS raw data supporting the conclusions of this article are included within the article as <u>Additional File 1</u>. The analysis notebook and required files to reproduce all statistics and figures is available as <u>Additional</u> <u>File 2</u>.

### Authors’ contributions

Conceptualization, LB and SJ; Formal Analysis, AR, PW, SJ; Investigation, LB, CG, TW, WPMB, EE, APMW, KK, MS; Writing LB, AR, SJ, Funding Acquisition LB, SJ. All authors read and approved the final manuscript.

## Supporting information

Additional File 1: Metabolites table

Additional File 2: Analysis repository

## Acknowledgments

We thank the anonymous reviewers for their careful reading of our manuscript and their many insightful comments and suggestions. We gratefully acknowledge Isabel Schäfer, Manuela Stockhausen and Sandra Plante for excellent technical assistance. We sincerely appreciate the exemplary means, which the authors of Robertson et al. invested in reproducibility and additional, prompt assistance from Dana Philpott and Heather Maughan. We furthermore want to express our cordial regret of the loss of Susan J. Robertson.

## Supplement

**Figure S1:**
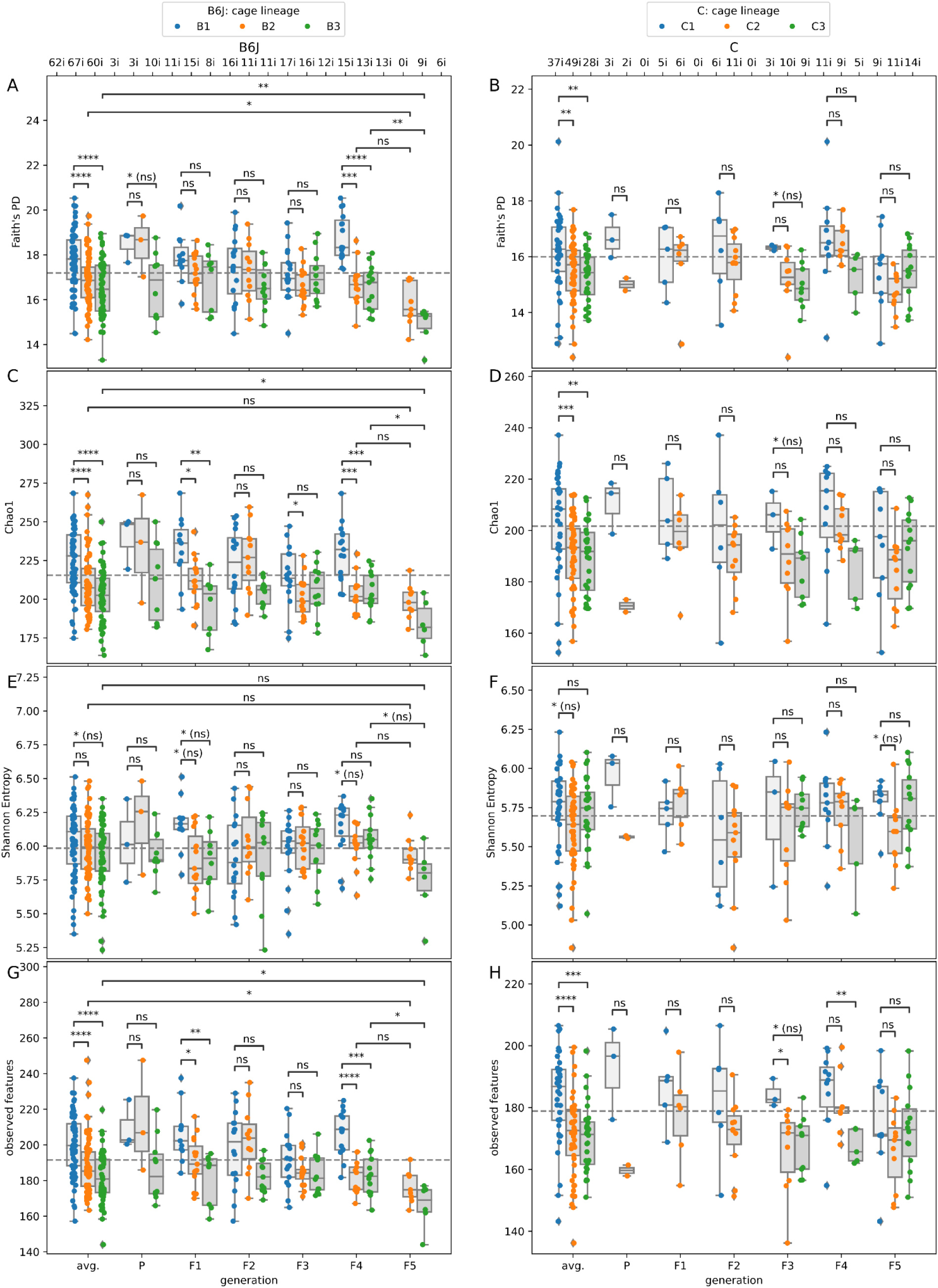
Individual Alpha Diversity of Gut Samples. Left and right columns display alpha diversity of B6J and C samples, respectively. Rows are different alpha diversity metrics: Faith’s PD, Chao1, Shannon Entropy and number of observed features, i.e. ASVs. The x-axis stratifies samples into generations, while hue indicates the three different cage lineages. Leftmost x-axis positions, entitled “avg.” are samples from all generations lumped together. We used two-sided Mann-Whitney-Wilcoxon tests with Benjamini-Hochberg correction to assess statistical significance.

**Figure S2:**
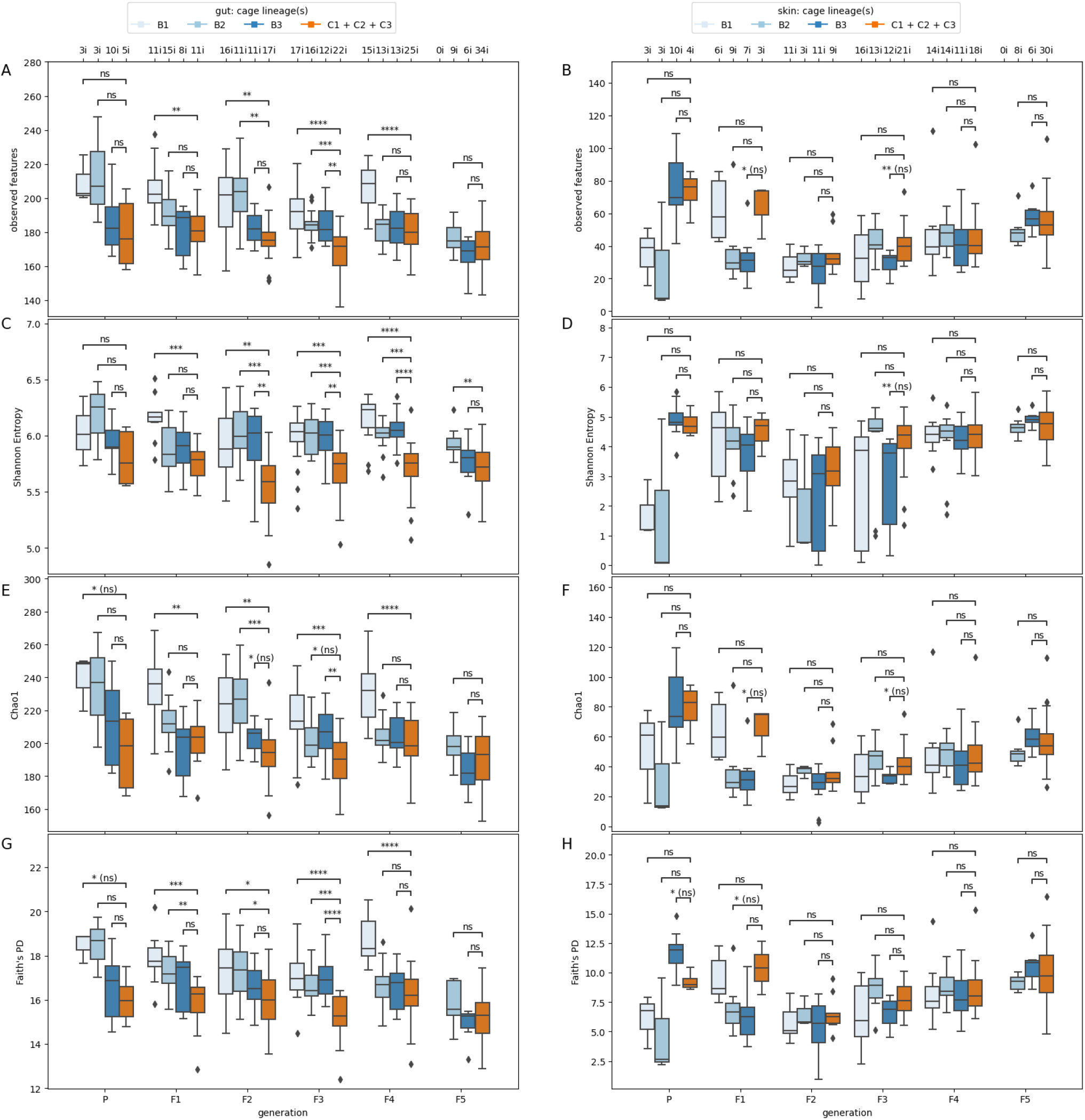
Impact of number of founding sires on gut alpha diversity. Left and right columns display alpha diversity of gut and skin samples, respectively. Rows are different alpha diversity metrics: Faith’s PD, Chao1, Shannon Entropy and number of observed features, i.e. ASVs. The x-axis stratifies samples into generations, while hue indicates samples originating from an individual founding sire in generation P, which coincides with cage lines for B6J and leads to lumping all three cage lines for C into one category. We used two-sided Mann-Whitney-Wilcoxon tests with Benjamini-Hochberg correction to assess statistical significance.

**Figure S3:**
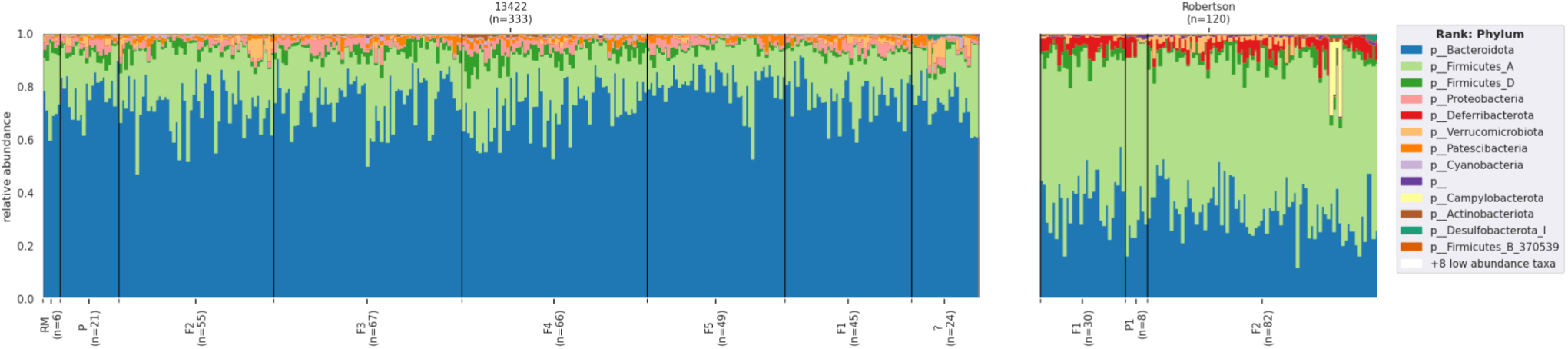
Taxonomic composition of our gut (n=333, labeled as Qiita study 13422) and Robertson’s et al. colon (n=120) samples. Feature counts have not been rarefied. Due to different variable 16S rRNA gene regions, we assume incompatible taxonomic assignments. Exemplary is the difference in the Bacteroidota / Firmicutes_A ratio.

**Figure S4:**
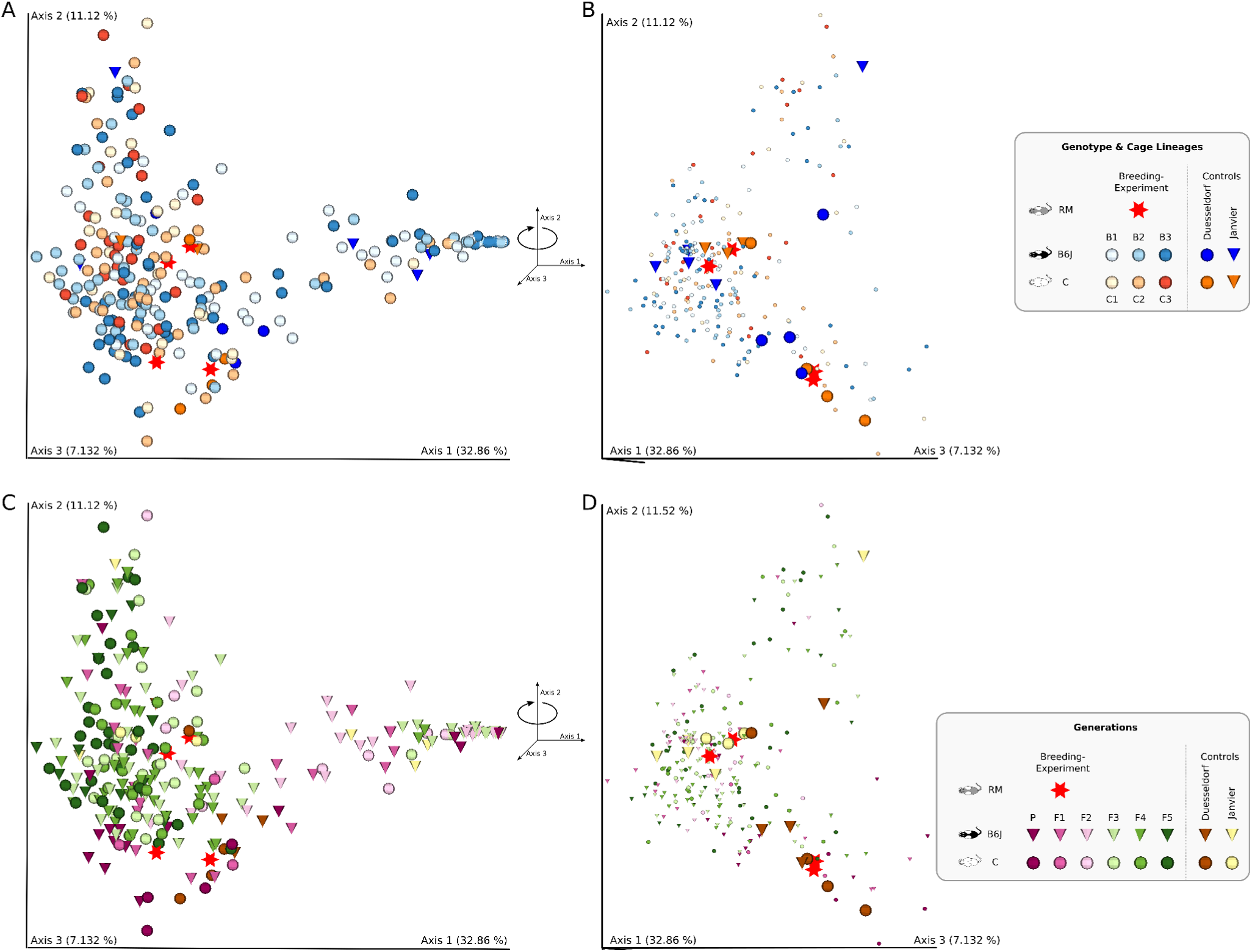
Skin Microbial Diversity. PCoA of weighted UniFrac distances for 262 external skin of the left earlobe samples. A) Colored by host genotype “host genotype” and cage lineage. Red stars indicate RM samples, while cones stand for Janvier control samples. B) Rotation of Panel A along Axis 2 with decreased icon size of Breeding-Experiment samples to highlight RM and control samples. C) Same PCoA as in Panel A, but color here indicates generation. Spheres indicate C samples; cones indicate B6J samples. D) Rotation of Panel C along Axis 2.

**Figure S5:**
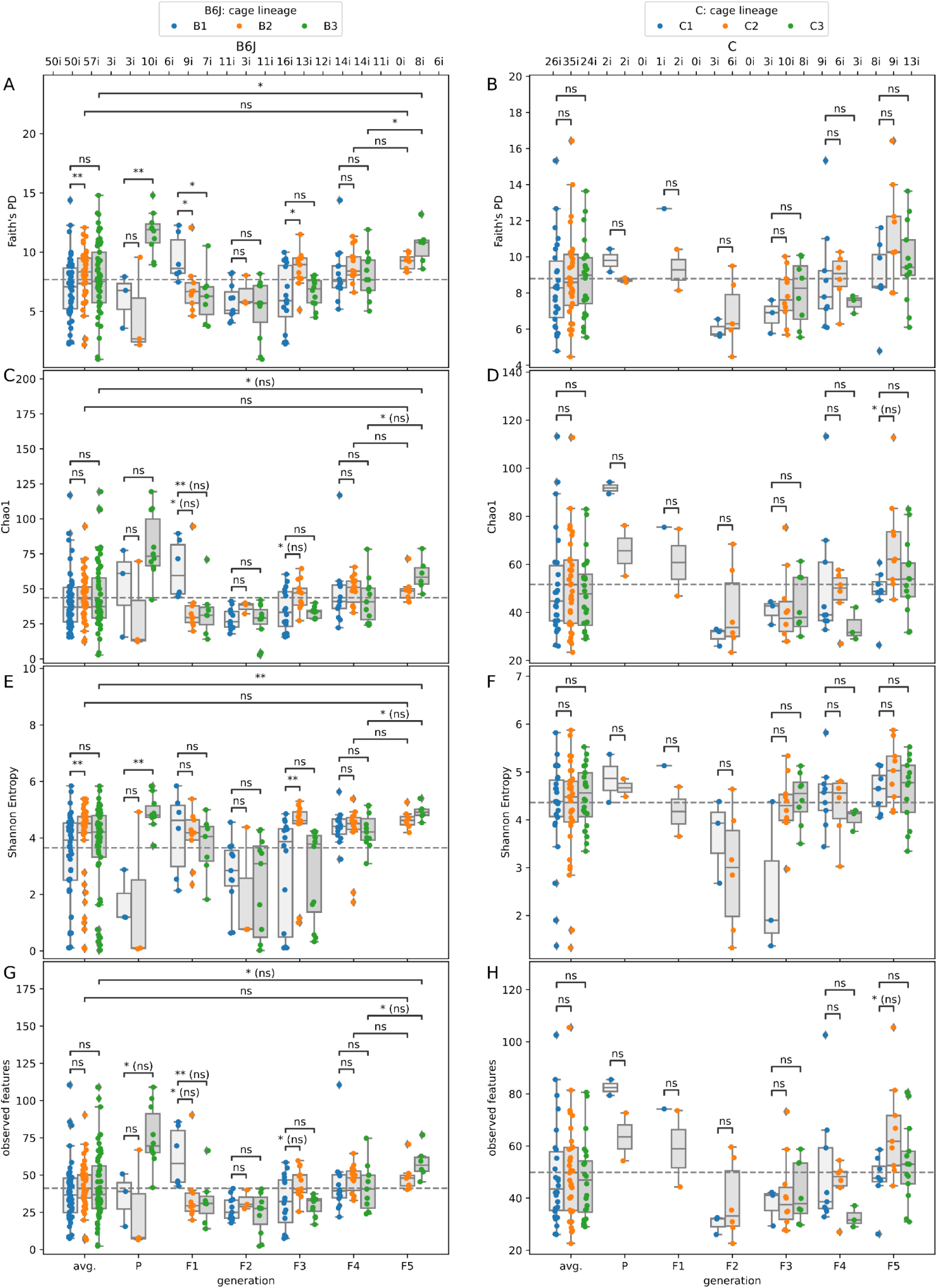
Individual Alpha Diversity of Skin samples. Left and right columns display alpha diversity of B6J and C samples, respectively. Rows are different alpha diversity metrics: Faith’s PD, Chao1, Shannon Entropy and number of observed features, i.e. ASVs. The x-axis stratifies samples into generations, while hue indicates the three different cage lineages. Leftmost x-axis positions, entitled “avg.” are samples from all generations lumped together. We used two-sided Mann-Whitney-Wilcoxon tests with Benjamini-Hochberg correction to assess statistical significance.

**Figure S6:**
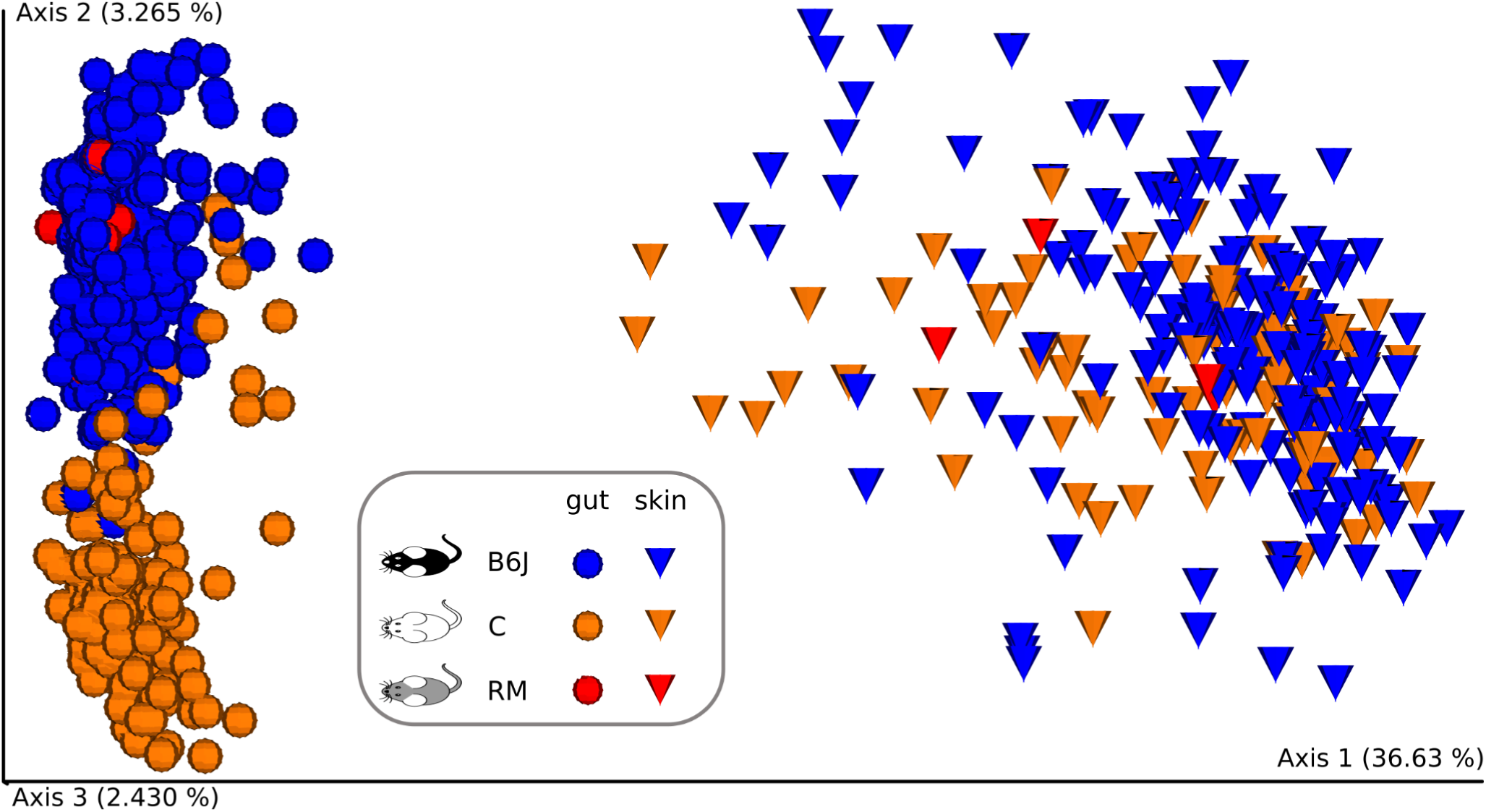
Differences in body sites. PCoA of samples of the gut n=333 (spheres) and of the external skin of the left earlobe n=268 (cones) based on unweighted UniFrac shown in two clusters. The samples are additionally stratified by their host genotype showing B6J (blue), C (orange), and B6CF1 (red) mice, which form two clusters in the gut samples and one cluster in the skin samples.

**Figure S7:**
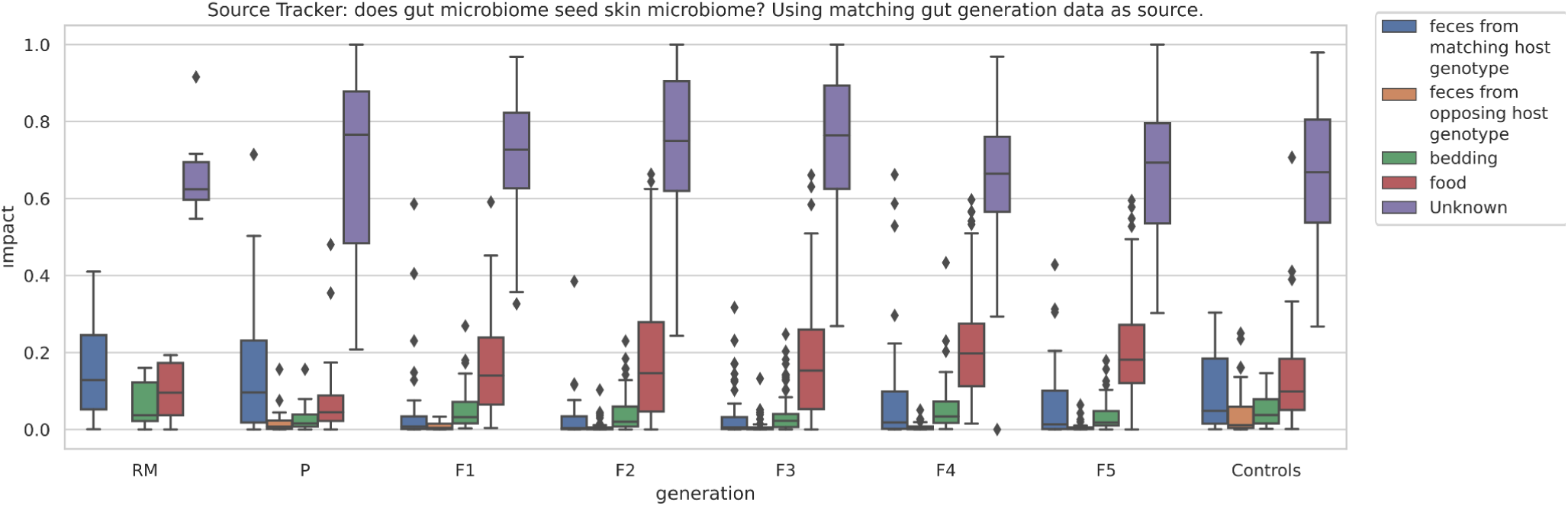
Impact of External Factors on the Skin Microbiome. We used SourceTracker to investigate whether the external factors of bedding (green) or food (red) had an impact on the skin microbiome of the mice. Additionally, we included if mice were impacted by microbiota from feces from matching host genotype“host genotype” (blue) or from opposing host genotype“host genotype” (orange). For variables outside the assigned categories SourceTracker assigns the label Unknown (purple). This analysis was stratified by generations as well as adding the control mice.

**Figure S8:**
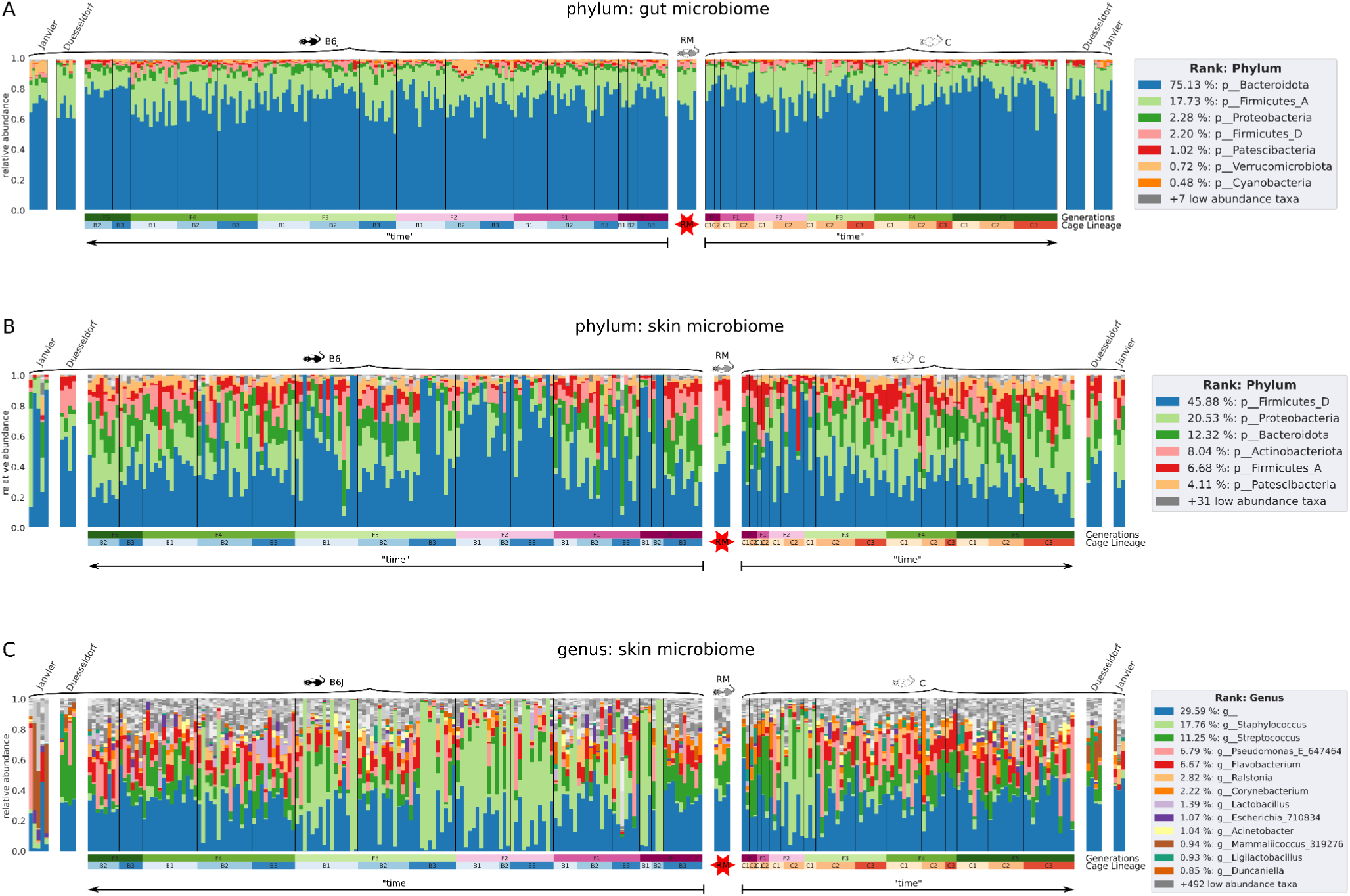
Taxonomy Barplots. A) Phylum composition of 333 gut samples, rarefied to 6000 reads. Purple to green bar indicates generations, blueish and orange bar indicate cage lineage. Six samples of recipient mothers (RM) of hybrid host genotype“host genotype” B6CF1 (gray) are marked by the red star. Offspring generations (P to F5) are temporarily aligned to the left for B6J (black) and to the right for C (white), respectively; followed by six control samples each from mice obtained from another facility Duesseldorf (Germany) and ordered from Janvier (France). Cage lineages B1-B3 and C1-C3 are color coded in blue and orange, respectively. B) Phylum composition of 2628 external skin of the left earlobe samples, rarefied to 1000 reads. C) Same as B, but on Genus level.

**Figure S9:**
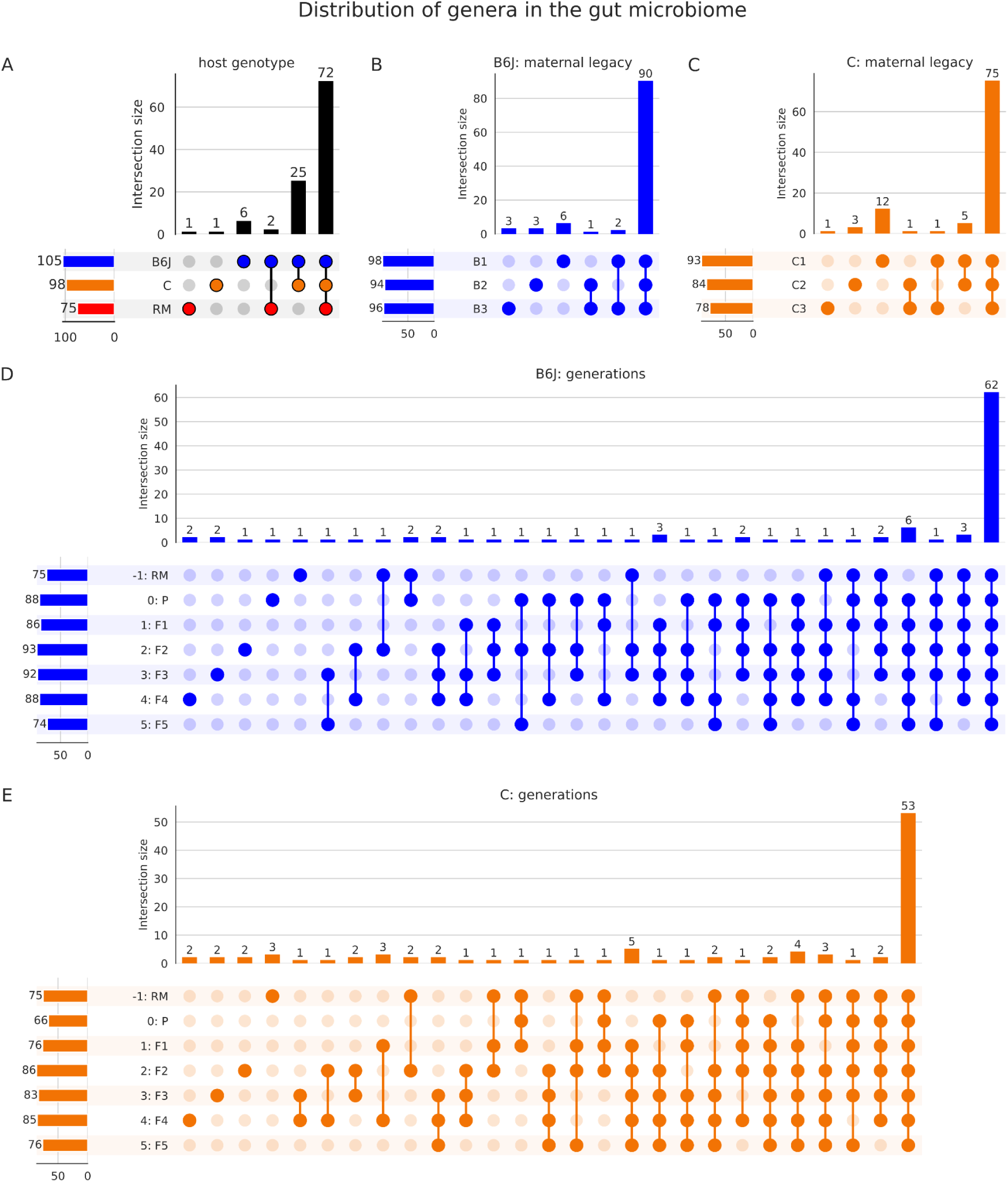
Distribution of Taxa in the gut. To investigate the size of the core microbiome and overlapping taxa in the gut we created Upsetplots. The intersection size indicates how many taxa exist in a group, the circles show which categories the group consists of. The bars on the left sum up the total amount of taxa per category. A) Upsetplot comparing the “host genotypes” RM, C and B6J. B) This Upsetplot shows the distribution of taxa from the perspective of maternal legacy of the B6J mice. C) Analogous to Panel B but for C mice. D) This Upsetplot shows B6J and RM samples sorted by generation including RM. E) Analogous to Panel D but for C mice.

**Figure S10:**
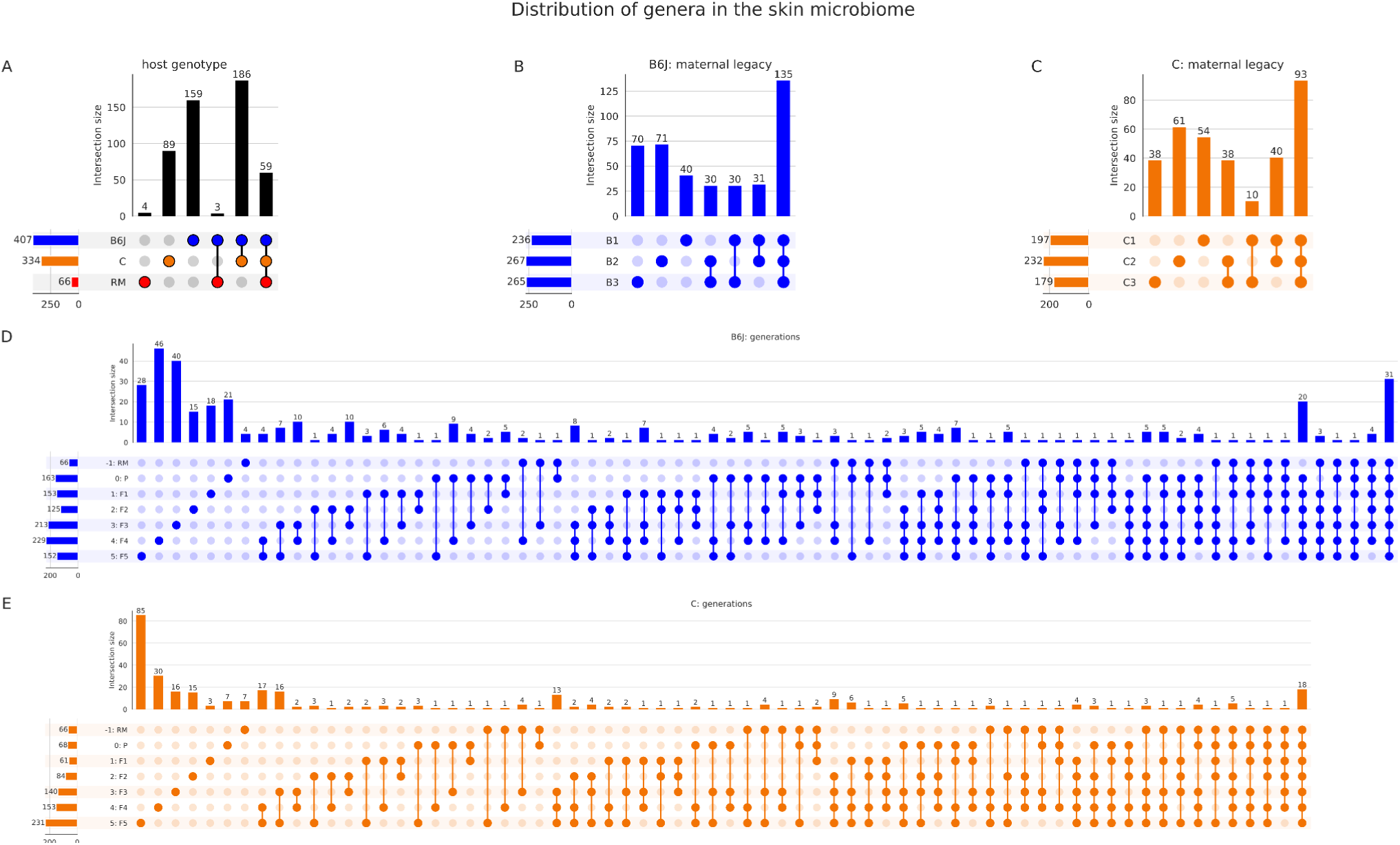
Distribution of Taxa in the Skin. To investigate the size of the core microbiome and overlapping taxa in the skin we created Upsetplots. The intersection size indicates how many taxa exist in a group, the circles show which categories the group consists of. The bars on the left sum up the total amount of taxa per category. A) Upsetplot comparing the “host genotypes” RM, C and B6J. B) This Upsetplot shows the distribution of taxa from the perspective of “maternal legacy” of the B6J mice. C) Analogous to Panel B but for C mice. D) This Upsetplot shows B6J and RM samples sorted by generation including RM. E) Analogous to Panel D but for C mice.

## Additional File 1

**Metabolites table.** Measures targeted metabolites for 40 blood serum samples (220511 Data_21-0025.xlsx).

## Additional File 2

**Analysis repository.** We performed all the presented analysis and graph generation through a single jupyter notebook. It lists necessary dependencies for full reproducibility. We have outsourced some of the functions into a public 16s rRNA gene analysis code repository: https://github.com/sjanssen2/ggmap. By default, external system calls are submitted to a Slurm cluster. You can instead run on your local machine by providing the optional argument `use_grid=False`. Additional conda environments can be conveniently created via “cd recipes && make all-env” after cloning the above repository and changing into its working directory. All files necessary for our analysis, the jupyter notebook and a static HTML version of the jupyter notebook for ease of readability are packaged as one zip compressed file (microbiome_benga_hostgenotype.zip).

