## Additional File 2: Analysis repository for "The host genotype actively shapes its microbiome across generations in laboratory mice": benga_hostgenotype.ipynb

```
# %% [markdown]
If you intent to re-execute this notebook, we recommend to
  1. create one conda environment into which you install the ggmap software collection: https://github.com/sjanssen2/ggmap/tree/master
  2. additionally install conda packages
     1. statannot
     2. tqdm
     3. upsetplot
     4. calour
     5. matplotlib-venn
  3. create all additional conda environments, necessary for data processing:
     1. `cd` into `ggmap/recipes`
     2. use makefile to create environments: `make all-env`

# %% [code]
# toggle between "notebook" and "inline", if graphs are not directly displayes
%matplotlib inline

# %% [code]
%load_ext autoreload
%autoreload 2

from matplotlib.patches import ConnectionPatch
from matplotlib.legend import Legend
from matplotlib.patches import Rectangle
from matplotlib_venn import venn2
from matplotlib import gridspec
import matplotlib.cm as cm

from IPython.display import Image
from IPython.core.display import HTML 

from collections import OrderedDict

from statannot import add_stat_annotation
from statannotations.Annotator import Annotator

import upsetplot
from skbio import TabularMSA, DNA
from tqdm import tqdm

import calour as ca
ca.set_log_level(40)

from ggmap.workflow import *

# %% [code]
settings.GRID_ACCOUNT = 'bcf'  # use specific Slurm cluster partition

# %% [code]
# an ugly mechanism to list my currently installes packages
cluster_run(['conda list > conda_list', 'rm -f dummy'], 
            'get_packages', 
            './dummy', 
            dry=False, use_grid=False, environment='notebookServer')
!cat conda_list

# %% [code]
# also report settings in the personal ~/.ggmaprc file
!cat $HOME/.ggmaprc

# %% [markdown]
## Definitions

# %% [code]
NA_VALUES = ['not applicable', 'not collected', 'not provided']
ORDER_GENERATIONS = ['RM','P','F1','F2','F3','F4','F5','Controls']
KEY_GENERATIONS = lambda x: x.apply(lambda a: ORDER_GENERATIONS.index(a))
ORDER_TAXPLOT = ['Janvier B6J', 'Duesseldorf B6J', 'F5_B2', 'F5_B3', 'F4_B1', 'F4_B2', 'F4_B3', 'F3_B1', 'F3_B2', 'F3_B3', 'F2_B1', 'F2_B2', 'F2_B3', 'F1_B1', 'F1_B2', 'F1_B3', 'P_B1', 'P_B2', 'P_B3', 'RM', 
                 'P_C1', 'P_C2', 'F1_C1', 'F1_C2', 'F2_C1', 'F2_C2', 'F3_C1', 'F3_C2', 'F3_C3', 'F4_C1', 'F4_C2', 'F4_C3', 'F5_C1', 'F5_C2', 'F5_C3', 'Duesseldorf C', 'Janvier C', ]
KEY_TAXPLOT = lambda x: x.apply(lambda a: ORDER_TAXPLOT.index(a))
ORDER_CONTROLS = ['Duesseldorf', 'Janvier']

ORDER_GENOTYPE = ['B6J', 'C']
COLORS = {
    'B6J': '#0000ff',
    'C': '#f27304',
    "RM": "#ff0000",
    
    'P': '#8e0152',
    'F1': '#d6589e',
    'F2': '#f5c4e1',
    'F3': '#c7e89f',
    'F4': '#6eae36',
    'F5': '#276419',
    
    'mJAX': '#ff0000',
    'mTAC': '#0000ff',
    
    "B1": "#deebf7", 
    "B2": "#9ecae1", 
    "B3": "#3182bd",
    "C1": "#fee8c8",
    "C2": "#fdbb84",
    "C3": "#e34a33",
}
DM_NAMES = {
    'PD_whole_tree': "Faith's PD",
    'unweighted_unifrac': 'unweighted UniFrac',
    'weighted_unifrac': 'weighted UniFrac',
    'observed_features': 'observed features',
    'shannon': 'Shannon Entropy',
    'bray_curtis': 'Bray-Curtis',
    'chao1': 'Chao1',
    'jaccard': 'Jaccard',
}

def alpha_blending(hex_color, alpha) :
    """ alpha blending as if on the white background.
    """
    foreground_tuple  = matplotlib.colors.hex2color(hex_color)
    foreground_arr = np.array(foreground_tuple)
    final = tuple( (1. -  alpha) + foreground_arr*alpha )
    return(final)

def _sample_level(sample, field='cage_line', just_host_genotype=False):
    """Assign sample categories based on multiple selected metadata fields"""
    if pd.notnull(sample[field]):
        if just_host_genotype:
            return sample['host_genotype']
        else:
            return sample[field]
    if sample['host_genotype'] == 'RM':
        return sample['host_genotype']
    if pd.notnull(sample['control']):
        return '%s %s' % (sample['control'], sample['host_genotype'])


# %% [markdown]
# Primary Analyses

# %% [markdown]
## Obtain data

Data are managed with Qiita as the public study with ID 13422. You can obtain all necessary data through three alternative ways:

(Note: the ITS sequencing to profile fungi was not successful, i.e. too low read qualities. Ignore this preparation in Qiita!)

1. per sample forward and reverse fastq.gz files from ENA, accession LINK
2. metadata and precomputed feature table (bacteria * samples) from our public Qiita study: [https://qiita.ucsd.edu/study/description/13422](https://qiita.ucsd.edu/study/description/13422). fastq.gz files are also available through Qiita.
3. **metadata and precomputed feature table (bacteria * samples) via [github](https://github.com/jlab/microbiome_benga_multiplegenerations), which are actually downloaded from Qiita for your convenience**

The per sample forward and reverse fastq.gz result from: base calling and simultaneous demultiplexing was done via bcl2fastq (v2.19.0.316) and primers were trimmed using cutadapt (v2.10, Martin 2011).

# %% [code]
# obtain metadata from Qiita as the combination of "Sample Info" and "Prep Info"
metadata = pd.concat([pd.read_csv("FromQiita/%s" % filename, sep="\t", dtype=str, na_values=NA_VALUES, index_col=0)
                      for filename in ["13422_prep_11649_20240705-141258.txt",  # Prep Info
                                       "13422_20231204-091505.txt"              # Sample Info
                                      ]], axis=1)

# make values true numbers
metadata['serum_triglyceride'] = metadata['serum_triglyceride'].astype(float)

metadata['gave_birth'] = metadata['gave_birth'].apply(lambda x: x == 'True')
metadata['mate_sire'] = metadata['mate_sire'].apply(lambda x: x == 'True')

# use abbreviations for "host genotype" as in the manuscript
metadata['host_genotype'] = metadata['host_genotype'].apply(lambda x: {'c57bl6/j': 'B6J',
                                                                       'balb/c': 'C',
                                                                       'b6cf1': 'RM'}.get(x, x))
metadata['cage_line'] = metadata['cage_line'].apply(lambda x: {'C57BL6/J_line1': 'B1',
                                                               'C57BL6/J_line2': 'B2',
                                                               'C57BL6/J_line3': 'B3',
                                                               'BALB/c_line4': 'C1',
                                                               'BALB/c_line5': 'C2',
                                                               'BALB/c_line6': 'C3',}.get(x,x))

# give locations shorter names
metadata['geo_loc_name'] = metadata['geo_loc_name'].apply(lambda x: {'France:Janvier': 'Janvier',
                                                                     'Germany:Duesseldorf': 'Duesseldorf'}.get(x,x))

# provide extra field to easily identify control (Janvier / Duesseldorf) samples
metadata['control'] = metadata[(metadata['host_scientific_name'] == 'Mus musculus') &
                               ((metadata['geo_loc_name'] == 'Janvier') |
                               (metadata['iacuc_institute'] == 'ZETT'))]['geo_loc_name'].apply(lambda x: {'Janvier': 'Janvier',
                                                                                                          'Duesseldorf': 'Duesseldorf'}.get(x, np.nan))

metadata['sample_category'] = metadata.apply(lambda x: _sample_level(x, 'cage_line', True), axis=1)
metadata['sample_category_cagelines'] = metadata.apply(lambda x: _sample_level(x, 'cage_line', False), axis=1)
metadata['sample_category_generation'] = metadata.apply(lambda x: _sample_level(x, 'generation', False), axis=1)
metadata['generation_cageline'] = metadata.apply(lambda row: '%s%s' % (row['generation'] if pd.notnull(row['generation']) else row['sample_category'],
                                                                       '_' + row['cage_line'] if pd.notnull(row['cage_line']) else ''), axis=1)
metadata['independent_cage_lineage'] = metadata['cage_line'].apply(lambda x: x[-1] if pd.notnull(x) else x)

metadata['serum_triglyceride'] = metadata['serum_triglyceride'].astype(float)

# %% [code]
# subset metadata to gut and skin, as we rarefy these samples differently to account for different 
# read coverage due to low biomass in skin

meta_gut = metadata[(metadata['host_scientific_name'] == 'Mus musculus') &  # ignore marmoset control samples
                    (metadata['host_body_site'] == 'UBERON:colon')]         # only select gut

# control samples are necessary for decontam filtering
meta_skin_withControls = metadata[((metadata['host_scientific_name'] == 'Mus musculus') & # ignore marmoset control samples
                                  (metadata['host_body_site'] == 'UBERON:sebum') &       # only select skin
                                  (metadata['batch'] != '2')) |                          # 12 skin samples have been sequenced twice, just use the better quality run
                                  (metadata['sample_type'] == 'sterile water')           # use sterile water samples as negative controls for decontam
                                 ]

# same for the feature table
counts = biom2pandas("FromQiita/128106_reference-hit.biom")
counts_gut = counts.loc[:, meta_gut.index]
counts_skin = counts.loc[:, [s for s in meta_skin_withControls.index if s in counts.columns]]

# %% [markdown]
## Rarefy, assign taxonomy & compute alpha- and beta- diversities

# %% [code]
sepp_tree = TreeNode.read("FromQiita/128106_insertion_tree.relabelled.tre", format="newick")

# %% [markdown]
**Note:** rarefaction is an inherently randomized process, i.e. if you re-compute from scratch, you will probably find small differences to the published version.

In order to provide you with almost identical results, we packaged cached results that are affected by rarefaction into the repository. Should you NOT want to use them, remove all file in the hidden subdirectory `.anacache`.

# %% [code]
res_gut = process_study(
    metadata=meta_gut,
    control_samples=set([]),  # we already filtered out all control samples
    fp_deblur_biom=counts_gut,  # "abuse" argument to provide a table instead of a filepath
    fp_insertiontree="FromQiita/128106_insertion_tree.relabelled.tre",
    rarefaction_depth=6000,
    fp_taxonomy_trained_classifier_gg138_chloroMitoRemoval='/vol/jlab/MicrobiomeAnalyses/References/Q2-Naive_Bayes_classifiers/gg-13-8-99-nb-classifier_2022.11.qza',  # as GG2 currently lacks labels for mitochondria/chloroplasts, see https://forum.qiime2.org/t/taxonomy-filtering-greengenes2/28334
    fp_taxonomy_trained_classifier="/vol/jlab/MicrobiomeAnalyses/References/Q2-Naive_Bayes_classifiers/gg_2022_10_backbone_full_length.nb_2023.5.qza",  # obtain this file from https://docs.qiime2.org/2023.9/data-resources/
    tree_insert=sepp_tree,
    is_v4_region=False,
    emperor_fp='Generated/Emperor',
    emperor_infix='_gut',
    dry=False,
    use_grid=True,
    fix_zero_len_branches=False, #'' since this has already been corrected via the project_sepp call
    emperor_skip_tsne_umap=True,
    
    beta_metrics=["unweighted_unifrac", "weighted_unifrac", "bray_curtis", "jaccard"],  # reviewer 1 specifically requested jaccard
    alpha_metrics=["PD_whole_tree", "shannon", "observed_features", "chao1"],           # reviewer 1 specifically requested chao1
)

# %% [code]
sns.set_style("whitegrid")
res_skin = process_study(
    metadata=meta_skin_withControls,
    control_samples=set(meta_skin_withControls[meta_skin_withControls['sample_type'] == 'sterile water'].index),
    fp_deblur_biom=counts_skin,  # "abuse" argument to provide a table instead of a filepath
    fp_insertiontree="FromQiita/128106_insertion_tree.relabelled.tre",
    rarefaction_depth=1000,
    fp_taxonomy_trained_classifier_gg138_chloroMitoRemoval='/vol/jlab/MicrobiomeAnalyses/References/Q2-Naive_Bayes_classifiers/gg-13-8-99-nb-classifier_2022.11.qza',  # as GG2 currently lacks labels for mitochondria/chloroplasts, see https://forum.qiime2.org/t/taxonomy-filtering-greengenes2/28334
    fp_taxonomy_trained_classifier="/vol/jlab/MicrobiomeAnalyses/References/Q2-Naive_Bayes_classifiers/gg_2022_10_backbone_full_length.nb_2023.5.qza",  # obtain this file from https://docs.qiime2.org/2023.9/data-resources/
    tree_insert=sepp_tree,
    is_v4_region=False,
    emperor_fp='Generated/Emperor',
    emperor_infix='_skinWithControls',
    dry=False,
    use_grid=True,
    fix_zero_len_branches=False, # since this has already been corrected via the project_sepp call
    emperor_skip_tsne_umap=True,
    
    decontam_col_concentration='concentration_ng_myl',
    decontam_col_sampletype='sample_type',
    decontam_name_control_sample='sterile water',
    decontam_lowbiomass_envs=['skin of external ear'],
    decontam_cols_batch=['flowcell', 'plate_number'],
    
    beta_metrics=["unweighted_unifrac", "weighted_unifrac", "bray_curtis", "jaccard"],  # reviewer 1 specifically requested jaccard
    alpha_metrics=["PD_whole_tree", "shannon", "observed_features", "chao1"],           # reviewer 1 specifically requested chao1
)

# %% [code]
# use DECONTAM treated feature-table for skin samples
counts_nokitome = pd.concat([
    counts.loc[:, [s for s in counts.columns if s not in res_skin['counts_decontaminated'].columns]],
    res_skin['counts_decontaminated']], axis=1)

idx_control_samples = set(metadata[metadata['sample_type'].isin(['sterile water', 'control bedding', 'control food'])].index)
res_both = process_study(
    metadata=metadata,
    control_samples=idx_control_samples,
    fp_deblur_biom=counts_nokitome,  # "abuse" argument to provide a table instead of a filepath
    fp_insertiontree="FromQiita/128106_insertion_tree.relabelled.tre",
    rarefaction_depth=1000,
    fp_taxonomy_trained_classifier_gg138_chloroMitoRemoval='/vol/jlab/MicrobiomeAnalyses/References/Q2-Naive_Bayes_classifiers/gg-13-8-99-nb-classifier_2022.11.qza',  # as GG2 currently lacks labels for mitochondria/chloroplasts, see https://forum.qiime2.org/t/taxonomy-filtering-greengenes2/28334
    fp_taxonomy_trained_classifier="/vol/jlab/MicrobiomeAnalyses/References/Q2-Naive_Bayes_classifiers/gg_2022_10_backbone_full_length.nb_2023.5.qza",  # obtain this file from https://docs.qiime2.org/2023.9/data-resources/
    tree_insert=sepp_tree,
    is_v4_region=False,
    emperor_fp='Generated/Emperor',
    emperor_infix='_both',
    dry=False,
    use_grid=True,
    fix_zero_len_branches=False, # since this has already been corrected via the project_sepp call
    emperor_skip_tsne_umap=True,
)

# %% [code]
# drop samples from metadata which did not pass QC, i.e. rarefaction
meta_gut = meta_gut.loc[res_gut['rarefaction']['results'].columns, :]
meta_skin = meta_skin_withControls.loc[list(set(meta_skin_withControls[meta_skin_withControls['sample_type'] == 'skin of external ear'].index) & 
                                            set(res_skin['rarefaction']['results'].columns)),:]

# %% [code]
# generate PCoA for skin samples, but without blank controls
res_skin_emp = emperor(meta_skin, res_skin['beta_diversity']['results'], 'Generated/Emperor', infix='_skin', dry=False, wait=False)

# %% [markdown]
## Figure 2
### Panels A,B,C

# %% [code]
def get_sample_numbers(meta):
    # collect number of samples: individuals, cages, cage_lines
    assert meta['host_subject_id'].value_counts().max() == 1
    
    numSamples = pd.concat([
        # breeding experiment
        pd.concat([
            meta.groupby(['generation', 'host_genotype']).size().rename('i'),  # individuals
            meta.groupby(['generation', 'host_genotype'])['cage_id'].unique().apply(len).rename('c'),  # cages
            meta.groupby(['generation', 'host_genotype'])['cage_line'].unique().apply(len).rename('l')  # cage-lines
        ], axis=1),
        # controls
        pd.concat([
            meta[pd.isnull(meta['generation'])].groupby(['geo_loc_name', 'host_genotype']).size().rename('i'),  # individuals
            meta[pd.isnull(meta['generation'])].groupby(['geo_loc_name', 'host_genotype'])['sex'].unique().apply(len).rename('c'),  # cages. Use sex as a proxy as we actually don't know the cage IDs
            meta[pd.isnull(meta['generation'])].groupby(['geo_loc_name', 'host_genotype'])['cage_line'].unique().apply(lambda x: np.nan).rename('l')  # we did not use cage lines for controls
        ], axis=1)
    ])
    
    return numSamples

def get_sample_label(grp, numSamples, genotypes=ORDER_GENOTYPE, justI=False):
    label = ""
    if grp == 'RM':
        genotypes = ['RM']
    for genotype in genotypes:
        cat = dict()
        for category in ['i','c','l']:
            if (grp, genotype) in numSamples.index:
                number = numSamples.loc[(grp, genotype), category]
            else:
                number = 0
            if pd.notnull(number):
                cat[category] = '%2i%s' % (number, category)
            else:
                cat[category] = number
        if justI:
            ns = '%s' % cat['i']
        else:
            if pd.isnull(cat['l']):
                ns = '%s (%s)' % (cat['i'], cat['c'])
            else:
                ns = '%s (%s,%s)' % (cat['i'], cat['c'], cat['l'])
        label += "\n" + ns
    return label

def compute_beta_distances(res_beta, meta, metric='weighted_unifrac'):
    results = dict()

    dm = res_beta[metric].to_data_frame()
    data = []
    for refgrp, g_ref in meta[~meta['sample_category'].isin(['C'])].groupby('sample_category'):
        for (host_genotype, generation), g in meta[meta['sample_category'].isin(['C','B6J','RM'])].groupby(['sample_category', 'generation']):
            subset_g_ref = g_ref
            if refgrp == host_genotype:
                continue
            if refgrp == 'B6J':
                subset_g_ref = g_ref[g_ref['generation'] == generation]
            assert len(set(subset_g_ref.index) & set(g.index)) == 0
            d = dm.loc[subset_g_ref.index, g.index]
            d.index.name = 'idx_reference'
            d.columns.name = 'idx_sample'
            d = d.stack().to_frame().reset_index().rename(columns={0: metric})
            d['generation'] = generation
            d['host_genotype'] = host_genotype
            d['reference'] = refgrp
            d['reference_host_genotype'] = refgrp.split()[-1]
            d['reference_host_geolocname'] = refgrp.split()[0]
            data.append(d)
    results['beta'] = pd.concat(data).sort_values(by='generation', key=KEY_GENERATIONS)
    
    cids = [cid for (cid, size) in meta.groupby('cage_id').size().items() if size > 1]
    subset = meta[meta['cage_id'].isin(cids)]
    results['intra_cage'] = {'num_cages': len(cids),
                             'sum_mice': subset.shape[0],
                             'avg_mice_per_cage': subset.groupby('cage_id').size().mean(),
                             'avg_%s' % metric: np.mean([dm.loc[a,b] for cage_id, g in subset.groupby('cage_id') for (a,b) in combinations(g.index, 2)])}

    distances = []
    for (generation, host_genotype, cage_line), g in meta.groupby(['generation', 'host_genotype', 'cage_line']):
        for (cageA, cageB) in combinations(g['cage_id'].unique(), 2):
            d = dm.loc[g[g['cage_id'] == cageA].index, g[g['cage_id'] == cageB].index]
            d.index.name = 'cageA'
            d.columns.name = 'cageB'
            d = d.stack().reset_index().rename(columns={0: metric})
            d[['generation', 'host_genotype', 'cage_line']] = [generation, host_genotype, cage_line]
            distances.append(d)
    distances = pd.concat(distances)
    results['intra_cage_line'] = {'num_cages': meta.groupby(['generation', 'host_genotype', 'cage_line'])['cage_id'].unique().apply(len).sum(),
                                  'avg_cages': meta.groupby(['generation', 'host_genotype', 'cage_line'])['cage_id'].unique().apply(len).mean(),
                                  'sum_mice_mice': distances.shape[0],
                                  'avg_mice_mice': distances.groupby(['generation', 'host_genotype', 'cage_line']).size().mean(),
                                  'avg_%s' % metric: distances[metric].mean(),
                                 }
    return results


def draw_temporals(res, meta, 
                   metric_alpha = 'PD_whole_tree',
                   METRIC_BETA = 'weighted_unifrac', 
                   ALPHA_LIM = (14.5,21.8), 
                   BETA_LIM=(0.085,0.169),
                   title='gut microbiome', num_permutations=999):
    #sns.set_style("white")
    numSamples = get_sample_numbers(meta)
    
    alpha = meta.merge(res['alpha_diversity']['results'], left_index=True, right_index=True)
    beta = compute_beta_distances(res['beta_diversity']['results'], meta, METRIC_BETA)

    sns.reset_orig()
    fig, axes = plt.subplots(3, 3, figsize=(15*1.4, 15), gridspec_kw={'width_ratios': [1, 6, 2]})
    fig.subplots_adjust(wspace=0)

    sns.boxplot(data=alpha[alpha['generation'] == 'RM'],
                y=metric_alpha, x='host_genotype',
                ax=axes[0,0], color=alpha_blending(COLORS['RM'], 0.4), width=0.4)
    sns.swarmplot(data=alpha[alpha['generation'] == 'RM'],
                  y=metric_alpha, x='host_genotype', 
                  ax=axes[0,0], color=COLORS['RM'], size=8, linewidth=1, edgecolor='white')


    data = alpha[(pd.notnull(alpha['generation'])) & (alpha['generation'] != 'RM')].sort_values(by='generation', key=KEY_GENERATIONS)
    sns.lineplot(data=data,
                 y=metric_alpha, x='generation', hue='host_genotype', 
                 ax=axes[0,1],sort=False, palette=COLORS, markers=True, marker="o")

    annotator = Annotator(axes[0,1], [
            ((generation, 'B6J'), (generation, 'C')) for generation in data['generation'].unique() if ((generation, 'B6J') in numSamples.index and numSamples.loc[(generation, 'B6J'), 'i'] > 0) and ((generation, 'C') in numSamples.index and numSamples.loc[(generation, 'C'), 'i'] > 0)
        ], data=data, y=metric_alpha, hue='host_genotype', x='generation')
    annotator.configure(test='Mann-Whitney', text_format='star', loc='outside', comparisons_correction="fdr_bh", correction_format="default")
    annotator.apply_and_annotate()


    ax = axes[0,2]
    data = alpha[pd.isnull(alpha['generation'])]
    sns.boxplot(data=data,
                y=metric_alpha, x='geo_loc_name', order=ORDER_CONTROLS, hue='host_genotype', 
                ax=axes[0,2], palette={n:alpha_blending(c,0.4) for n,c in COLORS.items()}, hue_order=ORDER_GENOTYPE)
    sns.swarmplot(data=data,
                  y=metric_alpha, x='geo_loc_name', order=ORDER_CONTROLS, hue='host_genotype', 
                  ax=axes[0,2], palette=COLORS, dodge=True, size=8, linewidth=1, edgecolor='white', hue_order=ORDER_GENOTYPE)
    annotator = Annotator(axes[0,2], [
            (('Duesseldorf', 'B6J'), ('Duesseldorf', 'C')),
            (('Janvier', 'B6J'), ('Janvier', 'C'))
        ], data=data, y=metric_alpha, hue='host_genotype', x='geo_loc_name', order=ORDER_CONTROLS)
    annotator.configure(test='Mann-Whitney', text_format='star', loc='outside', comparisons_correction="fdr_bh", correction_format="default")
    annotator.apply_and_annotate()

    with warnings.catch_warnings():
        warnings.simplefilter("ignore")
        # ignore warning, as we do not split data into hue category
        annotator = Annotator(axes[0,2], [
                    ('Duesseldorf', 'Janvier')
                ], data=data, y=metric_alpha, x='geo_loc_name')
    annotator.configure(test='Mann-Whitney', text_format='star', loc='outside', comparisons_correction="fdr_bh", correction_format="default")
    annotator.apply_and_annotate()

    axes[0,1].add_artist(ConnectionPatch(xyA=(1,ALPHA_LIM[0]+1), xyB=(0,ALPHA_LIM[0]+1), coordsA="data", coordsB="data",
                          axesA=axes[0,0], axesB=axes[0,1], color="black"))
    axes[0,0].text(0, ALPHA_LIM[0]+0.7, "p=%.3f" % kruskal(alpha[alpha['generation'] == 'RM'][metric_alpha], alpha[(alpha['generation'] == 'P')][metric_alpha]).pvalue)


    # figure aestetics
    axesTop = [axes[0,i].twiny() for i in range(len(axes[0]))]
    axesTop[0].set_xticks([0.5], ['%s\n\n' % get_sample_label('RM', numSamples)])
    axesTop[1].set_xlim(axes[0,1].get_xlim())
    axesTop[1].set_xticks(axes[0,1].get_xticks())
    axesTop[1].set_xticklabels(['\n%s' % get_sample_label(t.get_text(), numSamples) for t in axes[0,1].get_xticklabels()])
    axesTop[2].set_xlim(axes[0,2].get_xlim())
    offset = 0.2
    axesTop[2].set_xticks([0-offset,0+offset, 1-offset, 1+offset], 
                          ['\n%s\n' % get_sample_label('Duesseldorf', numSamples, genotypes=['B6J']),
                           '\n\n%s' % get_sample_label('Duesseldorf', numSamples, genotypes=['C']),
                           '\n%s\n' % get_sample_label('Janvier', numSamples, genotypes=['B6J']),
                           '\n\n%s' % get_sample_label('Janvier', numSamples, genotypes=['C'])])

    axes[0,2].set_title('Controls')
    axes[0,2].legend().remove()
    for ax in axes[0]:
        # same y limits for all three subplots
        ax.set_ylim(ALPHA_LIM)
        # add horizontal grid lines ...
        ax.set_axisbelow(True)
        # ... behind the plots
        ax.grid(axis='y')
        # no x axis label, as we already have tick labels
        ax.set_xlabel("")

        if ax == axes[0,0]:
            # nice label for Y axis
            ax.set_ylabel('α diversity\n%s' % DM_NAMES.get(metric_alpha))
        else:
            ax.set_ylabel("")
            ax.set_yticklabels([])

           
    ## SECOND ROW: beta diversity distances to RM
    for ax in [axes[1,0], axes[1,1]]:
        ax.axhline(y=beta['intra_cage']['avg_%s' % METRIC_BETA], linestyle='--', label='avg. intra cage', color=sns.color_palette()[7])
        ax.axhline(y=beta['intra_cage_line']['avg_%s' % METRIC_BETA], linestyle='dashdot', label='avg. intra cage linege', color=sns.color_palette()[6])

    axes[1,1].text(0.2, beta['intra_cage']['avg_%s' % METRIC_BETA]*1.02, '%ic (Σ %ii, Ø %.1fi)' % (
        beta['intra_cage']['num_cages'], 
        beta['intra_cage']['sum_mice'], 
        beta['intra_cage']['avg_mice_per_cage']))
    axes[1,1].text(3.2, beta['intra_cage_line']['avg_%s' % METRIC_BETA]*1.02, 'Σ %ic-c, Ø %.1fc-c  Σ %ii-i, Ø %.1fi-i' % (
        beta['intra_cage_line']['num_cages'], 
        beta['intra_cage_line']['avg_cages'], 
        beta['intra_cage_line']['sum_mice_mice'],
        beta['intra_cage_line']['avg_mice_mice']))

    data = beta['beta'][beta['beta']['reference'] == 'RM']
    sns.lineplot(data=data, y=METRIC_BETA, x='generation', hue='host_genotype', ax=axes[1,1], sort=False, palette=COLORS, markers=True, marker="o", hue_order=ORDER_GENOTYPE)
    
    annotator = Annotator(axes[1,1], [
        ((generation, 'B6J'), (generation, 'C')) for generation in data['generation'].unique() if ((generation, 'B6J') in numSamples.index and numSamples.loc[(generation, 'B6J'), 'i'] > 0) and ((generation, 'C') in numSamples.index and numSamples.loc[(generation, 'C'), 'i'] > 0)
        ], data=data, y=METRIC_BETA, hue='host_genotype', x='generation')
    annotator.configure(test='Mann-Whitney', text_format='star', loc='outside', comparisons_correction="fdr_bh", correction_format="default")
    annotator.apply_and_annotate()
    data = beta['beta'][beta['beta']['reference'] == 'B6J']
    sns.lineplot(data=data, y=METRIC_BETA, x='generation', ax=axes[1,1], sort=False, markers=True, marker="o", color='green', label='CG')

    axes[1,1].set_ylabel("")
    axes[1,1].legend().remove()
    
    idx_perm = meta[meta['generation'].isin(['RM', 'P'])].index
    res_perm = permanova(res['beta_diversity']['results'][METRIC_BETA].filter(idx_perm), meta.loc[idx_perm,:], column='generation', permutations=num_permutations)
    axes[1,1].add_artist(ConnectionPatch(xyA=(0.5, np.mean(list(BETA_LIM))), xyB=(0, np.mean(list(BETA_LIM))), coordsA="data", coordsB="data",
                      axesA=axes[1,0], axesB=axes[1,1], color="black"))
    axes[1,0].text(0, np.mean(list(BETA_LIM)), "PERMANOVA p=%.3f" % res_perm['p-value'], horizontalalignment="center")


    for ax in axes[1]:
        # same y limits for all three subplots
        ax.set_ylim(BETA_LIM)
        # add horizontal grid lines ...
        ax.set_axisbelow(True)
        # ... behind the plots
        ax.grid(axis='y')
        ax.set_xlabel("")

    axes[1,1].set_yticklabels([])
    for i in [1,2]:
        axes[i,0].set_ylabel('β diversity\n%s' % DM_NAMES.get(METRIC_BETA))
    axes[1,0].set_xticks([0.5], ["RM"])
    axes[1, 2].axis('off')


    ## THIRD ROW: beta diversity distances to Controls
    markers_control = {'Janvier': '^', 'Duesseldorf': 'o'}
    data = beta['beta'][~beta['beta']['reference'].isin(['B6J','RM']) & 
                        (beta['beta']['generation'] == 'RM')]
    #sns.lineplot(data=data, y=METRIC_BETA, x='generation', hue='reference_host_genotype', sort=False, ax=axes[2,0], marker="o", style='reference_host_geolocname', palette=COLORS)
    sns.lineplot(data=data, y=METRIC_BETA, x='generation', hue='reference_host_genotype', sort=False, ax=axes[2,0], markers=markers_control, style='reference_host_geolocname', style_order=['Duesseldorf', 'Janvier'], palette=COLORS)
    #sns.boxplot(data=data, y=METRIC_BETA, hue='reference_host_genotype', ax=axes[2,0], x='reference_host_geolocname', palette={n:alpha_blending(c,0.4) for n,c in COLORS.items()}, hue_order=ORDER_GENOTYPE)
    
    for (location, host_genotype), g in data.groupby(['reference_host_geolocname', 'reference_host_genotype']):
        axes[2,0].text(0.0, g[METRIC_BETA].mean(), get_sample_label(location, numSamples, genotypes=[host_genotype]), horizontalalignment='right')
    
    axes[2,0].legend().remove()
    axes[2,0].set_xlim((-1,1))
    for location, g in data.groupby('reference_host_geolocname'):
        #axes[2,0].text(0.01, 0.115 if location == 'Duesseldorf' else 0.145, 'p=%.3f' % kruskal(
        axes[2,0].text(0.01, g[METRIC_BETA].mean(), 'p=%.3f' % kruskal(
            g[(g['reference_host_genotype'] == 'B6J')][METRIC_BETA],
            g[(g['reference_host_genotype'] == 'C')][METRIC_BETA]).pvalue)

    data = beta['beta'][~beta['beta']['reference'].isin(['B6J','RM']) & 
                        (beta['beta']['reference'].apply(lambda x: x.split()[-1]) == beta['beta']['host_genotype'])]
    sns.lineplot(data=data, y=METRIC_BETA, x='generation', hue='reference_host_genotype', sort=False, ax=axes[2,1], markers=markers_control, style='reference_host_geolocname', style_order=['Duesseldorf', 'Janvier'], palette=COLORS)
    axes[2,1].legend().remove()
    
    for ax in axes[2]:
        # same y limits for all three subplots
        ax.set_ylim(BETA_LIM)
        # add horizontal grid lines ...
        ax.set_axisbelow(True)
        # ... behind the plots
        ax.grid(axis='y')
        if ax == axes[2,1]:
            ax.set_xlabel("Generations")
        else:
            ax.set_xlabel("")
    axes[2, 1].set_yticklabels([])

    axes[2, 1].set_ylabel("")
    axes[2, 2].axis('off')
    
    # legend for alpha diversity
    axes[0,1].legend(handles=[
        Line2D([0], [0], marker='o', color=COLORS['RM'], label='RM', linestyle=''),
        Line2D([0], [0], marker='o', color=COLORS['B6J'], label='B6J'),
        Line2D([0], [0], marker='o', color=COLORS['C'], label='C'),
        ],
        title='α diversity', loc='upper left')

    # legend for distance to RM
    axes[1,2].legend(handles=[
        Line2D([0], [0], color=sns.color_palette()[7], label='avg. intra cage', linestyle='--'),
        Line2D([0], [0], color=sns.color_palette()[6], label='avg. intra cage lineage', linestyle='dashdot'),
        Line2D([0], [0], marker='o', color=COLORS['B6J'], label='Δ(RM, B6J)'),
        Line2D([0], [0], marker='o', color=COLORS['C'], label='Δ(RM, C)'),
        Line2D([0], [0], marker='o', color='green', label='Δ(B6J, C)'),
        ],
        title='β distance to RM', loc='center')

    # legend for distance to Controls
    axes[2,2].legend(handles=[
        Line2D([0], [0], marker=markers_control['Duesseldorf'], color=COLORS['B6J'], label='Δ(Duesseldorf B6J, RM or B6J)'),
        Line2D([0], [0], marker=markers_control['Duesseldorf'], color=COLORS['C'],   label='Δ(Duesseldorf C, RM or C)'),
        Line2D([0], [0], marker=markers_control['Janvier'], color=COLORS['B6J'], label='Δ(Janvier B6J, RM or B6J)', linestyle='dotted'),
        Line2D([0], [0], marker=markers_control['Janvier'], color=COLORS['C'],   label='Δ(Janvier B6J, RM or C)', linestyle='dotted')],
        title='β distances to Controls', loc='center')

    plt.figtext(0.09, 0.87, 'A', fontsize=16)
    plt.figtext(0.09, 0.6, 'B', fontsize=16)
    plt.figtext(0.09, 0.33, 'C', fontsize=16)
    
    fig.suptitle(title)
    
    return fig

# %% [code]
def compute_beta_distances(res_beta, meta, metric='weighted_unifrac'):
    results = dict()

    dm = res_beta[metric].to_data_frame()
    data = []
    for refgrp, g_ref in meta[~meta['sample_category'].isin(['C'])].groupby('sample_category'):
        #print(refgrp)
        for (host_genotype, generation), g in meta[meta['sample_category'].isin(['C','B6J','RM'])].groupby(['sample_category', 'generation']):
            #print("  ", host_genotype, generation)
            subset_g_ref = g_ref
            if refgrp == host_genotype:
                continue
            if refgrp == 'B6J':
                subset_g_ref = g_ref[g_ref['generation'] == generation]
            assert len(set(subset_g_ref.index) & set(g.index)) == 0
            #print("    ", len(subset_g_ref.index), len(g.index))
            d = dm.loc[subset_g_ref.index, g.index].stack()
            d.index.rename(['reference_sample_name', 'biol_sample_name'], inplace=True)
            #display(d)
            d = d.to_frame().reset_index().rename(columns={0: metric, 'level_0': 'idx_reference', 'level_1': 'idx_sample'})
            d['generation'] = generation
            d['host_genotype'] = host_genotype
            d['reference'] = refgrp
            d['reference_host_genotype'] = refgrp.split()[-1]
            d['reference_host_geolocname'] = refgrp.split()[0]
            data.append(d)
    results['beta'] = pd.concat(data).sort_values(by='generation', key=KEY_GENERATIONS)
    
    cids = [cid for (cid, size) in meta.groupby('cage_id').size().items() if size > 1]
    subset = meta[meta['cage_id'].isin(cids)]
    results['intra_cage'] = {'num_cages': len(cids),
                             'sum_mice': subset.shape[0],
                             'avg_mice_per_cage': subset.groupby('cage_id').size().mean(),
                             'avg_%s' % metric: np.mean([dm.loc[a,b] for cage_id, g in subset.groupby('cage_id') for (a,b) in combinations(g.index, 2)])}

    distances = []
    for (generation, host_genotype, cage_line), g in meta.groupby(['generation', 'host_genotype', 'cage_line']):
        for (cageA, cageB) in combinations(g['cage_id'].unique(), 2):
            #display(dm.loc[g[g['cage_id'] == cageA].index, g[g['cage_id'] == cageB].index])
            d = dm.loc[g[g['cage_id'] == cageA].index, g[g['cage_id'] == cageB].index].stack()
            d.index.rename(['cageA', 'cageB'], inplace=True)
            d = d.reset_index().rename(columns={0: metric})
            d[['generation', 'host_genotype', 'cage_line']] = [generation, host_genotype, cage_line]
            distances.append(d)
    distances = pd.concat(distances)
    results['intra_cage_line'] = {'num_cages': meta.groupby(['generation', 'host_genotype', 'cage_line'])['cage_id'].unique().apply(len).sum(),
                                  'avg_cages': meta.groupby(['generation', 'host_genotype', 'cage_line'])['cage_id'].unique().apply(len).mean(),
                                  'sum_mice_mice': distances.shape[0],
                                  'avg_mice_mice': distances.groupby(['generation', 'host_genotype', 'cage_line']).size().mean(),
                                  'avg_%s' % metric: distances[metric].mean(),
                                 }
    return results

# %% [code]
%matplotlib inline
fig = draw_temporals(res_gut, meta_gut, num_permutations=9999)

# %% [markdown]
### Panel D

# %% [code]
def _get_maternal_type(row):
    if row['host_genotype_dam'] != row['host_genotype_offspring']:
        return 'other genotype'
    elif row['host_genotype_dam'] == 'Unknown':  # only for source tracker
        return 'Unknown'
    else:
        if row['cage_line_dam'] == row['cage_line_offspring']:
            return 'true dams'
        elif (row['generation_dam'] == 'F2') and (row['generation_offspring'] == 'F3') and \
             (row['cage_line_dam']  == 'C2') and (row['cage_line_offspring']  == 'C3'):
            return 'true dams'
        else:
            return 'other cage lineage'
        raise ValueError("should not happen")

def compute_maternalLegacy(meta, res_beta, metric):
    dm = res_beta['results'][metric].to_data_frame()

    dists = []
    for (genP, genO) in zip(ORDER_GENERATIONS[1:-1], ORDER_GENERATIONS[2:-1]):
        idx_dams = meta[(meta['generation'] == genP) & (meta['gave_birth'] == True)].index
        idx_offspring = meta[(meta['generation'] == genO)].index
        assert(len(set(idx_dams) & set(idx_offspring)) == 0)

        dist = dm.loc[idx_dams, idx_offspring]
        dist.index.name = 'dam'
        dist.columns.name = 'offspring'
        dist = dist.stack().reset_index().rename(
            columns={0: metric}).merge(
                meta[['generation', 'cage_line', 'host_genotype']], left_on='dam', right_index=True, how='left').merge(
                meta[['generation', 'cage_line', 'host_genotype']], left_on='offspring', right_index=True, how='left', suffixes=('_dam', '_offspring'))
        dist['maternal'] = dist.apply(_get_maternal_type, axis=1)
        dists.append(dist)
    dists = pd.concat(dists)

    return {'dists': dists, 
            'sample_numbers': dists.groupby(['generation_offspring','host_genotype_offspring','maternal']).size()}

def draw_maternalLegacy(meta, res_beta, metric='unweighted_unifrac'):
    data = compute_maternalLegacy(meta, res_beta, metric)
    dists = data['dists']
    nums = data['sample_numbers']
    
    fig, axes = plt.subplots(1, 2, figsize=(1.9*len(dists['generation_offspring'].unique()),5), sharey=True)
    fig.subplots_adjust(wspace=0)

    order_hue = ['true dams', 'other cage lineage', 'other genotype']

    for (ax, (host_genotype, g)) in zip(axes, list(dists.groupby('host_genotype_offspring'))):
        sns.boxplot(data=g, y=metric, x='generation_offspring', ax=ax, hue='maternal', hue_order=order_hue, palette={'other cage lineage': 'lightgray',
                                                                                                                     'other genotype': 'green',
                                                                                                                     'true dams': COLORS[host_genotype]})
        axTop = ax.twiny()
        axTop.set_xlim(ax.get_xlim())
        axTop.set_xticks(ax.get_xticks(), ['%ii-i\n%ii-i\n%ii-i' % ( 
                                               nums.loc[t.get_text(), host_genotype, 'true dams'],
                                               nums.loc[t.get_text(), host_genotype, 'other cage lineage'],
                                               nums.loc[t.get_text(), host_genotype, 'other genotype'],
                                         ) 
                                  for t in ax.get_xticklabels()])
        ax.legend().set_title("")

        annotator = Annotator(ax, 
            [((cat, 'true dams'), (cat, 'other cage lineage')) for cat in g['generation_offspring'].unique()] + \
            [((cat, 'other genotype'), (cat, 'other cage lineage')) for cat in g['generation_offspring'].unique()], 
            data=g, y=metric, x='generation_offspring', hue='maternal', hue_order=order_hue)
        annotator.configure(test='Mann-Whitney', text_format='star', loc='inside', comparisons_correction="fdr_bh", correction_format="default")
        _ = annotator.apply_and_annotate()

        ax.set_xlabel("")
        if host_genotype == 'B6J':
            ax.set_ylabel('β diversity\n%s' % DM_NAMES.get(metric))
        else:
            ax.set_ylabel("")
            ax.legend().remove()
        ax.text(0, ax.get_ylim()[0]*1.2, host_genotype)
    plt.figtext(0.09, 0.89, 'D', fontsize=16)
    return fig

# %% [code]
%matplotlib inline
fig = draw_maternalLegacy(meta_gut, res_gut['beta_diversity'], metric='unweighted_unifrac')

# %% [markdown]
### Panel E

# %% [code]
def compute_maternalLegacy_sourcetracker(meta, rawcounts):
    dists = []
    for (genP, genO) in zip(ORDER_GENERATIONS[1:-1], ORDER_GENERATIONS[2:-1]):
        meta_dams = meta[(meta['generation'] == genP) & (meta['gave_birth'] == True)].copy()
        meta_offspring = meta[(meta['generation'] == genO)].copy()
        assert(len(set(meta_dams.index) & set(meta_offspring.index)) == 0)

        meta_dams['st2_type'] = 'Source'
        meta_dams['st2_env'] = meta_dams.index
        meta_offspring['st2_type'] = 'Sink'
        meta_offspring['st2_env'] = np.nan
        
        meta_st = pd.concat([meta_dams[['st2_type', 'st2_env']],
                             meta_offspring[['st2_type', 'st2_env']]])
        
        res_st2 = sourcetracker2(rawcounts, meta_st, col_envname='st2_env', col_type='st2_type', dry=False, wait=False)        
        if res_st2['results'] is not None:
            dist = res_st2['results'].stack().reset_index().rename(
                columns={'Source': 'dam', 'Sink': 'offspring', 0: 'impact'}).merge(
                    meta[['generation', 'cage_line', 'host_genotype']], left_on='dam', right_index=True, how='left').merge(
                    meta[['generation', 'cage_line', 'host_genotype']], left_on='offspring', right_index=True, how='left', suffixes=('_dam', '_offspring'))
            dist['maternal'] = dist.apply(_get_maternal_type, axis=1)
            dists.append(dist)
    if len(dists) > 0:
        dists = pd.concat(dists)
    return dists

def draw_maternalLegacy_sourcetracker(meta, rawcounts):
    res_st = compute_maternalLegacy_sourcetracker(meta, rawcounts)
    
    fig, axes = plt.subplots(1, 2, figsize=(1.25*10, 5), sharey=True)
    fig.subplots_adjust(wspace=0)

    hue_order = ['other cage lineage', 'true dams', 'other genotype']
    numSamples = res_st[res_st['maternal'] != 'Unknown'].groupby(['host_genotype_offspring', 'generation_offspring', 'maternal']).size()

    for (ax, (host_genotype, g)) in zip(axes, res_st.groupby('host_genotype_offspring')):
        data = g[g['maternal'] != 'Unknown']
        sns.boxplot(data=data, y='impact', x='generation_offspring', hue='maternal',
                    ax=ax, palette={'true dams': COLORS[host_genotype], 'other cage lineage': 'lightgray', 'other genotype': 'green'},
                    hue_order=hue_order)

        annotator = Annotator(ax, [
                ((generation, 'true dams'), (generation, 'other cage lineage')) for generation in g['generation_offspring'].unique()
            ], data=data, y='impact', hue='maternal', x='generation_offspring', hue_order=hue_order)
        annotator.configure(test='Mann-Whitney', text_format='star', loc='inside', comparisons_correction="fdr_bh", correction_format="default")
        annotator.apply_and_annotate()


        if ax == axes[0]:
            ax.legend().set_title("")
        else:
            ax.legend().remove()
            ax.set_ylabel("")
        ax.set_xlabel("")
        ax.text(0, -0.205, host_genotype)

        axTop = ax.twiny()
        axTop.set_xlim(ax.get_xlim())
        axTop.set_xticks(ax.get_xticks(), 
                         ['\n'.join(['%ii-i' % numSamples.loc[(host_genotype, t.get_text(), maternal)]
                                     for maternal in reversed(hue_order)])
                          for t in ax.get_xticklabels()])
    return fig

# %% [code]
%matplotlib inline
fig = draw_maternalLegacy_sourcetracker(meta_gut, res_gut['counts_plantsremoved'])

# %% [markdown]
### Panel F

# %% [code]
ENAPROJECT = 'PRJEB28381'

prj_robertson = dict()
prj_robertson['paths'] = dict()

# %% [markdown]
#### Download SRA metadata for Robertson samples
Interestingly, it was easier to obtain metadata from NCBI than from ENA for us. Here is how we did it.
  1. Browse to `https://www.ncbi.nlm.nih.gov/search/all/?term=PRJEB28381`
  2. Select the 346 `SRA` "hits", not the single BioProject: ![image-9.png](attachment:image-9.png)
  3. In the resulting list page, hit `send to`, select `Run selector` as "format" and hit `go` ![image-7.png](attachment:image-7.png)
  4. Hit `metadata` and store the according text file `SraRunTable.txt` in `Incoming`: ![image-8.png](attachment:image-8.png) (we also provide this file through our repository)

# %% [markdown]
#### Normalize Robertson's metadata

# %% [code]
meta_robertson = pd.read_csv('Incoming/SraRunTable.txt', sep=",", index_col=0).rename(
    columns={'common_name': 'host_common_name',
             'Organism': 'host_scientific_name'})[[
    'host_common_name',
    'host_scientific_name',
    'Sample_name',
    'Sample Name',
    'sex',
    'Submitter_Id',  
]]

# obtain host body site as last character in sample name
meta_robertson['host_body_site'] = meta_robertson['Sample_name'].apply(lambda x: {
    'C': 'UBERON:colon',
    'I': 'UBERON:ileum',
    'P': 'UBERON:feces'}.get(x.split('.')[-1][0]))

meta_robertson['sample_type'] = meta_robertson['host_body_site'].apply(
    lambda x: 'feces' if x == 'UBERON:feces' else 'colon content')
meta_robertson['host_genotype'] = 'c57bl6'
meta_robertson['qiita_study_id'] = 'Robertson'
#meta_robertson['host_scientific_name'] = meta_robertson['host_scientific_name'].apply(str.lower)

# derive experiment from Sample_name, i.e. if name starts with 'CH' = 'co-housing', if starts 
# with either Js or Ts = 'sentinels' and all others are 'littermates'
def _get_experiment(row):
    if row['Sample_name'].startswith('CH'):
        return 'co-housing'
    if row['Sample_name'].startswith('Js') or row['Sample_name'].startswith('Ts'):
        return 'sentinels'
    return 'littermates'
meta_robertson['experiment'] = meta_robertson.apply(_get_experiment, axis=1)

# derive generation from Sample_name:
# 1) all sentinels are of generation P1, by definition
# 2) all co-housing animals are of generation F2, by definition
# 3) F2 generation samples of littermates are flagges by ".F2." in the sample name
meta_robertson.loc[meta_robertson[meta_robertson['experiment'] == 'sentinels'].index, 'generation'] = 'P1'
meta_robertson.loc[meta_robertson[meta_robertson['experiment'] == 'co-housing'].index, 'generation'] = 'F2'
meta_robertson.loc[meta_robertson[(meta_robertson['experiment'] == 'littermates') & 
                   meta_robertson['Sample_name'].apply(lambda x: '.F2.' not in x)].index, 'generation'] = 'F1'
meta_robertson.loc[meta_robertson[(meta_robertson['experiment'] == 'littermates') & 
                                  (pd.isnull(meta_robertson['generation']))].index, 'generation'] = 'F2'

# derive vendor from Sample_name:
meta_robertson.loc[
    meta_robertson[meta_robertson['experiment'] == 'sentinels'].index, 
    'vendor'] = meta_robertson[meta_robertson['experiment'] == 'sentinels']['Sample_name'].apply(
        lambda x: {'T': 'mTAC', 'J': 'mJAX'}.get(x[0]))
meta_robertson.loc[
    meta_robertson[meta_robertson['experiment'] == 'co-housing'].index, 
    'vendor'] = meta_robertson[meta_robertson['experiment'] == 'co-housing']['Sample_name'].apply(
        lambda x: 'mJAX' if '.JAX' in x else 'mTAC')
meta_robertson.loc[
    meta_robertson[meta_robertson['experiment'] == 'littermates'].index, 
    'vendor'] = meta_robertson[meta_robertson['experiment'] == 'littermates']['Sample_name'].apply(
        lambda x: 'mJAX' if x.startswith('J.T.') else 'mTAC')

# derive "litter" from Sample_name for littermates experiment
meta_robertson.loc[
    meta_robertson[(meta_robertson['experiment'] == 'littermates')].index, 
    'litter'] = meta_robertson[(meta_robertson['experiment'] == 'littermates')]['Sample_name'].apply(
        lambda x: x.split('.')[-2])

meta_robertson['sample_title'] = meta_robertson['experiment'].apply(
    lambda x: {'co-housing': "co-housed",
               'littermates': "littermate",
               'sentinels': "sentinel"}.get(x,x))
meta_robertson.loc[meta_robertson[(meta_robertson['experiment'] == 'co-housing') & 
                                  (meta_robertson['Sample_name'].apply(lambda x: x[3] == 's'))].index, 
                   'sample_title'] = 'non-co-housed'

# question by Stefan Janssen: 
# "In your metadata file, I do find 12 JAX and 13 TAC colon f2 littermate samples. It might be too hard to see, 
#  but in the supplemental Figure S6, I can only spot ~8 samples, which is in line with the below table. 
#  Is it possible, that you re-sequenced some samples due to bad quality?"
#
# answer from Heather Maughan (group of Dana Philpott)
# "Litter 3 ended up being excluded from the F2 analyses reported in the paper, so the sample size ended up 
#  being 8-10 samples for each (not sure why Sue only had n=8 in the table). Clearly we uploaded Litter 3's 
#  sequence data so that is why there appears to be more sequenced samples than those reported in the paper. 
#
#  Litter 3 was excluded because their microbiotas were quite divergent from other litters and the parents, 
#  and because they were the last of the 3 subsequent litters (1, 2, then 3 from the same breeding pair), the
#  lab was concerned about confounding factors (perhaps breeder age? really it was unknown) affecting the 
#  microbiota."
idx_skip = meta_robertson.loc[(meta_robertson['experiment'] == 'littermates') & 
                  (((meta_robertson['vendor'] == 'mJAX') & (meta_robertson['litter'] == '3')) |
                   ((meta_robertson['vendor'] == 'mTAC') & (meta_robertson['litter'] == '4')))].index

# flip indices of 6 samples to resolve a sample swap, see below 
idx_tac = list(meta_robertson[meta_robertson['Sample_name'].isin(
    ['CH7sa.JAX3.2.%s' % bs for bs in ['C','I','P']])].index)
idx_jax = list(meta_robertson[meta_robertson['Sample_name'].isin(
    ['CH7sa.TAC3.2.%s' % bs for bs in ['C','I','P']])].index)
meta_robertson.index = [{new: old 
                         for (new, old) 
                         in zip(idx_tac+idx_jax, idx_jax+idx_tac)}.get(i, i) 
                        for i 
                        in meta_robertson.index]

meta_robertson['skip_sample'] = False
meta_robertson.loc[idx_skip, 'skip_sample'] = True

# append _1 to sample names to match with sequencing file names
meta_robertson.index = list(map(lambda x: x + '_1', meta_robertson.index))

# %% [markdown]
**question by Stefan Janssen**:

"Here are six samples which seem to be undecided if they are TAX or JAX. Can you help me resolve this?
![image-10.png](attachment:image-10.png)

**answer from Heather Maughan (group of Dana Philpott)**:

Sue believed that these samples had been accidentally swapped during the extraction step; she changed the metadata but did not change sample names (because they are linked with the sequence read file names). However, it doesn’t
look like the “Breeding Pair” column was switched/updated. I fixed it in this screenshot. Does that make sense? 


# %% [markdown]
**Note** the below steps require downloaded 2.6GB of sequencing data, trimming and ASV calling, which in total will take some time. Therefore, we host the resulting feature-table in our repositors: `Incoming/robertson-hit.biom`. This notebook will use this file by default to save the extra compute. Should you want to reproduce these steps yourself, simply delete this file!

# %% [code]
prj_robertson['paths']['deblur_table'] = 'Incoming/robertson-hit.biom'

# %% [markdown]
#### Download Robertson's sequencing data
1. Browse to `https://www.ebi.ac.uk/ena/browser/view/PRJEB28381`
2. identify the `Download All` button (make sure to use the correct button, i.e. NOT `Submitted files: FTP` nor `SRA files: FTP` as this will complicate sequence-data to metadata mapping): ![image-11.png](attachment:image-11.png)
3. save the download script to obtain all forward and reverse sequncing files through `wget` (we also provide the resulting file at our repository `Incoming/ena-file-download-read_run-PRJEB28381-fastq_ftp-20231104-1013.sh`). Execution can be done in the following cell

# %% [code]
if not os.path.exists(prj_robertson['paths']['deblur_table']):
    fp_dlscript = 'Incoming/ena-file-download-read_run-PRJEB28381-fastq_ftp-20231104-1013.sh'
    ena_file_cmds = !cat $fp_dlscript | grep "_1.fastq.gz"  # as we discard all reverse reads, there is no need to download them in the first place

    enadir = os.path.join('tmp_dir/no_backup/', ENAPROJECT)
    cluster_run([
        'export ftp_proxy="http://proxy.computational.bio.uni-giessen.de:3128"',
        'mkdir -p %s' % enadir, 'cd %s' % enadir] + ena_file_cmds, 'dl_ENA', os.path.join(enadir, last_ena_file),
        dry=False, ppn=1, use_grid=False)

# %% [markdown]
#### Process Robertson's sequencing data
We are applying here the same pre-processing steps as with our own sequencing data, i.e.
  1. trimming primer sequences
  2. obtain 150nt long ASVs via Deblur

# %% [code]
prj_robertson['metadata'] = meta_robertson
prj_robertson['paths']['demux'] = os.path.abspath('tmp_dir/no_backup/%s/' % ENAPROJECT)
prj_robertson['paths']['tmp_workdir'] = os.path.abspath('./tmp_dir/no_backup')
prj_robertson['paths']['Generated/Emperor'] = os.path.abspath('./Generated/Emperor')

# %% [code]
if not os.path.exists(prj_robertson['paths']['deblur_table']):
    prj_robertson = project_trimprimers(
        "GTGCCAGCMGCCGCGGTAA", 
        "GGACTACHVGGGTWTCTAAT", prj_robertson, pattern_fwdfiles="*_1.fastq.gz", no_rev_seqs=True)

# %% [code]
if not os.path.exists(prj_robertson['paths']['deblur_table']):
    prj_robertson = project_deblur(prj_robertson, pattern_fwdfiles="*_1.fastq.gz", ppn=32, pmem='2GB')

# %% [code]
# this will produce an Bray-Curtis PCoA Emperor plot to only inspect the Robertson data
# we used this to find the above mentioned sample swap
rob_rare = rarefy(biom2pandas(prj_robertson['paths']['deblur_table']), rarefaction_depth=6000, dry=False)
if rob_rare['results'] is not None:
    rob_beta = beta_diversity(rob_rare['results'], metrics=['bray_curtis'], dry=False, wait=False)
    if rob_beta['results'] is not None:
        _ = emperor(meta_robertson, rob_beta['results'], fp_results='Generated/Emperor/', infix='_checkRob', dry=False, wait=False)

# %% [code]
# merge our and Robertson's metadata
meta_rob = pd.concat([meta_gut, prj_robertson['metadata']])
meta_rob

# %% [code]
# merge feature table
fp_counts_both = 'tmp_dir/counts_both.biom'
os.makedirs(os.path.dirname(fp_counts_both), exist_ok=True)
pandas2biom(fp_counts_both, pd.concat([
    res_gut['counts_plantsremoved'].loc[:, meta_gut.index],
    biom2pandas(prj_robertson['paths']['deblur_table'])], axis=1).fillna(0))

# %% [code]
# construct shared insertion tree
seppres = project_sepp({'paths': {'deblur_table': fp_counts_both,
                                  'tmp_workdir': 'tmp_dir/'}})

# %% [code]
idx_control_samples = set(meta_rob[meta_rob['sample_type'].isin(['sterile water', 'control bedding', 'control food'])].index)
res_rob = process_study(
    metadata=meta_rob,
    control_samples=idx_control_samples,
    fp_deblur_biom=fp_counts_both,
    fp_insertiontree=seppres['paths']['insertion_tree'],
    rarefaction_depth=6000,
    fp_taxonomy_trained_classifier_gg138_chloroMitoRemoval='/vol/jlab/MicrobiomeAnalyses/References/Q2-Naive_Bayes_classifiers/gg-13-8-99-nb-classifier_2022.11.qza',  # as GG2 currently lacks labels for mitochondria/chloroplasts, see https://forum.qiime2.org/t/taxonomy-filtering-greengenes2/28334
    fp_taxonomy_trained_classifier="/vol/jlab/MicrobiomeAnalyses/References/Q2-Naive_Bayes_classifiers/gg_2022_10_backbone_full_length.nb_2023.5.qza",
    tree_insert=seppres['insertion_tree'],
    is_v4_region=False,
    emperor_fp='Generated/Emperor',
    emperor_infix='_robertson',
    dry=False,
    use_grid=True, 
    fix_zero_len_branches=False, # since this has already been corrected via the project_sepp call
    emperor_skip_tsne_umap=True,
)

# %% [code]
def get_robertson_dists(betas: dict[str, DistanceMatrix], metric='unweighted_unifrac'):
    dists = []
    #dm = res_rob['beta_diversity']['results'][metric].to_data_frame()
    dm = betas[metric].to_data_frame()
    for (genotype_benga, generation_benga), g_benga in meta_rob[
                                           (meta_rob['skip_sample'] != True) &
                                           (meta_rob['host_scientific_name'] == "Mus musculus") & 
                                           (meta_rob['sample_type'] == "colon content") & 
                                           (meta_rob['host_body_site'] == "UBERON:colon") & 
                                           (meta_rob['qiita_study_id'] != 'Robertson')].fillna({'generation': 'control'}).groupby(['host_genotype', 'generation']):
        if genotype_benga == "b6cf1":
            continue
        for (genotype_rob, generation_rob, vendor_rob, experiment_rob, sample_title_rob), g_rob in meta_rob[
                                           (meta_rob['skip_sample'] != True) &
                                           (meta_rob['host_scientific_name'] == "Mus musculus") & 
                                           (meta_rob['sample_type'] == "colon content") & 
                                           (meta_rob['host_body_site'] == "UBERON:colon") & 
                                           (meta_rob['qiita_study_id'] == 'Robertson')].fillna({'sample_title':'?'}).groupby(['host_genotype', 'generation','vendor','experiment', 'sample_title']):
            # print(genotype_benga, generation_benga, g_benga.shape[0], genotype_rob, generation_rob, vendor_rob, experiment_rob, g_rob.shape[0])
            dm_sub = dm.loc[list(set(g_benga.index) & set(dm.index)),
                            list(set(g_rob.index)   & set(dm.index))]
            dm_sub = dm_sub.stack().reset_index()
            dm_sub['host_genotype_benga'] = genotype_benga
            dm_sub['generation_benga'] = generation_benga
            dm_sub['generation_rob'] = generation_rob
            dm_sub['vendor_rob'] = vendor_rob
            dm_sub['experiment_rob'] = experiment_rob
            dm_sub['sample_title'] = sample_title_rob
            dists.append(dm_sub)
    dists = pd.concat(dists)
    return dists

def get_robertson_combineddists(dists, metric):
    nums = dict()
    d1 = dists[dists['experiment_rob'].isin(['littermates','sentinels'])].copy()
    d1['cat'] = 'host_genotype : ' + d1['host_genotype_benga'] + " VS " + d1.apply(lambda row: 'both' if row['host_genotype_benga'] == 'C' else row['vendor_rob'], axis=1)

    d2 = dists[dists['experiment_rob'].isin(['littermates']) &
               (dists['host_genotype_benga'] == 'B6J') & 
               (dists['generation_rob'] == 'F2')].copy()
    d2['cat'] = 'maternal legacy : ' + d2['vendor_rob']

    d3 = dists[dists['experiment_rob'].isin(['littermates','sentinels']) &
               (dists['host_genotype_benga'] == 'B6J') & 
               (dists['vendor_rob'] == 'mJAX')].copy()
    d3['cat'] = 'generation : ' + d3['generation_rob']

    d4 = dists[dists['experiment_rob'].isin(['littermates','sentinels']) &
               (dists['host_genotype_benga'] == 'B6J') & 
               (dists['vendor_rob'] == 'mJAX') & 
               (dists['generation_rob'] == 'F2')].copy()
    d4['cat'] = 'breeding : ' + d4['generation_benga']

    d5 = dists[(dists['experiment_rob'] == 'co-housing') & 
               (dists['host_genotype_benga'] == 'B6J')].copy()
    d5['cat'] = 'co-housing : ' + d5['sample_title']

    comb_dists = pd.concat([d1, d2, d3, d4],axis=0)
    comb_dists = comb_dists.rename(columns={0: metric})

    nums = dict()
    for c, g in comb_dists.groupby('cat'):
        nums[c] = pd.concat([meta_rob.loc[g['level_0'].unique(),:].groupby('host_genotype').size(),
                             meta_rob.loc[g['level_1'].unique(),:].groupby('vendor').size()])
    
    return comb_dists, nums

def plot_robertson(beta_diversities: dict[str, DistanceMatrix], metric: str='unweighted_unifrac'):
    dists = get_robertson_dists(beta_diversities, metric)
    comb_dists, nums = get_robertson_combineddists(dists, metric)
    
    fig, axes = plt.subplots(1,1, figsize=(8/14*16,5)) #(8,5)

    o_cat = [
        'host_genotype : C VS both', 'host_genotype : B6J VS mTAC', 'host_genotype : B6J VS mJAX',
        'generation : P1', 'generation : F1', 'generation : F2',
        'maternal legacy : mTAC', 
        'breeding : P', 'breeding : F1', 'breeding : F2', 'breeding : F3', 'breeding : F4', 'breeding : F5','breeding : control', 
    ]
    pal = {
        'host_genotype : C VS both': COLORS['C'],
        'host_genotype : B6J VS mJAX': COLORS['mJAX'],
        'host_genotype : B6J VS mTAC': COLORS['B6J'],

        'generation : P1': COLORS['mJAX'], 
        'generation : F1': '#e7e7e7', 
        'generation : F2': '#e7e7e7',
        'maternal legacy : mTAC': '#e7e7e7',

        'breeding : P': COLORS['P'], 
        'breeding : F1': COLORS['F1'], 
        'breeding : F2': COLORS['F2'], 
        'breeding : F3': COLORS['F3'], 
        'breeding : F4': COLORS['F4'], 
        'breeding : F5': COLORS['F5'],
        'breeding : control': '#555555',
    }

    sns.boxplot(data=comb_dists, y=metric, x='cat', ax=axes, zorder=10, 
                order=o_cat, palette=pal,
               )
    axes.axhline(y=comb_dists.groupby('cat').mean(numeric_only=True).loc['generation : F2',metric], color='#e16d1a')
    axes.axvline(x=2.5, color='gray', linestyle='--')
    axes.axvline(x=6.5, color='gray', linestyle='--')
    axes.set_ylabel('β diversity\n%s' % DM_NAMES.get(metric))

    axTop = axes.twiny()
    axTop.set_xlim(axes.get_xlim())
    axTop.set_xticks(axes.get_xticks(), 
                     ['\n'.join(['' if (nums[t.get_text()].to_dict().get(hg, 0) == 0) else '%ii' % nums[t.get_text()].to_dict().get(hg, 0)
                      for hg in ['C', 'mTAC', 'mJAX', 'B6J']])
                      for t in axes.get_xticklabels()])
    axes.set_xticklabels(['C', 'TAC', 'JAX', 
                          'P1', 'F1', 'F2', 'F2',
                          'P', 'F1', 'F2', 'F3', 'F4', 'F5', 'Ctrl.'])
    axes.set_xlabel("")

    annotator = Annotator(axes, [
        ('host_genotype : B6J VS mTAC', 'host_genotype : B6J VS mJAX'),
        ('host_genotype : C VS both', 'host_genotype : B6J VS mTAC'),
        ('host_genotype : C VS both', 'host_genotype : B6J VS mJAX'),

        ('generation : F1', 'generation : F2'),
        ('generation : P1', 'generation : F1'),

        ('maternal legacy : mTAC', 'generation : F2'),

        ('breeding : P', 'breeding : F1'),
        ('breeding : F1', 'breeding : F2'),
        ('breeding : F2', 'breeding : F3'),
        ('breeding : F3', 'breeding : F4'),

    ], data=comb_dists, y=metric, x='cat', order=o_cat)
    annotator.configure(test='Mann-Whitney', text_format='star', loc='outside', comparisons_correction="fdr_bh", correction_format="default")
    annotator.apply_and_annotate()

    axes.text(1, 0.82, 'host genotype', horizontalalignment='center')
    axes.text(5.5, 0.82, 'maternal legacy', horizontalalignment='center')
    axes.text(5, 0.81, '♀JAX', horizontalalignment='center')
    axes.text(6, 0.68, '♀TAC', horizontalalignment='center')

    plt.figtext(0.09, 0.89, 'F', fontsize=16)
    
    return fig

# %% [code]
_ = plot_robertson(res_rob['beta_diversity']['results'], metric='unweighted_unifrac')

# %% [markdown]
## Figure 3
### Panel A
  1. Browse to view.qiime2.org and upload the file `Generated/Emperor/emperor-pcoa_gut_weighted_unifrac.qzv`. ![image-3.png](attachment:image-3.png)
  2. right click on the 3D animation to open the context menu ![image-1.png](attachment:image-1.png)
  3. choose `Load saved settings` and select the file `EmperorSettings/emp_settings_gut_panelA.json`
  
This is a screenshot of how your browser should render the interactive Emperor plot:
![image-2.png](attachment:image-2.png)

# %% [markdown]
### Panel B
see above, but load settings from file `EmperorSettings/emp_settings_gut_panelB.json`
![image-4.png](attachment:image-4.png)

# %% [markdown]
### Panel C
see above, but load settings from file `EmperorSettings/emp_settings_gut_panelC.json`
![image-5.png](attachment:image-5.png)

# %% [markdown]
### Panel D
see above, but load settings from file `EmperorSettings/emp_settings_gut_panelD.json`
![image-6.png](attachment:image-6.png)

# %% [markdown]
## Figure 4

# %% [code]
sns.set_style("ticks")
rred = redundancy_analysis_beta(meta_gut, 
                                    res_gut['beta_diversity']['results']['bray_curtis'], 
                                    'Bray-Curtis',
                                    categorials=['host_genotype','independent_cage_lineage','sex'], 
                                    intervals=[], dates={},
                                    ordinals={'generation': ORDER_GENERATIONS},
                                    title='Effect Sizes Gut Microbiome',
                                    dry=False, wait=False, #dirty=True,
                                    seed=42  # set a seed to the permutations engine of ordiR2step: https://stackoverflow.com/questions/30251636/reproducing-results-from-ordistep
                                   )

# %% [code]
# style the figure
axes = rred['results']['figure'].get_axes()[0]
axes.set_title('Effect Sizes Gut Microbiome')
axes.set_ylabel("")
axR = axes.twinx()
axR.set_ylabel("Bray-Curtis")
axR.set_yticks([])
axes.set_yticklabels([t.get_text().replace('independent_cage_lineage', 'maternal legacy').replace('host_genotype', 'host genotype') for t in axes.get_yticklabels()])
rred['results']['figure'].set_size_inches((6, 3))
rred['results']['figure'].savefig('Figures/fig4_gutEffectSizes.svg', bbox_inches='tight')
rred['results']['figure']

# %% [code]
rred['results']['table']

# %% [markdown]
## Figure 5
### Panels A,B,C

# %% [code]
%matplotlib inline
fig = draw_temporals(res_skin, meta_skin, 
                     ALPHA_LIM=(4.5,14.5),
                     METRIC_BETA='unweighted_unifrac', BETA_LIM=(0.51, 0.8),#78),
                     title='skin microbiome')

# %% [markdown]
### Panel D

# %% [code]
%matplotlib inline
fig = draw_maternalLegacy(meta_skin, res_skin['beta_diversity'], metric='unweighted_unifrac')

# %% [markdown]
### Panel E

# %% [code]
%matplotlib inline
fig = draw_maternalLegacy_sourcetracker(meta_skin, res_skin['counts_plantsremoved'])

# %% [markdown]
## Figure 6
### Panel A

# %% [code]
def drawTaxBarplot(res, meta, title='gut microbiome', minreadnr=60*270*0.5, rank='Genus'):
    fig, axes = plt.subplots(3,1,figsize=(40,5+2), gridspec_kw={'height_ratios': [10, 0.5, 0.5]})
    tax = plotTaxonomy(res['rarefaction']['results'], 
                       meta.fillna('Controls').sort_values(by='generation_cageline', key=KEY_TAXPLOT),
                       file_taxonomy=res['taxonomy']['RDP']['results']['Taxon'],
                       group_l1='sample_category', 
                       group_l2='generation_cageline', 
                       rank=rank,
                       min_abundance_grayscale=0.0, grayscale=True,
                       no_top_labels=True, no_sample_numbers=True,
                       print_meanrelabunances=True, 
                       minreadnr=minreadnr, 
                       ax=axes[0], horizontal_spacer=3)

    for ax in axes[1:]:
        ax.set_xlim(axes[0].get_xlim())
        ax.set_xticks([])
        ax.set_yticks([])
        ax.set_xlabel("")
        ax.set_ylabel("")
        ax.set_ylim((0,1))
        ax.set_facecolor("white")

    for _, row in tax[2].iterrows():
        axes[1].add_patch(Rectangle((row['xpos'], 0), 1, 1, linewidth=None, color=COLORS.get(row['group_l2'].split('_')[0], 'white')))
        axes[2].add_patch(Rectangle((row['xpos'], 0), 1, 1, linewidth=None, color=COLORS.get(row['group_l2'].split('_')[-1], 'white')))

    for (field, pos, ax) in [('generation', 0, axes[1]), ('cage_line', -1, axes[2])]:
        tax[2][field] = tax[2]['group_l2'].apply(lambda x: x.split('_')[pos])
        tax[2]['run_%s' % field] = tax[2].ne(tax[2].shift()).cumsum()[field]
        for run, g in tax[2].groupby('run_%s' % field, sort=False):
            text = g[field].unique()[0]
            if g['group_l2'].unique()[0] != text:
                ax.text(g['xpos'].mean(), 0.2, text, horizontalalignment='center')   
    fig.suptitle(title)
    return fig

# %% [code]
%matplotlib inline
fig = drawTaxBarplot(res_gut, meta_gut)

# %% [markdown]
### Panel B

# %% [code]
name_map = OrderedDict([
    ('s__UBA11471 sp000434215', 'species UBA11471 of order Bacteroidales'),
    ('s__Bacteroides_H acidifaciens', 'Bacteroides H acidifaciens'),
    ('s__Bacteroides_H caecimuris', 'Bacteroides H caecimuris'),
    ('s__UBA7173 sp002491305', 'a species of genus UBA7173, family Muribaculacea'),
    ('s__CAG-485 sp002493045', 'a species of genus CAG-485, family Muribaculacea'),
    ('s__UBA7173 sp900540205', 'a species of genus UBA7173, family Muribaculacea'),
    ('s__Paramuribaculum sp000431155', 'Paramuribaculum sp000431155'),
    ('s__Muribaculum intestinale', 'Muribaculum intestinale'),
    ('s__Muribaculum gordoncarteri', 'Muribaculum gordoncarteri'),
    ('s__Akkermansia muciniphila_D_776786', 'Akkermansia muciniphila D 776786'),
    ('s__Parasutterella sp009767915', 'Parasutterella sp009767915'),
    ('s__Ruminiclostridium_E siraeum', 'Ruminiclostridium E siraeum')
])

# %% [code]
%matplotlib inline
sns.set_style("ticks")
res_ancom = ancom(
    res_gut['rarefaction']['results'],
    #res_gut['counts_plantsremoved'],
    rank=('Species'),
    taxonomy=res_gut['taxonomy']['RDP']['results']['Taxon'],
    grouping=meta_gut[(meta_gut['host_genotype'] != 'RM')]['host_genotype'],
    dry=False, wait=False, min_mean_abundance_per_feature=0.007,
    post_cache_arguments={'palette': COLORS, 'feature_order': list(name_map.keys()), 'hue_order': ['B6J', 'C']})

# %% [code]
%matplotlib inline
ax = res_ancom['results']['figure'].get_axes()[0]
ax.set_yticks(ax.get_yticks(), [name_map.get(t.get_text(), t.get_text()) 
                                for t in ax.get_yticklabels()])
ax.set_ylabel("collapsed at species level")
ax.set_title("ANCOM on %i gut samples" % len(res_ancom['results']['features']['sample_name'].unique()))
ax.set_xlabel("relative abundance")
ax.get_legend().set_title("host genotype")
ax.axhline(y=3-0.5, linestyle='dashed', linewidth=1, color='lightgray', xmin=-1.55, xmax=1, clip_on = False)
ax.axhline(y=3+6-0.5, linestyle='dashed', linewidth=1, color='lightgray', xmin=-1.55, xmax=1, clip_on = False)
fig = res_ancom['results']['figure']
fig.set_size_inches((3, 5))
fig

# %% [markdown]
The following graph supports the statement in the discussion, that we can treat the significantly differentially abundant Genus "Ruminiclostridium" as equivalent to the Genus "Ruminococcus", which was discussed in Horne et al. 2019. The reason is a different taxonomic assignment, depending on the version of the GreenGenes database.

# %% [code]
def draw_Rumi_equivalence(tax_gg2, counts):
    # taxonomic assignment with GG1, i.e. release 13.8
    res_tax_gg1 = taxonomy_RDP(
        counts, 
        "/homes/sjanssen/MicrobiomeAnalyses/References/Q2-Naive_Bayes_classifiers/gg-13-8-99-nb-classifier_2022.11.qza", # obtained from https://data.qiime2.org/2022.11/common/gg-13-8-99-nb-classifier.qza
        dry=False, use_grid=True)
    gg1_featname = 'Ruminococcus'
    gg1_rumi = res_tax_gg1['results'][res_tax_gg1['results']['Taxon'].apply(lambda x: 'Ruminococcus' in x)]
    print("GG 13.8: Of the %i ASVs in our experiment, %s ASVs get assigned to Genus '%s'" % (
        counts.shape[0], 
        ', '.join(map(str, gg1_rumi.groupby('Taxon').size().unique())),
        gg1_featname))

    gg2_featname = 'Ruminiclostridium'
    gg2_rumi = tax_gg2[tax_gg2['Taxon'].apply(lambda x: gg2_featname in x)]
    print("GG 2   : Of the %i ASVs in our experiment,  %s ASVs get assigned to Genus '%s'" % (
        counts.shape[0], 
        ', '.join(map(str, gg2_rumi.groupby('Taxon').size().unique())),
        gg2_featname))

    size_gg2 = counts.loc[list((set(gg2_rumi.index) - set(gg1_rumi.index)) & set(counts.index)),:].sum(axis=1).sum()
    size_both = counts.loc[list((set(gg1_rumi.index) & set(gg2_rumi.index)) & set(counts.index)),:].sum(axis=1).sum()
    fig, axes = plt.subplots()
    venn2(subsets=(counts.loc[list((set(gg1_rumi.index) - set(gg2_rumi.index)) & set(counts.index)),:].sum(axis=1).sum(),
                   size_gg2,
                   size_both, 
                  ), set_labels=('GG1 (13.8): "%s"' % gg1_featname, 'GG 2: "%s"' % gg2_featname), ax=axes)
    _ = axes.set_title("%.2f%% of reads that\nGG2 assign to '%s'\nget the name '%s' with GG1 (13.8)" % (
        (1 - (size_gg2 / (size_both + size_gg2))) * 100, gg2_featname, gg1_featname))
    return fig

fig = draw_Rumi_equivalence(res_both['taxonomy']['RDP']['results'], res_both['counts_plantsStillIn'])

# %% [markdown]
### Panel C

# %% [code]
%matplotlib inline
sns.set_style("ticks")
feature = 's__Akkermansia muciniphila_D_776786'

cs = collapseCounts_objects(
    res_gut['rarefaction']['results'], 
    'Species', 
    taxonomy=res_gut['taxonomy']['RDP']['results']['Taxon'])[0]
cs /= cs.sum()
data = meta_gut[pd.notnull(meta_gut['generation'])].merge(
    cs.loc[[feature], :].T, 
    left_index=True, right_index=True, how='left')
numSamples = get_sample_numbers(data)

fig, axes = plt.subplots(1,1)#,figsize=(8,6))

sns.stripplot(data=data,
              y=feature, x='generation', hue='host_genotype', 
              palette=COLORS, dodge=True, order=ORDER_GENERATIONS[:-1],
              ax=axes)
axes.set_ylabel("relative abundance of\n%s" % name_map.get(feature))
axes.set_xlabel("")

axTop = axes.twiny()
axTop.set_xlim(axes.get_xlim())
_ = axTop.set_xticks(axes.get_xticks(), 
                     labels=[get_sample_label(generation, numSamples, justI=True) 
                             for generation in ORDER_GENERATIONS[:-1]])

# %% [code]
data.groupby('host_genotype')[feature].max()['B6J']

# %% [markdown]
## Figure 7

# %% [code]
fp_metabolites = 'Incoming/220511__Data_21-0025.xlsx'
x = pd.read_excel(fp_metabolites, sheet_name="01.AllData").rename(columns={'Sample.label.on.tube': 'sample_name'})
x['sample_name'] = 'sample.' + x['sample_name'].astype(str)
data = x[['sample_name', "ISTD", "Resp", "ISTD.Resp", "Resp.Ratio", "UserAnno.Peak", "MI.Peak", "MZ", "MZ.Ex.Left", "MZ.Ex.Right", "MZ.Ex.Units", "RT.Compound", "RT.Peak", "MS.Replicates", "data.source", "ISTD.Resp.Ratio", "Acq.Method.Condensed", "MS.project", "Report.Name", "data.dir", "relativeResponse", "IQR_1.5_outlier", "Compound"]].set_index(['sample_name', 'Compound'])
counts_chems = pd.pivot_table(data=data, columns='sample_name', index="Compound", values="relativeResponse").fillna(0)
# remove compounds containing 'unknown' in its name
counts_chems = counts_chems.loc[[c for c in counts_chems.index if 'unknown' not in c], :]

# also load metadata from Excel sheet that Philipp Westhoff provided
meta_chems = x[['sample_name', "date", "MS.Label", "Species", "Tissue.type", "Condition.1", "Generation", "Benga-Nr."]].set_index(['sample_name']).drop_duplicates().rename(columns={'Condition.1': 'genotype', 'Generation': 'generation'})
meta_chems['Benga-Nr.'] = meta_chems['Benga-Nr.'].astype(str)

# merge metadata from Philipp and Qiita
meta_chems = meta_chems.reset_index().merge(metadata.groupby('host_subject_id_bmfz').head(1), left_on='Benga-Nr.', right_on='host_subject_id_bmfz', how='left').set_index('sample_name')

res_beta_chems = beta_diversity(counts_chems, metrics=['bray_curtis'], dry=False, wait=False)
if res_beta_chems['results'] is not None:
    res_emp_chems = emperor(meta_chems, res_beta_chems['results'], fp_results='Generated/Emperor/', infix='_metabolites', run_tsne_umap=False, dry=False, wait=False)

# %% [markdown]
### Panel A
Use `Generated/Emperor/emperor-pcoa_metabolites_bray_curtis` to upload to `view.qiime2.org`

see Figure 2, but load settings from file `EmperorSettings/emp_settings_metabolites.json`
![image-12.png](attachment:image-12.png)


# %% [markdown]
### Panel B

# %% [code]
dm = res_beta_chems['results']['bray_curtis']
metacol = meta_chems['host_genotype']
fig, axes = plt.subplots(figsize=(4,2.5))
testres = detect_distant_groups(dm, 'Bray-Curtis', metacol)
p = plotGroup_permanovas(dm, metacol, **testres, 
                         ax=axes, horizontal=True, colors_boxplot={'left': COLORS['B6J'],
                                                                   'right': COLORS['C'],
                                                                   'between': 'gray'})
axes.set_ylabel('host genotype')
axes.yaxis.label.set_visible(True)
axes.set_title('pairwise PERMANOVA\n999 permutations')
offset = 0.26
_ = axes.set_yticks([-1*offset,0,offset], ['B6J (n=%i)' % meta_chems['host_genotype'].value_counts().loc['B6J'], 
                                           'p: %.2f' % testres['network']['B6J']['C']['p-value'], 
                                           'C (n=%i)' % meta_chems['host_genotype'].value_counts().loc['C']])
plt.figtext(0.09, 0.89, 'B', fontsize=16)

# %% [markdown]
### Panel C

# %% [code]
metacol = meta_chems['generation_y']
fig, axes = plt.subplots(figsize=(4,2.5))
testres = detect_distant_groups(dm, 'Bray-Curtis', metacol)
p = plotGroup_permanovas(dm, metacol, **testres, 
                         ax=axes, horizontal=True, colors_boxplot={'left': COLORS['F3'],
                                                                   'right': COLORS['F4'],
                                                                   'between': 'gray'})
axes.set_ylabel('generation')
axes.yaxis.label.set_visible(True)
axes.set_title('pairwise PERMANOVA\n999 permutations')
offset = 0.26
_ = axes.set_yticks([-1*offset,0,offset], ['F3 (n=%i)' % meta_chems['generation_x'].value_counts().loc['F3'], 
                                           'p: %.2f' % testres['network']['F3']['F4']['p-value'], 
                                           'F4 (n=%i)' % meta_chems['generation_x'].value_counts().loc['F4']])
plt.figtext(0.09, 0.89, 'C', fontsize=16)

# %% [markdown]
### Panel D

# %% [code]
dsFDR_experiment = dict()
# for dsFDR
pandas2biom('tmp_dir/tmp_dsfrd_metabolome.biom', counts_chems)
meta_chems.to_csv('tmp_dir/tmp_dsfdr_metabolome_meta.tsv', sep="\t", index=True)

dsFDR_experiment = ca.read_amplicon('tmp_dir/tmp_dsfrd_metabolome.biom', 
                                    'tmp_dir/tmp_dsfdr_metabolome_meta.tsv', min_reads=0, normalize=10000)

field = 'host_genotype'
res_dsFDR_genotype = find_diff_taxa(dsFDR_experiment, meta_chems[meta_chems['generation_x'] == 'F4'], [field],
                                    random_seed=43)

if len(res_dsFDR_genotype) > 0:
    p = plot_diff_taxa(counts_chems, meta_chems[meta_chems['generation_x'] == 'F4'][field], res_dsFDR_genotype,
                       min_mean_abundance=0.0,
                       colors_boxplot=COLORS, 
                       color_barplot='gray'
                      )
    p[0].get_axes()[0].set_ylabel("compound")
    
    p[0].delaxes(p[0].get_axes()[2])
    p[0].delaxes(p[0].get_axes()[1])
    
    p[0].get_axes()[0].set_ylabel("")
    p[0].get_axes()[0].set_title("relative abundance")
    p[0].get_axes()[0].set_xlabel("")
    p[0].suptitle("")
    plt.figtext(0.09, 0.89, 'D', fontsize=16)
    p[0].set_size_inches((4*3, 3))

# %% [markdown]
### Panel E

# %% [code]
%matplotlib inline
meta_tri = metadata[metadata['host_scientific_name'] == 'Mus musculus'].groupby('host_subject_id_bmfz').head(1).copy()
if meta_tri[meta_tri['generation'] == 'rm'].shape[0] < 12:
    meta_tri['category'] = meta_tri['generation'].apply(lambda x: 'RM & P mice' if x in ['P','RM'] else 'all mice')
    dupl_rm = meta_tri[meta_tri['generation'].isin(['RM','P'])].copy()
    dupl_rm['category'] = 'all mice'
    meta_tri = pd.concat([meta_tri, dupl_rm]).reset_index()

fig, axes = plt.subplots(1,1,figsize=(5,3))

hue_order = ['B6J','RM','C']
sns.boxplot(data=meta_tri, y='serum_triglyceride', x='category', hue='host_genotype', palette=COLORS, ax=axes, 
            hue_order=hue_order,
            dodge=True, fliersize=0)

sns.stripplot(data=meta_tri, y='serum_triglyceride', x='category', hue='host_genotype', palette=COLORS, ax=axes, 
              hue_order=hue_order,
              dodge=True, edgecolor='white', linewidth=1, legend=None)

annotator = Annotator(axes, 
    [(('RM & P mice', 'RM'), ('RM & P mice', 'B6J')), 
     (('RM & P mice', 'RM'), ('RM & P mice', 'C')), 
     
     (('all mice', 'RM'), ('all mice', 'C')), 
     (('all mice', 'RM'), ('all mice', 'B6J')), 
     
     (('RM & P mice', 'C'), ('all mice', 'C')), 
     ], 
    data=meta_tri, x='category', y='serum_triglyceride', hue='host_genotype', hue_order=hue_order)
annotator.configure(test='Mann-Whitney', text_format='star', loc='inside', comparisons_correction="fdr_bh", correction_format="default")
_ = annotator.apply_and_annotate()


axes.set_ylabel('nmol/well')
axes.set_xlabel('')
axes.set_title('serum triglyceride')

nums = meta_tri.dropna(subset=['serum_triglyceride']).groupby(['category', 'host_genotype']).size()
axTop = axes.twiny()
axTop.set_xlim(axes.get_xlim())
offset = 0.27
_ = axTop.set_xticks([0-offset, 0, 0+offset, 1-offset, 1, 1+offset],
                     ['n=%i' % x for x in  nums.loc[['RM & P mice', 'all mice'],hue_order].values])


# %% [markdown]
## Figure S1: gut alpha

# %% [code]
def draw_cageline_alphadiversity(meta, alpha, title):
    GENORDER = ['avg.'] + ORDER_GENERATIONS[1:-1]
    def _cageline_present(genotype, numbers, generation, numSamples):
        return all([genotype[0] + str(number) in numSamples.loc[generation].index for number in numbers])

    malpha = meta.merge(alpha, left_index=True, right_index=True)
    fig, axes = plt.subplots(alpha.shape[1], len(ORDER_GENOTYPE), figsize=(15,20), sharex=True)
    fig.subplots_adjust(hspace=0.01)

    for i, genotype in enumerate(ORDER_GENOTYPE):
        for j, metric in enumerate(reversed(alpha.columns)):
            data = malpha[malpha['host_genotype'] == genotype]
            avgdata = data.copy()
            avgdata['generation'] = 'avg.'
            avgdata.index = list(map(lambda x: 'avg.%s' % x, avgdata.index))
            data = pd.concat([data, avgdata], axis=0)
            hue_order = ["%s%i" % (genotype[0], n) for n in range(1,3+1)]
            sns.boxplot(data=data, y=metric, x="generation", order=GENORDER, hue='cage_line', hue_order=hue_order, ax=axes[j,i], color='lightgray')
            sns.stripplot(data=data, y=metric, x="generation", order=GENORDER, hue='cage_line', hue_order=hue_order, ax=axes[j,i], dodge=True)

            axes[j,i].axhline(y=data[metric].mean(), color='gray', linestyle='--')
            numSamples = data.groupby(['generation','cage_line']).size()
            
            annotator = Annotator(axes[j,i],
                ([(('F4', 'B2'), ('F5', 'B2')),
                  (('F4', 'B3'), ('F5', 'B3')),
                 
                  (('avg.', 'B2'), ('F5', 'B2')),
                  (('avg.', 'B3'), ('F5', 'B3'))
                 ] if (genotype == 'B6J') else []) + \
                [((generation, genotype[0]+'1'), (generation, genotype[0]+'2')) 
                 for generation 
                 in GENORDER 
                 if _cageline_present(genotype, [1, 2], generation, numSamples)] + \
                [((generation, genotype[0]+'1'), (generation, genotype[0]+'3')) 
                 for generation 
                 in GENORDER
                 if _cageline_present(genotype, [1, 3], generation, numSamples)]
                , data=data, y=metric, x="generation", order=GENORDER, hue='cage_line', hue_order=hue_order)
            annotator.configure(test='Mann-Whitney', text_format='star', loc='inside', comparisons_correction="fdr_bh", correction_format="default")
            annotator.apply_and_annotate()

            axes[j,i].get_legend().remove()
            #continue
            if j == 0:
                axTop = axes[j,i].twiny()
                axTop.set_xlim(axes[j,i].get_xlim())
                offset = 0.20
                axTop.set_xticks(sorted(list(axes[j,i].get_xticks() - 0.33) + \
                                        list(axes[j,i].get_xticks() + 0.33) + \
                                        list(axes[j,i].get_xticks())), ['%ii' % (numSamples.loc[generation, cage_line] if _cageline_present(genotype, [cage_line[-1]], generation, numSamples) else 0)
                                                                        for generation in GENORDER 
                                                                        for cage_line in sorted(list(data['cage_line'].dropna().unique())) 
                                                                       ])
                axes[j,i].set_title(genotype)

                axes[j,i].legend(handles=[
                    Line2D([0], [0], marker='o', color='#ffffff00', label='%s%i' % (genotype[0], n), markerfacecolor=sns.color_palette()[n-1]) for n in [1,2,3]
                    ],
                    title='%s: cage lineage' % genotype, ncol=3, bbox_to_anchor=(0.75, 1.25))


            if j < 2:
                axes[j,i].set_xlabel("")
            axes[j,i].set_ylabel(DM_NAMES.get(axes[j,i].get_ylabel(), axes[j,i].get_ylabel()))
    # fig.suptitle(title)
    
    plt.figtext(0.09, 0.87, 'A', fontsize=16)
    plt.figtext(0.09, 0.68, 'C', fontsize=16)
    plt.figtext(0.09, 0.48, 'E', fontsize=16)
    plt.figtext(0.09, 0.29, 'G', fontsize=16)
    
    plt.figtext(0.51, 0.87, 'B', fontsize=16)
    plt.figtext(0.51, 0.68, 'D', fontsize=16)
    plt.figtext(0.51, 0.48, 'F', fontsize=16)
    plt.figtext(0.51, 0.29, 'H', fontsize=16)

    return fig     

# %% [code]
%matplotlib inline
fig = draw_cageline_alphadiversity(meta_gut, res_gut['alpha_diversity']['results'], title='gut microbiome')
# fig.savefig("Figures/figS3_gutAlpha.svg", bbox_inches='tight')

# %% [markdown]
## Figure S2: Founding Sires
**Reviewer 1**: Was the different number of fathers used to produce the next generation between the two mouse strains taken into consideration/correct or in the analyses?

# %% [code]
metric = 'PD_whole_tree'
keyCL = 'C1 + C2 + C3'
order_clfather = ['B1','B2','B3',keyCL]
fig, axes = plt.subplots(res_gut['alpha_diversity']['results'].shape[1], 2,
                         figsize=(10 * 2, 5 * res_gut['alpha_diversity']['results'].shape[1]),
                         sharex=True)
fig.subplots_adjust(hspace=0.01)
COLORS.update({keyCL: COLORS['C']})

for col, (bodysite, alpha, meta) in enumerate([('gut', res_gut['alpha_diversity']['results'], meta_gut),
                                             ('skin', res_skin['alpha_diversity']['results'], meta_skin)]):
    data = meta.merge(alpha, left_index=True, right_index=True, how='right')
    data=data[(data['host_genotype'] != 'RM') & pd.notnull(data['generation'])]
    data['cl_father'] = data['cage_line'].apply(lambda x: keyCL if x.startswith('C') else x)
    numSamples = data.groupby(['generation', 'cl_father']).size().to_dict()

    for row, metric in enumerate(alpha.columns):
        ax = axes[row, col]
        sns.boxplot(data=data, y=metric, x='generation', hue='cl_father', palette=COLORS, hue_order=order_clfather, order=ORDER_GENERATIONS[1:-1], ax=ax)
        
        annotator = Annotator(ax, [
                ((generation, bcl), (generation, keyCL))
                    for generation in data['generation'].unique() for bcl in data[data['host_genotype'] == 'B6J']['cage_line'].unique()
                    if numSamples.get((generation, bcl), 0) > 0
                    if numSamples.get((generation, keyCL), 0) > 0
            ], data=data, y=metric, hue='cl_father', x='generation', hue_order=order_clfather, order=ORDER_GENERATIONS[1:-1])
        annotator.configure(test='Mann-Whitney', text_format='star', loc='inside', comparisons_correction="fdr_bh", correction_format="default")
        annotator.apply_and_annotate()

        ax.legend().remove() # set_title('cage line(s)')
        if row == 0:
            xticklabels = []
            for i, generation in enumerate(ORDER_GENERATIONS[1:-1]):
                for j, bcl in enumerate(order_clfather):
                    num = numSamples.get((generation, bcl), 0)
                    xticklabels.append('%ii' % num)
        
            xpos = []
            bias = 0.2
            for x in list(ax.get_xticks()):
                for i in range(4):
                    xpos.append(x - 1.5 * bias + bias*i)
                
            axTop = ax.twiny()
            axTop.set_xticks(xpos, xticklabels)
            axTop.set_xlim(ax.get_xlim())
            # ax.set_title(bodysite)

            ax.legend(handles=[
                    Line2D([0], [0], marker='s', color=COLORS[n], label=n, markerfacecolor=COLORS[n])
                        for n in order_clfather
                    ],
                    title='%s: cage lineage(s)' % bodysite, ncol=4, bbox_to_anchor=(0.75, 1.25))
            
        ax.set_ylabel(DM_NAMES.get(metric))


plt.figtext(0.09, 0.87, 'A', fontsize=16)
plt.figtext(0.09, 0.68, 'C', fontsize=16)
plt.figtext(0.09, 0.48, 'E', fontsize=16)
plt.figtext(0.09, 0.29, 'G', fontsize=16)

plt.figtext(0.51, 0.87, 'B', fontsize=16)
plt.figtext(0.51, 0.68, 'D', fontsize=16)
plt.figtext(0.51, 0.48, 'F', fontsize=16)
plt.figtext(0.51, 0.29, 'H', fontsize=16)

#_ = fig.suptitle('impact of number of founding sires on microbial diversity')
        
fig.savefig('tmp.svg', bbox_inches='tight')

# %% [markdown]
## Figure S3: taxonomy Robertson

# %% [code]
p = plotTaxonomy(res_rob['counts_plantsremoved'], meta_rob[meta_rob['host_body_site'] == 'UBERON:colon'].fillna('?'), file_taxonomy=res_rob['taxonomy']['RDP']['results']['Taxon'],
                 group_l1='qiita_study_id', rank='Phylum', group_l2='generation',
                 print_sample_labels=False, minreadnr=1000)
p[0].set_size_inches((25, 5))
p[0].savefig('Figures/figS9_taxonomyRobertson.svg')

# %% [markdown]
## Figure S4: skin diversity
Use `Generated/Emperor/emperor-pcoa_skin_weighted_unifrac.gzv` to upload to `view.qiime2.org`

### Panel A
see Figure 2, but load settings from file `EmperorSettings/emp_settings_skin_panelA.json`
![image-17.png](attachment:image-17.png)

### Panel B
see Figure 2, but load settings from file `EmperorSettings/emp_settings_skin_panelB.json`
![image-15.png](attachment:image-15.png)

### Panel C
see Figure 2, but load settings from file `EmperorSettings/emp_settings_skin_panelC.json`
![image-14.png](attachment:image-14.png)

### Panel D
see Figure 2, but load settings from file `EmperorSettings/emp_settings_skin_panelD.json`
![image-16.png](attachment:image-16.png)

# %% [markdown]
## Figure S5: skin alpha

# %% [code]
%matplotlib inline
fig = draw_cageline_alphadiversity(meta_skin, res_skin['alpha_diversity']['results'], title='skin microbiome')
fig.savefig("Figures/figS4_skinAlpha.svg", bbox_inches='tight')

# %% [markdown]
## Figure S6: Emperor skin and gut

# %% [code]
# create an emperor plot ONLY from samples of mice, bodysites gut or skin (skip resequenced batch 2 skin samples)
_ = emperor(metadata[(metadata['host_scientific_name'] == 'Mus musculus') & 
                     (metadata['host_body_site'] != 'UBERON:lung') & 
                     (~((metadata['batch'] == "2") & (metadata['host_body_site'] == 'UBERON:sebum')))
                    ],
            res_both['beta_diversity']['results'], fp_results='Generated/Emperor', infix='_gutskin', dry=False, wait=False)

# %% [markdown]
Use `Generated/Emperor/emperor-pcoa_gutskin_unweighted_unifrac` to upload to `view.qiime2.org`
see Figure 2, but load settings from file `EmperorSettings/emp_settings_gutANDskin.json`
![image-13.png](attachment:image-13.png)

# %% [markdown]
## Figure S7: skin source tracking

# %% [code]
def get_type(row):
    if row['Source'] == 'Unknown':
        return row['Source']
    elif row['Source'] == 'control_bedding':
        return "bedding"
    elif row['Source'] == 'control_food':
        return "food"
    elif row['Source'] == row['host_genotype']:
        return "feces from matching host genotype"
    elif row['Source'] != row['host_genotype']:
        return "feces from opposing host genotype"

meta_st2_gs = metadata[
                       (metadata['host_scientific_name'] != 'Callithrix jacchus') &
                        metadata['sample_type'].isin(['control bedding','control food','skin of external ear','colon content'])
                      ].copy().fillna({'generation': 'control'})

res_impact = []
res_st2_gs = dict()
for generation, g in meta_st2_gs[meta_st2_gs['sample_type'].isin(['colon content', 'skin of external ear'])].groupby('generation'):
    meta_gen = pd.concat([g, meta_st2_gs[meta_st2_gs['sample_type'].isin(['control bedding', 'control food'])]], axis=0)
    
    meta_gen['st2_type'] = 'Sink'
    meta_gen.loc[meta_gen[meta_gen['sample_type'].isin(['control bedding', 'control food', 'colon content'])].index, 'st2_type'] = 'Source'
    meta_gen['st2_env'] = meta_gen.apply(lambda r: r['host_genotype'] if pd.notnull(r['host_genotype']) else r['sample_type'], axis=1)
    res_st2_gs[generation] = sourcetracker2(res_both['counts_plantsremoved'], meta_gen, col_envname='st2_env', col_type='st2_type', dry=False, wait=False)
    
    if res_st2_gs[generation]['results'] is not None:
        d = res_st2_gs[generation]['results'].stack().reset_index().rename(columns={0: 'impact'}).merge(metadata[['host_genotype','generation','geo_loc_name']], left_on='Sink', right_index=True, how='left')
        d['type'] = d.apply(get_type, axis=1)
        d['generation'] = generation.replace('control', 'Controls')
       
        res_impact.append(d)
res_impact = pd.concat(res_impact)

# %% [code]
sns.set_style("whitegrid")
fig, axes = plt.subplots(1,1,figsize=(15,5))
sns.boxplot(data=res_impact, y='impact', hue='type', x='generation', order=ORDER_GENERATIONS, ax=axes,
            hue_order=['feces from matching host genotype', 'feces from opposing host genotype', 'bedding', 'food', 'Unknown'])
axes.legend(bbox_to_anchor=(1.1, 1.05))
_ = axes.set_title("Source Tracker: does gut microbiome seed skin microbiome? Using matching gut generation data as source.")
# fig.savefig('tmp.svg', bbox_inches='tight')

# %% [code]
res_impact.groupby(['Source','host_genotype'])['impact'].mean().loc[[('B6J','B6J'),('C','C')]].mean()

# %% [code]
res_impact.groupby(['Source','host_genotype'])['impact'].mean().loc[[('B6J','C'),('C','B6J')]].mean()

# %% [code]
res_impact.groupby(['Source','host_genotype'])['impact'].mean().loc['control_bedding'].mean()

# %% [code]
res_impact.groupby(['Source','host_genotype'])['impact'].mean().loc['control_food'].mean()

# %% [code]
res_impact.groupby(['Source','host_genotype'])['impact'].mean().loc['Unknown'].mean()

# %% [markdown]
## Figure S8: taxonomy barplots
### Panel A

# %% [code]
%matplotlib inline
fig = drawTaxBarplot(res_gut, meta_gut, rank='Phylum')
_ = plt.figtext(0.09, 0.89, 'A', fontsize=16)

# %% [markdown]
### Panel B

# %% [code]
%matplotlib inline
fig = drawTaxBarplot(res_skin, meta_skin,  title='skin microbiome', minreadnr=60*60*0.5, rank='Phylum')
_ = plt.figtext(0.09, 0.89, 'B', fontsize=16)
# fig.savefig('tmp.svg')

# %% [markdown]
### Panel C

# %% [code]
%matplotlib inline
fig = drawTaxBarplot(res_skin, meta_skin, title='skin microbiome', minreadnr=60*60*0.5)
_ = plt.figtext(0.09, 0.89, 'C', fontsize=16)
# fig.savefig('tmp.svg')

# %% [markdown]
## Figures S9: gut Genera

# %% [code]
#adding int to generation to properly list generations in upsetplots
conditions = [
    (meta_gut["generation"] == "RM"),
    (meta_gut["generation"] == "P"),
    (meta_gut["generation"] == "F1"),
    (meta_gut["generation"] == "F2"),
    (meta_gut["generation"] == "F3"),
    (meta_gut["generation"] == "F4"),
    (meta_gut["generation"] == "F5")
]

values = ["-1: RM", "0: P", "1: F1", "2: F2", "3: F3", "4: F4", "5: F5"]
meta_gut["i_gen"] = np.select(conditions, values, "other")
conditions = [
    (meta_skin["generation"] == "RM"),
    (meta_skin["generation"] == "P"),
    (meta_skin["generation"] == "F1"),
    (meta_skin["generation"] == "F2"),
    (meta_skin["generation"] == "F3"),
    (meta_skin["generation"] == "F4"),
    (meta_skin["generation"] == "F5")
]

values = ["-1: RM", "0: P", "1: F1", "2: F2", "3: F3", "4: F4", "5: F5"]
meta_skin["i_gen"] = np.select(conditions, values, "other")

# %% [code]
def pick_rank(counts, taxa, rank):
    # split information in taxa table per rank and assign proper names to each column
    taxa_split = taxa["Taxon"].str.split(pat=";", expand=True)
    taxa_split = taxa_split.rename(columns={0: "domain", 1: "phylum", 2: "class", 3: "order", 4: "family", 5: "genus", 6: "species"})
    # copy counts DataFrame to avoid chagnes being made to imported feature tables
    taxa_counts = counts.copy()
    taxa_counts[rank] = taxa_split[rank]
    # grouping taxa by rank, collapsing feature tables
    grouped_counts = taxa_counts.groupby(rank).sum(numeric_only=True)
    return(grouped_counts)

def make_upset(counts, meta, category, plotname, taxa=None, rank=None, title="", facecolor="black"):
    # if rank is give, collapse feature table to rank, else process ASVs
    if rank != None:
        upset = pick_rank(counts, taxa, rank)
        upset = upset.transpose()
    else:
        upset = counts.transpose()
    # adding category to feature table + control to filter out control samples
    upset[category] = meta[category]
    upset['control'] = meta['control']
    upset = upset.drop(upset[upset['control'].isin(["Duesseldorf","Janvier"])].index)
    # grouping by category and omitting taxa not present in all samples
    upset_grouped = upset.groupby(category).sum(numeric_only=True)
    upset_grouped = upset_grouped.loc[:, (upset_grouped != 0).any(axis=0)]
    # assign boolean values of True/False depending on whether values are > 0 or not, combining with the actual values and assigning the boolean columns as indeces
    upset_grouped = upset_grouped.transpose()
    over_zero = upset_grouped > 0
    over_zero = over_zero.rename(columns=lambda x: x + ' ')
    upset_combined = pd.concat([upset_grouped, over_zero],axis=1)
    upset_combined = upset_combined.set_index(list(over_zero.columns))
    # plotting upsetplot with the plot() function
    fig = upsetplot.plot(upset_combined, show_counts=True, subset_size="count", sort_categories_by="-input", facecolor=facecolor)
    plt.title(title)
    plt.figtext(0.09, 0.89, plotname, fontsize=16)
    # plt.savefig('tmp.svg')

# %% [markdown]
### Panel A

# %% [code]
sns.set_style("white")
make_upset(res_gut['counts_plantsremoved'], meta_gut, "host_genotype", "A", res_gut['taxonomy']['RDP']['results'], "genus", title='host genotype')

# %% [markdown]
### Panel B

# %% [code]
make_upset(res_gut['counts_plantsremoved'].loc[:, meta_gut[meta_gut['host_genotype'] == 'B6J'].index], 
           meta_gut, "cage_line", "B", res_gut['taxonomy']['RDP']['results'], "genus", title='B6J: maternal legacy', facecolor=COLORS['B6J'])

# %% [markdown]
### Panel C

# %% [code]
make_upset(res_gut['counts_plantsremoved'].loc[:, meta_gut[meta_gut['host_genotype'] == 'C'].index], 
           meta_gut, "cage_line", "C", res_gut['taxonomy']['RDP']['results'], "genus", title='C: maternal legacy', facecolor=COLORS['C'])

# %% [markdown]
### Panel D

# %% [code]
make_upset(res_gut['counts_plantsremoved'].loc[:, meta_gut[meta_gut['host_genotype'].isin(['B6J', 'RM'])].index], 
           meta_gut, "i_gen", "D", res_gut['taxonomy']['RDP']['results'], "genus", title="B6J: generations", facecolor=COLORS['B6J'])

# %% [markdown]
### Panel E

# %% [code]
make_upset(res_gut['counts_plantsremoved'].loc[:, meta_gut[meta_gut['host_genotype'].isin(['C', 'RM'])].index], 
           meta_gut, "i_gen", "E", res_gut['taxonomy']['RDP']['results'], "genus", title='C: generations', facecolor=COLORS['C'])

# %% [markdown]
## Figures S10: skin Genera

# %% [markdown]
### Panel A

# %% [code]
sns.set_style("white")
make_upset(res_skin['counts_plantsremoved'], meta_skin, "host_genotype", "A", res_skin['taxonomy']['RDP']['results'], "genus", title='host genotype')

# %% [markdown]
### Panel B

# %% [code]
make_upset(res_skin['counts_plantsremoved'].loc[:, meta_skin[meta_skin['host_genotype'] == 'B6J'].index], 
           meta_skin, "cage_line", "B", res_skin['taxonomy']['RDP']['results'], "genus", title='B6J: maternal legacy', facecolor=COLORS['B6J'])

# %% [markdown]
### Panel C

# %% [code]
make_upset(res_skin['counts_plantsremoved'].loc[:, meta_skin[meta_skin['host_genotype'] == 'C'].index], 
           meta_skin, "cage_line", "C", res_skin['taxonomy']['RDP']['results'], "genus", title='C: maternal legacy', facecolor=COLORS['C'])

# %% [markdown]
### Panel D

# %% [code]
make_upset(res_skin['counts_plantsremoved'].loc[:, meta_skin[meta_skin['host_genotype'].isin(['B6J', 'RM'])].index], 
           meta_skin, "i_gen", "D", res_skin['taxonomy']['RDP']['results'], "genus", title="B6J: generations", facecolor=COLORS['B6J'])

# %% [markdown]
### Panel E

# %% [code]
make_upset(res_skin['counts_plantsremoved'].loc[:, meta_skin[meta_skin['host_genotype'].isin(['C', 'RM'])].index], 
           meta_skin, "i_gen", "E", res_skin['taxonomy']['RDP']['results'], "genus", title='C: generations', facecolor=COLORS['C'])

# %% [markdown]
# Additional Analyses to address Reviewer concerns
## Joint analysis with Robertson et al. data
**Reviewer 2**: "Lines 655-656.  Speaking of this, how was this comparison conducted?  Were the reads of the separate study converted to genus-level abundances?  This is not stated in the methods."

# %% [markdown]
### Collapse on Genus

# %% [code]
# as not a single ASV overlap between Robertson's and our data, we need to collapse on higher taxonomic rank to 
# make the feature table accessible to e.g. Bray-Curtis or other non phylogenetic metrics
ranks = settings.RANKS + ['raw']
fig, axes = plt.subplots(1, len(ranks), figsize=(20, 20*len(ranks)))
counts_rob_collapsed = dict()
for ax, rank in zip(axes, ranks):
    counts_rob_collapsed[rank] = collapseCounts_objects(res_rob['rarefaction']['results'], rank, res_rob['taxonomy']['RDP']['results']['Taxon'])[0]
    asvs_rob = dict()
    for study in meta_rob['qiita_study_id'].unique():
        asvs_rob[study] = {asv for asv, count in counts_rob_collapsed[rank].loc[:, meta_rob[meta_rob['qiita_study_id'] == study].index].sum(axis=1).items() if count > 0}
    venn2(asvs_rob.values(), set_labels=list(map(lambda x: x.replace('13422', 'Benga'), asvs_rob.keys())), ax=ax)
    ax.set_title(rank.replace('raw', 'ASVs'))

# %% [code]
beta_rob_genus = beta_diversity(counts_rob_collapsed['Genus'], metrics=['bray_curtis'], dry=False, wait=True)

# %% [code]
sns.set_style('white')
fig = plot_robertson(beta_rob_genus['results'], metric='bray_curtis')

# %% [markdown]
### Determine variable region lengths

# %% [code]
# alignment positions obtained from http://qiime.org/home_static/nih-cloud-apr2012/variable_region_position_calculations.pdf
alnpos_gg138 = {
    'upstream': (0, 1916),
    'v3.v4': (1916, 4051),
    'v3': (1916, 2232),
    'v4': (2263, 4051),
    'downstream': (4051, 99999)}

# we have downloaded GreenGenes 13.8 from 
# ftp://greengenes.microbio.me/greengenes_release/gg_13_5/gg_13_8_otus.tar.gz and extracted into directory /vol/jlab/MicrobiomeAnalyses/References/gg_13_5_otus/
fp_gg138 = '/vol/jlab/MicrobiomeAnalyses/References/gg_13_5_otus'

# %% [code]
# read the GG13.8 alignment with 99322 sequences and 7682 columns
msa_gg138 = TabularMSA.read(os.path.join(fp_gg138, 'rep_set_aligned/97_otus.fasta'), format='fasta', constructor=DNA)

region_seqs = dict()
region_lengths = dict()
for region in alnpos_gg138.keys():
    # slice alignment to those columns that belong to a variable region
    submsa_gg138 = msa_gg138.iloc[:, alnpos_gg138[region][0]:alnpos_gg138[region][-1]]
    
    # degap alignment rows and store together with ID
    region_seqs[region] = dict()
    for row in submsa_gg138:
        seq = row.degap()
        region_seqs[region][row.metadata['id']] = str(seq)
    region_seqs[region] = pd.Series(region_seqs[region]).to_frame().rename(columns={0: 'amplicon'})
    
    region_lengths[region] = region_seqs[region]['amplicon'].apply(len).mean()
    print('region %s avg. length: %.0fbp' % (region, region_lengths[region]))

# %% [code]
sns.set_style('whitegrid')
def add_interval(ax, xdata, ydata, caps="  ", label=None, color=None, va_pos='bottom', ha_pos='center', text_xpos=None):
    line = ax.add_line(mpl.lines.Line2D(xdata, ydata, label=label, color=color))
    anno_args = {
        'ha': 'center',
        'va': 'center',
        'size': 20,
        'color': line.get_color() if color is None else color
    }
    a0 = ax.annotate(caps[0], xy=(xdata[0], ydata[0]), **anno_args)
    a1 = ax.annotate(caps[1], xy=(xdata[1], ydata[1]), **anno_args)
    xpos = xdata[0] + ((xdata[1]-xdata[0])/2)
    if text_xpos is not None:
        xpos = text_xpos
    atext = ax.annotate(label, xy=(xpos, ydata[0]), **{'ha': ha_pos, 'va': va_pos, 'size': 14})
    return (line,(a0,a1, atext))


fig, ax = plt.subplots()
offset = region_lengths['upstream']
add_interval(ax, (0, sum(region_lengths.values())), (4,4), "||", "16S rRNA", color='black', text_xpos=450)
add_interval(ax, (offset, offset + region_lengths['v3.v4']), (3,3), "||", "V3-V4")
add_interval(ax, (offset, offset + region_lengths['v3']), (2,2), "||", "V3")
add_interval(ax, (offset + region_lengths['v3.v4']-region_lengths['v4'], offset + region_lengths['v3.v4']), (2,2), "||", "V4")

add_interval(ax, (offset, offset + 250), (1,1), "||", "QC fwd reads", color='red')#, ha_pos='right')
add_interval(ax, (offset + region_lengths['v3.v4'] - 125, offset + region_lengths['v3.v4']), (1,1), "||", "QC rev reads", color='red')#, va_pos='top')#, ha_pos='left')

ax.set_xlim((250, 700))
ax.set_ylim((0,5))
_ = ax.set_xlabel('16S rRNA coordinates in bp')

# ax.legend()

# %% [markdown]
### determine read quality

# %% [code]
# we downloaded "All raw data" through Qiita's study 13422 as file FromQiita/study_raw_data_13422_082124-000334.zip

fp_tmp = 'tmp_dir/ReadQualities/'
os.makedirs(os.path.join(fp_tmp, 'per_sample_FASTQ/126766'), exist_ok=True)
# extract all read files. Note, this also contains unused ITS reads
cluster_run(['unzip FromQiita/study_raw_data_13422_082124-000334.zip -d %s' % fp_tmp], 'unzip', os.path.join(fp_tmp, 'per_sample_FASTQ/126766/46skin_S122_L001_R2_001.fastq.gz'), dry=False)

# %% [code]
# run fastqc for each 16S file
for fp in glob(os.path.join(fp_tmp, 'per_sample_FASTQ/126766/*.fastq.gz')):  # 149194 is ITS
    cmd = 'cd %s; /vol/software/bin/fastqc %s' % (os.path.dirname(os.path.abspath(fp)), os.path.basename(fp))
    fp_res = os.path.abspath(fp).replace('.fastq.gz', '_fastqc.html')
    cluster_run([cmd], 'fastqc', fp_res, ppn=1, dry=False, wait=False)

# %% [code]
# execute multiQC on fwd and rev read files separately
for orient in ['R1', 'R2']:
    fp_res = os.path.dirname(os.path.abspath(fp)) + '/res_multiqc_%s' % orient
    cmd = 'cd %s; multiqc --ignore-samples "*_%s_*" --outdir %s . ' % (os.path.dirname(os.path.abspath(fp)), 'R1' if orient == 'R2' else 'R2', fp_res)
    cluster_run([cmd], 'multiqc', fp_res, ppn=1, dry=True, wait=False, environment='ggmap_spike')

# %% [code]
# extract and display mean quality scores plot from multiQC report
for fp_report in sorted(glob(os.path.join(fp_tmp, 'per_sample_FASTQ/126766/res_multiqc_*/*.html'))):
    with open(fp_report, 'r') as f:
        for line in f.readlines():
            if 'id="mqc_fastqc_per_base_sequence_quality_plot' in line:
                png_search = re.search('img src="(.*)" /></div>', line)
                g = Image(url=png_search[1])
                print("Forward reads:" if '_R1/' in fp_report else 'Reverse reads:')
                display(g)

# %% [markdown]
## Quality control between Qiime1 and Qiime2
**Reviewer 3**: "In the methods section, it is indicated that QIIME 1.9.1 was used for QC, and downstream analysis was performed with QIIME2. The approaches for quality control in QIIME2 are much better than what was available in QIIME 1.9.1. It’s not clear what the rationale for this approach was. (See: [QIIME Forum](https://forum.qiime2.org/t/analyzed-data-with-qiime-1-9-1-should-we-go-back-and-reanalyze-with-qiime2/2028))"

# %% [code]
pick = [
    'Total number of input sequences',
    'Read too short after quality truncation',
    'Count of N characters exceeds limit',
    'Illumina quality digit',
    'Total number seqs written']
        
data = []
with open('FromQiita/126769_split_library_log.txt', 'r') as f:
    res = dict()
    for line in f.readlines():
        for field in pick:
            if field in line:
                res[field] = int(line.split()[-1].strip())
        if '---' in line:
            data.append(res)
            res = dict()
data = pd.DataFrame(data)

# %% [code]
data.sum()

# %% [markdown]
## check cross contamination

# %% [code]
for (flowcell, plate), g in metadata.groupby(['flowcell', 'plate_number']):
    #print(plate, g.shape, g['flowcell'].value_counts())
    #break
    #continue
    m = g.copy()
    m['st2_type'] = 'Source'
    m.loc[m[m['sample_type'] == 'sterile water'].index, 'st2_type'] = 'Sink'
    m['st2_env'] = m.index
    res_st2 = sourcetracker2(counts, m, 'st2_env', 'st2_type', dry=False, wait=False)
    s = res_st2['results'].sum(axis=1)
    s.name = 'sourcetracker'
    m = m.merge(s, left_index=True, right_index=True, how='left')
    m['well'] = m['plate_row'] + m['plate_column']
    m['name'] = list(map(lambda x: x[6:], m.index))
    
    x = cm.ScalarMappable(norm=mpl.colors.Normalize(vmin=0.0, vmax=1.0), cmap=cm.Reds)
    soi_col = {val: x.to_rgba(float(val)) for val in m['sourcetracker'].fillna(0.0).unique()}
    soi_col.update({'nan': x.to_rgba(0.0)})
    
    fig = plot_plate(m, col_position='well', col_label='sourcetracker', col_texts='name', colors=soi_col,
                     highlight_samples=set(m[m['sample_type'] == 'sterile water'].index))
    fig[0].get_axes()[0].set_xlabel(' '.join(['Flowcell: %s' % flowcell, ', Plate %s' % plate]))

# %% [markdown]
## Non-Phylogenetic Metrics
**Reviewer 1**: "Ecological and statistical analyses: The authors used exclusively phylogeny-based indices for measuring alpha, and beta diversity when assessing the impact of host genetics, and maternal legacy. It would be interesting to asses further indices which consider other aspects of community structure such as evenness, expected richness, structure/ presence – absence including Shannon, Chao1, Bray-Curtis, Jaccard indices when quantifying the role of host genotype, and maternal effect"

# %% [markdown]
### Figure 2
#### Panels A, B, C

# %% [code]
%matplotlib inline
fig = draw_temporals(res_gut, meta_gut, num_permutations=9999,
                     metric_alpha='shannon', ALPHA_LIM=(5.25,6.6),
                     METRIC_BETA='bray_curtis', BETA_LIM=(0.25, 0.8))

# %% [code]
%matplotlib inline
fig = draw_temporals(res_gut, meta_gut, num_permutations=9999,
                     metric_alpha='chao1', ALPHA_LIM=(160, 350),
                     METRIC_BETA='jaccard', BETA_LIM=(0.2, 0.8))

# %% [markdown]
#### Panel D

# %% [code]
%matplotlib inline
fig = draw_maternalLegacy(meta_gut, res_gut['beta_diversity'], metric='bray_curtis')

# %% [code]
%matplotlib inline
fig = draw_maternalLegacy(meta_gut, res_gut['beta_diversity'], metric='jaccard')

# %% [markdown]
### Figure 4

# %% [code]
for metric in res_gut['beta_diversity']['results'].keys():
    sns.set_style("ticks")
    rred = redundancy_analysis_beta(meta_gut, 
                                    res_gut['beta_diversity']['results'][metric], 
                                    metric,
                                    categorials=['host_genotype','independent_cage_lineage','sex'], 
                                    intervals=[], dates={},
                                    ordinals={'generation': ORDER_GENERATIONS},
                                    title='Effect Sizes Gut Microbiome',
                                    dry=False, wait=False, #dirty=True,, 
                                    seed=42  # set a seed to the permutations engine of ordiR2step: https://stackoverflow.com/questions/30251636/reproducing-results-from-ordistep
                                   )

    if rred['results'] is not None:
        # style the figure
        axes = rred['results']['figure'].get_axes()[0]
        axes.set_title('Effect Sizes Gut Microbiome')
        axes.set_ylabel("")
        axR = axes.twinx()
        axR.set_ylabel(DM_NAMES.get(metric))
        axR.set_yticks([])
        axes.set_yticklabels([t.get_text().replace('independent_cage_lineage', 'maternal legacy').replace('host_genotype', 'host genotype') for t in axes.get_yticklabels()])
        rred['results']['figure'].set_size_inches((6, 3))
    # rred['results']['figure'].savefig('Figures/fig6_gutEffectSizes.svg', bbox_inches='tight')
    #rred['results']['figure']

# %% [markdown]
### Figure 5
#### Panels A, B, C

# %% [code]
%matplotlib inline
fig = draw_temporals(res_skin, meta_skin, 
                     metric_alpha='shannon', ALPHA_LIM=(1.5,6.6),
                     METRIC_BETA='bray_curtis', BETA_LIM=(0.5, 1),
                     title='skin microbiome')

# %% [code]
%matplotlib inline
fig = draw_temporals(res_skin, meta_skin, 
                     metric_alpha='chao1', ALPHA_LIM=(12, 125),
                     METRIC_BETA='jaccard', BETA_LIM=(0.77, 0.98),
                     title='skin microbiome')

# %% [code]
%matplotlib inline
fig = draw_temporals(res_skin, meta_skin, 
                     metric_alpha='observed_features', ALPHA_LIM=(12, 125),
                     METRIC_BETA='jaccard', BETA_LIM=(0.77, 0.98),
                     title='skin microbiome')

# %% [markdown]
### Panel D

# %% [code]
%matplotlib inline
fig = draw_maternalLegacy(meta_skin, res_skin['beta_diversity'], metric='bray_curtis')

# %% [code]
%matplotlib inline
fig = draw_maternalLegacy(meta_skin, res_skin['beta_diversity'], metric='jaccard')

# %% [markdown]
## Dual Barcodes
**Reviewer 1**: "Did the authors use a dual barcoding approach to amplify and sequence the 16S rRNA?"

# %% [code]
metadata.groupby(['i5_index_id', 'i5_index']).size()

# %% [code]
sorted(metadata['i5_index'].dropna().apply(lambda x: x.split('-')[0]).unique())

# %% [code]
metadata.groupby(['i7_index_id', 'i7_index']).size()

# %% [code]
sorted(metadata['i7_index'].dropna().apply(lambda x: x.split('-')[0]).unique())

# %% [markdown]
## Short Sequences
**Reviewer 1**: "Why amplify a long 16S rRNA region (v3-v4) around 400 bp, use the 2x300 cycles sequencing technology, to then cut the sequences down to 150 bp? Which will impact the taxonomic resolution. Please explain why sequences needed to be cut this short"

### Merge through DADA2

# %% [code]
# run DADA2 on our data
# 1. assign samples to read files
manifest = metadata[['run_prefix']].dropna().copy()
for idx, row in tqdm(manifest.iterrows()):
    files = sorted(glob(os.path.join(fp_tmp, 'per_sample_FASTQ/126766/%s*.fastq.gz' % row['run_prefix'])))
    assert len(files) == 2, row['run_prefix']
    manifest.loc[idx, 'forward-absolute-filepath'] = os.path.abspath(files[0])
    manifest.loc[idx, 'reverse-absolute-filepath'] = os.path.abspath(files[1])
manifest

# %% [code]
# 2. create manifest files for qiime2 import
DADASTYLES = ['single', 'paired']
fps_manifest = dict()
for style in DADASTYLES:
    fps_manifest[style] = os.path.join('tmp_dir/SequenceLength/manifest_%s.tsv' % style)
    os.makedirs(os.path.dirname(fps_manifest[style]), exist_ok=True)
    if style == 'single':
        manifest.rename(columns={'forward-absolute-filepath': 'absolute-filepath'})[['absolute-filepath']].to_csv(fps_manifest[style], sep="\t", index_label='sample-id')
    else:
        manifest[['forward-absolute-filepath', 'reverse-absolute-filepath']].to_csv(fps_manifest[style], sep="\t", index_label='sample-id')

fps_manifest

# %% [code]
# 3. import read to qiime2
fps_demux = dict()
for style in DADASTYLES:
    _type = 'PairedEndSequencesWithQuality'
    _format = 'PairedEndFastqManifestPhred33V2'
    if style == 'single':
        _type = 'SequencesWithQuality'
        _format = 'SingleEndFastqManifestPhred33V2'
    fps_demux[style] = os.path.join(os.path.dirname(fps_manifest[style]), 'demux_%s.qza' % style)
    cmd = 'qiime tools import --type "SampleData[%s]" --input-format %s --input-path %s --output-path %s' % (_type, _format, os.path.abspath(fps_manifest[style]), os.path.abspath(fps_demux[style]))
    cluster_run([cmd], 'q2-import', fps_demux[style], 'qiime2-amplicon-2023.9', dry=False, wait=False, ppn=1)


# %% [code]
# 4. execute dada2
## fwd to rev gap for 389 long V3-V4 region is 389-250-125=14bp
## DADA2 requires 21bp overlap --> 14+21 = 35bp
##   add 1bp to make it even --> 36bp
##   36/2 = we need 18bp of both reads more than QC would indicate
fps_dada2 = dict()
for style in DADASTYLES:
    _method = 'denoise-paired'
    _revtrunc = ' --p-trunc-len-r 143 '  # this is 125 rev length plus 18
    _fwdtrunc = ' --p-trunc-len-f 268 '  # this is 250 fwd length plus 18
    if style == 'single':
        _method = 'denoise-single'
        _revtrunc = ''
        _fwdtrunc = ' --p-trunc-len 250 '
    fps_dada2[style] = os.path.join(os.path.dirname(fps_manifest[style]), 'dada2_%s/table.qza' % style)
    os.makedirs(os.path.dirname(fps_dada2[style]), exist_ok=True)
    fp_output = os.path.abspath(os.path.dirname(fps_dada2[style]))
    cmds = ['rm -rf %s' % fp_output, 
            'qiime dada2 %s --i-demultiplexed-seqs %s %s %s --p-n-threads 10 --output-dir %s ' % (_method, os.path.abspath(fps_demux[style]), _revtrunc, _fwdtrunc, fp_output)]
    print('; '.join(cmds))
    cluster_run(cmds, 'dada2', fps_dada2[style], 'qiime2-amplicon-2023.9', dry=False, wait=False, ppn=20)

# %% [code]
# 5. extract qiime2 dada2 results
res_dada2 = dict()
for style in DADASTYLES:
    for obj in ['table', 'denoising_stats']:
        fp_res = os.path.join(os.path.dirname(os.path.abspath(fps_dada2[style])), 'flat_%s' % obj)
        cmd = 'unzip %s -d %s' % (os.path.abspath(os.path.join(os.path.dirname(fps_dada2[style]), '%s.qza' % obj)), fp_res)
        if not os.path.exists(fp_res):
            !$cmd
        if obj == 'table':
            fp = glob(os.path.join(fp_res, '*/data/feature-table.biom'))[0]
            res_dada2[(style, obj)] = biom2pandas(fp)
        elif obj == 'denoising_stats':
            fp = glob(os.path.join(fp_res, '*/data/stats.tsv'))[0]
            res_dada2[(style, obj)] = pd.read_csv(fp, sep="\t", index_col=0, comment='#')

# %% [code]
cols = ['input', 'filtered', 'denoised', 'merged', 'non-chimeric']
data = []
for style in DADASTYLES:
    x = res_dada2[(style, 'denoising_stats')].stack().reset_index().rename(columns={'level_1': 'processing step', 0: 'number of reads'})
    x['style'] = style
    data.append(x)
data = pd.concat(data)
data = data[data['processing step'].isin(cols)]
sns.barplot(data, y='number of reads', x='processing step', order=cols, hue='style')

print('ratio of DADA2 reads from all reads in samples')
sumdata = pd.pivot_table(data, index='style', columns='processing step', values='number of reads', aggfunc=np.mean)
sumdata[cols[-1]] / sumdata[cols[0]]

# %% [code]
data = []
for style in DADASTYLES:
    counts = res_dada2[(style, 'table')].loc[:, meta_gut.index]
    counts = counts[counts.sum(axis=1) >= 10]
    data.append((counts > 0).sum().rename(style))
    print(style, counts.shape)
data = pd.concat(data, axis=1).stack().reset_index().rename(columns={'level_1': 'style', 0: 'number of ASVs'})

# %% [code]
sns.stripplot(data=data, y='number of ASVs', x='style', palette=sns.color_palette())
data.groupby('style')['number of ASVs'].mean()

# %% [markdown]
### 250bp via Deblur

# %% [code]
fp_sepp_tree250 = "FromQiita/204144_insertion_tree.relabelled.tre"
sepp_tree250 = TreeNode.read(fp_sepp_tree250, format="newick")

# %% [code]
res_gut250 = process_study(
    metadata=meta_gut,
    control_samples=set([]),  # we already filtered out all control samples
    fp_deblur_biom=biom2pandas('FromQiita/204144_reference-hit.biom'),  # "abuse" argument to provide a table instead of a filepath
    fp_insertiontree=fp_sepp_tree250,
    rarefaction_depth=3164, #6000,
    fp_taxonomy_trained_classifier_gg138_chloroMitoRemoval='/vol/jlab/MicrobiomeAnalyses/References/Q2-Naive_Bayes_classifiers/gg-13-8-99-nb-classifier_2022.11.qza',  # as GG2 currently lacks labels for mitochondria/chloroplasts, see https://forum.qiime2.org/t/taxonomy-filtering-greengenes2/28334
    fp_taxonomy_trained_classifier="/vol/jlab/MicrobiomeAnalyses/References/Q2-Naive_Bayes_classifiers/gg_2022_10_backbone_full_length.nb_2023.5.qza",  # obtain this file from https://docs.qiime2.org/2023.9/data-resources/
    tree_insert=sepp_tree250,
    is_v4_region=False,
    emperor_fp='Generated/Emperor',
    emperor_infix='_gut250',
    dry=False,
    use_grid=True,
    fix_zero_len_branches=False, # since this has already been corrected via the project_sepp call
    emperor_skip_tsne_umap=True,
    
    alpha_metrics=["PD_whole_tree", "shannon", "observed_features", "chao1"]
)

# %% [code]
data = pd.concat([(res_gut['counts_plantsremoved'] > 0).sum().rename('150bp'),
                  (res_gut250['counts_plantsremoved'] > 0).sum().rename('250bp')], axis=1).stack().reset_index().rename(
                    columns={0: 'number of ASVs', 'level_1': 'ASV length'})
sns.stripplot(data=data, y='number of ASVs', x='ASV length', palette=sns.color_palette())
data.groupby('ASV length')['number of ASVs'].mean()

# %% [code]
data = []
for l, res in [(150, res_gut), (250, res_gut250)]:
    x = res['alpha_diversity']['results'].copy()
    x.index.name = 'sample_name'
    x = x.stack().reset_index()
    x['ASV length'] = '%ibp' % l
    data.append(x)
data = pd.concat(data).rename(columns={'level_1': 'metric', 0: 'alpha diversity'})
data

# %% [code]
fig, axes = plt.subplots(1, len(data['metric'].unique()), figsize=(5*len(data['metric'].unique()), 5))
for (metric, g), ax in zip(data.groupby('metric'), axes):
    sns.barplot(g, y='alpha diversity', x='ASV length', ax=ax)
    ax.set_ylabel(DM_NAMES.get(metric, metric))

# %% [code]
for l, res in [(150, res_gut), (250, res_gut250)]:
    tax = res['taxonomy']['RDP']['results'].loc[res['rarefaction']['results'].index, :]
    print(l, tax['Confidence'].mean(), tax[tax['Taxon'].apply(lambda x: 's__' in x and x.split('s__')[-1] != "")].shape[0])

# %% [code]
859/327

# %% [markdown]
## F5 anomaly
**Reviewer 1**: "Results, Figure 1:  Please explain why in generation 5, also host genotype does not impact the microbiome"

# %% [code]
CGORDER = ['B1','B2','B3']
sns.barplot(
    data=meta_gut[meta_gut['host_genotype'] == 'B6J'].groupby(['generation', 'cage_line']).size().reset_index().rename(columns={0: 'number of samples'}),
    y='number of samples',
    x='generation', order=ORDER_GENERATIONS[1:-1], hue='cage_line', hue_order=CGORDER
)

# %% [code]
meta_gut[meta_gut['host_genotype'] == 'B6J'].groupby(['generation', 'cage_line']).size().reset_index().rename(columns={0: 'number of samples'})

# %% [code]
avgnums = meta_gut[meta_gut['host_genotype'] == 'B6J'].groupby(['generation', 'cage_line']).size().reset_index().rename(columns={0: 'number of samples'}).groupby('generation')['number of samples'].mean()
avgnums.mean()

# %% [markdown]
**Reviewer 3**: "What is the explanation that there are no differences in microbiome composition at F5? Would it be expected that microbiome changes are more divergent across generations? (L349)"

# %% [code]
idx_f5 = meta_gut[(meta_gut['host_genotype'] == 'B6J') & (meta_gut['generation'] == 'F5')].index
counts_f5 = res_gut['rarefaction']['results'].loc[:, idx_f5]
counts_f5 = counts_f5[counts_f5.sum(axis=1) > 0]

counts_nonf5 = res_gut['rarefaction']['results'].loc[:, list(set(meta_gut.index) - set(idx_f5))]
counts_nonf5 = counts_nonf5[counts_nonf5.sum(axis=1) > 0]

fig, axes = plt.subplots(1,1)
venn2([set(counts_f5.index), set(counts_nonf5.index)], ['F5', 'all but F5'], ax=axes)
axes.set_title('number of ASVs')

# %% [code]
ratio_f5asvs = (res_gut['rarefaction']['results'].loc[counts_f5.index, :].sum() / 6000).rename('relative abundance')
fig, axes = plt.subplots(1,1)
sns.boxplot(meta_gut[(meta_gut['host_genotype'] == 'B6J')].merge(ratio_f5asvs, left_index=True, right_index=True, how='right'),
            y='relative abundance',
            x='generation', order=ORDER_GENERATIONS[1:-1],
            hue='cage_line', hue_order=CGORDER,
            ax=axes
           )
axes.set_title('relative abundance of ASVs found in B6J-F5 samples')

# %% [markdown]
## control mouse difference
**Reviewer 3:** What is the explanation for no differences in microbiome composition between controls? Should these mice resemble the genetic effect over the environmental exposure? (Figure 2A, right panel)

# %% [code]
results = []
for col, (bodysite, res, meta) in enumerate([('gut', res_gut, meta_gut),
                                             ('skin', res_skin, meta_skin)]):
    meta = meta[pd.isnull(meta['generation'])]  # restrict to control samples
    for location, g in meta.groupby('geo_loc_name'):
        print(bodysite, location, g.shape)
        tres = plotNetworks('host_genotype', g, None, res['beta_diversity']['results'], summarize=True, b_min_num=1, permutations=9999)[0]
        tres['body site'] = bodysite
        tres['vendor'] = location
        results.append(tres)
        #tres.set_index(['
        #break
    #break
results = pd.concat(results)
results = results[['metric', 'p-value', 'test-statistic', 'num samples x', 'num samples y', 'body site', 'vendor']].set_index(['body site', 'vendor'])
results = results.rename(columns={'num samples x': '#i B6J', 'num samples y': '#i C'})
results['metric'] = results['metric'].apply(lambda x: DM_NAMES[x])
results

# %% [markdown]
## Paternal Legacy
**Reviewer 1**: Was the different number of fathers used to produce the next generation between the two mouse strains taken into consideration/correct or in the analyses?

# %% [code]
%matplotlib inline
for bcl in sorted(meta_gut[meta_gut['host_genotype'] == 'B6J']['cage_line'].dropna().unique()):
    subset = meta_gut[meta_gut['cage_line'].isin([bcl,'C1','C2','C3', np.nan])].index
    print(bcl, len(subset))
    fig = draw_temporals(res_gut, meta_gut.loc[subset, :], num_permutations=9999)
    fig.suptitle('single B6J cage line: %s\n%s' % (bcl, fig._suptitle.get_text()))

# %% [code]
meta_gut['flowcell'].unique()

# %% [code]
meta_skin['flowcell'].unique()

# %% [code]
```
