## Additional File 2: Analysis repository for "The host genotype actively shapes its microbiome across generations in laboratory mice": makefile

```
PNGDPI=300

SVGFILES = $(wildcard fig*.svg)

figures = PDF/GA.pdf \
	  PDF/fig1_breedingStrategy.pdf \
	  PDF/fig2_gutTemporal.pdf \
	  PDF/fig3_gutDiversity.pdf \
	  PDF/fig4_gutEffectSizes.pdf \
	  PDF/fig5_skinTemporal.pdf \
	  PDF/fig6_gutTaxonomy.pdf \
	  PDF/fig7_metabolites.pdf \
	  PDF/figS1_gutAlpha.pdf \
	  PDF/figS2_foundingSires.pdf \
	  PDF/figS3_taxonomyRobertson.pdf \
	  PDF/figS4_skinDiversity.pdf \
	  PDF/figS5_skinAlpha.pdf \
	  PDF/figS6_emperorSkinGut.pdf \
	  PDF/figS7_skinSourceTracking.pdf \
	  PDF/figS8_taxBarplots.pdf \
	  PDF/figS9_gutGenera.pdf \
	  PDF/figS10_skinGenera.pdf
figpng = $(figures:.pdf=.png)
figeps = $(figures:.pdf=.eps)

pdf: $(figures)
	pdfunite $(figures) figures.pdf;
png: $(subst PDF,PNG,$(figpng))
eps: $(subst PDF,EPS,$(figeps))


#PDF/fig1_breedingStrategy.pdf: fig1_breedingStrategy.pdf
#	cp fig1_breedingStrategy.pdf PDF/
#
#PNG/fig1_breedingStrategy.png: fig1_breedingStrategy.pdf
#	convert $< $@

PDF/%.pdf: %.svg
	mkdir -p `dirname $@`
	inkscape -d $(PNGDPI) -C --export-type=pdf -o $@ $<
	
PDF/%.pdf: %.jpg
	convert $< $@;

PNG/%.png: %.jpg
	convert $< $@;

PNG/%.png: %.svg
	mkdir -p `dirname $@`
	inkscape -d $(PNGDPI) -C --export-type=png -o $@ $<

EPS/%.eps: %.svg
	mkdir -p `dirname $@`
	inkscape -d $(PNGDPI) -C --export-type=eps -o $@ $<
	
EPS/%.eps: %.jpg
	convert $< $@;
```
