## Additional File 2: Analysis repository for "The host genotype actively shapes its microbiome across generations in laboratory mice": ena-file-download-read_run-PRJEB28381-fastq_ftp-20231104-1013.sh

```
wget -nc ftp://ftp.sra.ebi.ac.uk/vol1/fastq/ERR278/009/ERR2784489/ERR2784489_1.fastq.gz
wget -nc ftp://ftp.sra.ebi.ac.uk/vol1/fastq/ERR278/009/ERR2784559/ERR2784559_1.fastq.gz
wget -nc ftp://ftp.sra.ebi.ac.uk/vol1/fastq/ERR278/009/ERR2784629/ERR2784629_1.fastq.gz
wget -nc ftp://ftp.sra.ebi.ac.uk/vol1/fastq/ERR278/009/ERR2784419/ERR2784419_1.fastq.gz
wget -nc ftp://ftp.sra.ebi.ac.uk/vol1/fastq/ERR278/009/ERR2784349/ERR2784349_1.fastq.gz
wget -nc ftp://ftp.sra.ebi.ac.uk/vol1/fastq/ERR278/007/ERR2784457/ERR2784457_1.fastq.gz
wget -nc ftp://ftp.sra.ebi.ac.uk/vol1/fastq/ERR278/007/ERR2784597/ERR2784597_1.fastq.gz
wget -nc ftp://ftp.sra.ebi.ac.uk/vol1/fastq/ERR278/007/ERR2784527/ERR2784527_1.fastq.gz
wget -nc ftp://ftp.sra.ebi.ac.uk/vol1/fastq/ERR278/007/ERR2784317/ERR2784317_1.fastq.gz
wget -nc ftp://ftp.sra.ebi.ac.uk/vol1/fastq/ERR278/007/ERR2784387/ERR2784387_1.fastq.gz
wget -nc ftp://ftp.sra.ebi.ac.uk/vol1/fastq/ERR278/008/ERR2784358/ERR2784358_2.fastq.gz
wget -nc ftp://ftp.sra.ebi.ac.uk/vol1/fastq/ERR278/008/ERR2784428/ERR2784428_2.fastq.gz
wget -nc ftp://ftp.sra.ebi.ac.uk/vol1/fastq/ERR278/008/ERR2784638/ERR2784638_2.fastq.gz
wget -nc ftp://ftp.sra.ebi.ac.uk/vol1/fastq/ERR278/008/ERR2784498/ERR2784498_2.fastq.gz
wget -nc ftp://ftp.sra.ebi.ac.uk/vol1/fastq/ERR278/008/ERR2784568/ERR2784568_2.fastq.gz
wget -nc ftp://ftp.sra.ebi.ac.uk/vol1/fastq/ERR278/004/ERR2784644/ERR2784644_2.fastq.gz
wget -nc ftp://ftp.sra.ebi.ac.uk/vol1/fastq/ERR278/003/ERR2784323/ERR2784323_1.fastq.gz
wget -nc ftp://ftp.sra.ebi.ac.uk/vol1/fastq/ERR278/001/ERR2784361/ERR2784361_1.fastq.gz
wget -nc ftp://ftp.sra.ebi.ac.uk/vol1/fastq/ERR278/002/ERR2784402/ERR2784402_2.fastq.gz
wget -nc ftp://ftp.sra.ebi.ac.uk/vol1/fastq/ERR278/001/ERR2784501/ERR2784501_1.fastq.gz
wget -nc ftp://ftp.sra.ebi.ac.uk/vol1/fastq/ERR278/002/ERR2784332/ERR2784332_2.fastq.gz
wget -nc ftp://ftp.sra.ebi.ac.uk/vol1/fastq/ERR278/001/ERR2784431/ERR2784431_1.fastq.gz
wget -nc ftp://ftp.sra.ebi.ac.uk/vol1/fastq/ERR278/001/ERR2784611/ERR2784611_2.fastq.gz
wget -nc ftp://ftp.sra.ebi.ac.uk/vol1/fastq/ERR278/001/ERR2784541/ERR2784541_2.fastq.gz
wget -nc ftp://ftp.sra.ebi.ac.uk/vol1/fastq/ERR278/001/ERR2784471/ERR2784471_2.fastq.gz
wget -nc ftp://ftp.sra.ebi.ac.uk/vol1/fastq/ERR278/006/ERR2784396/ERR2784396_2.fastq.gz
wget -nc ftp://ftp.sra.ebi.ac.uk/vol1/fastq/ERR278/006/ERR2784466/ERR2784466_2.fastq.gz
wget -nc ftp://ftp.sra.ebi.ac.uk/vol1/fastq/ERR278/006/ERR2784536/ERR2784536_2.fastq.gz
wget -nc ftp://ftp.sra.ebi.ac.uk/vol1/fastq/ERR278/006/ERR2784606/ERR2784606_2.fastq.gz
wget -nc ftp://ftp.sra.ebi.ac.uk/vol1/fastq/ERR278/000/ERR2784570/ERR2784570_1.fastq.gz
wget -nc ftp://ftp.sra.ebi.ac.uk/vol1/fastq/ERR278/000/ERR2784640/ERR2784640_1.fastq.gz
wget -nc ftp://ftp.sra.ebi.ac.uk/vol1/fastq/ERR278/004/ERR2784354/ERR2784354_1.fastq.gz
wget -nc ftp://ftp.sra.ebi.ac.uk/vol1/fastq/ERR278/004/ERR2784424/ERR2784424_1.fastq.gz
wget -nc ftp://ftp.sra.ebi.ac.uk/vol1/fastq/ERR278/005/ERR2784565/ERR2784565_1.fastq.gz
wget -nc ftp://ftp.sra.ebi.ac.uk/vol1/fastq/ERR278/005/ERR2784495/ERR2784495_1.fastq.gz
wget -nc ftp://ftp.sra.ebi.ac.uk/vol1/fastq/ERR278/005/ERR2784635/ERR2784635_1.fastq.gz
wget -nc ftp://ftp.sra.ebi.ac.uk/vol1/fastq/ERR278/003/ERR2784503/ERR2784503_2.fastq.gz
wget -nc ftp://ftp.sra.ebi.ac.uk/vol1/fastq/ERR278/002/ERR2784602/ERR2784602_1.fastq.gz
wget -nc ftp://ftp.sra.ebi.ac.uk/vol1/fastq/ERR278/002/ERR2784532/ERR2784532_1.fastq.gz
wget -nc ftp://ftp.sra.ebi.ac.uk/vol1/fastq/ERR278/003/ERR2784363/ERR2784363_2.fastq.gz
wget -nc ftp://ftp.sra.ebi.ac.uk/vol1/fastq/ERR278/003/ERR2784433/ERR2784433_2.fastq.gz
wget -nc ftp://ftp.sra.ebi.ac.uk/vol1/fastq/ERR278/002/ERR2784392/ERR2784392_1.fastq.gz
wget -nc ftp://ftp.sra.ebi.ac.uk/vol1/fastq/ERR278/002/ERR2784462/ERR2784462_1.fastq.gz
wget -nc ftp://ftp.sra.ebi.ac.uk/vol1/fastq/ERR278/004/ERR2784574/ERR2784574_2.fastq.gz
wget -nc ftp://ftp.sra.ebi.ac.uk/vol1/fastq/ERR278/009/ERR2784599/ERR2784599_2.fastq.gz
wget -nc ftp://ftp.sra.ebi.ac.uk/vol1/fastq/ERR278/008/ERR2784488/ERR2784488_1.fastq.gz
wget -nc ftp://ftp.sra.ebi.ac.uk/vol1/fastq/ERR278/008/ERR2784558/ERR2784558_1.fastq.gz
wget -nc ftp://ftp.sra.ebi.ac.uk/vol1/fastq/ERR278/008/ERR2784628/ERR2784628_1.fastq.gz
wget -nc ftp://ftp.sra.ebi.ac.uk/vol1/fastq/ERR278/008/ERR2784348/ERR2784348_1.fastq.gz
wget -nc ftp://ftp.sra.ebi.ac.uk/vol1/fastq/ERR278/008/ERR2784418/ERR2784418_1.fastq.gz
wget -nc ftp://ftp.sra.ebi.ac.uk/vol1/fastq/ERR278/005/ERR2784325/ERR2784325_2.fastq.gz
wget -nc ftp://ftp.sra.ebi.ac.uk/vol1/fastq/ERR278/009/ERR2784569/ERR2784569_2.fastq.gz
wget -nc ftp://ftp.sra.ebi.ac.uk/vol1/fastq/ERR278/009/ERR2784429/ERR2784429_2.fastq.gz
wget -nc ftp://ftp.sra.ebi.ac.uk/vol1/fastq/ERR278/009/ERR2784499/ERR2784499_2.fastq.gz
wget -nc ftp://ftp.sra.ebi.ac.uk/vol1/fastq/ERR278/009/ERR2784639/ERR2784639_2.fastq.gz
wget -nc ftp://ftp.sra.ebi.ac.uk/vol1/fastq/ERR278/009/ERR2784359/ERR2784359_2.fastq.gz
wget -nc ftp://ftp.sra.ebi.ac.uk/vol1/fastq/ERR278/000/ERR2784370/ERR2784370_2.fastq.gz
wget -nc ftp://ftp.sra.ebi.ac.uk/vol1/fastq/ERR278/008/ERR2784318/ERR2784318_1.fastq.gz
wget -nc ftp://ftp.sra.ebi.ac.uk/vol1/fastq/ERR278/006/ERR2784456/ERR2784456_1.fastq.gz
wget -nc ftp://ftp.sra.ebi.ac.uk/vol1/fastq/ERR278/006/ERR2784596/ERR2784596_1.fastq.gz
wget -nc ftp://ftp.sra.ebi.ac.uk/vol1/fastq/ERR278/006/ERR2784526/ERR2784526_1.fastq.gz
wget -nc ftp://ftp.sra.ebi.ac.uk/vol1/fastq/ERR278/006/ERR2784386/ERR2784386_1.fastq.gz
wget -nc ftp://ftp.sra.ebi.ac.uk/vol1/fastq/ERR278/000/ERR2784440/ERR2784440_2.fastq.gz
wget -nc ftp://ftp.sra.ebi.ac.uk/vol1/fastq/ERR278/000/ERR2784580/ERR2784580_2.fastq.gz
wget -nc ftp://ftp.sra.ebi.ac.uk/vol1/fastq/ERR278/000/ERR2784510/ERR2784510_2.fastq.gz
wget -nc ftp://ftp.sra.ebi.ac.uk/vol1/fastq/ERR278/000/ERR2784650/ERR2784650_2.fastq.gz
wget -nc ftp://ftp.sra.ebi.ac.uk/vol1/fastq/ERR278/004/ERR2784324/ERR2784324_1.fastq.gz
wget -nc ftp://ftp.sra.ebi.ac.uk/vol1/fastq/ERR278/007/ERR2784397/ERR2784397_2.fastq.gz
wget -nc ftp://ftp.sra.ebi.ac.uk/vol1/fastq/ERR278/002/ERR2784472/ERR2784472_2.fastq.gz
wget -nc ftp://ftp.sra.ebi.ac.uk/vol1/fastq/ERR278/001/ERR2784401/ERR2784401_2.fastq.gz
wget -nc ftp://ftp.sra.ebi.ac.uk/vol1/fastq/ERR278/001/ERR2784331/ERR2784331_2.fastq.gz
wget -nc ftp://ftp.sra.ebi.ac.uk/vol1/fastq/ERR278/007/ERR2784467/ERR2784467_2.fastq.gz
wget -nc ftp://ftp.sra.ebi.ac.uk/vol1/fastq/ERR278/007/ERR2784537/ERR2784537_2.fastq.gz
wget -nc ftp://ftp.sra.ebi.ac.uk/vol1/fastq/ERR278/007/ERR2784607/ERR2784607_2.fastq.gz
wget -nc ftp://ftp.sra.ebi.ac.uk/vol1/fastq/ERR278/006/ERR2784326/ERR2784326_2.fastq.gz
wget -nc ftp://ftp.sra.ebi.ac.uk/vol1/fastq/ERR278/002/ERR2784362/ERR2784362_1.fastq.gz
wget -nc ftp://ftp.sra.ebi.ac.uk/vol1/fastq/ERR278/002/ERR2784542/ERR2784542_2.fastq.gz
wget -nc ftp://ftp.sra.ebi.ac.uk/vol1/fastq/ERR278/002/ERR2784612/ERR2784612_2.fastq.gz
wget -nc ftp://ftp.sra.ebi.ac.uk/vol1/fastq/ERR278/004/ERR2784564/ERR2784564_1.fastq.gz
wget -nc ftp://ftp.sra.ebi.ac.uk/vol1/fastq/ERR278/004/ERR2784494/ERR2784494_1.fastq.gz
wget -nc ftp://ftp.sra.ebi.ac.uk/vol1/fastq/ERR278/004/ERR2784634/ERR2784634_1.fastq.gz
wget -nc ftp://ftp.sra.ebi.ac.uk/vol1/fastq/ERR278/005/ERR2784645/ERR2784645_2.fastq.gz
wget -nc ftp://ftp.sra.ebi.ac.uk/vol1/fastq/ERR278/005/ERR2784575/ERR2784575_2.fastq.gz
wget -nc ftp://ftp.sra.ebi.ac.uk/vol1/fastq/ERR278/001/ERR2784391/ERR2784391_1.fastq.gz
wget -nc ftp://ftp.sra.ebi.ac.uk/vol1/fastq/ERR278/001/ERR2784461/ERR2784461_1.fastq.gz
wget -nc ftp://ftp.sra.ebi.ac.uk/vol1/fastq/ERR278/004/ERR2784434/ERR2784434_2.fastq.gz
wget -nc ftp://ftp.sra.ebi.ac.uk/vol1/fastq/ERR278/001/ERR2784601/ERR2784601_1.fastq.gz
wget -nc ftp://ftp.sra.ebi.ac.uk/vol1/fastq/ERR278/001/ERR2784531/ERR2784531_1.fastq.gz
wget -nc ftp://ftp.sra.ebi.ac.uk/vol1/fastq/ERR278/004/ERR2784364/ERR2784364_2.fastq.gz
wget -nc ftp://ftp.sra.ebi.ac.uk/vol1/fastq/ERR278/004/ERR2784504/ERR2784504_2.fastq.gz
wget -nc ftp://ftp.sra.ebi.ac.uk/vol1/fastq/ERR278/005/ERR2784595/ERR2784595_1.fastq.gz
wget -nc ftp://ftp.sra.ebi.ac.uk/vol1/fastq/ERR278/005/ERR2784455/ERR2784455_1.fastq.gz
wget -nc ftp://ftp.sra.ebi.ac.uk/vol1/fastq/ERR278/008/ERR2784318/ERR2784318_2.fastq.gz
wget -nc ftp://ftp.sra.ebi.ac.uk/vol1/fastq/ERR278/008/ERR2784458/ERR2784458_2.fastq.gz
wget -nc ftp://ftp.sra.ebi.ac.uk/vol1/fastq/ERR278/007/ERR2784557/ERR2784557_1.fastq.gz
wget -nc ftp://ftp.sra.ebi.ac.uk/vol1/fastq/ERR278/007/ERR2784417/ERR2784417_1.fastq.gz
wget -nc ftp://ftp.sra.ebi.ac.uk/vol1/fastq/ERR278/003/ERR2784493/ERR2784493_1.fastq.gz
wget -nc ftp://ftp.sra.ebi.ac.uk/vol1/fastq/ERR278/004/ERR2784394/ERR2784394_2.fastq.gz
wget -nc ftp://ftp.sra.ebi.ac.uk/vol1/fastq/ERR278/003/ERR2784423/ERR2784423_1.fastq.gz
wget -nc ftp://ftp.sra.ebi.ac.uk/vol1/fastq/ERR278/004/ERR2784534/ERR2784534_2.fastq.gz
wget -nc ftp://ftp.sra.ebi.ac.uk/vol1/fastq/ERR278/003/ERR2784633/ERR2784633_1.fastq.gz
wget -nc ftp://ftp.sra.ebi.ac.uk/vol1/fastq/ERR278/009/ERR2784319/ERR2784319_1.fastq.gz
wget -nc ftp://ftp.sra.ebi.ac.uk/vol1/fastq/ERR278/009/ERR2784389/ERR2784389_1.fastq.gz
wget -nc ftp://ftp.sra.ebi.ac.uk/vol1/fastq/ERR278/009/ERR2784529/ERR2784529_1.fastq.gz
wget -nc ftp://ftp.sra.ebi.ac.uk/vol1/fastq/ERR278/002/ERR2784502/ERR2784502_1.fastq.gz
wget -nc ftp://ftp.sra.ebi.ac.uk/vol1/fastq/ERR278/002/ERR2784642/ERR2784642_1.fastq.gz
wget -nc ftp://ftp.sra.ebi.ac.uk/vol1/fastq/ERR278/001/ERR2784441/ERR2784441_2.fastq.gz
wget -nc ftp://ftp.sra.ebi.ac.uk/vol1/fastq/ERR278/001/ERR2784581/ERR2784581_2.fastq.gz
wget -nc ftp://ftp.sra.ebi.ac.uk/vol1/fastq/ERR278/000/ERR2784540/ERR2784540_1.fastq.gz
wget -nc ftp://ftp.sra.ebi.ac.uk/vol1/fastq/ERR278/006/ERR2784356/ERR2784356_2.fastq.gz
wget -nc ftp://ftp.sra.ebi.ac.uk/vol1/fastq/ERR278/000/ERR2784330/ERR2784330_1.fastq.gz
wget -nc ftp://ftp.sra.ebi.ac.uk/vol1/fastq/ERR278/006/ERR2784496/ERR2784496_2.fastq.gz
wget -nc ftp://ftp.sra.ebi.ac.uk/vol1/fastq/ERR278/006/ERR2784636/ERR2784636_2.fastq.gz
wget -nc ftp://ftp.sra.ebi.ac.uk/vol1/fastq/ERR278/000/ERR2784330/ERR2784330_2.fastq.gz
wget -nc ftp://ftp.sra.ebi.ac.uk/vol1/fastq/ERR278/004/ERR2784464/ERR2784464_1.fastq.gz
wget -nc ftp://ftp.sra.ebi.ac.uk/vol1/fastq/ERR278/007/ERR2784567/ERR2784567_1.fastq.gz
wget -nc ftp://ftp.sra.ebi.ac.uk/vol1/fastq/ERR278/008/ERR2784468/ERR2784468_2.fastq.gz
wget -nc ftp://ftp.sra.ebi.ac.uk/vol1/fastq/ERR278/004/ERR2784604/ERR2784604_1.fastq.gz
wget -nc ftp://ftp.sra.ebi.ac.uk/vol1/fastq/ERR278/008/ERR2784608/ERR2784608_2.fastq.gz
wget -nc ftp://ftp.sra.ebi.ac.uk/vol1/fastq/ERR278/007/ERR2784327/ERR2784327_2.fastq.gz
wget -nc ftp://ftp.sra.ebi.ac.uk/vol1/fastq/ERR278/006/ERR2784426/ERR2784426_1.fastq.gz
wget -nc ftp://ftp.sra.ebi.ac.uk/vol1/fastq/ERR278/003/ERR2784543/ERR2784543_2.fastq.gz
wget -nc ftp://ftp.sra.ebi.ac.uk/vol1/fastq/ERR278/000/ERR2784390/ERR2784390_1.fastq.gz
wget -nc ftp://ftp.sra.ebi.ac.uk/vol1/fastq/ERR278/009/ERR2784379/ERR2784379_1.fastq.gz
wget -nc ftp://ftp.sra.ebi.ac.uk/vol1/fastq/ERR278/001/ERR2784431/ERR2784431_2.fastq.gz
wget -nc ftp://ftp.sra.ebi.ac.uk/vol1/fastq/ERR278/009/ERR2784519/ERR2784519_1.fastq.gz
wget -nc ftp://ftp.sra.ebi.ac.uk/vol1/fastq/ERR278/006/ERR2784646/ERR2784646_2.fastq.gz
wget -nc ftp://ftp.sra.ebi.ac.uk/vol1/fastq/ERR278/002/ERR2784572/ERR2784572_2.fastq.gz
wget -nc ftp://ftp.sra.ebi.ac.uk/vol1/fastq/ERR278/000/ERR2784530/ERR2784530_1.fastq.gz
wget -nc ftp://ftp.sra.ebi.ac.uk/vol1/fastq/ERR278/005/ERR2784365/ERR2784365_2.fastq.gz
wget -nc ftp://ftp.sra.ebi.ac.uk/vol1/fastq/ERR278/005/ERR2784505/ERR2784505_2.fastq.gz
wget -nc ftp://ftp.sra.ebi.ac.uk/vol1/fastq/ERR278/001/ERR2784321/ERR2784321_1.fastq.gz
wget -nc ftp://ftp.sra.ebi.ac.uk/vol1/fastq/ERR278/002/ERR2784492/ERR2784492_1.fastq.gz
wget -nc ftp://ftp.sra.ebi.ac.uk/vol1/fastq/ERR278/009/ERR2784319/ERR2784319_2.fastq.gz
wget -nc ftp://ftp.sra.ebi.ac.uk/vol1/fastq/ERR278/009/ERR2784459/ERR2784459_2.fastq.gz
wget -nc ftp://ftp.sra.ebi.ac.uk/vol1/fastq/ERR278/006/ERR2784556/ERR2784556_1.fastq.gz
wget -nc ftp://ftp.sra.ebi.ac.uk/vol1/fastq/ERR278/006/ERR2784416/ERR2784416_1.fastq.gz
wget -nc ftp://ftp.sra.ebi.ac.uk/vol1/fastq/ERR278/000/ERR2784480/ERR2784480_2.fastq.gz
wget -nc ftp://ftp.sra.ebi.ac.uk/vol1/fastq/ERR278/002/ERR2784632/ERR2784632_1.fastq.gz
wget -nc ftp://ftp.sra.ebi.ac.uk/vol1/fastq/ERR278/000/ERR2784620/ERR2784620_2.fastq.gz
wget -nc ftp://ftp.sra.ebi.ac.uk/vol1/fastq/ERR278/005/ERR2784395/ERR2784395_2.fastq.gz
wget -nc ftp://ftp.sra.ebi.ac.uk/vol1/fastq/ERR278/004/ERR2784404/ERR2784404_2.fastq.gz
wget -nc ftp://ftp.sra.ebi.ac.uk/vol1/fastq/ERR278/004/ERR2784544/ERR2784544_2.fastq.gz
wget -nc ftp://ftp.sra.ebi.ac.uk/vol1/fastq/ERR278/008/ERR2784528/ERR2784528_1.fastq.gz
wget -nc ftp://ftp.sra.ebi.ac.uk/vol1/fastq/ERR278/008/ERR2784388/ERR2784388_1.fastq.gz
wget -nc ftp://ftp.sra.ebi.ac.uk/vol1/fastq/ERR278/007/ERR2784497/ERR2784497_2.fastq.gz
wget -nc ftp://ftp.sra.ebi.ac.uk/vol1/fastq/ERR278/007/ERR2784637/ERR2784637_2.fastq.gz
wget -nc ftp://ftp.sra.ebi.ac.uk/vol1/fastq/ERR278/001/ERR2784331/ERR2784331_1.fastq.gz
wget -nc ftp://ftp.sra.ebi.ac.uk/vol1/fastq/ERR278/002/ERR2784442/ERR2784442_2.fastq.gz
wget -nc ftp://ftp.sra.ebi.ac.uk/vol1/fastq/ERR278/002/ERR2784582/ERR2784582_2.fastq.gz
wget -nc ftp://ftp.sra.ebi.ac.uk/vol1/fastq/ERR278/007/ERR2784357/ERR2784357_2.fastq.gz
wget -nc ftp://ftp.sra.ebi.ac.uk/vol1/fastq/ERR278/004/ERR2784454/ERR2784454_1.fastq.gz
wget -nc ftp://ftp.sra.ebi.ac.uk/vol1/fastq/ERR278/004/ERR2784594/ERR2784594_1.fastq.gz
wget -nc ftp://ftp.sra.ebi.ac.uk/vol1/fastq/ERR278/000/ERR2784470/ERR2784470_2.fastq.gz
wget -nc ftp://ftp.sra.ebi.ac.uk/vol1/fastq/ERR278/003/ERR2784463/ERR2784463_1.fastq.gz
wget -nc ftp://ftp.sra.ebi.ac.uk/vol1/fastq/ERR278/000/ERR2784610/ERR2784610_2.fastq.gz
wget -nc ftp://ftp.sra.ebi.ac.uk/vol1/fastq/ERR278/003/ERR2784603/ERR2784603_1.fastq.gz
wget -nc ftp://ftp.sra.ebi.ac.uk/vol1/fastq/ERR278/008/ERR2784328/ERR2784328_2.fastq.gz
wget -nc ftp://ftp.sra.ebi.ac.uk/vol1/fastq/ERR278/009/ERR2784609/ERR2784609_2.fastq.gz
wget -nc ftp://ftp.sra.ebi.ac.uk/vol1/fastq/ERR278/009/ERR2784469/ERR2784469_2.fastq.gz
wget -nc ftp://ftp.sra.ebi.ac.uk/vol1/fastq/ERR278/006/ERR2784566/ERR2784566_1.fastq.gz
wget -nc ftp://ftp.sra.ebi.ac.uk/vol1/fastq/ERR278/001/ERR2784641/ERR2784641_1.fastq.gz
wget -nc ftp://ftp.sra.ebi.ac.uk/vol1/fastq/ERR278/000/ERR2784360/ERR2784360_1.fastq.gz
wget -nc ftp://ftp.sra.ebi.ac.uk/vol1/fastq/ERR278/003/ERR2784403/ERR2784403_2.fastq.gz
wget -nc ftp://ftp.sra.ebi.ac.uk/vol1/fastq/ERR278/000/ERR2784500/ERR2784500_1.fastq.gz
wget -nc ftp://ftp.sra.ebi.ac.uk/vol1/fastq/ERR278/005/ERR2784535/ERR2784535_2.fastq.gz
wget -nc ftp://ftp.sra.ebi.ac.uk/vol1/fastq/ERR278/005/ERR2784425/ERR2784425_1.fastq.gz
wget -nc ftp://ftp.sra.ebi.ac.uk/vol1/fastq/ERR278/007/ERR2784647/ERR2784647_2.fastq.gz
wget -nc ftp://ftp.sra.ebi.ac.uk/vol1/fastq/ERR278/008/ERR2784518/ERR2784518_1.fastq.gz
wget -nc ftp://ftp.sra.ebi.ac.uk/vol1/fastq/ERR278/006/ERR2784366/ERR2784366_2.fastq.gz
wget -nc ftp://ftp.sra.ebi.ac.uk/vol1/fastq/ERR278/008/ERR2784378/ERR2784378_1.fastq.gz
wget -nc ftp://ftp.sra.ebi.ac.uk/vol1/fastq/ERR278/002/ERR2784432/ERR2784432_2.fastq.gz
wget -nc ftp://ftp.sra.ebi.ac.uk/vol1/fastq/ERR278/006/ERR2784506/ERR2784506_2.fastq.gz
wget -nc ftp://ftp.sra.ebi.ac.uk/vol1/fastq/ERR278/003/ERR2784573/ERR2784573_2.fastq.gz
wget -nc ftp://ftp.sra.ebi.ac.uk/vol1/fastq/ERR278/002/ERR2784322/ERR2784322_1.fastq.gz
wget -nc ftp://ftp.sra.ebi.ac.uk/vol1/fastq/ERR278/003/ERR2784513/ERR2784513_2.fastq.gz
wget -nc ftp://ftp.sra.ebi.ac.uk/vol1/fastq/ERR278/003/ERR2784653/ERR2784653_2.fastq.gz
wget -nc ftp://ftp.sra.ebi.ac.uk/vol1/fastq/ERR278/002/ERR2784402/ERR2784402_1.fastq.gz
wget -nc ftp://ftp.sra.ebi.ac.uk/vol1/fastq/ERR278/002/ERR2784542/ERR2784542_1.fastq.gz
wget -nc ftp://ftp.sra.ebi.ac.uk/vol1/fastq/ERR278/002/ERR2784472/ERR2784472_1.fastq.gz
wget -nc ftp://ftp.sra.ebi.ac.uk/vol1/fastq/ERR278/002/ERR2784322/ERR2784322_2.fastq.gz
wget -nc ftp://ftp.sra.ebi.ac.uk/vol1/fastq/ERR278/002/ERR2784392/ERR2784392_2.fastq.gz
wget -nc ftp://ftp.sra.ebi.ac.uk/vol1/fastq/ERR278/002/ERR2784462/ERR2784462_2.fastq.gz
wget -nc ftp://ftp.sra.ebi.ac.uk/vol1/fastq/ERR278/000/ERR2784580/ERR2784580_1.fastq.gz
wget -nc ftp://ftp.sra.ebi.ac.uk/vol1/fastq/ERR278/005/ERR2784615/ERR2784615_2.fastq.gz
wget -nc ftp://ftp.sra.ebi.ac.uk/vol1/fastq/ERR278/000/ERR2784440/ERR2784440_1.fastq.gz
wget -nc ftp://ftp.sra.ebi.ac.uk/vol1/fastq/ERR278/000/ERR2784510/ERR2784510_1.fastq.gz
wget -nc ftp://ftp.sra.ebi.ac.uk/vol1/fastq/ERR278/001/ERR2784351/ERR2784351_1.fastq.gz
wget -nc ftp://ftp.sra.ebi.ac.uk/vol1/fastq/ERR278/001/ERR2784621/ERR2784621_2.fastq.gz
wget -nc ftp://ftp.sra.ebi.ac.uk/vol1/fastq/ERR278/001/ERR2784631/ERR2784631_1.fastq.gz
wget -nc ftp://ftp.sra.ebi.ac.uk/vol1/fastq/ERR278/001/ERR2784411/ERR2784411_2.fastq.gz
wget -nc ftp://ftp.sra.ebi.ac.uk/vol1/fastq/ERR278/001/ERR2784551/ERR2784551_2.fastq.gz
wget -nc ftp://ftp.sra.ebi.ac.uk/vol1/fastq/ERR278/005/ERR2784485/ERR2784485_1.fastq.gz
wget -nc ftp://ftp.sra.ebi.ac.uk/vol1/fastq/ERR278/004/ERR2784434/ERR2784434_1.fastq.gz
wget -nc ftp://ftp.sra.ebi.ac.uk/vol1/fastq/ERR278/004/ERR2784644/ERR2784644_1.fastq.gz
wget -nc ftp://ftp.sra.ebi.ac.uk/vol1/fastq/ERR278/004/ERR2784574/ERR2784574_1.fastq.gz
wget -nc ftp://ftp.sra.ebi.ac.uk/vol1/fastq/ERR278/004/ERR2784504/ERR2784504_1.fastq.gz
wget -nc ftp://ftp.sra.ebi.ac.uk/vol1/fastq/ERR278/003/ERR2784383/ERR2784383_1.fastq.gz
wget -nc ftp://ftp.sra.ebi.ac.uk/vol1/fastq/ERR278/003/ERR2784313/ERR2784313_1.fastq.gz
wget -nc ftp://ftp.sra.ebi.ac.uk/vol1/fastq/ERR278/004/ERR2784354/ERR2784354_2.fastq.gz
wget -nc ftp://ftp.sra.ebi.ac.uk/vol1/fastq/ERR278/004/ERR2784424/ERR2784424_2.fastq.gz
wget -nc ftp://ftp.sra.ebi.ac.uk/vol1/fastq/ERR278/004/ERR2784564/ERR2784564_2.fastq.gz
wget -nc ftp://ftp.sra.ebi.ac.uk/vol1/fastq/ERR278/004/ERR2784494/ERR2784494_2.fastq.gz
wget -nc ftp://ftp.sra.ebi.ac.uk/vol1/fastq/ERR278/009/ERR2784329/ERR2784329_2.fastq.gz
wget -nc ftp://ftp.sra.ebi.ac.uk/vol1/fastq/ERR278/009/ERR2784339/ERR2784339_1.fastq.gz
wget -nc ftp://ftp.sra.ebi.ac.uk/vol1/fastq/ERR278/009/ERR2784409/ERR2784409_1.fastq.gz
wget -nc ftp://ftp.sra.ebi.ac.uk/vol1/fastq/ERR278/008/ERR2784578/ERR2784578_2.fastq.gz
wget -nc ftp://ftp.sra.ebi.ac.uk/vol1/fastq/ERR278/008/ERR2784648/ERR2784648_2.fastq.gz
wget -nc ftp://ftp.sra.ebi.ac.uk/vol1/fastq/ERR278/006/ERR2784456/ERR2784456_2.fastq.gz
wget -nc ftp://ftp.sra.ebi.ac.uk/vol1/fastq/ERR278/006/ERR2784526/ERR2784526_2.fastq.gz
wget -nc ftp://ftp.sra.ebi.ac.uk/vol1/fastq/ERR278/007/ERR2784327/ERR2784327_1.fastq.gz
wget -nc ftp://ftp.sra.ebi.ac.uk/vol1/fastq/ERR278/006/ERR2784536/ERR2784536_1.fastq.gz
wget -nc ftp://ftp.sra.ebi.ac.uk/vol1/fastq/ERR278/007/ERR2784367/ERR2784367_2.fastq.gz
wget -nc ftp://ftp.sra.ebi.ac.uk/vol1/fastq/ERR278/006/ERR2784606/ERR2784606_1.fastq.gz
wget -nc ftp://ftp.sra.ebi.ac.uk/vol1/fastq/ERR278/009/ERR2784569/ERR2784569_1.fastq.gz
wget -nc ftp://ftp.sra.ebi.ac.uk/vol1/fastq/ERR278/009/ERR2784499/ERR2784499_1.fastq.gz
wget -nc ftp://ftp.sra.ebi.ac.uk/vol1/fastq/ERR278/007/ERR2784377/ERR2784377_1.fastq.gz
wget -nc ftp://ftp.sra.ebi.ac.uk/vol1/fastq/ERR278/007/ERR2784447/ERR2784447_1.fastq.gz
wget -nc ftp://ftp.sra.ebi.ac.uk/vol1/fastq/ERR278/008/ERR2784418/ERR2784418_2.fastq.gz
wget -nc ftp://ftp.sra.ebi.ac.uk/vol1/fastq/ERR278/009/ERR2784489/ERR2784489_2.fastq.gz
wget -nc ftp://ftp.sra.ebi.ac.uk/vol1/fastq/ERR278/000/ERR2784490/ERR2784490_1.fastq.gz
wget -nc ftp://ftp.sra.ebi.ac.uk/vol1/fastq/ERR278/000/ERR2784560/ERR2784560_1.fastq.gz
wget -nc ftp://ftp.sra.ebi.ac.uk/vol1/fastq/ERR278/005/ERR2784635/ERR2784635_2.fastq.gz
wget -nc ftp://ftp.sra.ebi.ac.uk/vol1/fastq/ERR278/004/ERR2784374/ERR2784374_2.fastq.gz
wget -nc ftp://ftp.sra.ebi.ac.uk/vol1/fastq/ERR278/001/ERR2784611/ERR2784611_1.fastq.gz
wget -nc ftp://ftp.sra.ebi.ac.uk/vol1/fastq/ERR278/004/ERR2784444/ERR2784444_2.fastq.gz
wget -nc ftp://ftp.sra.ebi.ac.uk/vol1/fastq/ERR278/004/ERR2784584/ERR2784584_2.fastq.gz
wget -nc ftp://ftp.sra.ebi.ac.uk/vol1/fastq/ERR278/003/ERR2784333/ERR2784333_1.fastq.gz
wget -nc ftp://ftp.sra.ebi.ac.uk/vol1/fastq/ERR278/006/ERR2784546/ERR2784546_2.fastq.gz
wget -nc ftp://ftp.sra.ebi.ac.uk/vol1/fastq/ERR278/002/ERR2784422/ERR2784422_1.fastq.gz
wget -nc ftp://ftp.sra.ebi.ac.uk/vol1/fastq/ERR278/006/ERR2784476/ERR2784476_2.fastq.gz
wget -nc ftp://ftp.sra.ebi.ac.uk/vol1/fastq/ERR278/006/ERR2784336/ERR2784336_2.fastq.gz
wget -nc ftp://ftp.sra.ebi.ac.uk/vol1/fastq/ERR278/006/ERR2784406/ERR2784406_2.fastq.gz
wget -nc ftp://ftp.sra.ebi.ac.uk/vol1/fastq/ERR278/003/ERR2784603/ERR2784603_2.fastq.gz
wget -nc ftp://ftp.sra.ebi.ac.uk/vol1/fastq/ERR278/003/ERR2784533/ERR2784533_2.fastq.gz
wget -nc ftp://ftp.sra.ebi.ac.uk/vol1/fastq/ERR278/007/ERR2784387/ERR2784387_2.fastq.gz
wget -nc ftp://ftp.sra.ebi.ac.uk/vol1/fastq/ERR278/000/ERR2784340/ERR2784340_2.fastq.gz
wget -nc ftp://ftp.sra.ebi.ac.uk/vol1/fastq/ERR278/003/ERR2784363/ERR2784363_1.fastq.gz
wget -nc ftp://ftp.sra.ebi.ac.uk/vol1/fastq/ERR278/002/ERR2784452/ERR2784452_1.fastq.gz
wget -nc ftp://ftp.sra.ebi.ac.uk/vol1/fastq/ERR278/002/ERR2784522/ERR2784522_1.fastq.gz
wget -nc ftp://ftp.sra.ebi.ac.uk/vol1/fastq/ERR278/002/ERR2784592/ERR2784592_1.fastq.gz
wget -nc ftp://ftp.sra.ebi.ac.uk/vol1/fastq/ERR278/008/ERR2784638/ERR2784638_1.fastq.gz
wget -nc ftp://ftp.sra.ebi.ac.uk/vol1/fastq/ERR278/007/ERR2784357/ERR2784357_1.fastq.gz
wget -nc ftp://ftp.sra.ebi.ac.uk/vol1/fastq/ERR278/007/ERR2784427/ERR2784427_1.fastq.gz
wget -nc ftp://ftp.sra.ebi.ac.uk/vol1/fastq/ERR278/005/ERR2784465/ERR2784465_1.fastq.gz
wget -nc ftp://ftp.sra.ebi.ac.uk/vol1/fastq/ERR278/005/ERR2784395/ERR2784395_1.fastq.gz
wget -nc ftp://ftp.sra.ebi.ac.uk/vol1/fastq/ERR278/009/ERR2784549/ERR2784549_1.fastq.gz
wget -nc ftp://ftp.sra.ebi.ac.uk/vol1/fastq/ERR278/009/ERR2784479/ERR2784479_1.fastq.gz
wget -nc ftp://ftp.sra.ebi.ac.uk/vol1/fastq/ERR278/004/ERR2784554/ERR2784554_1.fastq.gz
wget -nc ftp://ftp.sra.ebi.ac.uk/vol1/fastq/ERR278/004/ERR2784624/ERR2784624_1.fastq.gz
wget -nc ftp://ftp.sra.ebi.ac.uk/vol1/fastq/ERR278/006/ERR2784316/ERR2784316_2.fastq.gz
wget -nc ftp://ftp.sra.ebi.ac.uk/vol1/fastq/ERR278/000/ERR2784320/ERR2784320_1.fastq.gz
wget -nc ftp://ftp.sra.ebi.ac.uk/vol1/fastq/ERR278/002/ERR2784482/ERR2784482_2.fastq.gz
wget -nc ftp://ftp.sra.ebi.ac.uk/vol1/fastq/ERR278/001/ERR2784641/ERR2784641_2.fastq.gz
wget -nc ftp://ftp.sra.ebi.ac.uk/vol1/fastq/ERR278/001/ERR2784571/ERR2784571_2.fastq.gz
wget -nc ftp://ftp.sra.ebi.ac.uk/vol1/fastq/ERR278/000/ERR2784360/ERR2784360_2.fastq.gz
wget -nc ftp://ftp.sra.ebi.ac.uk/vol1/fastq/ERR278/008/ERR2784438/ERR2784438_2.fastq.gz
wget -nc ftp://ftp.sra.ebi.ac.uk/vol1/fastq/ERR278/008/ERR2784508/ERR2784508_2.fastq.gz
wget -nc ftp://ftp.sra.ebi.ac.uk/vol1/fastq/ERR278/007/ERR2784587/ERR2784587_1.fastq.gz
wget -nc ftp://ftp.sra.ebi.ac.uk/vol1/fastq/ERR278/006/ERR2784516/ERR2784516_1.fastq.gz
wget -nc ftp://ftp.sra.ebi.ac.uk/vol1/fastq/ERR278/009/ERR2784349/ERR2784349_2.fastq.gz
wget -nc ftp://ftp.sra.ebi.ac.uk/vol1/fastq/ERR278/005/ERR2784385/ERR2784385_1.fastq.gz
wget -nc ftp://ftp.sra.ebi.ac.uk/vol1/fastq/ERR278/005/ERR2784315/ERR2784315_1.fastq.gz
wget -nc ftp://ftp.sra.ebi.ac.uk/vol1/fastq/ERR278/006/ERR2784506/ERR2784506_1.fastq.gz
wget -nc ftp://ftp.sra.ebi.ac.uk/vol1/fastq/ERR278/006/ERR2784646/ERR2784646_1.fastq.gz
wget -nc ftp://ftp.sra.ebi.ac.uk/vol1/fastq/ERR278/000/ERR2784390/ERR2784390_2.fastq.gz
wget -nc ftp://ftp.sra.ebi.ac.uk/vol1/fastq/ERR278/007/ERR2784487/ERR2784487_1.fastq.gz
wget -nc ftp://ftp.sra.ebi.ac.uk/vol1/fastq/ERR278/004/ERR2784544/ERR2784544_1.fastq.gz
wget -nc ftp://ftp.sra.ebi.ac.uk/vol1/fastq/ERR278/008/ERR2784598/ERR2784598_2.fastq.gz
wget -nc ftp://ftp.sra.ebi.ac.uk/vol1/fastq/ERR278/008/ERR2784528/ERR2784528_2.fastq.gz
wget -nc ftp://ftp.sra.ebi.ac.uk/vol1/fastq/ERR278/007/ERR2784347/ERR2784347_1.fastq.gz
wget -nc ftp://ftp.sra.ebi.ac.uk/vol1/fastq/ERR278/004/ERR2784324/ERR2784324_2.fastq.gz
wget -nc ftp://ftp.sra.ebi.ac.uk/vol1/fastq/ERR278/003/ERR2784353/ERR2784353_1.fastq.gz
wget -nc ftp://ftp.sra.ebi.ac.uk/vol1/fastq/ERR278/008/ERR2784608/ERR2784608_1.fastq.gz
wget -nc ftp://ftp.sra.ebi.ac.uk/vol1/fastq/ERR278/004/ERR2784464/ERR2784464_2.fastq.gz
wget -nc ftp://ftp.sra.ebi.ac.uk/vol1/fastq/ERR278/009/ERR2784599/ERR2784599_1.fastq.gz
wget -nc ftp://ftp.sra.ebi.ac.uk/vol1/fastq/ERR278/009/ERR2784459/ERR2784459_1.fastq.gz
wget -nc ftp://ftp.sra.ebi.ac.uk/vol1/fastq/ERR278/002/ERR2784572/ERR2784572_1.fastq.gz
wget -nc ftp://ftp.sra.ebi.ac.uk/vol1/fastq/ERR278/002/ERR2784432/ERR2784432_1.fastq.gz
wget -nc ftp://ftp.sra.ebi.ac.uk/vol1/fastq/ERR278/002/ERR2784352/ERR2784352_2.fastq.gz
wget -nc ftp://ftp.sra.ebi.ac.uk/vol1/fastq/ERR278/002/ERR2784492/ERR2784492_2.fastq.gz
wget -nc ftp://ftp.sra.ebi.ac.uk/vol1/fastq/ERR278/001/ERR2784511/ERR2784511_2.fastq.gz
wget -nc ftp://ftp.sra.ebi.ac.uk/vol1/fastq/ERR278/001/ERR2784651/ERR2784651_2.fastq.gz
wget -nc ftp://ftp.sra.ebi.ac.uk/vol1/fastq/ERR278/000/ERR2784400/ERR2784400_1.fastq.gz
wget -nc ftp://ftp.sra.ebi.ac.uk/vol1/fastq/ERR278/000/ERR2784470/ERR2784470_1.fastq.gz
wget -nc ftp://ftp.sra.ebi.ac.uk/vol1/fastq/ERR278/006/ERR2784426/ERR2784426_2.fastq.gz
wget -nc ftp://ftp.sra.ebi.ac.uk/vol1/fastq/ERR278/006/ERR2784566/ERR2784566_2.fastq.gz
wget -nc ftp://ftp.sra.ebi.ac.uk/vol1/fastq/ERR278/006/ERR2784416/ERR2784416_2.fastq.gz
wget -nc ftp://ftp.sra.ebi.ac.uk/vol1/fastq/ERR278/007/ERR2784497/ERR2784497_1.fastq.gz
wget -nc ftp://ftp.sra.ebi.ac.uk/vol1/fastq/ERR278/000/ERR2784410/ERR2784410_1.fastq.gz
wget -nc ftp://ftp.sra.ebi.ac.uk/vol1/fastq/ERR278/004/ERR2784534/ERR2784534_1.fastq.gz
wget -nc ftp://ftp.sra.ebi.ac.uk/vol1/fastq/ERR278/008/ERR2784538/ERR2784538_2.fastq.gz
wget -nc ftp://ftp.sra.ebi.ac.uk/vol1/fastq/ERR278/005/ERR2784325/ERR2784325_1.fastq.gz
wget -nc ftp://ftp.sra.ebi.ac.uk/vol1/fastq/ERR278/004/ERR2784454/ERR2784454_2.fastq.gz
wget -nc ftp://ftp.sra.ebi.ac.uk/vol1/fastq/ERR278/008/ERR2784618/ERR2784618_1.fastq.gz
wget -nc ftp://ftp.sra.ebi.ac.uk/vol1/fastq/ERR278/009/ERR2784379/ERR2784379_2.fastq.gz
wget -nc ftp://ftp.sra.ebi.ac.uk/vol1/fastq/ERR278/008/ERR2784588/ERR2784588_2.fastq.gz
wget -nc ftp://ftp.sra.ebi.ac.uk/vol1/fastq/ERR278/003/ERR2784613/ERR2784613_2.fastq.gz
wget -nc ftp://ftp.sra.ebi.ac.uk/vol1/fastq/ERR278/007/ERR2784337/ERR2784337_1.fastq.gz
wget -nc ftp://ftp.sra.ebi.ac.uk/vol1/fastq/ERR278/000/ERR2784460/ERR2784460_1.fastq.gz
wget -nc ftp://ftp.sra.ebi.ac.uk/vol1/fastq/ERR278/002/ERR2784342/ERR2784342_2.fastq.gz
wget -nc ftp://ftp.sra.ebi.ac.uk/vol1/fastq/ERR278/009/ERR2784449/ERR2784449_1.fastq.gz
wget -nc ftp://ftp.sra.ebi.ac.uk/vol1/fastq/ERR278/001/ERR2784501/ERR2784501_2.fastq.gz
wget -nc ftp://ftp.sra.ebi.ac.uk/vol1/fastq/ERR278/003/ERR2784623/ERR2784623_2.fastq.gz
wget -nc ftp://ftp.sra.ebi.ac.uk/vol1/fastq/ERR278/009/ERR2784369/ERR2784369_2.fastq.gz
wget -nc ftp://ftp.sra.ebi.ac.uk/vol1/fastq/ERR278/002/ERR2784582/ERR2784582_1.fastq.gz
wget -nc ftp://ftp.sra.ebi.ac.uk/vol1/fastq/ERR278/000/ERR2784380/ERR2784380_2.fastq.gz
wget -nc ftp://ftp.sra.ebi.ac.uk/vol1/fastq/ERR278/006/ERR2784576/ERR2784576_2.fastq.gz
wget -nc ftp://ftp.sra.ebi.ac.uk/vol1/fastq/ERR278/005/ERR2784375/ERR2784375_1.fastq.gz
wget -nc ftp://ftp.sra.ebi.ac.uk/vol1/fastq/ERR278/006/ERR2784656/ERR2784656_1.fastq.gz
wget -nc ftp://ftp.sra.ebi.ac.uk/vol1/fastq/ERR278/000/ERR2784420/ERR2784420_1.fastq.gz
wget -nc ftp://ftp.sra.ebi.ac.uk/vol1/fastq/ERR278/008/ERR2784408/ERR2784408_2.fastq.gz
wget -nc ftp://ftp.sra.ebi.ac.uk/vol1/fastq/ERR278/008/ERR2784548/ERR2784548_2.fastq.gz
wget -nc ftp://ftp.sra.ebi.ac.uk/vol1/fastq/ERR278/001/ERR2784531/ERR2784531_2.fastq.gz
wget -nc ftp://ftp.sra.ebi.ac.uk/vol1/fastq/ERR278/002/ERR2784562/ERR2784562_1.fastq.gz
wget -nc ftp://ftp.sra.ebi.ac.uk/vol1/fastq/ERR278/005/ERR2784335/ERR2784335_1.fastq.gz
wget -nc ftp://ftp.sra.ebi.ac.uk/vol1/fastq/ERR278/009/ERR2784389/ERR2784389_2.fastq.gz
wget -nc ftp://ftp.sra.ebi.ac.uk/vol1/fastq/ERR278/006/ERR2784626/ERR2784626_1.fastq.gz
wget -nc ftp://ftp.sra.ebi.ac.uk/vol1/fastq/ERR278/004/ERR2784334/ERR2784334_2.fastq.gz
wget -nc ftp://ftp.sra.ebi.ac.uk/vol1/fastq/ERR278/004/ERR2784474/ERR2784474_2.fastq.gz
wget -nc ftp://ftp.sra.ebi.ac.uk/vol1/fastq/ERR278/006/ERR2784446/ERR2784446_2.fastq.gz
wget -nc ftp://ftp.sra.ebi.ac.uk/vol1/fastq/ERR278/006/ERR2784586/ERR2784586_2.fastq.gz
wget -nc ftp://ftp.sra.ebi.ac.uk/vol1/fastq/ERR278/000/ERR2784590/ERR2784590_1.fastq.gz
wget -nc ftp://ftp.sra.ebi.ac.uk/vol1/fastq/ERR278/000/ERR2784450/ERR2784450_1.fastq.gz
wget -nc ftp://ftp.sra.ebi.ac.uk/vol1/fastq/ERR278/005/ERR2784365/ERR2784365_1.fastq.gz
wget -nc ftp://ftp.sra.ebi.ac.uk/vol1/fastq/ERR278/009/ERR2784439/ERR2784439_1.fastq.gz
wget -nc ftp://ftp.sra.ebi.ac.uk/vol1/fastq/ERR278/003/ERR2784633/ERR2784633_2.fastq.gz
wget -nc ftp://ftp.sra.ebi.ac.uk/vol1/fastq/ERR278/009/ERR2784579/ERR2784579_1.fastq.gz
wget -nc ftp://ftp.sra.ebi.ac.uk/vol1/fastq/ERR278/004/ERR2784524/ERR2784524_1.fastq.gz
wget -nc ftp://ftp.sra.ebi.ac.uk/vol1/fastq/ERR278/002/ERR2784372/ERR2784372_2.fastq.gz
wget -nc ftp://ftp.sra.ebi.ac.uk/vol1/fastq/ERR278/003/ERR2784393/ERR2784393_1.fastq.gz
wget -nc ftp://ftp.sra.ebi.ac.uk/vol1/fastq/ERR278/001/ERR2784521/ERR2784521_2.fastq.gz
wget -nc ftp://ftp.sra.ebi.ac.uk/vol1/fastq/ERR278/002/ERR2784552/ERR2784552_1.fastq.gz
wget -nc ftp://ftp.sra.ebi.ac.uk/vol1/fastq/ERR278/009/ERR2784399/ERR2784399_2.fastq.gz
wget -nc ftp://ftp.sra.ebi.ac.uk/vol1/fastq/ERR278/006/ERR2784636/ERR2784636_1.fastq.gz
wget -nc ftp://ftp.sra.ebi.ac.uk/vol1/fastq/ERR278/004/ERR2784314/ERR2784314_2.fastq.gz
wget -nc ftp://ftp.sra.ebi.ac.uk/vol1/fastq/ERR278/005/ERR2784605/ERR2784605_2.fastq.gz
wget -nc ftp://ftp.sra.ebi.ac.uk/vol1/fastq/ERR278/000/ERR2784430/ERR2784430_1.fastq.gz
wget -nc ftp://ftp.sra.ebi.ac.uk/vol1/fastq/ERR278/005/ERR2784345/ERR2784345_1.fastq.gz
wget -nc ftp://ftp.sra.ebi.ac.uk/vol1/fastq/ERR278/007/ERR2784477/ERR2784477_1.fastq.gz
wget -nc ftp://ftp.sra.ebi.ac.uk/vol1/fastq/ERR278/006/ERR2784596/ERR2784596_2.fastq.gz
wget -nc ftp://ftp.sra.ebi.ac.uk/vol1/fastq/ERR278/007/ERR2784467/ERR2784467_1.fastq.gz
wget -nc ftp://ftp.sra.ebi.ac.uk/vol1/fastq/ERR278/005/ERR2784355/ERR2784355_1.fastq.gz
wget -nc ftp://ftp.sra.ebi.ac.uk/vol1/fastq/ERR278/004/ERR2784484/ERR2784484_2.fastq.gz
wget -nc ftp://ftp.sra.ebi.ac.uk/vol1/fastq/ERR278/009/ERR2784589/ERR2784589_1.fastq.gz
wget -nc ftp://ftp.sra.ebi.ac.uk/vol1/fastq/ERR278/004/ERR2784514/ERR2784514_1.fastq.gz
wget -nc ftp://ftp.sra.ebi.ac.uk/vol1/fastq/ERR278/002/ERR2784362/ERR2784362_2.fastq.gz
wget -nc ftp://ftp.sra.ebi.ac.uk/vol1/fastq/ERR278/006/ERR2784436/ERR2784436_2.fastq.gz
wget -nc ftp://ftp.sra.ebi.ac.uk/vol1/fastq/ERR278/009/ERR2784429/ERR2784429_1.fastq.gz
wget -nc ftp://ftp.sra.ebi.ac.uk/vol1/fastq/ERR278/003/ERR2784643/ERR2784643_2.fastq.gz
wget -nc ftp://ftp.sra.ebi.ac.uk/vol1/fastq/ERR278/008/ERR2784558/ERR2784558_2.fastq.gz
wget -nc ftp://ftp.sra.ebi.ac.uk/vol1/fastq/ERR278/007/ERR2784517/ERR2784517_2.fastq.gz
wget -nc ftp://ftp.sra.ebi.ac.uk/vol1/fastq/ERR278/007/ERR2784587/ERR2784587_2.fastq.gz
wget -nc ftp://ftp.sra.ebi.ac.uk/vol1/fastq/ERR278/007/ERR2784447/ERR2784447_2.fastq.gz
wget -nc ftp://ftp.sra.ebi.ac.uk/vol1/fastq/ERR278/006/ERR2784616/ERR2784616_1.fastq.gz
wget -nc ftp://ftp.sra.ebi.ac.uk/vol1/fastq/ERR278/007/ERR2784377/ERR2784377_2.fastq.gz
wget -nc ftp://ftp.sra.ebi.ac.uk/vol1/fastq/ERR278/006/ERR2784476/ERR2784476_1.fastq.gz
wget -nc ftp://ftp.sra.ebi.ac.uk/vol1/fastq/ERR278/006/ERR2784336/ERR2784336_1.fastq.gz
wget -nc ftp://ftp.sra.ebi.ac.uk/vol1/fastq/ERR278/006/ERR2784406/ERR2784406_1.fastq.gz
wget -nc ftp://ftp.sra.ebi.ac.uk/vol1/fastq/ERR278/006/ERR2784546/ERR2784546_1.fastq.gz
wget -nc ftp://ftp.sra.ebi.ac.uk/vol1/fastq/ERR278/009/ERR2784619/ERR2784619_2.fastq.gz
wget -nc ftp://ftp.sra.ebi.ac.uk/vol1/fastq/ERR278/009/ERR2784549/ERR2784549_2.fastq.gz
wget -nc ftp://ftp.sra.ebi.ac.uk/vol1/fastq/ERR278/009/ERR2784339/ERR2784339_2.fastq.gz
wget -nc ftp://ftp.sra.ebi.ac.uk/vol1/fastq/ERR278/009/ERR2784409/ERR2784409_2.fastq.gz
wget -nc ftp://ftp.sra.ebi.ac.uk/vol1/fastq/ERR278/009/ERR2784479/ERR2784479_2.fastq.gz
wget -nc ftp://ftp.sra.ebi.ac.uk/vol1/fastq/ERR278/008/ERR2784368/ERR2784368_1.fastq.gz
wget -nc ftp://ftp.sra.ebi.ac.uk/vol1/fastq/ERR278/008/ERR2784578/ERR2784578_1.fastq.gz
wget -nc ftp://ftp.sra.ebi.ac.uk/vol1/fastq/ERR278/008/ERR2784648/ERR2784648_1.fastq.gz
wget -nc ftp://ftp.sra.ebi.ac.uk/vol1/fastq/ERR278/008/ERR2784438/ERR2784438_1.fastq.gz
wget -nc ftp://ftp.sra.ebi.ac.uk/vol1/fastq/ERR278/008/ERR2784508/ERR2784508_1.fastq.gz
wget -nc ftp://ftp.sra.ebi.ac.uk/vol1/fastq/ERR278/002/ERR2784382/ERR2784382_2.fastq.gz
wget -nc ftp://ftp.sra.ebi.ac.uk/vol1/fastq/ERR278/001/ERR2784551/ERR2784551_1.fastq.gz
wget -nc ftp://ftp.sra.ebi.ac.uk/vol1/fastq/ERR278/001/ERR2784621/ERR2784621_1.fastq.gz
wget -nc ftp://ftp.sra.ebi.ac.uk/vol1/fastq/ERR278/002/ERR2784452/ERR2784452_2.fastq.gz
wget -nc ftp://ftp.sra.ebi.ac.uk/vol1/fastq/ERR278/002/ERR2784522/ERR2784522_2.fastq.gz
wget -nc ftp://ftp.sra.ebi.ac.uk/vol1/fastq/ERR278/003/ERR2784313/ERR2784313_2.fastq.gz
wget -nc ftp://ftp.sra.ebi.ac.uk/vol1/fastq/ERR278/005/ERR2784625/ERR2784625_2.fastq.gz
wget -nc ftp://ftp.sra.ebi.ac.uk/vol1/fastq/ERR278/002/ERR2784342/ERR2784342_1.fastq.gz
wget -nc ftp://ftp.sra.ebi.ac.uk/vol1/fastq/ERR278/002/ERR2784412/ERR2784412_1.fastq.gz
wget -nc ftp://ftp.sra.ebi.ac.uk/vol1/fastq/ERR278/005/ERR2784485/ERR2784485_2.fastq.gz
wget -nc ftp://ftp.sra.ebi.ac.uk/vol1/fastq/ERR278/005/ERR2784555/ERR2784555_2.fastq.gz
wget -nc ftp://ftp.sra.ebi.ac.uk/vol1/fastq/ERR278/001/ERR2784481/ERR2784481_1.fastq.gz
wget -nc ftp://ftp.sra.ebi.ac.uk/vol1/fastq/ERR278/003/ERR2784593/ERR2784593_2.fastq.gz
wget -nc ftp://ftp.sra.ebi.ac.uk/vol1/fastq/ERR278/000/ERR2784490/ERR2784490_2.fastq.gz
wget -nc ftp://ftp.sra.ebi.ac.uk/vol1/fastq/ERR278/000/ERR2784560/ERR2784560_2.fastq.gz
wget -nc ftp://ftp.sra.ebi.ac.uk/vol1/fastq/ERR278/000/ERR2784630/ERR2784630_2.fastq.gz
wget -nc ftp://ftp.sra.ebi.ac.uk/vol1/fastq/ERR278/003/ERR2784373/ERR2784373_1.fastq.gz
wget -nc ftp://ftp.sra.ebi.ac.uk/vol1/fastq/ERR278/003/ERR2784443/ERR2784443_1.fastq.gz
wget -nc ftp://ftp.sra.ebi.ac.uk/vol1/fastq/ERR278/004/ERR2784344/ERR2784344_2.fastq.gz
wget -nc ftp://ftp.sra.ebi.ac.uk/vol1/fastq/ERR278/004/ERR2784414/ERR2784414_2.fastq.gz
wget -nc ftp://ftp.sra.ebi.ac.uk/vol1/fastq/ERR278/003/ERR2784513/ERR2784513_1.fastq.gz
wget -nc ftp://ftp.sra.ebi.ac.uk/vol1/fastq/ERR278/004/ERR2784654/ERR2784654_1.fastq.gz
wget -nc ftp://ftp.sra.ebi.ac.uk/vol1/fastq/ERR278/004/ERR2784584/ERR2784584_1.fastq.gz
wget -nc ftp://ftp.sra.ebi.ac.uk/vol1/fastq/ERR278/007/ERR2784367/ERR2784367_1.fastq.gz
wget -nc ftp://ftp.sra.ebi.ac.uk/vol1/fastq/ERR278/007/ERR2784507/ERR2784507_1.fastq.gz
wget -nc ftp://ftp.sra.ebi.ac.uk/vol1/fastq/ERR278/007/ERR2784647/ERR2784647_1.fastq.gz
wget -nc ftp://ftp.sra.ebi.ac.uk/vol1/fastq/ERR278/007/ERR2784437/ERR2784437_1.fastq.gz
wget -nc ftp://ftp.sra.ebi.ac.uk/vol1/fastq/ERR278/007/ERR2784577/ERR2784577_1.fastq.gz
wget -nc ftp://ftp.sra.ebi.ac.uk/vol1/fastq/ERR278/005/ERR2784475/ERR2784475_1.fastq.gz
wget -nc ftp://ftp.sra.ebi.ac.uk/vol1/fastq/ERR278/005/ERR2784545/ERR2784545_1.fastq.gz
wget -nc ftp://ftp.sra.ebi.ac.uk/vol1/fastq/ERR278/005/ERR2784615/ERR2784615_1.fastq.gz
wget -nc ftp://ftp.sra.ebi.ac.uk/vol1/fastq/ERR278/009/ERR2784399/ERR2784399_1.fastq.gz
wget -nc ftp://ftp.sra.ebi.ac.uk/vol1/fastq/ERR278/009/ERR2784469/ERR2784469_1.fastq.gz
wget -nc ftp://ftp.sra.ebi.ac.uk/vol1/fastq/ERR278/009/ERR2784329/ERR2784329_1.fastq.gz
wget -nc ftp://ftp.sra.ebi.ac.uk/vol1/fastq/ERR278/009/ERR2784609/ERR2784609_1.fastq.gz
wget -nc ftp://ftp.sra.ebi.ac.uk/vol1/fastq/ERR278/009/ERR2784539/ERR2784539_1.fastq.gz
wget -nc ftp://ftp.sra.ebi.ac.uk/vol1/fastq/ERR278/001/ERR2784491/ERR2784491_2.fastq.gz
wget -nc ftp://ftp.sra.ebi.ac.uk/vol1/fastq/ERR278/001/ERR2784561/ERR2784561_2.fastq.gz
wget -nc ftp://ftp.sra.ebi.ac.uk/vol1/fastq/ERR278/001/ERR2784631/ERR2784631_2.fastq.gz
wget -nc ftp://ftp.sra.ebi.ac.uk/vol1/fastq/ERR278/001/ERR2784351/ERR2784351_2.fastq.gz
wget -nc ftp://ftp.sra.ebi.ac.uk/vol1/fastq/ERR278/001/ERR2784421/ERR2784421_2.fastq.gz
wget -nc ftp://ftp.sra.ebi.ac.uk/vol1/fastq/ERR278/008/ERR2784378/ERR2784378_2.fastq.gz
wget -nc ftp://ftp.sra.ebi.ac.uk/vol1/fastq/ERR278/002/ERR2784312/ERR2784312_2.fastq.gz
wget -nc ftp://ftp.sra.ebi.ac.uk/vol1/fastq/ERR278/006/ERR2784486/ERR2784486_2.fastq.gz
wget -nc ftp://ftp.sra.ebi.ac.uk/vol1/fastq/ERR278/006/ERR2784556/ERR2784556_2.fastq.gz
wget -nc ftp://ftp.sra.ebi.ac.uk/vol1/fastq/ERR278/006/ERR2784626/ERR2784626_2.fastq.gz
wget -nc ftp://ftp.sra.ebi.ac.uk/vol1/fastq/ERR278/003/ERR2784383/ERR2784383_2.fastq.gz
wget -nc ftp://ftp.sra.ebi.ac.uk/vol1/fastq/ERR278/003/ERR2784453/ERR2784453_2.fastq.gz
wget -nc ftp://ftp.sra.ebi.ac.uk/vol1/fastq/ERR278/000/ERR2784550/ERR2784550_1.fastq.gz
wget -nc ftp://ftp.sra.ebi.ac.uk/vol1/fastq/ERR278/000/ERR2784620/ERR2784620_1.fastq.gz
wget -nc ftp://ftp.sra.ebi.ac.uk/vol1/fastq/ERR278/000/ERR2784480/ERR2784480_1.fastq.gz
wget -nc ftp://ftp.sra.ebi.ac.uk/vol1/fastq/ERR278/005/ERR2784415/ERR2784415_2.fastq.gz
wget -nc ftp://ftp.sra.ebi.ac.uk/vol1/fastq/ERR278/005/ERR2784345/ERR2784345_2.fastq.gz
wget -nc ftp://ftp.sra.ebi.ac.uk/vol1/fastq/ERR278/003/ERR2784523/ERR2784523_2.fastq.gz
wget -nc ftp://ftp.sra.ebi.ac.uk/vol1/fastq/ERR278/004/ERR2784594/ERR2784594_2.fastq.gz
wget -nc ftp://ftp.sra.ebi.ac.uk/vol1/fastq/ERR278/000/ERR2784350/ERR2784350_2.fastq.gz
wget -nc ftp://ftp.sra.ebi.ac.uk/vol1/fastq/ERR278/000/ERR2784420/ERR2784420_2.fastq.gz
wget -nc ftp://ftp.sra.ebi.ac.uk/vol1/fastq/ERR278/003/ERR2784413/ERR2784413_1.fastq.gz
wget -nc ftp://ftp.sra.ebi.ac.uk/vol1/fastq/ERR278/003/ERR2784343/ERR2784343_1.fastq.gz
wget -nc ftp://ftp.sra.ebi.ac.uk/vol1/fastq/ERR278/008/ERR2784448/ERR2784448_2.fastq.gz
wget -nc ftp://ftp.sra.ebi.ac.uk/vol1/fastq/ERR278/008/ERR2784518/ERR2784518_2.fastq.gz
wget -nc ftp://ftp.sra.ebi.ac.uk/vol1/fastq/ERR278/002/ERR2784372/ERR2784372_1.fastq.gz
wget -nc ftp://ftp.sra.ebi.ac.uk/vol1/fastq/ERR278/002/ERR2784442/ERR2784442_1.fastq.gz
wget -nc ftp://ftp.sra.ebi.ac.uk/vol1/fastq/ERR278/002/ERR2784512/ERR2784512_1.fastq.gz
wget -nc ftp://ftp.sra.ebi.ac.uk/vol1/fastq/ERR278/003/ERR2784583/ERR2784583_1.fastq.gz
wget -nc ftp://ftp.sra.ebi.ac.uk/vol1/fastq/ERR278/003/ERR2784653/ERR2784653_1.fastq.gz
wget -nc ftp://ftp.sra.ebi.ac.uk/vol1/fastq/ERR278/006/ERR2784576/ERR2784576_1.fastq.gz
wget -nc ftp://ftp.sra.ebi.ac.uk/vol1/fastq/ERR278/006/ERR2784436/ERR2784436_1.fastq.gz
wget -nc ftp://ftp.sra.ebi.ac.uk/vol1/fastq/ERR278/000/ERR2784320/ERR2784320_2.fastq.gz
wget -nc ftp://ftp.sra.ebi.ac.uk/vol1/fastq/ERR278/000/ERR2784460/ERR2784460_2.fastq.gz
wget -nc ftp://ftp.sra.ebi.ac.uk/vol1/fastq/ERR278/000/ERR2784600/ERR2784600_2.fastq.gz
wget -nc ftp://ftp.sra.ebi.ac.uk/vol1/fastq/ERR278/007/ERR2784617/ERR2784617_2.fastq.gz
wget -nc ftp://ftp.sra.ebi.ac.uk/vol1/fastq/ERR278/007/ERR2784337/ERR2784337_2.fastq.gz
wget -nc ftp://ftp.sra.ebi.ac.uk/vol1/fastq/ERR278/007/ERR2784477/ERR2784477_2.fastq.gz
wget -nc ftp://ftp.sra.ebi.ac.uk/vol1/fastq/ERR278/004/ERR2784614/ERR2784614_1.fastq.gz
wget -nc ftp://ftp.sra.ebi.ac.uk/vol1/fastq/ERR278/004/ERR2784404/ERR2784404_1.fastq.gz
wget -nc ftp://ftp.sra.ebi.ac.uk/vol1/fastq/ERR278/004/ERR2784474/ERR2784474_1.fastq.gz
wget -nc ftp://ftp.sra.ebi.ac.uk/vol1/fastq/ERR278/008/ERR2784538/ERR2784538_1.fastq.gz
wget -nc ftp://ftp.sra.ebi.ac.uk/vol1/fastq/ERR278/001/ERR2784381/ERR2784381_1.fastq.gz
wget -nc ftp://ftp.sra.ebi.ac.uk/vol1/fastq/ERR278/001/ERR2784311/ERR2784311_1.fastq.gz
wget -nc ftp://ftp.sra.ebi.ac.uk/vol1/fastq/ERR278/001/ERR2784521/ERR2784521_1.fastq.gz
wget -nc ftp://ftp.sra.ebi.ac.uk/vol1/fastq/ERR278/002/ERR2784422/ERR2784422_2.fastq.gz
wget -nc ftp://ftp.sra.ebi.ac.uk/vol1/fastq/ERR278/002/ERR2784562/ERR2784562_2.fastq.gz
wget -nc ftp://ftp.sra.ebi.ac.uk/vol1/fastq/ERR278/005/ERR2784655/ERR2784655_2.fastq.gz
wget -nc ftp://ftp.sra.ebi.ac.uk/vol1/fastq/ERR278/005/ERR2784515/ERR2784515_2.fastq.gz
wget -nc ftp://ftp.sra.ebi.ac.uk/vol1/fastq/ERR278/005/ERR2784375/ERR2784375_2.fastq.gz
wget -nc ftp://ftp.sra.ebi.ac.uk/vol1/fastq/ERR278/007/ERR2784487/ERR2784487_2.fastq.gz
wget -nc ftp://ftp.sra.ebi.ac.uk/vol1/fastq/ERR278/006/ERR2784346/ERR2784346_2.fastq.gz
wget -nc ftp://ftp.sra.ebi.ac.uk/vol1/fastq/ERR278/001/ERR2784591/ERR2784591_2.fastq.gz
wget -nc ftp://ftp.sra.ebi.ac.uk/vol1/fastq/ERR278/000/ERR2784340/ERR2784340_1.fastq.gz
wget -nc ftp://ftp.sra.ebi.ac.uk/vol1/fastq/ERR278/004/ERR2784384/ERR2784384_2.fastq.gz
wget -nc ftp://ftp.sra.ebi.ac.uk/vol1/fastq/ERR278/009/ERR2784449/ERR2784449_2.fastq.gz
wget -nc ftp://ftp.sra.ebi.ac.uk/vol1/fastq/ERR278/003/ERR2784623/ERR2784623_1.fastq.gz
wget -nc ftp://ftp.sra.ebi.ac.uk/vol1/fastq/ERR278/003/ERR2784483/ERR2784483_1.fastq.gz
wget -nc ftp://ftp.sra.ebi.ac.uk/vol1/fastq/ERR278/004/ERR2784524/ERR2784524_2.fastq.gz
wget -nc ftp://ftp.sra.ebi.ac.uk/vol1/fastq/ERR278/008/ERR2784548/ERR2784548_1.fastq.gz
wget -nc ftp://ftp.sra.ebi.ac.uk/vol1/fastq/ERR278/004/ERR2784414/ERR2784414_1.fastq.gz
wget -nc ftp://ftp.sra.ebi.ac.uk/vol1/fastq/ERR278/007/ERR2784407/ERR2784407_1.fastq.gz
wget -nc ftp://ftp.sra.ebi.ac.uk/vol1/fastq/ERR278/008/ERR2784398/ERR2784398_1.fastq.gz
wget -nc ftp://ftp.sra.ebi.ac.uk/vol1/fastq/ERR278/003/ERR2784553/ERR2784553_2.fastq.gz
wget -nc ftp://ftp.sra.ebi.ac.uk/vol1/fastq/ERR278/002/ERR2784412/ERR2784412_2.fastq.gz
wget -nc ftp://ftp.sra.ebi.ac.uk/vol1/fastq/ERR278/009/ERR2784439/ERR2784439_2.fastq.gz
wget -nc ftp://ftp.sra.ebi.ac.uk/vol1/fastq/ERR278/001/ERR2784511/ERR2784511_1.fastq.gz
wget -nc ftp://ftp.sra.ebi.ac.uk/vol1/fastq/ERR278/001/ERR2784371/ERR2784371_1.fastq.gz
wget -nc ftp://ftp.sra.ebi.ac.uk/vol1/fastq/ERR278/002/ERR2784652/ERR2784652_1.fastq.gz
wget -nc ftp://ftp.sra.ebi.ac.uk/vol1/fastq/ERR278/000/ERR2784450/ERR2784450_2.fastq.gz
wget -nc ftp://ftp.sra.ebi.ac.uk/vol1/fastq/ERR278/001/ERR2784311/ERR2784311_2.fastq.gz
wget -nc ftp://ftp.sra.ebi.ac.uk/vol1/fastq/ERR278/007/ERR2784627/ERR2784627_2.fastq.gz
wget -nc ftp://ftp.sra.ebi.ac.uk/vol1/fastq/ERR278/005/ERR2784445/ERR2784445_1.fastq.gz
wget -nc ftp://ftp.sra.ebi.ac.uk/vol1/fastq/ERR278/006/ERR2784586/ERR2784586_1.fastq.gz
wget -nc ftp://ftp.sra.ebi.ac.uk/vol1/fastq/ERR278/000/ERR2784350/ERR2784350_1.fastq.gz
wget -nc ftp://ftp.sra.ebi.ac.uk/vol1/fastq/ERR278/008/ERR2784338/ERR2784338_2.fastq.gz
wget -nc ftp://ftp.sra.ebi.ac.uk/vol1/fastq/ERR278/008/ERR2784478/ERR2784478_2.fastq.gz
wget -nc ftp://ftp.sra.ebi.ac.uk/vol1/fastq/ERR278/008/ERR2784618/ERR2784618_2.fastq.gz
wget -nc ftp://ftp.sra.ebi.ac.uk/vol1/fastq/ERR278/003/ERR2784613/ERR2784613_1.fastq.gz
wget -nc ftp://ftp.sra.ebi.ac.uk/vol1/fastq/ERR278/001/ERR2784601/ERR2784601_2.fastq.gz
wget -nc ftp://ftp.sra.ebi.ac.uk/vol1/fastq/ERR278/003/ERR2784473/ERR2784473_1.fastq.gz
wget -nc ftp://ftp.sra.ebi.ac.uk/vol1/fastq/ERR278/001/ERR2784461/ERR2784461_2.fastq.gz
wget -nc ftp://ftp.sra.ebi.ac.uk/vol1/fastq/ERR278/005/ERR2784405/ERR2784405_1.fastq.gz
wget -nc ftp://ftp.sra.ebi.ac.uk/vol1/fastq/ERR278/006/ERR2784516/ERR2784516_2.fastq.gz
wget -nc ftp://ftp.sra.ebi.ac.uk/vol1/fastq/ERR278/006/ERR2784656/ERR2784656_2.fastq.gz
wget -nc ftp://ftp.sra.ebi.ac.uk/vol1/fastq/ERR278/000/ERR2784520/ERR2784520_1.fastq.gz
wget -nc ftp://ftp.sra.ebi.ac.uk/vol1/fastq/ERR278/003/ERR2784423/ERR2784423_2.fastq.gz
wget -nc ftp://ftp.sra.ebi.ac.uk/vol1/fastq/ERR278/006/ERR2784376/ERR2784376_2.fastq.gz
wget -nc ftp://ftp.sra.ebi.ac.uk/vol1/fastq/ERR278/000/ERR2784380/ERR2784380_1.fastq.gz
wget -nc ftp://ftp.sra.ebi.ac.uk/vol1/fastq/ERR278/009/ERR2784369/ERR2784369_1.fastq.gz
wget -nc ftp://ftp.sra.ebi.ac.uk/vol1/fastq/ERR278/009/ERR2784509/ERR2784509_1.fastq.gz
wget -nc ftp://ftp.sra.ebi.ac.uk/vol1/fastq/ERR278/003/ERR2784563/ERR2784563_2.fastq.gz
wget -nc ftp://ftp.sra.ebi.ac.uk/vol1/fastq/ERR278/005/ERR2784435/ERR2784435_1.fastq.gz
wget -nc ftp://ftp.sra.ebi.ac.uk/vol1/fastq/ERR278/005/ERR2784575/ERR2784575_1.fastq.gz
wget -nc ftp://ftp.sra.ebi.ac.uk/vol1/fastq/ERR278/009/ERR2784649/ERR2784649_1.fastq.gz
wget -nc ftp://ftp.sra.ebi.ac.uk/vol1/fastq/ERR278/002/ERR2784312/ERR2784312_1.fastq.gz
wget -nc ftp://ftp.sra.ebi.ac.uk/vol1/fastq/ERR278/002/ERR2784592/ERR2784592_2.fastq.gz
wget -nc ftp://ftp.sra.ebi.ac.uk/vol1/fastq/ERR278/007/ERR2784347/ERR2784347_2.fastq.gz
wget -nc ftp://ftp.sra.ebi.ac.uk/vol1/fastq/ERR278/002/ERR2784622/ERR2784622_1.fastq.gz
wget -nc ftp://ftp.sra.ebi.ac.uk/vol1/fastq/ERR278/002/ERR2784482/ERR2784482_1.fastq.gz
wget -nc ftp://ftp.sra.ebi.ac.uk/vol1/fastq/ERR278/001/ERR2784341/ERR2784341_1.fastq.gz
wget -nc ftp://ftp.sra.ebi.ac.uk/vol1/fastq/ERR278/005/ERR2784415/ERR2784415_1.fastq.gz
wget -nc ftp://ftp.sra.ebi.ac.uk/vol1/fastq/ERR278/007/ERR2784547/ERR2784547_1.fastq.gz
wget -nc ftp://ftp.sra.ebi.ac.uk/vol1/fastq/ERR278/007/ERR2784397/ERR2784397_1.fastq.gz
wget -nc ftp://ftp.sra.ebi.ac.uk/vol1/fastq/ERR278/003/ERR2784413/ERR2784413_2.fastq.gz
wget -nc ftp://ftp.sra.ebi.ac.uk/vol1/fastq/ERR278/005/ERR2784525/ERR2784525_2.fastq.gz
wget -nc ftp://ftp.sra.ebi.ac.uk/vol1/fastq/ERR278/007/ERR2784537/ERR2784537_1.fastq.gz
wget -nc ftp://ftp.sra.ebi.ac.uk/vol1/fastq/ERR278/005/ERR2784385/ERR2784385_2.fastq.gz
wget -nc ftp://ftp.sra.ebi.ac.uk/vol1/fastq/ERR278/004/ERR2784554/ERR2784554_2.fastq.gz
wget -nc ftp://ftp.sra.ebi.ac.uk/vol1/fastq/ERR278/001/ERR2784651/ERR2784651_1.fastq.gz
wget -nc ftp://ftp.sra.ebi.ac.uk/vol1/fastq/ERR278/004/ERR2784444/ERR2784444_1.fastq.gz
wget -nc ftp://ftp.sra.ebi.ac.uk/vol1/fastq/ERR278/009/ERR2784359/ERR2784359_1.fastq.gz
wget -nc ftp://ftp.sra.ebi.ac.uk/vol1/fastq/ERR278/001/ERR2784451/ERR2784451_2.fastq.gz
wget -nc ftp://ftp.sra.ebi.ac.uk/vol1/fastq/ERR278/008/ERR2784488/ERR2784488_2.fastq.gz
wget -nc ftp://ftp.sra.ebi.ac.uk/vol1/fastq/ERR278/005/ERR2784585/ERR2784585_1.fastq.gz
wget -nc ftp://ftp.sra.ebi.ac.uk/vol1/fastq/ERR278/008/ERR2784628/ERR2784628_2.fastq.gz
wget -nc ftp://ftp.sra.ebi.ac.uk/vol1/fastq/ERR278/003/ERR2784443/ERR2784443_2.fastq.gz
wget -nc ftp://ftp.sra.ebi.ac.uk/vol1/fastq/ERR278/002/ERR2784612/ERR2784612_1.fastq.gz
wget -nc ftp://ftp.sra.ebi.ac.uk/vol1/fastq/ERR278/003/ERR2784583/ERR2784583_2.fastq.gz
wget -nc ftp://ftp.sra.ebi.ac.uk/vol1/fastq/ERR278/003/ERR2784373/ERR2784373_2.fastq.gz
wget -nc ftp://ftp.sra.ebi.ac.uk/vol1/fastq/ERR278/002/ERR2784332/ERR2784332_1.fastq.gz
wget -nc ftp://ftp.sra.ebi.ac.uk/vol1/fastq/ERR278/002/ERR2784602/ERR2784602_2.fastq.gz
wget -nc ftp://ftp.sra.ebi.ac.uk/vol1/fastq/ERR278/002/ERR2784532/ERR2784532_2.fastq.gz
wget -nc ftp://ftp.sra.ebi.ac.uk/vol1/fastq/ERR278/005/ERR2784545/ERR2784545_2.fastq.gz
wget -nc ftp://ftp.sra.ebi.ac.uk/vol1/fastq/ERR278/000/ERR2784650/ERR2784650_1.fastq.gz
wget -nc ftp://ftp.sra.ebi.ac.uk/vol1/fastq/ERR278/005/ERR2784335/ERR2784335_2.fastq.gz
wget -nc ftp://ftp.sra.ebi.ac.uk/vol1/fastq/ERR278/000/ERR2784370/ERR2784370_1.fastq.gz
wget -nc ftp://ftp.sra.ebi.ac.uk/vol1/fastq/ERR278/005/ERR2784475/ERR2784475_2.fastq.gz
wget -nc ftp://ftp.sra.ebi.ac.uk/vol1/fastq/ERR278/005/ERR2784405/ERR2784405_2.fastq.gz
wget -nc ftp://ftp.sra.ebi.ac.uk/vol1/fastq/ERR278/001/ERR2784491/ERR2784491_1.fastq.gz
wget -nc ftp://ftp.sra.ebi.ac.uk/vol1/fastq/ERR278/001/ERR2784561/ERR2784561_1.fastq.gz
wget -nc ftp://ftp.sra.ebi.ac.uk/vol1/fastq/ERR278/001/ERR2784421/ERR2784421_1.fastq.gz
wget -nc ftp://ftp.sra.ebi.ac.uk/vol1/fastq/ERR278/001/ERR2784481/ERR2784481_2.fastq.gz
wget -nc ftp://ftp.sra.ebi.ac.uk/vol1/fastq/ERR278/001/ERR2784341/ERR2784341_2.fastq.gz
wget -nc ftp://ftp.sra.ebi.ac.uk/vol1/fastq/ERR278/005/ERR2784625/ERR2784625_1.fastq.gz
wget -nc ftp://ftp.sra.ebi.ac.uk/vol1/fastq/ERR278/005/ERR2784555/ERR2784555_1.fastq.gz
wget -nc ftp://ftp.sra.ebi.ac.uk/vol1/fastq/ERR278/004/ERR2784364/ERR2784364_1.fastq.gz
wget -nc ftp://ftp.sra.ebi.ac.uk/vol1/fastq/ERR278/004/ERR2784634/ERR2784634_2.fastq.gz
wget -nc ftp://ftp.sra.ebi.ac.uk/vol1/fastq/ERR278/003/ERR2784523/ERR2784523_1.fastq.gz
wget -nc ftp://ftp.sra.ebi.ac.uk/vol1/fastq/ERR278/003/ERR2784593/ERR2784593_1.fastq.gz
wget -nc ftp://ftp.sra.ebi.ac.uk/vol1/fastq/ERR278/003/ERR2784453/ERR2784453_1.fastq.gz
wget -nc ftp://ftp.sra.ebi.ac.uk/vol1/fastq/ERR278/008/ERR2784358/ERR2784358_1.fastq.gz
wget -nc ftp://ftp.sra.ebi.ac.uk/vol1/fastq/ERR278/008/ERR2784428/ERR2784428_1.fastq.gz
wget -nc ftp://ftp.sra.ebi.ac.uk/vol1/fastq/ERR278/006/ERR2784386/ERR2784386_2.fastq.gz
wget -nc ftp://ftp.sra.ebi.ac.uk/vol1/fastq/ERR278/000/ERR2784640/ERR2784640_2.fastq.gz
wget -nc ftp://ftp.sra.ebi.ac.uk/vol1/fastq/ERR278/007/ERR2784507/ERR2784507_2.fastq.gz
wget -nc ftp://ftp.sra.ebi.ac.uk/vol1/fastq/ERR278/007/ERR2784437/ERR2784437_2.fastq.gz
wget -nc ftp://ftp.sra.ebi.ac.uk/vol1/fastq/ERR278/006/ERR2784466/ERR2784466_1.fastq.gz
wget -nc ftp://ftp.sra.ebi.ac.uk/vol1/fastq/ERR278/006/ERR2784396/ERR2784396_1.fastq.gz
wget -nc ftp://ftp.sra.ebi.ac.uk/vol1/fastq/ERR278/000/ERR2784570/ERR2784570_2.fastq.gz
wget -nc ftp://ftp.sra.ebi.ac.uk/vol1/fastq/ERR278/007/ERR2784517/ERR2784517_1.fastq.gz
wget -nc ftp://ftp.sra.ebi.ac.uk/vol1/fastq/ERR278/008/ERR2784588/ERR2784588_1.fastq.gz
wget -nc ftp://ftp.sra.ebi.ac.uk/vol1/fastq/ERR278/009/ERR2784629/ERR2784629_2.fastq.gz
wget -nc ftp://ftp.sra.ebi.ac.uk/vol1/fastq/ERR278/009/ERR2784639/ERR2784639_1.fastq.gz
wget -nc ftp://ftp.sra.ebi.ac.uk/vol1/fastq/ERR278/008/ERR2784348/ERR2784348_2.fastq.gz
wget -nc ftp://ftp.sra.ebi.ac.uk/vol1/fastq/ERR278/009/ERR2784559/ERR2784559_2.fastq.gz
wget -nc ftp://ftp.sra.ebi.ac.uk/vol1/fastq/ERR278/001/ERR2784321/ERR2784321_2.fastq.gz
wget -nc ftp://ftp.sra.ebi.ac.uk/vol1/fastq/ERR278/005/ERR2784565/ERR2784565_2.fastq.gz
wget -nc ftp://ftp.sra.ebi.ac.uk/vol1/fastq/ERR278/005/ERR2784495/ERR2784495_2.fastq.gz
wget -nc ftp://ftp.sra.ebi.ac.uk/vol1/fastq/ERR278/005/ERR2784355/ERR2784355_2.fastq.gz
wget -nc ftp://ftp.sra.ebi.ac.uk/vol1/fastq/ERR278/005/ERR2784425/ERR2784425_2.fastq.gz
wget -nc ftp://ftp.sra.ebi.ac.uk/vol1/fastq/ERR278/000/ERR2784630/ERR2784630_1.fastq.gz
wget -nc ftp://ftp.sra.ebi.ac.uk/vol1/fastq/ERR278/004/ERR2784654/ERR2784654_2.fastq.gz
wget -nc ftp://ftp.sra.ebi.ac.uk/vol1/fastq/ERR278/004/ERR2784514/ERR2784514_2.fastq.gz
wget -nc ftp://ftp.sra.ebi.ac.uk/vol1/fastq/ERR278/006/ERR2784616/ERR2784616_2.fastq.gz
wget -nc ftp://ftp.sra.ebi.ac.uk/vol1/fastq/ERR278/003/ERR2784403/ERR2784403_1.fastq.gz
wget -nc ftp://ftp.sra.ebi.ac.uk/vol1/fastq/ERR278/002/ERR2784352/ERR2784352_1.fastq.gz
wget -nc ftp://ftp.sra.ebi.ac.uk/vol1/fastq/ERR278/003/ERR2784393/ERR2784393_2.fastq.gz
wget -nc ftp://ftp.sra.ebi.ac.uk/vol1/fastq/ERR278/003/ERR2784463/ERR2784463_2.fastq.gz
wget -nc ftp://ftp.sra.ebi.ac.uk/vol1/fastq/ERR278/007/ERR2784317/ERR2784317_2.fastq.gz
wget -nc ftp://ftp.sra.ebi.ac.uk/vol1/fastq/ERR278/007/ERR2784457/ERR2784457_2.fastq.gz
wget -nc ftp://ftp.sra.ebi.ac.uk/vol1/fastq/ERR278/000/ERR2784410/ERR2784410_2.fastq.gz
wget -nc ftp://ftp.sra.ebi.ac.uk/vol1/fastq/ERR278/007/ERR2784527/ERR2784527_2.fastq.gz
wget -nc ftp://ftp.sra.ebi.ac.uk/vol1/fastq/ERR278/003/ERR2784503/ERR2784503_1.fastq.gz
wget -nc ftp://ftp.sra.ebi.ac.uk/vol1/fastq/ERR278/003/ERR2784433/ERR2784433_1.fastq.gz
wget -nc ftp://ftp.sra.ebi.ac.uk/vol1/fastq/ERR278/003/ERR2784573/ERR2784573_1.fastq.gz
wget -nc ftp://ftp.sra.ebi.ac.uk/vol1/fastq/ERR278/003/ERR2784643/ERR2784643_1.fastq.gz
wget -nc ftp://ftp.sra.ebi.ac.uk/vol1/fastq/ERR278/002/ERR2784382/ERR2784382_1.fastq.gz
wget -nc ftp://ftp.sra.ebi.ac.uk/vol1/fastq/ERR278/001/ERR2784541/ERR2784541_1.fastq.gz
wget -nc ftp://ftp.sra.ebi.ac.uk/vol1/fastq/ERR278/001/ERR2784471/ERR2784471_1.fastq.gz
wget -nc ftp://ftp.sra.ebi.ac.uk/vol1/fastq/ERR278/004/ERR2784314/ERR2784314_1.fastq.gz
wget -nc ftp://ftp.sra.ebi.ac.uk/vol1/fastq/ERR278/008/ERR2784498/ERR2784498_1.fastq.gz
wget -nc ftp://ftp.sra.ebi.ac.uk/vol1/fastq/ERR278/008/ERR2784568/ERR2784568_1.fastq.gz
wget -nc ftp://ftp.sra.ebi.ac.uk/vol1/fastq/ERR278/009/ERR2784619/ERR2784619_1.fastq.gz
wget -nc ftp://ftp.sra.ebi.ac.uk/vol1/fastq/ERR278/005/ERR2784605/ERR2784605_1.fastq.gz
wget -nc ftp://ftp.sra.ebi.ac.uk/vol1/fastq/ERR278/005/ERR2784535/ERR2784535_1.fastq.gz
wget -nc ftp://ftp.sra.ebi.ac.uk/vol1/fastq/ERR278/009/ERR2784589/ERR2784589_2.fastq.gz
wget -nc ftp://ftp.sra.ebi.ac.uk/vol1/fastq/ERR278/008/ERR2784338/ERR2784338_1.fastq.gz
wget -nc ftp://ftp.sra.ebi.ac.uk/vol1/fastq/ERR278/008/ERR2784408/ERR2784408_1.fastq.gz
wget -nc ftp://ftp.sra.ebi.ac.uk/vol1/fastq/ERR278/004/ERR2784484/ERR2784484_1.fastq.gz
wget -nc ftp://ftp.sra.ebi.ac.uk/vol1/fastq/ERR278/008/ERR2784328/ERR2784328_1.fastq.gz
wget -nc ftp://ftp.sra.ebi.ac.uk/vol1/fastq/ERR278/002/ERR2784552/ERR2784552_2.fastq.gz
wget -nc ftp://ftp.sra.ebi.ac.uk/vol1/fastq/ERR278/002/ERR2784622/ERR2784622_2.fastq.gz
wget -nc ftp://ftp.sra.ebi.ac.uk/vol1/fastq/ERR278/009/ERR2784579/ERR2784579_2.fastq.gz
wget -nc ftp://ftp.sra.ebi.ac.uk/vol1/fastq/ERR278/009/ERR2784649/ERR2784649_2.fastq.gz
wget -nc ftp://ftp.sra.ebi.ac.uk/vol1/fastq/ERR278/000/ERR2784430/ERR2784430_2.fastq.gz
wget -nc ftp://ftp.sra.ebi.ac.uk/vol1/fastq/ERR278/008/ERR2784368/ERR2784368_2.fastq.gz
wget -nc ftp://ftp.sra.ebi.ac.uk/vol1/fastq/ERR278/000/ERR2784500/ERR2784500_2.fastq.gz
wget -nc ftp://ftp.sra.ebi.ac.uk/vol1/fastq/ERR278/000/ERR2784590/ERR2784590_2.fastq.gz
wget -nc ftp://ftp.sra.ebi.ac.uk/vol1/fastq/ERR278/009/ERR2784419/ERR2784419_2.fastq.gz
wget -nc ftp://ftp.sra.ebi.ac.uk/vol1/fastq/ERR278/006/ERR2784446/ERR2784446_1.fastq.gz
wget -nc ftp://ftp.sra.ebi.ac.uk/vol1/fastq/ERR278/006/ERR2784376/ERR2784376_1.fastq.gz
wget -nc ftp://ftp.sra.ebi.ac.uk/vol1/fastq/ERR278/005/ERR2784525/ERR2784525_1.fastq.gz
wget -nc ftp://ftp.sra.ebi.ac.uk/vol1/fastq/ERR278/006/ERR2784366/ERR2784366_1.fastq.gz
wget -nc ftp://ftp.sra.ebi.ac.uk/vol1/fastq/ERR278/000/ERR2784530/ERR2784530_2.fastq.gz
wget -nc ftp://ftp.sra.ebi.ac.uk/vol1/fastq/ERR278/007/ERR2784547/ERR2784547_2.fastq.gz
wget -nc ftp://ftp.sra.ebi.ac.uk/vol1/fastq/ERR278/007/ERR2784407/ERR2784407_2.fastq.gz
wget -nc ftp://ftp.sra.ebi.ac.uk/vol1/fastq/ERR278/007/ERR2784627/ERR2784627_1.fastq.gz
wget -nc ftp://ftp.sra.ebi.ac.uk/vol1/fastq/ERR278/008/ERR2784388/ERR2784388_2.fastq.gz
wget -nc ftp://ftp.sra.ebi.ac.uk/vol1/fastq/ERR278/004/ERR2784334/ERR2784334_1.fastq.gz
wget -nc ftp://ftp.sra.ebi.ac.uk/vol1/fastq/ERR278/003/ERR2784563/ERR2784563_1.fastq.gz
wget -nc ftp://ftp.sra.ebi.ac.uk/vol1/fastq/ERR278/004/ERR2784604/ERR2784604_2.fastq.gz
wget -nc ftp://ftp.sra.ebi.ac.uk/vol1/fastq/ERR278/001/ERR2784591/ERR2784591_1.fastq.gz
wget -nc ftp://ftp.sra.ebi.ac.uk/vol1/fastq/ERR278/001/ERR2784451/ERR2784451_1.fastq.gz
wget -nc ftp://ftp.sra.ebi.ac.uk/vol1/fastq/ERR278/002/ERR2784632/ERR2784632_2.fastq.gz
wget -nc ftp://ftp.sra.ebi.ac.uk/vol1/fastq/ERR278/005/ERR2784585/ERR2784585_2.fastq.gz
wget -nc ftp://ftp.sra.ebi.ac.uk/vol1/fastq/ERR278/005/ERR2784445/ERR2784445_2.fastq.gz
wget -nc ftp://ftp.sra.ebi.ac.uk/vol1/fastq/ERR278/000/ERR2784610/ERR2784610_1.fastq.gz
wget -nc ftp://ftp.sra.ebi.ac.uk/vol1/fastq/ERR278/001/ERR2784371/ERR2784371_2.fastq.gz
wget -nc ftp://ftp.sra.ebi.ac.uk/vol1/fastq/ERR278/000/ERR2784400/ERR2784400_2.fastq.gz
wget -nc ftp://ftp.sra.ebi.ac.uk/vol1/fastq/ERR278/004/ERR2784394/ERR2784394_1.fastq.gz
wget -nc ftp://ftp.sra.ebi.ac.uk/vol1/fastq/ERR278/007/ERR2784557/ERR2784557_2.fastq.gz
wget -nc ftp://ftp.sra.ebi.ac.uk/vol1/fastq/ERR278/008/ERR2784398/ERR2784398_2.fastq.gz
wget -nc ftp://ftp.sra.ebi.ac.uk/vol1/fastq/ERR278/007/ERR2784637/ERR2784637_1.fastq.gz
wget -nc ftp://ftp.sra.ebi.ac.uk/vol1/fastq/ERR278/006/ERR2784356/ERR2784356_1.fastq.gz
wget -nc ftp://ftp.sra.ebi.ac.uk/vol1/fastq/ERR278/003/ERR2784553/ERR2784553_1.fastq.gz
wget -nc ftp://ftp.sra.ebi.ac.uk/vol1/fastq/ERR278/009/ERR2784519/ERR2784519_2.fastq.gz
wget -nc ftp://ftp.sra.ebi.ac.uk/vol1/fastq/ERR278/008/ERR2784478/ERR2784478_1.fastq.gz
wget -nc ftp://ftp.sra.ebi.ac.uk/vol1/fastq/ERR278/004/ERR2784344/ERR2784344_1.fastq.gz
wget -nc ftp://ftp.sra.ebi.ac.uk/vol1/fastq/ERR278/003/ERR2784473/ERR2784473_2.fastq.gz
wget -nc ftp://ftp.sra.ebi.ac.uk/vol1/fastq/ERR278/008/ERR2784468/ERR2784468_1.fastq.gz
wget -nc ftp://ftp.sra.ebi.ac.uk/vol1/fastq/ERR278/003/ERR2784483/ERR2784483_2.fastq.gz
wget -nc ftp://ftp.sra.ebi.ac.uk/vol1/fastq/ERR278/005/ERR2784595/ERR2784595_2.fastq.gz
wget -nc ftp://ftp.sra.ebi.ac.uk/vol1/fastq/ERR278/005/ERR2784315/ERR2784315_2.fastq.gz
wget -nc ftp://ftp.sra.ebi.ac.uk/vol1/fastq/ERR278/009/ERR2784509/ERR2784509_2.fastq.gz
wget -nc ftp://ftp.sra.ebi.ac.uk/vol1/fastq/ERR278/001/ERR2784361/ERR2784361_2.fastq.gz
wget -nc ftp://ftp.sra.ebi.ac.uk/vol1/fastq/ERR278/001/ERR2784441/ERR2784441_1.fastq.gz
wget -nc ftp://ftp.sra.ebi.ac.uk/vol1/fastq/ERR278/000/ERR2784520/ERR2784520_2.fastq.gz
wget -nc ftp://ftp.sra.ebi.ac.uk/vol1/fastq/ERR278/002/ERR2784642/ERR2784642_2.fastq.gz
wget -nc ftp://ftp.sra.ebi.ac.uk/vol1/fastq/ERR278/000/ERR2784600/ERR2784600_1.fastq.gz
wget -nc ftp://ftp.sra.ebi.ac.uk/vol1/fastq/ERR278/005/ERR2784435/ERR2784435_2.fastq.gz
wget -nc ftp://ftp.sra.ebi.ac.uk/vol1/fastq/ERR278/005/ERR2784515/ERR2784515_1.fastq.gz
wget -nc ftp://ftp.sra.ebi.ac.uk/vol1/fastq/ERR278/001/ERR2784391/ERR2784391_2.fastq.gz
wget -nc ftp://ftp.sra.ebi.ac.uk/vol1/fastq/ERR278/009/ERR2784529/ERR2784529_2.fastq.gz
wget -nc ftp://ftp.sra.ebi.ac.uk/vol1/fastq/ERR278/003/ERR2784543/ERR2784543_1.fastq.gz
wget -nc ftp://ftp.sra.ebi.ac.uk/vol1/fastq/ERR278/003/ERR2784323/ERR2784323_2.fastq.gz
wget -nc ftp://ftp.sra.ebi.ac.uk/vol1/fastq/ERR278/006/ERR2784486/ERR2784486_1.fastq.gz
wget -nc ftp://ftp.sra.ebi.ac.uk/vol1/fastq/ERR278/006/ERR2784346/ERR2784346_1.fastq.gz
wget -nc ftp://ftp.sra.ebi.ac.uk/vol1/fastq/ERR278/000/ERR2784550/ERR2784550_2.fastq.gz
wget -nc ftp://ftp.sra.ebi.ac.uk/vol1/fastq/ERR278/007/ERR2784597/ERR2784597_2.fastq.gz
wget -nc ftp://ftp.sra.ebi.ac.uk/vol1/fastq/ERR278/004/ERR2784614/ERR2784614_2.fastq.gz
wget -nc ftp://ftp.sra.ebi.ac.uk/vol1/fastq/ERR278/003/ERR2784353/ERR2784353_2.fastq.gz
wget -nc ftp://ftp.sra.ebi.ac.uk/vol1/fastq/ERR278/008/ERR2784598/ERR2784598_1.fastq.gz
wget -nc ftp://ftp.sra.ebi.ac.uk/vol1/fastq/ERR278/008/ERR2784458/ERR2784458_1.fastq.gz
wget -nc ftp://ftp.sra.ebi.ac.uk/vol1/fastq/ERR278/003/ERR2784493/ERR2784493_2.fastq.gz
wget -nc ftp://ftp.sra.ebi.ac.uk/vol1/fastq/ERR278/007/ERR2784567/ERR2784567_2.fastq.gz
wget -nc ftp://ftp.sra.ebi.ac.uk/vol1/fastq/ERR278/005/ERR2784645/ERR2784645_1.fastq.gz
wget -nc ftp://ftp.sra.ebi.ac.uk/vol1/fastq/ERR278/005/ERR2784505/ERR2784505_1.fastq.gz
wget -nc ftp://ftp.sra.ebi.ac.uk/vol1/fastq/ERR278/001/ERR2784401/ERR2784401_1.fastq.gz
wget -nc ftp://ftp.sra.ebi.ac.uk/vol1/fastq/ERR278/006/ERR2784316/ERR2784316_1.fastq.gz
wget -nc ftp://ftp.sra.ebi.ac.uk/vol1/fastq/ERR278/002/ERR2784652/ERR2784652_2.fastq.gz
wget -nc ftp://ftp.sra.ebi.ac.uk/vol1/fastq/ERR278/002/ERR2784512/ERR2784512_2.fastq.gz
wget -nc ftp://ftp.sra.ebi.ac.uk/vol1/fastq/ERR278/007/ERR2784427/ERR2784427_2.fastq.gz
wget -nc ftp://ftp.sra.ebi.ac.uk/vol1/fastq/ERR278/004/ERR2784384/ERR2784384_1.fastq.gz
wget -nc ftp://ftp.sra.ebi.ac.uk/vol1/fastq/ERR278/000/ERR2784540/ERR2784540_2.fastq.gz
wget -nc ftp://ftp.sra.ebi.ac.uk/vol1/fastq/ERR278/007/ERR2784417/ERR2784417_2.fastq.gz
wget -nc ftp://ftp.sra.ebi.ac.uk/vol1/fastq/ERR278/003/ERR2784533/ERR2784533_1.fastq.gz
wget -nc ftp://ftp.sra.ebi.ac.uk/vol1/fastq/ERR278/001/ERR2784411/ERR2784411_1.fastq.gz
wget -nc ftp://ftp.sra.ebi.ac.uk/vol1/fastq/ERR278/009/ERR2784539/ERR2784539_2.fastq.gz
wget -nc ftp://ftp.sra.ebi.ac.uk/vol1/fastq/ERR278/006/ERR2784496/ERR2784496_1.fastq.gz
wget -nc ftp://ftp.sra.ebi.ac.uk/vol1/fastq/ERR278/001/ERR2784571/ERR2784571_1.fastq.gz
wget -nc ftp://ftp.sra.ebi.ac.uk/vol1/fastq/ERR278/007/ERR2784617/ERR2784617_1.fastq.gz
wget -nc ftp://ftp.sra.ebi.ac.uk/vol1/fastq/ERR278/005/ERR2784465/ERR2784465_2.fastq.gz
wget -nc ftp://ftp.sra.ebi.ac.uk/vol1/fastq/ERR278/003/ERR2784333/ERR2784333_2.fastq.gz
wget -nc ftp://ftp.sra.ebi.ac.uk/vol1/fastq/ERR278/003/ERR2784343/ERR2784343_2.fastq.gz
wget -nc ftp://ftp.sra.ebi.ac.uk/vol1/fastq/ERR278/007/ERR2784607/ERR2784607_1.fastq.gz
wget -nc ftp://ftp.sra.ebi.ac.uk/vol1/fastq/ERR278/005/ERR2784455/ERR2784455_2.fastq.gz
wget -nc ftp://ftp.sra.ebi.ac.uk/vol1/fastq/ERR278/007/ERR2784577/ERR2784577_2.fastq.gz
wget -nc ftp://ftp.sra.ebi.ac.uk/vol1/fastq/ERR278/006/ERR2784326/ERR2784326_1.fastq.gz
wget -nc ftp://ftp.sra.ebi.ac.uk/vol1/fastq/ERR278/001/ERR2784581/ERR2784581_1.fastq.gz
wget -nc ftp://ftp.sra.ebi.ac.uk/vol1/fastq/ERR278/004/ERR2784624/ERR2784624_2.fastq.gz
wget -nc ftp://ftp.sra.ebi.ac.uk/vol1/fastq/ERR278/008/ERR2784448/ERR2784448_1.fastq.gz
wget -nc ftp://ftp.sra.ebi.ac.uk/vol1/fastq/ERR278/004/ERR2784374/ERR2784374_1.fastq.gz
wget -nc ftp://ftp.sra.ebi.ac.uk/vol1/fastq/ERR278/002/ERR2784502/ERR2784502_2.fastq.gz
wget -nc ftp://ftp.sra.ebi.ac.uk/vol1/fastq/ERR278/001/ERR2784381/ERR2784381_2.fastq.gz
wget -nc ftp://ftp.sra.ebi.ac.uk/vol1/fastq/ERR278/005/ERR2784655/ERR2784655_1.fastq.gz
```
