## Additional File 2: Analysis repository for "The host genotype actively shapes its microbiome across generations in laboratory mice": Benga_FiveGen.ipynb

```
# %% [markdown]
# Influence of the genetic background on murine gut microbiome composition and diversity over five generations

# %% [markdown]
The main goal of this study is to study the gut and skin microbiome of C75BL/6J and BALB/c mice, which have been naturally colonised with the same microbiome simultaneously.To achieve this the mice were born and weaned by B6CF1 foster mothers. The aim is to determine whether the genotype of the mice will have an impact on their microbiome. Therefore, the mice have been bred over 5 generations and kept in identical housing with identical breeding procedures, to minimise external effects on the microbiome. Additionally, sex and cage environment have been recorded during the experiment, to evaluate if they play a role in the composition of the microbiome as well.

This analysis aims to explore the differences and similarities of the mice microbiomes. It focuses on the calculation of alpha and beta diversities (Shannon, Faith's PD, observed OTUs, unweighted UniFrac) to carve out whether the genotype of a mouse has an impact on its gut microbiome. The alpha and beta diversities are visualised and used for statistical tests. Additionally, taxa that are significantly different between both genotypes and their respective generations are determined too by using ANCOM. Furthermore, we investigate whether the the parental microbiomes impacted their children's microbiomes by analyzing them with SourceTracker.

# %% [markdown]
## bcl2fastq and Trimming step

First, the raw sequencing data was transferred into FASTQ files using bcl2fastq (v2.19.0.316), with the parameter of `--barcode-mismatches 0` to ensure no false barcodes get assigned to the wrong sequence.

# %% [markdown]
`bcl2fastq` <br>
  `--runfolder-dir 210902_M04304_0266_000000000-JNGPR ` <br>
  `--output-dir /vol/jlab/MicrobiomeAnalyses/Projects/Benga_ThreeGen/Demux/ThirdBatch/`  <br>
  `--ignore-missing-bcls`  <br>
  `--sample-sheet /vol/jlab/MicrobiomeAnalyses/Projects/Benga_ThreeGen/Incoming/210902_M04304_0266_000000000-JNGPR.csv`  <br>
  `--loading-threads 6`  <br>
  `--processing-threads 6`  <br>
  `--writing-threads 6`  <br>
  `--barcode-mismatches 0` <br>

# %% [markdown]
Then, the adapters were trimmed using cutadapt version 2.10. Through the parameter `--minimum-length 1:1` no empty sequences will remain if only the adapter was included in the sequence.

`for forward in *R1_001.fastq.gz:` <br>
`do reverse=$(echo $forward | sed `s/R1/R2`);` <br>
`cutadapt` <br>
`-g CCTACGGGNGGCWGCAG` <br>
`-G GACTACHVGGGTATCTAATCC` <br>
`-n 2` <br>
`–minimum-length 1:1` <br>
`-o /vol/jlab/MicrobiomeAnalyses/Projects/Benga_ThreeGen/Trimming/CorrectedIDs/$forward` <br>
`-p /vol/jlab/MicrobiomeAnalyses/Projects/Benga_ThreeGen/Trimming/CorrectedIDs/$reverse $forward $reverse;` <br>
`done`

# %% [markdown]
## Adding additional columns to the metadata

# %% [code]
q_path='export PATH=/homes/arehm/miniconda3/envs/qiime2-2023.2/bin:$PATH'
work_dir='/homes/arehm/Benga_ThreeGen/Anna/Notebook'
meta_file='/homes/arehm/Benga_ThreeGen/Anna/Notebook/meta_updated.txt'
meta_file_stefan='/vol/jlab/MicrobiomeAnalyses/Projects/Benga_ThreeGen/FromQiita/13422_20230704-030402.txt'

# %% [code]
import pandas as pd
import numpy as np
from skbio.stats.distance import DistanceMatrix
from skbio.stats.distance import permanova
from skbio.tree import TreeNode
#from qiime2 import Artifact
import itertools
import pandas as pd
import seaborn as sns
import matplotlib.pyplot as plt

pd.set_option('display.max_columns', None)
pd.set_option('display.max_rows', None)

# %% [code]
%load_ext autoreload
%autoreload 2
%reload_ext autoreload
from ggmap.snippets import *
from ggmap.analyses import *
from ggmap.correlations import *
from ggmap.workflow import *

# %% [code]
meta[meta["cage_group"].isin(["CF1-1"])]["parental_cage"]

# %% [code]
meta = pd.read_csv("meta_full.txt", sep="\t", dtype=str).set_index('sample_name')

conditions = [
    meta["cage_group"] == "CF1-1",
    meta["cage_group"] == "CF1-2",
    meta["cage_group"] == "CF1-3",
    meta["cage_group"] == "CF2-1",
    meta["cage_group"] == "CF2-2",
    meta["cage_group"] == "CF2-3",
    meta["cage_group"] == "CF3-1",
    meta["cage_group"] == "CF3-2",
    meta["cage_group"] == "CF3-3",
    meta["cage_group"] == "CF4-1",
    meta["cage_group"] == "CF4-2",
    meta["cage_group"] == "CF4-3",
    meta["cage_group"] == "CF5-1",
    meta["cage_group"] == "CF5-2",
    meta["cage_group"] == "BF1-1",
    meta["cage_group"] == "BF1-2",
    meta["cage_group"] == "BF2-1",
    meta["cage_group"] == "BF2-2",
    meta["cage_group"] == "BF3-1",
    meta["cage_group"] == "BF3-2",
    meta["cage_group"] == "BF3-3",
    meta["cage_group"] == "BF4-1",
    meta["cage_group"] == "BF4-2",
    meta["cage_group"] == "BF4-3",
    meta["cage_group"] == "BF5-1",
    meta["cage_group"] == "BF5-2",
    meta["cage_group"] == "BF5-3"    
]

values = [
    "BF1-1",
    "BF1-2",
    "BF1-3",
    "BF2-1",
    "BF2-2",
    "BF2-3",
    "BF3-1",
    "BF3-2",
    "BF3-3",
    "BF4-1",
    "BF4-2",
    "BF4-3",
    "BF5-1",
    "BF5-2",
    "CF1-1",
    "CF1-2",
    "CF2-1",
    "CF2-2",
    "CF3-1",
    "CF3-2",
    "CF3-3",
    "CF4-1",
    "CF4-2",
    "CF4-3",
    "CF5-1",
    "CF5-2",
    "CF5-3"
]

meta["cage_group_new"] = np.select(conditions, values, "other")

new_meta = meta.drop(['cage_group'], axis=1)
#list(new_meta)
new_meta = new_meta.rename(columns={"cage_group_new": "cage_group"})
list(new_meta)


# %% [code]
meta = pd.read_csv(meta_file, sep="\t", dtype=str).set_index('sample_name')

meta.loc[meta.cage_id == "P1-10", 'cage_group'] = "PC-1"
meta.loc[meta.cage_id == "P1-11", 'cage_group'] = "PC-2"
meta.loc[meta.cage_id == "P1-1", 'cage_group'] = "PB-1"
meta.loc[meta.cage_id == "P1-3", 'cage_group'] = "PB-2"
meta.loc[meta.cage_id == "P1-5", 'cage_group'] = "PB-3"


indeces = []
for i in range (64,88):
    index = "13422.colon3" + str(i)
    indeces.append(index)
    index = "13422.skin3" + str(i)
    indeces.append(index)
    index = "13422.3" + str(i) + "skin"
    indeces.append(index)

conditions = [
    (meta["generation"] == "RM"),
    (meta["generation"] == "P1"),
    (meta["generation"] == "F1"),
    (meta["generation"] == "F2"),
    (meta["generation"] == "F3"),
    (meta["generation"] == "F4"),
    (meta["generation"] == "F5")
]

values = [-1, 0, 1, 2, 3, 4, 5]

meta["gen_number"] = np.select(conditions, values, -2)

conditions = [
    (meta["cage_group"] == "PB-1") | (meta["cage_group"] == "BF1-1") | (meta["cage_group"] == "BF2-1") | (meta["cage_group"] == "BF3-1") | (meta["cage_group"] == "BF4-1"),
    (meta["cage_group"] == "PB-2") | (meta["cage_group"] == "BF1-2") | (meta["cage_group"] == "BF2-2") | (meta["cage_group"] == "BF3-2") | (meta["cage_group"] == "BF4-2") | (meta["cage_group"] == "BF5-1"),
    (meta["cage_group"] == "PB-3") | (meta["cage_group"] == "BF1-3") | (meta["cage_group"] == "BF2-3") | (meta["cage_group"] == "BF3-3") | (meta["cage_group"] == "BF4-3") | (meta["cage_group"] == "BF5-2"),
    (meta["cage_group"] == "PC-1") | (meta["cage_group"] == "CF1-1") | (meta["cage_group"] == "CF2-1") | (meta["cage_group"] == "CF3-1") | (meta["cage_group"] == "CF4-1") | (meta["cage_group"] == "CF5-1"),
    (meta["cage_group"] == "PC-2") | (meta["cage_group"] == "CF1-2") | (meta["cage_group"] == "CF2-2") | (meta["cage_group"] == "CF3-2") | (meta["cage_group"] == "CF4-2") | (meta["cage_group"] == "CF5-2"),
    (meta["cage_group"] == "PC-2") | (meta["cage_group"] == "CF1-2") | (meta["cage_group"] == "CF2-2") | (meta["cage_group"] == "CF3-3") | (meta["cage_group"] == "CF4-3") | (meta["cage_group"] == "CF5-3")
]

values = ["B1", "B2", "B3", "C1", "C2", "C3"]

meta["cage_lineage"] = np.select(conditions, values, "other")

#meta_noRM = meta.loc[(meta['genotype'] != "B6CF1") & (meta["host_common_name"] == "mouse")]
meta_noRM = meta.loc[(meta["host_common_name"] == "mouse")]

meta['i_generation'] = meta['generation'].apply(lambda x: {'RM': '-1: RM',
                                    'P1': '0: P1',
                                    'F1': '1: F1',
                                    'F2': '2: F2',
                                    'F3': '3: F3',
                                    'F4': '4: F4',
                                    'F5': '5: F5',
                                   }.get(x, x))

meta['comparison'] = np.where(
    meta.index.isin(indeces), 1, 0) 

meta.to_csv("meta_updated.txt", header=True, sep="\t")

# %% [code]
duplicates = []
for i in range (64,88):
    index = "13422.skin3" + str(i)
    duplicates.append(index)
duplicate_indeces = str(duplicates)[1:-1]

# %% [code]
#filtering out duplicate skin samples from batch 2
! $q_path; qiime feature-table filter-samples \
--i-table /homes/arehm/Benga_ThreeGen/FromQiita/128106_feature-table.qza \
--m-metadata-file $meta_file \
--p-where "[sample_name] IN ($duplicate_indeces)" \
--p-exclude-ids \
--o-filtered-table $work_dir/no_duplicates_table.qza

# %% [code]
%time ! $q_path; qiime feature-table filter-features \
  --i-table $work_dir/no_duplicates_table.qza \
  --p-min-frequency 10 \
  --o-filtered-table $work_dir/filtered-10-feature-table.qza

# %% [code]
%time ! $q_path; qiime feature-table summarize \
  --i-table $work_dir/filtered-10-feature-table.qza\
  --o-visualization $work_dir/summary_filtered-10-table.qzv \
  --m-sample-metadata-file $meta_file

# %% [markdown]
Then, the sequence data from qiita after deblur was performed is imported into a QIIME2 artifact. This file is needed for the subsequent step of assigning taxonomy to the features using the Greengenes classifier provided on the QIIME2 docs page https://docs.qiime2.org/2022.2/data-resources/ . The Greengenes 13_8 99% OTUs full-length sequences (MD5: 6bbc9b3f2f9b51d663063a7979dd95f1)file was used.

# %% [code]
%time ! $q_path; qiime tools import \
  --input-path /homes/arehm/Benga_ThreeGen/FromQiita/128106_reference-hit.seqs.fa \
  --type 'FeatureData[Sequence]' \
  --output-path $work_dir/rep-seqs.qza

# %% [code]
%time ! $q_path ; qiime feature-classifier classify-sklearn \
  --i-reads $work_dir/rep-seqs.qza \
  --i-classifier $work_dir/2022.10.backbone.full-length.nb.qza \
  --o-classification $work_dir/taxonomy_full.qza

# %% [markdown]
In order to avoid features belonging to *Cyanobacteria* in the analysis, the need to be filtered out. Using the taxonomy from the previous step as a metadata file in `--m-metadata-file`.

# %% [code]
%time ! $q_path; qiime taxa filter-table \
  --i-table $work_dir/filtered-10-feature-table.qza \
  --i-taxonomy $work_dir/taxonomy_full.qza \
  --p-exclude mitochondria,chloroplast \
  --o-filtered-table $work_dir/filtered-noCyanobacteria-feature-table.qza

# %% [code]
%time ! $q_path; qiime tools import \
  --input-path /homes/arehm/Benga_ThreeGen/FromQiita/128106_insertion_tree.relabelled.tre \
  --type 'Phylogeny[Rooted]' \
  --output-path $work_dir/insertion-tree.qza

%time ! $q_path; qiime fragment-insertion filter-features \
--i-table $work_dir/filtered-noCyanobacteria-feature-table.qza \
--i-tree $work_dir/insertion-tree.qza \
--o-filtered-table $work_dir/full-filtered‐feature-table.qza \
--o-removed-table $work_dir/removed‐feature-table.qza

# %% [code]
%time ! $q_path; qiime feature-table summarize \
  --i-table $work_dir/full-filtered‐feature-table.qza\
  --o-visualization $work_dir/summary-full-filtered‐feature-table.qzv \
  --m-sample-metadata-file $meta_file

# %% [code]
%time ! $q_path; qiime tools export \
  --input-path $work_dir/full-filtered‐feature-table.qza\
  --output-path $work_dir/exported-full-filtered-feature-table

# %% [code]


# %% [markdown]
## Rarefaction

Rarefaction is important to avoid inflation of diversity. Rarefaction curves were created for all samples, as well as only gut and sebum samples.

# %% [markdown]
### Gut
Beofre following the previous steps for imloring a suitable sequencing depth, the table was filtered to only include gut samples.

# %% [code]
%time ! $q_path; qiime feature-table filter-samples \
  --i-table $work_dir/full-filtered‐feature-table.qza \
  --m-metadata-file $meta_file \
  --p-where "[host_body_product] IN ('UBERON:lumen of gut')" \
  --o-filtered-table $work_dir/full-filtered-gut-table.qza

# %% [code]
%time ! $q_path; qiime tools export \
  --input-path $work_dir/full-filtered-gut-table.qza \
  --output-path $work_dir/exported-full-filtered-gut-table

# %% [code]
%time ! $q_path; qiime feature-table summarize \
  --i-table $work_dir/full-filtered-gut-table.qza \
  --o-visualization $work_dir/summary-full-filtered-gut-table.qzv \
  --m-sample-metadata-file $meta_file

# %% [code]
%time ! $q_path; qiime diversity alpha-rarefaction \
  --i-table $work_dir/full-filtered-gut-table.qza \
  --i-phylogeny $work_dir/insertion-tree.qza \
  --p-max-depth 20000 \
  --p-min-depth 500 \
  --p-iterations 10 \
  --p-steps 50 \
  --m-metadata-file $meta_file \
  --o-visualization $work_dir/rarefaction_curves_gut.qzv

# %% [markdown]
After consulting the rarefaction plots, a sampling depth of 6000 was chosen.

# %% [code]
%time ! $q_path; qiime feature-table rarefy \
  --i-table $work_dir/full-filtered-gut-table.qza \
  --p-sampling-depth 6000 \
  --o-rarefied-table $work_dir/rarefied6000-filtered-gut-table.qza
%time ! $q_path; qiime feature-table summarize \
  --i-table $work_dir/rarefied6000-filtered-gut-table.qza \
  --o-visualization $work_dir/summary-rarefied6000-filtered-gut-table.qzv \
  --m-sample-metadata-file $meta_file

# %% [code]
%time ! $q_path; qiime tools export \
  --input-path $work_dir/rarefied6000-filtered-gut-table.qza \
  --output-path $work_dir/exported-rarefied6000-filtered-gut-table

# %% [markdown]
### Sebum
Analogous to the gut samples.

# %% [code]
%time ! $q_path; qiime feature-table filter-samples \
  --i-table $work_dir/full-filtered‐feature-table.qza \
  --m-metadata-file $meta_file \
  --p-where "[host_body_product] IN ('UBERON:sebum')" \
  --o-filtered-table $work_dir/full-filtered-sebum-table.qza

# %% [code]
%time ! $q_path; qiime tools export \
  --input-path $work_dir/full-filtered-sebum-table.qza \
  --output-path $work_dir/exported-full-filtered-sebum-table

# %% [code]
%time ! $q_path; qiime feature-table summarize \
  --i-table $work_dir/full-filtered-sebum-table.qza \
  --o-visualization $work_dir/summary-full-filtered-sebum-table.qzv \
  --m-sample-metadata-file $meta_file

# %% [code]
%time ! $q_path; qiime diversity alpha-rarefaction \
  --i-table $work_dir/full-filtered-sebum-table.qza \
  --i-phylogeny $work_dir/insertion-tree.qza \
  --p-max-depth 4000 \
  --p-min-depth 500 \
  --p-iterations 10 \
  --p-steps 50 \
  --m-metadata-file $meta_file \
  --o-visualization $work_dir/rarefaction_curves_sebum.qzv

# %% [code]
%time ! $q_path; qiime feature-table rarefy \
  --i-table $work_dir/full-filtered-sebum-table.qza \
  --p-sampling-depth 1000 \
  --o-rarefied-table $work_dir/rarefied-filtered-sebum-table.qza

# %% [code]
%time ! $q_path; qiime feature-table summarize \
  --i-table $work_dir/rarefied-filtered-sebum-table.qza \
  --o-visualization $work_dir/summary-rarefied-filtered-sebum-table.qzv \
  --m-sample-metadata-file $meta_file

# %% [code]
%time ! $q_path; qiime tools export \
  --input-path $work_dir/rarefied-filtered-sebum-table.qza \
  --output-path $work_dir/exported-rarified-filtered-sebum-table

# %% [markdown]
## Taxonomy

# %% [markdown]
### Taxonomy Barplots

# %% [code]
%time ! $q_path; qiime taxa barplot \
  --i-table $work_dir/full-filtered‐feature-table.qza \
  --i-taxonomy $work_dir/taxonomy_full.qza \
  --m-metadata-file $meta_file \
  --o-visualization $work_dir/taxa_barplot_full.qzv

# %% [code]
%time ! $q_path; qiime tools export \
  --input-path $work_dir/taxonomy_full.qza \
  --output-path $work_dir/exported-taxonomy-full

# %% [code]
exported_taxonomy = pd.read_csv("exported-taxonomy-full/taxonomy.tsv", sep="\t").set_index("Feature ID")

# %% [code]
exported_taxonomy[['Taxon1', 'Taxon2']] = exported_taxonomy["Taxon"].apply(lambda x: pd.Series(str(x).split(" c__")))
phylum_taxa= exported_taxonomy.drop(columns=["Taxon2", "Taxon"])
phylum_taxa.rename(columns={'Taxon1':'Taxon'}, inplace=True)
phylum_taxa = phylum_taxa[["Taxon", "Confidence"]]
phylum_taxa = phylum_taxa.drop(phylum_taxa[phylum_taxa.Taxon == "k__Bacteria"].index)
phylum_taxa.to_csv("exported-taxonomy-full/phylum.tsv", sep="\t")

# %% [code]
%matplotlib inline
fig = plotTaxonomy("exported-fragment-filtered-feature-table/feature-table.biom", meta[meta['host_body_product'].isin(['UBERON:lumen of gut', 'UBERON:sebum']) & meta['host_scientific_name'].isin(['Mus musculus']) & meta['genotype'].isin(['BALB/c', 'C57BL6/J'])],  file_taxonomy="exported-taxonomy-full/phylum.tsv", fct_aggregate=np.mean, group_l0="host_body_product",group_l1="genotype", rank="Phylum", minreadnr=100)
plt.show()

# %% [code]
exported_taxonomy[['Taxon1', 'Taxon2']] = exported_taxonomy["Taxon"].apply(lambda x: pd.Series(str(x).split(" g__")))
family_taxa= exported_taxonomy.drop(columns=["Taxon2", "Taxon"])
family_taxa.rename(columns={'Taxon1':'Taxon'}, inplace=True)
family_taxa = family_taxa[["Taxon", "Confidence"]]
family_taxa = family_taxa[family_taxa["Taxon"].str.contains("f__")]
family_taxa.to_csv("exported-taxonomy-full/family.tsv", sep="\t")

# %% [code]
%matplotlib inline
fig, rank_counts, graphinfo, vals, colors = plotTaxonomy("exported-fragment-filtered-feature-table/feature-table.biom", meta[meta['host_body_product'].isin(['UBERON:lumen of gut', 'UBERON:sebum']) & meta['host_scientific_name'].isin(['Mus musculus']) & meta['genotype'].isin(['BALB/c', 'C57BL6/J'])],  file_taxonomy="exported-taxonomy-full/family.tsv", fct_aggregate=np.mean, group_l0="host_body_product",group_l1="genotype", rank="Family", minreadnr=5000)

# %% [markdown]
### Scatterplots

# %% [code]
taxa_all_l6 = pd.read_csv("/homes/arehm/Benga_ThreeGen/Anna/Notebook_Final/exported-taxonomy-full/level-6.csv", sep=",").set_index('index')

# %% [code]
taxa_all_l6['d__Bacteria;p__Verrucomicrobiota;c__Verrucomicrobiae;o__Verrucomicrobiales;f__Akkermansiaceae;g__Akkermansia']

# %% [code]
#use level6 table as Akkermansia is a genus
taxa_gut = taxa_all_l6[taxa_all_l6['host_body_product'].isin(['UBERON:lumen of gut']) & taxa_all_l6['host_scientific_name'].isin(['Mus musculus']) & ~taxa_all_l6['generation'].isin(['not applicable'])]
taxa_gut['d__Bacteria;p__Verrucomicrobiota;c__Verrucomicrobiae;o__Verrucomicrobiales;f__Akkermansiaceae;g__Akkermansia'].astype(float)
total_abundance_akkermansia = taxa_gut['d__Bacteria;p__Verrucomicrobiota;c__Verrucomicrobiae;o__Verrucomicrobiales;f__Akkermansiaceae;g__Akkermansia'].sum()
taxa_gut['relative_abundance'] = taxa_gut['d__Bacteria;p__Verrucomicrobiota;c__Verrucomicrobiae;o__Verrucomicrobiales;f__Akkermansiaceae;g__Akkermansia'] / total_abundance_akkermansia

# %% [code]
#goal: scatterplot with genotype on x axis and relative abundance of akkermansia on the y axis
%matplotlib inline
ax = sns.stripplot(x= 'generation', y='relative_abundance', data= taxa_gut, hue='genotype', order=['RM','P1', 'F1', 'F2', 'F3', 'F4', 'F5'], hue_order=['C57BL6/J', 'BALB/c', 'B6CF1'],dodge=True)
plt.savefig(work_dir + "/scatter_akkermansia.png", transparent=False, bbox_inches='tight')
plt.show()

# %% [markdown]
### ANCOM

# %% [code]
%time !$q_path; qiime feature-table filter-samples \
  --i-table $work_dir/rarefied6000-filtered-gut-table.qza \
  --m-metadata-file $meta_file \
  --p-where "[host_common_name] IN ('mouse') AND [genotype] NOT IN ('B6CF1') AND [comparison] IN ('0')" \
  --o-filtered-table $work_dir/rarified-gut-nocompRM.qza

# %% [code]
%time !$q_path; qiime taxa collapse \
  --i-table $work_dir/rarified-gut-nocompRM.qza \
  --i-taxonomy $work_dir/taxonomy_full.qza \
  --p-level 7 \
  --o-collapsed-table $work_dir/collapsed-level7-mouse-gut-table-nocompRM.qza

# %% [code]
%time !$q_path; qiime composition add-pseudocount \
  --i-table $work_dir/collapsed-level7-mouse-gut-table-nocompRM.qza \
  --o-composition-table $work_dir/composition-table-mouse-gut-l7-nocompRM.qza

# %% [code]
%time !$q_path; qiime composition ancom \
  --i-table $work_dir/composition-table-mouse-gut-l7-nocompRM.qza \
  --m-metadata-file $meta_file \
  --m-metadata-column genotype \
  --o-visualization $work_dir/ancom-genotype-mouse-gut-l7-nocompRM.qzv

# %% [code]
!unzip $work_dir/composition-table-mouse-gut-l7-nocompRM.qza

# %% [code]
! $q_path; biom convert \
  --input-fp f3a6a978-a527-4588-a92d-2445cf953a0b/data/feature-table.biom       \
  --to-tsv \
  --output-fp $work_dir/ancom-biom-table-gut-l7

# %% [code]
!$q_path; biom summarize-table -i f3a6a978-a527-4588-a92d-2445cf953a0b/data/feature-table.biom

# %% [code]
ancom_biom_table_gut_l7 = pd.read_csv(work_dir+"/ancom-biom-table-gut-l7", 
            sep="\t",skiprows=1, index_col=0)

# %% [code]
%matplotlib inline

relative_abundance_gut= ancom_biom_table_gut_l7.div(6207)
list_C57B = meta[meta['genotype'].isin(['C57BL6/J'])].index
list_BALB = meta[meta['genotype'].isin(['BALB/c'])].index

ancom_C57B_gut = relative_abundance_gut[[col for col in relative_abundance_gut.columns if col in list_C57B]]
ancom_C57B_gut["average"]=ancom_C57B_gut.mean(axis=1)

ancom_BALB_gut = relative_abundance_gut[[col for col in relative_abundance_gut.columns if col in list_BALB]]
ancom_BALB_gut["average"] = ancom_BALB_gut.mean(axis=1)


with open('ancom_l6.txt', 'r') as f:
    ancom_taxa_gut_l6 = [line.rstrip() for line in f]
with open('ancom_l7.txt', 'r') as f:
    ancom_taxa_gut_l7 = [line.rstrip() for line in f]
    
taxa_C57B_gut = ancom_C57B_gut[ancom_C57B_gut.index.isin(ancom_taxa_gut_l7)]
taxa_BALB_gut = ancom_BALB_gut[ancom_BALB_gut.index.isin(ancom_taxa_gut_l7)]

ancom_plot = pd.DataFrame({'C57BL6/J': taxa_C57B_gut['average'],
                           'BALB/c': taxa_BALB_gut['average']},
                         index=taxa_C57B_gut.index)

# %% [markdown]
prepare lists wit indeces 

# %% [code]
import matplotlib.pyplot as plt
ancom_taxa_plot = ancom_plot.plot.barh(figsize=(20,15), width =0.8, fontsize=20).legend(loc=1, prop={'size':30})
plt.legend(labels=["C57BL/6J","BALB/c"], fontsize=30)
plt.xlim(right=0.1)
ancom_taxa_plot.figure.savefig(work_dir + '/ancom_plot_gut_species.svg', set_facecolor='white', bbox_inches='tight')

# %% [code]


# %% [markdown]
## Alpha Diversity

# %% [markdown]
### Gut

# %% [code]
filtered_gut_table = biom2pandas(work_dir + "/exported-full-filtered-gut-table/feature-table.biom").fillna(0)
alpha_div_gut = alpha_diversity(filtered_gut_table, 6000, metrics=["shannon", "observed_features", "PD_whole_tree"], num_iterations=10, reference_tree="/homes/arehm/Benga_ThreeGen/FromQiita/128106_insertion_tree.relabelled.tre", use_grid=True, dry=False)
alpha_results_merged_gut = alpha_div_gut['results'].merge(meta, left_index=True, right_index=True)
alpha_results_merged_gut[alpha_results_merged_gut['genotype'].isin(['C57BL6/J', 'BALB/c'])]

# %% [code]
RM_shannon_average = np.mean(alpha_results_merged_gut[alpha_results_merged_gut['generation'].isin(['RM'])]['shannon'])
RM_shannon_std = np.std(alpha_results_merged_gut[alpha_results_merged_gut['generation'].isin(['RM'])]['shannon'])
RM_PD_average = np.mean(alpha_results_merged_gut[alpha_results_merged_gut['generation'].isin(['RM'])]['PD_whole_tree'])
RM_PD_std = np.std(alpha_results_merged_gut[alpha_results_merged_gut['generation'].isin(['RM'])]['PD_whole_tree'])
RM_observed_average = np.mean(alpha_results_merged_gut[alpha_results_merged_gut['generation'].isin(['RM'])]['observed_features'])
RM_observed_std = np.std(alpha_results_merged_gut[alpha_results_merged_gut['generation'].isin(['RM'])]['observed_features'])

# %% [code]
meta[meta['genotype'].isin(['C57BL6/J','BALB/c', 'B6CF1']) & meta['host_common_name'].isin(['mouse']) & ~meta['generation'].isin(['not applicable'])].value_counts('i_generation')

# %% [code]
%matplotlib inline
import pylab
metric = 'shannon'
fig, axes = plt.subplots(1,1, figsize=(10,5))
a_data = plot_timecourse(meta[meta['genotype'].isin(['C57BL6/J','BALB/c', 'B6CF1']) & meta['host_common_name'].isin(['mouse']) & ~meta['generation'].isin(['not applicable']) & meta['comparison'].isin([0])],
                data=alpha_div_gut['results'][metric],
                col_events='i_generation', col_entities=None,
                cols_groups=['host_common_name', 'genotype'],
                #test_groups=[(('mouse', 'C57BL6/J'), ('mouse', 'BALB/c'))],
                pthreshold=0.05, print_samplesizes=True,
                ax=axes,
                colors_groups={('mouse', 'BALB/c'): "tab:orange", ('mouse', 'C57BL6/J'): "tab:blue"},
                test_groups=None
                )
#axes.legend(labels=['BALB/c', 'C57BL/6J'])
#plt.scatter("-1: RM", RM_shannon_average, color='green')
#plt.errorbar(["-1: RM"], [RM_shannon_average], yerr=RM_shannon_std, color='green')
#handles, labels = fig.get_legend_handles_labels()
labels = ['BALB/c', 'C57BL/6J', 'B6CF1']
#plt.legend(labels, loc=2)
plt.title('Darm')
plt.savefig(work_dir + "/alpha_gut_timeline_shannon.png", transparent=True, bbox_inches="tight", facecolor='white')

# %% [code]
%matplotlib inline

metric = 'observed_features'
fig, axes = plt.subplots(1,1, figsize=(10,5))
a_data = plot_timecourse(meta[meta['genotype'].isin(['C57BL6/J','BALB/c', 'B6CF1']) & meta['host_common_name'].isin(['mouse']) & ~meta['generation'].isin(['not applicable']) ],
                data=alpha_div_gut['results'][metric],
                col_events='i_generation', col_entities=None,
                cols_groups=['host_common_name', 'genotype'],
                #test_groups=[(('mouse', 'C57BL6/J'), ('mouse', 'BALB/c'))],
                test_groups=None,
                pthreshold=0.05, print_samplesizes=True,
                ax=axes,
                colors_groups={('mouse', 'BALB/c'): "tab:orange", ('mouse', 'C57BL6/J'): "tab:blue"})
#axes.legend(labels=['BALB/c', 'C57BL/6J'])
plt.scatter("-1: RM", RM_observed_average, color='green')
plt.errorbar(["-1: RM"], [RM_observed_average], yerr=RM_observed_std, color='green')
#handles, labels = fig.get_legend_handles_labels()
labels = ['BALB/c', 'C57BL/6J', 'B6CF1']
#plt.legend(handles, labels, loc=2)
plt.title('Darm')
plt.savefig(work_dir + "/alpha_gut_timeline_observed_features.png", transparent=False, bbox_inches="tight", facecolor='white')

# %% [code]
%matplotlib inline

metric = 'PD_whole_tree'
fig, axes = plt.subplots(1,1, figsize=(10,5))
a_data = plot_timecourse(meta[meta['genotype'].isin(['C57BL6/J','BALB/c', 'B6CF1']) & meta['host_common_name'].isin(['mouse']) & ~meta['generation'].isin(['not applicable'])],
                data=alpha_div_gut['results'][metric],
                col_events='i_generation', col_entities=None,
                cols_groups=['host_common_name', 'genotype'],
                #test_groups=[(('mouse', 'C57BL6/J'), ('mouse', 'BALB/c'))],
                test_groups=None,
                pthreshold=0.05, print_samplesizes=True,
                ax=axes,
                colors_groups={('mouse', 'BALB/c'): "tab:orange", ('mouse', 'C57BL6/J'): "tab:blue"})
#axes.legend(labels=['BALB/c', 'C57BL/6J'])
plt.scatter("-1: RM", RM_PD_average, color='green')
plt.errorbar(["-1: RM"], [RM_PD_average], yerr=RM_PD_std, color='green')
#handles, labels = fig.get_legend_handles_labels()
labels = ['BALB/c', 'C57BL/6J', 'B6CF1']
#plt.legend(handles, labels, loc=2)
plt.savefig(work_dir + "/alpha_gut_timeline_PD_whole_tree.png", transparent=False, bbox_inches="tight", facecolor='white')

# %% [markdown]
#### Scatterplots
##### Birth Offset

# %% [code]
meta_stefan = pd.read_csv(meta_file_stefan, sep="\t", dtype=str).set_index('sample_name')
meta_stefan.head()

# %% [code]
gut_indeces = meta_stefan[meta_stefan['genotype'].isin(['c57bl6/j']) & meta_stefan['host_body_product'].isin(['uberon:lumen of gut']) & ~meta_stefan['generation'].isin(['not applicable']) & ~meta_stefan['tube_id'].isin(['3-a8'])].index
gut_indeces
#gut_indeces = gut_indeces.delete('13422.217')

# %% [code]
alpha_div_gut['results']['PD_whole_tree'].loc[gut_indeces]

# %% [code]
alpha_results_merged_gut_stefan = alpha_div_gut['results'].merge(meta_stefan, left_index=True, right_index=True)
alpha_results_merged_gut_stefan[alpha_results_merged_gut_stefan['genotype'].isin(['c57bl6/j', 'balb/c'])]

# %% [code]
gut_indeces_C57B = meta_stefan[meta_stefan['genotype'].isin(['c57bl6/j']) & meta_stefan['host_body_product'].isin(['uberon:lumen of gut']) & ~meta_stefan['generation'].isin(['not applicable'])].index
data_gut_C57B_alpha = alpha_results_merged_gut.loc[alpha_results_merged_gut.index.intersection(gut_indeces_C57B)]
gut_indeces_BALB = meta_stefan[meta_stefan['genotype'].isin(['balb/c']) & meta_stefan['host_body_product'].isin(['uberon:lumen of gut']) & ~meta_stefan['generation'].isin(['not applicable'])].index
data_gut_BALB_alpha = alpha_results_merged_gut.loc[alpha_results_merged_gut.index.intersection(gut_indeces_BALB)]

# %% [code]
list(meta_stefan.columns)

# %% [code]
%matplotlib inline
import seaborn as sns
sns.stripplot(x= 'birth_offset_in_days', y='PD_whole_tree', data= alpha_results_merged_gut_stefan, hue='cage_line').set(title='Alpha Diversity per cage lineage by generation B6J')
#lt.show()
plt.savefig(work_dir + "/scatter_birth_offset.png", transparent=False, bbox_inches='tight')

# %% [code]
birth_offset = meta_stefan[(meta_stefan['sample_type'] == 'colon content') & 
                                    (meta_stefan['host_scientific_name'] != 'callithrix jacchus') & 
                                    (meta_stefan['iacuc_institute'] == 'LANUV')].groupby(['genotype', 'generation', 'cage_line'])['birth_timestamp'].unique().apply(lambda x: sorted(list(map(lambda d: datetime.strptime(d, '%Y-%m-%d'), x)))).apply(lambda dates: (dates[-1] - dates[0]).days).reset_index()
birth_offset = birth_offset.rename(columns={'birth_timestamp': 'birth_offset_in_days'})
if 'birth_offset_in_days' not in meta_stefan.columns:
    meta_stefan = meta_stefan.reset_index().merge(birth_offset, left_on=['genotype','generation','cage_line'], right_on=['genotype','generation','cage_line'], how='left').set_index('sample_name')
birth_offset.head()

# %% [code]
meta_stefan["birth_offset_average_generation"] = meta_stefan.groupby(['generation'])['birth_offset_in_days'].transform('mean')


# %% [code]
sns.barplot(data=birth_offset, orient='horizontal', x='birth_offset_in_days', y='cage_line')

# %% [markdown]
##### Animals per Cage

# %% [code]
stress = meta_stefan[(meta_stefan['sample_type'] == 'colon content') & 
                              (meta_stefan['host_scientific_name'] != 'callithrix jacchus') & 
                              (meta_stefan['iacuc_institute'] == 'LANUV')].groupby(['genotype', 'generation', 'cage_line', 'cage_id', 'sex']).size().reset_index()
stress = stress.rename(columns={0: 'animals_per_cage'})
if 'animals_per_cage' not in meta_stefan:
    meta_stefan = meta_stefan.reset_index().merge(stress, left_on=['genotype','generation','cage_line','cage_id','sex'], right_on=['genotype','generation','cage_line','cage_id','sex'], how='left').set_index('sample_name')
print(stress.shape[0])
stress.head()

# %% [code]
alpha_results_merged_gut_stefan = alpha_div_gut['results'].merge(meta_stefan, left_index=True, right_index=True)
alpha_results_merged_gut_stefan[alpha_results_merged_gut_stefan['genotype'].isin(['c57bl6/j', 'balb/c'])]

# %% [code]
gut_indeces_C57B = meta_stefan[meta_stefan['genotype'].isin(['c57bl6/j']) & meta_stefan['host_body_product'].isin(['uberon:lumen of gut']) & ~meta_stefan['generation'].isin(['not applicable'])].index
data_gut_C57B_alpha = alpha_results_merged_gut.loc[alpha_results_merged_gut.index.intersection(gut_indeces_C57B)]
gut_indeces_BALB = meta_stefan[meta_stefan['genotype'].isin(['balb/c']) & meta_stefan['host_body_product'].isin(['uberon:lumen of gut']) & ~meta_stefan['generation'].isin(['not applicable'])].index
data_gut_BALB_alpha = alpha_results_merged_gut.loc[alpha_results_merged_gut.index.intersection(gut_indeces_BALB)]

# %% [code]
%matplotlib inline
import seaborn as sns
sns.stripplot(x= 'animals_per_cage', y='PD_whole_tree', data= alpha_results_merged_gut_stefan, hue='cage_line').set(title='Alpha Diversity per cage lineage by generation B6J')
#lt.show()
plt.savefig(work_dir + "/scatter_animals_per_cage.png", transparent=False, bbox_inches='tight')

# %% [markdown]
### Sebum

# %% [code]
filtered_sebum_table = biom2pandas(work_dir + "/exported-full-filtered-sebum-table/feature-table.biom").fillna(0)
alpha_div_sebum = alpha_diversity(filtered_sebum_table, 1000, metrics=["shannon", "observed_features", "PD_whole_tree"], num_iterations=10, reference_tree="/homes/arehm/Benga_ThreeGen/FromQiita/128106_insertion_tree.relabelled.tre", use_grid=True, dry=False)
alpha_results_merged_sebum = alpha_div_sebum['results'].merge(meta, left_index=True, right_index=True)
alpha_results_merged_sebum[alpha_results_merged_sebum['genotype'].isin(['C57BL6/J', 'BALB/c'])]

# %% [code]
RM_shannon_average = np.mean(alpha_results_merged_sebum[alpha_results_merged_sebum['generation'].isin(['RM'])]['shannon'])
RM_shannon_std = np.std(alpha_results_merged_sebum[alpha_results_merged_sebum['generation'].isin(['RM'])]['shannon'])
RM_PD_average = np.mean(alpha_results_merged_sebum[alpha_results_merged_sebum['generation'].isin(['RM'])]['PD_whole_tree'])
RM_PD_std = np.std(alpha_results_merged_sebum[alpha_results_merged_sebum['generation'].isin(['RM'])]['PD_whole_tree'])
RM_observed_average = np.mean(alpha_results_merged_sebum[alpha_results_merged_sebum['generation'].isin(['RM'])]['observed_features'])
RM_observed_std = np.std(alpha_results_merged_sebum[alpha_results_merged_sebum['generation'].isin(['RM'])]['observed_features'])

# %% [code]
%matplotlib inline

metric = 'shannon'
fig, axes = plt.subplots(1,1, figsize=(10,5))
a_data = plot_timecourse(meta[meta['genotype'].isin(['C57BL6/J','BALB/c', 'B6CF1']) & meta['host_common_name'].isin(['mouse']) & ~meta['generation'].isin(['not applicable'])],
                data=alpha_div_sebum['results'][metric],
                col_events='i_generation', col_entities=None,
                cols_groups=['host_common_name', 'genotype'],
                #test_groups=[(('mouse', 'C57BL6/J'), ('mouse', 'BALB/c'))],
                test_groups=None,
                pthreshold=0.05, print_samplesizes=True,
                ax=axes,
                colors_groups={('mouse', 'BALB/c'): "tab:orange", ('mouse', 'C57BL6/J'): "tab:blue"})
plt.scatter("-1: RM", RM_shannon_average, color='green')
plt.errorbar(["-1: RM"], [RM_shannon_average], yerr=RM_shannon_std, color='green')
#handles, labels = fig.get_legend_handles_labels()
labels = ['BALB/c', 'C57BL/6J', 'B6CF1']
#plt.legend(handles, labels, loc=2)
#axes.legend(labels=['BALB/c', 'C57BL/6J', 'B6CF1'])
plt.savefig(work_dir + "/alpha_sebum_timeline_shannon.png", transparent=False, bbox_inches="tight", facecolor='white')

# %% [code]
%matplotlib inline

metric = 'observed_features'
fig, axes = plt.subplots(1,1, figsize=(10,5))
a_data = plot_timecourse(meta[meta['genotype'].isin(['C57BL6/J','BALB/c', 'B6CF1']) & meta['host_common_name'].isin(['mouse']) & ~meta['generation'].isin(['not applicable'])],
                data=alpha_div_sebum['results'][metric],
                col_events='i_generation', col_entities=None,
                cols_groups=['host_common_name', 'genotype'],
                #test_groups=[(('mouse', 'C57BL6/J'), ('mouse', 'BALB/c'))],
                test_groups=None,
                pthreshold=0.05, print_samplesizes=True,
                ax=axes,
                colors_groups={('mouse', 'BALB/c'): "tab:orange", ('mouse', 'C57BL6/J'): "tab:blue"})
plt.scatter("-1: RM", RM_observed_average, color='green')
plt.errorbar(["-1: RM"], [RM_observed_average], yerr=RM_observed_std, color='green')
#handles, labels = fig.get_legend_handles_labels()
labels = ['BALB/c', 'C57BL/6J', 'B6CF1']
#plt.legend(handles, labels, loc=2)
plt.title('Haut')
plt.savefig(work_dir + "/alpha_sebum_timeline_observed_features.png", transparent=False, bbox_inches="tight", facecolor='white')

# %% [code]
%matplotlib inline

metric = 'PD_whole_tree'
fig, axes = plt.subplots(1,1, figsize=(10,5))
a_data = plot_timecourse(meta[meta['genotype'].isin(['C57BL6/J','BALB/c', 'B6CF1']) & meta['host_common_name'].isin(['mouse']) & ~meta['generation'].isin(['not applicable'])],
                data=alpha_div_sebum['results'][metric],
                col_events='i_generation', col_entities=None,
                cols_groups=['host_common_name', 'genotype'],
                #test_groups=[(('mouse', 'C57BL6/J'), ('mouse', 'BALB/c'))],
                test_groups=None,
                pthreshold=0.05, print_samplesizes=True,
                ax=axes,
                colors_groups={('mouse', 'BALB/c'): "tab:orange", ('mouse', 'C57BL6/J'): "tab:blue"})
plt.scatter("-1: RM", RM_PD_average, color='green')
plt.errorbar(["-1: RM"], [RM_PD_average], yerr=RM_PD_std, color='green')
#handles, labels = fig.get_legend_handles_labels()
labels = ['BALB/c', 'C57BL/6J', 'B6CF1']
#plt.legend(handles, labels, loc=2)
#axes.legend(labels=['BALB/c', 'C57BL/6J'])
plt.savefig(work_dir + "/alpha_sebum_timeline_PD_whole_tree.png", transparent=False, bbox_inches="tight", facecolor='white')

# %% [markdown]
## Beta Diversity

# %% [markdown]
### Gut

# %% [markdown]
#### All samples

# %% [code]
rarified_gut_table = biom2pandas(work_dir + "/exported-rarefied6000-filtered-gut-table/feature-table.biom").fillna(0)

# %% [code]
res_beta_gut = beta_diversity(rarified_gut_table, metrics=["unweighted_unifrac","weighted_unifrac","bray_curtis"], reference_tree="/homes/arehm/Benga_ThreeGen/FromQiita/128106_insertion_tree.relabelled.tre", use_grid=True, dry=False)

# %% [code]
emperor(meta, res_beta_gut['results'], "Beta_Diversity", other_beta_diversities=None, infix="gut", run_tsne_umap=False, ppn=1, use_grid=True, dry=False)

# %% [code]
meta.loc[['13422.217']]

# %% [markdown]
#### Only Mice with Comparison Mice

# %% [code]
meta_rarified = meta.loc[list(rarified_gut_table.columns.values)]
condition = list(meta_rarified[meta_rarified['host_common_name'].isin(['mouse']) & meta_rarified['host_body_product'].isin(['UBERON:lumen of gut'])].index.values)
mice_with_comp = rarified_gut_table[condition]

# %% [code]
res_beta_gut_with_comp = beta_diversity(mice_with_comp, metrics=["unweighted_unifrac","weighted_unifrac","bray_curtis"], reference_tree="/homes/arehm/Benga_ThreeGen/FromQiita/128106_insertion_tree.relabelled.tre", use_grid=True, dry=False)

# %% [code]
emperor(meta, res_beta_gut_with_comp['results'], "Beta_Diversity", other_beta_diversities=None, infix="_gut_with_comp", run_tsne_umap=False, ppn=1, use_grid=True, dry=False)

# %% [code]


# %% [markdown]
#### Only Mice without Comparison

# %% [code]
meta_rarified = meta.loc[list(rarified_gut_table.columns.values)]
condition = list(meta_rarified[meta_rarified['host_common_name'].isin(['mouse']) & meta_rarified['host_body_product'].isin(['UBERON:lumen of gut']) & meta_rarified['comparison'].isin([0])].index.values)
mice_no_comp_gut = rarified_gut_table[condition]

# %% [code]
res_beta_gut_no_comp = beta_diversity(mice_no_comp_gut, metrics=["unweighted_unifrac","weighted_unifrac","bray_curtis"], reference_tree="/homes/arehm/Benga_ThreeGen/FromQiita/128106_insertion_tree.relabelled.tre", use_grid=True, dry=False)

# %% [code]
emperor(meta, res_beta_gut_no_comp['results'], "Beta_Diversity", other_beta_diversities=None, infix="_gut_no_comp", run_tsne_umap=False, ppn=1, use_grid=True, dry=False)

# %% [markdown]
### Sebum

# %% [markdown]
#### All Samples

# %% [code]
rarified_sebum_table = biom2pandas(work_dir + "/exported-rarified-filtered-sebum-table/feature-table.biom").fillna(0)

# %% [code]
res_beta_sebum = beta_diversity(rarified_sebum_table, metrics=["unweighted_unifrac","weighted_unifrac","bray_curtis"], reference_tree="/homes/arehm/Benga_ThreeGen/FromQiita/128106_insertion_tree.relabelled.tre", use_grid=True, dry=False)

# %% [code]
emperor(meta, res_beta_sebum['results'], "Beta_Diversity", other_beta_diversities=None, infix="_sebum", run_tsne_umap=False, ppn=1, use_grid=True, dry=False)

# %% [markdown]
#### Only Mice with Comparison Mice

# %% [code]
meta_rarified = meta.loc[list(rarified_sebum_table.columns.values)]
condition = list(meta_rarified[meta_rarified['host_common_name'].isin(['mouse']) & meta_rarified['host_body_product'].isin(['UBERON:sebum'])].index.values)
mice_with_comp_sebum = rarified_sebum_table[condition]

# %% [code]
res_beta_sebum_with_comp = beta_diversity(mice_with_comp_sebum, metrics=["unweighted_unifrac","weighted_unifrac","bray_curtis"], reference_tree="/homes/arehm/Benga_ThreeGen/FromQiita/128106_insertion_tree.relabelled.tre", use_grid=True, dry=False)

# %% [code]
emperor(meta, res_beta_sebum_with_comp['results'], "Beta_Diversity", other_beta_diversities=None, infix="_sebum_with_comp", run_tsne_umap=False, ppn=1, use_grid=True, dry=False)

# %% [code]


# %% [markdown]
#### Only Mice without Comparison

# %% [code]
meta_rarified = meta.loc[list(rarified_sebum_table.columns.values)]
condition = list(meta_rarified[meta_rarified['host_common_name'].isin(['mouse']) & meta_rarified['host_body_product'].isin(['UBERON:sebum']) & meta_rarified['comparison'].isin([0])].index.values)
mice_no_comp_sebum = rarified_sebum_table[condition]

# %% [code]
res_beta_sebum_no_comp = beta_diversity(mice_no_comp_sebum, metrics=["unweighted_unifrac","weighted_unifrac","bray_curtis"], reference_tree="/homes/arehm/Benga_ThreeGen/FromQiita/128106_insertion_tree.relabelled.tre", use_grid=True, dry=False)

# %% [code]
emperor(meta, res_beta_sebum_no_comp['results'], "Beta_Diversity", other_beta_diversities=None, infix="_sebum_no_comp", run_tsne_umap=False, ppn=1, use_grid=True, dry=False)

# %% [code]


# %% [markdown]
## Upsetplot

# %% [code]
import upsetplot
from upsetplot import UpSet
from upsetplot import from_memberships
from upsetplot import plot
from matplotlib import cm
import matplotlib.pyplot as plt

# %% [code]
def make_UpSet(category, upset_taxa_gut, plotname):
    position_category_column = upset_taxa_gut.columns.get_loc(category)
    up_to_gen = upset_taxa_gut.drop(upset_taxa_gut.iloc[:, (position_category_column+1):335],axis = 1)
    only_gen = up_to_gen.drop(up_to_gen.iloc[:, 277:position_category_column], axis = 1)
    taxa_sum = only_gen.groupby(category).sum()
    taxa_sum = taxa_sum.loc[:,~taxa_sum.columns.str.endswith("__")]
    taxa_sum = taxa_sum.loc[:, (taxa_sum != 0).any(axis=0)]
    taxa_sum = taxa_sum.transpose()
    over_zero = taxa_sum > 0
    over_zero = over_zero.rename(columns=lambda x: x + ' ')
    upset_combined = pd.concat([taxa_sum, over_zero],axis=1)
    upset_combined['taxa1'] = upset_combined.index
    upset_combined['taxa'] = upset_combined['taxa1'].str.split(';', expand=True)[5]
    upset_combined = upset_combined.drop(["taxa1"], axis=1)
    upset_combined = upset_combined.set_index(list(over_zero.columns))
    #upset = UpSet(upset_combined, subset_size='count', intersection_plot_elements=0, show_counts=True)
    plot(upset_combined, show_counts=True, subset_size="count", sort_categories_by="-input")
    #upset.add_stacked_bars(by="taxa")
    #upset.add_stacked_bars(by="taxa", elements=10, colors=cm.gist_stern)
    #upset.plot()
    #plt.legend(bbox_to_anchor=(1.04, 0.5), loc="center left", borderaxespad=0, title=f'Colours for taxa')
    plt.suptitle(f'UpSet plot for Genus divided by {category}')
    #plt.style.use("ggplot")
    plt.savefig(work_dir + f"/{plotname}.svg", transparent=False, bbox_inches='tight')
    plt.show()

# %% [markdown]
### UpSet Plots für Genotypen

# %% [code]
#prepare taxa_all_l6 table
upset_taxa_gut= taxa_all_l6[taxa_all_l6['host_body_product'].isin(['UBERON:lumen of gut']) & taxa_all_l6['host_scientific_name'].isin(['Mus musculus']) & ~taxa_all_l6['generation'].isin(['not applicable'])]

# %% [code]
make_UpSet("genotype", upset_taxa_gut, "upset_gut_genotype")

# %% [markdown]
#### BALB/c

# %% [code]
upset_taxa_gut_BALB= taxa_all_l6[taxa_all_l6['host_body_product'].isin(['UBERON:lumen of gut']) & taxa_all_l6['genotype'].isin(['BALB/c', 'B6CF1'])& taxa_all_l6['host_scientific_name'].isin(['Mus musculus']) & ~taxa_all_l6['generation'].isin(['not applicable'])]

# %% [code]
make_UpSet('generation', upset_taxa_gut_BALB, "upset_gut_BALB_generation")

# %% [markdown]
##### Cage Lineage

# %% [code]
make_UpSet("cage_lineage", upset_taxa_gut_BALB[upset_taxa_gut_BALB['genotype'].isin(['BALB/c'])], "upset_gut_BALB_cagelineage")

# %% [markdown]
#### B6J

# %% [code]
upset_taxa_gut_B6J= taxa_all_l6[taxa_all_l6['host_body_product'].isin(['UBERON:lumen of gut']) & taxa_all_l6['genotype'].isin(['C57BL6/J', 'B6CF1'])& taxa_all_l6['host_scientific_name'].isin(['Mus musculus']) & ~taxa_all_l6['generation'].isin(['not applicable'])]

# %% [code]
make_UpSet("generation", upset_taxa_gut_B6J, "upset_gut_B6J_generation")

# %% [markdown]
##### Cage Lineage

# %% [code]
make_UpSet("cage_lineage", upset_taxa_gut_B6J[upset_taxa_gut_B6J['genotype'].isin(['C57BL6/J'])], "upset_gut_B6J_cagelineage")

# %% [markdown]
### Sebum

# %% [code]
import matplotlib.pyplot as plt

# %% [code]
#prepare taxa_all_l6 table
upset_taxa_sebum= taxa_all_l6[taxa_all_l6['host_body_product'].isin(['UBERON:sebum']) & taxa_all_l6['host_scientific_name'].isin(['Mus musculus']) & ~taxa_all_l6['generation'].isin(['not applicable'])]

# %% [code]
make_UpSet("genotype", upset_taxa_sebum, "upset_sebum_genotype")

# %% [markdown]
#### BALB/c

# %% [code]
upset_taxa_sebum_BALB= taxa_all_l6[taxa_all_l6['host_body_product'].isin(['UBERON:sebum']) & taxa_all_l6['genotype'].isin(['BALB/c', 'B6CF1'])& taxa_all_l6['host_scientific_name'].isin(['Mus musculus']) & ~taxa_all_l6['generation'].isin(['not applicable'])]

# %% [code]
make_UpSet('generation', upset_taxa_sebum_BALB, "upset_sebum_BALB_generation")

# %% [markdown]
##### Cage Lineage

# %% [code]
make_UpSet("cage_lineage", upset_taxa_sebum_BALB[upset_taxa_sebum_BALB['genotype'].isin(['BALB/c'])], "upset_sebum_BALB_cagelineage")

# %% [markdown]
#### B6J

# %% [code]
upset_taxa_sebum_B6J= taxa_all_l6[taxa_all_l6['host_body_product'].isin(['UBERON:sebum']) & taxa_all_l6['genotype'].isin(['C57BL6/J'])& taxa_all_l6['host_scientific_name'].isin(['Mus musculus']) & ~taxa_all_l6['generation'].isin(['not applicable'])]

# %% [code]
make_UpSet("generation", upset_taxa_sebum_B6J, "upset_sebum_B6J_generation")

# %% [markdown]
##### Cage Lineage

# %% [code]
make_UpSet("cage_lineage", upset_taxa_sebum_B6J[upset_taxa_sebum_B6J["genotype"].isin(["C57BL6/J"])], "upset_sebum_B6J_cagelineage")

# %% [code]


# %% [code]


# %% [markdown]
## Statistics

# %% [code]
def permanova_beta (condition, dm, comp_group):
    counts = condition[comp_group].value_counts()
    idx = condition.index
    results = permanova(dm.filter(idx), meta.loc[idx, comp_group], permutations = 999)
    return counts, results

# %% [markdown]
### Gut
Wir vergleichen alle Betadiversitätergebnisse von Maussamples aus dem Darm, mit und ohne Vergleichsmäuse. Dabei werden die Genotypen verglichen!
#### Gut with Comparison

# %% [code]
dm_gut_comp = res_beta_gut_with_comp['results']['weighted_unifrac']

# %% [code]
genotype_dm = meta.loc[dm_gut_comp.ids,:][meta['genotype'].isin(['C57BL6/J', 'BALB/c'])]

# %% [code]
permanova_beta(genotype_dm, dm_gut_comp, "genotype")

# %% [markdown]
#### Janvier vs. Düsseldorf

# %% [code]
comparison = meta.loc[dm_gut_comp.ids,:][meta['comparison'].isin([1])]
permanova_beta(comparison, dm_gut_comp, "genotype")

# %% [markdown]
Im Vergleich sind die Unterschiede zwischen den Genotypen aus Dússeldorf und Janvier nicht signifikant.

# %% [markdown]
Vergleich zwischen den Balb und C57B Mäusen aus Janvier und aus Düsseldorf.

# %% [code]
for genotype in ["C57BL6/J", "BALB/c"]:
    comparison = meta.loc[dm_gut_comp.ids,:][meta['comparison'].isin([1]) & meta['genotype'].isin([genotype])]
    result = permanova_beta(comparison, dm_gut_comp, "geo_loc_name")
    print (f"{genotype} \n {result}")

# %% [code]
for city in ["France:Janvier", "Germany:Duesseldorf"]:
    comparison = meta.loc[dm_gut_comp.ids,:][meta['comparison'].isin([1]) & meta['geo_loc_name'].isin([city])]
    result = permanova_beta(comparison, dm_gut_comp, "genotype")
    print (f"{city} \n {result}")

# %% [markdown]
##### BALB/c

# %% [code]
for genotype in ['BALB/c', 'C57BL6/J']:
    comparison = meta.loc[dm_gut_comp.ids,:][(meta['comparison'].isin([0]) & meta['genotype'].isin(['BALB/c'])) | (meta['comparison'].isin([1]) & meta['genotype'].isin([genotype]))]
    result = permanova_beta(comparison, dm_gut_comp, "comparison")
    print (f"BALB/c experiment + {genotype} comparison \n {result}")
    for city in ['France:Janvier', 'Germany:Duesseldorf']:
        comparison = meta.loc[dm_gut_comp.ids,:][(meta['comparison'].isin([0]) & meta['genotype'].isin(['BALB/c'])) | (meta['comparison'].isin([1]) & meta['geo_loc_name'].isin([city]) & meta['genotype'].isin([genotype]))]
        result = permanova_beta(comparison, dm_gut_comp, "comparison")
        print (f"BALB/c experiment + {genotype}-{city} comparison \n {result}")

# %% [code]
for city in ['France:Janvier', 'Germany:Duesseldorf']:
    comparison = meta.loc[dm_gut_comp.ids,:][(meta['comparison'].isin([0]) & meta['genotype'].isin(['BALB/c'])) | (meta['comparison'].isin([1]) & meta['geo_loc_name'].isin([city]))]
    result = permanova_beta(comparison, dm_gut_comp, "comparison")
    print (f"BALB/c experiment + whole {city} comparison \n {result}")

# %% [code]
comparison = meta.loc[dm_gut_comp.ids,:][(meta['comparison'].isin([0]) & meta['genotype'].isin(['BALB/c'])) | meta['comparison'].isin([1])]
result = permanova_beta(comparison, dm_gut_comp, "comparison")
print (f"BALB/c experiment + whole comparison \n {result}")

# %% [code]
comparison = meta.loc[dm_gut_comp.ids,:][meta['comparison'].isin([0])]
result = permanova_beta(comparison, dm_gut_comp, "genotype")
print (f"BALB/c experiment + C57BL6/J \n {result}")

# %% [markdown]
##### C57BL6/J

# %% [code]
for genotype in ['BALB/c', 'C57BL6/J']:
    comparison = meta.loc[dm_gut_comp.ids,:][(meta['comparison'].isin([0]) & meta['genotype'].isin(['C57BL6/J'])) | (meta['comparison'].isin([1]) & meta['genotype'].isin([genotype]))]
    result = permanova_beta(comparison, dm_gut_comp, "comparison")
    print (f"C57BL6/J experiment + {genotype} comparison \n {result}")
    for city in ['France:Janvier', 'Germany:Duesseldorf']:
        comparison = meta.loc[dm_gut_comp.ids,:][(meta['comparison'].isin([0]) & meta['genotype'].isin(['C57BL6/J'])) | (meta['comparison'].isin([1]) & meta['geo_loc_name'].isin([city]) & meta['genotype'].isin([genotype]))]
        result = permanova_beta(comparison, dm_gut_comp, "comparison")
        print (f"C57BL6/J experiment + {genotype}-{city} comparison \n {result}")

# %% [code]
for city in ['France:Janvier', 'Germany:Duesseldorf']:
    comparison = meta.loc[dm_gut_comp.ids,:][(meta['comparison'].isin([0]) & meta['genotype'].isin(['C57BL6/J'])) | (meta['comparison'].isin([1]) & meta['geo_loc_name'].isin([city]))]
    result = permanova_beta(comparison, dm_gut_comp, "comparison")
    print (f"C57BL6/J experiment + whole {city} comparison \n {result}")

# %% [code]
comparison = meta.loc[dm_gut_comp.ids,:][(meta['comparison'].isin([0]) & meta['genotype'].isin(['C57BL6/J'])) | meta['comparison'].isin([1])]
result = permanova_beta(comparison, dm_gut_comp, "comparison")
print (f"C57BL6/J experiment + whole comparison \n {result}")

# %% [code]
comparison = meta.loc[dm_gut_comp.ids,:][meta['comparison'].isin([0])]
result = permanova_beta(comparison, dm_gut_comp, "genotype")
print (f"C57BL6/J experiment + BALB/c \n {result}")

# %% [markdown]
#### Gut without comparison

# %% [code]
dm_gut_no_comp = res_beta_gut_no_comp['results']['weighted_unifrac']

# %% [code]
meta['genotype'].value_counts()

# %% [code]
genotype_dm_no_comp = meta.loc[dm_gut_no_comp.ids,:][meta['genotype'].isin(['C57BL6/J', 'BALB/c'])]

# %% [code]
permanova_beta(genotype_dm_no_comp, dm_gut_no_comp, 'genotype')

# %% [markdown]
##### cage lineage

# %% [code]
#all
lineage_combination = list(itertools.combinations(['C1','C2','C3','B1','B2','B3'], 2))
count = 0
for lineage in lineage_combination:
    count += 1
    comparison = meta.loc[dm_gut_no_comp.ids,:][meta['comparison'].isin([0]) & meta['cage_lineage'].isin(lineage)]
    result = permanova_beta(comparison, dm_gut_no_comp, 'cage_lineage')
    print(f"Cage lineage pair {lineage} \n {result} \n {count}")

# %% [code]
#B6J
lineage_combination = list(itertools.combinations(['B1','B2','B3'], 2))
count = 0
for lineage in lineage_combination:
    count += 1
    comparison = meta.loc[dm_gut_no_comp.ids,:][meta['comparison'].isin([0]) & meta['genotype'].isin(['C57BL6/J']) & meta['cage_lineage'].isin(lineage)]
    result = permanova_beta(comparison, dm_gut_no_comp, 'cage_lineage')
    print(f"Cage lineage pair {lineage} \n {result} \n {count}")

# %% [code]
#B6J
lineage_combination = list(itertools.combinations(['C1','C2','C3'], 2))
count = 0
for lineage in lineage_combination:
    count += 1
    comparison = meta.loc[dm_gut_no_comp.ids,:][meta['comparison'].isin([0]) & meta['genotype'].isin(['BALB/c']) & meta['cage_lineage'].isin(lineage)]
    result = permanova_beta(comparison, dm_gut_no_comp, 'cage_lineage')
    print(f"Cage lineage pair {lineage} \n {result} \n {count}")

# %% [markdown]
##### Animals per cage

# %% [code]
def permanova_beta_stefan (condition, dm, comp_group):
    counts = condition[comp_group].value_counts()
    idx = condition.index
    results = permanova(dm.filter(idx), meta_stefan.loc[idx, comp_group], permutations = 9999)
    return counts, results

# %% [code]
animals_combination = list(itertools.combinations([1.0,2.0,3.0, 4.0, 5.0], 2))
count = 0
for animal_number in animals_combination:
    count += 1
    comparison = meta_stefan.loc[dm_gut_no_comp.ids,:][meta_stefan['iacuc_institute'].isin(['LANUV']) & meta_stefan['genotype'].isin(['balb/c']) & meta_stefan['animals_per_cage'].isin(animal_number)]
    result = permanova_beta_stefan(comparison, dm_gut_no_comp, 'animals_per_cage')
    print(f"Animals per cage pair {animal_number} \n {result} \n {count}")

# %% [code]
animals_combination = list(itertools.combinations([1.0,2.0,3.0, 4.0, 5.0], 2))
count = 0
for animal_number in animals_combination:
    count += 1
    comparison = meta_stefan.loc[dm_gut_no_comp.ids,:][meta_stefan['iacuc_institute'].isin(['LANUV']) & meta_stefan['genotype'].isin(['c57bl6/j']) & meta_stefan['animals_per_cage'].isin(animal_number)]
    result = permanova_beta_stefan(comparison, dm_gut_no_comp, 'animals_per_cage')
    print(f"Animals per cage pair {animal_number} \n {result} \n {count}")

# %% [code]
meta_stefan[meta_stefan['cage_line'].isin(['BALB/c_line6'])]['animals_per_cage'].value_counts()

# %% [code]
line_animals = {
    'C57BL6/J_line1': [1.0,2.0,3.0, 4.0],
    'C57BL6/J_line2': [1.0,2.0,3.0, 4.0,5.0],
    'C57BL6/J_line3': [1.0,2.0,3.0, 4.0],
    'BALB/c_line4': [1.0,2.0,3.0, 4.0],
    'BALB/c_line5': [1.0,2.0,3.0, 4.0,5.0],
    'BALB/c_line6': [1.0,2.0,3.0, 4.0]
}

# %% [code]
for cageline in line_animals.keys():
    print(cageline)
    animals_combination = list(itertools.combinations(line_animals[cageline], 2))
    count = 0
    for animal_number in animals_combination:
        count += 1
        comparison = meta_stefan.loc[dm_gut_no_comp.ids,:][meta_stefan['iacuc_institute'].isin(['LANUV']) & meta_stefan['cage_line'].isin([cageline]) & meta_stefan['animals_per_cage'].isin(animal_number)]
        result = permanova_beta_stefan(comparison, dm_gut_no_comp, 'animals_per_cage')
        print(f"Animals per cage pair {animal_number} \n {result} \n {count}")

# %% [code]
meta_stefan[meta_stefan['cage_line'].isin(['BALB/c_line6']) & meta_stefan['animals_per_cage'].isin([4.0])]['generation'].value_counts()

# %% [markdown]
###### Swarmplot Stefan

# %% [code]
list(meta[meta['generation'].isin(['F1']) & meta['cage_lineage'].isin(['C2']) & meta['host_body_product'].isin(['UBERON:lumen of gut'])].index)

# %% [code]
C1_to_C2_list = list(meta[meta['generation'].isin(['F1']) & meta['cage_lineage'].isin(['C1', 'C2']) & meta['host_body_product'].isin(['UBERON:lumen of gut'])].index)
C12_iter = list(itertools.combinations(C1_to_C2_list, 2))
B1_to_B2_list = list(meta[meta['generation'].isin(['F1']) & meta['cage_lineage'].isin(['B1', 'B2']) & meta['host_body_product'].isin(['UBERON:lumen of gut'])].index)
B12_iter = list(itertools.combinations(B1_to_B2_list, 2))
B1_to_B3_list = list(meta[meta['generation'].isin(['F1']) & meta['cage_lineage'].isin(['B1', 'B3']) & meta['host_body_product'].isin(['UBERON:lumen of gut'])].index)
B13_iter = list(itertools.combinations(B1_to_B3_list, 2))
B2_to_B3_list = list(meta[meta['generation'].isin(['F1']) & meta['cage_lineage'].isin(['B2', 'B3']) & meta['host_body_product'].isin(['UBERON:lumen of gut'])].index)
B23_iter = list(itertools.combinations(B2_to_B3_list, 2))

# %% [code]
C1= list(meta[meta['generation'].isin(['F1']) & meta['cage_lineage'].isin(['C1']) & meta['host_body_product'].isin(['UBERON:lumen of gut'])].index)
C2= list(meta[meta['generation'].isin(['F1']) & meta['cage_lineage'].isin(['C2']) & meta['host_body_product'].isin(['UBERON:lumen of gut'])].index)
B1= list(meta[meta['generation'].isin(['F1']) & meta['cage_lineage'].isin(['B1']) & meta['host_body_product'].isin(['UBERON:lumen of gut'])].index)
B2= list(meta[meta['generation'].isin(['F1']) & meta['cage_lineage'].isin(['B2']) & meta['host_body_product'].isin(['UBERON:lumen of gut'])].index)
B3= list(meta[meta['generation'].isin(['F1']) & meta['cage_lineage'].isin(['B3']) & meta['host_body_product'].isin(['UBERON:lumen of gut'])].index)

# %% [code]
combinations = [[C1,C2], [B1, B2], [B1,B3], [B2,B3]]
def get_combinations(pair):
    couples = []
    for i in range(len(pair[0])):
        for j in range(len(pair[1])):
            couples.append((pair[0][i], pair[1][j]))
    return couples

# %% [code]
#this is very ugly right now but i dont care
C12_iter=get_combinations(combinations[0])
B12_iter=get_combinations(combinations[1])
B13_iter=get_combinations(combinations[2])
B23_iter=get_combinations(combinations[3])

# %% [code]
def all_distances(iter_list, dm):
    distances = []
    for combination in iter_list:
        distance = dm[combination]
        distances.append(distance)
    return distances

# %% [code]
C12_distances=all_distances(C12_iter,dm_gut_no_comp)
B12_distances=all_distances(B12_iter,dm_gut_no_comp)
B13_distances=all_distances(B13_iter,dm_gut_no_comp)
B23_distances=all_distances(B23_iter,dm_gut_no_comp)
df_distances =pd.DataFrame({'C1-C2' : pd.Series(C12_distances),
                            'B1-B2' : pd.Series(B12_distances),
                            'B2-B3' : pd.Series(B23_distances),
                            'B1-B3' : pd.Series(B13_distances)})
df_distances

# %% [code]
%matplotlib inline
ax = sns.swarmplot(df_distances)
ax.set(xlabel='Cage Lineage Combinations', ylabel='unweighted unifrac distance')
plt.title('Distances between Cage lineages in F1 - uw unifrac')
plt.savefig(work_dir + f"/swarm_F1.png", transparent=False, bbox_inches='tight')
plt.show()

# %% [code]
%matplotlib inline
ax = sns.boxplot(df_distances)
ax.set(xlabel='Cage Lineage Combinations', ylabel='unweighted unifrac distance')
plt.title('Distances between Cage lineages in F1 - uw unifrac')
plt.savefig(work_dir + f"/box_F1.png", transparent=False, bbox_inches='tight')
plt.show()

# %% [code]
%matplotlib inline
ax = sns.violinplot(df_distances)
ax.set(xlabel='Cage Lineage Combinations', ylabel='unweighted unifrac distance')
plt.title('Distances between Cage lineages in F1 - uw unifrac')
plt.savefig(work_dir + f"/violin_F1.png", transparent=False, bbox_inches='tight')
plt.show()

# %% [markdown]
##### Generations

# %% [code]
generations_combination = list(itertools.combinations(['P1', 'F1', 'F2', 'F3', 'F4', 'F5'], 2))

# %% [code]
count = 0
for generation in generations_combination:
    count += 1
    comparison = meta.loc[dm_gut_no_comp.ids,:][meta['comparison'].isin([0]) & meta['genotype'].isin(['BALB/c']) & meta['generation'].isin(generation)]
    result = permanova_beta(comparison, dm_gut_no_comp, 'generation')
    print(f"BALB/c generation pair {generation} \n {result} \n {count}")

# %% [code]
count = 0
for generation in generations_combination:
    count += 1
    comparison = meta.loc[dm_gut_no_comp.ids,:][meta['comparison'].isin([0]) & meta['genotype'].isin(['C57BL6/J']) & meta['generation'].isin(generation)]
    result = permanova_beta(comparison, dm_gut_no_comp, 'generation')
    print(f"C57BL6/J generation pair {generation} \n {result} \n {count}")

# %% [markdown]
##### Genotype Per Generation

# %% [code]
for generation in ['P1', 'F1', 'F2', 'F3', 'F4', 'F5']:
    comparison = meta.loc[dm_gut_no_comp.ids,:][meta['comparison'].isin([0]) & meta['generation'].isin([generation])]
    result = permanova_beta(comparison, dm_gut_no_comp, 'genotype')
    print(f"{generation} \n {result}")

# %% [code]


# %% [markdown]
##### Sex in each generation per genotype

# %% [code]
count = 0
for generation in ['P1', 'F1', 'F2', 'F3', 'F4', 'F5']:
    comparison = meta.loc[dm_gut_no_comp.ids,:][meta['generation'].isin([generation]) & meta['genotype'].isin(['BALB/c'])]
    result = permanova_beta(comparison, dm_gut_no_comp, 'sex')
    count += 1
    print(f"{generation} \n {result}")
print(count)

# %% [code]
count = 0
for generation in ['P1', 'F1', 'F2', 'F3', 'F4', 'F5']:
    comparison = meta.loc[dm_gut_no_comp.ids,:][meta['generation'].isin([generation]) & meta['genotype'].isin(['C57BL6/J'])]
    result = permanova_beta(comparison, dm_gut_no_comp, 'sex')
    count += 1
    print(f"{generation} \n {result}")
print(count)

# %% [markdown]
##### Genotype in each generation per sex

# %% [code]


# %% [code]
count = 0
for generation in ['P1', 'F1', 'F2', 'F3', 'F4', 'F5']:
    comparison = meta.loc[dm_gut_no_comp.ids,:][meta['generation'].isin([generation]) & meta['sex'].isin(['female'])]
    result = permanova_beta(comparison, dm_gut_no_comp, 'genotype')
    count += 1
    print(f"{generation} \n {result}")
print(count)

# %% [code]
count = 0
for generation in ['P1', 'F1', 'F2', 'F3', 'F4', 'F5']:
    comparison = meta.loc[dm_gut_no_comp.ids,:][meta['generation'].isin([generation]) & meta['sex'].isin(['male'])]
    result = permanova_beta(comparison, dm_gut_no_comp, 'genotype')
    count += 1
    print(f"{generation} \n {result}")
print(count)

# %% [code]


# %% [markdown]
### Sebum

# %% [markdown]
#### Sebum with Comparison

# %% [code]
dm_sebum_comp = res_beta_sebum_with_comp['results']['unweighted_unifrac']
gen_comparison_sebum = meta.loc[dm_sebum_comp.ids,:][meta['genotype'].isin(['C57BL6/J', 'BALB/c'])]
permanova_beta(gen_comparison_sebum, dm_sebum_comp, "genotype")

# %% [markdown]
#### Janvier vs. Düsseldorf

# %% [code]
test = meta[meta['comparison'].isin([1]) & meta['host_body_product'].isin(['UBERON:sebum'])]
test['geo_loc_name'].value_counts()

# %% [code]
comparison = meta.loc[dm_sebum_comp.ids,:][meta['comparison'].isin([1])]
permanova_beta(comparison, dm_sebum_comp, "genotype")

# %% [code]
for genotype in ["C57BL6/J", "BALB/c"]:
    comparison = meta.loc[dm_sebum_comp.ids,:][meta['comparison'].isin([1]) & meta['genotype'].isin([genotype])]
    result = permanova_beta(comparison, dm_sebum_comp, "geo_loc_name")
    print (f"{genotype} \n {result}")

# %% [code]
for city in ["France:Janvier", "Germany:Duesseldorf"]:
    comparison = meta.loc[dm_sebum_comp.ids,:][meta['comparison'].isin([1]) & meta['geo_loc_name'].isin([city])]
    result = permanova_beta(comparison, dm_sebum_comp, "genotype")
    print (f"{city} \n {result}")

# %% [code]


# %% [markdown]
##### BALB/c

# %% [code]
for genotype in ['BALB/c', 'C57BL6/J']:
    comparison = meta.loc[dm_sebum_comp.ids,:][(meta['comparison'].isin([0]) & meta['genotype'].isin(['BALB/c'])) | (meta['comparison'].isin([1]) & meta['genotype'].isin([genotype]))]
    result = permanova_beta(comparison, dm_sebum_comp, "comparison")
    print (f"BALB/c experiment + {genotype} comparison \n {result}")
    for city in ['France:Janvier', 'Germany:Duesseldorf']:
        comparison = meta.loc[dm_sebum_comp.ids,:][(meta['comparison'].isin([0]) & meta['genotype'].isin(['BALB/c'])) | (meta['comparison'].isin([1]) & meta['geo_loc_name'].isin([city]) & meta['genotype'].isin([genotype]))]
        result = permanova_beta(comparison, dm_sebum_comp, "comparison")
        print (f"BALB/c experiment + {genotype}-{city} comparison \n {result}")

# %% [code]
for city in ['France:Janvier', 'Germany:Duesseldorf']:
    comparison = meta.loc[dm_sebum_comp.ids,:][(meta['comparison'].isin([0]) & meta['genotype'].isin(['BALB/c'])) | (meta['comparison'].isin([1]) & meta['geo_loc_name'].isin([city]))]
    result = permanova_beta(comparison, dm_sebum_comp, "comparison")
    print (f"BALB/c experiment + whole {city} comparison \n {result}")

# %% [code]
comparison = meta.loc[dm_sebum_comp.ids,:][(meta['comparison'].isin([0]) & meta['genotype'].isin(['BALB/c'])) | meta['comparison'].isin([1])]
result = permanova_beta(comparison, dm_sebum_comp, "comparison")
print (f"BALB/c experiment + whole comparison \n {result}")

# %% [code]
comparison = meta.loc[dm_sebum_comp.ids,:][meta['comparison'].isin([0]) & meta['genotype'].isin(['C57BL6/J', 'BALB/c'])]
result = permanova_beta(comparison, dm_sebum_comp, "genotype")
print (f"BALB/c experiment + C57BL6/J \n {result}")

# %% [markdown]
##### C57BL6/J

# %% [code]
for genotype in ['BALB/c', 'C57BL6/J']:
    comparison = meta.loc[dm_sebum_comp.ids,:][(meta['comparison'].isin([0]) & meta['genotype'].isin(['C57BL6/J'])) | (meta['comparison'].isin([1]) & meta['genotype'].isin([genotype]))]
    result = permanova_beta(comparison, dm_sebum_comp, "comparison")
    print (f"C57BL6/J experiment + {genotype} comparison \n {result}")
    for city in ['France:Janvier', 'Germany:Duesseldorf']:
        comparison = meta.loc[dm_sebum_comp.ids,:][(meta['comparison'].isin([0]) & meta['genotype'].isin(['C57BL6/J'])) | (meta['comparison'].isin([1]) & meta['geo_loc_name'].isin([city]) & meta['genotype'].isin([genotype]))]
        result = permanova_beta(comparison, dm_sebum_comp, "comparison")
        print (f"C57BL6/J experiment + {genotype}-{city} comparison \n {result}")

# %% [code]
for city in ['France:Janvier', 'Germany:Duesseldorf']:
    comparison = meta.loc[dm_sebum_comp.ids,:][(meta['comparison'].isin([0]) & meta['genotype'].isin(['C57BL6/J'])) | (meta['comparison'].isin([1]) & meta['geo_loc_name'].isin([city]))]
    result = permanova_beta(comparison, dm_sebum_comp, "comparison")
    print (f"C57BL6/J experiment + whole {city} comparison \n {result}")

# %% [code]
comparison = meta.loc[dm_sebum_comp.ids,:][(meta['comparison'].isin([0]) & meta['genotype'].isin(['C57BL6/J'])) | meta['comparison'].isin([1])]
result = permanova_beta(comparison, dm_sebum_comp, "comparison")
print (f"C57BL6/J experiment + whole comparison \n {result}")

# %% [code]
comparison = meta.loc[dm_sebum_comp.ids,:][meta['comparison'].isin([0]) & meta['genotype'].isin(['C57BL6/J', 'BALB/c'])]
result = permanova_beta(comparison, dm_sebum_comp, "genotype")
print (f"C57BL6/J experiment + BALB/c experiment \n {result}")

# %% [markdown]
#### Sebum without Comparison

# %% [code]
dm_sebum_no_comp = res_beta_sebum_no_comp['results']['unweighted_unifrac']
gen_comparison_sebum = meta.loc[dm_sebum_no_comp.ids,:][meta['genotype'].isin(['C57BL6/J', 'BALB/c'])]
permanova_beta(gen_comparison_sebum, dm_sebum_no_comp, "genotype")

# %% [markdown]
##### cage lineage

# %% [code]
#all
lineage_combination = list(itertools.combinations(['C1','C2','C3','B1','B2','B3'], 2))
count = 0
for lineage in lineage_combination:
    count += 1
    comparison = meta.loc[dm_sebum_no_comp.ids,:][meta['comparison'].isin([0]) & meta['cage_lineage'].isin(lineage)]
    result = permanova_beta(comparison, dm_sebum_no_comp, 'cage_lineage')
    print(f"Cage lineage pair {lineage} \n {result} \n {count}")

# %% [code]
#B6J
lineage_combination = list(itertools.combinations(['B1','B2','B3'], 2))
count = 0
for lineage in lineage_combination:
    count += 1
    comparison = meta.loc[dm_sebum_no_comp.ids,:][meta['comparison'].isin([0]) & meta['genotype'].isin(['C57BL6/J']) & meta['cage_lineage'].isin(lineage)]
    result = permanova_beta(comparison, dm_sebum_no_comp, 'cage_lineage')
    print(f"Cage lineage pair {lineage} \n {result} \n {count}")

# %% [code]
#BALB
lineage_combination = list(itertools.combinations(['C1','C2','C3'], 2))
count = 0
for lineage in lineage_combination:
    count += 1
    comparison = meta.loc[dm_sebum_no_comp.ids,:][meta['comparison'].isin([0]) & meta['genotype'].isin(['BALB/c']) & meta['cage_lineage'].isin(lineage)]
    result = permanova_beta(comparison, dm_sebum_no_comp, 'cage_lineage')
    print(f"Cage lineage pair {lineage} \n {result} \n {count}")

# %% [markdown]
##### Generation per Genotype

# %% [code]
count = 0
for generation in generations_combination:
    count += 1
    comparison = meta.loc[dm_sebum_no_comp.ids,:][meta['comparison'].isin([0]) & meta['genotype'].isin(['BALB/c']) & meta['generation'].isin(generation)]
    result = permanova_beta(comparison, dm_sebum_no_comp, 'generation')
    print(f"BALB/c generation pair {generation} \n {result} \n {count}")

# %% [code]
count = 0
for generation in generations_combination:
    count += 1
    comparison = meta.loc[dm_sebum_no_comp.ids,:][meta['comparison'].isin([0]) & meta['genotype'].isin(['C57BL6/J']) & meta['generation'].isin(generation)]
    result = permanova_beta(comparison, dm_sebum_no_comp, 'generation')
    print(f"C57BL6/J generation pair {generation} \n {result} \n {count}")

# %% [markdown]
##### Genotype per Generation

# %% [code]
for generation in ['P1', 'F1', 'F2', 'F3', 'F4', 'F5']:
    comparison = meta.loc[dm_sebum_no_comp.ids,:][meta['comparison'].isin([0]) & meta['generation'].isin([generation])]
    result = permanova_beta(comparison, dm_sebum_no_comp, 'genotype')
    print(f"{generation} \n {result}")

# %% [markdown]
##### Sex in each generation per genotype

# %% [code]
count = 0
for generation in ['P1', 'F1', 'F2', 'F3', 'F4', 'F5']:
    comparison = meta.loc[dm_sebum_no_comp.ids,:][meta['generation'].isin([generation]) & meta['genotype'].isin(['BALB/c'])]
    result = permanova_beta(comparison, dm_sebum_no_comp, 'sex')
    count += 1
    print(f"{generation} \n {result}")
print(count)

# %% [code]
count = 0
for generation in ['P1', 'F1', 'F2', 'F3', 'F4', 'F5']:
    comparison = meta.loc[dm_sebum_no_comp.ids,:][meta['generation'].isin([generation]) & meta['genotype'].isin(['C57BL6/J'])]
    result = permanova_beta(comparison, dm_sebum_no_comp, 'sex')
    count += 1
    print(f"{generation} \n {result}")
print(count)

# %% [code]


# %% [markdown]
##### Genotype in each generation per sex

# %% [code]
count = 0
for generation in ['P1', 'F1', 'F2', 'F3', 'F4', 'F5']:
    comparison = meta.loc[dm_sebum_no_comp.ids,:][meta['generation'].isin([generation]) & meta['sex'].isin(['female'])]
    result = permanova_beta(comparison, dm_sebum_no_comp, 'genotype')
    count += 1
    print(f"{generation} \n {result}")
print(count)

# %% [code]
count = 0
for generation in ['P1', 'F1', 'F2', 'F3', 'F4', 'F5']:
    comparison = meta.loc[dm_sebum_no_comp.ids,:][meta['generation'].isin([generation]) & meta['sex'].isin(['male'])]
    result = permanova_beta(comparison, dm_sebum_no_comp, 'genotype')
    count += 1
    print(f"{generation} \n {result}")
print(count)

# %% [markdown]
## Alpha Statistics

# %% [code]
def mannwhitneyu_table(pair, pairlist_str, count):
    generation_pair = (pairlist_str[count])
    print(generation_pair)
    statistic, p = mannwhitneyu(pair[0], pair[1], alternative='greater')
    statistics.append(statistic)
    p_values.append(p)
    groups.append(generation_pair)
    print(f'Statistik: {statistic} \n p-Wert: {p} \n')
    bonferroni.append("Y") if p < alpha else bonferroni.append("N")
    zerofive.append("Y") if p< 0.05 else zerofive.append("N")
    
    

# %% [markdown]
### Gut

# %% [markdown]
#### Differences within the comparisons

# %% [code]
alpha_C57B_comp = alpha_results_merged_gut[alpha_results_merged_gut['genotype'].isin(['C57BL6/J', 'B6CF1']) & alpha_results_merged_gut['comparison'].isin([1])]
alpha_BALB_comp = alpha_results_merged_gut[alpha_results_merged_gut['genotype'].isin(['BALB/c', 'B6CF1']) & alpha_results_merged_gut['comparison'].isin([1])]

# %% [code]
Janvier_faith_comp = alpha_results_merged_gut[(alpha_results_merged_gut['comparison'].isin([1])) & (alpha_results_merged_gut['geo_loc_name'].isin(['France:Janvier']))]['PD_whole_tree']
Duess_faith_comp = alpha_results_merged_gut[alpha_results_merged_gut['comparison'].isin([1]) & (alpha_results_merged_gut['geo_loc_name'].isin(['Germany:Duesseldorf']))]['PD_whole_tree']
exp_faith_comp = alpha_results_merged_gut[alpha_results_merged_gut['comparison'].isin([1])]['PD_whole_tree']

# %% [code]
#all
faith_groups_comp = list(itertools.combinations([Janvier_faith_comp, Duess_faith_comp, exp_faith_comp],2))
faith_groups_comp_str = list(itertools.combinations(["Janvier_faith_comp", "Duess_faith_comp", "exp_faith_comp"],2))

alpha = 0.05/len(faith_groups_comp)
count = 0
statistics = []
p_values = []
significant = []
groups = []
bonferroni = []
zerofive = []
for pair in faith_groups_comp:
    mannwhitneyu_table(pair, faith_groups_comp_str, count)
    count +=1

alpha_results_df = pd.DataFrame({'Comparisons': groups,
                                'Statistics': statistics,
                                'p_values': p_values,
                                'Alpha': alpha,
                                'Reject H0?': bonferroni,
                                'No Bonferroni reject H0': zerofive})
alpha_results_df

# %% [code]
Janvier_faith_C57B_comp = alpha_C57B_comp[(alpha_C57B_comp['comparison'].isin([1])) & (alpha_C57B_comp['geo_loc_name'].isin(['France:Janvier']))]['PD_whole_tree']
Duess_faith_C57B_comp = alpha_C57B_comp[alpha_C57B_comp['comparison'].isin([1]) & (alpha_C57B_comp['geo_loc_name'].isin(['Germany:Duesseldorf']))]['PD_whole_tree']
exp_faith_C57B_comp = alpha_C57B_comp[alpha_C57B_comp['comparison'].isin([1])]['PD_whole_tree']

faith_C57B_groups_comp = list(itertools.combinations([Janvier_faith_C57B_comp, Duess_faith_C57B_comp, exp_faith_C57B_comp],2))
faith_C57B_groups_comp_str = list(itertools.combinations(["Janvier_faith_C57B", "Duess_faith_C57B", "exp_faith_C57B"],2))

# %% [code]
alpha = 0.05/len(faith_C57B_groups_comp)
count = 0
statistics = []
p_values = []
significant = []
groups = []
bonferroni = []
zerofive = []
for pair in faith_C57B_groups_comp:
    mannwhitneyu_table(pair, faith_C57B_groups_comp_str, count)
    count +=1

# %% [code]
alpha_results_df = pd.DataFrame({'Comparisons': groups,
                                'Statistics': statistics,
                                'p_values': p_values,
                                'Alpha': alpha,
                                'Reject H0?': bonferroni,
                                'No Bonferroni reject H0': zerofive})
alpha_results_df

# %% [code]
#BALB/c

# %% [code]
Janvier_faith_BALB = alpha_BALB_comp[(alpha_BALB_comp['comparison'].isin([1])) & (alpha_BALB_comp['geo_loc_name'].isin(['France:Janvier']))]['PD_whole_tree']
Duess_faith_BALB = alpha_BALB_comp[alpha_BALB_comp['comparison'].isin([1]) & (alpha_BALB_comp['geo_loc_name'].isin(['Germany:Duesseldorf']))]['PD_whole_tree']
exp_faith_BALB = alpha_BALB_comp[alpha_BALB_comp['comparison'].isin([1])]['PD_whole_tree']

faith_BALB_groups_comp = list(itertools.combinations([Janvier_faith_BALB, Duess_faith_BALB, exp_faith_BALB],2))
faith_BALB_groups_comp_str = list(itertools.combinations(["Janvier_faith_BALB", "Duess_faith_BALB", "exp_faith_BALB"],2))

# %% [code]
alpha = 0.05/len(faith_BALB_groups_comp)
count = 0
statistics = []
p_values = []
significant = []
groups = []
bonferroni = []
zerofive = []
for pair in faith_BALB_groups_comp:
    mannwhitneyu_table(pair, faith_BALB_groups_comp_str, count)
    count +=1

# %% [code]
alpha_results_df = pd.DataFrame({'Comparisons': groups,
                                'Statistics': statistics,
                                'p_values': p_values,
                                'Alpha': alpha,
                                'Reject H0?': bonferroni,
                                'No Bonferroni reject H0': zerofive})
alpha_results_df

# %% [code]
#genotype per category
statistic, p = mannwhitneyu(Janvier_faith_C57B_comp, Janvier_faith_BALB, alternative='less')
print(f'Statistic: {statistic} \n p={p}')
statistic, p = mannwhitneyu(Duess_faith_C57B_comp, Duess_faith_BALB, alternative='less')
print(f'Statistic: {statistic} \n p={p}')
statistic, p = mannwhitneyu(exp_faith_C57B_comp, exp_faith_BALB, alternative='less')
print(f'Statistic: {statistic} \n p={p}')

# %% [markdown]
#### Differences in Alpha Diversity per Genotype between generations

# %% [code]
alpha_C57B = alpha_results_merged_gut[alpha_results_merged_gut['genotype'].isin(['C57BL6/J', 'B6CF1'])]
alpha_BALB = alpha_results_merged_gut[alpha_results_merged_gut['genotype'].isin(['BALB/c', 'B6CF1'])]

# %% [code]
F1_faith_C57B = alpha_C57B[alpha_C57B['generation'].isin(['F1'])]['PD_whole_tree']
F2_faith_C57B = alpha_C57B[alpha_C57B['generation'].isin(['F2'])]['PD_whole_tree']
F3_faith_C57B = alpha_C57B[alpha_C57B['generation'].isin(['F3'])]['PD_whole_tree']
F4_faith_C57B = alpha_C57B[alpha_C57B['generation'].isin(['F4'])]['PD_whole_tree']
F5_faith_C57B = alpha_C57B[alpha_C57B['generation'].isin(['F5'])]['PD_whole_tree']
P1_faith_C57B = alpha_C57B[alpha_C57B['generation'].isin(['P1'])]['PD_whole_tree']
RM_faith_C57B = alpha_C57B[alpha_C57B['generation'].isin(['RM'])]['PD_whole_tree']

faith_C57B_groups = list(itertools.combinations([RM_faith_C57B, P1_faith_C57B, F1_faith_C57B, F2_faith_C57B, F3_faith_C57B, F4_faith_C57B, F5_faith_C57B],2))
faith_C57B_groups_str = list(itertools.combinations(['RM_faith_C57B','P1_faith_C57B', 'F1_faith_C57B', 'F2_faith_C57B', 'F3_faith_C57B', 'F4_faith_C57B', 'F5_faith_C57B'],2))

# %% [code]
F1_faith_BALB = alpha_BALB[alpha_BALB['generation'].isin(['F1'])]['PD_whole_tree']
F2_faith_BALB = alpha_BALB[alpha_BALB['generation'].isin(['F2'])]['PD_whole_tree']
F3_faith_BALB = alpha_BALB[alpha_BALB['generation'].isin(['F3'])]['PD_whole_tree']
F4_faith_BALB = alpha_BALB[alpha_BALB['generation'].isin(['F4'])]['PD_whole_tree']
F5_faith_BALB = alpha_BALB[alpha_BALB['generation'].isin(['F5'])]['PD_whole_tree']
P1_faith_BALB = alpha_BALB[alpha_BALB['generation'].isin(['P1'])]['PD_whole_tree']
RM_faith_BALB = alpha_BALB[alpha_BALB['generation'].isin(['RM'])]['PD_whole_tree']

faith_BALB_groups = list(itertools.combinations([RM_faith_BALB, P1_faith_BALB, F1_faith_BALB, F2_faith_BALB, F3_faith_BALB, F4_faith_BALB, F5_faith_BALB],2))
faith_BALB_groups_str = list(itertools.combinations(['RM_faith_BALB','P1_faith_BALB', 'F1_faith_BALB', 'F2_faith_BALB', 'F3_faith_BALB', 'F4_faith_BALB', 'F5_faith_BALB'],2))

# %% [code]
alpha = 0.05/len(faith_C57B_groups)
count = 0
statistics = []
p_values = []
significant = []
groups = []
bonferroni = []
zerofive = []
for pair in faith_C57B_groups:
    mannwhitneyu_table(pair, faith_C57B_groups_str, count)
    count +=1

# %% [code]
alpha = 0.05/len(faith_BALB_groups)
count = 0
for pair in faith_BALB_groups:
    mannwhitneyu_table(pair, faith_BALB_groups_str, count)
    count +=1

# %% [code]
alpha_results_df = pd.DataFrame({'Comparisons': groups,
                                'Statistics': statistics,
                                'p_values': p_values,
                                'Alpha': alpha,
                                'Reject H0?': bonferroni,
                                'No Bonferroni reject H0': zerofive})
alpha_results_df

# %% [markdown]
##### Differences in Alpha Diversity per Generation between genotypes

# %% [code]
generation_list_alpha = [(RM_faith_C57B, RM_faith_BALB),
                        (P1_faith_C57B, P1_faith_BALB),
                        (F1_faith_C57B, F1_faith_BALB),
                        (F2_faith_C57B, F2_faith_BALB),
                        (F3_faith_C57B, F3_faith_BALB),
                        (F4_faith_C57B, F4_faith_BALB),
                        (F5_faith_C57B, F5_faith_BALB)]
generation_list_alpha_str = [('RM_faith_C57B', 'RM_faith_BALB'),
                        ('P1_faith_C57B', 'P1_faith_BALB'),
                        ('F1_faith_C57B', 'F1_faith_BALB'),
                        ('F2_faith_C57B', 'F2_faith_BALB'),
                        ('F3_faith_C57B', 'F3_faith_BALB'),
                        ('F4_faith_C57B', 'F4_faith_BALB'),
                        ('F5_faith_C57B', 'F5_faith_BALB')]
statistics = []
p_values = []
significant = []
groups = []
bonferroni = []
zerofive = []
generations=6
alpha = 0.05/generations
count = 0
for pair in generation_list_alpha:
    mannwhitneyu_table(pair, generation_list_alpha_str, count)
    count += 1

# %% [code]
alpha_results_gen_df = pd.DataFrame({'Comparisons': groups,
                                'Statistics': statistics,
                                'p_values': p_values,
                                'Alpha': alpha,
                                'Reject H0: x<=y?': bonferroni,
                                'No Bonferroni reject H0': zerofive})
alpha_results_gen_df

# %% [markdown]
##### cage groups

# %% [code]
faith_B1 = alpha_BALB[alpha_BALB['cage_lineage'].isin(['C1'])]['PD_whole_tree']
faith_B2 = alpha_BALB[alpha_BALB['cage_lineage'].isin(['C2'])]['PD_whole_tree']
faith_B3 = alpha_BALB[alpha_BALB['cage_lineage'].isin(['C3'])]['PD_whole_tree']

faith_BALB_Cage = list(itertools.combinations([faith_B1, faith_B2, faith_B3],2))
faith_BALB_Cage_str = list(itertools.combinations(['faith_B1', 'faith_B2', 'faith_B3'],2))

# %% [code]
alpha = 0.05/len(faith_BALB_Cage)
count = 0
statistics = []
p_values = []
significant = []
groups = []
bonferroni = []
zerofive = []
for pair in faith_BALB_Cage:
    mannwhitneyu_table(pair, faith_BALB_Cage_str, count)
    count +=1

alpha_results_df = pd.DataFrame({'Comparisons': groups,
                                'Statistics': statistics,
                                'p_values': p_values,
                                'Alpha': alpha,
                                'Reject H0?': bonferroni,
                                'No Bonferroni reject H0': zerofive})
alpha_results_df

# %% [code]
faith_C1 = alpha_C57B[alpha_C57B['cage_lineage'].isin(['B1'])]['PD_whole_tree']
faith_C2 = alpha_C57B[alpha_C57B['cage_lineage'].isin(['B2'])]['PD_whole_tree']
faith_C3 = alpha_C57B[alpha_C57B['cage_lineage'].isin(['B3'])]['PD_whole_tree']

faith_C57B_Cage = list(itertools.combinations([faith_C1, faith_C2, faith_C3],2))
faith_C57B_Cage_str = list(itertools.combinations(['faith_C1', 'faith_C2', 'faith_C3'],2))

# %% [code]
alpha = 0.05/len(faith_C57B_Cage)
count = 0
statistics = []
p_values = []
significant = []
groups = []
bonferroni = []
zerofive = []
for pair in faith_C57B_Cage:
    mannwhitneyu_table(pair, faith_C57B_Cage_str, count)
    count +=1

alpha_results_df = pd.DataFrame({'Comparisons': groups,
                                'Statistics': statistics,
                                'p_values': p_values,
                                'Alpha': alpha,
                                'Reject H0?': bonferroni,
                                'No Bonferroni reject H0': zerofive})
alpha_results_df

# %% [markdown]
##### cage lines, animals per cage

# %% [code]
alpha_C57B_stefan = alpha_results_merged_gut_stefan[alpha_results_merged_gut_stefan['genotype'].isin(['c57bl6/j', 'b6cf1'])]
alpha_BALB_stefan = alpha_results_merged_gut_stefan[alpha_results_merged_gut_stefan['genotype'].isin(['balb/c', 'b6cf1'])]

# %% [code]
#cage line BALB/c_line6
faith_l6_4 = alpha_BALB_stefan[alpha_BALB_stefan['cage_line'].isin(['BALB/c_line6']) & alpha_BALB_stefan['animals_per_cage'].isin([4.0])]['PD_whole_tree']
faith_l6_1 = alpha_BALB_stefan[alpha_BALB_stefan['cage_line'].isin(['BALB/c_line6']) & alpha_BALB_stefan['animals_per_cage'].isin([1.0])]['PD_whole_tree']
faith_l6_2 = alpha_BALB_stefan[alpha_BALB_stefan['cage_line'].isin(['BALB/c_line6']) & alpha_BALB_stefan['animals_per_cage'].isin([2.0])]['PD_whole_tree']
faith_l6_3 = alpha_BALB_stefan[alpha_BALB_stefan['cage_line'].isin(['BALB/c_line6']) & alpha_BALB_stefan['animals_per_cage'].isin([3.0])]['PD_whole_tree']

faith_l6_animal = list(itertools.combinations([faith_l6_1, faith_l6_2, faith_l6_3, faith_l6_4],2))
faith_l6_animal_str = list(itertools.combinations(['faith_l6_1', 'faith_l6_2', 'faith_l6_3', 'faith_l6_4'],2))

# %% [code]
alpha = 0.05/len(faith_l6_animal)
count = 0
statistics = []
p_values = []
significant = []
groups = []
bonferroni = []
zerofive = []
for pair in faith_l6_animal:
    mannwhitneyu_table(pair, faith_l6_animal_str, count)
    count +=1

alpha_results_df = pd.DataFrame({'Comparisons': groups,
                                'Statistics': statistics,
                                'p_values': p_values,
                                'Alpha': alpha,
                                'Reject H0?': bonferroni,
                                'No Bonferroni reject H0': zerofive})
alpha_results_df

# %% [code]
#cage line BALB/c_line5
faith_l5_4 = alpha_BALB_stefan[alpha_BALB_stefan['cage_line'].isin(['BALB/c_line5']) & alpha_BALB_stefan['animals_per_cage'].isin([4.0])]['PD_whole_tree']
faith_l5_1 = alpha_BALB_stefan[alpha_BALB_stefan['cage_line'].isin(['BALB/c_line5']) & alpha_BALB_stefan['animals_per_cage'].isin([1.0])]['PD_whole_tree']
faith_l5_2 = alpha_BALB_stefan[alpha_BALB_stefan['cage_line'].isin(['BALB/c_line5']) & alpha_BALB_stefan['animals_per_cage'].isin([2.0])]['PD_whole_tree']
faith_l5_3 = alpha_BALB_stefan[alpha_BALB_stefan['cage_line'].isin(['BALB/c_line5']) & alpha_BALB_stefan['animals_per_cage'].isin([3.0])]['PD_whole_tree']
faith_l5_5 = alpha_BALB_stefan[alpha_BALB_stefan['cage_line'].isin(['BALB/c_line5']) & alpha_BALB_stefan['animals_per_cage'].isin([5.0])]['PD_whole_tree']

faith_l5_animal = list(itertools.combinations([faith_l5_1, faith_l5_2, faith_l5_3, faith_l5_4, faith_l5_5],2))
faith_l5_animal_str = list(itertools.combinations(['faith_l5_1', 'faith_l5_2', 'faith_l5_3', 'faith_l5_4', 'faith_l5_5'],2))

# %% [code]
alpha = 0.05/len(faith_l5_animal)
count = 0
statistics = []
p_values = []
significant = []
groups = []
bonferroni = []
zerofive = []
for pair in faith_l5_animal:
    mannwhitneyu_table(pair, faith_l5_animal_str, count)
    count +=1

alpha_results_df = pd.DataFrame({'Comparisons': groups,
                                'Statistics': statistics,
                                'p_values': p_values,
                                'Alpha': alpha,
                                'Reject H0?': bonferroni,
                                'No Bonferroni reject H0': zerofive})
alpha_results_df

# %% [code]
#cage line BALB/c_line4
faith_l4_4 = alpha_BALB_stefan[alpha_BALB_stefan['cage_line'].isin(['BALB/c_line4']) & alpha_BALB_stefan['animals_per_cage'].isin([4.0])]['PD_whole_tree']
faith_l4_1 = alpha_BALB_stefan[alpha_BALB_stefan['cage_line'].isin(['BALB/c_line4']) & alpha_BALB_stefan['animals_per_cage'].isin([1.0])]['PD_whole_tree']
faith_l4_2 = alpha_BALB_stefan[alpha_BALB_stefan['cage_line'].isin(['BALB/c_line4']) & alpha_BALB_stefan['animals_per_cage'].isin([2.0])]['PD_whole_tree']
faith_l4_3 = alpha_BALB_stefan[alpha_BALB_stefan['cage_line'].isin(['BALB/c_line4']) & alpha_BALB_stefan['animals_per_cage'].isin([3.0])]['PD_whole_tree']

faith_l4_animal = list(itertools.combinations([faith_l4_1, faith_l4_2, faith_l4_3, faith_l4_4],2))
faith_l4_animal_str = list(itertools.combinations(['faith_l4_1', 'faith_l4_2', 'faith_l4_3', 'faith_l4_4'],2))

# %% [code]
alpha = 0.05/len(faith_l4_animal)
count = 0
statistics = []
p_values = []
significant = []
groups = []
bonferroni = []
zerofive = []
for pair in faith_l4_animal:
    mannwhitneyu_table(pair, faith_l4_animal_str, count)
    count +=1

alpha_results_df = pd.DataFrame({'Comparisons': groups,
                                'Statistics': statistics,
                                'p_values': p_values,
                                'Alpha': alpha,
                                'Reject H0?': bonferroni,
                                'No Bonferroni reject H0': zerofive})
alpha_results_df

# %% [code]
#cage line B6_line3
faith_l3_4 = alpha_C57B_stefan[alpha_C57B_stefan['cage_line'].isin(['C57BL6/J_line3']) & alpha_C57B_stefan['animals_per_cage'].isin([4.0])]['PD_whole_tree']
faith_l3_1 = alpha_C57B_stefan[alpha_C57B_stefan['cage_line'].isin(['C57BL6/J_line3']) & alpha_C57B_stefan['animals_per_cage'].isin([1.0])]['PD_whole_tree']
faith_l3_2 = alpha_C57B_stefan[alpha_C57B_stefan['cage_line'].isin(['C57BL6/J_line3']) & alpha_C57B_stefan['animals_per_cage'].isin([2.0])]['PD_whole_tree']
faith_l3_3 = alpha_C57B_stefan[alpha_C57B_stefan['cage_line'].isin(['C57BL6/J_line3']) & alpha_C57B_stefan['animals_per_cage'].isin([3.0])]['PD_whole_tree']

faith_l3_animal = list(itertools.combinations([faith_l3_1, faith_l3_2, faith_l3_3, faith_l3_4],2))
faith_l3_animal_str = list(itertools.combinations(['faith_l3_1', 'faith_l3_2', 'faith_l3_3', 'faith_l3_4'],2))

# %% [code]
alpha = 0.05/len(faith_l3_animal)
count = 0
statistics = []
p_values = []
significant = []
groups = []
bonferroni = []
zerofive = []
for pair in faith_l3_animal:
    mannwhitneyu_table(pair, faith_l3_animal_str, count)
    count +=1

alpha_results_df = pd.DataFrame({'Comparisons': groups,
                                'Statistics': statistics,
                                'p_values': p_values,
                                'Alpha': alpha,
                                'Reject H0?': bonferroni,
                                'No Bonferroni reject H0': zerofive})
alpha_results_df

# %% [code]
#cage line B6_line2
faith_l2_4 = alpha_C57B_stefan[alpha_C57B_stefan['cage_line'].isin(['C57BL6/J_line2']) & alpha_C57B_stefan['animals_per_cage'].isin([4.0])]['PD_whole_tree']
faith_l2_1 = alpha_C57B_stefan[alpha_C57B_stefan['cage_line'].isin(['C57BL6/J_line2']) & alpha_C57B_stefan['animals_per_cage'].isin([1.0])]['PD_whole_tree']
faith_l2_2 = alpha_C57B_stefan[alpha_C57B_stefan['cage_line'].isin(['C57BL6/J_line2']) & alpha_C57B_stefan['animals_per_cage'].isin([2.0])]['PD_whole_tree']
faith_l2_3 = alpha_C57B_stefan[alpha_C57B_stefan['cage_line'].isin(['C57BL6/J_line2']) & alpha_C57B_stefan['animals_per_cage'].isin([3.0])]['PD_whole_tree']
faith_l2_5 = alpha_C57B_stefan[alpha_C57B_stefan['cage_line'].isin(['C57BL6/J_line2']) & alpha_C57B_stefan['animals_per_cage'].isin([5.0])]['PD_whole_tree']


faith_l2_animal = list(itertools.combinations([faith_l2_1, faith_l2_2, faith_l2_3, faith_l2_4, faith_l2_5],2))
faith_l2_animal_str = list(itertools.combinations(['faith_l2_1', 'faith_l2_2', 'faith_l2_3', 'faith_l2_4', 'faith_l2_5'],2))

# %% [code]
alpha = 0.05/len(faith_l2_animal)
count = 0
statistics = []
p_values = []
significant = []
groups = []
bonferroni = []
zerofive = []
for pair in faith_l2_animal:
    mannwhitneyu_table(pair, faith_l2_animal_str, count)
    count +=1

alpha_results_df = pd.DataFrame({'Comparisons': groups,
                                'Statistics': statistics,
                                'p_values': p_values,
                                'Alpha': alpha,
                                'Reject H0?': bonferroni,
                                'No Bonferroni reject H0': zerofive})
alpha_results_df

# %% [code]
#cage line B6_line1
faith_l1_4 = alpha_C57B_stefan[alpha_C57B_stefan['cage_line'].isin(['C57BL6/J_line1']) & alpha_C57B_stefan['animals_per_cage'].isin([4.0])]['PD_whole_tree']
faith_l1_1 = alpha_C57B_stefan[alpha_C57B_stefan['cage_line'].isin(['C57BL6/J_line1']) & alpha_C57B_stefan['animals_per_cage'].isin([1.0])]['PD_whole_tree']
faith_l1_2 = alpha_C57B_stefan[alpha_C57B_stefan['cage_line'].isin(['C57BL6/J_line1']) & alpha_C57B_stefan['animals_per_cage'].isin([2.0])]['PD_whole_tree']
faith_l1_3 = alpha_C57B_stefan[alpha_C57B_stefan['cage_line'].isin(['C57BL6/J_line1']) & alpha_C57B_stefan['animals_per_cage'].isin([3.0])]['PD_whole_tree']

faith_l1_animal = list(itertools.combinations([faith_l1_1, faith_l1_2, faith_l1_3, faith_l1_4],2))
faith_l1_animal_str = list(itertools.combinations(['faith_l1_1', 'faith_l1_2', 'faith_l1_3', 'faith_l1_4'],2))

# %% [code]
alpha = 0.05/len(faith_l1_animal)
count = 0
statistics = []
p_values = []
significant = []
groups = []
bonferroni = []
zerofive = []
for pair in faith_l1_animal:
    mannwhitneyu_table(pair, faith_l1_animal_str, count)
    count +=1

alpha_results_df = pd.DataFrame({'Comparisons': groups,
                                'Statistics': statistics,
                                'p_values': p_values,
                                'Alpha': alpha,
                                'Reject H0?': bonferroni,
                                'No Bonferroni reject H0': zerofive})
alpha_results_df

# %% [code]


# %% [markdown]
### Sebum

# %% [markdown]
##### Differences within the comparisons

# %% [code]
alpha_C57B_comp_sebum = alpha_results_merged_sebum[alpha_results_merged_sebum['genotype'].isin(['C57BL6/J', 'B6CF1'])]
alpha_BALB_comp_sebum = alpha_results_merged_sebum[alpha_results_merged_sebum['genotype'].isin(['BALB/c', 'B6CF1'])]

# %% [code]
Janvier_faith_C57B_sebum = alpha_C57B_comp_sebum[(alpha_C57B_comp_sebum['comparison'].isin([1])) & (alpha_C57B_comp_sebum['geo_loc_name'].isin(['France:Janvier']))]['PD_whole_tree']
Duess_faith_C57B_sebum = alpha_C57B_comp_sebum[alpha_C57B_comp_sebum['comparison'].isin([1]) & (alpha_C57B_comp_sebum['geo_loc_name'].isin(['Germany:Duesseldorf']))]['PD_whole_tree']
exp_faith_C57B_sebum = alpha_C57B_comp_sebum[alpha_C57B_comp_sebum['comparison'].isin([1])]['PD_whole_tree']

faith_C57B_groups_comp_sebum = list(itertools.combinations([Janvier_faith_C57B_comp, Duess_faith_C57B_comp, exp_faith_C57B_comp],2))
faith_C57B_groups_comp_sebum_str = list(itertools.combinations(["Janvier_faith_C57B", "Duess_faith_C57B", "exp_faith_C57B"],2))

# %% [code]
alpha = 0.05/len(faith_C57B_groups_comp_sebum)
count = 0
statistics = []
p_values = []
significant = []
groups = []
bonferroni = []
zerofive = []
for pair in faith_C57B_groups_comp_sebum:
    mannwhitneyu_table(pair, faith_C57B_groups_comp_sebum_str, count)
    count +=1

# %% [code]
alpha_results_df = pd.DataFrame({'Comparisons': groups,
                                'Statistics': statistics,
                                'p_values': p_values,
                                'Alpha': alpha,
                                'Reject H0?': bonferroni,
                                'No Bonferroni reject H0': zerofive})
alpha_results_df

# %% [code]
#BALB/c

# %% [code]
Janvier_faith_BALB_comp = alpha_BALB_comp_sebum[(alpha_BALB_comp_sebum['comparison'].isin([1])) & (alpha_BALB_comp_sebum['geo_loc_name'].isin(['France:Janvier']))]['PD_whole_tree']
Duess_faith_BALB_comp = alpha_BALB_comp_sebum[alpha_BALB_comp_sebum['comparison'].isin([1]) & (alpha_BALB_comp_sebum['geo_loc_name'].isin(['Germany:Duesseldorf']))]['PD_whole_tree']
exp_faith_BALB_comp = alpha_BALB_comp_sebum[alpha_BALB_comp_sebum['comparison'].isin([1])]['PD_whole_tree']

faith_BALB_groups_comp_sebum = list(itertools.combinations([Janvier_faith_BALB_comp, Duess_faith_BALB_comp, exp_faith_BALB_comp],2))
faith_BALB_groups_comp_sebum_str = list(itertools.combinations(["Janvier_faith_BALB", "Duess_faith_BALB", "exp_faith_BALB"],2))

# %% [code]
alpha = 0.05/len(faith_BALB_groups_comp_sebum)
count = 0
statistics = []
p_values = []
significant = []
groups = []
bonferroni = []
zerofive = []
for pair in faith_BALB_groups_comp_sebum:
    mannwhitneyu_table(pair, faith_BALB_groups_comp_sebum_str, count)
    count +=1

# %% [code]
alpha_results_df = pd.DataFrame({'Comparisons': groups,
                                'Statistics': statistics,
                                'p_values': p_values,
                                'Alpha': alpha,
                                'Reject H0?': bonferroni,
                                'No Bonferroni reject H0': zerofive})
alpha_results_df

# %% [markdown]
##### Differences in Alpha Diversity per Genotype between generations

# %% [code]
alpha_C57B = alpha_results_merged_sebum[alpha_results_merged_sebum['genotype'].isin(['C57BL6/J', 'B6CF1'])]
alpha_BALB = alpha_results_merged_sebum[alpha_results_merged_sebum['genotype'].isin(['BALB/c', 'B6CF1'])]

# %% [code]
F1_faith_C57B = alpha_C57B[alpha_C57B['generation'].isin(['F1'])]['PD_whole_tree']
F2_faith_C57B = alpha_C57B[alpha_C57B['generation'].isin(['F2'])]['PD_whole_tree']
F3_faith_C57B = alpha_C57B[alpha_C57B['generation'].isin(['F3'])]['PD_whole_tree']
F4_faith_C57B = alpha_C57B[alpha_C57B['generation'].isin(['F4'])]['PD_whole_tree']
F5_faith_C57B = alpha_C57B[alpha_C57B['generation'].isin(['F5'])]['PD_whole_tree']
P1_faith_C57B = alpha_C57B[alpha_C57B['generation'].isin(['P1'])]['PD_whole_tree']
RM_faith_C57B = alpha_C57B[alpha_C57B['generation'].isin(['RM'])]['PD_whole_tree']

faith_C57B_groups = list(itertools.combinations([RM_faith_C57B, P1_faith_C57B, F1_faith_C57B, F2_faith_C57B, F3_faith_C57B, F4_faith_C57B, F5_faith_C57B],2))
faith_C57B_groups_str = list(itertools.combinations(['RM_faith_C57B','P1_faith_C57B', 'F1_faith_C57B', 'F2_faith_C57B', 'F3_faith_C57B', 'F4_faith_C57B', 'F5_faith_C57B'],2))

# %% [code]
F1_faith_BALB = alpha_BALB[alpha_BALB['generation'].isin(['F1'])]['PD_whole_tree']
F2_faith_BALB = alpha_BALB[alpha_BALB['generation'].isin(['F2'])]['PD_whole_tree']
F3_faith_BALB = alpha_BALB[alpha_BALB['generation'].isin(['F3'])]['PD_whole_tree']
F4_faith_BALB = alpha_BALB[alpha_BALB['generation'].isin(['F4'])]['PD_whole_tree']
F5_faith_BALB = alpha_BALB[alpha_BALB['generation'].isin(['F5'])]['PD_whole_tree']
P1_faith_BALB = alpha_BALB[alpha_BALB['generation'].isin(['P1'])]['PD_whole_tree']
RM_faith_BALB = alpha_BALB[alpha_BALB['generation'].isin(['RM'])]['PD_whole_tree']

faith_BALB_groups = list(itertools.combinations([RM_faith_BALB, P1_faith_BALB, F1_faith_BALB, F2_faith_BALB, F3_faith_BALB, F4_faith_BALB, F5_faith_BALB],2))
faith_BALB_groups_str = list(itertools.combinations(['RM_faith_BALB','P1_faith_BALB', 'F1_faith_BALB', 'F2_faith_BALB', 'F3_faith_BALB', 'F4_faith_BALB', 'F5_faith_BALB'],2))

# %% [code]
alpha = 0.05/len(faith_C57B_groups)
count = 0
statistics = []
p_values = []
significant = []
groups = []
bonferroni = []
zerofive = []
for pair in faith_C57B_groups:
    mannwhitneyu_table(pair, faith_C57B_groups_str, count)
    count +=1

# %% [code]
alpha = 0.05/len(faith_BALB_groups)
count = 0
for pair in faith_BALB_groups:
    mannwhitneyu_table(pair, faith_BALB_groups_str, count)
    count +=1

# %% [code]
alpha_results_df = pd.DataFrame({'Comparisons': groups,
                                'Statistics': statistics,
                                'p_values': p_values,
                                'Alpha': alpha,
                                'Reject H0?': bonferroni,
                                'No Bonferroni reject H0': zerofive})
alpha_results_df

# %% [markdown]
##### Differences in Alpha Diversity per Generation between genotypes

# %% [code]
generation_list_alpha = [(RM_faith_C57B, RM_faith_BALB),
                        (P1_faith_C57B, P1_faith_BALB),
                        (F1_faith_C57B, F1_faith_BALB),
                        (F2_faith_C57B, F2_faith_BALB),
                        (F3_faith_C57B, F3_faith_BALB),
                        (F4_faith_C57B, F4_faith_BALB),
                        (F5_faith_C57B, F5_faith_BALB)]
generation_list_alpha_str = [('RM_faith_C57B', 'RM_faith_BALB'),
                        ('P1_faith_C57B', 'P1_faith_BALB'),
                        ('F1_faith_C57B', 'F1_faith_BALB'),
                        ('F2_faith_C57B', 'F2_faith_BALB'),
                        ('F3_faith_C57B', 'F3_faith_BALB'),
                        ('F4_faith_C57B', 'F4_faith_BALB'),
                        ('F5_faith_C57B', 'F5_faith_BALB')]
statistics = []
p_values = []
significant = []
groups = []
bonferroni = []
zerofive = []
generations=6
alpha = 0.05/generations
count = 0
for pair in generation_list_alpha:
    mannwhitneyu_table(pair, generation_list_alpha_str, count)
    count += 1

# %% [code]
alpha_results_gen_df = pd.DataFrame({'Comparisons': groups,
                                'Statistics': statistics,
                                'p_values': p_values,
                                'Alpha': alpha,
                                'Reject H0: x<=y?': bonferroni,
                                'No Bonferroni reject H0': zerofive})
alpha_results_gen_df

# %% [markdown]
##### cage groups

# %% [code]
faith_B1 = alpha_BALB[alpha_BALB['cage_lineage'].isin(['C1'])]['PD_whole_tree']
faith_B2 = alpha_BALB[alpha_BALB['cage_lineage'].isin(['C2'])]['PD_whole_tree']
faith_B3 = alpha_BALB[alpha_BALB['cage_lineage'].isin(['C3'])]['PD_whole_tree']

faith_BALB_Cage = list(itertools.combinations([faith_B1, faith_B2, faith_B3],2))
faith_BALB_Cage_str = list(itertools.combinations(['faith_B1', 'faith_B2', 'faith_B3'],2))

# %% [code]
alpha = 0.05/len(faith_BALB_Cage)
count = 0
statistics = []
p_values = []
significant = []
groups = []
bonferroni = []
zerofive = []
for pair in faith_BALB_Cage:
    mannwhitneyu_table(pair, faith_BALB_Cage_str, count)
    count +=1

alpha_results_df = pd.DataFrame({'Comparisons': groups,
                                'Statistics': statistics,
                                'p_values': p_values,
                                'Alpha': alpha,
                                'Reject H0?': bonferroni,
                                'No Bonferroni reject H0': zerofive})
alpha_results_df

# %% [code]
faith_C1 = alpha_C57B[alpha_C57B['cage_lineage'].isin(['B1'])]['PD_whole_tree']
faith_C2 = alpha_C57B[alpha_C57B['cage_lineage'].isin(['B2'])]['PD_whole_tree']
faith_C3 = alpha_C57B[alpha_C57B['cage_lineage'].isin(['B3'])]['PD_whole_tree']

faith_C57B_Cage = list(itertools.combinations([faith_C1, faith_C2, faith_C3],2))
faith_C57B_Cage_str = list(itertools.combinations(['faith_C1', 'faith_C2', 'faith_C3'],2))

# %% [code]
alpha = 0.05/len(faith_C57B_Cage)
count = 0
statistics = []
p_values = []
significant = []
groups = []
bonferroni = []
zerofive = []
for pair in faith_C57B_Cage:
    mannwhitneyu_table(pair, faith_C57B_Cage_str, count)
    count +=1

alpha_results_df = pd.DataFrame({'Comparisons': groups,
                                'Statistics': statistics,
                                'p_values': p_values,
                                'Alpha': alpha,
                                'Reject H0?': bonferroni,
                                'No Bonferroni reject H0': zerofive})
alpha_results_df

# %% [code]


# %% [markdown]
## Effect Sizes, Forward Step Redundancy analysis
### Gut

# %% [code]
conditions = [
    (meta["cage_lineage"] == "B1"),
    (meta["cage_lineage"] == "B2"),
    (meta["cage_lineage"] == "B3"),
    (meta["cage_lineage"] == "C1"),
    (meta["cage_lineage"] == "C2"),
    (meta["cage_lineage"] == "C3"),
]

values = ["A", "B", "C", "A", "B", "C"]
 
meta['independent_cage_lineage']= np.select(conditions, values, "other")

# %% [markdown]
#### BALB

# %% [code]
%matplotlib inline
#categorials
categorials = ['independent_cage_lineage', 'sex']

#ordinals
ordinals = {'generation': ['P1', 'F1', 'F2', 'F3', 'F4', 'F5']}

#intervals
intervals = []

#dates
dates = {}

#metrics
metrics_alpha = ['PD_whole_tree', 'shannon', 'observed_features']
metrics_beta = ['bray_curtis', 'unweighted_unifrac', 'weighted_unifrac']

#max_rows = max(len(alpha_div_gut['results'].keys()), len(gut_beta_div['results'].keys()))
#fig, axes = plt.subplots(max_rows, 2, figsize=(5*2, 5*max_rows))
for i, metric in enumerate(metrics_alpha):
    alpha=redundancy_analysis_alpha(meta[meta['genotype'].isin(['BALB/c']) & meta['comparison'].isin([0])],
                             alpha_div_gut['results'][metric],
                             categorials=categorials,
                             ordinals=ordinals,
                             intervals=intervals,
                             dates=dates,
                             #ax=axes[i][0],
                             title='alpha %s' % metric,
                             dry=False,
                             wait=True,
                             use_grid=True,
                             dirty=True,
                             nocache=True)
    plt.savefig(work_dir + "/gut_redundancies_seperate_%s.png" % metric, transparent=False, bbox_inches='tight')
for i, metric in enumerate(metrics_beta):
    beta=redundancy_analysis_beta(meta[meta['genotype'].isin(['BALB/c']) & meta['comparison'].isin([0])],
                             res_beta_gut_no_comp['results'][metric],
                             metric,
                             categorials=categorials,
                             ordinals=ordinals,
                             intervals=intervals,
                             dates=dates,
                             #ax=axes[i][1],
                             title='beta %s' % metric,
                             dry=False,
                             wait=True,
                             use_grid=True,
                             dirty=True,
                             nocache=True)
    plt.savefig(work_dir + "/gut_redundancies_seperate_%s.png" % metric, transparent=False, bbox_inches='tight')

# %% [code]
%matplotlib inline
#categorials
categorials = ['independent_cage_lineage', 'sex']

#ordinals
ordinals = {'generation': ['P1', 'F1', 'F2', 'F3', 'F4', 'F5']}

#intervals
intervals = []

#dates
dates = {}

#metrics
metrics_alpha = ['PD_whole_tree', 'shannon', 'observed_features']
metrics_beta = ['bray_curtis', 'unweighted_unifrac', 'weighted_unifrac']

#max_rows = max(len(alpha_div_gut['results'].keys()), len(gut_beta_div['results'].keys()))
#fig, axes = plt.subplots(max_rows, 2, figsize=(5*2, 5*max_rows))
for i, metric in enumerate(metrics_alpha):
    alpha=redundancy_analysis_alpha(meta[meta['genotype'].isin(['C57BL6/J']) & meta['comparison'].isin([0])],
                             alpha_div_gut['results'][metric],
                             categorials=categorials,
                             ordinals=ordinals,
                             intervals=intervals,
                             dates=dates,
                             #ax=axes[i][0],
                             title='alpha %s' % metric,
                             dry=False,
                             wait=True,
                             use_grid=True,
                             dirty=True,
                             nocache=True)
    plt.savefig(work_dir + "/gut_redundancies_seperate_B6_%s.png" % metric, transparent=False, bbox_inches='tight')
for i, metric in enumerate(metrics_beta):
    beta=redundancy_analysis_beta(meta[meta['genotype'].isin(['C57BL6/J']) & meta['comparison'].isin([0])],
                             res_beta_gut_no_comp['results'][metric],
                             metric,
                             categorials=categorials,
                             ordinals=ordinals,
                             intervals=intervals,
                             dates=dates,
                             #ax=axes[i][1],
                             title='beta %s' % metric,
                             dry=False,
                             wait=True,
                             use_grid=True,
                             dirty=True,
                             nocache=True)
    plt.savefig(work_dir + "/gut_redundancies_seperate_B6_%s.png" % metric, transparent=False, bbox_inches='tight')

# %% [markdown]
#### All

# %% [code]
%matplotlib inline
#categorials
categorials = ['genotype', 'independent_cage_lineage', 'sex']

#ordinals
ordinals = {'generation': ['P1', 'F1', 'F2', 'F3', 'F4', 'F5']}

#intervals
intervals = []

#dates
dates = {}

#metrics
metrics_alpha = ['PD_whole_tree', 'shannon', 'observed_features']
metrics_beta = ['bray_curtis', 'unweighted_unifrac', 'weighted_unifrac']

#max_rows = max(len(alpha_div_gut['results'].keys()), len(gut_beta_div['results'].keys()))
#fig, axes = plt.subplots(max_rows, 2, figsize=(5*2, 5*max_rows))
for i, metric in enumerate(metrics_alpha):
    alpha=redundancy_analysis_alpha(meta[meta['comparison'].isin([0])],
                             alpha_div_gut['results'][metric],
                             categorials=categorials,
                             ordinals=ordinals,
                             intervals=intervals,
                             dates=dates,
                             #ax=axes[i][0],
                             title='alpha %s' % metric,
                             dry=False,
                             wait=True,
                             use_grid=True,
                             dirty=True,
                             nocache=True)
    plt.savefig(work_dir + "/gut_redundancies_seperate_all_%s.png" % metric, transparent=False, bbox_inches='tight')
for i, metric in enumerate(metrics_beta):
    beta=redundancy_analysis_beta(meta[meta['comparison'].isin([0])],
                             res_beta_gut_no_comp['results'][metric],
                             metric,
                             categorials=categorials,
                             ordinals=ordinals,
                             intervals=intervals,
                             dates=dates,
                             #ax=axes[i][1],
                             title='beta %s' % metric,
                             dry=False,
                             wait=True,
                             use_grid=True,
                             dirty=True,
                             nocache=True)
    plt.savefig(work_dir + "/gut_redundancies_seperate_all_%s.png" % metric, transparent=False, bbox_inches='tight')

# %% [code]
alpha['results']

# %% [markdown]
### Sebum

# %% [code]
%matplotlib inline
#categorials
categorials = ['genotype', 'independent_cage_lineage', 'sex']

#ordinals
ordinals = {'generation': ['P1', 'F1', 'F2', 'F3', 'F4', 'F5']}

#intervals
intervals = []

#dates
dates = {}

#metrics
metrics_alpha = ['PD_whole_tree', 'shannon', 'observed_features']
metrics_beta = ['bray_curtis', 'unweighted_unifrac', 'weighted_unifrac']

#max_rows = max(len(alpha_div_sebum['results'].keys()), len(sebum_beta_div['results'].keys()))
#fig, axes = plt.subplots(max_rows, 2, figsize=(5*2, 5*max_rows))
for i, metric in enumerate(metrics_alpha):
    redundancy_analysis_alpha(meta[meta['comparison'].isin([0])],
                             alpha_div_sebum['results'][metric],
                             categorials=categorials,
                             ordinals=ordinals,
                             intervals=intervals,
                             dates=dates,
                             #ax=axes[i][0],
                             title='alpha %s' % metric,
                             dry=False,
                             wait=True,
                             use_grid=True)
    plt.savefig(work_dir + "/sebum_redundancies_seperate_all_%s.png" % metric, transparent=False, bbox_inches='tight')
for i, metric in enumerate(metrics_beta):
    redundancy_analysis_beta(meta[meta['comparison'].isin([0])],
                             res_beta_sebum_no_comp['results'][metric],
                             metric,
                             categorials=categorials,
                             ordinals=ordinals,
                             intervals=intervals,
                             dates=dates,
                             #ax=axes[i][1],
                             title='beta %s' % metric,
                             dry=False,
                             wait=True,
                             use_grid=True)
    plt.savefig(work_dir + "/sebum_redundancies_seperate_all_%s.png" % metric, transparent=False, bbox_inches='tight')

# %% [code]


# %% [code]
%matplotlib inline
#categorials
categorials = ['independent_cage_lineage', 'sex']

#ordinals
ordinals = {'generation': ['P1', 'F1', 'F2', 'F3', 'F4', 'F5']}

#intervals
intervals = []

#dates
dates = {}

#metrics
metrics_alpha = ['PD_whole_tree', 'shannon', 'observed_features']
metrics_beta = ['bray_curtis', 'unweighted_unifrac', 'weighted_unifrac']

#max_rows = max(len(alpha_div_gut['results'].keys()), len(gut_beta_div['results'].keys()))
#fig, axes = plt.subplots(max_rows, 2, figsize=(5*2, 5*max_rows))
for i, metric in enumerate(metrics_alpha):
    alpha=redundancy_analysis_alpha(meta[meta['genotype'].isin(['BALB/c']) & meta['comparison'].isin([0])],
                             alpha_div_sebum['results'][metric],
                             categorials=categorials,
                             ordinals=ordinals,
                             intervals=intervals,
                             dates=dates,
                             #ax=axes[i][0],
                             title='alpha %s' % metric,
                             dry=False,
                             wait=True,
                             use_grid=True,
                             dirty=True,
                             nocache=True)
    plt.savefig(work_dir + "/gut_redundancies_seperate_BALB_%s.png" % metric, transparent=False, bbox_inches='tight')
for i, metric in enumerate(metrics_beta):
    beta=redundancy_analysis_beta(meta[meta['genotype'].isin(['BALB/c']) & meta['comparison'].isin([0])],
                             res_beta_sebum_no_comp['results'][metric],
                             metric,
                             categorials=categorials,
                             ordinals=ordinals,
                             intervals=intervals,
                             dates=dates,
                             #ax=axes[i][1],
                             title='beta %s' % metric,
                             dry=False,
                             wait=True,
                             use_grid=True,
                             dirty=True,
                             nocache=True)
    plt.savefig(work_dir + "/gut_redundancies_seperate_BALB_%s.png" % metric, transparent=False, bbox_inches='tight')

# %% [code]
%matplotlib inline
#categorials
categorials = ['independent_cage_lineage', 'sex']

#ordinals
ordinals = {'generation': ['P1', 'F1', 'F2', 'F3', 'F4', 'F5']}

#intervals
intervals = []

#dates
dates = {}

#metrics
metrics_alpha = ['PD_whole_tree', 'shannon', 'observed_features']
metrics_beta = ['bray_curtis', 'unweighted_unifrac', 'weighted_unifrac']

#max_rows = max(len(alpha_div_gut['results'].keys()), len(gut_beta_div['results'].keys()))
#fig, axes = plt.subplots(max_rows, 2, figsize=(5*2, 5*max_rows))
for i, metric in enumerate(metrics_alpha):
    alpha=redundancy_analysis_alpha(meta[meta['genotype'].isin(['C57BL6/J']) & meta['comparison'].isin([0])],
                             alpha_div_gut['results'][metric],
                             categorials=categorials,
                             ordinals=ordinals,
                             intervals=intervals,
                             dates=dates,
                             #ax=axes[i][0],
                             title='alpha %s' % metric,
                             dry=False,
                             wait=True,
                             use_grid=True,
                             dirty=True,
                             nocache=True)
    plt.savefig(work_dir + "/gut_redundancies_seperate_B6_%s.png" % metric, transparent=False, bbox_inches='tight')
for i, metric in enumerate(metrics_beta):
    beta=redundancy_analysis_beta(meta[meta['genotype'].isin(['C57BL6/J']) & meta['comparison'].isin([0])],
                             res_beta_sebum_no_comp['results'][metric],
                             metric,
                             categorials=categorials,
                             ordinals=ordinals,
                             intervals=intervals,
                             dates=dates,
                             #ax=axes[i][1],
                             title='beta %s' % metric,
                             dry=False,
                             wait=True,
                             use_grid=True,
                             dirty=True,
                             nocache=True)
    plt.savefig(work_dir + "/gut_redundancies_seperate_B6_%s.png" % metric, transparent=False, bbox_inches='tight')

# %% [code]


# %% [markdown]
## Source Tracking
### Gut

# %% [code]


# %% [code]
for sink_group in ["RM"]:
    meta[meta['genotype'].isin(["BALB/c"])]
    conditions = (
        (meta["cage_group"] == "PC-1") | (meta["cage_group"] == "PC-2") | (meta["cage_group"] == "PB-1") | (meta["cage_group"] == "PB-2") | (meta["cage_group"] == "PB-3"),
        meta["generation"] == sink_group
    )
    values = ["Source", "Sink"]
    column_name ="source_sink_P1-%s" % str(sink_group)
    meta[column_name] = np.select(conditions, values, "None")

# %% [code]
P1_RM_source_tracking = sourcetracker2(
    mice_no_comp_gut, #FeatureTable
    meta,
    col_envname="cage_group",
    col_type="source_sink_P1-RM",
    dry=False,
    wait=True
    )

# %% [code]
%matplotlib inline
ax= P1_RM_source_tracking["results"].transpose().plot.barh(stacked=True)
plt.xlabel('relative impact')   
plt.title(f'Impact on RM by P1 generation')
plt.legend(bbox_to_anchor=(1.05, 1), loc=2)
plt.savefig(work_dir + "/source_tracker_RM_P1.png", transparent=False, bbox_inches='tight')

# %% [markdown]
#### BALB/c
##### Impact of CF1 cage groups on CF2 cage groups

# %% [code]
for sink_group in ["CF2-1", "CF2-2"]:
    meta[meta['genotype'].isin(["BALB/c"])]
    conditions = (
        (meta["cage_group"] == "CF1-1") | (meta["cage_group"] == "CF1-2"),
        meta["cage_group"] == sink_group
    )
    values = ["Source", "Sink"]
    column_name ="source_sink_CF1-%s" % str(sink_group)
    meta[column_name] = np.select(conditions, values, "None")

# %% [code]
CF1_2_1_source_tracking = sourcetracker2(
    mice_no_comp_gut, #FeatureTable
    meta,
    col_envname="cage_group",
    col_type="source_sink_CF1-CF2-1",
    dry=False,
    wait=True
    )
CF1_2_2_source_tracking = sourcetracker2(
    mice_no_comp_gut, #FeatureTable
    meta,
    col_envname="cage_group",
    col_type="source_sink_CF1-CF2-2",
    dry=False,
    wait=True
    )

# %% [code]
%matplotlib inline
for i,group in enumerate([CF1_2_1_source_tracking, CF1_2_2_source_tracking], 1):
    ax= group["results"].transpose().plot.barh(stacked=True)
    if i == 1:
        for tick in ax.get_yticklabels():
            cage_ID = meta.loc[tick.get_text()]["cage_id"]
            if cage_ID in ("F2-17"):
                tick.set_color('green')
            elif cage_ID in ("F2-18"):
                tick.set_color('blue')
            elif cage_ID in ("F2-19"):
                tick.set_color('darkred')
            elif cage_ID in ("F2-20"):
                tick.set_color('magenta')
            else:
                tick.set_color('black')
        
    else:
        for tick in ax.get_yticklabels():
            cage_ID = meta.loc[tick.get_text()]["cage_id"]
            if cage_ID in ("F2-21"):
                tick.set_color('green')
            elif cage_ID in ("F2-22"):
                tick.set_color('blue')
            elif cage_ID in ("F2-23"):
                tick.set_color('darkred')
            elif cage_ID in ("F2-24"):
                tick.set_color('magenta')
            elif cage_ID in ("F2-25"):
                tick.set_color('teal')
            else:
                tick.set_color('black')
    plt.xlabel('relative impact')   
    plt.title(f'Impact on CF2-{i} by CF1 generation')
    plt.legend(bbox_to_anchor=(1.05, 1), loc=2)
    plt.savefig(work_dir + "/source_tracker_%s.png" %i, transparent=False, bbox_inches='tight')

# %% [markdown]
##### Impact of CF2 cage groups on CF3 cage groups

# %% [code]
for sink_group in ["CF3-1", "CF3-2", "CF3-3"]:
    meta[meta['genotype'].isin(["BALB/c"])]
    conditions = (
        (meta["cage_group"] == "CF2-1") | (meta["cage_group"] == "CF2-2") | (meta["cage_group"] == "CF2-3"),
        meta["cage_group"] == sink_group
    )
    values = ["Source", "Sink"]
    column_name ="source_sink_CF2-" + str(sink_group)
    meta[column_name] = np.select(conditions, values, "None")

# %% [code]
CF2_3_1_source_tracking = sourcetracker2(
    mice_no_comp_gut, #FeatureTable
    meta,
    col_envname="cage_group",
    col_type="source_sink_CF2-CF3-1",
    dry=False,
    wait=True
)

CF2_3_2_source_tracking = sourcetracker2(
    mice_no_comp_gut, #FeatureTable
    meta,
    col_envname="cage_group",
    col_type="source_sink_CF2-CF3-2",
    dry=False,
    wait=True
)

CF2_3_3_source_tracking = sourcetracker2(
    mice_no_comp_gut, #FeatureTable
    meta,
    col_envname="cage_group",
    col_type="source_sink_CF2-CF3-3",
    dry=False,
    wait=True
)

# %% [code]
for i,group in enumerate([CF2_3_1_source_tracking, CF2_3_2_source_tracking, CF2_3_3_source_tracking], 1):
    ax= group["results"].transpose().plot.barh(stacked=True)
    if i == 1:
        for tick in ax.get_yticklabels():
            cage_ID = meta.loc[tick.get_text()]["cage_id"]
            if cage_ID in ("F3-21"):
                tick.set_color('green')
            elif cage_ID in ("F3-22"):
                tick.set_color('blue')
            else:
                tick.set_color('black')
    elif i==2:
        for tick in ax.get_yticklabels():
            cage_ID = meta.loc[tick.get_text()]["cage_id"]
            if cage_ID in ("F3-23"):
                tick.set_color('green')
            elif cage_ID in ("F3-24"):
                tick.set_color('blue')
            elif cage_ID in ("F3-25"):
                tick.set_color('red')
            elif cage_ID in ("F3-26"):
                tick.set_color('magenta')
            elif cage_ID in ("F3-27"):
                tick.set_color('teal')
            else:
                tick.set_color('black')       
    else:
        for tick in ax.get_yticklabels():
            cage_ID = meta.loc[tick.get_text()]["cage_id"]
            if cage_ID in ("F3-28"):
                tick.set_color('green')
            elif cage_ID in ("F3-29"):
                tick.set_color('blue')
            elif cage_ID in ("F3-30"):
                tick.set_color('red')
            elif cage_ID in ("F3-31"):
                tick.set_color('magenta')
            elif cage_ID in ("F3-32"):
                tick.set_color('teal')
            else:
                tick.set_color('black')
    plt.title(f'Impact on CF3-{i} by CF2 generation')
    plt.legend(bbox_to_anchor=(1.05, 1), loc=2)
    plt.savefig(work_dir + "/gut_BALB_source_tracker_%s.png" %i, transparent=False, bbox_inches='tight')

# %% [markdown]
##### Impact of CF3 cage groups on CF4 cage groups

# %% [code]
for sink_group in ["CF4-1", "CF4-2", "CF4-3"]:
    meta[meta['genotype'].isin(["BALB/c"])]
    conditions = (
        (meta["cage_group"] == "CF3-1") | (meta["cage_group"] == "CF3-2") | (meta["cage_group"] == "CF3-3"),
        meta["cage_group"] == sink_group
    )
    values = ["Source", "Sink"]
    column_name ="source_sink_CF3-" + str(sink_group)
    meta[column_name] = np.select(conditions, values, "None")

# %% [code]
CF3_4_1_source_tracking = sourcetracker2(
    mice_no_comp_gut, #FeatureTable
    meta,
    col_envname="cage_group",
    col_type="source_sink_CF3-CF4-1",
    dry=False,
    wait=True
)

CF3_4_2_source_tracking = sourcetracker2(
    mice_no_comp_gut, #FeatureTable
    meta,
    col_envname="cage_group",
    col_type="source_sink_CF3-CF4-2",
    dry=False,
    wait=True
)

CF3_4_3_source_tracking = sourcetracker2(
    mice_no_comp_gut, #FeatureTable
    meta,
    col_envname="cage_group",
    col_type="source_sink_CF3-CF4-3",
    dry=False,
    wait=True
)

# %% [code]
for i,group in enumerate([CF3_4_1_source_tracking, CF3_4_2_source_tracking, CF3_4_3_source_tracking], 1):
    ax= group["results"].transpose().plot.barh(stacked=True)
    if i == 1:
        for tick in ax.get_yticklabels():
            cage_ID = meta.loc[tick.get_text()]["cage_id"]
            if cage_ID in ("F4-19"):
                tick.set_color('green')
            elif cage_ID in ("F4-20"):
                tick.set_color('blue')
            elif cage_ID in ("F4-21"):
                tick.set_color('red')
            elif cage_ID in ("F4-22"):
                tick.set_color('magenta')
            elif cage_ID in ("F4-23"):
                tick.set_color('teal')
            else:
                tick.set_color('black')
    elif i==2:
        for tick in ax.get_yticklabels():
            cage_ID = meta.loc[tick.get_text()]["cage_id"]
            if cage_ID in ("F4-24"):
                tick.set_color('green')
            elif cage_ID in ("F4-25"):
                tick.set_color('blue')
            elif cage_ID in ("F4-26"):
                tick.set_color('red')
            elif cage_ID in ("F4-27"):
                tick.set_color('magenta')
            else:
                tick.set_color('black')       
    else:
        for tick in ax.get_yticklabels():
            cage_ID = meta.loc[tick.get_text()]["cage_id"]
            if cage_ID in ("F4-28"):
                tick.set_color('green')
            elif cage_ID in ("F4-29"):
                tick.set_color('blue')
            elif cage_ID in ("F4-30"):
                tick.set_color('red')
            elif cage_ID in ("F4-31"):
                tick.set_color('magenta')
            else:
                tick.set_color('black')
    plt.title(f'Impact on CF4-{i} by CF3 generation')
    plt.legend(bbox_to_anchor=(1.05, 1), loc=2)

# %% [code]


# %% [markdown]
##### Impact of CF4 cage groups on CF5 cage groups

# %% [code]
meta["cage_group"]

# %% [code]
_5_1_source_tracking = sourcetracker2(
    mice_no_comp_gut, #FeatureTable
    meta,
    col_envname="cage_group",
    col_type="source_sink_CF4-CF5-1",
    dry=False,
    wait=True
)

CF4_5_2_source_tracking = sourcetracker2(
    mice_no_comp_gut, #FeatureTable
    meta,
    col_envname="cage_group",
    col_type="source_sink_CF4-CF5-2",
    dry=False,
    wait=True
)

CF4_5_3_source_tracking = sourcetracker2(
    mice_no_comp_gut, #FeatureTable
    meta,
    col_envname="cage_group",
    col_type="source_sink_CF4-CF5-3",
    dry=False,
    wait=True
)

# %% [code]
for i,group in enumerate([CF4_5_1_source_tracking, CF4_5_2_source_tracking, CF4_5_3_source_tracking], 1):
    ax= group["results"].transpose().plot.barh(stacked=True)
    if i == 1:
        for tick in ax.get_yticklabels():
            cage_ID = meta.loc[tick.get_text()]["cage_id"]
            if cage_ID in ("F5-8"):
                tick.set_color('green')
            elif cage_ID in ("F5-9"):
                tick.set_color('blue')
            elif cage_ID in ("F5-10"):
                tick.set_color('red')
            else:
                tick.set_color('black')
    elif i==2:
        for tick in ax.get_yticklabels():
            cage_ID = meta.loc[tick.get_text()]["cage_id"]
            if cage_ID in ("F5-11"):
                tick.set_color('green')
            elif cage_ID in ("F5-12"):
                tick.set_color('blue')
            elif cage_ID in ("F5-13"):
                tick.set_color('red')
            elif cage_ID in ("F5-14"):
                tick.set_color('magenta')
            else:
                tick.set_color('black')       
    else:
        for tick in ax.get_yticklabels():
            cage_ID = meta.loc[tick.get_text()]["cage_id"]
            if cage_ID in ("F5-15"):
                tick.set_color('green')
            elif cage_ID in ("F5-16"):
                tick.set_color('blue')
            elif cage_ID in ("F5-17"):
                tick.set_color('red')
            elif cage_ID in ("F5-18"):
                tick.set_color('magenta')
            elif cage_ID in ("F5-19"):
                tick.set_color('teal')
            else:
                tick.set_color('black')
    plt.title(f'Impact on CF5-{i} by CF4 generation')
    plt.legend(bbox_to_anchor=(1.05, 1), loc=2)
    plt.savefig(work_dir + "/gut_BALB_source_tracker_%s.png" %i, transparent=False, bbox_inches='tight')

# %% [markdown]
##### Impact of P1 on F generations

# %% [markdown]
###### F1

# %% [code]
for sink_group in ["CF1-1","CF1-2"]:
    meta[meta['genotype'].isin(["BALB/c"])]
    conditions = (
        (meta["cage_group"] == "PC-1") | (meta["cage_group"] == "PC-2"),
        meta["cage_group"] == sink_group
    )
    values = ["Source", "Sink"]
    column_name ="source_sink_PC-%s" % str(sink_group)
    meta[column_name] = np.select(conditions, values, "None")

# %% [code]
PC_CF1_1_source_tracking = sourcetracker2(
    mice_no_comp_gut, #FeatureTable
    meta,
    col_envname="cage_group",
    col_type="source_sink_PC-CF1-1",
    dry=False,
    wait=True
    )
PC_CF1_2_source_tracking = sourcetracker2(
    mice_no_comp_gut, #FeatureTable
    meta,
    col_envname="cage_group",
    col_type="source_sink_PC-CF1-2",
    dry=False,
    wait=True
    )

# %% [code]
meta[meta["cage_group"].isin(["CF1-2"])]["cage_id"]

# %% [code]
%matplotlib inline
for i,group in enumerate([PC_CF1_1_source_tracking, PC_CF1_2_source_tracking], 1):
    ax= group["results"].transpose().plot.barh(stacked=True)
    if i == 1:
        for tick in ax.get_yticklabels():
            cage_ID = meta.loc[tick.get_text()]["cage_id"]
            if cage_ID in ("F1-15"):
                tick.set_color('green')
            elif cage_ID in ("F1-16"):
                tick.set_color('blue')
            elif cage_ID in ("F1-17"):
                tick.set_color('red')
            elif cage_ID in ("F1-18"):
                tick.set_color('magenta')
            elif cage_ID in ("F1-19"):
                tick.set_color('teal')
            else:
                tick.set_color('black')
    else:
        for tick in ax.get_yticklabels():
            cage_ID = meta.loc[tick.get_text()]["cage_id"]
            if cage_ID in ("F1-20"):
                tick.set_color('green')
            elif cage_ID in ("F1-21"):
                tick.set_color('blue')
            elif cage_ID in ("F1-22"):
                tick.set_color('red')
            elif cage_ID in ("F1-23"):
                tick.set_color('magenta')
            else:
                tick.set_color('black')       
    plt.title(f'Impact on CF1-{i} by PC generation')
    plt.legend(bbox_to_anchor=(1.05, 1), loc=2)
    plt.savefig(work_dir + "/gut_BALB_source_tracker_PC-F1%s.png" %i, transparent=False, bbox_inches='tight')

# %% [markdown]
###### F2

# %% [code]
for sink_group in ["CF2-1","CF2-2"]:
    meta[meta['genotype'].isin(["BALB/c"])]
    conditions = (
        (meta["cage_group"] == "PC-1") | (meta["cage_group"] == "PC-2"),
        meta["cage_group"] == sink_group
    )
    values = ["Source", "Sink"]
    column_name ="source_sink_PC-%s" % str(sink_group)
    meta[column_name] = np.select(conditions, values, "None")

# %% [code]
PC_CF2_1_source_tracking = sourcetracker2(
    mice_no_comp_gut, #FeatureTable
    meta,
    col_envname="cage_group",
    col_type="source_sink_PC-CF2-1",
    dry=False,
    wait=True
    )
PC_CF2_2_source_tracking = sourcetracker2(
    mice_no_comp_gut, #FeatureTable
    meta,
    col_envname="cage_group",
    col_type="source_sink_PC-CF2-2",
    dry=False,
    wait=True
    )

# %% [code]
meta[meta["cage_group"].isin(["CF2-2"])]["cage_id"]

# %% [code]
%matplotlib inline
for i,group in enumerate([PC_CF2_1_source_tracking, PC_CF2_2_source_tracking], 1):
    ax= group["results"].transpose().plot.barh(stacked=True)
    if i == 1:
        for tick in ax.get_yticklabels():
            cage_ID = meta.loc[tick.get_text()]["cage_id"]
            if cage_ID in ("F2-17"):
                tick.set_color('green')
            elif cage_ID in ("F2-18"):
                tick.set_color('blue')
            elif cage_ID in ("F2-19"):
                tick.set_color('red')
            elif cage_ID in ("F2-20"):
                tick.set_color('magenta')
            else:
                tick.set_color('black')
    else:
        for tick in ax.get_yticklabels():
            cage_ID = meta.loc[tick.get_text()]["cage_id"]
            if cage_ID in ("F2-21"):
                tick.set_color('green')
            elif cage_ID in ("F2-22"):
                tick.set_color('blue')
            elif cage_ID in ("F2-23"):
                tick.set_color('red')
            elif cage_ID in ("F2-24"):
                tick.set_color('magenta')
            elif cage_ID in ("F2-25"):
                tick.set_color('teal')
            else:
                tick.set_color('black')       
    plt.title(f'Impact on CF2-{i} by PC generation')
    plt.legend(bbox_to_anchor=(1.05, 1), loc=2)
    plt.savefig(work_dir + "/gut_BALB_source_tracker_PC-F2%s.png" %i, transparent=False, bbox_inches='tight')

# %% [markdown]
###### F3

# %% [code]
for sink_group in ["CF3-1","CF3-2","CF3-3"]:
    meta[meta['genotype'].isin(["BALB/c"])]
    conditions = (
        (meta["cage_group"] == "PC-1") | (meta["cage_group"] == "PC-2"),
        meta["cage_group"] == sink_group
    )
    values = ["Source", "Sink"]
    column_name ="source_sink_PC-%s" % str(sink_group)
    meta[column_name] = np.select(conditions, values, "None")

# %% [code]
PC_CF3_1_source_tracking = sourcetracker2(
    mice_no_comp_gut, #FeatureTable
    meta,
    col_envname="cage_group",
    col_type="source_sink_PC-CF3-1",
    dry=False,
    wait=True
    )
PC_CF3_2_source_tracking = sourcetracker2(
    mice_no_comp_gut, #FeatureTable
    meta,
    col_envname="cage_group",
    col_type="source_sink_PC-CF3-2",
    dry=False,
    wait=True
    )
PC_CF3_3_source_tracking = sourcetracker2(
    mice_no_comp_gut, #FeatureTable
    meta,
    col_envname="cage_group",
    col_type="source_sink_PC-CF3-3",
    dry=False,
    wait=True
    )

# %% [code]
meta[meta["cage_group"].isin(["CF3-1"])]["cage_id"]

# %% [code]
%matplotlib inline
for i,group in enumerate([PC_CF3_1_source_tracking, PC_CF3_2_source_tracking, PC_CF3_3_source_tracking], 1):
    ax= group["results"].transpose().plot.barh(stacked=True)
    if i == 1:
        for tick in ax.get_yticklabels():
            cage_ID = meta.loc[tick.get_text()]["cage_id"]
            if cage_ID in ("F3-21"):
                tick.set_color('green')
            elif cage_ID in ("F3-22"):
                tick.set_color('blue')
            else:
                tick.set_color('black')
    elif i==2:
        for tick in ax.get_yticklabels():
            cage_ID = meta.loc[tick.get_text()]["cage_id"]
            if cage_ID in ("F3-23"):
                tick.set_color('green')
            elif cage_ID in ("F3-24"):
                tick.set_color('blue')
            elif cage_ID in ("F3-25"):
                tick.set_color('red')
            elif cage_ID in ("F3-26"):
                tick.set_color('magenta')
            elif cage_ID in ("F3-27"):
                tick.set_color('teal')
            else:
                tick.set_color('black')
    else:
        for tick in ax.get_yticklabels():
            cage_ID = meta.loc[tick.get_text()]["cage_id"]
            if cage_ID in ("F3-28"):
                tick.set_color('green')
            elif cage_ID in ("F3-29"):
                tick.set_color('blue')
            elif cage_ID in ("F3-30"):
                tick.set_color('red')
            elif cage_ID in ("F3-31"):
                tick.set_color('magenta')
            elif cage_ID in ("F3-32"):
                tick.set_color('teal')
            else:
                tick.set_color('black')
    plt.title(f'Impact on CF3-{i} by PC generation')
    plt.legend(bbox_to_anchor=(1.05, 1), loc=2)
    plt.savefig(work_dir + "/gut_BALB_source_tracker_PC-F3%s.png" %i, transparent=False, bbox_inches='tight')

# %% [code]


# %% [markdown]
###### F4

# %% [code]
for sink_group in ["CF4-1","CF4-2","CF4-3"]:
    meta[meta['genotype'].isin(["BALB/c"])]
    conditions = (
        (meta["cage_group"] == "PC-1") | (meta["cage_group"] == "PC-2"),
        meta["cage_group"] == sink_group
    )
    values = ["Source", "Sink"]
    column_name ="source_sink_PC-%s" % str(sink_group)
    meta[column_name] = np.select(conditions, values, "None")

# %% [code]
PC_CF4_1_source_tracking = sourcetracker2(
    mice_no_comp_gut, #FeatureTable
    meta,
    col_envname="cage_group",
    col_type="source_sink_PC-CF4-1",
    dry=False,
    wait=True
    )
PC_CF4_2_source_tracking = sourcetracker2(
    mice_no_comp_gut, #FeatureTable
    meta,
    col_envname="cage_group",
    col_type="source_sink_PC-CF4-2",
    dry=False,
    wait=True
    )
PC_CF4_3_source_tracking = sourcetracker2(
    mice_no_comp_gut, #FeatureTable
    meta,
    col_envname="cage_group",
    col_type="source_sink_PC-CF4-3",
    dry=False,
    wait=True
    )

# %% [code]
meta[meta["cage_group"].isin(["CF4-3"])]["cage_id"]

# %% [code]
%matplotlib inline
for i,group in enumerate([PC_CF4_1_source_tracking, PC_CF4_2_source_tracking, PC_CF4_3_source_tracking], 1):
    ax= group["results"].transpose().plot.barh(stacked=True)
    if i == 1:
        for tick in ax.get_yticklabels():
            cage_ID = meta.loc[tick.get_text()]["cage_id"]
            if cage_ID in ("F4-19"):
                tick.set_color('green')
            elif cage_ID in ("F4-20"):
                tick.set_color('blue')
            elif cage_ID in ("F4-21"):
                tick.set_color('red')
            elif cage_ID in ("F4-22"):
                tick.set_color('magenta')
            elif cage_ID in ("F4-23"):
                tick.set_color('teal')
            else:
                tick.set_color('black')
    elif i==2:
        for tick in ax.get_yticklabels():
            cage_ID = meta.loc[tick.get_text()]["cage_id"]
            if cage_ID in ("F4-24"):
                tick.set_color('green')
            elif cage_ID in ("F4-25"):
                tick.set_color('blue')
            elif cage_ID in ("F4-26"):
                tick.set_color('red')
            elif cage_ID in ("F4-27"):
                tick.set_color('magenta')
            else:
                tick.set_color('black')
    else:
        for tick in ax.get_yticklabels():
            cage_ID = meta.loc[tick.get_text()]["cage_id"]
            if cage_ID in ("F4-28"):
                tick.set_color('green')
            elif cage_ID in ("F4-29"):
                tick.set_color('blue')
            elif cage_ID in ("F4-30"):
                tick.set_color('red')
            elif cage_ID in ("F4-31"):
                tick.set_color('magenta')
            else:
                tick.set_color('black')
    plt.title(f'Impact on CF4-{i} by PC generation')
    plt.legend(bbox_to_anchor=(1.05, 1), loc=2)
    plt.savefig(work_dir + "/gut_BALB_source_tracker_PC-F4%s.png" %i, transparent=False, bbox_inches='tight')

# %% [code]


# %% [markdown]
###### F5

# %% [code]
for sink_group in ["CF5-1","CF5-2","CF5-3"]:
    meta[meta['genotype'].isin(["BALB/c"])]
    conditions = (
        (meta["cage_group"] == "PC-1") | (meta["cage_group"] == "PC-2"),
        meta["cage_group"] == sink_group
    )
    values = ["Source", "Sink"]
    column_name ="source_sink_PC-%s" % str(sink_group)
    meta[column_name] = np.select(conditions, values, "None")

# %% [code]
PC_CF5_1_source_tracking = sourcetracker2(
    mice_no_comp_gut, #FeatureTable
    meta,
    col_envname="cage_group",
    col_type="source_sink_PC-CF5-1",
    dry=False,
    wait=True
    )
PC_CF5_2_source_tracking = sourcetracker2(
    mice_no_comp_gut, #FeatureTable
    meta,
    col_envname="cage_group",
    col_type="source_sink_PC-CF5-2",
    dry=False,
    wait=True
    )
PC_CF5_3_source_tracking = sourcetracker2(
    mice_no_comp_gut, #FeatureTable
    meta,
    col_envname="cage_group",
    col_type="source_sink_PC-CF5-3",
    dry=False,
    wait=True
    )

# %% [code]
meta[meta["cage_group"].isin(["CF5-3"])]["cage_id"]

# %% [code]
%matplotlib inline
for i,group in enumerate([PC_CF5_1_source_tracking, PC_CF5_2_source_tracking, PC_CF5_3_source_tracking], 1):
    ax= group["results"].transpose().plot.barh(stacked=True)
    if i == 1:
        for tick in ax.get_yticklabels():
            cage_ID = meta.loc[tick.get_text()]["cage_id"]
            if cage_ID in ("F5-8"):
                tick.set_color('green')
            elif cage_ID in ("F5-9"):
                tick.set_color('blue')
            elif cage_ID in ("F5-10"):
                tick.set_color('red')
            else:
                tick.set_color('black')
    elif i==2:
        for tick in ax.get_yticklabels():
            cage_ID = meta.loc[tick.get_text()]["cage_id"]
            if cage_ID in ("F5-11"):
                tick.set_color('green')
            elif cage_ID in ("F5-12"):
                tick.set_color('blue')
            elif cage_ID in ("F5-13"):
                tick.set_color('red')
            elif cage_ID in ("F5-14"):
                tick.set_color('magenta')
            else:
                tick.set_color('black')
    else:
        for tick in ax.get_yticklabels():
            cage_ID = meta.loc[tick.get_text()]["cage_id"]
            if cage_ID in ("F5-15"):
                tick.set_color('green')
            elif cage_ID in ("F5-16"):
                tick.set_color('blue')
            elif cage_ID in ("F5-17"):
                tick.set_color('red')
            elif cage_ID in ("F5-18"):
                tick.set_color('magenta')
            elif cage_ID in ("F5-19"):
                tick.set_color('teal')
            else:
                tick.set_color('black')
    plt.title(f'Impact on CF5-{i} by PC generation')
    plt.legend(bbox_to_anchor=(1.05, 1), loc=2)
    plt.savefig(work_dir + "/gut_BALB_source_tracker_PC-F5%s.png" %i, transparent=False, bbox_inches='tight')

# %% [code]


# %% [code]


# %% [markdown]
#### C57BL6/J
##### Impact of BF1 cage groups on BF2 cage groups

# %% [code]


# %% [code]
for sink_group in ["BF2-1", "BF2-2", "BF2-3"]:
    meta[meta['genotype'].isin(["C57BL6/J"])]
    conditions = (
        (meta["cage_group"] == "BF1-1") | (meta["cage_group"] == "BF1-2") | (meta["cage_group"] == "BF1-3"),
        meta["cage_group"] == sink_group
    )
    values = ["Source", "Sink"]
    column_name ="source_sink_BF1-%s" % str(sink_group)
    meta[column_name] = np.select(conditions, values, "None")

# %% [code]
BF1_2_1_source_tracking = sourcetracker2(
    mice_no_comp_gut, #FeatureTaCle
    meta,
    col_envname="cage_group",
    col_type="source_sink_BF1-BF2-1",
    dry=False,
    wait=True
    )
BF1_2_2_source_tracking = sourcetracker2(
    mice_no_comp_gut, #FeatureTable
    meta,
    col_envname="cage_group",
    col_type="source_sink_BF1-BF2-2",
    dry=False,
    wait=True
    )
BF1_2_3_source_tracking = sourcetracker2(
    mice_no_comp_gut, #FeatureTable
    meta,
    col_envname="cage_group",
    col_type="source_sink_BF1-BF2-3",
    dry=False,
    wait=True
    )

# %% [code]
meta.groupby(["source_sink_BF1-BF2-1", "cage_group"]).size()

# %% [code]
for i,group in enumerate([BF1_2_1_source_tracking, BF1_2_2_source_tracking, BF1_2_3_source_tracking], 1):
    ax= group["results"].transpose().plot.barh(stacked=True)
    if i == 1:
        for tick in ax.get_yticklabels():
            cage_ID = meta.loc[tick.get_text()]["cage_id"]
            if cage_ID in ("F2-1"):
                tick.set_color('green')
            elif cage_ID in ("F2-2"):
                tick.set_color('blue')
            elif cage_ID in ("F2-3"):
                tick.set_color('red')
            elif cage_ID in ("F2-4"):
                tick.set_color('magenta')
            elif cage_ID in ("F2-5"):
                tick.set_color('teal')
            elif cage_ID in ("F2-6"):
                tick.set_color('darkolivegreen')
            elif cage_ID in ("F2-7"):
                tick.set_color('darkorange')
            else:
                tick.set_color('black')
    elif i==2:
        for tick in ax.get_yticklabels():
            cage_ID = meta.loc[tick.get_text()]["cage_id"]
            if cage_ID in ("F2-8"):
                tick.set_color('green')
            elif cage_ID in ("F2-9"):
                tick.set_color('blue')
            elif cage_ID in ("F2-10"):
                tick.set_color('red')
            elif cage_ID in ("F2-11"):
                tick.set_color('magenta')
            else:
                tick.set_color('black')       
    else:
        for tick in ax.get_yticklabels():
            cage_ID = meta.loc[tick.get_text()]["cage_id"]
            if cage_ID in ("F2-12"):
                tick.set_color('green')
            elif cage_ID in ("F2-13"):
                tick.set_color('blue')
            elif cage_ID in ("F2-14"):
                tick.set_color('red')
            elif cage_ID in ("F2-15"):
                tick.set_color('magenta')
            elif cage_ID in ("F2-16"):
                tick.set_color('teal')
            else:
                tick.set_color('black')
    plt.title(f'Impact on BF2-{i} by BF1 generation')
    plt.legend(bbox_to_anchor=(1.05, 1), loc=2)

# %% [markdown]
##### Impact of BF2 cage groups on BF3 cage groups

# %% [code]
for sink_group in ["BF3-1", "BF3-2", "BF3-3"]:
    meta[meta['genotype'].isin(["C57BL6/J"])]
    conditions = (
        (meta["cage_group"] == "BF2-1") | (meta["cage_group"] == "BF2-2") | (meta["cage_group"] == "BF2-3"),
        meta["cage_group"] == sink_group
    )
    values = ["Source", "Sink"]
    column_name ="source_sink_BF2-" + str(sink_group)
    meta[column_name] = np.select(conditions, values, "None")

# %% [code]
BF2_3_1_source_tracking = sourcetracker2(
    mice_no_comp_gut, #FeatureTable
    meta,
    col_envname="cage_group",
    col_type="source_sink_BF2-BF3-1",
    dry=False,
    wait=True
)

BF2_3_2_source_tracking = sourcetracker2(
    mice_no_comp_gut, #FeatureTable
    meta,
    col_envname="cage_group",
    col_type="source_sink_BF2-BF3-2",
    dry=False,
    wait=True
)

BF2_3_3_source_tracking = sourcetracker2(
    mice_no_comp_gut, #FeatureTable
    meta,
    col_envname="cage_group",
    col_type="source_sink_BF2-BF3-3",
    dry=False,
    wait=True
)

# %% [code]
for i,group in enumerate([BF2_3_1_source_tracking, BF2_3_2_source_tracking, BF2_3_3_source_tracking], 1):
    ax= group["results"].transpose().plot.barh(stacked=True)
    if i == 1:
        for tick in ax.get_yticklabels():
            cage_ID = meta.loc[tick.get_text()]["cage_id"]
            if cage_ID in ("F3-1"):
                tick.set_color('green')
            elif cage_ID in ("F3-2"):
                tick.set_color('blue')
            elif cage_ID in ("F3-3"):
                tick.set_color('red')
            elif cage_ID in ("F3-4"):
                tick.set_color('magenta')
            elif cage_ID in ("F3-5"):
                tick.set_color('teal')
            elif cage_ID in ("F3-6"):
                tick.set_color('darkorange')
            else:
                tick.set_color('black')
    elif i==2:
        for tick in ax.get_yticklabels():
            cage_ID = meta.loc[tick.get_text()]["cage_id"]
            if cage_ID in ("F3-7"):
                tick.set_color('green')
            elif cage_ID in ("F3-8"):
                tick.set_color('blue')
            elif cage_ID in ("F3-9"):
                tick.set_color('red')
            elif cage_ID in ("F3-10"):
                tick.set_color('magenta')
            elif cage_ID in ("F3-11"):
                tick.set_color('teal')
            elif cage_ID in ("F3-12"):
                tick.set_color('darkolivegreen')
            elif cage_ID in ("F3-13"):
                tick.set_color('darkorange')
            elif cage_ID in ("F3-14"):
                tick.set_color('dodgerblue')
            else:
                tick.set_color('black')       
    else:
        for tick in ax.get_yticklabels():
            cage_ID = meta.loc[tick.get_text()]["cage_id"]
            if cage_ID in ("F3-15"):
                tick.set_color('green')
            elif cage_ID in ("F3-16"):
                tick.set_color('blue')
            elif cage_ID in ("F3-17"):
                tick.set_color('red')
            elif cage_ID in ("F3-18"):
                tick.set_color('magenta')
            elif cage_ID in ("F3-19"):
                tick.set_color('teal')
            elif cage_ID in ("F3-20"):
                tick.set_color('darkorange')
            else:
                tick.set_color('black')
    plt.title(f'Impact on BF3-{i} by BF2 generation')
    plt.legend(bbox_to_anchor=(1.05, 1), loc=2)

# %% [markdown]
##### Impact of BF3 cage groups on BF4 cage groups

# %% [code]
for sink_group in ["BF4-1", "BF4-2", "BF4-3"]:
    meta[meta['genotype'].isin(["C57BL6/J"])]
    conditions = (
        (meta["cage_group"] == "BF3-1") | (meta["cage_group"] == "BF3-2") | (meta["cage_group"] == "BF3-3"),
        meta["cage_group"] == sink_group
    )
    values = ["Source", "Sink"]
    column_name ="source_sink_BF3-" + str(sink_group)
    meta[column_name] = np.select(conditions, values, "None")

# %% [code]
BF3_4_1_source_tracking = sourcetracker2(
    mice_no_comp_gut, #FeatureTable
    meta,
    col_envname="cage_group",
    col_type="source_sink_BF3-BF4-1",
    dry=False,
    wait=True
)

BF3_4_2_source_tracking = sourcetracker2(
    mice_no_comp_gut, #FeatureTable
    meta,
    col_envname="cage_group",
    col_type="source_sink_BF3-BF4-2",
    dry=False,
    wait=True
)

BF3_4_3_source_tracking = sourcetracker2(
    mice_no_comp_gut, #FeatureTable
    meta,
    col_envname="cage_group",
    col_type="source_sink_BF3-BF4-3",
    dry=False,
    wait=True
)

# %% [code]
for i,group in enumerate([BF3_4_1_source_tracking, BF3_4_2_source_tracking, BF3_4_3_source_tracking], 1):
    ax= group["results"].transpose().plot.barh(stacked=True)
    if i == 1:
        for tick in ax.get_yticklabels():
            cage_ID = meta.loc[tick.get_text()]["cage_id"]
            if cage_ID in ("F4-1"):
                tick.set_color('green')
            elif cage_ID in ("F4-2"):
                tick.set_color('blue')
            elif cage_ID in ("F4-3"):
                tick.set_color('red')
            elif cage_ID in ("F4-4"):
                tick.set_color('magenta')
            elif cage_ID in ("F4-5"):
                tick.set_color('teal')
            elif cage_ID in ("F4-6"):
                tick.set_color('darkorange')
            else:
                tick.set_color('black')
    elif i==2:
        for tick in ax.get_yticklabels():
            cage_ID = meta.loc[tick.get_text()]["cage_id"]
            if cage_ID in ("F4-8"):
                tick.set_color('green')
            elif cage_ID in ("F4-9"):
                tick.set_color('blue')
            elif cage_ID in ("F4-10"):
                tick.set_color('red')
            elif cage_ID in ("F4-11"):
                tick.set_color('magenta')
            elif cage_ID in ("F4-12"):
                tick.set_color('teal')
            elif cage_ID in ("F4-13"):
                tick.set_color('darkorange')
            else:
                tick.set_color('black')       
    else:
        for tick in ax.get_yticklabels():
            cage_ID = meta.loc[tick.get_text()]["cage_id"]
            if cage_ID in ("F4-14"):
                tick.set_color('green')
            elif cage_ID in ("F4-15"):
                tick.set_color('blue')
            elif cage_ID in ("F4-16"):
                tick.set_color('red')
            elif cage_ID in ("F4-17"):
                tick.set_color('magenta')
            elif cage_ID in ("F4-18"):
                tick.set_color('teal')
            else:
                tick.set_color('black')
    plt.title(f'Impact on BF4-{i} by BF3 generation')
    plt.legend(bbox_to_anchor=(1.05, 1), loc=2)

# %% [code]


# %% [markdown]
##### Impact of BF4 cage groups on BF5 cage groups

# %% [code]
for sink_group in ["BF5-1", "BF5-2"]:
    meta[meta['genotype'].isin(["C57BL6/J"])]
    conditions = (
        (meta["cage_group"] == "BF4-1") | (meta["cage_group"] == "BF4-2") | (meta["cage_group"] == "BF4-3"),
        meta["cage_group"] == sink_group
    )
    values = ["Source", "Sink"]
    column_name ="source_sink_BF4-" + str(sink_group)
    meta[column_name] = np.select(conditions, values, "None")

# %% [code]
BF4_5_1_source_tracking = sourcetracker2(
    mice_no_comp_gut, #FeatureTable
    meta,
    col_envname="cage_group",
    col_type="source_sink_BF4-BF5-1",
    dry=False,
    wait=True
)

BF4_5_2_source_tracking = sourcetracker2(
    mice_no_comp_gut, #FeatureTable
    meta,
    col_envname="cage_group",
    col_type="source_sink_BF4-BF5-2",
    dry=False,
    wait=True
)

# %% [code]
for i,group in enumerate([BF4_5_1_source_tracking, BF4_5_2_source_tracking], 1):
    ax= group["results"].transpose().plot.barh(stacked=True)
    if i == 1:
        for tick in ax.get_yticklabels():
            cage_ID = meta.loc[tick.get_text()]["cage_id"]
            if cage_ID in ("F5-1"):
                tick.set_color('green')
            elif cage_ID in ("F5-2"):
                tick.set_color('blue')
            elif cage_ID in ("F5-3"):
                tick.set_color('red')
            elif cage_ID in ("F5-4"):
                tick.set_color('magenta')
            else:
                tick.set_color('black')  
    else:
        for tick in ax.get_yticklabels():
            cage_ID = meta.loc[tick.get_text()]["cage_id"]
            if cage_ID in ("F5-5"):
                tick.set_color('green')
            elif cage_ID in ("F5-6"):
                tick.set_color('blue')
            elif cage_ID in ("F5-7"):
                tick.set_color('red')
            else:
                tick.set_color('black')
    plt.title(f'Impact on BF5-{i} by BF4 generation')
    plt.legend(bbox_to_anchor=(1.05, 1), loc=2)

# %% [code]


# %% [markdown]
##### Impact of P1 on F generations

# %% [markdown]
###### F1

# %% [code]
meta["cage_group"].values()

# %% [code]
for sink_group in ["BF1-1","BF1-2","BF1-3"]:
    meta[meta['genotype'].isin(["C57BL6/J"])]
    conditions = (
        (meta["cage_group"] == "PB-1") | (meta["cage_group"] == "PB-2") | (meta["cage_group"] == "PB-3"),
        meta["cage_group"] == sink_group
    )
    values = ["Source", "Sink"]
    column_name ="source_sink_PB-%s" % str(sink_group)
    meta[column_name] = np.select(conditions, values, "None")

# %% [code]
PB_BF1_1_source_tracking = sourcetracker2(
    mice_no_comp_gut, #FeatureTaCle
    meta,
    col_envname="cage_group",
    col_type="source_sink_PB-BF1-1",
    dry=False,
    wait=True
    )
PB_BF1_2_source_tracking = sourcetracker2(
    mice_no_comp_gut, #FeatureTable
    meta,
    col_envname="cage_group",
    col_type="source_sink_PB-BF1-2",
    dry=False,
    wait=True
    )
PB_BF1_3_source_tracking = sourcetracker2(
    mice_no_comp_gut, #FeatureTable
    meta,
    col_envname="cage_group",
    col_type="source_sink_PB-BF1-3",
    dry=False,
    wait=True
    )

# %% [code]
meta[meta["cage_group"].isin(["BF1-3"])]["cage_id"]

# %% [code]
%matplotlib inline
for i,group in enumerate([PB_BF1_1_source_tracking, PB_BF1_2_source_tracking, PB_BF1_3_source_tracking], 1):
    ax= group["results"].transpose().plot.barh(stacked=True)
    if i == 1:
        for tick in ax.get_yticklabels():
            cage_ID = meta.loc[tick.get_text()]["cage_id"]
            if cage_ID in ("F1-1"):
                tick.set_color('green')
            elif cage_ID in ("F1-2"):
                tick.set_color('blue')
            elif cage_ID in ("F1-3"):
                tick.set_color('red')
            elif cage_ID in ("F1-4"):
                tick.set_color('magenta')
            elif cage_ID in ("F1-5"):
                tick.set_color('teal')
            else:
                tick.set_color('black')
    elif i==2:
        for tick in ax.get_yticklabels():
            cage_ID = meta.loc[tick.get_text()]["cage_id"]
            if cage_ID in ("F1-6"):
                tick.set_color('green')
            elif cage_ID in ("F1-7"):
                tick.set_color('blue')
            elif cage_ID in ("F1-8"):
                tick.set_color('red')
            elif cage_ID in ("F1-9"):
                tick.set_color('magenta')
            elif cage_ID in ("F1-10"):
                tick.set_color('teal')
            else:
                tick.set_color('black')      
    else:
        for tick in ax.get_yticklabels():
            cage_ID = meta.loc[tick.get_text()]["cage_id"]
            if cage_ID in ("F1-11"):
                tick.set_color('green')
            elif cage_ID in ("F1-12"):
                tick.set_color('blue')
            elif cage_ID in ("F1-13"):
                tick.set_color('red')
            elif cage_ID in ("F1-14"):
                tick.set_color('magenta')
            else:
                tick.set_color('black') 
    plt.title(f'Impact on BC1-{i} by PB generation')
    plt.legend(bbox_to_anchor=(1.05, 1), loc=2)
    plt.savefig(work_dir + "/gut_C57B_source_tracker_PB-F1%s.png" %i, transparent=False, bbox_inches='tight')

# %% [markdown]
###### F2

# %% [code]
for sink_group in ["BF2-1","BF2-2", "BF2-3"]:
    meta[meta['genotype'].isin(["C57BL6/J"])]
    conditions = (
        (meta["cage_group"] == "PB-1") | (meta["cage_group"] == "PB-2") | (meta["cage_group"] == "PB-3"),
        meta["cage_group"] == sink_group
    )
    values = ["Source", "Sink"]
    column_name ="source_sink_PB-%s" % str(sink_group)
    meta[column_name] = np.select(conditions, values, "None")

# %% [code]
PB_BF2_1_source_tracking = sourcetracker2(
    mice_no_comp_gut, #FeatureTable
    meta,
    col_envname="cage_group",
    col_type="source_sink_PB-BF2-1",
    dry=False,
    wait=True
    )
PB_BF2_2_source_tracking = sourcetracker2(
    mice_no_comp_gut, #FeatureTable
    meta,
    col_envname="cage_group",
    col_type="source_sink_PB-BF2-2",
    dry=False,
    wait=True
    )
PB_BF2_3_source_tracking = sourcetracker2(
    mice_no_comp_gut, #FeatureTable
    meta,
    col_envname="cage_group",
    col_type="source_sink_PB-BF2-3",
    dry=False,
    wait=True
    )

# %% [code]
meta[meta["cage_group"].isin(["BF2-3"])]["cage_id"]

# %% [code]
%matplotlib inline
for i,group in enumerate([PB_BF2_1_source_tracking, PB_BF2_2_source_tracking, PB_BF2_3_source_tracking], 1):
    ax= group["results"].transpose().plot.barh(stacked=True)
    if i == 1:
        for tick in ax.get_yticklabels():
            cage_ID = meta.loc[tick.get_text()]["cage_id"]
            if cage_ID in ("F2-1"):
                tick.set_color('green')
            elif cage_ID in ("F2-2"):
                tick.set_color('blue')
            elif cage_ID in ("F2-3"):
                tick.set_color('red')
            elif cage_ID in ("F2-4"):
                tick.set_color('magenta')
            elif cage_ID in ("F2-5"):
                tick.set_color('teal')
            elif cage_ID in ("F2-6"):
                tick.set_color('orange')
            else:
                tick.set_color('black')
    elif i==2:
        for tick in ax.get_yticklabels():
            cage_ID = meta.loc[tick.get_text()]["cage_id"]
            if cage_ID in ("F2-8"):
                tick.set_color('green')
            elif cage_ID in ("F2-9"):
                tick.set_color('blue')
            elif cage_ID in ("F2-10"):
                tick.set_color('red')
            elif cage_ID in ("F2-11"):
                tick.set_color('magenta')
            else:
                tick.set_color('black') 
    else:
        for tick in ax.get_yticklabels():
            cage_ID = meta.loc[tick.get_text()]["cage_id"]
            if cage_ID in ("F2-12"):
                tick.set_color('green')
            elif cage_ID in ("F2-13"):
                tick.set_color('blue')
            elif cage_ID in ("F2-14"):
                tick.set_color('red')
            elif cage_ID in ("F2-15"):
                tick.set_color('magenta')
            elif cage_ID in ("F2-16"):
                tick.set_color('teal')
            else:
                tick.set_color('black')  
    plt.title(f'Impact on BF2-{i} by PB generation')
    plt.legend(bbox_to_anchor=(1.05, 1), loc=2)
    plt.savefig(work_dir + "/gut_C57B_source_tracker_PB-F2%s.png" %i, transparent=False, bbox_inches='tight')

# %% [markdown]
###### F3

# %% [code]
for sink_group in ["BF3-1","BF3-2","BF3-3"]:
    meta[meta['genotype'].isin(["C57BL6/J"])]
    conditions = (
        (meta["cage_group"] == "PB-1") | (meta["cage_group"] == "PB-2") | (meta["cage_group"] == "PB-3"),
        meta["cage_group"] == sink_group
    )
    values = ["Source", "Sink"]
    column_name ="source_sink_PB-%s" % str(sink_group)
    meta[column_name] = np.select(conditions, values, "None")

# %% [code]
PB_BF3_1_source_tracking = sourcetracker2(
    mice_no_comp_gut, #FeatureTable
    meta,
    col_envname="cage_group",
    col_type="source_sink_PB-BF3-1",
    dry=False,
    wait=True
    )
PB_BF3_2_source_tracking = sourcetracker2(
    mice_no_comp_gut, #FeatureTable
    meta,
    col_envname="cage_group",
    col_type="source_sink_PB-BF3-2",
    dry=False,
    wait=True
    )
PB_BF3_3_source_tracking = sourcetracker2(
    mice_no_comp_gut, #FeatureTable
    meta,
    col_envname="cage_group",
    col_type="source_sink_PB-BF3-3",
    dry=False,
    wait=True
    )

# %% [code]
meta[meta["cage_group"].isin(["BF3-3"])]["cage_id"]

# %% [code]
%matplotlib inline
for i,group in enumerate([PB_BF3_1_source_tracking, PB_BF3_2_source_tracking, PB_BF3_3_source_tracking], 1):
    ax= group["results"].transpose().plot.barh(stacked=True)
    if i == 1:
        for tick in ax.get_yticklabels():
            cage_ID = meta.loc[tick.get_text()]["cage_id"]
            if cage_ID in ("F3-1"):
                tick.set_color('green')
            elif cage_ID in ("F3-2"):
                tick.set_color('blue')
            elif cage_ID in ("F3-3"):
                tick.set_color('red')
            elif cage_ID in ("F3-4"):
                tick.set_color('magenta')
            elif cage_ID in ("F3-5"):
                tick.set_color('teal')
            else:
                tick.set_color('black')
    elif i==2:
        for tick in ax.get_yticklabels():
            cage_ID = meta.loc[tick.get_text()]["cage_id"]
            if cage_ID in ("F3-7"):
                tick.set_color('green')
            elif cage_ID in ("F3-8"):
                tick.set_color('blue')
            elif cage_ID in ("F3-9"):
                tick.set_color('red')
            elif cage_ID in ("F3-10"):
                tick.set_color('magenta')
            elif cage_ID in ("F3-11"):
                tick.set_color('teal')
            elif cage_ID in ("F3-12"):
                tick.set_color('purple')
            elif cage_ID in ("F3-13"):
                tick.set_color('orange')
            else:
                tick.set_color('black')
    else:
        for tick in ax.get_yticklabels():
            cage_ID = meta.loc[tick.get_text()]["cage_id"]
            if cage_ID in ("F3-15"):
                tick.set_color('green')
            elif cage_ID in ("F3-16"):
                tick.set_color('blue')
            elif cage_ID in ("F3-17"):
                tick.set_color('red')
            elif cage_ID in ("F3-18"):
                tick.set_color('magenta')
            elif cage_ID in ("F3-19"):
                tick.set_color('teal')
            else:
                tick.set_color('black')
    plt.title(f'Impact on BF3-{i} by PB generation')
    plt.legend(bbox_to_anchor=(1.05, 1), loc=2)
    plt.savefig(work_dir + "/gut_C57B_source_tracker_PB-F3%s.png" %i, transparent=False, bbox_inches='tight')

# %% [code]


# %% [markdown]
###### F4

# %% [code]
for sink_group in ["BF4-1","BF4-2","BF4-3"]:
    meta[meta['genotype'].isin(["C57BL6/J"])]
    conditions = (
        (meta["cage_group"] == "PB-1") | (meta["cage_group"] == "PB-2") | (meta["cage_group"] == "PB-3"),
        meta["cage_group"] == sink_group
    )
    values = ["Source", "Sink"]
    column_name ="source_sink_PB-%s" % str(sink_group)
    meta[column_name] = np.select(conditions, values, "None")

# %% [code]
PB_BF4_1_source_tracking = sourcetracker2(
    mice_no_comp_gut, #FeatureTable
    meta,
    col_envname="cage_group",
    col_type="source_sink_PB-BF4-1",
    dry=False,
    wait=True
    )
PB_BF4_2_source_tracking = sourcetracker2(
    mice_no_comp_gut, #FeatureTable
    meta,
    col_envname="cage_group",
    col_type="source_sink_PB-BF4-2",
    dry=False,
    wait=True
    )
PB_BF4_3_source_tracking = sourcetracker2(
    mice_no_comp_gut, #FeatureTable
    meta,
    col_envname="cage_group",
    col_type="source_sink_PB-BF4-3",
    dry=False,
    wait=True
    )

# %% [code]
meta[meta["cage_group"].isin(["BF4-3"])]["cage_id"]

# %% [code]
%matplotlib inline
for i,group in enumerate([PB_BF4_1_source_tracking, PB_BF4_2_source_tracking, PB_BF4_3_source_tracking], 1):
    ax= group["results"].transpose().plot.barh(stacked=True)
    if i == 1:
        for tick in ax.get_yticklabels():
            cage_ID = meta.loc[tick.get_text()]["cage_id"]
            if cage_ID in ("F4-19"):
                tick.set_color('green')
            elif cage_ID in ("F4-20"):
                tick.set_color('blue')
            elif cage_ID in ("F4-21"):
                tick.set_color('red')
            elif cage_ID in ("F4-22"):
                tick.set_color('magenta')
            elif cage_ID in ("F4-23"):
                tick.set_color('teal')
            else:
                tick.set_color('black')
    elif i==2:
        for tick in ax.get_yticklabels():
            cage_ID = meta.loc[tick.get_text()]["cage_id"]
            if cage_ID in ("F4-24"):
                tick.set_color('green')
            elif cage_ID in ("F4-25"):
                tick.set_color('blue')
            elif cage_ID in ("F4-26"):
                tick.set_color('red')
            elif cage_ID in ("F4-27"):
                tick.set_color('magenta')
            else:
                tick.set_color('black')
    else:
        for tick in ax.get_yticklabels():
            cage_ID = meta.loc[tick.get_text()]["cage_id"]
            if cage_ID in ("F4-28"):
                tick.set_color('green')
            elif cage_ID in ("F4-29"):
                tick.set_color('blue')
            elif cage_ID in ("F4-30"):
                tick.set_color('red')
            elif cage_ID in ("F4-31"):
                tick.set_color('magenta')
            else:
                tick.set_color('black')
    plt.title(f'Impact on BF4-{i} by PB generation')
    plt.legend(bbox_to_anchor=(1.05, 1), loc=2)
    plt.savefig(work_dir + "/gut_C57B_source_tracker_PB-F4%s.png" %i, transparent=False, bbox_inches='tight')

# %% [code]


# %% [markdown]
###### F5

# %% [code]
for sink_group in ["BF5-1","BF5-2"]:
    meta[meta['genotype'].isin(["BALB/c"])]
    conditions = (
        (meta["cage_group"] == "PB-1") | (meta["cage_group"] == "PB-2") | (meta["cage_group"] == "PB-3"),
        meta["cage_group"] == sink_group
    )
    values = ["Source", "Sink"]
    column_name ="source_sink_PB-%s" % str(sink_group)
    meta[column_name] = np.select(conditions, values, "None")

# %% [code]
PB_BF5_1_source_tracking = sourcetracker2(
    mice_no_comp_gut, #FeatureTable
    meta,
    col_envname="cage_group",
    col_type="source_sink_PB-BF5-1",
    dry=False,
    wait=True
    )
PB_BF5_2_source_tracking = sourcetracker2(
    mice_no_comp_gut, #FeatureTable
    meta,
    col_envname="cage_group",
    col_type="source_sink_PB-BF5-2",
    dry=False,
    wait=True
    )

# %% [code]
meta[meta["cage_group"].isin(["BF5-2"])]["cage_id"]

# %% [code]
%matplotlib inline
for i,group in enumerate([PB_BF5_1_source_tracking, PB_BF5_2_source_tracking], 1):
    ax= group["results"].transpose().plot.barh(stacked=True)
    if i == 1:
        for tick in ax.get_yticklabels():
            cage_ID = meta.loc[tick.get_text()]["cage_id"]
            if cage_ID in ("F5-1"):
                tick.set_color('green')
            elif cage_ID in ("F5-2"):
                tick.set_color('blue')
            elif cage_ID in ("F5-3"):
                tick.set_color('red')
            elif cage_ID in ("F5-4"):
                tick.set_color('magenta')
            else:
                tick.set_color('black')
    else:
        for tick in ax.get_yticklabels():
            cage_ID = meta.loc[tick.get_text()]["cage_id"]
            if cage_ID in ("F5-5"):
                tick.set_color('green')
            elif cage_ID in ("F5-6"):
                tick.set_color('blue')
            elif cage_ID in ("F5-7"):
                tick.set_color('red')
            else:
                tick.set_color('black')
    plt.title(f'Impact on BF5-{i} by PB generation')
    plt.legend(bbox_to_anchor=(1.05, 1), loc=2)
    plt.savefig(work_dir + "/gut_C57B_source_tracker_PB-F5%s.png" %i, transparent=False, bbox_inches='tight')

# %% [markdown]
### Sebum

# %% [code]
rarefied_sebum = biom2pandas(work_dir + "/rarefied-filtered-sebum-table_export/feature-table.biom").fillna(0)

# %% [markdown]
#### BALB/c
##### Impact of BF1 cage groups on BF2 cage groups

# %% [code]
BF1_2_1_source_tracking_sebum = sourcetracker2(
    rarefied_sebum, #FeatureTable
    meta,
    col_envname="cage_group",
    col_type="source_sink_BF1-BF2-1",
    dry=False,
    wait=True
    )
BF1_2_2_source_tracking_sebum = sourcetracker2(
    rarefied_sebum, #FeatureTable
    meta,
    col_envname="cage_group",
    col_type="source_sink_BF1-BF2-2",
    dry=False,
    wait=True
    )

# %% [code]
for i,group in enumerate([BF1_2_1_source_tracking_sebum, BF1_2_2_source_tracking_sebum], 1):
    ax= group["results"].transpose().plot.barh(stacked=True)
    if i == 1:
        for tick in ax.get_yticklabels():
            cage_ID = meta.loc[tick.get_text()]["cage_id"]
            if cage_ID in ("F2-17"):
                tick.set_color('green')
            elif cage_ID in ("F2-18"):
                tick.set_color('blue')
            elif cage_ID in ("F2-19"):
                tick.set_color('darkred')
            elif cage_ID in ("F2-20"):
                tick.set_color('magenta')
            else:
                tick.set_color('black')
        
    else:
        for tick in ax.get_yticklabels():
            cage_ID = meta.loc[tick.get_text()]["cage_id"]
            if cage_ID in ("F2-21"):
                tick.set_color('green')
            elif cage_ID in ("F2-22"):
                tick.set_color('blue')
            elif cage_ID in ("F2-23"):
                tick.set_color('darkred')
            elif cage_ID in ("F2-24"):
                tick.set_color('magenta')
            elif cage_ID in ("F2-25"):
                tick.set_color('teal')
            else:
                tick.set_color('black')
    plt.title(f'Impact on BF2-{i} by BF1 generation')
    plt.legend(bbox_to_anchor=(1.05, 1), loc=2)

# %% [markdown]
##### Impact of BF2 cage groups on BF3 cage groups

# %% [code]
BF2_3_1_source_tracking_sebum = sourcetracker2(
    rarefied_sebum, #FeatureTable
    meta,
    col_envname="cage_group",
    col_type="source_sink_BF2-BF3-1",
    dry=False,
    wait=True
)

BF2_3_2_source_tracking_sebum = sourcetracker2(
    rarefied_sebum, #FeatureTable
    meta,
    col_envname="cage_group",
    col_type="source_sink_BF2-BF3-2",
    dry=False,
    wait=True
)

BF2_3_3_source_tracking_sebum = sourcetracker2(
    rarefied_sebum, #FeatureTable
    meta,
    col_envname="cage_group",
    col_type="source_sink_BF2-BF3-3",
    dry=False,
    wait=True
)

# %% [code]
for i,group in enumerate([BF2_3_1_source_tracking_sebum, BF2_3_2_source_tracking_sebum, BF2_3_3_source_tracking_sebum], 1):
    ax= group["results"].transpose().plot.barh(stacked=True)
    if i == 1:
        for tick in ax.get_yticklabels():
            cage_ID = meta.loc[tick.get_text()]["cage_id"]
            if cage_ID in ("F3-21"):
                tick.set_color('green')
            elif cage_ID in ("F3-22"):
                tick.set_color('blue')
            else:
                tick.set_color('black')
    elif i==2:
        for tick in ax.get_yticklabels():
            cage_ID = meta.loc[tick.get_text()]["cage_id"]
            if cage_ID in ("F3-23"):
                tick.set_color('green')
            elif cage_ID in ("F3-24"):
                tick.set_color('blue')
            elif cage_ID in ("F3-25"):
                tick.set_color('red')
            elif cage_ID in ("F3-26"):
                tick.set_color('magenta')
            elif cage_ID in ("F3-27"):
                tick.set_color('teal')
            else:
                tick.set_color('black')       
    else:
        for tick in ax.get_yticklabels():
            cage_ID = meta.loc[tick.get_text()]["cage_id"]
            if cage_ID in ("F3-28"):
                tick.set_color('green')
            elif cage_ID in ("F3-29"):
                tick.set_color('blue')
            elif cage_ID in ("F3-30"):
                tick.set_color('red')
            elif cage_ID in ("F3-31"):
                tick.set_color('magenta')
            elif cage_ID in ("F3-32"):
                tick.set_color('teal')
            else:
                tick.set_color('black')
    plt.title(f'Impact on BF3-{i} by BF2 generation')
    plt.legend(bbox_to_anchor=(1.05, 1), loc=2)

# %% [markdown]
##### Impact of BF3 cage groups on BF4 cage groups

# %% [code]
BF3_4_1_source_tracking_sebum = sourcetracker2(
    rarefied_sebum, #FeatureTable
    meta,
    col_envname="cage_group",
    col_type="source_sink_BF3-BF4-1",
    dry=False,
    wait=True
)

BF3_4_2_source_tracking_sebum = sourcetracker2(
    rarefied_sebum, #FeatureTable
    meta,
    col_envname="cage_group",
    col_type="source_sink_BF3-BF4-2",
    dry=False,
    wait=True
)

BF3_4_3_source_tracking_sebum = sourcetracker2(
    rarefied_sebum, #FeatureTable
    meta,
    col_envname="cage_group",
    col_type="source_sink_BF3-BF4-3",
    dry=False,
    wait=True
)

# %% [code]
for i,group in enumerate([BF3_4_1_source_tracking_sebum, BF3_4_2_source_tracking_sebum, BF3_4_3_source_tracking_sebum], 1):
    ax= group["results"].transpose().plot.barh(stacked=True)
    if i == 1:
        for tick in ax.get_yticklabels():
            cage_ID = meta.loc[tick.get_text()]["cage_id"]
            if cage_ID in ("F4-19"):
                tick.set_color('green')
            elif cage_ID in ("F4-20"):
                tick.set_color('blue')
            elif cage_ID in ("F4-21"):
                tick.set_color('red')
            elif cage_ID in ("F4-22"):
                tick.set_color('magenta')
            elif cage_ID in ("F4-23"):
                tick.set_color('teal')
            else:
                tick.set_color('black')
    elif i==2:
        for tick in ax.get_yticklabels():
            cage_ID = meta.loc[tick.get_text()]["cage_id"]
            if cage_ID in ("F4-24"):
                tick.set_color('green')
            elif cage_ID in ("F4-25"):
                tick.set_color('blue')
            elif cage_ID in ("F4-26"):
                tick.set_color('red')
            elif cage_ID in ("F4-27"):
                tick.set_color('magenta')
            else:
                tick.set_color('black')       
    else:
        for tick in ax.get_yticklabels():
            cage_ID = meta.loc[tick.get_text()]["cage_id"]
            if cage_ID in ("F4-28"):
                tick.set_color('green')
            elif cage_ID in ("F4-29"):
                tick.set_color('blue')
            elif cage_ID in ("F4-30"):
                tick.set_color('red')
            elif cage_ID in ("F4-31"):
                tick.set_color('magenta')
            else:
                tick.set_color('black')
    plt.title(f'Impact on BF4-{i} by BF3 generation')
    plt.legend(bbox_to_anchor=(1.05, 1), loc=2)

# %% [code]


# %% [markdown]
##### Impact of BF4 cage groups on BF5 cage groups

# %% [code]
BF4_5_1_source_tracking_sebum = sourcetracker2(
    rarefied_sebum, #FeatureTable
    meta,
    col_envname="cage_group",
    col_type="source_sink_BF4-BF5-1",
    dry=False,
    wait=True
)

BF4_5_2_source_tracking_sebum = sourcetracker2(
    rarefied_sebum, #FeatureTable
    meta,
    col_envname="cage_group",
    col_type="source_sink_BF4-BF5-2",
    dry=False,
    wait=True
)

BF4_5_3_source_tracking_sebum = sourcetracker2(
    rarefied_sebum, #FeatureTable
    meta,
    col_envname="cage_group",
    col_type="source_sink_BF4-BF5-3",
    dry=False,
    wait=True
)

# %% [code]
for i,group in enumerate([BF4_5_1_source_tracking_sebum, BF4_5_2_source_tracking_sebum, BF4_5_3_source_tracking_sebum], 1):
    ax= group["results"].transpose().plot.barh(stacked=True)
    if i == 1:
        for tick in ax.get_yticklabels():
            cage_ID = meta.loc[tick.get_text()]["cage_id"]
            if cage_ID in ("F5-8"):
                tick.set_color('green')
            elif cage_ID in ("F5-9"):
                tick.set_color('blue')
            elif cage_ID in ("F5-10"):
                tick.set_color('red')
            else:
                tick.set_color('black')
    elif i==2:
        for tick in ax.get_yticklabels():
            cage_ID = meta.loc[tick.get_text()]["cage_id"]
            if cage_ID in ("F5-11"):
                tick.set_color('green')
            elif cage_ID in ("F5-12"):
                tick.set_color('blue')
            elif cage_ID in ("F5-13"):
                tick.set_color('red')
            elif cage_ID in ("F5-14"):
                tick.set_color('magenta')
            else:
                tick.set_color('black')       
    else:
        for tick in ax.get_yticklabels():
            cage_ID = meta.loc[tick.get_text()]["cage_id"]
            if cage_ID in ("F5-15"):
                tick.set_color('green')
            elif cage_ID in ("F5-16"):
                tick.set_color('blue')
            elif cage_ID in ("F5-17"):
                tick.set_color('red')
            elif cage_ID in ("F5-18"):
                tick.set_color('magenta')
            elif cage_ID in ("F5-19"):
                tick.set_color('teal')
            else:
                tick.set_color('black')
    plt.title(f'Impact on BF5-{i} by BF4 generation')
    plt.legend(bbox_to_anchor=(1.05, 1), loc=2)

# %% [code]


# %% [markdown]
#### C57BL6/J
##### Impact of BF1 cage groups on BF2 cage groups

# %% [code]
CF1_2_1_source_tracking_sebum = sourcetracker2(
    rarefied_sebum, #FeatureTaCle
    meta,
    col_envname="cage_group",
    col_type="source_sink_CF1-CF2-1",
    dry=False,
    wait=True
    )
CF1_2_2_source_tracking_sebum = sourcetracker2(
    rarefied_sebum, #FeatureTable
    meta,
    col_envname="cage_group",
    col_type="source_sink_CF1-CF2-2",
    dry=False,
    wait=True
    )
CF1_2_3_source_tracking_sebum = sourcetracker2(
    rarefied_sebum, #FeatureTable
    meta,
    col_envname="cage_group",
    col_type="source_sink_CF1-CF2-3",
    dry=False,
    wait=True
    )

# %% [code]
for i,group in enumerate([CF1_2_1_source_tracking_sebum, CF1_2_2_source_tracking_sebum, CF1_2_3_source_tracking_sebum], 1):
    ax= group["results"].transpose().plot.barh(stacked=True)
    if i == 1:
        for tick in ax.get_yticklabels():
            cage_ID = meta.loc[tick.get_text()]["cage_id"]
            if cage_ID in ("F2-1"):
                tick.set_color('green')
            elif cage_ID in ("F2-2"):
                tick.set_color('blue')
            elif cage_ID in ("F2-3"):
                tick.set_color('red')
            elif cage_ID in ("F2-4"):
                tick.set_color('magenta')
            elif cage_ID in ("F2-5"):
                tick.set_color('teal')
            elif cage_ID in ("F2-6"):
                tick.set_color('darkolivegreen')
            elif cage_ID in ("F2-7"):
                tick.set_color('darkorange')
            else:
                tick.set_color('black')
    elif i==2:
        for tick in ax.get_yticklabels():
            cage_ID = meta.loc[tick.get_text()]["cage_id"]
            if cage_ID in ("F2-8"):
                tick.set_color('green')
            elif cage_ID in ("F2-9"):
                tick.set_color('blue')
            elif cage_ID in ("F2-10"):
                tick.set_color('red')
            elif cage_ID in ("F2-11"):
                tick.set_color('magenta')
            else:
                tick.set_color('black')       
    else:
        for tick in ax.get_yticklabels():
            cage_ID = meta.loc[tick.get_text()]["cage_id"]
            if cage_ID in ("F2-12"):
                tick.set_color('green')
            elif cage_ID in ("F2-13"):
                tick.set_color('blue')
            elif cage_ID in ("F2-14"):
                tick.set_color('red')
            elif cage_ID in ("F2-15"):
                tick.set_color('magenta')
            elif cage_ID in ("F2-16"):
                tick.set_color('teal')
            else:
                tick.set_color('black')
    plt.title(f'Impact on CF2-{i} by CF1 generation')
    plt.legend(bbox_to_anchor=(1.05, 1), loc=2)

# %% [markdown]
##### Impact of CF2 cage groups on CF3 cage groups

# %% [code]
CF2_3_1_source_tracking_sebum = sourcetracker2(
    rarefied_sebum, #FeatureTable
    meta,
    col_envname="cage_group",
    col_type="source_sink_CF2-CF3-1",
    dry=False,
    wait=True
)

CF2_3_2_source_tracking_sebum = sourcetracker2(
    rarefied_sebum, #FeatureTable
    meta,
    col_envname="cage_group",
    col_type="source_sink_CF2-CF3-2",
    dry=False,
    wait=True
)

CF2_3_3_source_tracking_sebum = sourcetracker2(
    rarefied_sebum, #FeatureTable
    meta,
    col_envname="cage_group",
    col_type="source_sink_CF2-CF3-3",
    dry=False,
    wait=True
)

# %% [code]
meta[meta['cage_group'].isin(['CF3-1'])]['cage_id'].value_counts()

# %% [code]
for i,group in enumerate([CF2_3_1_source_tracking_sebum, CF2_3_2_source_tracking_sebum, CF2_3_3_source_tracking_sebum], 1):
    ax= group["results"].transpose().plot.barh(stacked=True)
    if i == 1:
        for tick in ax.get_yticklabels():
            cage_ID = meta.loc[tick.get_text()]["cage_id"]
            if cage_ID in ("F3-1"):
                tick.set_color('green')
            elif cage_ID in ("F3-2"):
                tick.set_color('blue')
            elif cage_ID in ("F3-3"):
                tick.set_color('red')
            elif cage_ID in ("F3-4"):
                tick.set_color('magenta')
            elif cage_ID in ("F3-5"):
                tick.set_color('teal')
            elif cage_ID in ("F3-6"):
                tick.set_color('darkorange')
            else:
                tick.set_color('black')
    elif i==2:
        for tick in ax.get_yticklabels():
            cage_ID = meta.loc[tick.get_text()]["cage_id"]
            if cage_ID in ("F3-7"):
                tick.set_color('green')
            elif cage_ID in ("F3-8"):
                tick.set_color('blue')
            elif cage_ID in ("F3-9"):
                tick.set_color('red')
            elif cage_ID in ("F3-10"):
                tick.set_color('magenta')
            elif cage_ID in ("F3-11"):
                tick.set_color('teal')
            elif cage_ID in ("F3-12"):
                tick.set_color('darkolivegreen')
            elif cage_ID in ("F3-13"):
                tick.set_color('darkorange')
            elif cage_ID in ("F3-14"):
                tick.set_color('dodgerblue')
            else:
                tick.set_color('black')       
    else:
        for tick in ax.get_yticklabels():
            cage_ID = meta.loc[tick.get_text()]["cage_id"]
            if cage_ID in ("F3-15"):
                tick.set_color('green')
            elif cage_ID in ("F3-16"):
                tick.set_color('blue')
            elif cage_ID in ("F3-17"):
                tick.set_color('red')
            elif cage_ID in ("F3-18"):
                tick.set_color('magenta')
            elif cage_ID in ("F3-19"):
                tick.set_color('teal')
            elif cage_ID in ("F3-20"):
                tick.set_color('darkorange')
            else:
                tick.set_color('black')
    plt.title(f'Impact on CF3-{i} by CF2 generation')
    plt.legend(bbox_to_anchor=(1.05, 1), loc=2)

# %% [markdown]
##### Impact of CF3 cage groups on CF4 cage groups

# %% [code]
CF3_4_1_source_tracking_sebum = sourcetracker2(
    rarefied_sebum, #FeatureTable
    meta,
    col_envname="cage_group",
    col_type="source_sink_CF3-CF4-1",
    dry=False,
    wait=True
)

CF3_4_2_source_tracking_sebum = sourcetracker2(
    rarefied_sebum, #FeatureTable
    meta,
    col_envname="cage_group",
    col_type="source_sink_CF3-CF4-2",
    dry=False,
    wait=True
)

CF3_4_3_source_tracking_sebum = sourcetracker2(
    rarefied_sebum, #FeatureTable
    meta,
    col_envname="cage_group",
    col_type="source_sink_CF3-CF4-3",
    dry=False,
    wait=True
)

# %% [code]
for i,group in enumerate([CF3_4_1_source_tracking_sebum, CF3_4_2_source_tracking_sebum, CF3_4_3_source_tracking_sebum], 1):
    ax= group["results"].transpose().plot.barh(stacked=True)
    if i == 1:
        for tick in ax.get_yticklabels():
            cage_ID = meta.loc[tick.get_text()]["cage_id"]
            if cage_ID in ("F4-1"):
                tick.set_color('green')
            elif cage_ID in ("F4-2"):
                tick.set_color('blue')
            elif cage_ID in ("F4-3"):
                tick.set_color('red')
            elif cage_ID in ("F4-4"):
                tick.set_color('magenta')
            elif cage_ID in ("F4-5"):
                tick.set_color('teal')
            elif cage_ID in ("F4-6"):
                tick.set_color('darkorange')
            else:
                tick.set_color('black')
    elif i==2:
        for tick in ax.get_yticklabels():
            cage_ID = meta.loc[tick.get_text()]["cage_id"]
            if cage_ID in ("F4-8"):
                tick.set_color('green')
            elif cage_ID in ("F4-9"):
                tick.set_color('blue')
            elif cage_ID in ("F4-10"):
                tick.set_color('red')
            elif cage_ID in ("F4-11"):
                tick.set_color('magenta')
            elif cage_ID in ("F4-12"):
                tick.set_color('teal')
            elif cage_ID in ("F4-13"):
                tick.set_color('darkorange')
            else:
                tick.set_color('black')       
    else:
        for tick in ax.get_yticklabels():
            cage_ID = meta.loc[tick.get_text()]["cage_id"]
            if cage_ID in ("F4-14"):
                tick.set_color('green')
            elif cage_ID in ("F4-15"):
                tick.set_color('blue')
            elif cage_ID in ("F4-16"):
                tick.set_color('red')
            elif cage_ID in ("F4-17"):
                tick.set_color('magenta')
            elif cage_ID in ("F4-18"):
                tick.set_color('teal')
            else:
                tick.set_color('black')
    plt.title(f'Impact on CF4-{i} by CF3 generation')
    plt.legend(bbox_to_anchor=(1.05, 1), loc=2)

# %% [code]


# %% [markdown]
##### Impact of CF4 cage groups on CF5 cage groups

# %% [code]
CF4_5_1_source_tracking_sebum = sourcetracker2(
    rarefied_sebum, #FeatureTable
    meta,
    col_envname="cage_group",
    col_type="source_sink_CF4-CF5-1",
    dry=False,
    wait=True
)

CF4_5_2_source_tracking_sebum = sourcetracker2(
    rarefied_sebum, #FeatureTable
    meta,
    col_envname="cage_group",
    col_type="source_sink_CF4-CF5-2",
    dry=False,
    wait=True
)

# %% [code]
for i,group in enumerate([CF4_5_1_source_tracking_sebum, CF4_5_2_source_tracking_sebum], 1):
    ax= group["results"].transpose().plot.barh(stacked=True)
    if i == 1:
        for tick in ax.get_yticklabels():
            cage_ID = meta.loc[tick.get_text()]["cage_id"]
            if cage_ID in ("F5-1"):
                tick.set_color('green')
            elif cage_ID in ("F5-2"):
                tick.set_color('blue')
            elif cage_ID in ("F5-3"):
                tick.set_color('red')
            elif cage_ID in ("F5-4"):
                tick.set_color('magenta')
            else:
                tick.set_color('black')  
    else:
        for tick in ax.get_yticklabels():
            cage_ID = meta.loc[tick.get_text()]["cage_id"]
            if cage_ID in ("F5-5"):
                tick.set_color('green')
            elif cage_ID in ("F5-6"):
                tick.set_color('blue')
            elif cage_ID in ("F5-7"):
                tick.set_color('red')
            else:
                tick.set_color('black')
    plt.title(f'Impact on CF5-{i} by CF4 generation')
    plt.legend(bbox_to_anchor=(1.05, 1), loc=2)
```
