## Additional File 2: Analysis repository for "The host genotype actively shapes its microbiome across generations in laboratory mice": Benga_Thesis_notebook.ipynb

```
# %% [markdown]
# Influence of the genetic background on murine gut microbiome composition and diversity over five generations

# %% [markdown]
The main goal of this study is to study the gut and skin microbiome of C75BL/6J and BALB/c mice, which have been naturally colonised with the same microbiome simultaneously.To achieve this the mice were born and weaned by B6CF1 foster mothers. The aim is to determine whether the genotype of the mice will have an impact on their microbiome. Therefore, the mice have been bred over 5 generations and kept in identical housing with identical breeding procedures, to minimise external effects on the microbiome. Additionally, sex and cage environment have been recorded during the experiment, to evaluate if they play a role in the composition of the microbiome as well.

This analysis aims to explore the differences and similarities of the mice microbiomes. It focuses on the calculation of alpha and beta diversities (Shannon, Faith's PD, observed OTUs, unweighted UniFrac) to carve out whether the genotype of a mouse has an impact on its gut microbiome. The alpha and beta diversities are visualised and used for statistical tests. Additionally, taxa that are significantly different between both genotypes and their respective generations are determined too by using ANCOM. Furthermore, we investigate whether the the parental microbiomes impacted their children's microbiomes by analyzing them with SourceTracker.

# %% [markdown]
## bcl2fastq and Trimming step

First, the raw sequencing data was transferred into FASTQ files using bcl2fastq (v2.19.0.316), with the parameter of `--barcode-mismatches 0` to ensure no false barcodes get assigned to the wrong sequence.

# %% [markdown]
`bcl2fastq` <br>
  `--runfolder-dir 210902_M04304_0266_000000000-JNGPR ` <br>
  `--output-dir /vol/jlab/MicrobiomeAnalyses/Projects/Benga_ThreeGen/Demux/ThirdBatch/`  <br>
  `--ignore-missing-bcls`  <br>
  `--sample-sheet /vol/jlab/MicrobiomeAnalyses/Projects/Benga_ThreeGen/Incoming/210902_M04304_0266_000000000-JNGPR.csv`  <br>
  `--loading-threads 6`  <br>
  `--processing-threads 6`  <br>
  `--writing-threads 6`  <br>
  `--barcode-mismatches 0` <br>

# %% [markdown]
Then, the adapters were trimmed using cutadapt version 2.10. Through the parameter `--minimum-length 1:1` no empty sequences will remain if only the adapter was included in the sequence.

`for forward in *R1_001.fastq.gz:` <br>
`do reverse=$(echo $forward | sed `s/R1/R2`);` <br>
`cutadapt` <br>
`-g CCTACGGGNGGCWGCAG` <br>
`-G GACTACHVGGGTATCTAATCC` <br>
`-n 2` <br>
`–minimum-length 1:1` <br>
`-o /vol/jlab/MicrobiomeAnalyses/Projects/Benga_ThreeGen/Trimming/CorrectedIDs/$forward` <br>
`-p /vol/jlab/MicrobiomeAnalyses/Projects/Benga_ThreeGen/Trimming/CorrectedIDs/$reverse $forward $reverse;` <br>
`done`

# %% [markdown]
## Adding additional columns to the metadata

# %% [code]
q_path='export PATH=/homes/arehm/miniconda3/envs/qiime2-2022.8/bin:$PATH'
work_dir='/homes/arehm/Documents/Thesis/JupyterNotebook'
meta_file='/homes/arehm/Documents/Thesis/13422_20210909-070806.txt'

# %% [markdown]
For all the following analysis to run smoothly, all the additions to the existing metadata sheet from qiita are compiled in the following cell.

# %% [code]
import pandas as pd
import numpy as np

meta = pd.read_csv(meta_file, sep="\t", dtype=str).set_index('sample_name')

indeces = []
for i in range (64,88):
    index = "13422.colon3" + str(i)
    indeces.append(index)
    index = "13422.skin3" + str(i)
    indeces.append(index)
    index = "13422.3" + str(i) + "skin"
    indeces.append(index)

conditions = [
    (meta["generation"] == "RM"),
    (meta["generation"] == "P1"),
    (meta["generation"] == "F1"),
    (meta["generation"] == "F2"),
    (meta["generation"] == "F3"),
    (meta["generation"] == "F4"),
    (meta["generation"] == "F5")
]

values = [-1, 0, 1, 2, 3, 4, 5]

meta["gen_number"] = np.select(conditions, values, -2)

conditions = [
    (meta["cage_group"] == "PC") | (meta["cage_group"] == "CF1-1") | (meta["cage_group"] == "CF2-1") | (meta["cage_group"] == "CF3-1") | (meta["cage_group"] == "CF4-1"),
    (meta["cage_group"] == "PC") | (meta["cage_group"] == "CF1-2") | (meta["cage_group"] == "CF2-2") | (meta["cage_group"] == "CF3-2") | (meta["cage_group"] == "CF4-2") | (meta["cage_group"] == "CF5-1"),
    (meta["cage_group"] == "PC") | (meta["cage_group"] == "CF1-3") | (meta["cage_group"] == "CF2-3") | (meta["cage_group"] == "CF3-3") | (meta["cage_group"] == "CF4-3") | (meta["cage_group"] == "CF5-2"),
    (meta["cage_group"] == "PB") | (meta["cage_group"] == "BF1-1") | (meta["cage_group"] == "BF2-1") | (meta["cage_group"] == "BF3-1") | (meta["cage_group"] == "BF4-1") | (meta["cage_group"] == "BF5-1"),
    (meta["cage_group"] == "PB") | (meta["cage_group"] == "BF1-2") | (meta["cage_group"] == "BF2-2") | (meta["cage_group"] == "BF3-2") | (meta["cage_group"] == "BF4-2") | (meta["cage_group"] == "BF5-2"),
    (meta["cage_group"] == "PB") | (meta["cage_group"] == "BF1-2") | (meta["cage_group"] == "BF2-2") | (meta["cage_group"] == "BF3-3") | (meta["cage_group"] == "BF4-3") | (meta["cage_group"] == "BF5-3")
]

values = ["C1", "C2", "C3", "B1", "B2", "B3"]

meta["cage_lineage"] = np.select(conditions, values, "other")

#meta_noRM = meta.loc[(meta['genotype'] != "B6CF1") & (meta["host_common_name"] == "mouse")]
meta_noRM = meta.loc[(meta["host_common_name"] == "mouse")]

meta['i_generation'] = meta['generation'].apply(lambda x: {'RM': '-1: RM',
                                    'P1': '0: P1',
                                    'F1': '1: F1',
                                    'F2': '2: F2',
                                    'F3': '3: F3',
                                    'F4': '4: F4',
                                    'F5': '5: F5',
                                   }.get(x, x))

meta['comparison'] = np.where(
    meta.index.isin(indeces), 1, 0) 

meta.to_csv("meta_gen_num.txt", header=True, sep="\t")

# %% [code]
pd.set_option('display.max_columns', None)
pd.set_option('display.max_rows', 400)
display(meta[(meta["batch"]=="3") & (meta["host_body_product"]=="UBERON:sebum") & (meta["comparison"]==1)])

# %% [code]
batch2 = meta[(meta["batch"]=="2") & (meta["host_body_product"]=="UBERON:sebum")]
pd.set_option('display.max_columns', None)
meta.loc[batch2.index,:]

# %% [code]
duplicates = []
for i in range (64,88):
    index = "13422.skin3" + str(i)
    duplicates.append(index)
duplicate_indeces = str(duplicates)[1:-1]

# %% [markdown]
## Filtering
First, the feature table is filtered to drop features with an abundance lower than 10. 

# %% [code]
#filtering out duplicate skin samples from batch 2
! $q_path; qiime feature-table filter-samples \
--i-table /homes/arehm/Benga_ThreeGen/FromQiita/128106_feature-table.qza \
--m-metadata-file $meta_file \
--p-where "[sample_name] IN ($duplicate_indeces)" \
--p-exclude-ids \
--o-filtered-table $work_dir/no_duplicates_table.qza

# %% [code]
%time ! $q_path; qiime feature-table filter-features \
  --i-table $work_dir/no_duplicates_table.qza \
  --p-min-frequency 10 \
  --o-filtered-table $work_dir/filtered-10-feature-table.qza

# %% [code]
%time ! $q_path; qiime feature-table summarize \
  --i-table $work_dir/filtered-10-feature-table.qza\
  --o-visualization $work_dir/summary_filtered-10-table.qzv \
  --m-sample-metadata-file $meta_file

# %% [markdown]
Then, the sequence data from qiita after deblur was performed is imported into a QIIME2 artifact. This file is needed for the subsequent step of assigning taxonomy to the features using the Greengenes classifier provided on the QIIME2 docs page https://docs.qiime2.org/2022.2/data-resources/ . The Greengenes 13_8 99% OTUs full-length sequences (MD5: 6bbc9b3f2f9b51d663063a7979dd95f1)file was used.

# %% [code]
%time ! $q_path; qiime tools import \
  --input-path /homes/arehm/Benga_ThreeGen/FromQiita/128106_reference-hit.seqs.fa \
  --type 'FeatureData[Sequence]' \
  --output-path $work_dir/rep-seqs.qza

# %% [code]
%time ! $q_path ; qiime feature-classifier classify-sklearn \
  --i-reads $work_dir/rep-seqs.qza \
  --i-classifier $work_dir/gg-13-8-99-nb-classifier.qza \
  --o-classification $work_dir/taxonomy_full.qza

# %% [markdown]
In order to avoid features belonging to *Cyanobacteria* in the analysis, the need to be filtered out. Using the taxonomy from the previous step as a metadata file in `--m-metadata-file`.

# %% [code]
%time ! $q_path; qiime taxa filter-table \
  --i-table $work_dir/filtered-10-feature-table.qza \
  --i-taxonomy $work_dir/taxonomy_full.qza \
  --p-exclude mitochondria,chloroplast \
  --o-filtered-table $work_dir/filtered-noCyanobacteria-feature-table.qza

# %% [code]
%time ! $q_path; qiime tools import \
  --input-path /homes/arehm/Benga_ThreeGen/FromQiita/128106_insertion_tree.relabelled.tre \
  --type 'Phylogeny[Rooted]' \
  --output-path $work_dir/insertion-tree.qza

%time ! $q_path; qiime fragment-insertion filter-features \
--i-table $work_dir/filtered-noCyanobacteria-feature-table.qza \
--i-tree $work_dir/insertion-tree.qza \
--o-filtered-table $work_dir/fragment-filtered‐feature-table.qza \
--o-removed-table $work_dir/removed‐feature-table.qza

# %% [code]
%time ! $q_path; qiime feature-table summarize \
  --i-table $work_dir/fragment-filtered‐feature-table.qza\
  --o-visualization $work_dir/summary-fragment-filtered‐feature-table.qzv \
  --m-sample-metadata-file $meta_file

# %% [code]
%time ! $q_path; qiime tools export \
  --input-path $work_dir/fragment-filtered‐feature-table.qza\
  --output-path $work_dir/exported-fragment-filtered-feature-table

# %% [markdown]
## Rarefaction

Rarefaction is important to avoid inflation of diversity. Rarefaction curves were created for all samples, as well as only gut and sebum samples.

# %% [markdown]
### Gut
Beofre following the previous steps for imloring a suitable sequencing depth, the table was filtered to only include gut samples.

# %% [code]
%time ! $q_path; qiime feature-table filter-samples \
  --i-table $work_dir/fragment-filtered‐feature-table.qza \
  --m-metadata-file $meta_file \
  --p-where "[host_body_product] IN ('UBERON:lumen of gut')" \
  --o-filtered-table $work_dir/fragment-filtered-gut-table.qza

# %% [code]
%time ! $q_path; qiime feature-table summarize \
  --i-table $work_dir/fragment-filtered-gut-table.qza \
  --o-visualization $work_dir/summary-fragment-filtered-gut-table.qzv \
  --m-sample-metadata-file $meta_file

# %% [code]
%time ! $q_path; qiime diversity alpha-rarefaction \
  --i-table $work_dir/fragment-filtered-gut-table.qza \
  --i-phylogeny $work_dir/insertion-tree.qza \
  --p-max-depth 20000 \
  --p-min-depth 500 \
  --p-iterations 10 \
  --p-steps 50 \
  --m-metadata-file $meta_file \
  --o-visualization $work_dir/rarefaction_curves_gut.qzv

# %% [markdown]
After consulting the rarefaction plots, a sampling depth of 6000 was chosen.

# %% [code]
%time ! $q_path; qiime feature-table rarefy \
  --i-table $work_dir/fragment-filtered-gut-table.qza \
  --p-sampling-depth 6000 \
  --o-rarefied-table $work_dir/rarefied6000-filtered-gut-table.qza
%time ! $q_path; qiime feature-table summarize \
  --i-table $work_dir/rarefied6000-filtered-gut-table.qza \
  --o-visualization $work_dir/summary-rarefied6000-filtered-gut-table.qzv \
  --m-sample-metadata-file $meta_file

# %% [code]
biom2pandas('/homes/arehm/Documents/Thesis/JupyterNotebook/feature-table_export/feature-table.biom')

# %% [markdown]
### Sebum
Analogous to the gut samples.

# %% [code]
%time ! $q_path; qiime feature-table filter-samples \
  --i-table $work_dir/fragment-filtered‐feature-table.qza \
  --m-metadata-file $meta_file \
  --p-where "[host_body_product] IN ('UBERON:sebum')" \
  --o-filtered-table $work_dir/fragment-filtered-sebum-table.qza

# %% [code]
%time ! $q_path; qiime feature-table summarize \
  --i-table $work_dir/fragment-filtered-sebum-table.qza \
  --o-visualization $work_dir/summary-fragment-filtered-sebum-table.qzv \
  --m-sample-metadata-file $meta_file

# %% [code]
%time ! $q_path; qiime diversity alpha-rarefaction \
  --i-table $work_dir/fragment-filtered-sebum-table.qza \
  --i-phylogeny $work_dir/insertion-tree.qza \
  --p-max-depth 4000 \
  --p-min-depth 500 \
  --p-iterations 10 \
  --p-steps 50 \
  --m-metadata-file $meta_file \
  --o-visualization $work_dir/rarefaction_curves_sebum.qzv

# %% [code]
%time ! $q_path; qiime feature-table rarefy \
  --i-table $work_dir/fragment-filtered-sebum-table.qza \
  --p-sampling-depth 1000 \
  --o-rarefied-table $work_dir/rarefied-filtered-sebum-table.qza

# %% [code]
%time ! $q_path; qiime feature-table summarize \
  --i-table $work_dir/rarefied-filtered-sebum-table.qza \
  --o-visualization $work_dir/summary-rarefied-filtered-sebum-table.qzv \
  --m-sample-metadata-file $meta_file

# %% [markdown]
# PICRUSt2
## nicht rarifiziert

# %% [code]
#exportieren der Biom tabellen -> verwenden derer die für Alphadiversität exportiert wurden, also OHNE rarefaction und OHNE Vergleichsproben
from ggmap.snippets import biom2pandas

picrust_gut = biom2pandas(work_dir + "/exported-filtered-gut-table-for-alpha/feature-table.biom").fillna(0)


# %% [code]
#run picrust2
res_picrust2_gut = picrust2(picrust_gut, dry=False, dirty=True)

# %% [code]
# you can list all different results, they come as feature tables=counts, speaking names=taxonomy and ~importance=NSTI
res_picrust2_gut['results'].keys()

# %% [code]
res_picrust2_gut['results']['counts'].keys()

# %% [code]
res_picrust2_gut['results']['counts']['COG']

# %% [code]
res_picrust2_gut['results']['taxonomy']['COG']

# %% [code]
! rm -r /homes/arehm/TMP/ana_bdiv_945a7wzv

# %% [code]
# you can then compute e.g. beta diversity (but no UniFrac of course, since we lack a tree)
res_beta_picrust_gut = beta_diversity(res_picrust2_gut['results']['counts']['COG'], metrics=['bray_curtis'], dry=False, wait=True)

# %% [code]
# and compute emperor plots from these distances
emperor(meta, res_beta_picrust_gut['results'], fp_results="./", dry=False)

# %% [code]
sample_subset = meta[meta['genotype'].isin(["C57BL6/J","BALB/c"])
                     & (meta['host_body_product'] == 'UBERON:lumen of gut') 
                     & (meta['comparison'] == 0)
                     & meta['generation'].isin(['F3','F4'])
                    ]

# %% [code]
sample_subset

# %% [code]
res_beta['results']['bray_curtis']

# %% [code]
#emperor for each of the categories
for t in res_picrust2_gut['results']['counts'].keys():
    print(t)
    res_beta = beta_diversity(res_picrust2_gut['results']['counts'][t], metrics=['bray_curtis'], dry=False, wait=False)
    if res_beta['results'] is not None:
        res_emp = emperor(sample_subset, res_beta['results'], fp_results="./", infix=t + 'f3f4', dry=False, wait=False, )
        display(plotNetworks('genotype', sample_subset, alpha=None, beta=res_beta['results'], summarize=True)[0])

# %% [code]
import calour as ca
ca.set_log_level(40)

# %% [code]
%matplotlib inline
for t in res_picrust2_gut['results']['counts'].keys():
    print(t)
    pandas2biom("tmp.biom", res_picrust2_gut['results']['counts'][t].loc[:, set(sample_subset.index) & set(picrust_gut.columns)])
    sample_subset.to_csv("tmp.meta", sep="\t")
    e = ca.read_amplicon("tmp.biom", "tmp.meta", min_reads=0, normalize=10000)
    print(e)

    res_dsFDR = find_diff_taxa(e, sample_subset, ['genotype'], random_seed=42)
    
    p = plot_diff_taxa(res_picrust2_gut['results']['counts'][t], 
                   sample_subset['genotype'], 
                   res_dsFDR, #onlyusetaxa=taxa_rf_dsfdr, 
                   #taxonomy=res_all['taxonomy']['RDP']['results']['Taxon'],
                   min_mean_abundance=0.01)

# %% [markdown]
## rarifiziert
### Gut

# %% [code]
%time ! $q_path; qiime tools export \
  --input-path $work_dir/rarefied6000-filtered-gut-table.qza\
  --output-path $work_dir/exported-rarified6000-gut-table

# %% [code]
#exportieren der Biom tabellen -> verwenden derer die für Alphadiversität exportiert wurden, also OHNE rarefaction und OHNE Vergleichsproben
from ggmap.snippets import biom2pandas

picrust_gut = biom2pandas(work_dir + "/exported-rarified6000-gut-table/feature-table.biom").fillna(0)


# %% [code]
#run picrust2
res_picrust2_gut = picrust2(picrust_gut, dry=False, dirty=True, use_grid=False)

# %% [code]
# you can list all different results, they come as feature tables=counts, speaking names=taxonomy and ~importance=NSTI
res_picrust2_gut['results'].keys()

# %% [code]
res_picrust2_gut['results']['counts'].keys()

# %% [code]
res_picrust2_gut['results']['counts']['COG']

# %% [code]
res_picrust2_gut['results']['taxonomy']['COG']

# %% [code]
! rm -r /homes/arehm/TMP/ana_bdiv_945a7wzv

# %% [code]
# you can then compute e.g. beta diversity (but no UniFrac of course, since we lack a tree)
res_beta_picrust_gut = beta_diversity(res_picrust2_gut['results']['counts']['COG'], metrics=['bray_curtis'], dry=False, wait=True, use_grid=False)

# %% [code]
# and compute emperor plots from these distances
emperor(meta, res_beta_picrust_gut['results'], fp_results="./", dry=False, use_grid=False)

# %% [code]
sample_subset = meta[meta['genotype'].isin(["C57BL6/J","BALB/c"])
                     & (meta['host_body_product'] == 'UBERON:lumen of gut') 
                     & (meta['comparison'] == 0)
                     & meta['generation'].isin(['F3','F4'])
                    ]

# %% [code]
sample_subset

# %% [code]
!rm -r /homes/arehm/TMP/ana_bdiv_setznxkn

# %% [code]
#emperor for each of the categories
for t in res_picrust2_gut['results']['counts'].keys():
    print(t)
    res_beta_2 = beta_diversity(res_picrust2_gut['results']['counts'][t], metrics=['bray_curtis'], dry=False, wait=False, use_grid=False)
    if res_beta_2['results'] is not None:
        res_emp = emperor(sample_subset, res_beta_2['results'], fp_results="./", infix=t + 'f3f4', dry=False, wait=False, use_grid=False )
        display(plotNetworks('genotype', sample_subset, alpha=None, beta=res_beta_2['results'], summarize=True)[0], use_grid=False)

# %% [code]
import calour as ca
ca.set_log_level(40)

# %% [code]
%matplotlib inline
for t in res_picrust2_gut['results']['counts'].keys():
    print(t)
    pandas2biom("tmp.biom", res_picrust2_gut['results']['counts'][t].loc[:, set(sample_subset.index) & set(picrust_gut.columns)])
    sample_subset.to_csv("tmp.meta", sep="\t")
    e = ca.read_amplicon("tmp.biom", "tmp.meta", min_reads=0, normalize=10000)
    print(e)

    res_dsFDR = find_diff_taxa(e, sample_subset, ['genotype'], random_seed=42)
    
    p = plot_diff_taxa(res_picrust2_gut['results']['counts'][t], 
                   sample_subset['genotype'], 
                   res_dsFDR, #onlyusetaxa=taxa_rf_dsfdr, 
                   #taxonomy=res_all['taxonomy']['RDP']['results']['Taxon'],
                   min_mean_abundance=0.01)

# %% [code]


# %% [code]


# %% [code]


# %% [markdown]
# Alpha Diversity

# %% [markdown]
### Lumen of the Gut

# %% [markdown]
Table needs to be filtered first by bodysite as to avoid artificial diversity by combining skin and gut samples.

# %% [code]
meta = pd.read_csv(meta_file, sep="\t", dtype=str).set_index('sample_name')

# %% [code]
test = meta[meta['host_body_product'].isin(['UBERON:lumen of gut'])]
test['genotype'].value_counts()

# %% [code]
indeces = []
for i in range (64,88):
    index = "13422.colon3" + str(i)
    indeces.append(index)
    index = "13422.skin3" + str(i)
    indeces.append(index)
    index = "13422.3" + str(i) + "skin"
    indeces.append(index)


# %% [code]
comparison_mice_indeces = str(indeces)[1:-1]

# %% [code]
%time ! $q_path; qiime feature-table filter-samples \
  --i-table $work_dir/fragment-filtered‐feature-table.qza \
  --m-metadata-file $meta_file \
  --p-where "[sample_name] NOT IN ($comparison_mice_indeces) AND [host_body_product] IN ('UBERON:lumen of gut')" \
  --o-filtered-table $work_dir/filtered-gut-table-for-alpha.qza

# %% [code]
%time ! $q_path; qiime tools export \
  --input-path $work_dir/filtered-gut-table-for-alpha.qza \
  --output-path $work_dir/exported-filtered-gut-table-for-alpha

# %% [markdown]
Diversity was calculated using the ggmap package

# %% [code]
from ggmap.snippets import biom2pandas

filtered_gut_table = biom2pandas(work_dir + "/exported-filtered-gut-table-for-alpha/feature-table.biom").fillna(0)

# %% [code]
alpha_div_gut = alpha_diversity(filtered_gut_table, 6000, metrics=["shannon", "observed_features", "PD_whole_tree"], num_iterations=10, reference_tree="/homes/arehm/Benga_ThreeGen/FromQiita/128106_insertion_tree.relabelled.tre", use_grid=True, dry=False)

# %% [code]
alpha_div_gut['results']

# %% [code]
%time ! $q_path; qiime feature-table filter-samples \
  --i-table $work_dir/fragment-filtered‐feature-table.qza \
  --m-metadata-file $meta_file \
  --p-where "[host_body_product] IN ('UBERON:lumen of gut')" \
  --o-filtered-table $work_dir/filtered-gut-table-for-alpha_compincluded.qza

# %% [code]
%time ! $q_path; qiime tools export \
  --input-path $work_dir/filtered-gut-table-for-alpha_compincluded.qza \
  --output-path $work_dir/exported-filtered-gut-table-for-alpha_compincluded

# %% [code]
filtered_gut_table_comp_included = biom2pandas(work_dir + "/exported-filtered-gut-table-for-alpha_compincluded/feature-table.biom").fillna(0)
alpha_div_gut_comp_included = alpha_diversity(filtered_gut_table_comp_included, 6000, metrics=["shannon", "observed_features", "PD_whole_tree"], num_iterations=10, reference_tree="/homes/arehm/Benga_ThreeGen/FromQiita/128106_insertion_tree.relabelled.tre", use_grid=True, dry=False)

# %% [code]
alpha_results_merged_gut_comp = alpha_div_gut_comp_included['results'].merge(meta, left_index=True, right_index=True)
alpha_results_merged_gut_comp[alpha_results_merged_gut_comp['genotype'].isin(['C57BL6/J', 'BALB/c'])]

# %% [markdown]
##### Visual Comparison of Genotype Alpha Diversity

# %% [code]
#import pandas as pd
#alpha_shannon_gut = pd.read_csv(work_dir + "/alpha_shannon_gut.tsv", sep="\t", index_col=0)
#alpha_observed_gut = pd.read_csv(work_dir + "/alpha_observed_features_gut.tsv", sep="\t", index_col=0)
#alpha_faithpd_gut = pd.read_csv(work_dir + "/alpha_faithpd_gut.tsv", sep="\t", index_col=0)
#alpha_shannon_gut = alpha_shannon_gut.drop('#q2:types')
#alpha_observed_gut = alpha_observed_gut.drop('#q2:types')
#alpha_faithpd_gut = alpha_faithpd_gut.drop('#q2:types')

# %% [code]
alpha_results_merged_gut = alpha_div_gut['results'].merge(meta, left_index=True, right_index=True)
alpha_results_merged_gut[alpha_results_merged_gut['genotype'].isin(['C57BL6/J', 'BALB/c'])]

# %% [code]
%matplotlib inline

import seaborn as sns
import matplotlib.pyplot as plt

no_RM = alpha_results_merged_gut[alpha_results_merged_gut['genotype'].isin(['C57BL6/J', 'BALB/c'])]

cdf = no_RM.explode("shannon")
cdf["shannon"] = cdf["shannon"].astype("float")
order_list = ['P1', 'F1', 'F2', 'F3', 'F4', 'F5']

ax = sns.boxplot(x="genotype", y="shannon", hue="generation", data=cdf, hue_order=order_list)
plt.title("Shannon Distance over generations per Genotype")
plt.ylabel("Shannon Distance")
plt.legend(bbox_to_anchor=(1.05, 1), loc=2)
plt.show()


cdf = no_RM.explode("PD_whole_tree")
cdf["PD_whole_tree"] = cdf["PD_whole_tree"].astype("float")

ax = sns.boxplot(x="genotype", y="PD_whole_tree", hue="generation", data=cdf, hue_order=order_list)
plt.title("Faith's PD over generations per Genotype")
plt.ylabel("Faith's PD")
plt.legend(bbox_to_anchor=(1.05, 1), loc=2)
plt.show()

cdf = no_RM.explode("observed_features")
cdf["observed_features"] = cdf["observed_features"].astype("float")

ax = sns.boxplot(x="genotype", y="observed_features", hue="generation", data=cdf, hue_order=order_list)
plt.title("Observed OTUs over generations per Genotype")
plt.ylabel("Observed OTUs")
plt.legend(bbox_to_anchor=(1.05, 1), loc=2)
plt.show()

# %% [markdown]
#### Skin

# %% [code]
%time ! $q_path; qiime feature-table filter-samples \
  --i-table $work_dir/fragment-filtered‐feature-table.qza \
  --m-metadata-file $meta_file \
  --p-where "[sample_name] NOT IN ($comparison_mice_indeces) AND [host_body_product] IN ('UBERON:sebum')" \
  --o-filtered-table $work_dir/filtered-sebum-table-for-alpha.qza

%time ! $q_path; qiime tools export \
  --input-path $work_dir/filtered-sebum-table-for-alpha.qza \
  --output-path $work_dir/exported-filtered-sebum-table-for-alpha

# %% [code]
filtered_sebum_table = biom2pandas(work_dir + "/exported-filtered-sebum-table-for-alpha/feature-table.biom").fillna(0)
alpha_div_sebum = alpha_diversity(filtered_sebum_table, 1000, metrics=["shannon", "observed_features", "PD_whole_tree"], num_iterations=10, reference_tree="/homes/arehm/Benga_ThreeGen/FromQiita/128106_insertion_tree.relabelled.tre", use_grid=True, dry=False)

# %% [code]
alpha_results_merged_sebum[alpha_results_merged_sebum['comparison'].isin(['1'])]

# %% [code]
alpha_results_merged_sebum = alpha_div_sebum['results'].merge(meta, left_index=True, right_index=True)
alpha_results_merged_sebum[alpha_results_merged_sebum['genotype'].isin(['C57BL6/J', 'BALB/c'])]

# %% [code]
%time ! $q_path; qiime feature-table filter-samples \
  --i-table $work_dir/fragment-filtered‐feature-table.qza \
  --m-metadata-file $meta_file \
  --p-where "[host_body_product] IN ('UBERON:sebum')" \
  --o-filtered-table $work_dir/filtered-sebum-table-for-alpha_compincluded.qza

%time ! $q_path; qiime tools export \
  --input-path $work_dir/filtered-sebum-table-for-alpha_compincluded.qza \
  --output-path $work_dir/exported-filtered-sebum-table-for-alpha_compincluded

# %% [code]
filtered_sebum_table_comp_included = biom2pandas(work_dir + "/exported-filtered-sebum-table-for-alpha_compincluded/feature-table.biom").fillna(0)
alpha_div_sebum_comp = alpha_diversity(filtered_sebum_table_comp_included, 1000, metrics=["shannon", "observed_features", "PD_whole_tree"], num_iterations=10, reference_tree="/homes/arehm/Benga_ThreeGen/FromQiita/128106_insertion_tree.relabelled.tre", use_grid=True, dry=False)

# %% [code]
alpha_results_merged_sebum_comp = alpha_div_sebum_comp['results'].merge(meta, left_index=True, right_index=True)
alpha_results_merged_sebum_comp[alpha_results_merged_sebum_comp['genotype'].isin(['C57BL6/J', 'BALB/c'])]

# %% [markdown]
##### Visual Comparison of Genotype Alpha Diversity Skin

# %% [code]
#alpha_shannon_sebum = pd.read_csv(work_dir + "/alpha_shannon_sebum.tsv", sep="\t", index_col=0)
#alpha_observed_sebum = pd.read_csv(work_dir + "/alpha_observed_features_sebum.tsv", sep="\t", index_col=0)
#alpha_faithpd_sebum = pd.read_csv(work_dir + "/alpha_faithpd_sebum.tsv", sep="\t", index_col=0)
#alpha_shannon_sebum = alpha_shannon_sebum.drop('#q2:types')
#alpha_observed_sebum = alpha_observed_sebum.drop('#q2:types')
#alpha_faithpd_sebum = alpha_faithpd_sebum.drop('#q2:types')

# %% [code]
%matplotlib inline
no_RM = alpha_results_merged_sebum[alpha_results_merged_sebum['genotype'].isin(['C57BL6/J', 'BALB/c'])]
cdf = no_RM.explode("shannon")
cdf["shannon"] = cdf["shannon"].astype("float")
order_list = ['P1', 'F1', 'F2', 'F3', 'F4', 'F5']

ax = sns.boxplot(x="genotype", y="shannon", hue="generation", data=cdf, hue_order=order_list)
plt.title("Shannon Distance over generations per Genotype")
plt.ylabel("Shannon Distance")
plt.legend(bbox_to_anchor=(1.05, 1), loc=2)
plt.show()


cdf = no_RM.explode("PD_whole_tree")
cdf["PD_whole_tree"] = cdf["PD_whole_tree"].astype("float")

ax = sns.boxplot(x="genotype", y="PD_whole_tree", hue="generation", data=cdf, hue_order=order_list)
plt.title("Faith's PD over generations per Genotype")
plt.ylabel("Faith's PD")
plt.legend(bbox_to_anchor=(1.05, 1), loc=2)
plt.show()

cdf = no_RM.explode("observed_features")
cdf["observed_features"] = cdf["observed_features"].astype("float")

ax = sns.boxplot(x="genotype", y="observed_features", hue="generation", data=cdf, hue_order=order_list)
plt.title("Observed OTUs over generations per Genotype")
plt.ylabel("Observed OTUs")
plt.legend(bbox_to_anchor=(1.05, 1), loc=2)
plt.show()

# %% [markdown]
# Beta diversity

# %% [code]
%time ! $q_path; qiime diversity beta-phylogenetic \
  --i-table $work_dir/rarefied-filtered-feature-table.qza \
  --i-phylogeny $work_dir/insertion-tree.qza \
  --p-metric "unweighted_unifrac" \
  --o-distance-matrix $work_dir/beta_matrix.qza

%time ! $q_path; qiime diversity pcoa \
  --i-distance-matrix $work_dir/beta_matrix.qza \
  --o-pcoa $work_dir/beta_pcoa_unweighted.qza

%time ! $q_path; qiime emperor plot \
  --i-pcoa $work_dir/beta_pcoa_unweighted.qza \
  --m-metadata-file $meta_file \
  --o-visualization $work_dir/unweighted_emperor.qzv

# %% [markdown]
### Only mice

# %% [code]
%time ! $q_path; qiime feature-table filter-samples \
  --i-table $work_dir/rarefied-filtered-feature-table.qza \
  --m-metadata-file $meta_file \
  --p-where "[host_common_name] IN ('marmoset') OR [host_common_name] IN ('not applicable') OR [host_body_product] IN ('UBERON:lung')" \
  --p-exclude-ids True\
  --o-filtered-table $work_dir/mouse-filtered-table-new.qza

%time ! $q_path; qiime diversity beta-phylogenetic \
  --i-table $work_dir/mouse-filtered-table-new.qza \
  --i-phylogeny $work_dir/insertion-tree.qza \
  --p-metric "unweighted_unifrac" \
  --o-distance-matrix $work_dir/mouse_beta_matrix-new.qza

%time ! $q_path; qiime diversity pcoa \
  --i-distance-matrix $work_dir/mouse_beta_matrix-new.qza \
  --o-pcoa $work_dir/mouse_beta_pcoa_unweighted-new.qza

%time ! $q_path; qiime emperor plot \
  --i-pcoa $work_dir/mouse_beta_pcoa_unweighted-new.qza \
  --m-metadata-file $meta_file \
  --o-visualization $work_dir/mouse_unweighted_emperor-new.qzv

# %% [markdown]
### Per Batch

# %% [code]
%time ! $q_path; qiime feature-table filter-samples \
  --i-table $work_dir/rarefied-filtered-feature-table.qza \
  --m-metadata-file $meta_file \
  --p-where "[host_common_name] IN ('mouse') AND [batch] IN ('1') " \
  --o-filtered-table $work_dir/mouse-filtered-table-batch1.qza

%time ! $q_path; qiime diversity beta-phylogenetic \
  --i-table $work_dir/mouse-filtered-table-batch1.qza \
  --i-phylogeny $work_dir/insertion-tree.qza \
  --p-metric "unweighted_unifrac" \
  --o-distance-matrix $work_dir/mouse_beta_matrix_batch1.qza

%time ! $q_path; qiime diversity pcoa \
  --i-distance-matrix $work_dir/mouse_beta_matrix_batch1.qza \
  --o-pcoa $work_dir/mouse_beta_pcoa_unweighted_batch1.qza

%time ! $q_path; qiime emperor plot \
  --i-pcoa $work_dir/mouse_beta_pcoa_unweighted_batch1.qza \
  --m-metadata-file $meta_file \
  --o-visualization $work_dir/mouse_unweighted_emperor_batch1.qzv

# %% [code]
%time ! $q_path; qiime feature-table filter-samples \
  --i-table $work_dir/rarefied-filtered-feature-table.qza \
  --m-metadata-file $meta_file \
  --p-where "[host_common_name] IN ('mouse') AND [batch] IN ('2') AND [sample_name] NOT IN ($comparison_mice_indeces)" \
  --o-filtered-table $work_dir/mouse-filtered-table-batch2.qza

%time ! $q_path; qiime diversity beta-phylogenetic \
  --i-table $work_dir/mouse-filtered-table-batch2.qza \
  --i-phylogeny $work_dir/insertion-tree.qza \
  --p-metric "unweighted_unifrac" \
  --o-distance-matrix $work_dir/mouse_beta_matrix_batch2.qza

%time ! $q_path; qiime diversity pcoa \
  --i-distance-matrix $work_dir/mouse_beta_matrix_batch2.qza \
  --o-pcoa $work_dir/mouse_beta_pcoa_unweighted_batch2.qza

%time ! $q_path; qiime emperor plot \
  --i-pcoa $work_dir/mouse_beta_pcoa_unweighted_batch2.qza \
  --m-metadata-file $meta_file \
  --o-visualization $work_dir/mouse_unweighted_emperor_batch2.qzv

# %% [code]
%time ! $q_path; qiime feature-table filter-samples \
  --i-table $work_dir/rarefied-filtered-feature-table.qza \
  --m-metadata-file $meta_file \
  --p-where "[host_common_name] IN ('mouse') AND [batch] IN ('3') AND [sample_name] NOT IN ($comparison_mice_indeces)" \
  --o-filtered-table $work_dir/mouse-filtered-table-batch3.qza

%time ! $q_path; qiime diversity beta-phylogenetic \
  --i-table $work_dir/mouse-filtered-table-batch3.qza \
  --i-phylogeny $work_dir/insertion-tree.qza \
  --p-metric "unweighted_unifrac" \
  --o-distance-matrix $work_dir/mouse_beta_matrix_batch3.qza

%time ! $q_path; qiime diversity pcoa \
  --i-distance-matrix $work_dir/mouse_beta_matrix_batch3.qza \
  --o-pcoa $work_dir/mouse_beta_pcoa_unweighted_batch3.qza

%time ! $q_path; qiime emperor plot \
  --i-pcoa $work_dir/mouse_beta_pcoa_unweighted_batch3.qza \
  --m-metadata-file $meta_file \
  --o-visualization $work_dir/mouse_unweighted_emperor_batch3.qzv

# %% [markdown]
### Per Body Product
#### Gut with comparison

# %% [code]
%time ! $q_path; qiime feature-table filter-samples \
  --i-table $work_dir/rarefied6000-filtered-gut-table.qza \
  --m-metadata-file $meta_file \
  --p-where "[host_common_name]='mouse'" \
  --o-filtered-table $work_dir/gut6000-mouse-filtered-table.qza

# %% [code]
%time ! $q_path; qiime diversity beta-phylogenetic \
  --i-table $work_dir/gut6000-mouse-filtered-table.qza \
  --i-phylogeny $work_dir/insertion-tree.qza \
  --p-metric "unweighted_unifrac" \
  --o-distance-matrix $work_dir/gut6000_beta_matrix.qza

%time ! $q_path; qiime diversity pcoa \
  --i-distance-matrix $work_dir/gut6000_beta_matrix.qza \
  --o-pcoa $work_dir/gut6000_beta_pcoa_unweighted.qza

# %% [code]
%time ! $q_path; qiime emperor plot \
  --i-pcoa $work_dir/gut6000_beta_pcoa_unweighted.qza \
  --m-metadata-file meta_gen_num.txt \
  --o-visualization $work_dir/gut6000_unweighted_emperor.qzv

# %% [markdown]
#### Gut without comparison

# %% [markdown]
2022/11/15: interesting development here on the beta diversity front for gut samples. The cage lineage C3 seems to be moving into a different direction than C1 and C2 -> instead of "moving down" it seems to be "moving up" or rather staying in the same place. this might be the duality of samples Celeste was talking about 

# %% [code]
%time ! $q_path; qiime feature-table filter-samples \
  --i-table $work_dir/gut6000-mouse-filtered-table.qza \
  --m-metadata-file $meta_file \
  --p-where "[sample_name] NOT IN ($comparison_mice_indeces)" \
  --o-filtered-table $work_dir/gut6000-mouse-filtered-table-nocomp.qza

# %% [code]
%time ! $q_path; qiime diversity beta-phylogenetic \
  --i-table $work_dir/gut6000-mouse-filtered-table-nocomp.qza \
  --i-phylogeny $work_dir/insertion-tree.qza \
  --p-metric "unweighted_unifrac" \
  --o-distance-matrix $work_dir/gut6000_beta_matrix-nocomp.qza

%time ! $q_path; qiime diversity pcoa \
  --i-distance-matrix $work_dir/gut6000_beta_matrix-nocomp.qza \
  --o-pcoa $work_dir/gut6000_beta_pcoa_unweighted-nocomp.qza

%time ! $q_path; qiime emperor plot \
  --i-pcoa $work_dir/gut6000_beta_pcoa_unweighted-nocomp.qza \
  --m-metadata-file meta_gen_num.txt \
  --o-visualization $work_dir/gut6000_unweighted_emperor-nocomp.qzv

# %% [code]
%time ! $q_path; qiime diversity beta\
  --i-table $work_dir/gut6000-mouse-filtered-table-nocomp.qza \
  --p-metric "braycurtis" \
  --o-distance-matrix $work_dir/gut6000_beta_braycurtis_matrix-nocomp.qza

%time ! $q_path; qiime diversity beta-phylogenetic \
  --i-table $work_dir/gut6000-mouse-filtered-table-nocomp.qza \
  --i-phylogeny $work_dir/insertion-tree.qza \
  --p-metric "weighted_unifrac" \
  --o-distance-matrix $work_dir/gut6000_beta_weighted_unifrac_matrix-nocomp.qza

# %% [code]
%time ! $q_path; qiime diversity pcoa \
  --i-distance-matrix $work_dir/gut6000_beta_braycurtis_matrix-nocomp.qza \
  --o-pcoa $work_dir/gut6000_beta_pcoa_braycurtis-nocomp.qza

%time ! $q_path; qiime diversity pcoa \
  --i-distance-matrix $work_dir/gut6000_beta_weighted_unifrac_matrix-nocomp.qza \
  --o-pcoa $work_dir/gut6000_beta_pcoa_weighted-nocomp.qza

# %% [code]
%time ! $q_path; qiime emperor plot \
  --i-pcoa  $work_dir/gut6000_beta_pcoa_weighted-nocomp.qza \
  --m-metadata-file $meta_file \
  --o-visualization $work_dir/gut6000_unweighted_emperor-nocomp.qzv

# %% [code]
%time ! $q_path; qiime emperor plot \
  --i-pcoa $work_dir/gut_beta_pcoa_unweighted-nocomp.qza \
  --m-metadata-file meta_gen_num.txt \
  --p-custom-axes gen_number \
  --o-visualization $work_dir/gut_unweighted_emperor-nocomp-time.qzv

# %% [code]
%time ! $q_path; qiime diversity beta-group-significance \
  --i-distance-matrix $work_dir/gut_beta_matrix-nocomp.qza \
  --m-metadata-file meta_gen_num.txt \
  --m-metadata-column i_generation \
  --p-pairwise \
  --p-permutations 9999 \
  --o-visualization $work_dir/gut_beta_group_significance_nopair.qzv

# %% [markdown]
#### Sebum with comparison

# %% [code]
%time ! $q_path; qiime feature-table filter-samples \
  --i-table $work_dir/rarefied-filtered-sebum-table.qza \
  --m-metadata-file $meta_file \
  --p-where "[host_common_name]='mouse'" \
  --o-filtered-table $work_dir/sebum-mouse-filtered-table.qza

%time ! $q_path; qiime diversity beta-phylogenetic \
  --i-table $work_dir/sebum-mouse-filtered-table.qza \
  --i-phylogeny $work_dir/insertion-tree.qza \
  --p-metric "unweighted_unifrac" \
  --o-distance-matrix $work_dir/sebum_beta_matrix.qza

%time ! $q_path; qiime diversity pcoa \
  --i-distance-matrix $work_dir/sebum_beta_matrix.qza \
  --o-pcoa $work_dir/sebum_beta_pcoa_unweighted.qza

# %% [code]
%time ! $q_path; qiime emperor plot \
  --i-pcoa $work_dir/sebum_beta_pcoa_unweighted.qza \
  --m-metadata-file meta_gen_num.txt \
  --o-visualization $work_dir/sebum_unweighted_emperor.qzv

# %% [markdown]
#### Sebum without comparison

# %% [code]
%time ! $q_path; qiime feature-table filter-samples \
  --i-table $work_dir/sebum-mouse-filtered-table.qza \
  --m-metadata-file $meta_file \
  --p-where "[sample_name] NOT IN ($comparison_mice_indeces)" \
  --o-filtered-table $work_dir/sebum-mouse-filtered-table-nocomp.qza

%time ! $q_path; qiime diversity beta-phylogenetic \
  --i-table $work_dir/sebum-mouse-filtered-table-nocomp.qza \
  --i-phylogeny $work_dir/insertion-tree.qza \
  --p-metric "unweighted_unifrac" \
  --o-distance-matrix $work_dir/sebum_beta_matrix-nocomp.qza

%time ! $q_path; qiime diversity pcoa \
  --i-distance-matrix $work_dir/sebum_beta_matrix-nocomp.qza \
  --o-pcoa $work_dir/sebum_beta_pcoa_unweighted-nocomp.qza

%time ! $q_path; qiime emperor plot \
  --i-pcoa $work_dir/sebum_beta_pcoa_unweighted-nocomp.qza \
  --m-metadata-file meta_gen_num.txt \
  --o-visualization $work_dir/sebum_unweighted_emperor-nocomp.qzv

# %% [code]
%time ! $q_path; qiime diversity beta \
  --i-table $work_dir/sebum-mouse-filtered-table-nocomp.qza \
  --p-metric "braycurtis" \
  --o-distance-matrix $work_dir/sebum_beta_braycurtis_matrix-nocomp.qza

%time ! $q_path; qiime diversity beta-phylogenetic \
  --i-table $work_dir/sebum-mouse-filtered-table-nocomp.qza \
  --i-phylogeny $work_dir/insertion-tree.qza \
  --p-metric "weighted_unifrac" \
  --o-distance-matrix $work_dir/sebum_beta_weighted_unifrac_matrix-nocomp.qza

# %% [code]
%time ! $q_path; qiime diversity pcoa \
  --i-distance-matrix $work_dir/sebum_beta_braycurtis_matrix-nocomp.qza \
  --o-pcoa $work_dir/sebum_beta_pcoa_braycurtis-nocomp.qza

%time ! $q_path; qiime diversity pcoa \
  --i-distance-matrix $work_dir/sebum_beta_weighted_unifrac_matrix-nocomp.qza \
  --o-pcoa $work_dir/sebum_beta_pcoa_weighted-nocomp.qza

# %% [code]
%time ! $q_path; qiime emperor plot \
  --i-pcoa $work_dir/sebum_beta_pcoa_unweighted-nocomp.qza \
  --m-metadata-file meta_gen_num.txt \
  --p-custom-axes gen_number \
  --o-visualization $work_dir/sebum_unweighted_emperor-nocomp-time.qzv

# %% [code]
%time ! $q_path; qiime diversity beta-group-significance \
  --i-distance-matrix $work_dir/sebum_beta_matrix-nocomp.qza \
  --m-metadata-file meta_gen_num.txt \
  --m-metadata-column i_generation \
  --p-permutations 9999 \
  --o-visualization $work_dir/sebum_beta_group_significance_nopair.qzv

# %% [markdown]
## Taxonomy barplots

# %% [code]
%time ! $q_path; qiime feature-table summarize \
  --i-table $work_dir/rarefied-filtered-noCyanobacteria-feature-table-all.qza \
  --o-visualization $work_dir/summary-rarefied-filtered-noCyanobacteria-feature-table.qzv \
  --m-sample-metadata-file $meta_file

# %% [code]
%time ! $q_path; qiime taxa barplot \
  --i-table $work_dir/fragment-filtered‐feature-table.qza \
  --i-taxonomy $work_dir/taxonomy_full.qza \
  --m-metadata-file meta_gen_num.txt \
  --o-visualization $work_dir/taxa_barplot_full.qzv

# %% [code]
%time ! $q_path; qiime tools export \
  --input-path $work_dir/taxonomy_full.qza \
  --output-path $work_dir/exported-taxonomy-full

# %% [code]
%time ! $q_path; qiime tools export \
  --input-path $work_dir/fragment-filtered‐feature-table.qza \
  --output-path $work_dir/exported-fragment-filtered-feature-table

# %% [code]
#taxa_all_l2 = pd.read_csv("/homes/arehm/Documents/Thesis/JupyterNotebook/taxa/level-2.csv", sep=",").set_index('index')
#taxa_all_l3 = pd.read_csv("/homes/arehm/Documents/Thesis/JupyterNotebook/taxa/level-3.csv", sep=",").set_index('index')
#taxa_all_l4 = pd.read_csv("/homes/arehm/Documents/Thesis/JupyterNotebook/taxa/level-4.csv", sep=",").set_index('index')
#taxa_all_l5 = pd.read_csv("/homes/arehm/Documents/Thesis/JupyterNotebook/taxa/level-5.csv", sep=",").set_index('index')
taxa_all_l6 = pd.read_csv("/homes/arehm/Documents/Thesis/JupyterNotebook/taxa/level-6.csv", sep=",").set_index('index')
#taxa_all_l7 = pd.read_csv("/homes/arehm/Documents/Thesis/JupyterNotebook/taxa/level-7.csv", sep=",").set_index('index')

# %% [code]
taxa_all_l5 = taxa_all_l5[taxa_all_l5['host_body_product'].isin(['UBERON:lumen of gut', 'UBERON:sebum']) & taxa_all_l5['host_scientific_name'].isin(['Mus musculus']) & taxa_all_l5['genotype'].isin(['BALB/c', 'C57BL6/J'])]
filtered_taxa = taxa_all_l5.groupby(["host_body_product","genotype"]).sum()
filtered_taxa = filtered_taxa.drop(columns=['batch', 'dna_extracted', 'elevation', 'latitude', 'longitude', 'physical_specimen_remaining', 'qiita_study_id', 'taxon_id', 'gen_number', 'comparison'])
#taxa_all_l5['percent'] = (taxa_all_l5['sum']/taxa_all_l5['sum'].sum())*100
#taxa_all_l5 = taxa_all_l5[~taxa_all_l5.index.str.endswith("__")]
filtered_taxa = filtered_taxa.loc[:,~filtered_taxa.columns.str.endswith('__')]

# %% [code]


# %% [code]
exported_taxonomy = pd.read_csv("exported-taxonomy-full/taxonomy.tsv", sep="\t").set_index("Feature ID")

# %% [code]
exported_taxonomy[['Taxon1', 'Taxon2']] = exported_taxonomy["Taxon"].apply(lambda x: pd.Series(str(x).split(" c__")))

# %% [code]
phylum_taxa= exported_taxonomy.drop(columns=["Taxon2", "Taxon"])
#phylum_taxa = phylum_taxa.loc[:,~phylum_taxa.columns.str.endswith('k__Bacteria')]
phylum_taxa.rename(columns={'Taxon1':'Taxon'}, inplace=True)
phylum_taxa = phylum_taxa[["Taxon", "Confidence"]]
phylum_taxa = phylum_taxa.drop(phylum_taxa[phylum_taxa.Taxon == "k__Bacteria"].index)
phylum_taxa.to_csv("exported-taxonomy-full/phylum.tsv", sep="\t")

# %% [code]


# %% [code]
filtered_exported_taxonomy = exported_taxonomy.loc[:,~exported_taxonomy.columns.str.endswith('__')]

# %% [code]
filtered_exported_taxonomy.to_csv("exported-taxonomy-full/filtered-taxonomy.tsv", header=True, sep="\t")

# %% [code]
exported_taxonomy.to_csv("exported-taxonomy-full/test.tsv", header=True, sep="\t")

# %% [code]
%matplotlib inline
fig = plotTaxonomy("exported-fragment-filtered-feature-table/feature-table.biom", meta[meta['host_body_product'].isin(['UBERON:lumen of gut', 'UBERON:sebum']) & meta['host_scientific_name'].isin(['Mus musculus']) & meta['genotype'].isin(['BALB/c', 'C57BL6/J'])],  file_taxonomy="exported-taxonomy-full/phylum.tsv", fct_aggregate=np.mean, group_l0="host_body_product",group_l1="genotype", rank="Phylum", minreadnr=100)
plt.show()

# %% [code]
exported_taxonomy[['Taxon1', 'Taxon2']] = exported_taxonomy["Taxon"].apply(lambda x: pd.Series(str(x).split(" g__")))
family_taxa= exported_taxonomy.drop(columns=["Taxon2", "Taxon"])
#phylum_taxa = phylum_taxa.loc[:,~phylum_taxa.columns.str.endswith('k__Bacteria')]
family_taxa.rename(columns={'Taxon1':'Taxon'}, inplace=True)
family_taxa = family_taxa[["Taxon", "Confidence"]]
#family_taxa = family_taxa.drop(family_taxa[family_taxa.Taxon == "k__Bacteria"].index)
#family_taxa = family_taxa[~(family_taxa.Taxon == "k__Bacteria") | (family_taxa.Taxon == "k__Bacteria; p__Firmicutes; c__Clostridia; o__Clostridiales")]
#family_taxa = family_taxa[~(family_taxa.Taxon == "k__Bacteria; p__Firmicutes; c__Clostridia; o__Clostridiales") & ~(family_taxa.Taxon == "k__Bacteria") & ~(family_taxa.Taxon == "k__Bacteria; p__Firmicutes") & ~(family_taxa.Taxon == "k__Bacteria; p__Firmicutes; c__Clostridia")]
family_taxa = family_taxa[~(family_taxa['Taxon'].str.endswith('f__;')) & ~(family_taxa['Taxon'].str.endswith('f__;'))]
family_taxa = family_taxa[family_taxa["Taxon"].str.contains("f__")]
family_taxa.to_csv("exported-taxonomy-full/family.tsv", sep="\t")

# %% [code]
family_taxa["Taxon"].value_counts()

# %% [code]
%matplotlib inline
fig, rank_counts, graphinfo, vals, colors = plotTaxonomy("exported-fragment-filtered-feature-table/feature-table.biom", meta[meta['host_body_product'].isin(['UBERON:lumen of gut', 'UBERON:sebum']) & meta['host_scientific_name'].isin(['Mus musculus']) & meta['genotype'].isin(['BALB/c', 'C57BL6/J'])],  file_taxonomy="exported-taxonomy-full/family.tsv", fct_aggregate=np.mean, group_l0="host_body_product",group_l1="genotype", rank="Family", minreadnr=5000)

# %% [markdown]
###### all levels

# %% [code]
for taxa_level in [taxa_all_l2, taxa_all_l3,taxa_all_l4,taxa_all_l5,taxa_all_l6,taxa_all_l7]:
    taxa_gut = taxa_level[taxa_level['host_body_product'].isin(['UBERON:lumen of gut']) & taxa_level['host_scientific_name'].isin(['Mus musculus'])]
    taxa_sebum = taxa_level[taxa_level['host_body_product'].isin(['UBERON:sebum']) & taxa_level['host_scientific_name'].isin(['Mus musculus'])]
    for i, body_product in enumerate([taxa_gut, taxa_sebum]):
        product = "Gut" if i == 0 else "Sebum"
        taxa_body_sum_all =body_product.sum(axis=0, skipna=True)[:-58].to_frame(name='sum').sort_values('sum', ascending=False)
        taxa_body_sum_all['percent'] = (taxa_body_sum_all['sum'] / taxa_body_sum_all['sum'].sum())*100
        taxa_body_sum_all = taxa_body_sum_all[~taxa_body_sum_all.index.str.endswith("__")]
        plt.figure(figsize=(15,8))
        ax = sns.barplot(x=taxa_body_sum_all.index, y=taxa_body_sum_all['sum'])
        ax.set(yscale="log", title = f"Absolute abundance of taxa in the {product}")
        plt.xticks(rotation=90, ha="right")
        plt.show()
        plt.figure(figsize=(15,8))
        ax = sns.barplot(x=taxa_body_sum_all.index, y=taxa_body_sum_all['percent'])
        ax.set(yscale = "log", title = f"Relative abundance of taxa in the {product}")
        plt.xticks(rotation=90, ha="right")
        plt.show()

# %% [code]
taxa_l2_gut = taxa_all_l2[taxa_all_l2['host_body_product'].isin(['UBERON:lumen of gut']) & taxa_all_l2['host_scientific_name'].isin(['Mus musculus'])]
taxa_l2_sebum = taxa_all_l2[taxa_all_l2['host_body_product'].isin(['UBERON:sebum']) & taxa_all_l2['host_scientific_name'].isin(['Mus musculus'])]

# %% [code]
taxa_gut_sum_all =taxa_gut.sum(axis=0, skipna=True)[:-58].to_frame(name='sum').sort_values('sum', ascending=False)
taxa_gut_sum_all['percent'] = (taxa_gut_sum_all['sum'] / taxa_gut_sum_all['sum'].sum())*100

# %% [code]


# %% [markdown]
##### Sebum

# %% [code]
%time ! $q_path; qiime feature-table filter-samples \
  --i-table /homes/arehm/Benga_ThreeGen/FromQiita/128106_feature-table.qza \
  --m-metadata-file $meta_file \
  --p-where "[sample_name] NOT IN ($comparison_mice_indeces) AND [host_body_product] IN ('UBERON:sebum') AND [genotype] IN ('C57BL6/J', 'BALB/c')" \
  --o-filtered-table $work_dir/dsFDR_table_sebum.qza

%time ! $q_path; qiime tools export \
  --input-path $work_dir/dsFDR_table_sebum.qza \
  --output-path $work_dir/dsFDR_table_sebum_export

# %% [markdown]
## Taxonomy Scatterplots

# %% [code]
import seaborn as sns

# %% [code]
v1 = taxa_gut.iloc[1]['k__Bacteria;p__Verrucomicrobia;c__Verrucomicrobiae;o__Verrucomicrobiales;f__Verrucomicrobiaceae;g__Akkermansia']
otal_abundance_akkermansia = taxa_gut['k__Bacteria;p__Verrucomicrobia;c__Verrucomicrobiae;o__Verrucomicrobiales;f__Verrucomicrobiaceae;g__Akkermansia'].sum()
result = v1 / otal_abundance_akkermansia
print(result)

# %% [code]
#use level6 table as Akkermansia is a genus
taxa_gut = taxa_all_l6[taxa_all_l6['host_body_product'].isin(['UBERON:lumen of gut']) & taxa_all_l6['host_scientific_name'].isin(['Mus musculus']) & ~taxa_all_l6['generation'].isin(['not applicable'])]
taxa_gut['k__Bacteria;p__Verrucomicrobia;c__Verrucomicrobiae;o__Verrucomicrobiales;f__Verrucomicrobiaceae;g__Akkermansia'].astype(float)
total_abundance_akkermansia = taxa_gut['k__Bacteria;p__Verrucomicrobia;c__Verrucomicrobiae;o__Verrucomicrobiales;f__Verrucomicrobiaceae;g__Akkermansia'].sum()
taxa_gut['relative_abundance'] = taxa_gut['k__Bacteria;p__Verrucomicrobia;c__Verrucomicrobiae;o__Verrucomicrobiales;f__Verrucomicrobiaceae;g__Akkermansia'] / total_abundance_akkermansia

# %% [code]
%matplotlib inline
taxa_all_l6[taxa_all_l6['comparison'].isin(['1'])]

# %% [code]
#goal: scatterplot with genotype on x axis and relative abundance of akkermansia on the y axis
%matplotlib inline
ax = sns.stripplot(x= 'generation', y='relative_abundance', data= taxa_gut, hue='genotype', order=['RM','P1', 'F1', 'F2', 'F3', 'F4', 'F5'], hue_order=['C57BL6/J', 'BALB/c', 'B6CF1'],dodge=True)
plt.savefig(work_dir + "/scatter_akkermansia.png", transparent=False, bbox_inches='tight')
plt.show()

# %% [code]


# %% [code]
taxa_gut_P1 = taxa_all_l6[taxa_all_l6['host_body_product'].isin(['UBERON:lumen of gut']) & taxa_all_l6['host_scientific_name'].isin(['Mus musculus']) & ~taxa_all_l6['generation'].isin(['not applicable']) & taxa_all_l6['generation'].isin(['P1'])]
taxa_gut_F1 = taxa_all_l6[taxa_all_l6['host_body_product'].isin(['UBERON:lumen of gut']) & taxa_all_l6['host_scientific_name'].isin(['Mus musculus']) & ~taxa_all_l6['generation'].isin(['not applicable']) & taxa_all_l6['generation'].isin(['F1'])]
taxa_gut_F2 = taxa_all_l6[taxa_all_l6['host_body_product'].isin(['UBERON:lumen of gut']) & taxa_all_l6['host_scientific_name'].isin(['Mus musculus']) & ~taxa_all_l6['generation'].isin(['not applicable']) & taxa_all_l6['generation'].isin(['F2'])]
taxa_gut_F3 = taxa_all_l6[taxa_all_l6['host_body_product'].isin(['UBERON:lumen of gut']) & taxa_all_l6['host_scientific_name'].isin(['Mus musculus']) & ~taxa_all_l6['generation'].isin(['not applicable']) & taxa_all_l6['generation'].isin(['F3'])]
taxa_gut_F4 = taxa_all_l6[taxa_all_l6['host_body_product'].isin(['UBERON:lumen of gut']) & taxa_all_l6['host_scientific_name'].isin(['Mus musculus']) & ~taxa_all_l6['generation'].isin(['not applicable']) & taxa_all_l6['generation'].isin(['F4'])]
taxa_gut_F5 = taxa_all_l6[taxa_all_l6['host_body_product'].isin(['UBERON:lumen of gut']) & taxa_all_l6['host_scientific_name'].isin(['Mus musculus']) & ~taxa_all_l6['generation'].isin(['not applicable']) & taxa_all_l6['generation'].isin(['F5'])]

# %% [code]
gen_dict={
    '0': 'P1',
    '1': 'F1',
    '2': 'F2',
    '3': 'F3',
    '4': 'F4',
    '5':'F5'
}

for i,generation in enumerate([taxa_gut_P1, taxa_gut_F1, taxa_gut_F2, taxa_gut_F3, taxa_gut_F4, taxa_gut_F5]):
    %matplotlib inline
    gen = gen_dict[str(i)]
    generation['k__Bacteria;p__Verrucomicrobia;c__Verrucomicrobiae;o__Verrucomicrobiales;f__Verrucomicrobiaceae;g__Akkermansia'].astype(float)
    total_abundance_akkermansia = generation['k__Bacteria;p__Verrucomicrobia;c__Verrucomicrobiae;o__Verrucomicrobiales;f__Verrucomicrobiaceae;g__Akkermansia'].sum()
    generation['relative_abundance'] = generation['k__Bacteria;p__Verrucomicrobia;c__Verrucomicrobiae;o__Verrucomicrobiales;f__Verrucomicrobiaceae;g__Akkermansia'] / total_abundance_akkermansia
    ax = sns.stripplot(x= 'genotype', y='relative_abundance', data= generation).set(title=f'Relative Abundance of Akkermansia per Genotype in {gen}')
    plt.show()

# %% [markdown]
### ANCOM
#### Only mice without comparison mice (GUT)

# %% [code]
import matplotlib.pyplot as plt

# %% [code]
#ANCOM gut total
%time !$q_path; qiime taxa collapse \
  --i-table $work_dir/gut6000-mouse-filtered-table-nocomp.qza \
  --i-taxonomy $work_dir/taxonomy_full.qza \
  --p-level 7 \
  --o-collapsed-table $work_dir/collapsed-level7-mouse-gut-table-nocomp.qza
%time !$q_path; qiime composition add-pseudocount \
  --i-table $work_dir/collapsed-level7-mouse-gut-table-nocompRM.qza \
  --o-composition-table $work_dir/composition-table-mouse-gut-l7-nocompRM.qza
%time !$q_path; qiime composition ancom \
  --i-table $work_dir/composition-table-mouse-gut-l7-nocompRM.qza \
  --m-metadata-file $meta_file \
  --m-metadata-column genotype \
  --o-visualization $work_dir/ancom-genotype-mouse-gut-l7-nocompRM.qzv

# %% [code]
#ANCOM for specific generations
%time !$q_path; qiime taxa collapse \
  --i-table $work_dir/mouse-table-gut6000-nocompRM_F1.qza \
  --i-taxonomy $work_dir/taxonomy_full.qza \
  --p-level 7 \
  --o-collapsed-table $work_dir/collapsed-level7-mouse-gut-table-nocompRM_F1.qza
%time !$q_path; qiime composition add-pseudocount \
  --i-table $work_dir/collapsed-level7-mouse-gut-table-nocompRM_F1.qza \
  --o-composition-table $work_dir/composition-table-mouse-gut-l7-nocompRM_F1.qza
%time !$q_path; qiime composition ancom \
  --i-table $work_dir/composition-table-mouse-gut-l7-nocompRM_F1.qza \
  --m-metadata-file $meta_file \
  --m-metadata-column genotype \
  --o-visualization $work_dir/ancom-genotype-mouse-gut-l7-nocompRM_F1.qzv

# %% [code]
!unzip $work_dir/composition-table-mouse-gut-l7-nocompRM_F1.qza

# %% [code]
! $q_path; biom convert \
  --input-fp 0847fa0f-7390-4411-a385-ffd3996c3c15/data/feature-table.biom       \
  --to-tsv \
  --output-fp $work_dir/ancom-biom-table-gut6-F1 

# %% [code]
ancom_biom_table_gut = pd.read_csv(work_dir+"/ancom-biom-table-gut6-F1", 
            sep="\t",skiprows=1, index_col=0)
ancom_biom_table_gut.sum()

# %% [code]
relative_abundance_gut= ancom_biom_table_gut.div(6053)
list_C57B = meta[meta['genotype'].isin(['C57BL6/J'])].index
list_BALB = meta[meta['genotype'].isin(['BALB/c'])].index

ancom_C57B_gut=relative_abundance_gut[[col for col in relative_abundance_gut.columns if col in list_C57B]]
ancom_C57B_gut["average"]=ancom_C57B_gut.mean(axis=1)

ancom_BALB_gut = relative_abundance_gut[[col for col in relative_abundance_gut.columns if col in list_BALB]]
ancom_BALB_gut["average"] = ancom_BALB_gut.mean(axis=1)

%matplotlib inline
#ancom_taxa_gut_F1 = ['k__Bacteria;p__Verrucomicrobia;c__Verrucomicrobiae;o__Verrucomicrobiales;f__Verrucomicrobiaceae;g__Akkermansia;s__muciniphila']
#ancom_taxa_gut_F2=['k__Bacteria;p__Verrucomicrobia;c__Verrucomicrobiae;o__Verrucomicrobiales;f__Verrucomicrobiaceae;g__Akkermansia;s__muciniphila']
#ancom_taxa_gut_F3=['k__Bacteria;p__Bacteroidetes;c__Bacteroidia;o__Bacteroidales;f__Bacteroidaceae;g__Bacteroides;s__acidifaciens']
#ancom_taxa_gut_F4 = ['k__Bacteria;p__Bacteroidetes;c__Bacteroidia;o__Bacteroidales;f__Bacteroidaceae;g__Bacteroides;s__acidifaciens', 'k__Bacteria;p__Actinobacteria;c__Actinobacteria;o__Bifidobacteriales;f__Bifidobacteriaceae;g__Bifidobacterium;s__pseudolongum']
#ancom_taxa_gut_F5=['k__Bacteria;p__Bacteroidetes;c__Bacteroidia;o__Bacteroidales;f__Bacteroidaceae;g__Bacteroides;s__acidifaciens']
#ancom_taxa_gut_F5-6=['k__Bacteria;p__Firmicutes;c__Erysipelotrichi;o__Erysipelotrichales;f__Erysipelotrichaceae;g__Allobaculum']
#ancom_taxa_gut_F4-6 = ['k__Bacteria;p__Firmicutes;c__Erysipelotrichi;o__Erysipelotrichales;f__Erysipelotrichaceae;g__Allobaculum', 'k__Bacteria;p__Bacteroidetes;c__Bacteroidia;o__Bacteroidales;f__Bacteroidaceae;g__Bacteroides', 'k__Bacteria;p__Actinobacteria;c__Actinobacteria;o__Bifidobacteriales;f__Bifidobacteriaceae;g__Bifidobacterium', 'k__Bacteria;p__Bacteroidetes;c__Bacteroidia;o__Bacteroidales;f__[Odoribacteraceae];g__Odoribacter', 'k__Bacteria;p__Firmicutes;c__Bacilli;o__Turicibacterales;f__Turicibacteraceae;g__Turicibacter', 'k__Bacteria;p__Proteobacteria;c__Betaproteobacteria;o__Burkholderiales;f__Alcaligenaceae;g__Sutterella', 'k__Bacteria;p__Firmicutes;c__Clostridia;o__Clostridiales;f__Ruminococcaceae;g__Ruminococcus']
#ancom_taxa_gut_F3-6 = ['k__Bacteria;p__Firmicutes;c__Erysipelotrichi;o__Erysipelotrichales;f__Erysipelotrichaceae;g__Allobaculum', 'k__Bacteria;p__Bacteroidetes;c__Bacteroidia;o__Bacteroidales;f__[Odoribacteraceae];g__Odoribacter', 'k__Bacteria;p__Proteobacteria;c__Betaproteobacteria;o__Burkholderiales;f__Alcaligenaceae;g__Sutterella', 'k__Bacteria;p__Firmicutes;c__Clostridia;o__Clostridiales;f__Ruminococcaceae;g__Ruminococcus', 'k__Bacteria;p__Bacteroidetes;c__Bacteroidia;o__Bacteroidales;f__Bacteroidaceae;g__Bacteroides']
#ancom_taxa_gut_F2-6 = ['k__Bacteria;p__Verrucomicrobia;c__Verrucomicrobiae;o__Verrucomicrobiales;f__Verrucomicrobiaceae;g__Akkermansia', 'k__Bacteria;p__Bacteroidetes;c__Bacteroidia;o__Bacteroidales;f__[Odoribacteraceae];g__Odoribacter', 'k__Bacteria;p__Firmicutes;c__Clostridia;o__Clostridiales;f__Ruminococcaceae;g__Ruminococcus', 'k__Bacteria;p__Proteobacteria;c__Betaproteobacteria;o__Burkholderiales;f__Alcaligenaceae;g__Sutterella', 'k__Bacteria;p__Firmicutes;c__Erysipelotrichi;o__Erysipelotrichales;f__Erysipelotrichaceae;g__Allobaculum']
ancom_taxa_gut = ['k__Bacteria;p__Verrucomicrobia;c__Verrucomicrobiae;o__Verrucomicrobiales;f__Verrucomicrobiaceae;g__Akkermansia', 'k__Bacteria;p__Firmicutes;c__Erysipelotrichi;o__Erysipelotrichales;f__Erysipelotrichaceae;g__Allobaculum']
taxa_C57B_gut = ancom_C57B_gut[ancom_C57B_gut.index.isin(ancom_taxa_gut)]
taxa_BALB_gut = ancom_BALB_gut[ancom_BALB_gut.index.isin(ancom_taxa_gut)]

ancom_plot = pd.DataFrame({'C57BL6/J': taxa_C57B_gut['average'],
                           'BALB/c': taxa_BALB_gut['average']},
                         index=taxa_C57B_gut.index)
ancom_taxa_plot = ancom_plot.plot.barh(figsize=(20,15), width =0.8, fontsize=20).legend(loc=1, prop={'size':30})
plt.legend(labels=["C57BL/6J","BALB/c"], fontsize=30)
ancom_taxa_plot.figure.savefig(work_dir + '/ancom_plot_gut6_F1.png', set_facecolor='white', bbox_inches='tight', facecolor='white')

# %% [code]
%time !$q_path; qiime feature-table filter-samples \
  --i-table $work_dir/gut6000-mouse-filtered-table-nocomp.qza \
  --m-metadata-file $meta_file \
  --p-where "[host_common_name] IN ('mouse') AND [genotype] NOT IN ('B6CF1')" \
  --o-filtered-table $work_dir/mouse-table-gut6000-nocompRM.qza

# %% [code]
%time !$q_path; qiime taxa collapse \
  --i-table $work_dir/mouse-table-gut6000-nocompRM.qza \
  --i-taxonomy $work_dir/taxonomy_full.qza \
  --p-level 6 \
  --o-collapsed-table $work_dir/collapsed-level6-mouse-gut-table-nocompRM.qza

# %% [code]
%time !$q_path; qiime composition add-pseudocount \
  --i-table $work_dir/collapsed-level6-mouse-gut-table-nocompRM.qza \
  --o-composition-table $work_dir/composition-table-mouse-gut-l6-nocompRM.qza

# %% [code]
%time !$q_path; qiime composition ancom \
  --i-table $work_dir/composition-table-mouse-gut-l6-nocompRM.qza \
  --m-metadata-file meta_gen_num.txt \
  --m-metadata-column genotype \
  --o-visualization $work_dir/ancom-genotype-mouse-gut-l6-nocompRM.qzv

# %% [code]
!unzip $work_dir/composition-table-mouse-gut-l6-nocompRM.qza

# %% [code]
! $q_path; biom convert \
  --input-fp d3386047-2e2a-466d-92a1-107610ae2a54/data/feature-table.biom       \
  --to-tsv \
  --output-fp $work_dir/ancom-biom-table-gut-l6 

# %% [code]
!$q_path; biom summarize-table -i d3386047-2e2a-466d-92a1-107610ae2a54/data/feature-table.biom  

# %% [code]
ancom_biom_table_gut_l6 = pd.read_csv(work_dir+"/ancom-biom-table-gut-l6", 
            sep="\t",skiprows=1, index_col=0)

# %% [code]
    k__Bacteria;p__Firmicutes;c__Erysipelotrichi;o__Erysipelotrichales;f__Erysipelotrichaceae;g__Allobaculum
    k__Bacteria;p__Bacteroidetes;c__Bacteroidia;o__Bacteroidales;f__Bacteroidaceae;g__Bacteroides
k__Bacteria;p__Bacteroidetes;c__Bacteroidia;o__Bacteroidales;f__[Odoribacteraceae];g__Odoribacter
    k__Bacteria;p__Verrucomicrobia;c__Verrucomicrobiae;o__Verrucomicrobiales;f__Verrucomicrobiaceae;g__Akkermansia
    k__Bacteria;p__Proteobacteria;c__Betaproteobacteria;o__Burkholderiales;f__Alcaligenaceae;g__Sutterella
    k__Bacteria;p__Firmicutes;c__Clostridia;o__Clostridiales;f__Ruminococcaceae;g__Ruminococcus
    k__Bacteria;p__Actinobacteria;c__Actinobacteria;o__Bifidobacteriales;f__Bifidobacteriaceae;g__Bifidobacterium
    k__Bacteria;p__Firmicutes;c__Bacilli;o__Turicibacterales;f__Turicibacteraceae;g__Turicibacter
    k__Bacteria;p__Proteobacteria;c__Deltaproteobacteria;o__Desulfovibrionales;f__Desulfovibrionaceae;g__Desulfovibrio

# %% [code]
%matplotlib inline

relative_abundance_gut= ancom_biom_table_gut.div(6081)
list_C57B = meta[meta['genotype'].isin(['C57BL6/J'])].index
list_BALB = meta[meta['genotype'].isin(['BALB/c'])].index

ancom_C57B_gut = relative_abundance_gut[[col for col in relative_abundance_gut.columns if col in list_C57B]]
ancom_C57B_gut["average"]=ancom_C57B_gut.mean(axis=1)

ancom_BALB_gut = relative_abundance_gut[[col for col in relative_abundance_gut.columns if col in list_BALB]]
ancom_BALB_gut["average"] = ancom_BALB_gut.mean(axis=1)


ancom_taxa_gut = ['k__Bacteria;p__Proteobacteria;c__Deltaproteobacteria;o__Desulfovibrionales;f__Desulfovibrionaceae;g__Desulfovibrio;s__C21_c20','k__Bacteria;p__Firmicutes;c__Bacilli;o__Lactobacillales;f__Lactobacillaceae;g__Lactobacillus;s__reuteri','k__Bacteria;p__Actinobacteria;c__Actinobacteria;o__Bifidobacteriales;f__Bifidobacteriaceae;g__Bifidobacterium;s__pseudolongum','k__Bacteria;p__Bacteroidetes;c__Bacteroidia;o__Bacteroidales;f__Porphyromonadaceae;g__Parabacteroides;s__gordonii','k__Bacteria;p__Bacteroidetes;c__Bacteroidia;o__Bacteroidales;f__Bacteroidaceae;g__Bacteroides;s__acidifaciens','k__Bacteria;p__Verrucomicrobia;c__Verrucomicrobiae;o__Verrucomicrobiales;f__Verrucomicrobiaceae;g__Akkermansia;s__muciniphila']
ancom_taxa_gut_l6 = ['k__Bacteria;p__Actinobacteria;c__Actinobacteria;o__Bifidobacteriales;f__Bifidobacteriaceae;g__Bifidobacterium','k__Bacteria;p__Proteobacteria;c__Deltaproteobacteria;o__Desulfovibrionales;f__Desulfovibrionaceae;g__Desulfovibrio', 'k__Bacteria;p__Firmicutes;c__Erysipelotrichi;o__Erysipelotrichales;f__Erysipelotrichaceae;g__Allobaculum', 'k__Bacteria;p__Firmicutes;c__Bacilli;o__Turicibacterales;f__Turicibacteraceae;g__Turicibacter', 'k__Bacteria;p__Verrucomicrobia;c__Verrucomicrobiae;o__Verrucomicrobiales;f__Verrucomicrobiaceae;g__Akkermansia', 'k__Bacteria;p__Proteobacteria;c__Betaproteobacteria;o__Burkholderiales;f__Alcaligenaceae;g__Sutterella', 'k__Bacteria;p__Firmicutes;c__Clostridia;o__Clostridiales;f__Ruminococcaceae;g__Ruminococcus', 'k__Bacteria;p__Bacteroidetes;c__Bacteroidia;o__Bacteroidales;f__Bacteroidaceae;g__Bacteroides', 'k__Bacteria;p__Bacteroidetes;c__Bacteroidia;o__Bacteroidales;f__[Odoribacteraceae];g__Odoribacter']

taxa_C57B_gut = ancom_C57B_gut[ancom_C57B_gut.index.isin(ancom_taxa_gut)]
taxa_BALB_gut = ancom_BALB_gut[ancom_BALB_gut.index.isin(ancom_taxa_gut)]

ancom_plot = pd.DataFrame({'C57BL6/J': taxa_C57B_gut['average'],
                           'BALB/c': taxa_BALB_gut['average']},
                         index=taxa_C57B_gut.index)

# %% [markdown]
prepare lists wit indeces 

# %% [code]
import matplotlib.pyplot as plt
ancom_taxa_plot = ancom_plot.plot.barh(figsize=(20,15), width =0.8, fontsize=20).legend(loc=1, prop={'size':30})
plt.legend(labels=["C57BL/6J","BALB/c"], fontsize=30)
plt.xlim(right=0.07)
ancom_taxa_plot.figure.savefig(work_dir + '/ancom_plot_gut_species.svg', set_facecolor='white', bbox_inches='tight')

# %% [markdown]
#### Only Mice without comparison mice (sebum)

# %% [code]
%time !$q_path; qiime feature-table filter-samples \
  --i-table $work_dir/sebum-mouse-filtered-table-nocomp.qza \
  --m-metadata-file $meta_file \
  --p-where "[host_common_name] IN ('mouse') AND [genotype] NOT IN ('B6CF1')" \
  --o-filtered-table $work_dir/mouse-table-sebum-nocompRM.qza

# %% [code]
%time !$q_path; qiime feature-table filter-samples \
  --i-table $work_dir/mouse-table-sebum-nocompRM.qza \
  --m-metadata-file $meta_file \
  --p-where "[host_common_name] IN ('mouse') AND [genotype] NOT IN ('B6CF1') AND [generation] IN ('F5')" \
  --o-filtered-table $work_dir/mouse-table-sebum-nocompRM_F5.qza

# %% [code]
%time !$q_path; qiime taxa collapse \
  --i-table $work_dir/mouse-table-sebum-nocompRM_F1.qza \
  --i-taxonomy $work_dir/taxonomy_full.qza \
  --p-level 6 \
  --o-collapsed-table $work_dir/collapsed-level6-mouse-sebum-table-nocompRM_F1.qza

%time !$q_path; qiime composition add-pseudocount \
  --i-table $work_dir/collapsed-level6-mouse-sebum-table-nocompRM_F1.qza \
  --o-composition-table $work_dir/composition-table-mouse-sebum-l6-nocompRM_F1.qza

%time !$q_path; qiime composition ancom \
  --i-table $work_dir/composition-table-mouse-sebum-l6-nocompRM_F1.qza \
  --m-metadata-file $meta_file \
  --m-metadata-column genotype \
  --o-visualization $work_dir/ancom-genotype-mouse-sebum-l6-nocompRM_F1.qzv
!unzip $work_dir/composition-table-mouse-sebum-l6-nocompRM_F1.qza

# %% [code]
! $q_path; biom convert \
  --input-fp 8322a6d9-60c4-4a35-9f3a-d880bfed4211/data/feature-table.biom       \
  --to-tsv \
  --output-fp $work_dir/ancom-biom-table-sebum6-F2 

# %% [code]
ancom_biom_table_sebum = pd.read_csv(work_dir+"/ancom-biom-table-sebum6-F2", 
            sep="\t",skiprows=1, index_col=0)
ancom_biom_table_sebum.sum()

# %% [code]
relative_abundance_sebum= ancom_biom_table_sebum.div(1279)
list_C57B = meta[meta['genotype'].isin(['C57BL6/J'])].index
list_BALB = meta[meta['genotype'].isin(['BALB/c'])].index

ancom_C57B_sebum=relative_abundance_sebum[[col for col in relative_abundance_sebum.columns if col in list_C57B]]
ancom_C57B_sebum["average"]=ancom_C57B_sebum.mean(axis=1)

ancom_BALB_sebum = relative_abundance_sebum[[col for col in relative_abundance_sebum.columns if col in list_BALB]]
ancom_BALB_sebum["average"] = ancom_BALB_sebum.mean(axis=1)

%matplotlib inline
#ancom_taxa_sebum_F1 = ['']
#ancom_taxa_sebum_F2=['']
#ancom_taxa_sebum_F3=['']
#ancom_taxa_sebum_F4 = ['k__Bacteria;p__Firmicutes;c__Bacilli;o__Lactobacillales;f__Enterococcaceae;g__Enterococcus;s__casseliflavus']
#ancom_taxa_sebum_F5 =['']
#ancom_taxa_sebum_F5-6=['']
#ancom_taxa_sebum_F4-6 = ['k__Bacteria;p__Firmicutes;c__Bacilli;o__Lactobacillales;f__Enterococcaceae;g__Enterococcus']
#ancom_taxa_sebum_F3-6 = ['']
#ancom_taxa_sebum_F2-6 = ['']
#ancom_taxa_sebum = ['k__Bacteria;p__Verrucomicrobia;c__Verrucomicrobiae;o__Verrucomicrobiales;f__Verrucomicrobiaceae;g__Akkermansia', 'k__Bacteria;p__Firmicutes;c__Erysipelotrichi;o__Erysipelotrichales;f__Erysipelotrichaceae;g__Allobaculum']
taxa_C57B_sebum = ancom_C57B_sebum[ancom_C57B_sebum.index.isin(ancom_taxa_sebum)]
taxa_BALB_sebum = ancom_BALB_sebum[ancom_BALB_sebum.index.isin(ancom_taxa_sebum)]

ancom_plot = pd.DataFrame({'C57BL6/J': taxa_C57B_sebum['average'],
                           'BALB/c': taxa_BALB_sebum['average']},
                         index=taxa_C57B_sebum.index)
ancom_taxa_plot = ancom_plot.plot.barh(figsize=(20,15), width =0.8, fontsize=20).legend(loc=1, prop={'size':30})
plt.legend(labels=["C57BL/6J","BALB/c"], fontsize=30)
ancom_taxa_plot.figure.savefig(work_dir + '/ancom_plot_sebum6_F3.png', set_facecolor='white', bbox_inches='tight', facecolor='white')

# %% [code]
%time !$q_path; qiime taxa collapse \
  --i-table $work_dir/mouse-table-sebum-nocompRM.qza \
  --i-taxonomy $work_dir/taxonomy_full.qza \
  --p-level 7 \
  --o-collapsed-table $work_dir/collapsed-level7-mouse-sebum-table-nocompRM.qza

%time !$q_path; qiime composition add-pseudocount \
  --i-table $work_dir/collapsed-level7-mouse-sebum-table-nocompRM.qza \
  --o-composition-table $work_dir/composition-table-mouse-sebum-l7-nocompRM.qza

%time !$q_path; qiime composition ancom \
  --i-table $work_dir/composition-table-mouse-sebum-l7-nocompRM.qza \
  --m-metadata-file $meta_file \
  --m-metadata-column genotype \
  --o-visualization $work_dir/ancom-genotype-mouse-sebum-l7-nocompRM.qzv

# %% [code]
!unzip $work_dir/composition-table-mouse-sebum-l7-nocompRM.qza

# %% [code]
!$q_path; biom summarize-table -i 6c422394-92ca-43ef-846f-39f870386ff8/data/feature-table.biom  

# %% [code]
! $q_path; biom convert \
  --input-fp fcf55a1b-6e91-4838-925f-9f8982d868b5/data/feature-table.biom      \
  --to-tsv \
  --output-fp $work_dir/ancom-biom-table-sebum 

# %% [code]
ancom_biom_table_sebum = pd.read_csv(work_dir+"/ancom-biom-table-sebum", 
            sep="\t",skiprows=1, index_col=0)

relative_abundance_sebum= ancom_biom_table_sebum.div(1582)

ancom_C57B_sebum = relative_abundance_sebum[[col for col in relative_abundance_sebum.columns if col in list_C57B]]
ancom_C57B_sebum["average"]=ancom_C57B_sebum.mean(axis=1)
ancom_BALB_sebum = relative_abundance_sebum[[col for col in relative_abundance_sebum.columns if col in list_BALB]]
ancom_BALB_sebum["average"] = ancom_BALB_sebum.mean(axis=1)

# %% [code]
k__Bacteria;p__Verrucomicrobia;c__Verrucomicrobiae;o__Verrucomicrobiales;f__Verrucomicrobiaceae;g__Akkermansia;s__muciniphila 	589
k__Bacteria;p__Firmicutes;c__Clostridia;o__Clostridiales;f__Ruminococcaceae;__;__ 	587
k__Bacteria;p__Firmicutes;c__Clostridia;o__Clostridiales;f__Ruminococcaceae;g__Ruminococcus;s__ 	585
k__Bacteria;p__Firmicutes;c__Bacilli;o__Turicibacterales;f__Turicibacteraceae;g__Turicibacter;s__ 	584
k__Bacteria;p__Firmicutes;__;__;__;__;__ 	583
k__Bacteria;p__Bacteroidetes;c__Bacteroidia;o__Bacteroidales;f__Bacteroidaceae;g__Bacteroides;s__acidifaciens 	582
k__Bacteria;p__Bacteroidetes;c__Bacteroidia;o__Bacteroidales;f__;g__;s__ 	582
k__Bacteria;p__Proteobacteria;c__Alphaproteobacteria;o__RF32;f__;g__;s__ 	579
k__Bacteria;p__Bacteroidetes;c__Bacteroidia;o__Bacteroidales;f__Porphyromonadaceae;g__Parabacteroides;s__ 	575
k__Bacteria;p__Actinobacteria;c__Actinobacteria;o__Bifidobacteriales;f__Bifidobacteriaceae;g__Bifidobacterium;s__pseudolongum

# %% [code]
ancom_taxa_sebum = ['k__Bacteria;p__Actinobacteria;c__Actinobacteria;o__Bifidobacteriales;f__Bifidobacteriaceae;g__Bifidobacterium;s__pseudolongum','k__Bacteria;p__Bacteroidetes;c__Bacteroidia;o__Bacteroidales;f__Bacteroidaceae;g__Bacteroides;s__acidifaciens','k__Bacteria;p__Verrucomicrobia;c__Verrucomicrobiae;o__Verrucomicrobiales;f__Verrucomicrobiaceae;g__Akkermansia;s__muciniphila']
taxa_C57B_sebum = ancom_C57B_sebum[ancom_C57B_sebum.index.isin(ancom_taxa_sebum)]
taxa_BALB_sebum = ancom_BALB_sebum[ancom_BALB_sebum.index.isin(ancom_taxa_sebum)]


ancom_plot = pd.DataFrame({'C57BL6/J': taxa_C57B_sebum['average'],
                           'BALB/c': taxa_BALB_sebum['average']},
                         index=taxa_C57B_sebum.index)
import matplotlib.pyplot as plt
ancom_taxa_plot = ancom_plot.plot.barh(figsize=(20,15), width =0.8, fontsize=20).legend(loc=1, prop={'size':30})
ancom_taxa_plot.figure.savefig(work_dir + '/ancom_plot-sebum7.png', set_facecolor='white', bbox_inches='tight')

# %% [markdown]
## Upsetplot

# %% [code]
import upsetplot
from upsetplot import UpSet
from upsetplot import from_memberships
from upsetplot import plot
from matplotlib import cm

# %% [code]
def make_UpSet(category, upset_taxa_gut, plotname):
    position_category_column = upset_taxa_gut.columns.get_loc(category)
    up_to_gen = upset_taxa_gut.drop(upset_taxa_gut.iloc[:, (position_category_column+1):335],axis = 1)
    only_gen = up_to_gen.drop(up_to_gen.iloc[:, 277:position_category_column], axis = 1)
    taxa_sum = only_gen.groupby(category).sum()
    taxa_sum = taxa_sum.loc[:,~taxa_sum.columns.str.endswith("__")]
    taxa_sum = taxa_sum.loc[:, (taxa_sum != 0).any(axis=0)]
    taxa_sum = taxa_sum.transpose()
    over_zero = taxa_sum > 0
    over_zero = over_zero.rename(columns=lambda x: x + ' ')
    upset_combined = pd.concat([taxa_sum, over_zero],axis=1)
    upset_combined['taxa1'] = upset_combined.index
    upset_combined['taxa'] = upset_combined['taxa1'].str.split(';', expand=True)[5]
    upset_combined = upset_combined.drop(["taxa1"], axis=1)
    upset_combined = upset_combined.set_index(list(over_zero.columns))
    #upset = UpSet(upset_combined, subset_size='count', intersection_plot_elements=0, show_counts=True)
    plot(upset_combined, show_counts=True, subset_size="count", sort_categories_by="-input")
    #upset.add_stacked_bars(by="taxa")
    #upset.add_stacked_bars(by="taxa", elements=10, colors=cm.gist_stern)
    #upset.plot()
    #plt.legend(bbox_to_anchor=(1.04, 0.5), loc="center left", borderaxespad=0, title=f'Colours for taxa')
    plt.suptitle(f'UpSet plot for Genus divided by {category}')
    #plt.style.use("ggplot")
    plt.savefig(work_dir + f"/{plotname}.svg", transparent=False, bbox_inches='tight')
    plt.show()

# %% [code]
taxa_all_l6[taxa_all_l6['host_body_product'].isin(['UBERON:lumen of gut']) & taxa_all_l6['host_scientific_name'].isin(['Mus musculus']) & ~taxa_all_l6['generation'].isin(['not applicable'])][-60:]

# %% [markdown]
### UpSet Plots für Genotypen

# %% [code]
#prepare taxa_all_l6 table
upset_taxa_gut= taxa_all_l6[taxa_all_l6['host_body_product'].isin(['UBERON:lumen of gut']) & taxa_all_l6['host_scientific_name'].isin(['Mus musculus']) & ~taxa_all_l6['generation'].isin(['not applicable'])]

# %% [code]
make_UpSet("genotype", upset_taxa_gut, "upset_gut_genotype")

# %% [markdown]
#### BALB/c

# %% [code]
upset_taxa_gut_BALB= taxa_all_l6[taxa_all_l6['host_body_product'].isin(['UBERON:lumen of gut']) & taxa_all_l6['genotype'].isin(['BALB/c', 'B6CF1'])& taxa_all_l6['host_scientific_name'].isin(['Mus musculus']) & ~taxa_all_l6['generation'].isin(['not applicable'])]

# %% [code]
make_UpSet('generation', upset_taxa_gut_BALB, "upset_gut_BALB_generation")

# %% [markdown]
##### Cage Lineage

# %% [code]
make_UpSet("cage_lineage", upset_taxa_gut_BALB[upset_taxa_gut_BALB['genotype'].isin(['BALB/c'])], "upset_gut_BALB_cagelineage")

# %% [markdown]
#### B6J

# %% [code]
upset_taxa_gut_B6J= taxa_all_l6[taxa_all_l6['host_body_product'].isin(['UBERON:lumen of gut']) & taxa_all_l6['genotype'].isin(['C57BL6/J', 'B6CF1'])& taxa_all_l6['host_scientific_name'].isin(['Mus musculus']) & ~taxa_all_l6['generation'].isin(['not applicable'])]

# %% [code]
make_UpSet("generation", upset_taxa_gut_B6J, "upset_gut_B6J_generation")

# %% [markdown]
##### Cage Lineage

# %% [code]
make_UpSet("cage_lineage", upset_taxa_gut_B6J[upset_taxa_gut_B6J['genotype'].isin(['C57BL6/J'])], "upset_gut_B6J_cagelineage")

# %% [markdown]
### Sebum

# %% [code]
import matplotlib.pyplot as plt

# %% [code]
#prepare taxa_all_l6 table
upset_taxa_sebum= taxa_all_l6[taxa_all_l6['host_body_product'].isin(['UBERON:sebum']) & taxa_all_l6['host_scientific_name'].isin(['Mus musculus']) & ~taxa_all_l6['generation'].isin(['not applicable'])]

# %% [code]
make_UpSet("genotype", upset_taxa_sebum, "upset_sebum_genotype")

# %% [markdown]
#### BALB/c

# %% [code]
upset_taxa_sebum_BALB= taxa_all_l6[taxa_all_l6['host_body_product'].isin(['UBERON:sebum']) & taxa_all_l6['genotype'].isin(['BALB/c', 'B6CF1'])& taxa_all_l6['host_scientific_name'].isin(['Mus musculus']) & ~taxa_all_l6['generation'].isin(['not applicable'])]

# %% [code]
make_UpSet('generation', upset_taxa_sebum_BALB, "upset_sebum_BALB_generation")

# %% [markdown]
##### Cage Lineage

# %% [code]
make_UpSet("cage_lineage", upset_taxa_sebum_BALB[upset_taxa_sebum_BALB['genotype'].isin(['BALB/c'])], "upset_sebum_BALB_cagelineage")

# %% [markdown]
#### B6J

# %% [code]


# %% [code]
upset_taxa_sebum_B6J= taxa_all_l6[taxa_all_l6['host_body_product'].isin(['UBERON:sebum']) & taxa_all_l6['genotype'].isin(['C57BL6/J'])& taxa_all_l6['host_scientific_name'].isin(['Mus musculus']) & ~taxa_all_l6['generation'].isin(['not applicable'])]

# %% [code]
make_UpSet("generation", upset_taxa_sebum_B6J, "upset_sebum_B6J_generation")

# %% [markdown]
##### Cage Lineage

# %% [code]
make_UpSet("cage_lineage", upset_taxa_sebum_B6J[upset_taxa_sebum_B6J["genotype"].isin(["C57BL6/J"])], "upset_sebum_B6J_cagelineage")

# %% [markdown]
## GGMAP Timeline

# %% [code]
from skbio.tree import TreeNode
%load_ext autoreload
%autoreload 2
%reload_ext autoreload
from ggmap.snippets import *
from ggmap.analyses import *
from ggmap.correlations import *
from ggmap.workflow import *
import seaborn as sns
import matplotlib.pyplot as plt

# %% [code]
meta['i_generation'] = meta['generation'].apply(lambda x: {'RM': '-1: RM',
                                    'P1': '0: P1',
                                    'F1': '1: F1',
                                    'F2': '2: F2',
                                    'F3': '3: F3',
                                    'F4': '4: F4',
                                    'F5': '5: F5',
                                   }.get(x, x))

# %% [markdown]
#### Alpha Diversity
##### Gut

# %% [code]
RM_shannon_average = np.mean(alpha_results_merged_gut[alpha_results_merged_gut['generation'].isin(['RM'])]['shannon'])
RM_shannon_std = np.std(alpha_results_merged_gut[alpha_results_merged_gut['generation'].isin(['RM'])]['shannon'])
RM_PD_average = np.mean(alpha_results_merged_gut[alpha_results_merged_gut['generation'].isin(['RM'])]['PD_whole_tree'])
RM_PD_std = np.std(alpha_results_merged_gut[alpha_results_merged_gut['generation'].isin(['RM'])]['PD_whole_tree'])
RM_observed_average = np.mean(alpha_results_merged_gut[alpha_results_merged_gut['generation'].isin(['RM'])]['observed_features'])
RM_observed_std = np.std(alpha_results_merged_gut[alpha_results_merged_gut['generation'].isin(['RM'])]['observed_features'])

# %% [code]
%matplotlib inline
import pylab
metric = 'shannon'
fig, axes = plt.subplots(1,1, figsize=(10,5))
a_data = plot_timecourse(meta[meta['genotype'].isin(['C57BL6/J','BALB/c', 'B6CF1']) & meta['host_common_name'].isin(['mouse']) & ~meta['generation'].isin(['not applicable'])],
                data=alpha_div_gut['results'][metric],
                col_events='i_generation', col_entities=None,
                cols_groups=['host_common_name', 'genotype'],
                #test_groups=[(('mouse', 'C57BL6/J'), ('mouse', 'BALB/c'))],
                pthreshold=0.05, print_samplesizes=True,
                ax=axes,
                colors_groups={('mouse', 'BALB/c'): "tab:orange", ('mouse', 'C57BL6/J'): "tab:blue"},
                test_groups=None
                )
#axes.legend(labels=['BALB/c', 'C57BL/6J'])
plt.scatter("-1: RM", RM_shannon_average, color='green')
plt.errorbar(["-1: RM"], [RM_shannon_average], yerr=RM_shannon_std, color='green')
#handles, labels = fig.get_legend_handles_labels()
labels = ['BALB/c', 'C57BL/6J', 'B6CF1']
#plt.legend(labels, loc=2)
plt.title('Darm')
plt.savefig(work_dir + "/alpha_gut_timeline_shannon.png", transparent=True, bbox_inches="tight", facecolor='white')

# %% [code]
%matplotlib inline

metric = 'observed_features'
fig, axes = plt.subplots(1,1, figsize=(10,5))
a_data = plot_timecourse(meta[meta['genotype'].isin(['C57BL6/J','BALB/c', 'B6CF1']) & meta['host_common_name'].isin(['mouse']) & ~meta['generation'].isin(['not applicable']) ],
                data=alpha_div_gut['results'][metric],
                col_events='i_generation', col_entities=None,
                cols_groups=['host_common_name', 'genotype'],
                #test_groups=[(('mouse', 'C57BL6/J'), ('mouse', 'BALB/c'))],
                test_groups=None,
                pthreshold=0.05, print_samplesizes=True,
                ax=axes,
                colors_groups={('mouse', 'BALB/c'): "tab:orange", ('mouse', 'C57BL6/J'): "tab:blue"})
#axes.legend(labels=['BALB/c', 'C57BL/6J'])
plt.scatter("-1: RM", RM_observed_average, color='green')
plt.errorbar(["-1: RM"], [RM_observed_average], yerr=RM_observed_std, color='green')
#handles, labels = fig.get_legend_handles_labels()
labels = ['BALB/c', 'C57BL/6J', 'B6CF1']
#plt.legend(handles, labels, loc=2)
plt.title('Darm')
plt.savefig(work_dir + "/alpha_gut_timeline_observed_features.png", transparent=False, bbox_inches="tight", facecolor='white')

# %% [code]
%matplotlib inline

metric = 'PD_whole_tree'
fig, axes = plt.subplots(1,1, figsize=(10,5))
a_data = plot_timecourse(meta[meta['genotype'].isin(['C57BL6/J','BALB/c', 'B6CF1']) & meta['host_common_name'].isin(['mouse']) & ~meta['generation'].isin(['not applicable'])],
                data=alpha_div_gut['results'][metric],
                col_events='i_generation', col_entities=None,
                cols_groups=['host_common_name', 'genotype'],
                #test_groups=[(('mouse', 'C57BL6/J'), ('mouse', 'BALB/c'))],
                test_groups=None,
                pthreshold=0.05, print_samplesizes=True,
                ax=axes,
                colors_groups={('mouse', 'BALB/c'): "tab:orange", ('mouse', 'C57BL6/J'): "tab:blue"})
#axes.legend(labels=['BALB/c', 'C57BL/6J'])
plt.scatter("-1: RM", RM_PD_average, color='green')
plt.errorbar(["-1: RM"], [RM_PD_average], yerr=RM_PD_std, color='green')
#handles, labels = fig.get_legend_handles_labels()
labels = ['BALB/c', 'C57BL/6J', 'B6CF1']
#plt.legend(handles, labels, loc=2)
plt.savefig(work_dir + "/alpha_gut_timeline_PD_whole_tree.png", transparent=False, bbox_inches="tight", facecolor='white')

# %% [markdown]
##### Sebum

# %% [code]
RM_shannon_average = np.mean(alpha_results_merged_sebum[alpha_results_merged_sebum['generation'].isin(['RM'])]['shannon'])
RM_shannon_std = np.std(alpha_results_merged_sebum[alpha_results_merged_sebum['generation'].isin(['RM'])]['shannon'])
RM_PD_average = np.mean(alpha_results_merged_sebum[alpha_results_merged_sebum['generation'].isin(['RM'])]['PD_whole_tree'])
RM_PD_std = np.std(alpha_results_merged_sebum[alpha_results_merged_sebum['generation'].isin(['RM'])]['PD_whole_tree'])
RM_observed_average = np.mean(alpha_results_merged_sebum[alpha_results_merged_sebum['generation'].isin(['RM'])]['observed_features'])
RM_observed_std = np.std(alpha_results_merged_sebum[alpha_results_merged_sebum['generation'].isin(['RM'])]['observed_features'])

# %% [code]
%matplotlib inline

metric = 'shannon'
fig, axes = plt.subplots(1,1, figsize=(10,5))
a_data = plot_timecourse(meta[meta['genotype'].isin(['C57BL6/J','BALB/c', 'B6CF1']) & meta['host_common_name'].isin(['mouse']) & ~meta['generation'].isin(['not applicable'])],
                data=alpha_div_sebum['results'][metric],
                col_events='i_generation', col_entities=None,
                cols_groups=['host_common_name', 'genotype'],
                #test_groups=[(('mouse', 'C57BL6/J'), ('mouse', 'BALB/c'))],
                test_groups=None,
                pthreshold=0.05, print_samplesizes=True,
                ax=axes,
                colors_groups={('mouse', 'BALB/c'): "tab:orange", ('mouse', 'C57BL6/J'): "tab:blue"})
plt.scatter("-1: RM", RM_shannon_average, color='green')
plt.errorbar(["-1: RM"], [RM_shannon_average], yerr=RM_shannon_std, color='green')
#handles, labels = fig.get_legend_handles_labels()
labels = ['BALB/c', 'C57BL/6J', 'B6CF1']
#plt.legend(handles, labels, loc=2)
#axes.legend(labels=['BALB/c', 'C57BL/6J', 'B6CF1'])
plt.savefig(work_dir + "/alpha_sebum_timeline_shannon.png", transparent=False, bbox_inches="tight", facecolor='white')

# %% [code]
%matplotlib inline

metric = 'observed_features'
fig, axes = plt.subplots(1,1, figsize=(10,5))
a_data = plot_timecourse(meta[meta['genotype'].isin(['C57BL6/J','BALB/c', 'B6CF1']) & meta['host_common_name'].isin(['mouse']) & ~meta['generation'].isin(['not applicable'])],
                data=alpha_div_sebum['results'][metric],
                col_events='i_generation', col_entities=None,
                cols_groups=['host_common_name', 'genotype'],
                #test_groups=[(('mouse', 'C57BL6/J'), ('mouse', 'BALB/c'))],
                test_groups=None,
                pthreshold=0.05, print_samplesizes=True,
                ax=axes,
                colors_groups={('mouse', 'BALB/c'): "tab:orange", ('mouse', 'C57BL6/J'): "tab:blue"})
plt.scatter("-1: RM", RM_observed_average, color='green')
plt.errorbar(["-1: RM"], [RM_observed_average], yerr=RM_observed_std, color='green')
#handles, labels = fig.get_legend_handles_labels()
labels = ['BALB/c', 'C57BL/6J', 'B6CF1']
#plt.legend(handles, labels, loc=2)
plt.title('Haut')
plt.savefig(work_dir + "/alpha_sebum_timeline_observed_features.png", transparent=False, bbox_inches="tight", facecolor='white')

# %% [code]
%matplotlib inline

metric = 'PD_whole_tree'
fig, axes = plt.subplots(1,1, figsize=(10,5))
a_data = plot_timecourse(meta[meta['genotype'].isin(['C57BL6/J','BALB/c', 'B6CF1']) & meta['host_common_name'].isin(['mouse']) & ~meta['generation'].isin(['not applicable'])],
                data=alpha_div_sebum['results'][metric],
                col_events='i_generation', col_entities=None,
                cols_groups=['host_common_name', 'genotype'],
                #test_groups=[(('mouse', 'C57BL6/J'), ('mouse', 'BALB/c'))],
                test_groups=None,
                pthreshold=0.05, print_samplesizes=True,
                ax=axes,
                colors_groups={('mouse', 'BALB/c'): "tab:orange", ('mouse', 'C57BL6/J'): "tab:blue"})
plt.scatter("-1: RM", RM_PD_average, color='green')
plt.errorbar(["-1: RM"], [RM_PD_average], yerr=RM_PD_std, color='green')
#handles, labels = fig.get_legend_handles_labels()
labels = ['BALB/c', 'C57BL/6J', 'B6CF1']
#plt.legend(handles, labels, loc=2)
#axes.legend(labels=['BALB/c', 'C57BL/6J'])
plt.savefig(work_dir + "/alpha_sebum_timeline_PD_whole_tree.png", transparent=False, bbox_inches="tight", facecolor='white')

# %% [markdown]
##### Scatter Plots Aplha Diversity
###### Gut

# %% [code]
gut_indeces = meta[meta['genotype'].isin(['C57BL6/J']) & meta['host_body_product'].isin(['UBERON:lumen of gut']) & ~meta['generation'].isin(['not applicable']) & ~meta['tube_id'].isin(['3-a8'])].index
gut_indeces
#gut_indeces = gut_indeces.delete('13422.217')

# %% [code]
alpha_div_gut['results']['PD_whole_tree'].loc[gut_indeces]

# %% [code]
%matplotlib inline
import seaborn as sns
sns.stripplot(x= 'generation', y='PD_whole_tree', data= alpha_results_merged_gut.loc[gut_indeces],order=['P1', 'F1', 'F2', 'F3', 'F4', 'F5'], hue='cage_lineage').set(title='Alpha Diversity per cage lineage by generation B6J')
#lt.show()

# %% [markdown]
it seems as if the cage lineage C1 adds a lot of diversity in F4 to the overall alpha diversity in C75B. This is then missing in F5 as the C1 is merged with C2 to make up for missing samples like we talked about with Laurentiu. The drop in Alphadiversity may thus not originate from a simple lack of number of samples but from the lack of C1 specifically!

# %% [code]
gut_indeces_BALB = meta[meta['genotype'].isin(['BALB/c']) & meta['host_body_product'].isin(['UBERON:lumen of gut']) & ~meta['generation'].isin(['not applicable'])].index
sns.stripplot(x= 'generation', y='PD_whole_tree', data= alpha_results_merged_gut.loc[gut_indeces_BALB],order=['P1', 'F1', 'F2', 'F3', 'F4', 'F5'],  hue='cage_lineage').set(title='Alpha Diversity per cage lineage by generation BALB/c')
plt.show()

# %% [code]
gut_indeces_C57B = meta[meta['genotype'].isin(['C57BL6/J']) & meta['host_body_product'].isin(['UBERON:lumen of gut']) & ~meta['generation'].isin(['not applicable'])].index
data_gut_C57B_alpha = alpha_results_merged_gut.loc[alpha_results_merged_gut.index.intersection(gut_indeces_C57B)]
gut_indeces_BALB = meta[meta['genotype'].isin(['BALB/c']) & meta['host_body_product'].isin(['UBERON:lumen of gut']) & ~meta['generation'].isin(['not applicable'])].index
data_gut_BALB_alpha = alpha_results_merged_gut.loc[alpha_results_merged_gut.index.intersection(gut_indeces_BALB)]
for count, data in enumerate([data_gut_BALB_alpha, data_gut_C57B_alpha]):
    genotype='BALB/c' if count == 0 else 'C57BL/6J'
    for metric in ['PD_whole_tree', 'shannon', 'observed_features']:
        sns.stripplot(x= 'generation', y=metric, data= data,order=['P1', 'F1', 'F2', 'F3', 'F4', 'F5'],  hue='cage_lineage')
        plt.title(f'Alpha Diversity per cage lineage by generation {genotype}')
        plt.legend(bbox_to_anchor=(1.02, 1), loc='upper left', borderaxespad=0)
        plt.show()

# %% [markdown]
###### Sebum

# %% [code]
sebum_indeces_C57B = meta[meta['genotype'].isin(['C57BL6/J']) & meta['host_body_product'].isin(['UBERON:sebum']) & ~meta['generation'].isin(['not applicable']) & ~meta['tube_id'].isin(['3-a8'])].index
alpha_results_merged_sebum.loc[alpha_results_merged_sebum.index.intersection(sebum_indeces_C57B)]

# %% [code]
sebum_indeces_C57B = meta[meta['genotype'].isin(['C57BL6/J']) & meta['host_body_product'].isin(['UBERON:sebum']) & ~meta['generation'].isin(['not applicable'])].index
data_sebum_C57B_alpha = alpha_results_merged_sebum.loc[alpha_results_merged_sebum.index.intersection(sebum_indeces_C57B)]
sebum_indeces_BALB = meta[meta['genotype'].isin(['BALB/c']) & meta['host_body_product'].isin(['UBERON:sebum']) & ~meta['generation'].isin(['not applicable'])].index
data_sebum_BALB_alpha = alpha_results_merged_sebum.loc[alpha_results_merged_sebum.index.intersection(sebum_indeces_BALB)]
for count, data in enumerate([data_sebum_BALB_alpha, data_sebum_C57B_alpha]):
    genotype='BALB/c' if count == 0 else 'C57BL/6J'
    for metric in ['PD_whole_tree', 'shannon', 'observed_features']:
        sns.stripplot(x= 'generation', y=metric, data= data,order=['P1', 'F1', 'F2', 'F3', 'F4', 'F5'],  hue='cage_lineage')
        plt.title(f'Alpha Diversity per cage lineage by generation {genotype}')
        plt.legend(bbox_to_anchor=(1.02, 1), loc='upper left', borderaxespad=0)
        plt.show()

# %% [markdown]
###### Sebumcatplot

# %% [code]
sebum_indeces_C57B = meta[meta['genotype'].isin(['C57BL6/J']) & meta['host_body_product'].isin(['UBERON:sebum']) & ~meta['generation'].isin(['not applicable'])].index
data_sebum_C57B_alpha = alpha_results_merged_sebum.loc[alpha_results_merged_sebum.index.intersection(sebum_indeces_C57B)]
sebum_indeces_BALB = meta[meta['genotype'].isin(['BALB/c']) & meta['host_body_product'].isin(['UBERON:sebum']) & ~meta['generation'].isin(['not applicable'])].index
data_sebum_BALB_alpha = alpha_results_merged_sebum.loc[alpha_results_merged_sebum.index.intersection(sebum_indeces_BALB)]
for count, data in enumerate([data_sebum_BALB_alpha, data_sebum_C57B_alpha]):
    genotype='BALB' if count == 0 else 'C57B'
    order=['B1', 'B2', 'B3'] if genotype == 'BALB' else ['C1', 'C2', 'C3']
    for metric in ['PD_whole_tree', 'shannon', 'observed_features']:
        sns.catplot(data=data, x='cage_lineage', y=metric, hue="sex", col="generation", aspect=.5, order=order)
        #sns.stripplot(x= 'generation', y=metric, data= data,order=['P1', 'F1', 'F2', 'F3', 'F4', 'F5'],  hue='cage_lineage')
        plt.title(f'{genotype} metric {metric}')
        plt.legend(bbox_to_anchor=(1.02, 1), loc='upper left', borderaxespad=0)
        plt.show()

# %% [code]
for count, data in enumerate([data_sebum_BALB_alpha, data_sebum_C57B_alpha]):
    genotype='BALB' if count == 0 else 'C57B'
    order=['B1', 'B2', 'B3'] if genotype == 'BALB' else ['C1', 'C2', 'C3']
    for metric in ['PD_whole_tree', 'shannon', 'observed_features']:
        sns.catplot(data=data, x='cage_lineage', y=metric, hue="", col="generation", aspect=.5, order=order)
        #sns.stripplot(x= 'generation', y=metric, data= data,order=['P1', 'F1', 'F2', 'F3', 'F4', 'F5'],  hue='cage_lineage')
        plt.title(f'{genotype} metric {metric}')
        plt.legend(bbox_to_anchor=(1.02, 1), loc='upper left', borderaxespad=0)
        plt.show()

# %% [markdown]
#### Beta Diversity
##### Gut

# %% [code]
significance_gut = pd.read_csv(work_dir + "/beta_significance_raw_gut.tsv", sep="\t", dtype=str)

# %% [code]
P1_significance_C57B_gut = significance_gut[significance_gut["Group1"].isin(["0: P1"]) & significance_gut["Group2"].isin(["1: F1", "2: F2", "3: F3", "4: F4", "5: F5"]) & significance_gut["SubjectID1"].isin(list_C57B) & significance_gut["SubjectID2"].isin(list_C57B)]

# %% [code]
P1_significance_C57B_gut["genotype"] = "C57BL6/J"

# %% [code]
P1_significance_BALB_gut = significance_gut[significance_gut["Group1"].isin(["0: P1"]) & significance_gut["Group2"].isin(["1: F1", "2: F2", "3: F3", "4: F4", "5: F5"]) & significance_gut["SubjectID1"].isin(list_BALB) & significance_gut["SubjectID2"].isin(list_BALB)]
P1_significance_BALB_gut["genotype"] = "BALB/c"

# %% [code]
P1_significance_gut = P1_significance_C57B_gut.append(P1_significance_BALB_gut)
P1_significance_gut["Distance"] = P1_significance_gut["Distance"].astype(float)

# %% [code]
sns.lineplot(x="Group2", y="Distance", hue="genotype", err_style="band", data=P1_significance_gut)
plt.legend(bbox_to_anchor=(1.05, 1), loc=2)
plt.savefig(work_dir + "/beta_significance_distance_generations_gut.png", transparent=False, bbox_inches='tight')

# %% [code]
distance_significance_gut_BALB= mannwhitneyu(P1_significance_gut[P1_significance_gut['Group2'].isin(["1: F1"]) & P1_significance_gut["genotype"].isin(["BALB/c"])]["Distance"] , P1_significance_gut[P1_significance_gut['Group2'].isin(["5: F5"]) & P1_significance_gut["genotype"].isin(["BALB/c"])]["Distance"])
distance_significance_gut_C57B= mannwhitneyu(P1_significance_gut[P1_significance_gut['Group2'].isin(["1: F1"]) & P1_significance_gut["genotype"].isin(["C57BL6/J"])]["Distance"] , P1_significance_gut[P1_significance_gut['Group2'].isin(["5: F5"]) & P1_significance_gut["genotype"].isin(["C57BL6/J"])]["Distance"])

# %% [code]
P1_significance_gut["Group2"].value_counts()

# %% [code]
distance_significance_gut_BALB

# %% [code]
distance_significance_gut_C57B

# %% [markdown]
##### Sebum

# %% [code]
significance_sebum = pd.read_csv(work_dir + "/beta_significance_raw_sebum.tsv", sep="\t", dtype=str)

# %% [code]
P1_significance_C57B_sebum = significance_sebum[significance_sebum["Group1"].isin(["0: P1"]) & significance_sebum["Group2"].isin(["1: F1", "2: F2", "3: F3", "4: F4", "5: F5"]) & significance_sebum["SubjectID1"].isin(list_C57B) & significance_sebum["SubjectID2"].isin(list_C57B)]

P1_significance_C57B_sebum["genotype"] = "C57BL6/J"

P1_significance_BALB_sebum = significance_sebum[significance_sebum["Group1"].isin(["0: P1"]) & significance_sebum["Group2"].isin(["1: F1", "2: F2", "3: F3", "4: F4", "5: F5"]) & significance_sebum["SubjectID1"].isin(list_BALB) & significance_sebum["SubjectID2"].isin(list_BALB)]
P1_significance_BALB_sebum["genotype"] = "BALB/c"

P1_significance_sebum = P1_significance_C57B_sebum.append(P1_significance_BALB_sebum)
P1_significance_sebum["Distance"] = P1_significance_sebum["Distance"].astype(float)

# %% [code]
sns.lineplot(x="Group2", y="Distance", hue="genotype", err_style="band", data=P1_significance_sebum)
plt.legend(bbox_to_anchor=(1.05, 1), loc=2)
plt.savefig(work_dir + "/beta_significance_distance_generations_sebum.png", transparent=False, bbox_inches='tight')

# %% [code]
distance_significance_sebum_BALB = mannwhitneyu(P1_significance_sebum[P1_significance_sebum['Group2'].isin(["1: F1"]) & P1_significance_sebum["genotype"].isin(["BALB/c"])]["Distance"], P1_significance_sebum[P1_significance_sebum['Group2'].isin(["5: F5"])& P1_significance_sebum["genotype"].isin(["BALB/c"])]["Distance"])
distance_significance_sebum_C57B = mannwhitneyu(P1_significance_sebum[P1_significance_sebum['Group2'].isin(["1: F1"])& P1_significance_sebum["genotype"].isin(["C57BL6/J"])]["Distance"], P1_significance_sebum[P1_significance_sebum['Group2'].isin(["5: F5"])& P1_significance_sebum["genotype"].isin(["C57BL6/J"])]["Distance"])

# %% [code]
distance_significance_sebum_BALB

# %% [code]
distance_significance_sebum_C57B

# %% [markdown]


# %% [code]


# %% [code]


# %% [code]


# %% [code]


# %% [markdown]
# Statistics

# %% [code]
#from skbio.stats.distance import DistanceMatrix
#from skbio.stats.distance import permanova
from qiime2 import Artifact
import itertools
import pandas as pd
import seaborn as sns
import matplotlib.pyplot as plt
pd.set_option('display.max_columns', None)
pd.set_option('display.max_rows', None)

# %% [markdown]
#### Seaborn Clustermap for Determination of HPLC samples

# %% [code]
%time ! $q_path; qiime diversity filter-distance-matrix \
  --i-distance-matrix $work_dir/gut6000_beta_matrix-nocomp.qza \
  --m-metadata-file $work_dir/meta_gen_num.txt \
  --p-where "[genotype] IN ('C57BL6/J') AND [generation] IN ('F4')" \
  --o-filtered-distance-matrix $work_dir/gut6000_C57B_beta_matrix-nocomp.qza

# %% [code]
dm_6000gut = Artifact.load(work_dir + "/gut6000_C57B_beta_matrix-nocomp.qza").view(DistanceMatrix)

# %% [code]
%time ! $q_path; qiime tools export \
  --input-path $work_dir/gut6000_C57B_beta_matrix-nocomp.qza \
  --output-path $work_dir/gut6000_C57B_beta_matrix-nocomp_export

# %% [code]
dm_6000gut_C57B = pd.read_csv(work_dir + "/gut6000_C57B_beta_matrix-nocomp_export/distance-matrix.tsv", sep="\t", dtype=str, index_col=0)

# %% [code]
dm_6000gut_C57B_float = dm_6000gut_C57B.apply(pd.to_numeric)

# %% [code]
meta['cage_group'].value_counts()

# %% [code]
dm_6000gut_C57B

# %% [code]
C57B_F4 = sns.clustermap(dm_6000gut_C57B_float, yticklabels = True, xticklabels= True)
#sns.set_style('white')
for tick in C57B_F4.ax_heatmap.get_yticklabels():
    group = meta.loc[tick.get_text()]["cage_group"]
    if group in ("CF4-1"):
        tick.set_color('green')
    elif group in ("CF4-2"):
        tick.set_color('blue')
    elif group in ("CF4-3"):
        tick.set_color('red')
    else:
        tick.set_color('black')
for tick in C57B_F4.ax_heatmap.get_xticklabels():
    group = meta.loc[tick.get_text()]["cage_group"]
    if group in ("CF4-1"):
        tick.set_color('green')
    elif group in ("CF4-2"):
        tick.set_color('blue')
    elif group in ("CF4-3"):
        tick.set_color('red')
    else:
        tick.set_color('black')
plt.savefig(work_dir + "/C57B_F4.png", transparent=False)

# %% [code]
%time ! $q_path; qiime diversity filter-distance-matrix \
  --i-distance-matrix $work_dir/gut6000_beta_matrix-nocomp.qza \
  --m-metadata-file $work_dir/meta_gen_num.txt \
  --p-where "[genotype] IN ('BALB/c') AND [generation] IN ('F4')" \
  --o-filtered-distance-matrix $work_dir/gut6000_BALB_beta_matrix-nocomp.qza

%time ! $q_path; qiime tools export \
  --input-path $work_dir/gut6000_BALB_beta_matrix-nocomp.qza \
  --output-path $work_dir/gut6000_BALB_beta_matrix-nocomp_export

dm_6000gut_BALB = pd.read_csv(work_dir + "/gut6000_BALB_beta_matrix-nocomp_export/distance-matrix.tsv", sep="\t", dtype=str, index_col=0)

dm_6000gut_BALB_float = dm_6000gut_BALB.apply(pd.to_numeric)

# %% [code]
BALB_F4 = sns.clustermap(dm_6000gut_BALB_float, yticklabels = True, xticklabels= True)
for tick in BALB_F4.ax_heatmap.get_yticklabels():
    group = meta.loc[tick.get_text()]["cage_group"]
    if group in ("BF4-1"):
        tick.set_color('green')
    elif group in ("BF4-2"):
        tick.set_color('blue')
    elif group in ("BF4-3"):
        tick.set_color('red')
    else:
        tick.set_color('black')
for tick in BALB_F4.ax_heatmap.get_xticklabels():
    group = meta.loc[tick.get_text()]["cage_group"]
    if group in ("BF4-1"):
        tick.set_color('green')
    elif group in ("BF4-2"):
        tick.set_color('blue')
    elif group in ("BF4-3"):
        tick.set_color('red')
    else:
        tick.set_color('black')
plt.savefig(work_dir + "/BALB_F4.png", transparent=False)

# %% [code]
meta['comparison'] = np.where(
    meta.index.isin(indeces), 1, 0) 

# %% [code]
meta[meta['host_body_product'].isin(['UBERON:lumen of gut'])]['comparison'].value_counts()

# %% [code]
meta.head()

# %% [code]
def permanova_beta (condition, dm, comp_group):
    counts = condition[comp_group].value_counts()
    idx = condition.index
    results = permanova(dm.filter(idx), meta.loc[idx, comp_group], permutations = 9999)
    return counts, results

# %% [markdown]
## Gut
Wir vergleichen alle Betadiversitätergebnisse von Maussamples aus dem Darm, mit und ohne Vergleichsmäuse. Dabei werden die Genotypen verglichen!
#### Gut with Comparison

# %% [code]
dm_gut_comp = Artifact.load(work_dir + "/gut6000_beta_matrix.qza").view(DistanceMatrix)

# %% [code]
genotype_dm = meta.loc[dm_gut_comp.ids,:][meta['genotype'].isin(['C57BL6/J', 'BALB/c'])]

# %% [code]
permanova_beta(genotype_dm, dm_gut_comp, "genotype")

# %% [markdown]
#### Janvier vs. Düsseldorf

# %% [code]
comparison = meta.loc[dm_gut_comp.ids,:][meta['comparison'].isin([1])]
permanova_beta(comparison, dm_gut_comp, "genotype")

# %% [markdown]
Im Vergleich sind die Unterschiede zwischen den Genotypen aus Dússeldorf und Janvier nicht signifikant.

# %% [markdown]
Vergleich zwischen den Balb und C57B Mäusen aus Janvier und aus Düsseldorf.

# %% [code]
for genotype in ["C57BL6/J", "BALB/c"]:
    comparison = meta.loc[dm_gut_comp.ids,:][meta['comparison'].isin([1]) & meta['genotype'].isin([genotype])]
    result = permanova_beta(comparison, dm_gut_comp, "geo_loc_name")
    print (f"{genotype} \n {result}")

# %% [code]
for city in ["France:Janvier", "Germany:Duesseldorf"]:
    comparison = meta.loc[dm_gut_comp.ids,:][meta['comparison'].isin([1]) & meta['geo_loc_name'].isin([city])]
    result = permanova_beta(comparison, dm_gut_comp, "genotype")
    print (f"{city} \n {result}")

# %% [markdown]
##### BALB/c

# %% [code]
for genotype in ['BALB/c', 'C57BL6/J']:
    comparison = meta.loc[dm_gut_comp.ids,:][(meta['comparison'].isin([0]) & meta['genotype'].isin(['BALB/c'])) | (meta['comparison'].isin([1]) & meta['genotype'].isin([genotype]))]
    result = permanova_beta(comparison, dm_gut_comp, "comparison")
    print (f"BALB/c experiment + {genotype} comparison \n {result}")
    for city in ['France:Janvier', 'Germany:Duesseldorf']:
        comparison = meta.loc[dm_gut_comp.ids,:][(meta['comparison'].isin([0]) & meta['genotype'].isin(['BALB/c'])) | (meta['comparison'].isin([1]) & meta['geo_loc_name'].isin([city]) & meta['genotype'].isin([genotype]))]
        result = permanova_beta(comparison, dm_gut_comp, "comparison")
        print (f"BALB/c experiment + {genotype}-{city} comparison \n {result}")

# %% [code]
for city in ['France:Janvier', 'Germany:Duesseldorf']:
    comparison = meta.loc[dm_gut_comp.ids,:][(meta['comparison'].isin([0]) & meta['genotype'].isin(['BALB/c'])) | (meta['comparison'].isin([1]) & meta['geo_loc_name'].isin([city]))]
    result = permanova_beta(comparison, dm_gut_comp, "comparison")
    print (f"BALB/c experiment + whole {city} comparison \n {result}")

# %% [code]
comparison = meta.loc[dm_gut_comp.ids,:][(meta['comparison'].isin([0]) & meta['genotype'].isin(['BALB/c'])) | meta['comparison'].isin([1])]
result = permanova_beta(comparison, dm_gut_comp, "comparison")
print (f"BALB/c experiment + whole comparison \n {result}")

# %% [code]
comparison = meta.loc[dm_gut_comp.ids,:][meta['comparison'].isin([0])]
result = permanova_beta(comparison, dm_gut_comp, "genotype")
print (f"BALB/c experiment + C57BL6/J \n {result}")

# %% [markdown]
##### C57BL6/J

# %% [code]
for genotype in ['BALB/c', 'C57BL6/J']:
    comparison = meta.loc[dm_gut_comp.ids,:][(meta['comparison'].isin([0]) & meta['genotype'].isin(['C57BL6/J'])) | (meta['comparison'].isin([1]) & meta['genotype'].isin([genotype]))]
    result = permanova_beta(comparison, dm_gut_comp, "comparison")
    print (f"C57BL6/J experiment + {genotype} comparison \n {result}")
    for city in ['France:Janvier', 'Germany:Duesseldorf']:
        comparison = meta.loc[dm_gut_comp.ids,:][(meta['comparison'].isin([0]) & meta['genotype'].isin(['C57BL6/J'])) | (meta['comparison'].isin([1]) & meta['geo_loc_name'].isin([city]) & meta['genotype'].isin([genotype]))]
        result = permanova_beta(comparison, dm_gut_comp, "comparison")
        print (f"C57BL6/J experiment + {genotype}-{city} comparison \n {result}")

# %% [code]
for city in ['France:Janvier', 'Germany:Duesseldorf']:
    comparison = meta.loc[dm_gut_comp.ids,:][(meta['comparison'].isin([0]) & meta['genotype'].isin(['C57BL6/J'])) | (meta['comparison'].isin([1]) & meta['geo_loc_name'].isin([city]))]
    result = permanova_beta(comparison, dm_gut_comp, "comparison")
    print (f"C57BL6/J experiment + whole {city} comparison \n {result}")

# %% [code]
comparison = meta.loc[dm_gut_comp.ids,:][(meta['comparison'].isin([0]) & meta['genotype'].isin(['C57BL6/J'])) | meta['comparison'].isin([1])]
result = permanova_beta(comparison, dm_gut_comp, "comparison")
print (f"C57BL6/J experiment + whole comparison \n {result}")

# %% [code]
comparison = meta.loc[dm_gut_comp.ids,:][meta['comparison'].isin([0])]
result = permanova_beta(comparison, dm_gut_comp, "genotype")
print (f"C57BL6/J experiment + BALB/c \n {result}")

# %% [markdown]
#### Gut without comparison

# %% [code]
dm_gut_no_comp = Artifact.load(work_dir + "/gut6000_beta_matrix-nocomp.qza").view(DistanceMatrix)

# %% [code]
meta['genotype'].value_counts()

# %% [code]
genotype_dm_no_comp = meta.loc[dm_gut_no_comp.ids,:][meta['genotype'].isin(['C57BL6/J', 'BALB/c'])]

# %% [code]
permanova_beta(genotype_dm_no_comp, dm_gut_no_comp, 'genotype')

# %% [markdown]
##### cage lineage

# %% [code]
#all
lineage_combination = list(itertools.combinations(['C1','C2','C3','B1','B2','B3'], 2))
count = 0
for lineage in lineage_combination:
    count += 1
    comparison = meta.loc[dm_gut_no_comp.ids,:][meta['comparison'].isin([0]) & meta['cage_lineage'].isin(lineage)]
    result = permanova_beta(comparison, dm_gut_no_comp, 'cage_lineage')
    print(f"Cage lineage pair {lineage} \n {result} \n {count}")

# %% [code]
#B6J
lineage_combination = list(itertools.combinations(['C1','C2','C3'], 2))
count = 0
for lineage in lineage_combination:
    count += 1
    comparison = meta.loc[dm_gut_no_comp.ids,:][meta['comparison'].isin([0]) & meta['genotype'].isin(['C57BL6/J']) & meta['cage_lineage'].isin(lineage)]
    result = permanova_beta(comparison, dm_gut_no_comp, 'cage_lineage')
    print(f"Cage lineage pair {lineage} \n {result} \n {count}")

# %% [code]
#B6J
lineage_combination = list(itertools.combinations(['B1','B2','B3'], 2))
count = 0
for lineage in lineage_combination:
    count += 1
    comparison = meta.loc[dm_gut_no_comp.ids,:][meta['comparison'].isin([0]) & meta['genotype'].isin(['BALB/c']) & meta['cage_lineage'].isin(lineage)]
    result = permanova_beta(comparison, dm_gut_no_comp, 'cage_lineage')
    print(f"Cage lineage pair {lineage} \n {result} \n {count}")

# %% [code]
genotype_dm_no_comp = meta.loc[dm_gut_no_comp.ids,:][meta['cage_lineage'].isin(['C1', 'C3'])]
permanova_beta(genotype_dm_no_comp, dm_gut_no_comp, 'cage_lineage')

# %% [markdown]
##### Generations

# %% [code]
generations_combination = list(itertools.combinations(['P1', 'F1', 'F2', 'F3', 'F4', 'F5'], 2))

# %% [code]
generations_combination

# %% [code]
test = meta.loc[dm_gut_no_comp.ids,:][meta['comparison'].isin([0]) & meta['genotype'].isin(['BALB/c'])]
test['generation'].value_counts()
test1 = meta.loc[dm_gut_no_comp.ids,:][meta['comparison'].isin([0]) & meta['genotype'].isin(['C57BL6/J'])]
test1['generation'].value_counts()

# %% [code]
count = 0
for generation in generations_combination:
    count += 1
    comparison = meta.loc[dm_gut_no_comp.ids,:][meta['comparison'].isin([0]) & meta['genotype'].isin(['BALB/c']) & meta['generation'].isin(generation)]
    result = permanova_beta(comparison, dm_gut_no_comp, 'generation')
    print(f"BALB/c generation pair {generation} \n {result} \n {count}")

# %% [code]
print(0.05 /  15)

# %% [code]
count = 0
for generation in generations_combination:
    count += 1
    comparison = meta.loc[dm_gut_no_comp.ids,:][meta['comparison'].isin([0]) & meta['genotype'].isin(['C57BL6/J']) & meta['generation'].isin(generation)]
    result = permanova_beta(comparison, dm_gut_no_comp, 'generation')
    print(f"C57BL6/J generation pair {generation} \n {result} \n {count}")

# %% [markdown]
##### Genotype Per Generation

# %% [code]
for generation in ['P1', 'F1', 'F2', 'F3', 'F4', 'F5']:
    comparison = meta.loc[dm_gut_no_comp.ids,:][meta['comparison'].isin([0]) & meta['generation'].isin([generation])]
    result = permanova_beta(comparison, dm_gut_no_comp, 'genotype')
    print(f"{generation} \n {result}")

# %% [code]


# %% [markdown]
##### Cage Groups BALB/c

# %% [code]
cg_1 = list(itertools.combinations(['BF1-1', 'BF1-2'], 2))
cg_2 = list(itertools.combinations(['BF2-1', 'BF2-2'], 2))
cg_3 = list(itertools.combinations(['BF3-1', 'BF3-2', 'BF3-3'], 2))
cg_4 = list(itertools.combinations(['BF4-1', 'BF4-2', 'BF4-3'], 2))
cg_5 = list(itertools.combinations(['BF5-1', 'BF5-2', 'BF5-3'], 2))
cgB_list = [cg_1, cg_2, cg_3, cg_4, cg_5]

# %% [code]
for cg_generation in cgB_list:
    for cg in cg_generation:
        comparison = meta.loc[dm_gut_no_comp.ids,:][meta['cage_group'].isin(cg) & meta['genotype'].isin(['BALB/c'])]
        result = permanova_beta(comparison, dm_gut_no_comp, 'cage_group')
        print(f"{cg} \n {result}")

# %% [markdown]
##### Cage Groups C57BL6/J

# %% [code]
cg_1 = list(itertools.combinations(['CF1-1', 'CF1-2', 'CF1-3'], 2))
cg_2 = list(itertools.combinations(['CF2-1', 'CF2-2', 'CF2-3'], 2))
cg_3 = list(itertools.combinations(['CF3-1', 'CF3-2', 'CF3-3'], 2))
cg_4 = list(itertools.combinations(['CF4-1', 'CF4-2', 'CF4-3'], 2))
cg_5 = list(itertools.combinations(['CF5-1', 'CF5-2'], 2))
cgC_list = [cg_1, cg_2, cg_3, cg_4, cg_5]

# %% [code]
for cg_generation in cgC_list:
    for cg in cg_generation:
        comparison = meta.loc[dm_gut_no_comp.ids,:][meta['cage_group'].isin(cg) & meta['genotype'].isin(['C57BL6/J'])]
        result = permanova_beta(comparison, dm_gut_no_comp, 'cage_group')
        print(f"{cg} \n {result}")

# %% [code]


# %% [markdown]
##### Cage-Columns BALB/c

# %% [code]
cb_1 = list(itertools.combinations(['BF1-1', 'BF2-1', 'BF3-1', 'BF4-1', 'BF5-1', 'PB', 'RM'], 2))
cb_2 = list(itertools.combinations(['BF1-2', 'BF2-2', 'BF3-2', 'BF4-2', 'BF5-2', 'PB', 'RM'], 2))
cb_3 = list(itertools.combinations(['BF1-2', 'BF2-2', 'BF3-3', 'BF4-3', 'BF5-3', 'PB', 'RM'], 2))
ccB_list = [cb_1, cb_2, cb_3]

# %% [code]
for cg_column in ccB_list:
    for column in cg_column:
        print(column)
        comparison = meta.loc[dm_gut_no_comp.ids,:][meta['cage_group'].isin(column) & meta['genotype'].isin(['BALB/c', 'B6CF1'])]
        result = permanova_beta(comparison, dm_gut_no_comp, 'cage_group')
        print(result)

# %% [markdown]
##### Cage-Columns C57BL6/J

# %% [code]
cc_1 = list(itertools.combinations(['CF1-1', 'CF2-1', 'CF3-1', 'CF4-1', 'CF5-1', 'PC', 'RM'], 2))
cc_2 = list(itertools.combinations(['CF1-2', 'CF2-2', 'CF3-2', 'CF4-2', 'CF5-1', 'PC', 'RM'], 2))
cc_3 = list(itertools.combinations(['CF1-3', 'CF2-3', 'CF3-3', 'CF4-3', 'CF5-2', 'PC', 'RM'], 2))
ccC_list = [cc_1, cc_2, cc_3]

# %% [code]
for cg_column in ccC_list:
    for column in cg_column:
        comparison = meta.loc[dm_gut_no_comp.ids,:][meta['cage_group'].isin(column) & meta['genotype'].isin(['C57BL6/J', 'B6CF1'])]
        result = permanova_beta(comparison, dm_gut_no_comp, 'cage_group')
        print(f"{column} \n {result}")

# %% [code]


# %% [markdown]
##### Sex in each generation per genotype

# %% [code]
count = 0
for generation in ['P1', 'F1', 'F2', 'F3', 'F4', 'F5']:
    comparison = meta.loc[dm_gut_no_comp.ids,:][meta['generation'].isin([generation]) & meta['genotype'].isin(['BALB/c'])]
    result = permanova_beta(comparison, dm_gut_no_comp, 'sex')
    count += 1
    print(f"{generation} \n {result}")
print(count)

# %% [code]
count = 0
for generation in ['P1', 'F1', 'F2', 'F3', 'F4', 'F5']:
    comparison = meta.loc[dm_gut_no_comp.ids,:][meta['generation'].isin([generation]) & meta['genotype'].isin(['C57BL6/J'])]
    result = permanova_beta(comparison, dm_gut_no_comp, 'sex')
    count += 1
    print(f"{generation} \n {result}")
print(count)

# %% [markdown]
##### Genotype in each generation per sex

# %% [code]


# %% [code]
count = 0
for generation in ['P1', 'F1', 'F2', 'F3', 'F4', 'F5']:
    comparison = meta.loc[dm_gut_no_comp.ids,:][meta['generation'].isin([generation]) & meta['sex'].isin(['female'])]
    result = permanova_beta(comparison, dm_gut_no_comp, 'genotype')
    count += 1
    print(f"{generation} \n {result}")
print(count)

# %% [code]
count = 0
for generation in ['P1', 'F1', 'F2', 'F3', 'F4', 'F5']:
    comparison = meta.loc[dm_gut_no_comp.ids,:][meta['generation'].isin([generation]) & meta['sex'].isin(['male'])]
    result = permanova_beta(comparison, dm_gut_no_comp, 'genotype')
    count += 1
    print(f"{generation} \n {result}")
print(count)

# %% [code]


# %% [markdown]
## Sebum

# %% [markdown]
#### Sebum with Comparison

# %% [code]
dm_sebum_comp = Artifact.load(work_dir + "/sebum_beta_matrix.qza").view(DistanceMatrix)
gen_comparison_sebum = meta.loc[dm_sebum_comp.ids,:][meta['genotype'].isin(['C57BL6/J', 'BALB/c'])]
permanova_beta(gen_comparison_sebum, dm_sebum_comp, "genotype")

# %% [markdown]
#### Janvier vs. Düsseldorf

# %% [code]
test = meta[meta['comparison'].isin([1]) & meta['host_body_product'].isin(['UBERON:sebum'])]
test['geo_loc_name'].value_counts()

# %% [code]
comparison = meta.loc[dm_sebum_comp.ids,:][meta['comparison'].isin([1])]
permanova_beta(comparison, dm_sebum_comp, "genotype")

# %% [code]
for genotype in ["C57BL6/J", "BALB/c"]:
    comparison = meta.loc[dm_sebum_comp.ids,:][meta['comparison'].isin([1]) & meta['genotype'].isin([genotype])]
    result = permanova_beta(comparison, dm_sebum_comp, "geo_loc_name")
    print (f"{genotype} \n {result}")

# %% [code]
for city in ["France:Janvier", "Germany:Duesseldorf"]:
    comparison = meta.loc[dm_sebum_comp.ids,:][meta['comparison'].isin([1]) & meta['geo_loc_name'].isin([city])]
    result = permanova_beta(comparison, dm_sebum_comp, "genotype")
    print (f"{city} \n {result}")

# %% [code]


# %% [markdown]
##### BALB/c

# %% [code]
for genotype in ['BALB/c', 'C57BL6/J']:
    comparison = meta.loc[dm_sebum_comp.ids,:][(meta['comparison'].isin([0]) & meta['genotype'].isin(['BALB/c'])) | (meta['comparison'].isin([1]) & meta['genotype'].isin([genotype]))]
    result = permanova_beta(comparison, dm_sebum_comp, "comparison")
    print (f"BALB/c experiment + {genotype} comparison \n {result}")
    for city in ['France:Janvier', 'Germany:Duesseldorf']:
        comparison = meta.loc[dm_sebum_comp.ids,:][(meta['comparison'].isin([0]) & meta['genotype'].isin(['BALB/c'])) | (meta['comparison'].isin([1]) & meta['geo_loc_name'].isin([city]) & meta['genotype'].isin([genotype]))]
        result = permanova_beta(comparison, dm_sebum_comp, "comparison")
        print (f"BALB/c experiment + {genotype}-{city} comparison \n {result}")

# %% [code]
for city in ['France:Janvier', 'Germany:Duesseldorf']:
    comparison = meta.loc[dm_sebum_comp.ids,:][(meta['comparison'].isin([0]) & meta['genotype'].isin(['BALB/c'])) | (meta['comparison'].isin([1]) & meta['geo_loc_name'].isin([city]))]
    result = permanova_beta(comparison, dm_sebum_comp, "comparison")
    print (f"BALB/c experiment + whole {city} comparison \n {result}")

# %% [code]
comparison = meta.loc[dm_sebum_comp.ids,:][(meta['comparison'].isin([0]) & meta['genotype'].isin(['BALB/c'])) | meta['comparison'].isin([1])]
result = permanova_beta(comparison, dm_sebum_comp, "comparison")
print (f"BALB/c experiment + whole comparison \n {result}")

# %% [code]
comparison = meta.loc[dm_sebum_comp.ids,:][meta['comparison'].isin([0]) & meta['genotype'].isin(['C57BL6/J', 'BALB/c'])]
result = permanova_beta(comparison, dm_sebum_comp, "genotype")
print (f"BALB/c experiment + C57BL6/J \n {result}")

# %% [markdown]
##### C57BL6/J

# %% [code]
for genotype in ['BALB/c', 'C57BL6/J']:
    comparison = meta.loc[dm_sebum_comp.ids,:][(meta['comparison'].isin([0]) & meta['genotype'].isin(['C57BL6/J'])) | (meta['comparison'].isin([1]) & meta['genotype'].isin([genotype]))]
    result = permanova_beta(comparison, dm_sebum_comp, "comparison")
    print (f"C57BL6/J experiment + {genotype} comparison \n {result}")
    for city in ['France:Janvier', 'Germany:Duesseldorf']:
        comparison = meta.loc[dm_sebum_comp.ids,:][(meta['comparison'].isin([0]) & meta['genotype'].isin(['C57BL6/J'])) | (meta['comparison'].isin([1]) & meta['geo_loc_name'].isin([city]) & meta['genotype'].isin([genotype]))]
        result = permanova_beta(comparison, dm_sebum_comp, "comparison")
        print (f"C57BL6/J experiment + {genotype}-{city} comparison \n {result}")

# %% [code]
for city in ['France:Janvier', 'Germany:Duesseldorf']:
    comparison = meta.loc[dm_sebum_comp.ids,:][(meta['comparison'].isin([0]) & meta['genotype'].isin(['C57BL6/J'])) | (meta['comparison'].isin([1]) & meta['geo_loc_name'].isin([city]))]
    result = permanova_beta(comparison, dm_sebum_comp, "comparison")
    print (f"C57BL6/J experiment + whole {city} comparison \n {result}")

# %% [code]
comparison = meta.loc[dm_sebum_comp.ids,:][(meta['comparison'].isin([0]) & meta['genotype'].isin(['C57BL6/J'])) | meta['comparison'].isin([1])]
result = permanova_beta(comparison, dm_sebum_comp, "comparison")
print (f"C57BL6/J experiment + whole comparison \n {result}")

# %% [code]
comparison = meta.loc[dm_sebum_comp.ids,:][meta['comparison'].isin([0]) & meta['genotype'].isin(['C57BL6/J', 'BALB/c'])]
result = permanova_beta(comparison, dm_sebum_comp, "genotype")
print (f"C57BL6/J experiment + BALB/c experiment \n {result}")

# %% [code]


# %% [code]


# %% [code]


# %% [code]


# %% [code]


# %% [code]


# %% [markdown]
#### Sebum without Comparison

# %% [code]
dm_sebum_no_comp = Artifact.load(work_dir + "/sebum_beta_matrix-nocomp.qza").view(DistanceMatrix)
gen_comparison_sebum = meta.loc[dm_sebum_no_comp.ids,:][meta['genotype'].isin(['C57BL6/J', 'BALB/c'])]
permanova_beta(gen_comparison_sebum, dm_sebum_no_comp, "genotype")

# %% [code]
dm_sebum_no_comp = Artifact.load(work_dir + "/sebum_beta_matrix-nocomp.qza").view(DistanceMatrix)
foster_comparison_sebum = meta.loc[dm_sebum_no_comp.ids,:][meta['generation'].isin(['RM', 'P1'])]
permanova_beta(foster_comparison_sebum, dm_sebum_no_comp, "generation")

# %% [markdown]
##### Generation per Genotype

# %% [code]
count = 0
for generation in generations_combination:
    count += 1
    comparison = meta.loc[dm_sebum_no_comp.ids,:][meta['comparison'].isin([0]) & meta['genotype'].isin(['BALB/c']) & meta['generation'].isin(generation)]
    result = permanova_beta(comparison, dm_sebum_no_comp, 'generation')
    print(f"BALB/c generation pair {generation} \n {result} \n {count}")

# %% [code]
count = 0
for generation in generations_combination:
    count += 1
    comparison = meta.loc[dm_sebum_no_comp.ids,:][meta['comparison'].isin([0]) & meta['genotype'].isin(['C57BL6/J']) & meta['generation'].isin(generation)]
    result = permanova_beta(comparison, dm_sebum_no_comp, 'generation')
    print(f"C57BL6/J generation pair {generation} \n {result} \n {count}")

# %% [markdown]
##### Genotype per Generation

# %% [code]
for generation in ['P1', 'F1', 'F2', 'F3', 'F4', 'F5']:
    comparison = meta.loc[dm_sebum_no_comp.ids,:][meta['comparison'].isin([0]) & meta['generation'].isin([generation])]
    result = permanova_beta(comparison, dm_sebum_no_comp, 'genotype')
    print(f"{generation} \n {result}")

# %% [markdown]
##### Cage Groups BALB/c

# %% [code]
for cg_generation in cgB_list:
    for cg in cg_generation:
        comparison = meta.loc[dm_sebum_no_comp.ids,:][meta['cage_group'].isin(cg) & meta['genotype'].isin(['BALB/c'])]
        result = permanova_beta(comparison, dm_sebum_no_comp, 'cage_group')
        print(f"{cg} \n {result}")

# %% [code]
for cg_generation in cgC_list:
    for cg in cg_generation:
        comparison = meta.loc[dm_sebum_no_comp.ids,:][meta['cage_group'].isin(cg) & meta['genotype'].isin(['C57BL6/J'])]
        result = permanova_beta(comparison, dm_sebum_no_comp, 'cage_group')
        print(f"{cg} \n {result}")

# %% [code]


# %% [code]


# %% [markdown]
##### Cage-Columns BALB/c

# %% [code]
cb_1 = list(itertools.combinations(['BF1-1', 'BF2-1', 'BF3-1', 'BF4-1', 'BF5-1', 'PB', 'RM'], 2))
cb_2 = list(itertools.combinations(['BF1-2', 'BF2-2', 'BF3-2', 'BF4-2', 'BF5-2', 'PB', 'RM'], 2))
cb_3 = list(itertools.combinations(['BF1-2', 'BF2-2', 'BF3-3', 'BF4-3', 'BF5-3', 'PB', 'RM'], 2))
ccB_list = [cb_1, cb_2, cb_3]

# %% [code]
for cg_column in ccB_list:
    for column in cg_column:
        print(column)
        comparison = meta.loc[dm_sebum_no_comp.ids,:][meta['cage_group'].isin(column) & meta['genotype'].isin(['BALB/c', 'B6CF1'])]
        result = permanova_beta(comparison, dm_sebum_no_comp, 'cage_group')
        print(result)

# %% [markdown]
##### Cage-Columns C57BL6/J

# %% [code]
cc_1 = list(itertools.combinations(['CF1-1', 'CF2-1', 'CF3-1', 'CF4-1', 'CF5-1', 'PC', 'RM'], 2))
cc_2 = list(itertools.combinations(['CF1-2', 'CF2-2', 'CF3-2', 'CF4-2', 'CF5-1', 'PC', 'RM'], 2))
cc_3 = list(itertools.combinations(['CF1-3', 'CF2-3', 'CF3-3', 'CF4-3', 'CF5-2', 'PC', 'RM'], 2))
ccC_list = [cc_1, cc_2, cc_3]

# %% [code]
for cg_column in ccC_list:
    for column in cg_column:
        comparison = meta.loc[dm_sebum_no_comp.ids,:][meta['cage_group'].isin(column) & meta['genotype'].isin(['C57BL6/J', 'B6CF1'])]
        result = permanova_beta(comparison, dm_sebum_no_comp, 'cage_group')
        print(f"{column} \n {result}")

# %% [code]


# %% [markdown]
##### Sex in each generation per genotype

# %% [code]
count = 0
for generation in ['P1', 'F1', 'F2', 'F3', 'F4', 'F5']:
    comparison = meta.loc[dm_sebum_no_comp.ids,:][meta['generation'].isin([generation]) & meta['genotype'].isin(['BALB/c'])]
    result = permanova_beta(comparison, dm_sebum_no_comp, 'sex')
    count += 1
    print(f"{generation} \n {result}")
print(count)

# %% [code]
count = 0
for generation in ['P1', 'F1', 'F2', 'F3', 'F4', 'F5']:
    comparison = meta.loc[dm_sebum_no_comp.ids,:][meta['generation'].isin([generation]) & meta['genotype'].isin(['C57BL6/J'])]
    result = permanova_beta(comparison, dm_sebum_no_comp, 'sex')
    count += 1
    print(f"{generation} \n {result}")
print(count)

# %% [code]


# %% [markdown]
##### Genotype in each generation per sex

# %% [code]
count = 0
for generation in ['P1', 'F1', 'F2', 'F3', 'F4', 'F5']:
    comparison = meta.loc[dm_sebum_no_comp.ids,:][meta['generation'].isin([generation]) & meta['sex'].isin(['female'])]
    result = permanova_beta(comparison, dm_sebum_no_comp, 'genotype')
    count += 1
    print(f"{generation} \n {result}")
print(count)

# %% [code]
count = 0
for generation in ['P1', 'F1', 'F2', 'F3', 'F4', 'F5']:
    comparison = meta.loc[dm_sebum_no_comp.ids,:][meta['generation'].isin([generation]) & meta['sex'].isin(['male'])]
    result = permanova_beta(comparison, dm_sebum_no_comp, 'genotype')
    count += 1
    print(f"{generation} \n {result}")
print(count)

# %% [code]


# %% [markdown]
## Alpha Diversity Statistics

# %% [markdown]
#### Gut

# %% [markdown]
##### Differences within the comparisons

# %% [code]
#genotype per category
statistic, p = mannwhitneyu(Janvier_faith_C57B_comp, Janvier_faith_BALB, alternative='less')
print(f'Statistic: {statistic} \n p={p}')
statistic, p = mannwhitneyu(Duess_faith_C57B_comp, Duess_faith_BALB, alternative='less')
print(f'Statistic: {statistic} \n p={p}')
statistic, p = mannwhitneyu(exp_faith_C57B_comp, exp_faith_BALB, alternative='less')
print(f'Statistic: {statistic} \n p={p}')

# %% [code]
alpha_C57B_comp = alpha_results_merged_gut_comp[alpha_results_merged_gut_comp['genotype'].isin(['C57BL6/J', 'B6CF1'])]
alpha_BALB_comp = alpha_results_merged_gut_comp[alpha_results_merged_gut_comp['genotype'].isin(['BALB/c', 'B6CF1'])]

# %% [code]
#all
Janvier_faith_comp = alpha_results_merged_gut_comp[(alpha_results_merged_gut_comp['comparison'].isin(['1'])) & (alpha_results_merged_gut_comp['geo_loc_name'].isin(['France:Janvier']))]['PD_whole_tree']
Duess_faith_comp = alpha_results_merged_gut_comp[alpha_results_merged_gut_comp['comparison'].isin(['1']) & (alpha_results_merged_gut_comp['geo_loc_name'].isin(['Germany:Duesseldorf']))]['PD_whole_tree']
exp_faith_comp = alpha_results_merged_gut_comp[alpha_results_merged_gut_comp['comparison'].isin(['1'])]['PD_whole_tree']

faith_groups_comp = list(itertools.combinations([Janvier_faith_comp, Duess_faith_comp, exp_faith_comp],2))
faith_groups_comp_str = list(itertools.combinations(["Janvier_faith_comp", "Duess_faith_comp", "exp_faith_comp"],2))

alpha = 0.05/len(faith_groups_comp)
count = 0
statistics = []
p_values = []
significant = []
groups = []
bonferroni = []
zerofive = []
for pair in faith_groups_comp:
    mannwhitneyu_table(pair, faith_groups_comp_str, count)
    count +=1

alpha_results_df = pd.DataFrame({'Comparisons': groups,
                                'Statistics': statistics,
                                'p_values': p_values,
                                'Alpha': alpha,
                                'Reject H0?': bonferroni,
                                'No Bonferroni reject H0': zerofive})
alpha_results_df

# %% [code]
Janvier_faith_C57B_comp = alpha_C57B_comp[(alpha_C57B_comp['comparison'].isin(['1'])) & (alpha_C57B_comp['geo_loc_name'].isin(['France:Janvier']))]['PD_whole_tree']
Duess_faith_C57B_comp = alpha_C57B_comp[alpha_C57B_comp['comparison'].isin(['1']) & (alpha_C57B_comp['geo_loc_name'].isin(['Germany:Duesseldorf']))]['PD_whole_tree']
exp_faith_C57B_comp = alpha_C57B_comp[alpha_C57B_comp['comparison'].isin(['1'])]['PD_whole_tree']

faith_C57B_groups_comp = list(itertools.combinations([Janvier_faith_C57B_comp, Duess_faith_C57B_comp, exp_faith_C57B_comp],2))
faith_C57B_groups_comp_str = list(itertools.combinations(["Janvier_faith_C57B", "Duess_faith_C57B", "exp_faith_C57B"],2))

# %% [code]
alpha = 0.05/len(faith_C57B_groups_comp)
count = 0
statistics = []
p_values = []
significant = []
groups = []
bonferroni = []
zerofive = []
for pair in faith_C57B_groups_comp:
    mannwhitneyu_table(pair, faith_C57B_groups_comp_str, count)
    count +=1

# %% [code]
alpha_results_df = pd.DataFrame({'Comparisons': groups,
                                'Statistics': statistics,
                                'p_values': p_values,
                                'Alpha': alpha,
                                'Reject H0?': bonferroni,
                                'No Bonferroni reject H0': zerofive})
alpha_results_df

# %% [code]
#BALB/c

# %% [code]
Janvier_faith_BALB = alpha_BALB_comp[(alpha_BALB_comp['comparison'].isin(['1'])) & (alpha_BALB_comp['geo_loc_name'].isin(['France:Janvier']))]['PD_whole_tree']
Duess_faith_BALB = alpha_BALB_comp[alpha_BALB_comp['comparison'].isin(['1']) & (alpha_BALB_comp['geo_loc_name'].isin(['Germany:Duesseldorf']))]['PD_whole_tree']
exp_faith_BALB = alpha_BALB_comp[alpha_BALB_comp['comparison'].isin(['1'])]['PD_whole_tree']

faith_BALB_groups_comp = list(itertools.combinations([Janvier_faith_BALB, Duess_faith_BALB, exp_faith_BALB],2))
faith_BALB_groups_comp_str = list(itertools.combinations(["Janvier_faith_BALB", "Duess_faith_BALB", "exp_faith_BALB"],2))

# %% [code]
alpha = 0.05/len(faith_BALB_groups_comp)
count = 0
statistics = []
p_values = []
significant = []
groups = []
bonferroni = []
zerofive = []
for pair in faith_BALB_groups_comp:
    mannwhitneyu_table(pair, faith_BALB_groups_comp_str, count)
    count +=1

# %% [code]
alpha_results_df = pd.DataFrame({'Comparisons': groups,
                                'Statistics': statistics,
                                'p_values': p_values,
                                'Alpha': alpha,
                                'Reject H0?': bonferroni,
                                'No Bonferroni reject H0': zerofive})
alpha_results_df

# %% [code]


# %% [code]


# %% [code]


# %% [code]


# %% [code]


# %% [markdown]
##### Differences in Alpha Diversity per Genotype between generations

# %% [code]
alpha_C57B = alpha_results_merged_gut[alpha_results_merged_gut['genotype'].isin(['C57BL6/J', 'B6CF1'])]
alpha_BALB = alpha_results_merged_gut[alpha_results_merged_gut['genotype'].isin(['BALB/c', 'B6CF1'])]

# %% [code]
F1_faith_C57B = alpha_C57B[alpha_C57B['generation'].isin(['F1'])]['PD_whole_tree']
F2_faith_C57B = alpha_C57B[alpha_C57B['generation'].isin(['F2'])]['PD_whole_tree']
F3_faith_C57B = alpha_C57B[alpha_C57B['generation'].isin(['F3'])]['PD_whole_tree']
F4_faith_C57B = alpha_C57B[alpha_C57B['generation'].isin(['F4'])]['PD_whole_tree']
F5_faith_C57B = alpha_C57B[alpha_C57B['generation'].isin(['F5'])]['PD_whole_tree']
P1_faith_C57B = alpha_C57B[alpha_C57B['generation'].isin(['P1'])]['PD_whole_tree']
RM_faith_C57B = alpha_C57B[alpha_C57B['generation'].isin(['RM'])]['PD_whole_tree']

faith_C57B_groups = list(itertools.combinations([RM_faith_C57B, P1_faith_C57B, F1_faith_C57B, F2_faith_C57B, F3_faith_C57B, F4_faith_C57B, F5_faith_C57B],2))
faith_C57B_groups_str = list(itertools.combinations(['RM_faith_C57B','P1_faith_C57B', 'F1_faith_C57B', 'F2_faith_C57B', 'F3_faith_C57B', 'F4_faith_C57B', 'F5_faith_C57B'],2))

# %% [code]
F1_faith_BALB = alpha_BALB[alpha_BALB['generation'].isin(['F1'])]['PD_whole_tree']
F2_faith_BALB = alpha_BALB[alpha_BALB['generation'].isin(['F2'])]['PD_whole_tree']
F3_faith_BALB = alpha_BALB[alpha_BALB['generation'].isin(['F3'])]['PD_whole_tree']
F4_faith_BALB = alpha_BALB[alpha_BALB['generation'].isin(['F4'])]['PD_whole_tree']
F5_faith_BALB = alpha_BALB[alpha_BALB['generation'].isin(['F5'])]['PD_whole_tree']
P1_faith_BALB = alpha_BALB[alpha_BALB['generation'].isin(['P1'])]['PD_whole_tree']
RM_faith_BALB = alpha_BALB[alpha_BALB['generation'].isin(['RM'])]['PD_whole_tree']

faith_BALB_groups = list(itertools.combinations([RM_faith_BALB, P1_faith_BALB, F1_faith_BALB, F2_faith_BALB, F3_faith_BALB, F4_faith_BALB, F5_faith_BALB],2))
faith_BALB_groups_str = list(itertools.combinations(['RM_faith_BALB','P1_faith_BALB', 'F1_faith_BALB', 'F2_faith_BALB', 'F3_faith_BALB', 'F4_faith_BALB', 'F5_faith_BALB'],2))

# %% [code]
from scipy.stats import mannwhitneyu

# %% [code]
def mannwhitneyu_table(pair, pairlist_str, count):
    generation_pair = (pairlist_str[count])
    print(generation_pair)
    statistic, p = mannwhitneyu(pair[0], pair[1], alternative='greater')
    statistics.append(statistic)
    p_values.append(p)
    groups.append(generation_pair)
    print(f'Statistik: {statistic} \n p-Wert: {p} \n')
    bonferroni.append("Y") if p < alpha else bonferroni.append("N")
    zerofive.append("Y") if p< 0.05 else zerofive.append("N")
    
    

# %% [code]
alpha = 0.05/len(faith_C57B_groups)
count = 0
statistics = []
p_values = []
significant = []
groups = []
bonferroni = []
zerofive = []
for pair in faith_C57B_groups:
    mannwhitneyu_table(pair, faith_C57B_groups_str, count)
    count +=1

# %% [code]
alpha = 0.05/len(faith_BALB_groups)
count = 0
for pair in faith_BALB_groups:
    mannwhitneyu_table(pair, faith_BALB_groups_str, count)
    count +=1

# %% [code]
alpha_results_df = pd.DataFrame({'Comparisons': groups,
                                'Statistics': statistics,
                                'p_values': p_values,
                                'Alpha': alpha,
                                'Reject H0?': bonferroni,
                                'No Bonferroni reject H0': zerofive})
alpha_results_df

# %% [markdown]
##### Differences in Alpha Diversity per Generation between genotypes

# %% [code]
generation_list_alpha = [(RM_faith_C57B, RM_faith_BALB),
                        (P1_faith_C57B, P1_faith_BALB),
                        (F1_faith_C57B, F1_faith_BALB),
                        (F2_faith_C57B, F2_faith_BALB),
                        (F3_faith_C57B, F3_faith_BALB),
                        (F4_faith_C57B, F4_faith_BALB),
                        (F5_faith_C57B, F5_faith_BALB)]
generation_list_alpha_str = [('RM_faith_C57B', 'RM_faith_BALB'),
                        ('P1_faith_C57B', 'P1_faith_BALB'),
                        ('F1_faith_C57B', 'F1_faith_BALB'),
                        ('F2_faith_C57B', 'F2_faith_BALB'),
                        ('F3_faith_C57B', 'F3_faith_BALB'),
                        ('F4_faith_C57B', 'F4_faith_BALB'),
                        ('F5_faith_C57B', 'F5_faith_BALB')]
statistics = []
p_values = []
significant = []
groups = []
bonferroni = []
zerofive = []
generations=6
alpha = 0.05/generations
count = 0
for pair in generation_list_alpha:
    mannwhitneyu_table(pair, generation_list_alpha_str, count)
    count += 1

# %% [code]
alpha_results_gen_df = pd.DataFrame({'Comparisons': groups,
                                'Statistics': statistics,
                                'p_values': p_values,
                                'Alpha': alpha,
                                'Reject H0: x<=y?': bonferroni,
                                'No Bonferroni reject H0': zerofive})
alpha_results_gen_df

# %% [markdown]
##### Horizontal cage groups comparisons per genotype

# %% [code]
faith_CF1_1 = alpha_C57B[alpha_C57B['cage_group'].isin(['CF1-1'])]['PD_whole_tree']
faith_CF1_2 = alpha_C57B[alpha_C57B['cage_group'].isin(['CF1-2'])]['PD_whole_tree']
faith_CF1_3 = alpha_C57B[alpha_C57B['cage_group'].isin(['CF1-3'])]['PD_whole_tree']

faith_CF2_1 = alpha_C57B[alpha_C57B['cage_group'].isin(['CF2-1'])]['PD_whole_tree']
faith_CF2_2 = alpha_C57B[alpha_C57B['cage_group'].isin(['CF2-2'])]['PD_whole_tree']
faith_CF2_3 = alpha_C57B[alpha_C57B['cage_group'].isin(['CF2-3'])]['PD_whole_tree']

faith_CF3_1 = alpha_C57B[alpha_C57B['cage_group'].isin(['CF3-1'])]['PD_whole_tree']
faith_CF3_2 = alpha_C57B[alpha_C57B['cage_group'].isin(['CF3-2'])]['PD_whole_tree']
faith_CF3_3 = alpha_C57B[alpha_C57B['cage_group'].isin(['CF3-3'])]['PD_whole_tree']

faith_CF4_1 = alpha_C57B[alpha_C57B['cage_group'].isin(['CF4-1'])]['PD_whole_tree']
faith_CF4_2 = alpha_C57B[alpha_C57B['cage_group'].isin(['CF4-2'])]['PD_whole_tree']
faith_CF4_3 = alpha_C57B[alpha_C57B['cage_group'].isin(['CF4-3'])]['PD_whole_tree']


faith_CF5_1 = alpha_C57B[alpha_C57B['cage_group'].isin(['CF5-1'])]['PD_whole_tree']
faith_CF5_2 = alpha_C57B[alpha_C57B['cage_group'].isin(['CF5-2'])]['PD_whole_tree']

faith_CP1 = alpha_C57B[alpha_C57B['cage_group'].isin(['PC'])]['PD_whole_tree']
faith_CRM = alpha_C57B[alpha_C57B['cage_group'].isin(['RM'])]['PD_whole_tree']

faith_C57B_CF1 = list(itertools.combinations([faith_CF1_1, faith_CF1_2, faith_CF1_3],2))
faith_C57B_CF1_str = list(itertools.combinations(['faith_CF1_1', 'faith_CF1_2', 'faith_CF1_3'],2))

faith_C57B_CF2 = list(itertools.combinations([faith_CF2_1, faith_CF2_2, faith_CF2_3],2))
faith_C57B_CF2_str = list(itertools.combinations(['faith_CF2_1', 'faith_CF2_2', 'faith_CF2_3'],2))

faith_C57B_CF3 = list(itertools.combinations([faith_CF3_1, faith_CF3_2, faith_CF3_3],2))
faith_C57B_CF3_str = list(itertools.combinations(['faith_CF3_1', 'faith_CF3_2', 'faith_CF3_3'],2))

faith_C57B_CF4 = list(itertools.combinations([faith_CF4_1, faith_CF4_2, faith_CF4_3],2))
faith_C57B_CF4_str = list(itertools.combinations(['faith_CF4_1', 'faith_CF4_2', 'faith_CF4_3'],2))

faith_C57B_CF5 = list(itertools.combinations([faith_CF5_1, faith_CF5_2],2))
faith_C57B_CF5_str = list(itertools.combinations(['faith_CF5_1', 'faith_CF5_2'],2))

# %% [code]
faith_BF1_1 = alpha_BALB[alpha_BALB['cage_group'].isin(['BF1-1'])]['PD_whole_tree']
faith_BF1_2 = alpha_BALB[alpha_BALB['cage_group'].isin(['BF1-2'])]['PD_whole_tree']

faith_BF2_1 = alpha_BALB[alpha_BALB['cage_group'].isin(['BF2-1'])]['PD_whole_tree']
faith_BF2_2 = alpha_BALB[alpha_BALB['cage_group'].isin(['BF2-2'])]['PD_whole_tree']

faith_BF3_1 = alpha_BALB[alpha_BALB['cage_group'].isin(['BF3-1'])]['PD_whole_tree']
faith_BF3_2 = alpha_BALB[alpha_BALB['cage_group'].isin(['BF3-2'])]['PD_whole_tree']
faith_BF3_3 = alpha_BALB[alpha_BALB['cage_group'].isin(['BF3-3'])]['PD_whole_tree']

faith_BF4_1 = alpha_BALB[alpha_BALB['cage_group'].isin(['BF4-1'])]['PD_whole_tree']
faith_BF4_2 = alpha_BALB[alpha_BALB['cage_group'].isin(['BF4-2'])]['PD_whole_tree']
faith_BF4_3 = alpha_BALB[alpha_BALB['cage_group'].isin(['BF4-3'])]['PD_whole_tree']


faith_BF5_1 = alpha_BALB[alpha_BALB['cage_group'].isin(['BF5-1'])]['PD_whole_tree']
faith_BF5_2 = alpha_BALB[alpha_BALB['cage_group'].isin(['BF5-2'])]['PD_whole_tree']
faith_BF5_3 = alpha_BALB[alpha_BALB['cage_group'].isin(['BF5-3'])]['PD_whole_tree']

faith_CP1 = alpha_BALB[alpha_BALB['cage_group'].isin(['PC'])]['PD_whole_tree']
faith_CRM = alpha_BALB[alpha_BALB['cage_group'].isin(['RM'])]['PD_whole_tree']

faith_BALB_BF1 = list(itertools.combinations([faith_BF1_1, faith_BF1_2],2))
faith_BALB_BF1_str = list(itertools.combinations(['faith_BF1_1', 'faith_BF1_2'],2))

faith_BALB_BF2 = list(itertools.combinations([faith_BF2_1, faith_BF2_2],2))
faith_BALB_BF2_str = list(itertools.combinations(['faith_BF2_1', 'faith_BF2_2'],2))

faith_BALB_BF3 = list(itertools.combinations([faith_BF3_1, faith_BF3_2, faith_BF3_3],2))
faith_BALB_BF3_str = list(itertools.combinations(['faith_BF3_1', 'faith_BF3_2', 'faith_BF3_3'],2))

faith_BALB_BF4 = list(itertools.combinations([faith_BF4_1, faith_BF4_2, faith_BF4_3],2))
faith_BALB_BF4_str = list(itertools.combinations(['faith_BF4_1', 'faith_BF4_2', 'faith_BF4_3'],2))

faith_BALB_BF5 = list(itertools.combinations([faith_BF5_1, faith_BF5_2, faith_BF5_3],2))
faith_BALB_BF5_str = list(itertools.combinations(['faith_BF5_1', 'faith_BF5_2', 'faith_BF5_3'],2))

# %% [code]
statistics = []
p_values = []
significant = []
groups = []
bonferroni = []
zerofive = []
alpha_list=[]

list_count = 0
pair_count = 0
list_of_cages=[faith_C57B_CF1, faith_C57B_CF2, faith_C57B_CF3, faith_C57B_CF4, faith_C57B_CF5,
              faith_BALB_BF1, faith_BALB_BF2, faith_BALB_BF3, faith_BALB_BF4, faith_BALB_BF5]
list_of_cages_str=[faith_C57B_CF1_str, faith_C57B_CF2_str, faith_C57B_CF3_str, faith_C57B_CF4_str, faith_C57B_CF5_str,
                  faith_BALB_BF1_str, faith_BALB_BF2_str, faith_BALB_BF3_str, faith_BALB_BF4_str, faith_BALB_BF5_str]
for cage_list in list_of_cages:
    alpha = 0.05/len(cage_list)
    for pair in cage_list:
        pair_list_str = list_of_cages_str[list_count]
        mannwhitneyu_table(pair, pair_list_str, pair_count)
        alpha_list.append(alpha)
        pair_count += 1
    list_count +=1
    pair_count = 0

# %% [code]
alpha_results_cage_group_horizontal_C57B_df = pd.DataFrame({'Comparisons': groups,
                                'Statistics': statistics,
                                'p_values': p_values,
                                'Alpha': alpha_list,
                                'Reject H0: x<=y?': bonferroni,
                                'No Bonferroni reject H0': zerofive})
print('Results of horizontal compariosons of cage-groups')
alpha_results_cage_group_horizontal_C57B_df

# %% [markdown]
##### Vertical cage groups comparisons per genotype

# %% [code]
faith_CP1 = alpha_C57B[alpha_C57B['cage_group'].isin(['PC'])]['PD_whole_tree']
faith_CRM = alpha_C57B[alpha_C57B['cage_group'].isin(['RM'])]['PD_whole_tree']

faith_C57B_CF_1 = list(itertools.combinations([faith_CRM, faith_CP1, faith_CF1_1, faith_CF2_1, faith_CF3_1, faith_CF4_1, faith_CF5_1],2))
faith_C57B_CF_1_str = list(itertools.combinations(['faith_CRM', 'faith_CP1', 'faith_CF1_1', 'faith_CF2_1', 'faith_CF3_1', 'faith_CF4_1', 'faith_CF5_1'],2))

faith_C57B_CF_2 = list(itertools.combinations([faith_CRM, faith_CP1, faith_CF1_2, faith_CF2_2, faith_CF3_2, faith_CF4_2, faith_CF5_1],2))
faith_C57B_CF_2_str = list(itertools.combinations(['faith_CRM', 'faith_CP1', 'faith_CF1_2', 'faith_CF2_2', 'faith_CF3_2', 'faith_CF4_2', 'faith_CF5_1'],2))

faith_C57B_CF_3 = list(itertools.combinations([faith_CRM, faith_CP1, faith_CF1_3, faith_CF2_3, faith_CF3_3, faith_CF4_3, faith_CF5_2],2))
faith_C57B_CF_3_str = list(itertools.combinations(['faith_CRM', 'faith_CP1', 'faith_CF1_3', 'faith_CF2_3', 'faith_CF3_3', 'faith_CF4_3', 'faith_CF5_2'],2))

# %% [code]
faith_BP1 = alpha_BALB[alpha_BALB['cage_group'].isin(['PB'])]['PD_whole_tree']
faith_BRM = alpha_BALB[alpha_BALB['cage_group'].isin(['RM'])]['PD_whole_tree']

faith_BALB_BF_1 = list(itertools.combinations([faith_BRM, faith_BP1, faith_BF1_1, faith_BF2_1, faith_BF3_1, faith_BF4_1, faith_BF5_1],2))
faith_BALB_BF_1_str = list(itertools.combinations(['faith_BRM', 'faith_BP1', 'faith_BF1_1', 'faith_BF2_1', 'faith_BF3_1', 'faith_BF4_1', 'faith_BF5_1'],2))

faith_BALB_BF_2 = list(itertools.combinations([faith_BRM, faith_BP1, faith_BF1_2, faith_BF2_2, faith_BF3_2, faith_BF4_2, faith_BF5_2],2))
faith_BALB_BF_2_str = list(itertools.combinations(['faith_BRM', 'faith_BP1', 'faith_BF1_2', 'faith_BF2_2', 'faith_BF3_2', 'faith_BF4_2', 'faith_BF5_2'],2))

faith_BALB_BF_3 = list(itertools.combinations([faith_BRM, faith_BP1, faith_BF1_2, faith_BF2_2, faith_BF3_3, faith_BF4_3, faith_BF5_3],2))
faith_BALB_BF_3_str = list(itertools.combinations(['faith_BRM', 'faith_BP1', 'faith_BF1_2', 'faith_BF2_2', 'faith_BF3_3', 'faith_BF4_3', 'faith_BF5_3'],2))

# %% [code]
statistics = []
p_values = []
significant = []
groups = []
bonferroni = []
zerofive = []
alpha_list=[]

list_count = 0
pair_count = 0
list_of_cages=[faith_C57B_CF_1, faith_C57B_CF_2, faith_C57B_CF_3,
              faith_BALB_BF_1, faith_BALB_BF_2, faith_BALB_BF_3]
list_of_cages_str=[faith_C57B_CF_1_str, faith_C57B_CF_2_str, faith_C57B_CF_3_str,
                  faith_BALB_BF_1_str, faith_BALB_BF_2_str, faith_BALB_BF_3_str]
for cage_list in list_of_cages:
    alpha = 0.05/len(cage_list)
    for pair in cage_list:
        pair_list_str = list_of_cages_str[list_count]
        mannwhitneyu_table(pair, pair_list_str, pair_count)
        alpha_list.append(alpha)
        pair_count += 1
    list_count +=1
    pair_count = 0

# %% [code]
alpha_results_cage_group_vertical_C57B_df = pd.DataFrame({'Comparisons': groups,
                                'Statistics': statistics,
                                'p_values': p_values,
                                'Alpha': alpha_list,
                                'Reject H0: x<=y?': bonferroni,
                                'No Bonferroni reject H0': zerofive})
print('Results of vertical comparisons of cage-groups')
alpha_results_cage_group_vertical_C57B_df

# %% [markdown]
##### cage groups

# %% [code]
faith_B1 = alpha_BALB[alpha_BALB['cage_lineage'].isin(['B1'])]['PD_whole_tree']
faith_B2 = alpha_BALB[alpha_BALB['cage_lineage'].isin(['B2'])]['PD_whole_tree']
faith_B3 = alpha_BALB[alpha_BALB['cage_lineage'].isin(['B3'])]['PD_whole_tree']

faith_BALB_Cage = list(itertools.combinations([faith_B1, faith_B2, faith_B3],2))
faith_BALB_Cage_str = list(itertools.combinations(['faith_B1', 'faith_B2', 'faith_B3'],2))

# %% [code]
alpha = 0.05/len(faith_BALB_Cage)
count = 0
statistics = []
p_values = []
significant = []
groups = []
bonferroni = []
zerofive = []
for pair in faith_BALB_Cage:
    mannwhitneyu_table(pair, faith_BALB_Cage_str, count)
    count +=1

alpha_results_df = pd.DataFrame({'Comparisons': groups,
                                'Statistics': statistics,
                                'p_values': p_values,
                                'Alpha': alpha,
                                'Reject H0?': bonferroni,
                                'No Bonferroni reject H0': zerofive})
alpha_results_df

# %% [code]
faith_C1 = alpha_C57B[alpha_C57B['cage_lineage'].isin(['C1'])]['PD_whole_tree']
faith_C2 = alpha_C57B[alpha_C57B['cage_lineage'].isin(['C2'])]['PD_whole_tree']
faith_C3 = alpha_C57B[alpha_C57B['cage_lineage'].isin(['C3'])]['PD_whole_tree']

faith_C57B_Cage = list(itertools.combinations([faith_C1, faith_C2, faith_C3],2))
faith_C57B_Cage_str = list(itertools.combinations(['faith_C1', 'faith_C2', 'faith_C3'],2))

# %% [code]
alpha = 0.05/len(faith_C57B_Cage)
count = 0
statistics = []
p_values = []
significant = []
groups = []
bonferroni = []
zerofive = []
for pair in faith_C57B_Cage:
    mannwhitneyu_table(pair, faith_C57B_Cage_str, count)
    count +=1

alpha_results_df = pd.DataFrame({'Comparisons': groups,
                                'Statistics': statistics,
                                'p_values': p_values,
                                'Alpha': alpha,
                                'Reject H0?': bonferroni,
                                'No Bonferroni reject H0': zerofive})
alpha_results_df

# %% [code]


# %% [markdown]
#### Sebum

# %% [markdown]
##### Differences within the comparisons

# %% [code]
alpha_C57B_comp_sebum = alpha_results_merged_sebum_comp[alpha_results_merged_sebum_comp['genotype'].isin(['C57BL6/J', 'B6CF1'])]
alpha_BALB_comp_sebum = alpha_results_merged_sebum_comp[alpha_results_merged_sebum_comp['genotype'].isin(['BALB/c', 'B6CF1'])]

# %% [code]
Janvier_faith_C57B_sebum = alpha_C57B_comp_sebum[(alpha_C57B_comp_sebum['comparison'].isin(['1'])) & (alpha_C57B_comp_sebum['geo_loc_name'].isin(['France:Janvier']))]['PD_whole_tree']
Duess_faith_C57B_sebum = alpha_C57B_comp_sebum[alpha_C57B_comp_sebum['comparison'].isin(['1']) & (alpha_C57B_comp_sebum['geo_loc_name'].isin(['Germany:Duesseldorf']))]['PD_whole_tree']
exp_faith_C57B_sebum = alpha_C57B_comp_sebum[alpha_C57B_comp_sebum['comparison'].isin(['1'])]['PD_whole_tree']

faith_C57B_groups_comp_sebum = list(itertools.combinations([Janvier_faith_C57B_comp, Duess_faith_C57B_comp, exp_faith_C57B_comp],2))
faith_C57B_groups_comp_sebum_str = list(itertools.combinations(["Janvier_faith_C57B", "Duess_faith_C57B", "exp_faith_C57B"],2))

# %% [code]
alpha = 0.05/len(faith_C57B_groups_comp_sebum)
count = 0
statistics = []
p_values = []
significant = []
groups = []
bonferroni = []
zerofive = []
for pair in faith_C57B_groups_comp_sebum:
    mannwhitneyu_table(pair, faith_C57B_groups_comp_sebum_str, count)
    count +=1

# %% [code]
alpha_results_df = pd.DataFrame({'Comparisons': groups,
                                'Statistics': statistics,
                                'p_values': p_values,
                                'Alpha': alpha,
                                'Reject H0?': bonferroni,
                                'No Bonferroni reject H0': zerofive})
alpha_results_df

# %% [code]
#BALB/c

# %% [code]
Janvier_faith_BALB_comp = alpha_BALB_comp_sebum[(alpha_BALB_comp_sebum['comparison'].isin(['1'])) & (alpha_BALB_comp_sebum['geo_loc_name'].isin(['France:Janvier']))]['PD_whole_tree']
Duess_faith_BALB_comp = alpha_BALB_comp_sebum[alpha_BALB_comp_sebum['comparison'].isin(['1']) & (alpha_BALB_comp_sebum['geo_loc_name'].isin(['Germany:Duesseldorf']))]['PD_whole_tree']
exp_faith_BALB_comp = alpha_BALB_comp_sebum[alpha_BALB_comp_sebum['comparison'].isin(['1'])]['PD_whole_tree']

faith_BALB_groups_comp_sebum = list(itertools.combinations([Janvier_faith_BALB_comp, Duess_faith_BALB_comp, exp_faith_BALB_comp],2))
faith_BALB_groups_comp_sebum_str = list(itertools.combinations(["Janvier_faith_BALB", "Duess_faith_BALB", "exp_faith_BALB"],2))

# %% [code]
alpha = 0.05/len(faith_BALB_groups_comp_sebum)
count = 0
statistics = []
p_values = []
significant = []
groups = []
bonferroni = []
zerofive = []
for pair in faith_BALB_groups_comp_sebum:
    mannwhitneyu_table(pair, faith_BALB_groups_comp_sebum_str, count)
    count +=1

# %% [code]
alpha_results_df = pd.DataFrame({'Comparisons': groups,
                                'Statistics': statistics,
                                'p_values': p_values,
                                'Alpha': alpha,
                                'Reject H0?': bonferroni,
                                'No Bonferroni reject H0': zerofive})
alpha_results_df

# %% [markdown]
##### Differences in Alpha Diversity per Genotype between generations

# %% [code]
alpha_C57B = alpha_results_merged_sebum[alpha_results_merged_sebum['genotype'].isin(['C57BL6/J', 'B6CF1'])]
alpha_BALB = alpha_results_merged_sebum[alpha_results_merged_sebum['genotype'].isin(['BALB/c', 'B6CF1'])]

# %% [code]
F1_faith_C57B = alpha_C57B[alpha_C57B['generation'].isin(['F1'])]['PD_whole_tree']
F2_faith_C57B = alpha_C57B[alpha_C57B['generation'].isin(['F2'])]['PD_whole_tree']
F3_faith_C57B = alpha_C57B[alpha_C57B['generation'].isin(['F3'])]['PD_whole_tree']
F4_faith_C57B = alpha_C57B[alpha_C57B['generation'].isin(['F4'])]['PD_whole_tree']
F5_faith_C57B = alpha_C57B[alpha_C57B['generation'].isin(['F5'])]['PD_whole_tree']
P1_faith_C57B = alpha_C57B[alpha_C57B['generation'].isin(['P1'])]['PD_whole_tree']
RM_faith_C57B = alpha_C57B[alpha_C57B['generation'].isin(['RM'])]['PD_whole_tree']

faith_C57B_groups = list(itertools.combinations([RM_faith_C57B, P1_faith_C57B, F1_faith_C57B, F2_faith_C57B, F3_faith_C57B, F4_faith_C57B, F5_faith_C57B],2))
faith_C57B_groups_str = list(itertools.combinations(['RM_faith_C57B','P1_faith_C57B', 'F1_faith_C57B', 'F2_faith_C57B', 'F3_faith_C57B', 'F4_faith_C57B', 'F5_faith_C57B'],2))

# %% [code]
F1_faith_BALB = alpha_BALB[alpha_BALB['generation'].isin(['F1'])]['PD_whole_tree']
F2_faith_BALB = alpha_BALB[alpha_BALB['generation'].isin(['F2'])]['PD_whole_tree']
F3_faith_BALB = alpha_BALB[alpha_BALB['generation'].isin(['F3'])]['PD_whole_tree']
F4_faith_BALB = alpha_BALB[alpha_BALB['generation'].isin(['F4'])]['PD_whole_tree']
F5_faith_BALB = alpha_BALB[alpha_BALB['generation'].isin(['F5'])]['PD_whole_tree']
P1_faith_BALB = alpha_BALB[alpha_BALB['generation'].isin(['P1'])]['PD_whole_tree']
RM_faith_BALB = alpha_BALB[alpha_BALB['generation'].isin(['RM'])]['PD_whole_tree']

faith_BALB_groups = list(itertools.combinations([RM_faith_BALB, P1_faith_BALB, F1_faith_BALB, F2_faith_BALB, F3_faith_BALB, F4_faith_BALB, F5_faith_BALB],2))
faith_BALB_groups_str = list(itertools.combinations(['RM_faith_BALB','P1_faith_BALB', 'F1_faith_BALB', 'F2_faith_BALB', 'F3_faith_BALB', 'F4_faith_BALB', 'F5_faith_BALB'],2))

# %% [code]
from scipy.stats import mannwhitneyu

# %% [code]
def mannwhitneyu_table(pair, pairlist_str, count):
    generation_pair = (pairlist_str[count])
    print(generation_pair)
    statistic, p = mannwhitneyu(pair[0], pair[1], alternative='greater')
    statistics.append(statistic)
    p_values.append(p)
    groups.append(generation_pair)
    print(f'Statistik: {statistic} \n p-Wert: {p} \n')
    bonferroni.append("Y") if p < alpha else bonferroni.append("N")
    zerofive.append("Y") if p< 0.05 else zerofive.append("N")
    
    

# %% [code]
alpha = 0.05/len(faith_C57B_groups)
count = 0
statistics = []
p_values = []
significant = []
groups = []
bonferroni = []
zerofive = []
for pair in faith_C57B_groups:
    mannwhitneyu_table(pair, faith_C57B_groups_str, count)
    count +=1

# %% [code]
alpha = 0.05/len(faith_BALB_groups)
count = 0
for pair in faith_BALB_groups:
    mannwhitneyu_table(pair, faith_BALB_groups_str, count)
    count +=1

# %% [code]
alpha_results_df = pd.DataFrame({'Comparisons': groups,
                                'Statistics': statistics,
                                'p_values': p_values,
                                'Alpha': alpha,
                                'Reject H0?': bonferroni,
                                'No Bonferroni reject H0': zerofive})
alpha_results_df

# %% [markdown]
##### Differences in Alpha Diversity per Generation between genotypes

# %% [code]
generation_list_alpha = [(RM_faith_C57B, RM_faith_BALB),
                        (P1_faith_C57B, P1_faith_BALB),
                        (F1_faith_C57B, F1_faith_BALB),
                        (F2_faith_C57B, F2_faith_BALB),
                        (F3_faith_C57B, F3_faith_BALB),
                        (F4_faith_C57B, F4_faith_BALB),
                        (F5_faith_C57B, F5_faith_BALB)]
generation_list_alpha_str = [('RM_faith_C57B', 'RM_faith_BALB'),
                        ('P1_faith_C57B', 'P1_faith_BALB'),
                        ('F1_faith_C57B', 'F1_faith_BALB'),
                        ('F2_faith_C57B', 'F2_faith_BALB'),
                        ('F3_faith_C57B', 'F3_faith_BALB'),
                        ('F4_faith_C57B', 'F4_faith_BALB'),
                        ('F5_faith_C57B', 'F5_faith_BALB')]
statistics = []
p_values = []
significant = []
groups = []
bonferroni = []
zerofive = []
generations=6
alpha = 0.05/generations
count = 0
for pair in generation_list_alpha:
    mannwhitneyu_table(pair, generation_list_alpha_str, count)
    count += 1

# %% [code]
alpha_results_gen_df = pd.DataFrame({'Comparisons': groups,
                                'Statistics': statistics,
                                'p_values': p_values,
                                'Alpha': alpha,
                                'Reject H0: x<=y?': bonferroni,
                                'No Bonferroni reject H0': zerofive})
alpha_results_gen_df

# %% [markdown]
##### Horizontal cage groups comparisons per genotype

# %% [code]
faith_CF1_1 = alpha_C57B[alpha_C57B['cage_group'].isin(['CF1-1'])]['PD_whole_tree']
faith_CF1_2 = alpha_C57B[alpha_C57B['cage_group'].isin(['CF1-2'])]['PD_whole_tree']
faith_CF1_3 = alpha_C57B[alpha_C57B['cage_group'].isin(['CF1-3'])]['PD_whole_tree']

faith_CF2_1 = alpha_C57B[alpha_C57B['cage_group'].isin(['CF2-1'])]['PD_whole_tree']
faith_CF2_2 = alpha_C57B[alpha_C57B['cage_group'].isin(['CF2-2'])]['PD_whole_tree']
faith_CF2_3 = alpha_C57B[alpha_C57B['cage_group'].isin(['CF2-3'])]['PD_whole_tree']

faith_CF3_1 = alpha_C57B[alpha_C57B['cage_group'].isin(['CF3-1'])]['PD_whole_tree']
faith_CF3_2 = alpha_C57B[alpha_C57B['cage_group'].isin(['CF3-2'])]['PD_whole_tree']
faith_CF3_3 = alpha_C57B[alpha_C57B['cage_group'].isin(['CF3-3'])]['PD_whole_tree']

faith_CF4_1 = alpha_C57B[alpha_C57B['cage_group'].isin(['CF4-1'])]['PD_whole_tree']
faith_CF4_2 = alpha_C57B[alpha_C57B['cage_group'].isin(['CF4-2'])]['PD_whole_tree']
faith_CF4_3 = alpha_C57B[alpha_C57B['cage_group'].isin(['CF4-3'])]['PD_whole_tree']


faith_CF5_1 = alpha_C57B[alpha_C57B['cage_group'].isin(['CF5-1'])]['PD_whole_tree']
faith_CF5_2 = alpha_C57B[alpha_C57B['cage_group'].isin(['CF5-2'])]['PD_whole_tree']

faith_CP1 = alpha_C57B[alpha_C57B['cage_group'].isin(['PC'])]['PD_whole_tree']
faith_CRM = alpha_C57B[alpha_C57B['cage_group'].isin(['RM'])]['PD_whole_tree']

faith_C57B_CF1 = list(itertools.combinations([faith_CF1_1, faith_CF1_2, faith_CF1_3],2))
faith_C57B_CF1_str = list(itertools.combinations(['faith_CF1_1', 'faith_CF1_2', 'faith_CF1_3'],2))

faith_C57B_CF2 = list(itertools.combinations([faith_CF2_1, faith_CF2_2, faith_CF2_3],2))
faith_C57B_CF2_str = list(itertools.combinations(['faith_CF2_1', 'faith_CF2_2', 'faith_CF2_3'],2))

faith_C57B_CF3 = list(itertools.combinations([faith_CF3_1, faith_CF3_2, faith_CF3_3],2))
faith_C57B_CF3_str = list(itertools.combinations(['faith_CF3_1', 'faith_CF3_2', 'faith_CF3_3'],2))

faith_C57B_CF4 = list(itertools.combinations([faith_CF4_1, faith_CF4_2, faith_CF4_3],2))
faith_C57B_CF4_str = list(itertools.combinations(['faith_CF4_1', 'faith_CF4_2', 'faith_CF4_3'],2))

faith_C57B_CF5 = list(itertools.combinations([faith_CF5_1, faith_CF5_2],2))
faith_C57B_CF5_str = list(itertools.combinations(['faith_CF5_1', 'faith_CF5_2'],2))

# %% [code]
faith_BF1_1 = alpha_BALB[alpha_BALB['cage_group'].isin(['BF1-1'])]['PD_whole_tree']
faith_BF1_2 = alpha_BALB[alpha_BALB['cage_group'].isin(['BF1-2'])]['PD_whole_tree']

faith_BF2_1 = alpha_BALB[alpha_BALB['cage_group'].isin(['BF2-1'])]['PD_whole_tree']
faith_BF2_2 = alpha_BALB[alpha_BALB['cage_group'].isin(['BF2-2'])]['PD_whole_tree']

faith_BF3_1 = alpha_BALB[alpha_BALB['cage_group'].isin(['BF3-1'])]['PD_whole_tree']
faith_BF3_2 = alpha_BALB[alpha_BALB['cage_group'].isin(['BF3-2'])]['PD_whole_tree']
faith_BF3_3 = alpha_BALB[alpha_BALB['cage_group'].isin(['BF3-3'])]['PD_whole_tree']

faith_BF4_1 = alpha_BALB[alpha_BALB['cage_group'].isin(['BF4-1'])]['PD_whole_tree']
faith_BF4_2 = alpha_BALB[alpha_BALB['cage_group'].isin(['BF4-2'])]['PD_whole_tree']
faith_BF4_3 = alpha_BALB[alpha_BALB['cage_group'].isin(['BF4-3'])]['PD_whole_tree']


faith_BF5_1 = alpha_BALB[alpha_BALB['cage_group'].isin(['BF5-1'])]['PD_whole_tree']
faith_BF5_2 = alpha_BALB[alpha_BALB['cage_group'].isin(['BF5-2'])]['PD_whole_tree']
faith_BF5_3 = alpha_BALB[alpha_BALB['cage_group'].isin(['BF5-3'])]['PD_whole_tree']

faith_CP1 = alpha_BALB[alpha_BALB['cage_group'].isin(['PC'])]['PD_whole_tree']
faith_CRM = alpha_BALB[alpha_BALB['cage_group'].isin(['RM'])]['PD_whole_tree']

faith_BALB_BF1 = list(itertools.combinations([faith_BF1_1, faith_BF1_2],2))
faith_BALB_BF1_str = list(itertools.combinations(['faith_BF1_1', 'faith_BF1_2'],2))

faith_BALB_BF2 = list(itertools.combinations([faith_BF2_1, faith_BF2_2],2))
faith_BALB_BF2_str = list(itertools.combinations(['faith_BF2_1', 'faith_BF2_2'],2))

faith_BALB_BF3 = list(itertools.combinations([faith_BF3_1, faith_BF3_2, faith_BF3_3],2))
faith_BALB_BF3_str = list(itertools.combinations(['faith_BF3_1', 'faith_BF3_2', 'faith_BF3_3'],2))

faith_BALB_BF4 = list(itertools.combinations([faith_BF4_1, faith_BF4_2, faith_BF4_3],2))
faith_BALB_BF4_str = list(itertools.combinations(['faith_BF4_1', 'faith_BF4_2', 'faith_BF4_3'],2))

faith_BALB_BF5 = list(itertools.combinations([faith_BF5_1, faith_BF5_2, faith_BF5_3],2))
faith_BALB_BF5_str = list(itertools.combinations(['faith_BF5_1', 'faith_BF5_2', 'faith_BF5_3'],2))

# %% [code]
statistics = []
p_values = []
significant = []
groups = []
bonferroni = []
zerofive = []
alpha_list=[]

list_count = 0
pair_count = 0
list_of_cages=[faith_C57B_CF1, faith_C57B_CF2, faith_C57B_CF3, faith_C57B_CF4, faith_C57B_CF5,
              faith_BALB_BF1, faith_BALB_BF2, faith_BALB_BF3, faith_BALB_BF4, faith_BALB_BF5]
list_of_cages_str=[faith_C57B_CF1_str, faith_C57B_CF2_str, faith_C57B_CF3_str, faith_C57B_CF4_str, faith_C57B_CF5_str,
                  faith_BALB_BF1_str, faith_BALB_BF2_str, faith_BALB_BF3_str, faith_BALB_BF4_str, faith_BALB_BF5_str]
for cage_list in list_of_cages:
    alpha = 0.05/len(cage_list)
    for pair in cage_list:
        pair_list_str = list_of_cages_str[list_count]
        mannwhitneyu_table(pair, pair_list_str, pair_count)
        alpha_list.append(alpha)
        pair_count += 1
    list_count +=1
    pair_count = 0

# %% [code]
alpha_results_cage_group_horizontal_C57B_df = pd.DataFrame({'Comparisons': groups,
                                'Statistics': statistics,
                                'p_values': p_values,
                                'Alpha': alpha_list,
                                'Reject H0: x<=y?': bonferroni,
                                'No Bonferroni reject H0': zerofive})
print('Results of horizontal compariosons of cage-groups')
alpha_results_cage_group_horizontal_C57B_df

# %% [markdown]
##### Vertical cage groups comparisons per genotype

# %% [code]
faith_CP1 = alpha_C57B[alpha_C57B['cage_group'].isin(['PC'])]['PD_whole_tree']
faith_CRM = alpha_C57B[alpha_C57B['cage_group'].isin(['RM'])]['PD_whole_tree']

faith_C57B_CF_1 = list(itertools.combinations([faith_CRM, faith_CP1, faith_CF1_1, faith_CF2_1, faith_CF3_1, faith_CF4_1, faith_CF5_1],2))
faith_C57B_CF_1_str = list(itertools.combinations(['faith_CRM', 'faith_CP1', 'faith_CF1_1', 'faith_CF2_1', 'faith_CF3_1', 'faith_CF4_1', 'faith_CF5_1'],2))

faith_C57B_CF_2 = list(itertools.combinations([faith_CRM, faith_CP1, faith_CF1_2, faith_CF2_2, faith_CF3_2, faith_CF4_2, faith_CF5_1],2))
faith_C57B_CF_2_str = list(itertools.combinations(['faith_CRM', 'faith_CP1', 'faith_CF1_2', 'faith_CF2_2', 'faith_CF3_2', 'faith_CF4_2', 'faith_CF5_1'],2))

faith_C57B_CF_3 = list(itertools.combinations([faith_CRM, faith_CP1, faith_CF1_3, faith_CF2_3, faith_CF3_3, faith_CF4_3, faith_CF5_2],2))
faith_C57B_CF_3_str = list(itertools.combinations(['faith_CRM', 'faith_CP1', 'faith_CF1_3', 'faith_CF2_3', 'faith_CF3_3', 'faith_CF4_3', 'faith_CF5_2'],2))

# %% [code]
faith_BP1 = alpha_BALB[alpha_BALB['cage_group'].isin(['PB'])]['PD_whole_tree']
faith_BRM = alpha_BALB[alpha_BALB['cage_group'].isin(['RM'])]['PD_whole_tree']

faith_BALB_BF_1 = list(itertools.combinations([faith_BRM, faith_BP1, faith_BF1_1, faith_BF2_1, faith_BF3_1, faith_BF4_1, faith_BF5_1],2))
faith_BALB_BF_1_str = list(itertools.combinations(['faith_BRM', 'faith_BP1', 'faith_BF1_1', 'faith_BF2_1', 'faith_BF3_1', 'faith_BF4_1', 'faith_BF5_1'],2))

faith_BALB_BF_2 = list(itertools.combinations([faith_BRM, faith_BP1, faith_BF1_2, faith_BF2_2, faith_BF3_2, faith_BF4_2, faith_BF5_2],2))
faith_BALB_BF_2_str = list(itertools.combinations(['faith_BRM', 'faith_BP1', 'faith_BF1_2', 'faith_BF2_2', 'faith_BF3_2', 'faith_BF4_2', 'faith_BF5_2'],2))

faith_BALB_BF_3 = list(itertools.combinations([faith_BRM, faith_BP1, faith_BF1_2, faith_BF2_2, faith_BF3_3, faith_BF4_3, faith_BF5_3],2))
faith_BALB_BF_3_str = list(itertools.combinations(['faith_BRM', 'faith_BP1', 'faith_BF1_2', 'faith_BF2_2', 'faith_BF3_3', 'faith_BF4_3', 'faith_BF5_3'],2))

# %% [code]
statistics = []
p_values = []
significant = []
groups = []
bonferroni = []
zerofive = []
alpha_list=[]

list_count = 0
pair_count = 0
list_of_cages=[faith_C57B_CF_1, faith_C57B_CF_2, faith_C57B_CF_3,
              faith_BALB_BF_1, faith_BALB_BF_2, faith_BALB_BF_3]
list_of_cages_str=[faith_C57B_CF_1_str, faith_C57B_CF_2_str, faith_C57B_CF_3_str,
                  faith_BALB_BF_1_str, faith_BALB_BF_2_str, faith_BALB_BF_3_str]
for cage_list in list_of_cages:
    alpha = 0.05/len(cage_list)
    for pair in cage_list:
        pair_list_str = list_of_cages_str[list_count]
        mannwhitneyu_table(pair, pair_list_str, pair_count)
        alpha_list.append(alpha)
        pair_count += 1
    list_count +=1
    pair_count = 0

# %% [code]
alpha_results_cage_group_vertical_C57B_df = pd.DataFrame({'Comparisons': groups,
                                'Statistics': statistics,
                                'p_values': p_values,
                                'Alpha': alpha_list,
                                'Reject H0: x<=y?': bonferroni,
                                'No Bonferroni reject H0': zerofive})
print('Results of vertical comparisons of cage-groups')
alpha_results_cage_group_vertical_C57B_df

# %% [markdown]
##### cage groups

# %% [code]
faith_B1 = alpha_BALB[alpha_BALB['cage_lineage'].isin(['B1'])]['PD_whole_tree']
faith_B2 = alpha_BALB[alpha_BALB['cage_lineage'].isin(['B2'])]['PD_whole_tree']
faith_B3 = alpha_BALB[alpha_BALB['cage_lineage'].isin(['B3'])]['PD_whole_tree']

faith_BALB_Cage = list(itertools.combinations([faith_B1, faith_B2, faith_B3],2))
faith_BALB_Cage_str = list(itertools.combinations(['faith_B1', 'faith_B2', 'faith_B3'],2))

# %% [code]
alpha = 0.05/len(faith_BALB_Cage)
count = 0
statistics = []
p_values = []
significant = []
groups = []
bonferroni = []
zerofive = []
for pair in faith_BALB_Cage:
    mannwhitneyu_table(pair, faith_BALB_Cage_str, count)
    count +=1

alpha_results_df = pd.DataFrame({'Comparisons': groups,
                                'Statistics': statistics,
                                'p_values': p_values,
                                'Alpha': alpha,
                                'Reject H0?': bonferroni,
                                'No Bonferroni reject H0': zerofive})
alpha_results_df

# %% [code]
faith_C1 = alpha_C57B[alpha_C57B['cage_lineage'].isin(['C1'])]['PD_whole_tree']
faith_C2 = alpha_C57B[alpha_C57B['cage_lineage'].isin(['C2'])]['PD_whole_tree']
faith_C3 = alpha_C57B[alpha_C57B['cage_lineage'].isin(['C3'])]['PD_whole_tree']

faith_C57B_Cage = list(itertools.combinations([faith_C1, faith_C2, faith_C3],2))
faith_C57B_Cage_str = list(itertools.combinations(['faith_C1', 'faith_C2', 'faith_C3'],2))

# %% [code]
alpha = 0.05/len(faith_C57B_Cage)
count = 0
statistics = []
p_values = []
significant = []
groups = []
bonferroni = []
zerofive = []
for pair in faith_C57B_Cage:
    mannwhitneyu_table(pair, faith_C57B_Cage_str, count)
    count +=1

alpha_results_df = pd.DataFrame({'Comparisons': groups,
                                'Statistics': statistics,
                                'p_values': p_values,
                                'Alpha': alpha,
                                'Reject H0?': bonferroni,
                                'No Bonferroni reject H0': zerofive})
alpha_results_df

# %% [code]


# %% [markdown]
# Effect Sizes, Forward Step Redundancy analysis
### Gut

# %% [code]
meta_nocomp = meta[meta['comparison'].isin([0])]

# %% [code]
conditions = [
    (meta["cage_lineage"] == "C1"),
    (meta["cage_lineage"] == "C2"),
    (meta["cage_lineage"] == "C3"),
    (meta["cage_lineage"] == "B1"),
    (meta["cage_lineage"] == "B2"),
    (meta["cage_lineage"] == "B3"),
]

values = ["A", "B", "C", "A", "B", "C"]
 
meta['independent_cage_lineage']= np.select(conditions, values, "other")

# %% [code]
%matplotlib inline
#categorials
categorials = ['season', 'independent_cage_lineage', 'sex']

#ordinals
ordinals = {'generation': ['P1', 'F1', 'F2', 'F3', 'F4', 'F5']}

#intervals
intervals = []

#dates
dates = {}

#metrics
metrics_alpha = ['PD_whole_tree', 'shannon', 'observed_features']
metrics_beta = ['bray_curtis', 'unweighted_unifrac', 'weighted_unifrac']

#max_rows = max(len(alpha_div_gut['results'].keys()), len(gut_beta_div['results'].keys()))
#fig, axes = plt.subplots(max_rows, 2, figsize=(5*2, 5*max_rows))
for i, metric in enumerate(metrics_alpha):
    alpha=redundancy_analysis_alpha(meta_nocomp[meta_nocomp['genotype'].isin(['C57BL6/J'])],
                             alpha_div_gut['results'][metric],
                             categorials=categorials,
                             ordinals=ordinals,
                             intervals=intervals,
                             dates=dates,
                             #ax=axes[i][0],
                             title='alpha %s' % metric,
                             dry=False,
                             wait=True,
                             use_grid=True,
                             dirty=True,
                             nocache=True)
    plt.savefig(work_dir + "/gut_redundancies_seperate_%s.png" % metric, transparent=False, bbox_inches='tight')
for i, metric in enumerate(metrics_beta):
    beta=redundancy_analysis_beta(meta_nocomp[meta_nocomp['genotype'].isin(['C57BL6/J'])],
                             gut_beta_div['results'][metric],
                             metric,
                             categorials=categorials,
                             ordinals=ordinals,
                             intervals=intervals,
                             dates=dates,
                             #ax=axes[i][1],
                             title='beta %s' % metric,
                             dry=False,
                             wait=True,
                             use_grid=True,
                             dirty=True,
                             nocache=True)
    plt.savefig(work_dir + "/gut_redundancies_seperate_%s.png" % metric, transparent=False, bbox_inches='tight')

# %% [code]
%matplotlib inline
#categorials
categorials = ['genotype', 'independent_cage_lineage', 'sex', 'season']

#ordinals
ordinals = {'generation': ['P1', 'F1', 'F2', 'F3', 'F4', 'F5']}

#intervals
intervals = []

#dates
dates = {}

#metrics
metrics_alpha = ['PD_whole_tree', 'shannon', 'observed_features']
metrics_beta = ['bray_curtis', 'unweighted_unifrac', 'weighted_unifrac']

#max_rows = max(len(alpha_div_gut['results'].keys()), len(gut_beta_div['results'].keys()))
#fig, axes = plt.subplots(max_rows, 2, figsize=(5*2, 5*max_rows))
for i, metric in enumerate(metrics_alpha):
    alpha=redundancy_analysis_alpha(meta_nocomp,
                             alpha_div_gut['results'][metric],
                             categorials=categorials,
                             ordinals=ordinals,
                             intervals=intervals,
                             dates=dates,
                             #ax=axes[i][0],
                             title='alpha %s' % metric,
                             dry=False,
                             wait=True,
                             use_grid=True,
                             dirty=True,
                             nocache=True)
    plt.savefig(work_dir + "/gut_redundancies_seperate_%s.png" % metric, transparent=False, bbox_inches='tight')
for i, metric in enumerate(metrics_beta):
    beta=redundancy_analysis_beta(meta_nocomp,
                             gut_beta_div['results'][metric],
                             metric,
                             categorials=categorials,
                             ordinals=ordinals,
                             intervals=intervals,
                             dates=dates,
                             #ax=axes[i][1],
                             title='beta %s' % metric,
                             dry=False,
                             wait=True,
                             use_grid=True,
                             dirty=True,
                             nocache=True)
    plt.savefig(work_dir + "/gut_redundancies_seperate_%s.png" % metric, transparent=False, bbox_inches='tight')

# %% [code]
%time ! $q_path; qiime tools export \
  --input-path $work_dir/rarefied6000-filtered-gut-table.qza \
  --output-path $work_dir/rarefied6000-filtered-gut-table_export

# %% [code]
#OLD DO NOT USE

# %% [code]
rarefied6000_gut = biom2pandas(work_dir + "/rarefied6000-filtered-gut-table_export/feature-table.biom").fillna(0)

# %% [code]
gut_beta_div = beta_diversity(rarefied6000_gut, metrics=["unweighted_unifrac", "weighted_unifrac", "bray_curtis"], reference_tree="/homes/arehm/Benga_ThreeGen/FromQiita/128106_insertion_tree.relabelled.tre", dry=False, use_grid=True)

# %% [markdown]
### Sebum

# %% [code]
%time ! $q_path; qiime tools export \
  --input-path $work_dir/rarefied-filtered-sebum-table.qza \
  --output-path $work_dir/rarefied-filtered-sebum-table_export

# %% [code]
rarefied_sebum = biom2pandas(work_dir + "/rarefied-filtered-sebum-table_export/feature-table.biom").fillna(0)

# %% [code]
sebum_beta_div = beta_diversity(rarefied_sebum, metrics=["unweighted_unifrac", "weighted_unifrac", "bray_curtis"], reference_tree="/homes/arehm/Benga_ThreeGen/FromQiita/128106_insertion_tree.relabelled.tre", dry=False, use_grid=True)

# %% [code]
%matplotlib inline
#categorials
categorials = ['genotype', 'independent_cage_lineage', 'sex', 'season']

#ordinals
ordinals = {'generation': ['P1', 'F1', 'F2', 'F3', 'F4', 'F5']}

#intervals
intervals = []

#dates
dates = {}

#metrics
metrics_alpha = ['PD_whole_tree', 'shannon', 'observed_features']
metrics_beta = ['bray_curtis', 'unweighted_unifrac', 'weighted_unifrac']

#max_rows = max(len(alpha_div_sebum['results'].keys()), len(sebum_beta_div['results'].keys()))
#fig, axes = plt.subplots(max_rows, 2, figsize=(5*2, 5*max_rows))
for i, metric in enumerate(metrics_alpha):
    redundancy_analysis_alpha(meta_nocomp,
                             alpha_div_sebum['results'][metric],
                             categorials=categorials,
                             ordinals=ordinals,
                             intervals=intervals,
                             dates=dates,
                             #ax=axes[i][0],
                             title='alpha %s' % metric,
                             dry=False,
                             wait=True,
                             use_grid=True)
    plt.savefig(work_dir + "/sebum_redundancies_seperate_%s.png" % metric, transparent=False, bbox_inches='tight')
for i, metric in enumerate(metrics_beta):
    redundancy_analysis_beta(meta_nocomp,
                             sebum_beta_div['results'][metric],
                             metric,
                             categorials=categorials,
                             ordinals=ordinals,
                             intervals=intervals,
                             dates=dates,
                             #ax=axes[i][1],
                             title='beta %s' % metric,
                             dry=False,
                             wait=True,
                             use_grid=True)
    plt.savefig(work_dir + "/sebum_redundancies_seperate_%s.png" % metric, transparent=False, bbox_inches='tight')

# %% [code]
%matplotlib inline
#categorials
categorials = ['genotype', 'independent_cage_lineage', 'sex', 'season']

#ordinals
ordinals = {'generation': ['P1', 'F1', 'F2', 'F3', 'F4', 'F5']}

#intervals
intervals = []

#dates
dates = {}

#metrics
metrics_alpha = ['PD_whole_tree', 'shannon', 'observed_features']
metrics_beta = ['bray_curtis', 'unweighted_unifrac', 'weighted_unifrac']

#max_rows = max(len(alpha_div_sebum['results'].keys()), len(sebum_beta_div['results'].keys()))
#fig, axes = plt.subplots(max_rows, 2, figsize=(5*2, 5*max_rows))
for i, metric in enumerate(metrics_alpha):
    redundancy_analysis_alpha(meta_nocomp,
                             alpha_div_sebum['results'][metric],
                             categorials=categorials,
                             ordinals=ordinals,
                             intervals=intervals,
                             dates=dates,
                             #ax=axes[i][0],
                             title='alpha %s' % metric,
                             dry=False,
                             wait=True,
                             use_grid=True)
    plt.savefig(work_dir + "/sebum_redundancies_seperate_%s.png" % metric, transparent=False, bbox_inches='tight')

# %% [code]


# %% [code]
%matplotlib inline
#categorials
categorials = ['season', 'independent_cage_lineage', 'sex']

#ordinals
ordinals = {'generation': ['P1', 'F1', 'F2', 'F3', 'F4', 'F5']}

#intervals
intervals = []

#dates
dates = {}

#metrics
metrics_alpha = ['PD_whole_tree', 'shannon', 'observed_features']
metrics_beta = ['bray_curtis', 'unweighted_unifrac', 'weighted_unifrac']

#max_rows = max(len(alpha_div_gut['results'].keys()), len(gut_beta_div['results'].keys()))
#fig, axes = plt.subplots(max_rows, 2, figsize=(5*2, 5*max_rows))
for i, metric in enumerate(metrics_alpha):
    alpha=redundancy_analysis_alpha(meta_nocomp[meta_nocomp['genotype'].isin(['BALB/c'])],
                             alpha_div_sebum['results'][metric],
                             categorials=categorials,
                             ordinals=ordinals,
                             intervals=intervals,
                             dates=dates,
                             #ax=axes[i][0],
                             title='alpha %s' % metric,
                             dry=False,
                             wait=True,
                             use_grid=True,
                             dirty=True,
                             nocache=True)
    plt.savefig(work_dir + "/gut_redundancies_seperate_%s.png" % metric, transparent=False, bbox_inches='tight')
for i, metric in enumerate(metrics_beta):
    beta=redundancy_analysis_beta(meta_nocomp[meta_nocomp['genotype'].isin(['BALB/c'])],
                             sebum_beta_div['results'][metric],
                             metric,
                             categorials=categorials,
                             ordinals=ordinals,
                             intervals=intervals,
                             dates=dates,
                             #ax=axes[i][1],
                             title='beta %s' % metric,
                             dry=False,
                             wait=True,
                             use_grid=True,
                             dirty=True,
                             nocache=True)
    plt.savefig(work_dir + "/gut_redundancies_seperate_%s.png" % metric, transparent=False, bbox_inches='tight')

# %% [code]
%matplotlib inline
#categorials
categorials = ['season', 'independent_cage_lineage', 'sex']

#ordinals
ordinals = {'generation': ['P1', 'F1', 'F2', 'F3', 'F4', 'F5']}

#intervals
intervals = []

#dates
dates = {}

#metrics
metrics_alpha = ['PD_whole_tree', 'shannon', 'observed_features']
metrics_beta = ['bray_curtis', 'unweighted_unifrac', 'weighted_unifrac']

#max_rows = max(len(alpha_div_gut['results'].keys()), len(gut_beta_div['results'].keys()))
#fig, axes = plt.subplots(max_rows, 2, figsize=(5*2, 5*max_rows))
for i, metric in enumerate(metrics_alpha):
    alpha=redundancy_analysis_alpha(meta_nocomp[meta_nocomp['genotype'].isin(['C57BL6/J'])],
                             alpha_div_gut['results'][metric],
                             categorials=categorials,
                             ordinals=ordinals,
                             intervals=intervals,
                             dates=dates,
                             #ax=axes[i][0],
                             title='alpha %s' % metric,
                             dry=False,
                             wait=True,
                             use_grid=True,
                             dirty=True,
                             nocache=True)
    plt.savefig(work_dir + "/gut_redundancies_seperate_%s.png" % metric, transparent=False, bbox_inches='tight')
for i, metric in enumerate(metrics_beta):
    beta=redundancy_analysis_beta(meta_nocomp[meta_nocomp['genotype'].isin(['C57BL6/J'])],
                             gut_beta_div['results'][metric],
                             metric,
                             categorials=categorials,
                             ordinals=ordinals,
                             intervals=intervals,
                             dates=dates,
                             #ax=axes[i][1],
                             title='beta %s' % metric,
                             dry=False,
                             wait=True,
                             use_grid=True,
                             dirty=True,
                             nocache=True)
    plt.savefig(work_dir + "/gut_redundancies_seperate_%s.png" % metric, transparent=False, bbox_inches='tight')

# %% [markdown]
# Source Tracking
### Gut

# %% [code]
rarefied6000_gut = biom2pandas(work_dir + "/rarefied6000-filtered-gut-table_export/feature-table.biom").fillna(0)

# %% [markdown]
#### BALB/c
##### Impact of BF1 cage groups on BF2 cage groups

# %% [code]
for sink_group in ["BF2-1", "BF2-2"]:
    meta[meta['genotype'].isin(["BALB/c"])]
    conditions = (
        (meta["cage_group"] == "BF1-1") | (meta["cage_group"] == "BF1-2"),
        meta["cage_group"] == sink_group
    )
    values = ["Source", "Sink"]
    column_name ="source_sink_BF1-%s" % str(sink_group)
    meta[column_name] = np.select(conditions, values, "None")

# %% [code]
BF1_2_1_source_tracking = sourcetracker2(
    rarefied6000_gut, #FeatureTable
    meta,
    col_envname="cage_group",
    col_type="source_sink_BF1-BF2-1",
    dry=False,
    wait=True
    )
BF1_2_2_source_tracking = sourcetracker2(
    rarefied6000_gut, #FeatureTable
    meta,
    col_envname="cage_group",
    col_type="source_sink_BF1-BF2-2",
    dry=False,
    wait=True
    )

# %% [code]
for i,group in enumerate([BF1_2_1_source_tracking, BF1_2_2_source_tracking], 1):
    ax= group["results"].transpose().plot.barh(stacked=True)
    if i == 1:
        for tick in ax.get_yticklabels():
            cage_ID = meta.loc[tick.get_text()]["cage_id"]
            if cage_ID in ("F2-17"):
                tick.set_color('green')
            elif cage_ID in ("F2-18"):
                tick.set_color('blue')
            elif cage_ID in ("F2-19"):
                tick.set_color('darkred')
            elif cage_ID in ("F2-20"):
                tick.set_color('magenta')
            else:
                tick.set_color('black')
        
    else:
        for tick in ax.get_yticklabels():
            cage_ID = meta.loc[tick.get_text()]["cage_id"]
            if cage_ID in ("F2-21"):
                tick.set_color('green')
            elif cage_ID in ("F2-22"):
                tick.set_color('blue')
            elif cage_ID in ("F2-23"):
                tick.set_color('darkred')
            elif cage_ID in ("F2-24"):
                tick.set_color('magenta')
            elif cage_ID in ("F2-25"):
                tick.set_color('teal')
            else:
                tick.set_color('black')
    plt.xlabel('relative impact')   
    plt.title(f'Impact on BF2-{i} by BF1 generation')
    plt.legend(bbox_to_anchor=(1.05, 1), loc=2)
    plt.savefig(work_dir + "/source_tracker_%s.png" %i, transparent=False, bbox_inches='tight')

# %% [markdown]
##### Impact of BF2 cage groups on BF3 cage groups

# %% [code]
for sink_group in ["BF3-1", "BF3-2", "BF3-3"]:
    meta[meta['genotype'].isin(["BALB/c"])]
    conditions = (
        (meta["cage_group"] == "BF2-1") | (meta["cage_group"] == "BF2-2") | (meta["cage_group"] == "BF2-3"),
        meta["cage_group"] == sink_group
    )
    values = ["Source", "Sink"]
    column_name ="source_sink_BF2-" + str(sink_group)
    meta[column_name] = np.select(conditions, values, "None")

# %% [code]
BF2_3_1_source_tracking = sourcetracker2(
    rarefied6000_gut, #FeatureTable
    meta,
    col_envname="cage_group",
    col_type="source_sink_BF2-BF3-1",
    dry=False,
    wait=True
)

BF2_3_2_source_tracking = sourcetracker2(
    rarefied6000_gut, #FeatureTable
    meta,
    col_envname="cage_group",
    col_type="source_sink_BF2-BF3-2",
    dry=False,
    wait=True
)

BF2_3_3_source_tracking = sourcetracker2(
    rarefied6000_gut, #FeatureTable
    meta,
    col_envname="cage_group",
    col_type="source_sink_BF2-BF3-3",
    dry=False,
    wait=True
)

# %% [code]
for i,group in enumerate([BF2_3_1_source_tracking, BF2_3_2_source_tracking, BF2_3_3_source_tracking], 1):
    ax= group["results"].transpose().plot.barh(stacked=True)
    if i == 1:
        for tick in ax.get_yticklabels():
            cage_ID = meta.loc[tick.get_text()]["cage_id"]
            if cage_ID in ("F3-21"):
                tick.set_color('green')
            elif cage_ID in ("F3-22"):
                tick.set_color('blue')
            else:
                tick.set_color('black')
    elif i==2:
        for tick in ax.get_yticklabels():
            cage_ID = meta.loc[tick.get_text()]["cage_id"]
            if cage_ID in ("F3-23"):
                tick.set_color('green')
            elif cage_ID in ("F3-24"):
                tick.set_color('blue')
            elif cage_ID in ("F3-25"):
                tick.set_color('red')
            elif cage_ID in ("F3-26"):
                tick.set_color('magenta')
            elif cage_ID in ("F3-27"):
                tick.set_color('teal')
            else:
                tick.set_color('black')       
    else:
        for tick in ax.get_yticklabels():
            cage_ID = meta.loc[tick.get_text()]["cage_id"]
            if cage_ID in ("F3-28"):
                tick.set_color('green')
            elif cage_ID in ("F3-29"):
                tick.set_color('blue')
            elif cage_ID in ("F3-30"):
                tick.set_color('red')
            elif cage_ID in ("F3-31"):
                tick.set_color('magenta')
            elif cage_ID in ("F3-32"):
                tick.set_color('teal')
            else:
                tick.set_color('black')
    plt.title(f'Impact on BF3-{i} by BF2 generation')
    plt.legend(bbox_to_anchor=(1.05, 1), loc=2)
    plt.savefig(work_dir + "/gut_BALB_source_tracker_%s.png" %i, transparent=False, bbox_inches='tight')

# %% [markdown]
##### Impact of BF3 cage groups on BF4 cage groups

# %% [code]
for sink_group in ["BF4-1", "BF4-2", "BF4-3"]:
    meta[meta['genotype'].isin(["BALB/c"])]
    conditions = (
        (meta["cage_group"] == "BF3-1") | (meta["cage_group"] == "BF3-2") | (meta["cage_group"] == "BF3-3"),
        meta["cage_group"] == sink_group
    )
    values = ["Source", "Sink"]
    column_name ="source_sink_BF3-" + str(sink_group)
    meta[column_name] = np.select(conditions, values, "None")

# %% [code]
BF3_4_1_source_tracking = sourcetracker2(
    rarefied6000_gut, #FeatureTable
    meta,
    col_envname="cage_group",
    col_type="source_sink_BF3-BF4-1",
    dry=False,
    wait=True
)

BF3_4_2_source_tracking = sourcetracker2(
    rarefied6000_gut, #FeatureTable
    meta,
    col_envname="cage_group",
    col_type="source_sink_BF3-BF4-2",
    dry=False,
    wait=True
)

BF3_4_3_source_tracking = sourcetracker2(
    rarefied6000_gut, #FeatureTable
    meta,
    col_envname="cage_group",
    col_type="source_sink_BF3-BF4-3",
    dry=False,
    wait=True
)

# %% [code]
for i,group in enumerate([BF3_4_1_source_tracking, BF3_4_2_source_tracking, BF3_4_3_source_tracking], 1):
    ax= group["results"].transpose().plot.barh(stacked=True)
    if i == 1:
        for tick in ax.get_yticklabels():
            cage_ID = meta.loc[tick.get_text()]["cage_id"]
            if cage_ID in ("F4-19"):
                tick.set_color('green')
            elif cage_ID in ("F4-20"):
                tick.set_color('blue')
            elif cage_ID in ("F4-21"):
                tick.set_color('red')
            elif cage_ID in ("F4-22"):
                tick.set_color('magenta')
            elif cage_ID in ("F4-23"):
                tick.set_color('teal')
            else:
                tick.set_color('black')
    elif i==2:
        for tick in ax.get_yticklabels():
            cage_ID = meta.loc[tick.get_text()]["cage_id"]
            if cage_ID in ("F4-24"):
                tick.set_color('green')
            elif cage_ID in ("F4-25"):
                tick.set_color('blue')
            elif cage_ID in ("F4-26"):
                tick.set_color('red')
            elif cage_ID in ("F4-27"):
                tick.set_color('magenta')
            else:
                tick.set_color('black')       
    else:
        for tick in ax.get_yticklabels():
            cage_ID = meta.loc[tick.get_text()]["cage_id"]
            if cage_ID in ("F4-28"):
                tick.set_color('green')
            elif cage_ID in ("F4-29"):
                tick.set_color('blue')
            elif cage_ID in ("F4-30"):
                tick.set_color('red')
            elif cage_ID in ("F4-31"):
                tick.set_color('magenta')
            else:
                tick.set_color('black')
    plt.title(f'Impact on BF4-{i} by BF3 generation')
    plt.legend(bbox_to_anchor=(1.05, 1), loc=2)

# %% [code]


# %% [markdown]
##### Impact of BF4 cage groups on BF5 cage groups

# %% [code]
for sink_group in ["BF5-1", "BF5-2", "BF5-3"]:
    meta[meta['genotype'].isin(["BALB/c"])]
    conditions = (
        (meta["cage_group"] == "BF4-1") | (meta["cage_group"] == "BF4-2") | (meta["cage_group"] == "BF4-3"),
        meta["cage_group"] == sink_group
    )
    values = ["Source", "Sink"]
    column_name ="source_sink_BF4-" + str(sink_group)
    meta[column_name] = np.select(conditions, values, "None")

# %% [code]
BF4_5_1_source_tracking = sourcetracker2(
    rarefied6000_gut, #FeatureTable
    meta,
    col_envname="cage_group",
    col_type="source_sink_BF4-BF5-1",
    dry=False,
    wait=True
)

BF4_5_2_source_tracking = sourcetracker2(
    rarefied6000_gut, #FeatureTable
    meta,
    col_envname="cage_group",
    col_type="source_sink_BF4-BF5-2",
    dry=False,
    wait=True
)

BF4_5_3_source_tracking = sourcetracker2(
    rarefied6000_gut, #FeatureTable
    meta,
    col_envname="cage_group",
    col_type="source_sink_BF4-BF5-3",
    dry=False,
    wait=True
)

# %% [code]
for i,group in enumerate([BF4_5_1_source_tracking, BF4_5_2_source_tracking, BF4_5_3_source_tracking], 1):
    ax= group["results"].transpose().plot.barh(stacked=True)
    if i == 1:
        for tick in ax.get_yticklabels():
            cage_ID = meta.loc[tick.get_text()]["cage_id"]
            if cage_ID in ("F5-8"):
                tick.set_color('green')
            elif cage_ID in ("F5-9"):
                tick.set_color('blue')
            elif cage_ID in ("F5-10"):
                tick.set_color('red')
            else:
                tick.set_color('black')
    elif i==2:
        for tick in ax.get_yticklabels():
            cage_ID = meta.loc[tick.get_text()]["cage_id"]
            if cage_ID in ("F5-11"):
                tick.set_color('green')
            elif cage_ID in ("F5-12"):
                tick.set_color('blue')
            elif cage_ID in ("F5-13"):
                tick.set_color('red')
            elif cage_ID in ("F5-14"):
                tick.set_color('magenta')
            else:
                tick.set_color('black')       
    else:
        for tick in ax.get_yticklabels():
            cage_ID = meta.loc[tick.get_text()]["cage_id"]
            if cage_ID in ("F5-15"):
                tick.set_color('green')
            elif cage_ID in ("F5-16"):
                tick.set_color('blue')
            elif cage_ID in ("F5-17"):
                tick.set_color('red')
            elif cage_ID in ("F5-18"):
                tick.set_color('magenta')
            elif cage_ID in ("F5-19"):
                tick.set_color('teal')
            else:
                tick.set_color('black')
    plt.title(f'Impact on BF5-{i} by BF4 generation')
    plt.legend(bbox_to_anchor=(1.05, 1), loc=2)
    plt.savefig(work_dir + "/gut_BALB_source_tracker_%s.png" %i, transparent=False, bbox_inches='tight')

# %% [code]


# %% [markdown]
#### C57BL6/J
##### Impact of CF1 cage groups on CF2 cage groups

# %% [code]
for sink_group in ["CF2-1", "CF2-2", "CF2-3"]:
    meta[meta['genotype'].isin(["C57BL6/J"])]
    conditions = (
        (meta["cage_group"] == "CF1-1") | (meta["cage_group"] == "CF1-2") | (meta["cage_group"] == "CF1-3"),
        meta["cage_group"] == sink_group
    )
    values = ["Source", "Sink"]
    column_name ="source_sink_CF1-%s" % str(sink_group)
    meta[column_name] = np.select(conditions, values, "None")

# %% [code]
CF1_2_1_source_tracking = sourcetracker2(
    rarefied6000_gut, #FeatureTaCle
    meta,
    col_envname="cage_group",
    col_type="source_sink_CF1-CF2-1",
    dry=False,
    wait=True
    )
CF1_2_2_source_tracking = sourcetracker2(
    rarefied6000_gut, #FeatureTable
    meta,
    col_envname="cage_group",
    col_type="source_sink_CF1-CF2-2",
    dry=False,
    wait=True
    )
CF1_2_3_source_tracking = sourcetracker2(
    rarefied6000_gut, #FeatureTable
    meta,
    col_envname="cage_group",
    col_type="source_sink_CF1-CF2-3",
    dry=False,
    wait=True
    )

# %% [code]
meta.groupby(["source_sink_CF1-CF2-1", "cage_group"]).size()

# %% [code]
for i,group in enumerate([CF1_2_1_source_tracking, CF1_2_2_source_tracking, CF1_2_3_source_tracking], 1):
    ax= group["results"].transpose().plot.barh(stacked=True)
    if i == 1:
        for tick in ax.get_yticklabels():
            cage_ID = meta.loc[tick.get_text()]["cage_id"]
            if cage_ID in ("F2-1"):
                tick.set_color('green')
            elif cage_ID in ("F2-2"):
                tick.set_color('blue')
            elif cage_ID in ("F2-3"):
                tick.set_color('red')
            elif cage_ID in ("F2-4"):
                tick.set_color('magenta')
            elif cage_ID in ("F2-5"):
                tick.set_color('teal')
            elif cage_ID in ("F2-6"):
                tick.set_color('darkolivegreen')
            elif cage_ID in ("F2-7"):
                tick.set_color('darkorange')
            else:
                tick.set_color('black')
    elif i==2:
        for tick in ax.get_yticklabels():
            cage_ID = meta.loc[tick.get_text()]["cage_id"]
            if cage_ID in ("F2-8"):
                tick.set_color('green')
            elif cage_ID in ("F2-9"):
                tick.set_color('blue')
            elif cage_ID in ("F2-10"):
                tick.set_color('red')
            elif cage_ID in ("F2-11"):
                tick.set_color('magenta')
            else:
                tick.set_color('black')       
    else:
        for tick in ax.get_yticklabels():
            cage_ID = meta.loc[tick.get_text()]["cage_id"]
            if cage_ID in ("F2-12"):
                tick.set_color('green')
            elif cage_ID in ("F2-13"):
                tick.set_color('blue')
            elif cage_ID in ("F2-14"):
                tick.set_color('red')
            elif cage_ID in ("F2-15"):
                tick.set_color('magenta')
            elif cage_ID in ("F2-16"):
                tick.set_color('teal')
            else:
                tick.set_color('black')
    plt.title(f'Impact on CF2-{i} by CF1 generation')
    plt.legend(bbox_to_anchor=(1.05, 1), loc=2)

# %% [markdown]
##### Impact of CF2 cage groups on CF3 cage groups

# %% [code]
for sink_group in ["CF3-1", "CF3-2", "CF3-3"]:
    meta[meta['genotype'].isin(["C57BL6/J"])]
    conditions = (
        (meta["cage_group"] == "CF2-1") | (meta["cage_group"] == "CF2-2") | (meta["cage_group"] == "CF2-3"),
        meta["cage_group"] == sink_group
    )
    values = ["Source", "Sink"]
    column_name ="source_sink_CF2-" + str(sink_group)
    meta[column_name] = np.select(conditions, values, "None")

# %% [code]
CF2_3_1_source_tracking = sourcetracker2(
    rarefied6000_gut, #FeatureTable
    meta,
    col_envname="cage_group",
    col_type="source_sink_CF2-CF3-1",
    dry=False,
    wait=True
)

CF2_3_2_source_tracking = sourcetracker2(
    rarefied6000_gut, #FeatureTable
    meta,
    col_envname="cage_group",
    col_type="source_sink_CF2-CF3-2",
    dry=False,
    wait=True
)

CF2_3_3_source_tracking = sourcetracker2(
    rarefied6000_gut, #FeatureTable
    meta,
    col_envname="cage_group",
    col_type="source_sink_CF2-CF3-3",
    dry=False,
    wait=True
)

# %% [code]
for i,group in enumerate([CF2_3_1_source_tracking, CF2_3_2_source_tracking, CF2_3_3_source_tracking], 1):
    ax= group["results"].transpose().plot.barh(stacked=True)
    if i == 1:
        for tick in ax.get_yticklabels():
            cage_ID = meta.loc[tick.get_text()]["cage_id"]
            if cage_ID in ("F3-1"):
                tick.set_color('green')
            elif cage_ID in ("F3-2"):
                tick.set_color('blue')
            elif cage_ID in ("F3-3"):
                tick.set_color('red')
            elif cage_ID in ("F3-4"):
                tick.set_color('magenta')
            elif cage_ID in ("F3-5"):
                tick.set_color('teal')
            elif cage_ID in ("F3-6"):
                tick.set_color('darkorange')
            else:
                tick.set_color('black')
    elif i==2:
        for tick in ax.get_yticklabels():
            cage_ID = meta.loc[tick.get_text()]["cage_id"]
            if cage_ID in ("F3-7"):
                tick.set_color('green')
            elif cage_ID in ("F3-8"):
                tick.set_color('blue')
            elif cage_ID in ("F3-9"):
                tick.set_color('red')
            elif cage_ID in ("F3-10"):
                tick.set_color('magenta')
            elif cage_ID in ("F3-11"):
                tick.set_color('teal')
            elif cage_ID in ("F3-12"):
                tick.set_color('darkolivegreen')
            elif cage_ID in ("F3-13"):
                tick.set_color('darkorange')
            elif cage_ID in ("F3-14"):
                tick.set_color('dodgerblue')
            else:
                tick.set_color('black')       
    else:
        for tick in ax.get_yticklabels():
            cage_ID = meta.loc[tick.get_text()]["cage_id"]
            if cage_ID in ("F3-15"):
                tick.set_color('green')
            elif cage_ID in ("F3-16"):
                tick.set_color('blue')
            elif cage_ID in ("F3-17"):
                tick.set_color('red')
            elif cage_ID in ("F3-18"):
                tick.set_color('magenta')
            elif cage_ID in ("F3-19"):
                tick.set_color('teal')
            elif cage_ID in ("F3-20"):
                tick.set_color('darkorange')
            else:
                tick.set_color('black')
    plt.title(f'Impact on CF3-{i} by CF2 generation')
    plt.legend(bbox_to_anchor=(1.05, 1), loc=2)

# %% [markdown]
##### Impact of CF3 cage groups on CF4 cage groups

# %% [code]
for sink_group in ["CF4-1", "CF4-2", "CF4-3"]:
    meta[meta['genotype'].isin(["C57BL6/J"])]
    conditions = (
        (meta["cage_group"] == "CF3-1") | (meta["cage_group"] == "CF3-2") | (meta["cage_group"] == "CF3-3"),
        meta["cage_group"] == sink_group
    )
    values = ["Source", "Sink"]
    column_name ="source_sink_CF3-" + str(sink_group)
    meta[column_name] = np.select(conditions, values, "None")

# %% [code]
CF3_4_1_source_tracking = sourcetracker2(
    rarefied6000_gut, #FeatureTable
    meta,
    col_envname="cage_group",
    col_type="source_sink_CF3-CF4-1",
    dry=False,
    wait=True
)

CF3_4_2_source_tracking = sourcetracker2(
    rarefied6000_gut, #FeatureTable
    meta,
    col_envname="cage_group",
    col_type="source_sink_CF3-CF4-2",
    dry=False,
    wait=True
)

CF3_4_3_source_tracking = sourcetracker2(
    rarefied6000_gut, #FeatureTable
    meta,
    col_envname="cage_group",
    col_type="source_sink_CF3-CF4-3",
    dry=False,
    wait=True
)

# %% [code]
for i,group in enumerate([CF3_4_1_source_tracking, CF3_4_2_source_tracking, CF3_4_3_source_tracking], 1):
    ax= group["results"].transpose().plot.barh(stacked=True)
    if i == 1:
        for tick in ax.get_yticklabels():
            cage_ID = meta.loc[tick.get_text()]["cage_id"]
            if cage_ID in ("F4-1"):
                tick.set_color('green')
            elif cage_ID in ("F4-2"):
                tick.set_color('blue')
            elif cage_ID in ("F4-3"):
                tick.set_color('red')
            elif cage_ID in ("F4-4"):
                tick.set_color('magenta')
            elif cage_ID in ("F4-5"):
                tick.set_color('teal')
            elif cage_ID in ("F4-6"):
                tick.set_color('darkorange')
            else:
                tick.set_color('black')
    elif i==2:
        for tick in ax.get_yticklabels():
            cage_ID = meta.loc[tick.get_text()]["cage_id"]
            if cage_ID in ("F4-8"):
                tick.set_color('green')
            elif cage_ID in ("F4-9"):
                tick.set_color('blue')
            elif cage_ID in ("F4-10"):
                tick.set_color('red')
            elif cage_ID in ("F4-11"):
                tick.set_color('magenta')
            elif cage_ID in ("F4-12"):
                tick.set_color('teal')
            elif cage_ID in ("F4-13"):
                tick.set_color('darkorange')
            else:
                tick.set_color('black')       
    else:
        for tick in ax.get_yticklabels():
            cage_ID = meta.loc[tick.get_text()]["cage_id"]
            if cage_ID in ("F4-14"):
                tick.set_color('green')
            elif cage_ID in ("F4-15"):
                tick.set_color('blue')
            elif cage_ID in ("F4-16"):
                tick.set_color('red')
            elif cage_ID in ("F4-17"):
                tick.set_color('magenta')
            elif cage_ID in ("F4-18"):
                tick.set_color('teal')
            else:
                tick.set_color('black')
    plt.title(f'Impact on CF4-{i} by CF3 generation')
    plt.legend(bbox_to_anchor=(1.05, 1), loc=2)

# %% [code]


# %% [markdown]
##### Impact of CF4 cage groups on CF5 cage groups

# %% [code]
for sink_group in ["CF5-1", "CF5-2"]:
    meta[meta['genotype'].isin(["C57BL6/J"])]
    conditions = (
        (meta["cage_group"] == "CF4-1") | (meta["cage_group"] == "CF4-2") | (meta["cage_group"] == "CF4-3"),
        meta["cage_group"] == sink_group
    )
    values = ["Source", "Sink"]
    column_name ="source_sink_CF4-" + str(sink_group)
    meta[column_name] = np.select(conditions, values, "None")

# %% [code]
CF4_5_1_source_tracking = sourcetracker2(
    rarefied6000_gut, #FeatureTable
    meta,
    col_envname="cage_group",
    col_type="source_sink_CF4-CF5-1",
    dry=False,
    wait=True
)

CF4_5_2_source_tracking = sourcetracker2(
    rarefied6000_gut, #FeatureTable
    meta,
    col_envname="cage_group",
    col_type="source_sink_CF4-CF5-2",
    dry=False,
    wait=True
)

# %% [code]
for i,group in enumerate([CF4_5_1_source_tracking, CF4_5_2_source_tracking], 1):
    ax= group["results"].transpose().plot.barh(stacked=True)
    if i == 1:
        for tick in ax.get_yticklabels():
            cage_ID = meta.loc[tick.get_text()]["cage_id"]
            if cage_ID in ("F5-1"):
                tick.set_color('green')
            elif cage_ID in ("F5-2"):
                tick.set_color('blue')
            elif cage_ID in ("F5-3"):
                tick.set_color('red')
            elif cage_ID in ("F5-4"):
                tick.set_color('magenta')
            else:
                tick.set_color('black')  
    else:
        for tick in ax.get_yticklabels():
            cage_ID = meta.loc[tick.get_text()]["cage_id"]
            if cage_ID in ("F5-5"):
                tick.set_color('green')
            elif cage_ID in ("F5-6"):
                tick.set_color('blue')
            elif cage_ID in ("F5-7"):
                tick.set_color('red')
            else:
                tick.set_color('black')
    plt.title(f'Impact on CF5-{i} by CF4 generation')
    plt.legend(bbox_to_anchor=(1.05, 1), loc=2)

# %% [code]


# %% [markdown]
### Sebum

# %% [code]
rarefied_sebum = biom2pandas(work_dir + "/rarefied-filtered-sebum-table_export/feature-table.biom").fillna(0)

# %% [markdown]
#### BALB/c
##### Impact of BF1 cage groups on BF2 cage groups

# %% [code]
BF1_2_1_source_tracking_sebum = sourcetracker2(
    rarefied_sebum, #FeatureTable
    meta,
    col_envname="cage_group",
    col_type="source_sink_BF1-BF2-1",
    dry=False,
    wait=True
    )
BF1_2_2_source_tracking_sebum = sourcetracker2(
    rarefied_sebum, #FeatureTable
    meta,
    col_envname="cage_group",
    col_type="source_sink_BF1-BF2-2",
    dry=False,
    wait=True
    )

# %% [code]
for i,group in enumerate([BF1_2_1_source_tracking_sebum, BF1_2_2_source_tracking_sebum], 1):
    ax= group["results"].transpose().plot.barh(stacked=True)
    if i == 1:
        for tick in ax.get_yticklabels():
            cage_ID = meta.loc[tick.get_text()]["cage_id"]
            if cage_ID in ("F2-17"):
                tick.set_color('green')
            elif cage_ID in ("F2-18"):
                tick.set_color('blue')
            elif cage_ID in ("F2-19"):
                tick.set_color('darkred')
            elif cage_ID in ("F2-20"):
                tick.set_color('magenta')
            else:
                tick.set_color('black')
        
    else:
        for tick in ax.get_yticklabels():
            cage_ID = meta.loc[tick.get_text()]["cage_id"]
            if cage_ID in ("F2-21"):
                tick.set_color('green')
            elif cage_ID in ("F2-22"):
                tick.set_color('blue')
            elif cage_ID in ("F2-23"):
                tick.set_color('darkred')
            elif cage_ID in ("F2-24"):
                tick.set_color('magenta')
            elif cage_ID in ("F2-25"):
                tick.set_color('teal')
            else:
                tick.set_color('black')
    plt.title(f'Impact on BF2-{i} by BF1 generation')
    plt.legend(bbox_to_anchor=(1.05, 1), loc=2)

# %% [markdown]
##### Impact of BF2 cage groups on BF3 cage groups

# %% [code]
BF2_3_1_source_tracking_sebum = sourcetracker2(
    rarefied_sebum, #FeatureTable
    meta,
    col_envname="cage_group",
    col_type="source_sink_BF2-BF3-1",
    dry=False,
    wait=True
)

BF2_3_2_source_tracking_sebum = sourcetracker2(
    rarefied_sebum, #FeatureTable
    meta,
    col_envname="cage_group",
    col_type="source_sink_BF2-BF3-2",
    dry=False,
    wait=True
)

BF2_3_3_source_tracking_sebum = sourcetracker2(
    rarefied_sebum, #FeatureTable
    meta,
    col_envname="cage_group",
    col_type="source_sink_BF2-BF3-3",
    dry=False,
    wait=True
)

# %% [code]
for i,group in enumerate([BF2_3_1_source_tracking_sebum, BF2_3_2_source_tracking_sebum, BF2_3_3_source_tracking_sebum], 1):
    ax= group["results"].transpose().plot.barh(stacked=True)
    if i == 1:
        for tick in ax.get_yticklabels():
            cage_ID = meta.loc[tick.get_text()]["cage_id"]
            if cage_ID in ("F3-21"):
                tick.set_color('green')
            elif cage_ID in ("F3-22"):
                tick.set_color('blue')
            else:
                tick.set_color('black')
    elif i==2:
        for tick in ax.get_yticklabels():
            cage_ID = meta.loc[tick.get_text()]["cage_id"]
            if cage_ID in ("F3-23"):
                tick.set_color('green')
            elif cage_ID in ("F3-24"):
                tick.set_color('blue')
            elif cage_ID in ("F3-25"):
                tick.set_color('red')
            elif cage_ID in ("F3-26"):
                tick.set_color('magenta')
            elif cage_ID in ("F3-27"):
                tick.set_color('teal')
            else:
                tick.set_color('black')       
    else:
        for tick in ax.get_yticklabels():
            cage_ID = meta.loc[tick.get_text()]["cage_id"]
            if cage_ID in ("F3-28"):
                tick.set_color('green')
            elif cage_ID in ("F3-29"):
                tick.set_color('blue')
            elif cage_ID in ("F3-30"):
                tick.set_color('red')
            elif cage_ID in ("F3-31"):
                tick.set_color('magenta')
            elif cage_ID in ("F3-32"):
                tick.set_color('teal')
            else:
                tick.set_color('black')
    plt.title(f'Impact on BF3-{i} by BF2 generation')
    plt.legend(bbox_to_anchor=(1.05, 1), loc=2)

# %% [markdown]
##### Impact of BF3 cage groups on BF4 cage groups

# %% [code]
BF3_4_1_source_tracking_sebum = sourcetracker2(
    rarefied_sebum, #FeatureTable
    meta,
    col_envname="cage_group",
    col_type="source_sink_BF3-BF4-1",
    dry=False,
    wait=True
)

BF3_4_2_source_tracking_sebum = sourcetracker2(
    rarefied_sebum, #FeatureTable
    meta,
    col_envname="cage_group",
    col_type="source_sink_BF3-BF4-2",
    dry=False,
    wait=True
)

BF3_4_3_source_tracking_sebum = sourcetracker2(
    rarefied_sebum, #FeatureTable
    meta,
    col_envname="cage_group",
    col_type="source_sink_BF3-BF4-3",
    dry=False,
    wait=True
)

# %% [code]
for i,group in enumerate([BF3_4_1_source_tracking_sebum, BF3_4_2_source_tracking_sebum, BF3_4_3_source_tracking_sebum], 1):
    ax= group["results"].transpose().plot.barh(stacked=True)
    if i == 1:
        for tick in ax.get_yticklabels():
            cage_ID = meta.loc[tick.get_text()]["cage_id"]
            if cage_ID in ("F4-19"):
                tick.set_color('green')
            elif cage_ID in ("F4-20"):
                tick.set_color('blue')
            elif cage_ID in ("F4-21"):
                tick.set_color('red')
            elif cage_ID in ("F4-22"):
                tick.set_color('magenta')
            elif cage_ID in ("F4-23"):
                tick.set_color('teal')
            else:
                tick.set_color('black')
    elif i==2:
        for tick in ax.get_yticklabels():
            cage_ID = meta.loc[tick.get_text()]["cage_id"]
            if cage_ID in ("F4-24"):
                tick.set_color('green')
            elif cage_ID in ("F4-25"):
                tick.set_color('blue')
            elif cage_ID in ("F4-26"):
                tick.set_color('red')
            elif cage_ID in ("F4-27"):
                tick.set_color('magenta')
            else:
                tick.set_color('black')       
    else:
        for tick in ax.get_yticklabels():
            cage_ID = meta.loc[tick.get_text()]["cage_id"]
            if cage_ID in ("F4-28"):
                tick.set_color('green')
            elif cage_ID in ("F4-29"):
                tick.set_color('blue')
            elif cage_ID in ("F4-30"):
                tick.set_color('red')
            elif cage_ID in ("F4-31"):
                tick.set_color('magenta')
            else:
                tick.set_color('black')
    plt.title(f'Impact on BF4-{i} by BF3 generation')
    plt.legend(bbox_to_anchor=(1.05, 1), loc=2)

# %% [code]


# %% [markdown]
##### Impact of BF4 cage groups on BF5 cage groups

# %% [code]
BF4_5_1_source_tracking_sebum = sourcetracker2(
    rarefied_sebum, #FeatureTable
    meta,
    col_envname="cage_group",
    col_type="source_sink_BF4-BF5-1",
    dry=False,
    wait=True
)

BF4_5_2_source_tracking_sebum = sourcetracker2(
    rarefied_sebum, #FeatureTable
    meta,
    col_envname="cage_group",
    col_type="source_sink_BF4-BF5-2",
    dry=False,
    wait=True
)

BF4_5_3_source_tracking_sebum = sourcetracker2(
    rarefied_sebum, #FeatureTable
    meta,
    col_envname="cage_group",
    col_type="source_sink_BF4-BF5-3",
    dry=False,
    wait=True
)

# %% [code]
for i,group in enumerate([BF4_5_1_source_tracking_sebum, BF4_5_2_source_tracking_sebum, BF4_5_3_source_tracking_sebum], 1):
    ax= group["results"].transpose().plot.barh(stacked=True)
    if i == 1:
        for tick in ax.get_yticklabels():
            cage_ID = meta.loc[tick.get_text()]["cage_id"]
            if cage_ID in ("F5-8"):
                tick.set_color('green')
            elif cage_ID in ("F5-9"):
                tick.set_color('blue')
            elif cage_ID in ("F5-10"):
                tick.set_color('red')
            else:
                tick.set_color('black')
    elif i==2:
        for tick in ax.get_yticklabels():
            cage_ID = meta.loc[tick.get_text()]["cage_id"]
            if cage_ID in ("F5-11"):
                tick.set_color('green')
            elif cage_ID in ("F5-12"):
                tick.set_color('blue')
            elif cage_ID in ("F5-13"):
                tick.set_color('red')
            elif cage_ID in ("F5-14"):
                tick.set_color('magenta')
            else:
                tick.set_color('black')       
    else:
        for tick in ax.get_yticklabels():
            cage_ID = meta.loc[tick.get_text()]["cage_id"]
            if cage_ID in ("F5-15"):
                tick.set_color('green')
            elif cage_ID in ("F5-16"):
                tick.set_color('blue')
            elif cage_ID in ("F5-17"):
                tick.set_color('red')
            elif cage_ID in ("F5-18"):
                tick.set_color('magenta')
            elif cage_ID in ("F5-19"):
                tick.set_color('teal')
            else:
                tick.set_color('black')
    plt.title(f'Impact on BF5-{i} by BF4 generation')
    plt.legend(bbox_to_anchor=(1.05, 1), loc=2)

# %% [code]


# %% [markdown]
#### C57BL6/J
##### Impact of CF1 cage groups on CF2 cage groups

# %% [code]
CF1_2_1_source_tracking_sebum = sourcetracker2(
    rarefied_sebum, #FeatureTaCle
    meta,
    col_envname="cage_group",
    col_type="source_sink_CF1-CF2-1",
    dry=False,
    wait=True
    )
CF1_2_2_source_tracking_sebum = sourcetracker2(
    rarefied_sebum, #FeatureTable
    meta,
    col_envname="cage_group",
    col_type="source_sink_CF1-CF2-2",
    dry=False,
    wait=True
    )
CF1_2_3_source_tracking_sebum = sourcetracker2(
    rarefied_sebum, #FeatureTable
    meta,
    col_envname="cage_group",
    col_type="source_sink_CF1-CF2-3",
    dry=False,
    wait=True
    )

# %% [code]
for i,group in enumerate([CF1_2_1_source_tracking_sebum, CF1_2_2_source_tracking_sebum, CF1_2_3_source_tracking_sebum], 1):
    ax= group["results"].transpose().plot.barh(stacked=True)
    if i == 1:
        for tick in ax.get_yticklabels():
            cage_ID = meta.loc[tick.get_text()]["cage_id"]
            if cage_ID in ("F2-1"):
                tick.set_color('green')
            elif cage_ID in ("F2-2"):
                tick.set_color('blue')
            elif cage_ID in ("F2-3"):
                tick.set_color('red')
            elif cage_ID in ("F2-4"):
                tick.set_color('magenta')
            elif cage_ID in ("F2-5"):
                tick.set_color('teal')
            elif cage_ID in ("F2-6"):
                tick.set_color('darkolivegreen')
            elif cage_ID in ("F2-7"):
                tick.set_color('darkorange')
            else:
                tick.set_color('black')
    elif i==2:
        for tick in ax.get_yticklabels():
            cage_ID = meta.loc[tick.get_text()]["cage_id"]
            if cage_ID in ("F2-8"):
                tick.set_color('green')
            elif cage_ID in ("F2-9"):
                tick.set_color('blue')
            elif cage_ID in ("F2-10"):
                tick.set_color('red')
            elif cage_ID in ("F2-11"):
                tick.set_color('magenta')
            else:
                tick.set_color('black')       
    else:
        for tick in ax.get_yticklabels():
            cage_ID = meta.loc[tick.get_text()]["cage_id"]
            if cage_ID in ("F2-12"):
                tick.set_color('green')
            elif cage_ID in ("F2-13"):
                tick.set_color('blue')
            elif cage_ID in ("F2-14"):
                tick.set_color('red')
            elif cage_ID in ("F2-15"):
                tick.set_color('magenta')
            elif cage_ID in ("F2-16"):
                tick.set_color('teal')
            else:
                tick.set_color('black')
    plt.title(f'Impact on CF2-{i} by CF1 generation')
    plt.legend(bbox_to_anchor=(1.05, 1), loc=2)

# %% [markdown]
##### Impact of CF2 cage groups on CF3 cage groups

# %% [code]
CF2_3_1_source_tracking_sebum = sourcetracker2(
    rarefied_sebum, #FeatureTable
    meta,
    col_envname="cage_group",
    col_type="source_sink_CF2-CF3-1",
    dry=False,
    wait=True
)

CF2_3_2_source_tracking_sebum = sourcetracker2(
    rarefied_sebum, #FeatureTable
    meta,
    col_envname="cage_group",
    col_type="source_sink_CF2-CF3-2",
    dry=False,
    wait=True
)

CF2_3_3_source_tracking_sebum = sourcetracker2(
    rarefied_sebum, #FeatureTable
    meta,
    col_envname="cage_group",
    col_type="source_sink_CF2-CF3-3",
    dry=False,
    wait=True
)

# %% [code]
meta[meta['cage_group'].isin(['CF3-1'])]['cage_id'].value_counts()

# %% [code]
for i,group in enumerate([CF2_3_1_source_tracking_sebum, CF2_3_2_source_tracking_sebum, CF2_3_3_source_tracking_sebum], 1):
    ax= group["results"].transpose().plot.barh(stacked=True)
    if i == 1:
        for tick in ax.get_yticklabels():
            cage_ID = meta.loc[tick.get_text()]["cage_id"]
            if cage_ID in ("F3-1"):
                tick.set_color('green')
            elif cage_ID in ("F3-2"):
                tick.set_color('blue')
            elif cage_ID in ("F3-3"):
                tick.set_color('red')
            elif cage_ID in ("F3-4"):
                tick.set_color('magenta')
            elif cage_ID in ("F3-5"):
                tick.set_color('teal')
            elif cage_ID in ("F3-6"):
                tick.set_color('darkorange')
            else:
                tick.set_color('black')
    elif i==2:
        for tick in ax.get_yticklabels():
            cage_ID = meta.loc[tick.get_text()]["cage_id"]
            if cage_ID in ("F3-7"):
                tick.set_color('green')
            elif cage_ID in ("F3-8"):
                tick.set_color('blue')
            elif cage_ID in ("F3-9"):
                tick.set_color('red')
            elif cage_ID in ("F3-10"):
                tick.set_color('magenta')
            elif cage_ID in ("F3-11"):
                tick.set_color('teal')
            elif cage_ID in ("F3-12"):
                tick.set_color('darkolivegreen')
            elif cage_ID in ("F3-13"):
                tick.set_color('darkorange')
            elif cage_ID in ("F3-14"):
                tick.set_color('dodgerblue')
            else:
                tick.set_color('black')       
    else:
        for tick in ax.get_yticklabels():
            cage_ID = meta.loc[tick.get_text()]["cage_id"]
            if cage_ID in ("F3-15"):
                tick.set_color('green')
            elif cage_ID in ("F3-16"):
                tick.set_color('blue')
            elif cage_ID in ("F3-17"):
                tick.set_color('red')
            elif cage_ID in ("F3-18"):
                tick.set_color('magenta')
            elif cage_ID in ("F3-19"):
                tick.set_color('teal')
            elif cage_ID in ("F3-20"):
                tick.set_color('darkorange')
            else:
                tick.set_color('black')
    plt.title(f'Impact on CF3-{i} by CF2 generation')
    plt.legend(bbox_to_anchor=(1.05, 1), loc=2)

# %% [markdown]
##### Impact of CF3 cage groups on CF4 cage groups

# %% [code]
CF3_4_1_source_tracking_sebum = sourcetracker2(
    rarefied_sebum, #FeatureTable
    meta,
    col_envname="cage_group",
    col_type="source_sink_CF3-CF4-1",
    dry=False,
    wait=True
)

CF3_4_2_source_tracking_sebum = sourcetracker2(
    rarefied_sebum, #FeatureTable
    meta,
    col_envname="cage_group",
    col_type="source_sink_CF3-CF4-2",
    dry=False,
    wait=True
)

CF3_4_3_source_tracking_sebum = sourcetracker2(
    rarefied_sebum, #FeatureTable
    meta,
    col_envname="cage_group",
    col_type="source_sink_CF3-CF4-3",
    dry=False,
    wait=True
)

# %% [code]
for i,group in enumerate([CF3_4_1_source_tracking_sebum, CF3_4_2_source_tracking_sebum, CF3_4_3_source_tracking_sebum], 1):
    ax= group["results"].transpose().plot.barh(stacked=True)
    if i == 1:
        for tick in ax.get_yticklabels():
            cage_ID = meta.loc[tick.get_text()]["cage_id"]
            if cage_ID in ("F4-1"):
                tick.set_color('green')
            elif cage_ID in ("F4-2"):
                tick.set_color('blue')
            elif cage_ID in ("F4-3"):
                tick.set_color('red')
            elif cage_ID in ("F4-4"):
                tick.set_color('magenta')
            elif cage_ID in ("F4-5"):
                tick.set_color('teal')
            elif cage_ID in ("F4-6"):
                tick.set_color('darkorange')
            else:
                tick.set_color('black')
    elif i==2:
        for tick in ax.get_yticklabels():
            cage_ID = meta.loc[tick.get_text()]["cage_id"]
            if cage_ID in ("F4-8"):
                tick.set_color('green')
            elif cage_ID in ("F4-9"):
                tick.set_color('blue')
            elif cage_ID in ("F4-10"):
                tick.set_color('red')
            elif cage_ID in ("F4-11"):
                tick.set_color('magenta')
            elif cage_ID in ("F4-12"):
                tick.set_color('teal')
            elif cage_ID in ("F4-13"):
                tick.set_color('darkorange')
            else:
                tick.set_color('black')       
    else:
        for tick in ax.get_yticklabels():
            cage_ID = meta.loc[tick.get_text()]["cage_id"]
            if cage_ID in ("F4-14"):
                tick.set_color('green')
            elif cage_ID in ("F4-15"):
                tick.set_color('blue')
            elif cage_ID in ("F4-16"):
                tick.set_color('red')
            elif cage_ID in ("F4-17"):
                tick.set_color('magenta')
            elif cage_ID in ("F4-18"):
                tick.set_color('teal')
            else:
                tick.set_color('black')
    plt.title(f'Impact on CF4-{i} by CF3 generation')
    plt.legend(bbox_to_anchor=(1.05, 1), loc=2)

# %% [code]


# %% [markdown]
##### Impact of CF4 cage groups on CF5 cage groups

# %% [code]
CF4_5_1_source_tracking_sebum = sourcetracker2(
    rarefied_sebum, #FeatureTable
    meta,
    col_envname="cage_group",
    col_type="source_sink_CF4-CF5-1",
    dry=False,
    wait=True
)

CF4_5_2_source_tracking_sebum = sourcetracker2(
    rarefied_sebum, #FeatureTable
    meta,
    col_envname="cage_group",
    col_type="source_sink_CF4-CF5-2",
    dry=False,
    wait=True
)

# %% [code]
for i,group in enumerate([CF4_5_1_source_tracking_sebum, CF4_5_2_source_tracking_sebum], 1):
    ax= group["results"].transpose().plot.barh(stacked=True)
    if i == 1:
        for tick in ax.get_yticklabels():
            cage_ID = meta.loc[tick.get_text()]["cage_id"]
            if cage_ID in ("F5-1"):
                tick.set_color('green')
            elif cage_ID in ("F5-2"):
                tick.set_color('blue')
            elif cage_ID in ("F5-3"):
                tick.set_color('red')
            elif cage_ID in ("F5-4"):
                tick.set_color('magenta')
            else:
                tick.set_color('black')  
    else:
        for tick in ax.get_yticklabels():
            cage_ID = meta.loc[tick.get_text()]["cage_id"]
            if cage_ID in ("F5-5"):
                tick.set_color('green')
            elif cage_ID in ("F5-6"):
                tick.set_color('blue')
            elif cage_ID in ("F5-7"):
                tick.set_color('red')
            else:
                tick.set_color('black')
    plt.title(f'Impact on CF5-{i} by CF4 generation')
    plt.legend(bbox_to_anchor=(1.05, 1), loc=2)

# %% [code]


# %% [markdown]
# Seasonal Changes?

# %% [code]
meta.loc[meta.collection_timestamp.str.contains('\d{4}-0[456789]-\d{2}'), :].index

# %% [code]
meta['season'] = np.where(
    meta.index.isin(meta.loc[meta.collection_timestamp.str.contains('\d{4}-0[456789]-\d{2}'), :].index), 'summer', 'winter') 

# %% [markdown]
### Gut
#### BALB/c

# %% [code]
categorials=["season"]

season_gut_beta_group_significance = correlation_diversity_metacolumns(
    meta[meta['genotype'].isin(['BALB/c'])], 
    categorial=categorials, 
    alpha_diversities=alpha_div_gut['results'],
    beta_diversities=gut_beta_div['results'],
    dry=False,
    use_grid=True
)

# %% [code]
season_gut_beta_group_significance['results'][season_gut_beta_group_significance['results']['column'].isin(['season'])]

# %% [markdown]
#### C57B

# %% [code]
categorials=["season"]

season_gut_beta_group_significance = correlation_diversity_metacolumns(
    meta[meta['genotype'].isin(['C57BL6/J'])], 
    categorial=categorials, 
    alpha_diversities=alpha_div_gut['results'],
    beta_diversities=gut_beta_div['results'],
    dry=False,
    use_grid=True
)

# %% [code]
season_gut_beta_group_significance['results'][season_gut_beta_group_significance['results']['column'].isin(['season'])]

# %% [code]


# %% [code]


# %% [markdown]
### Sebum
#### BALB/c

# %% [code]
categorials=["season"]

season_sebum_beta_group_significance_BALB = correlation_diversity_metacolumns(
    meta[meta['genotype'].isin(['BALB/c'])], 
    categorial=categorials, 
    alpha_diversities=alpha_div_sebum['results'],
    beta_diversities=sebum_beta_div['results'],
    dry=False,
    use_grid=True
)

# %% [code]
season_sebum_beta_group_significance_BALB['results'][season_sebum_beta_group_significance_BALB['results']['column'].isin(['season'])]

# %% [markdown]
#### C57B

# %% [code]
categorials=["season"]

season_sebum_beta_group_significance_C57B = correlation_diversity_metacolumns(
    meta[meta['genotype'].isin(['C57BL6/J'])], 
    categorial=categorials, 
    alpha_diversities=alpha_div_sebum['results'],
    beta_diversities=sebum_beta_div['results'],
    dry=False,
    use_grid=True
)

# %% [code]
! rm -r /homes/arehm/TMP/ana_corr-divmeta_lgyo__dm

# %% [code]
season_sebum_beta_group_significance_C57B['results'][season_sebum_beta_group_significance_C57B['results']['column'].isin(['season'])]

# %% [code]


# %% [markdown]
# Metadata correlation

# %% [code]
meta['description'].value_counts()

# %% [code]
%matplotlib inline
#categorials
categorials=['sex', 
             'genotype',
             'season',
             'cage_group',
             'cage_id',
             'env_material',
             'geo_loc_name',
             'host_body_habitat',
             'host_body_product',
             'host_body_site',
             'batch',
             'description',
             'empo_1',
             'empo_2',
             'empo_3',
             'mother_subject_id',
             'parental_cage',
             'cage_lineage',
             'comparison',
             'generation'
            ]

#ordinals
ordinals = {}

#intervals
intervals = []

#dates
dates = {'collection_timestamp': 'yyyy-mm-dd', 'birth_timestamp': 'yyyy-mm-dd'}


correlate_metadata(meta, categorials=categorials, intervals=intervals, dates=dates, ordinals=ordinals)

# %% [code]
meta_groupby= pd.DataFrame(meta.groupby(["host_body_product","genotype", "season", "generation", "cage_group", "cage_id", "collection_timestamp"]).size())

# %% [code]
meta_groupby.to_csv(work_dir + '/seasonality_table.csv', sep="\t")

# %% [code]
meta.groupby(["host_body_product","genotype", "season", "generation", "cage_group", "cage_id", "collection_timestamp"]).size()

# %% [code]
meta_groupby

# %% [code]
meta.groupby(["host_body_product","genotype", "season", "generation", "cage_group", "cage_id"]).size()

# %% [code]
meta['comparison'].value_counts()

# %% [code]
meta.groupby(["host_body_product","genotype", "sex"]).size()

# %% [markdown]
# QURRO

# %% [markdown]
### Run DEICODE

# %% [code]
! $q_path; qiime deicode --help

# %% [code]
meta['host_common_name']

# %% [code]
%time ! $q_path; qiime feature-table filter-samples \
  --i-table $work_dir/fragment-filtered‐feature-table.qza \
  --m-metadata-file $meta_file \
  --p-where "[sample_name] NOT IN ($comparison_mice_indeces) AND [host_body_product] IN ('UBERON:lumen of gut') AND [host_common_name] IN ('mouse')" \
  --o-filtered-table $work_dir/filtered-gut-table-mouse-for-deicode.qza

# %% [markdown]
Tutorial: https://forum.qiime2.org/t/robust-aitchison-pca-beta-diversity-with-deicode/8333

# %% [code]
#running DEICODE on gut samples (filtered for contaminants)
! $q_path; qiime deicode rpca \
    --i-table $work_dir/filtered-gut-table-mouse-for-deicode.qza \
    --p-min-feature-count 10 \
    --p-min-sample-count 500 \
    --o-biplot $work_dir/gut_deicode_ordination.qza \
    --o-distance-matrix $work_dir/gut_deicode_distance.qza

# %% [markdown]
### Running Qurro

# %% [code]
! $q_path; qiime qurro loading-plot \
    --i-table $work_dir/filtered-gut-table-mouse-for-deicode.qza \
    --i-ranks $work_dir/gut_deicode_ordination.qza \
    --m-sample-metadata-file meta_gen_num.txt \
    --m-feature-metadata-file $work_dir/taxonomy_full.qza \
    --o-visualization $work_dir/gut-qurro-plot.qzv

# %% [markdown]
# ITS
import of ITS Sequences

# %% [code]
! $q_path; qiime tools import \
  --type 'SampleData[PairedEndSequencesWithQuality]' \
  --input-path /vol/jlab/MicrobiomeAnalyses/Projects/Benga_ThreeGen/Trimming/ITS_sequences/ \
  --input-format CasavaOneEightSingleLanePerSampleDirFmt \
  --output-path $work_dir/ITS-demux-paired-end.qza

# %% [markdown]
quality control with DADA2, NOT USED

# %% [code]
! $q_path; qiime dada2 denoise-single \
  --i-demultiplexed-seqs $work_dir/ITS-demux-paired-end.qza \
  --p-trim-left 13 \
  --p-trunc-len 150 \
  --o-representative-sequences $work_dir/its-dada2-single-end-rep-seqs.qza \
  --o-table $work_dir/its-dada2-single-end-table.qza \
  --o-denoising-stats $work_dir/its-dada2-single-end-stats.qza

# %% [code]
! $q_path; qiime metadata tabulate \
  --m-input-file $work_dir/its-dada2-single-end-stats.qza \
  --o-visualization $work_dir/its-dada2-single-end-stats.qzv

# %% [markdown]
## deblur
quality control with deblur outside of QIIME2

# %% [code]
! $q_path; deblur workflow --help

# %% [code]
! $q_path; deblur workflow \
--seqs-fp /vol/jlab/MicrobiomeAnalyses/Projects/Benga_ThreeGen/Trimming/ITS_sequences/ \
--output-dir $work_dir/deblur-its \
--trim-length 150


# %% [markdown]
import of `all.biom` file from deblur

# %% [code]
! $q_path; qiime tools import \
  --input-path $work_dir/deblur-its/all.biom \
  --type 'FeatureTable[Frequency]' \
  --input-format BIOMV210Format \
  --output-path $work_dir/deblur-its/its-feature-table-deblur.qza

# %% [code]
%time ! $q_path; qiime feature-table summarize \
  --i-table $work_dir/deblur-its/its-feature-table-deblur.qza\
  --o-visualization $work_dir/deblur-its/its-feature-table-deblur-summary.qzv

# %% [markdown]
## taxonomy with UNITE
import sequences from unite database ver8 99% 10.05.2021

# %% [code]
%time ! $q_path; qiime tools import \
  --input-path /homes/arehm/Documents/Thesis/JupyterNotebook/UNITE_8_3/sh_qiime_release_10.05.2021/sh_refs_qiime_ver8_99_10.05.2021.fasta \
  --type 'FeatureData[Sequence]' \
  --output-path $work_dir/unite_refs_qiime_ver8_99_10.05.2021_rep-seqs.qza

# %% [markdown]
import taxonomy for QIIME2 from UNITE

# %% [code]
%time ! $q_path; qiime tools import \
  --input-path /homes/arehm/Documents/Thesis/JupyterNotebook/UNITE_8_3/sh_qiime_release_10.05.2021/sh_taxonomy_qiime_ver8_99_10.05.2021.txt \
  --type FeatureData[Taxonomy] \
  --input-format HeaderlessTSVTaxonomyFormat \
  --output-path $work_dir/unite_taxonomy_qiime_ver8_99_10.05.2021_taxonomy.qza

# %% [markdown]
train classifier to fit naive bayes with UNITE data

# %% [code]
%time ! $q_path; qiime feature-classifier fit-classifier-naive-bayes \
  --i-reference-reads $work_dir/unite_refs_qiime_ver8_99_10.05.2021_rep-seqs.qza \
  --i-reference-taxonomy $work_dir/unite_taxonomy_qiime_ver8_99_10.05.2021_taxonomy.qza \
  --o-classifier $work_dir/unite-ver8_99_10.05.2021_classifier.qza 

# %% [markdown]
import sequence data after deblur

# %% [code]
%time ! $q_path; qiime tools import \
  --input-path $work_dir/deblur-its/all.seqs.fa \
  --type 'FeatureData[Sequence]' \
  --output-path $work_dir/its-rep-seqs.qza

# %% [markdown]
run classify-sklearn on ITS sequence data after deblur with unite classifier

# %% [code]
%time ! $q_path ; qiime feature-classifier classify-sklearn \
  --i-reads $work_dir/its-rep-seqs.qza \
  --i-classifier $work_dir/unite-ver8_99_10.05.2021_classifier.qza \
  --o-classification $work_dir/its_taxonomy_full.qza

# %% [markdown]
create barplot

# %% [code]
%time ! $q_path; qiime taxa barplot \
  --i-table $work_dir/deblur-its/its-feature-table-deblur.qza \
  --i-taxonomy $work_dir/its_taxonomy_full.qza \
  --o-visualization $work_dir/its-taxa_barplot_full.qzv

# %% [markdown]
OUTPUT: 
- majority of sequences only gets kingdom assignment -> same with the SILVA database, little bit worse
- some samples show assignment up to the species level -> these samples might be of interest
- connect with metadata to implore which samples these are

# %% [markdown]
## Files from Qiita

# %% [code]
%time ! $q_path; qiime tools import \
  --input-path $work_dir/deblur-its/qiita-its-deblur-final-reference-hit.seqs.fa \
  --type 'FeatureData[Sequence]' \
  --output-path $work_dir/qiita-deblur-final-its-rep-seqs.qza

# %% [code]
%time ! $q_path ; qiime feature-classifier classify-sklearn \
  --i-reads $work_dir/qiita-deblur-final-its-rep-seqs.qza \
  --i-classifier $work_dir/unite-ver8_99_10.05.2021_classifier.qza \
  --o-classification $work_dir/qiita-its_taxonomy_full.qza

# %% [code]
%time ! $q_path; qiime taxa barplot \
  --i-table $work_dir/deblur-its/qiita-its-deblur-final-feature-table.qza \
  --i-taxonomy $work_dir/qiita-its_taxonomy_full.qza \
  --m-metadata-file $work_dir/deblur-its/13422_20220524-060106.txt \
  --o-visualization $work_dir/qiita-its-taxa_barplot_full.qzv

# %% [markdown]
# Metabolome

# %% [code]
#gc_data = pd.read_csv('/homes/arehm/Benga_ThreeGen/Incoming/GC_results/220511__Data_21-0025.csv', sep="\t").rename(columns={'Sample.label.on.tube': 'sample_name'}).set_index("sample_name")
gc_data = pd.read_csv('/homes/arehm/Benga_ThreeGen/Incoming/GC_results/220511__Data_21-0025.csv', sep="\t").rename(columns={'Sample.label.on.tube': 'sample_name'})

# %% [code]
gc_pivot = pd.pivot_table(gc_data, values = "relativeResponse", index = "Compound", columns = "sample_name").fillna(0)

# %% [code]
gc_pivot.loc["3-P-glycerol_UK",["1.1.", "2.1.", "3.1.", "4.1.", "5.1."]]
BALB_list = ["11.1.", "12.1.", "13.1.", "14.1.", "15.1.", "16.1.", "17.1.", "18.1.", "19.1.", "20.1."]
C57B_list = ["1.1.", "2.1.", "3.1.", "4.1.", "5.1.", "6.1.", "7.1.", "8.1.", "9.1.", "10.1."]
F3_list = ["1.1.", "2.1.", "3.1.", "4.1.", "5.1.", "11.1.", "12.1.", "13.1.", "14.1.", "15.1."]
F4_list = ["6.1.", "7.1.", "8.1.", "9.1.", "10.1.", "16.1.", "17.1.", "18.1.", "19.1.", "20.1."]

BALB_set = set(["11.1.", "12.1.", "13.1.", "14.1.", "15.1.", "16.1.", "17.1.", "18.1.", "19.1.", "20.1."])
C57B_set = set(["1.1.", "2.1.", "3.1.", "4.1.", "5.1.", "6.1.", "7.1.", "8.1.", "9.1.", "10.1."])
F3_set = set(["1.1.", "2.1.", "3.1.", "4.1.", "5.1.", "11.1.", "12.1.", "13.1.", "14.1.", "15.1."])
F4_set = set(["6.1.", "7.1.", "8.1.", "9.1.", "10.1.", "16.1.", "17.1.", "18.1.", "19.1.", "20.1."])
list((BALB_set.intersection(F3_set)))

# %% [code]
gc_pivot.shape

# %% [code]
gc_pivot

# %% [code]
meta_gc = gc_data[['sample_name', "date", "MS.Label", "Species", "Tissue.type", "Condition.1", "Generation", "Benga-Nr."]].set_index(['sample_name']).drop_duplicates().rename(columns={'Condition.1': 'genotype', 'Generation': 'generation'})

# %% [code]
meta_gc

# %% [code]
gc_alpha_div_results = dict()
gc_alpha_div_results = alpha_diversity(gc_pivot, metrics=['shannon'], dry=False, wait=True, rarefaction_depth=None)

# %% [code]
gc_alpha_div_results['results']

# %% [code]
ax = gc_alpha_div_results['results'].sort_values(by = 'shannon').plot.bar(y = 'shannon')

# %% [code]
gc_beta_div_results = beta_diversity(gc_pivot, metrics=["bray_curtis"], reference_tree=None, dry=True, use_grid=True)

# %% [code]
gc_data["sample_name"].loc[gc_data["Generation"] == "F3"]

# %% [code]
group_dm

# %% [markdown]
## Beta Diversity - Genotypes in Generation

# %% [code]
res_all_chems = {
    'alpha_diversity': alpha_diversity(gc_pivot, metrics=['shannon'], dry=False, wait=False, rarefaction_depth=None),
    'beta_diversity': beta_diversity(gc_pivot, metrics=['bray_curtis'], dry=False, wait=False)
}


# %% [code]
for generation in ['F3', 'F4']:
    field = 'genotype'
    field_meta = meta_gc[meta_gc['generation'] == generation][field]

    b = detect_distant_groups(res_all_chems['beta_diversity']['results']['bray_curtis'], 'Bray-Curtis', field_meta)
    ax, _ = plotGroup_permanovas(res_all_chems['beta_diversity']['results']['bray_curtis'], field_meta, **b,
                                 horizontal=True, print_sample_numbers=True, colors_boxplot={'between': 'gray', 'left': 'red', 'right': 'blue'})
    ax.set_ylabel(field+ "\n(%i healthy mice)" % field_meta.shape[0])
    ax.yaxis.label.set_visible(True)
    _ = ax.set_title('%s\npairwise %s\n%s permutations' % (generation, str.upper(b['fct_name']), b['num_permutations']))

# %% [markdown]
## Beta Diversity - Generations in Genotype

# %% [code]
for genotype in ['BALB/c', 'C57BL/6J']:
    field = 'generation'
    field_meta = meta_gc[meta_gc['genotype'] == genotype][field]

    b = detect_distant_groups(res_all_chems['beta_diversity']['results']['bray_curtis'], 'Bray-Curtis', field_meta)
    ax, _ = plotGroup_permanovas(res_all_chems['beta_diversity']['results']['bray_curtis'], field_meta, **b,
                                 horizontal=True, print_sample_numbers=True, colors_boxplot={'between': 'gray', 'left': 'red', 'right': 'blue'})
    ax.set_ylabel(field+ "\n(%i healthy mice)" % field_meta.shape[0])
    ax.yaxis.label.set_visible(True)
    _ = ax.set_title('%s\npairwise %s\n%s permutations' % (genotype, str.upper(b['fct_name']), b['num_permutations']))

# %% [markdown]
## Beta Diversity - Genotypes combined / Generations combined

# %% [code]
field = 'genotype'
field_meta = meta_gc[field]

b = detect_distant_groups(res_all_chems['beta_diversity']['results']['bray_curtis'], 'Bray-Curtis', field_meta)
ax, _ = plotGroup_permanovas(res_all_chems['beta_diversity']['results']['bray_curtis'], field_meta, **b,
                             horizontal=True, print_sample_numbers=True, colors_boxplot={'between': 'gray', 'left': 'red', 'right': 'blue'})
ax.set_ylabel(field+ "\n(%i healthy mice)" % field_meta.shape[0])
ax.yaxis.label.set_visible(True)
_ = ax.set_title('%s\npairwise %s\n%s permutations' % (field, str.upper(b['fct_name']), b['num_permutations']))

field = 'generation'
field_meta = meta_gc[field]

b = detect_distant_groups(res_all_chems['beta_diversity']['results']['bray_curtis'], 'Bray-Curtis', field_meta)
ax, _ = plotGroup_permanovas(res_all_chems['beta_diversity']['results']['bray_curtis'], field_meta, **b,
                             horizontal=True, print_sample_numbers=True, colors_boxplot={'between': 'gray', 'left': 'red', 'right': 'blue'})
ax.set_ylabel(field+ "\n(%i healthy mice)" % field_meta.shape[0])
ax.yaxis.label.set_visible(True)
_ = ax.set_title('%s\npairwise %s\n%s permutations' % (field, str.upper(b['fct_name']), b['num_permutations']))

# %% [code]
_ = plotNetworks('genotype', meta_gc, 
                 alpha=res_all_chems["alpha_diversity"]['results'], 
                 beta=res_all_chems["beta_diversity"]['results'], minnumalpha=1)

# %% [markdown]
## dsFDR

# %% [code]
import calour as ca
ca.set_log_level(40)
pandas2biom(work_dir + '/dsfrd_metabolome.biom', gc_pivot)
meta_gc.to_csv(work_dir + '/dsfdr_metabolome_meta.tsv', sep="\t", index=True)
   
dsFDR_experiment = ca.read_amplicon(work_dir + '/dsfrd_metabolome.biom', work_dir + '/dsfdr_metabolome_meta.tsv', min_reads=0, normalize=10000)

# %% [code]
dsFDR_experiment

# %% [code]
field = 'genotype'
res_dsFDR_genotype = find_diff_taxa(dsFDR_experiment, meta_gc[meta_gc['generation'] == 'F4'], [field])

if len(res_dsFDR_genotype) > 0:
    p = plot_diff_taxa(gc_pivot, meta_gc[meta_gc['generation'] == 'F4'][field], res_dsFDR_genotype,
                       min_mean_abundance=0.0,
#                       colors_boxplot={'Pax5+/-': 'red', 'WT': 'blue'}, 
                       color_barplot='gray'
                      )
    p[0].get_axes()[0].set_ylabel("compound")

# %% [code]
for generation in ['F3', 'F4']:
    print(generation)
    field = 'genotype'
    res_dsFDR_generation = find_diff_taxa(dsFDR_experiment, meta_gc[meta_gc['generation'] == generation], [field])

    if len(res_dsFDR_generation) > 0:
        
        p = plot_diff_taxa(gc_pivot, meta_gc[meta_gc['generation'] == generation][field], res_dsFDR_generation,
                           min_mean_abundance=0.0,
    #                       colors_boxplot={'Pax5+/-': 'red', 'WT': 'blue'}, 
                           color_barplot='gray'
                          )
        p[0].get_axes()[0].set_ylabel("compound")
        print(generation)

# %% [code]
for genotype in ['BALB/c', 'C57BL/6J']:
    print(genotype)
    field = 'generation'
    res_dsFDR_genotype = find_diff_taxa(dsFDR_experiment, meta_gc[meta_gc['genotype'] == genotype], [field])

    if len(res_dsFDR_genotype) > 0:
        
        p = plot_diff_taxa(gc_pivot, meta_gc[meta_gc['genotype'] == genotype][field], res_dsFDR_genotype,
                           min_mean_abundance=0.0,
    #                       colors_boxplot={'Pax5+/-': 'red', 'WT': 'blue'}, 
                           color_barplot='gray'
                          )
        p[0].get_axes()[0].set_ylabel("compound")
        print(genotype)

# %% [markdown]
## Triglyceride values

# %% [code]
tg_values = pd.read_csv('/homes/arehm/Benga_ThreeGen/Incoming/Triglyceride.csv', sep="\t").set_index("sample_name")

# %% [code]
tg_values

# %% [code]
type(tg_values_only['triglyceride_values'][1])

# %% [code]
tg_values_only = pd.DataFrame(tg_values['triglyceride_values']).dropna()
df_keinSerum = tg_values_only.index.isin(["13422.286","13422.286skin"])

tg_values_only = tg_values_only[~df_keinSerum]
tg_values_only['genotype'] = tg_values['genotype']
tg_values_only['generation'] = tg_values['generation']

tg_values_only['triglyceride_values'] = pd.to_numeric(tg_values_only['triglyceride_values'])
tg_values_only = tg_values_only.dropna()
tg_values_only

# %% [code]
tg_plot = tg_values_only.groupby('genotype').agg([np.mean])
tg_plot

# %% [code]
ax = tg_plot.plot.bar(legend=False, title='Average Triglyceride concentration per genotype')
ax.set_ylabel("nmol/well")

# %% [code]
grouped = tg_values_only.groupby(['genotype'])

# %% [code]
tg_values_only[tg_values_only['genotype'].isin(['BALB/c'])]['triglyceride_values']

# %% [code]
mannwhitneyu(tg_values_only[tg_values_only['genotype'].isin(['BALB/c'])]['triglyceride_values'],tg_values_only[tg_values_only['genotype'].isin(['C57BL6/J'])]['triglyceride_values'])

# %% [code]
mannwhitneyu(tg_values_only[tg_values_only['genotype'].isin(['BALB/c'])]['triglyceride_values'],tg_values_only[tg_values_only['genotype'].isin(['C57BL6/J'])]['triglyceride_values'])

# %% [code]
F1_tg_C57B = tg_values_only[(tg_values_only['generation'].isin(['F1'])) & (tg_values_only['genotype'].isin(['C57BL6/J']))]['triglyceride_values']
F2_tg_C57B = tg_values_only[(tg_values_only['generation'].isin(['F2'])) & (tg_values_only['genotype'].isin(['C57BL6/J']))]['triglyceride_values']
F3_tg_C57B = tg_values_only[(tg_values_only['generation'].isin(['F3'])) & (tg_values_only['genotype'].isin(['C57BL6/J']))]['triglyceride_values']
F4_tg_C57B = tg_values_only[(tg_values_only['generation'].isin(['F4'])) & (tg_values_only['genotype'].isin(['C57BL6/J']))]['triglyceride_values']
F5_tg_C57B = tg_values_only[(tg_values_only['generation'].isin(['F5'])) & (tg_values_only['genotype'].isin(['C57BL6/J']))]['triglyceride_values']
P1_tg_C57B = tg_values_only[(tg_values_only['generation'].isin(['P1'])) & (tg_values_only['genotype'].isin(['C57BL6/J']))]['triglyceride_values']
#RM_tg_C57B = tg_values_only[(tg_values_only['generation'].isin(['RM'])) & (tg_values_only['genotype'].isin(['C57BL6/J']))]['triglyceride_values']

tg_C57B_groups = list(itertools.combinations([P1_tg_C57B, F1_tg_C57B, F2_tg_C57B, F3_tg_C57B, F4_tg_C57B, F5_tg_C57B],2))
tg_C57B_groups_str = list(itertools.combinations(['P1_tg_C57B', 'F1_tg_C57B', 'F2_tg_C57B', 'F3_tg_C57B', 'F4_tg_C57B', 'F5_tg_C57B'],2))

# %% [code]
F1_tg_BALB = tg_values_only[(tg_values_only['generation'].isin(['F1'])) & (tg_values_only['genotype'].isin(['BALB/c']))]['triglyceride_values']
F2_tg_BALB = tg_values_only[(tg_values_only['generation'].isin(['F2'])) & (tg_values_only['genotype'].isin(['BALB/c']))]['triglyceride_values']
F3_tg_BALB = tg_values_only[(tg_values_only['generation'].isin(['F3'])) & (tg_values_only['genotype'].isin(['BALB/c']))]['triglyceride_values']
F4_tg_BALB = tg_values_only[(tg_values_only['generation'].isin(['F4'])) & (tg_values_only['genotype'].isin(['BALB/c']))]['triglyceride_values']
F5_tg_BALB = tg_values_only[(tg_values_only['generation'].isin(['F5'])) & (tg_values_only['genotype'].isin(['BALB/c']))]['triglyceride_values']
P1_tg_BALB = tg_values_only[(tg_values_only['generation'].isin(['P1'])) & (tg_values_only['genotype'].isin(['BALB/c']))]['triglyceride_values']
#RM_tg_BALB = tg_values_only[(tg_values_only['generation'].isin(['RM'])) & (tg_values_only['genotype'].isin(['BALB/c']))]['triglyceride_values']

tg_BALB_groups = list(itertools.combinations([P1_tg_BALB, F1_tg_BALB, F2_tg_BALB, F3_tg_BALB, F4_tg_BALB, F5_tg_BALB],2))
tg_BALB_groups_str = list(itertools.combinations(['P1_tg_BALB', 'F1_tg_BALB', 'F2_tg_BALB', 'F3_tg_BALB', 'F4_tg_BALB', 'F5_tg_BALB'],2))

# %% [code]
alpha = 0.05/len(tg_C57B_groups)
count = 0
statistics = []
p_values = []
significant = []
groups = []
bonferroni = []
zerofive = []
for pair in tg_C57B_groups:
    mannwhitneyu_table(pair, tg_C57B_groups_str, count)
    count +=1

# %% [code]
alpha_results_cage_group_vertical_C57B_df = pd.DataFrame({'Comparisons': groups,
                                'Statistics': statistics,
                                'p_values': p_values,
                                'Alpha': alpha,
                                'Reject H0: x<=y?': bonferroni,
                                'No Bonferroni reject H0': zerofive})
print('Results of vertical comparisons of cage-groups')
alpha_results_cage_group_vertical_C57B_df

# %% [code]


# %% [code]
alpha = 0.05/len(tg_BALB_groups)
count = 0
statistics = []
p_values = []
significant = []
groups = []
bonferroni = []
zerofive = []
for pair in tg_BALB_groups:
    mannwhitneyu_table(pair, tg_BALB_groups_str, count)
    count +=1

# %% [code]
alpha_results_cage_group_vertical_BALB_df = pd.DataFrame({'Comparisons': groups,
                                'Statistics': statistics,
                                'p_values': p_values,
                                'Alpha': alpha,
                                'Reject H0: x<=y?': bonferroni,
                                'No Bonferroni reject H0': zerofive})
print('Results of vertical comparisons of cage-groups')
alpha_results_cage_group_vertical_BALB_df

# %% [code]
T = tg_values_only.transpose()
#T.columns
T['13422.1'].fillna(0).value_counts().shape[0]

# %% [code]
res_tg = {
    'alpha_diversity': alpha_diversity(tg_values_only.transpose(), metrics=['shannon'], dry=False, wait=True, rarefaction_depth=None)
    #,    'beta_diversity': beta_diversity(tg_values_only.transpose(), metrics=['bray_curtis'], dry=False, wait=True)
}

# %% [code]
! rm -r /homes/arehm/TMP/ana_adiv_ia_ydsfo

# %% [code]
```
