## Additional File 2: Analysis repository for "The host genotype actively shapes its microbiome across generations in laboratory mice": benga_threegen_analysis.ipynb

```
# %% [markdown]
<h1>Table of Contents<span class="tocSkip"></span></h1>
<div class="toc"><ul class="toc-item"><li><span><a href="#Data-Processing" data-toc-modified-id="Data-Processing-1"><span class="toc-item-num">1&nbsp;&nbsp;</span>Data Processing</a></span><ul class="toc-item"><li><span><a href="#Metadata-correlation" data-toc-modified-id="Metadata-correlation-1.1"><span class="toc-item-num">1.1&nbsp;&nbsp;</span>Metadata correlation</a></span></li></ul></li><li><span><a href="#Time-Course" data-toc-modified-id="Time-Course-2"><span class="toc-item-num">2&nbsp;&nbsp;</span>Time Course</a></span></li><li><span><a href="#Metabolites" data-toc-modified-id="Metabolites-3"><span class="toc-item-num">3&nbsp;&nbsp;</span>Metabolites</a></span><ul class="toc-item"><li><span><a href="#Serum-Trigylceride" data-toc-modified-id="Serum-Trigylceride-3.1"><span class="toc-item-num">3.1&nbsp;&nbsp;</span>Serum Trigylceride</a></span></li><li><span><a href="#genotype-across-generations" data-toc-modified-id="genotype-across-generations-3.2"><span class="toc-item-num">3.2&nbsp;&nbsp;</span>genotype across generations</a></span><ul class="toc-item"><li><span><a href="#genotype-stratefied-by-generation" data-toc-modified-id="genotype-stratefied-by-generation-3.2.1"><span class="toc-item-num">3.2.1&nbsp;&nbsp;</span>genotype stratefied by generation</a></span><ul class="toc-item"><li><span><a href="#genotype-in-F4" data-toc-modified-id="genotype-in-F4-3.2.1.1"><span class="toc-item-num">3.2.1.1&nbsp;&nbsp;</span>genotype in F4</a></span></li></ul></li></ul></li><li><span><a href="#generations-across-genotype" data-toc-modified-id="generations-across-genotype-3.3"><span class="toc-item-num">3.3&nbsp;&nbsp;</span>generations across genotype</a></span><ul class="toc-item"><li><span><a href="#generations-stratefied-by-genotype" data-toc-modified-id="generations-stratefied-by-genotype-3.3.1"><span class="toc-item-num">3.3.1&nbsp;&nbsp;</span>generations stratefied by genotype</a></span></li></ul></li></ul></li><li><span><a href="#Repeat-from-Scratch" data-toc-modified-id="Repeat-from-Scratch-4"><span class="toc-item-num">4&nbsp;&nbsp;</span>Repeat from Scratch</a></span><ul class="toc-item"><li><ul class="toc-item"><li><span><a href="#beta-dist-within-cage" data-toc-modified-id="beta-dist-within-cage-4.0.1"><span class="toc-item-num">4.0.1&nbsp;&nbsp;</span>beta dist within cage</a></span></li><li><span><a href="#beta-dist-within-cage-groups" data-toc-modified-id="beta-dist-within-cage-groups-4.0.2"><span class="toc-item-num">4.0.2&nbsp;&nbsp;</span>beta dist within cage groups</a></span></li></ul></li></ul></li><li><span><a href="#Eco-Filter?" data-toc-modified-id="Eco-Filter?-5"><span class="toc-item-num">5&nbsp;&nbsp;</span>Eco Filter?</a></span><ul class="toc-item"><li><span><a href="#add-host_subject_id,-parents,-cages-to-metadata-(only-run-once!)" data-toc-modified-id="add-host_subject_id,-parents,-cages-to-metadata-(only-run-once!)-5.1"><span class="toc-item-num">5.1&nbsp;&nbsp;</span>add host_subject_id, parents, cages to metadata (only run once!)</a></span></li><li><span><a href="#Cage-effect:-are-mice-of-same-cage-more-similar-than-mice-of-different-cages-in-same-cage_line/generation/genotype?---->YES" data-toc-modified-id="Cage-effect:-are-mice-of-same-cage-more-similar-than-mice-of-different-cages-in-same-cage_line/generation/genotype?---->YES-5.2"><span class="toc-item-num">5.2&nbsp;&nbsp;</span>Cage effect: are mice of same cage more similar than mice of different cages in same cage_line/generation/genotype? <br> --&gt;YES</a></span></li><li><span><a href="#Cage-lineage-effect:-are-mice-of-same-cage_lineage-more-similar-than-mice-of-different-cage_lines/generation/C57BL6/J?---->YES" data-toc-modified-id="Cage-lineage-effect:-are-mice-of-same-cage_lineage-more-similar-than-mice-of-different-cage_lines/generation/C57BL6/J?---->YES-5.3"><span class="toc-item-num">5.3&nbsp;&nbsp;</span>Cage lineage effect: are mice of same cage_lineage more similar than mice of different cage_lines/generation/C57BL6/J? <br> --&gt;YES</a></span></li><li><span><a href="#do-genotype-microbiomes-become-more-distant-over-time?---->-Yes,-but-only-up-to-f2" data-toc-modified-id="do-genotype-microbiomes-become-more-distant-over-time?---->-Yes,-but-only-up-to-f2-5.4"><span class="toc-item-num">5.4&nbsp;&nbsp;</span>do genotype microbiomes become more distant over time? <br> --&gt; Yes, but only up to f2</a></span><ul class="toc-item"><li><span><a href="#across-genotype" data-toc-modified-id="across-genotype-5.4.1"><span class="toc-item-num">5.4.1&nbsp;&nbsp;</span>across genotype</a></span></li><li><span><a href="#within-genotype" data-toc-modified-id="within-genotype-5.4.2"><span class="toc-item-num">5.4.2&nbsp;&nbsp;</span>within genotype</a></span></li></ul></li><li><span><a href="#ASVs" data-toc-modified-id="ASVs-5.5"><span class="toc-item-num">5.5&nbsp;&nbsp;</span>ASVs</a></span><ul class="toc-item"><li><span><a href="#Venn" data-toc-modified-id="Venn-5.5.1"><span class="toc-item-num">5.5.1&nbsp;&nbsp;</span>Venn</a></span></li></ul></li><li><span><a href="#what-is-the-most-prototypic-generation/cage_line-per-genotype?" data-toc-modified-id="what-is-the-most-prototypic-generation/cage_line-per-genotype?-5.6"><span class="toc-item-num">5.6&nbsp;&nbsp;</span>what is the most prototypic generation/cage_line per genotype?</a></span></li><li><span><a href="#RF-to-predict-genotype--->-83%,-i.e.-&quot;filtered-microbiom&quot;-is-not-prototypic" data-toc-modified-id="RF-to-predict-genotype--->-83%,-i.e.-&quot;filtered-microbiom&quot;-is-not-prototypic-5.7"><span class="toc-item-num">5.7&nbsp;&nbsp;</span>RF to predict genotype <br>--&gt; 83%, i.e. "filtered microbiom" is not prototypic</a></span></li></ul></li><li><span><a href="#Temporal-Summary" data-toc-modified-id="Temporal-Summary-6"><span class="toc-item-num">6&nbsp;&nbsp;</span>Temporal Summary</a></span><ul class="toc-item"><li><span><a href="#rarefie-mice" data-toc-modified-id="rarefie-mice-6.1"><span class="toc-item-num">6.1&nbsp;&nbsp;</span>rarefie mice</a></span></li></ul></li><li><span><a href="#Quantify-Stress" data-toc-modified-id="Quantify-Stress-7"><span class="toc-item-num">7&nbsp;&nbsp;</span>Quantify Stress</a></span><ul class="toc-item"><li><span><a href="#animals-per-cage--->-makes-a-difference!" data-toc-modified-id="animals-per-cage--->-makes-a-difference!-7.1"><span class="toc-item-num">7.1&nbsp;&nbsp;</span>animals per cage --&gt; makes a difference!</a></span><ul class="toc-item"><li><span><a href="#gut" data-toc-modified-id="gut-7.1.1"><span class="toc-item-num">7.1.1&nbsp;&nbsp;</span>gut</a></span></li><li><span><a href="#skin" data-toc-modified-id="skin-7.1.2"><span class="toc-item-num">7.1.2&nbsp;&nbsp;</span>skin</a></span></li></ul></li><li><span><a href="#birth-offset" data-toc-modified-id="birth-offset-7.2"><span class="toc-item-num">7.2&nbsp;&nbsp;</span>birth offset</a></span><ul class="toc-item"><li><span><a href="#are-cousins-born-on-same-day-closer-to-each-other-than-cousins-born-on-different-days?--->-yes,-but-only-in-unweighted-UniFrac" data-toc-modified-id="are-cousins-born-on-same-day-closer-to-each-other-than-cousins-born-on-different-days?--->-yes,-but-only-in-unweighted-UniFrac-7.2.1"><span class="toc-item-num">7.2.1&nbsp;&nbsp;</span>are cousins born on same day closer to each other than cousins born on different days? --&gt; yes, but only in unweighted UniFrac</a></span></li></ul></li><li><span><a href="#mothers-deliver-in-sync--->-no-difference" data-toc-modified-id="mothers-deliver-in-sync--->-no-difference-7.3"><span class="toc-item-num">7.3&nbsp;&nbsp;</span>mothers deliver in sync --&gt; no difference</a></span></li></ul></li><li><span><a href="#food-/-beeding-as-taxa-sources" data-toc-modified-id="food-/-beeding-as-taxa-sources-8"><span class="toc-item-num">8&nbsp;&nbsp;</span>food / beeding as taxa sources</a></span></li><li><span><a href="#maternal-legacy" data-toc-modified-id="maternal-legacy-9"><span class="toc-item-num">9&nbsp;&nbsp;</span>maternal legacy</a></span><ul class="toc-item"><li><span><a href="#gut" data-toc-modified-id="gut-9.1"><span class="toc-item-num">9.1&nbsp;&nbsp;</span>gut</a></span></li><li><span><a href="#skin" data-toc-modified-id="skin-9.2"><span class="toc-item-num">9.2&nbsp;&nbsp;</span>skin</a></span></li></ul></li><li><span><a href="#Main-Text-Statement" data-toc-modified-id="Main-Text-Statement-10"><span class="toc-item-num">10&nbsp;&nbsp;</span>Main Text Statement</a></span><ul class="toc-item"><li><span><a href="#for-skin" data-toc-modified-id="for-skin-10.1"><span class="toc-item-num">10.1&nbsp;&nbsp;</span>for skin</a></span></li></ul></li><li><span><a href="#Meta:-Robertsen" data-toc-modified-id="Meta:-Robertsen-11"><span class="toc-item-num">11&nbsp;&nbsp;</span>Meta: Robertsen</a></span></li><li><span><a href="#Source-tracking-skin" data-toc-modified-id="Source-tracking-skin-12"><span class="toc-item-num">12&nbsp;&nbsp;</span>Source tracking skin</a></span><ul class="toc-item"><li><span><a href="#prototypic-gut-microbiome-+-food/bedding" data-toc-modified-id="prototypic-gut-microbiome-+-food/bedding-12.1"><span class="toc-item-num">12.1&nbsp;&nbsp;</span>prototypic gut microbiome + food/bedding</a></span></li><li><span><a href="#gut-seeds-skin?" data-toc-modified-id="gut-seeds-skin?-12.2"><span class="toc-item-num">12.2&nbsp;&nbsp;</span>gut seeds skin?</a></span><ul class="toc-item"><li><span><a href="#per-generation,-i.e.-cross-genotypes" data-toc-modified-id="per-generation,-i.e.-cross-genotypes-12.2.1"><span class="toc-item-num">12.2.1&nbsp;&nbsp;</span>per generation, i.e. cross genotypes</a></span></li><li><span><a href="#per-cage-lineage" data-toc-modified-id="per-cage-lineage-12.2.2"><span class="toc-item-num">12.2.2&nbsp;&nbsp;</span>per cage lineage</a></span></li></ul></li></ul></li></ul></div>

# %% [code]
%matplotlib inline
%load_ext autoreload
%autoreload 2

import pandas as pd
from skbio.tree import TreeNode

from ggmap.snippets import *
from ggmap.analyses import *
from ggmap.correlations import *
from ggmap.workflow import *
from statannot import add_stat_annotation
from statannotations.Annotator import Annotator
from skbio.stats.distance import MissingIDError
from matplotlib_venn import venn3, venn3_circles, venn3_unweighted

import calour as ca
ca.set_log_level(40)

def alpha_blending(hex_color, alpha) :
    """ alpha blending as if on the white background.
    """
    foreground_tuple  = matplotlib.colors.hex2color(hex_color)
    foreground_arr = np.array(foreground_tuple)
    final = tuple( (1. -  alpha) + foreground_arr*alpha )
    return(final)

# %% [code]
fp_workdir = 'Workdir/'
fp_generated = 'Generated/'
for fp in [fp_workdir, fp_generated]:
    os.makedirs(fp, exist_ok=True)

fps = {'meta': {
    '5gen': 'meta_swap.csv',
               },
       'prep': {'5gen': '../FromQiita/13422_prep_9771_20201201-075435.txt',
               },
       'deblur_150': {'5gen': '../FromQiita/102006_reference-hit.biom',
                     },
       'closedref_150': {'5gen': '../FromQiita/102007_otu_table.biom'},
       'workdir': {
           'combined_bioms_deblur':          'deblur_all.biom',
           'combined_bioms_closedref':       'closedref_all.biom',
           'insertion_tree':                 'insertion_tree_all.newick',
        },
       
      }
assert len(set(fps['workdir'].values())) == len(list(fps['workdir'].values())), 'two workdir paths are identical'
for f in fps['workdir']:
    fps['workdir'][f] = fp_workdir + fps['workdir'][f]

na_values=['not applicable', 'not provided', 'missing', 'not collected']
    
def _savefig(fig, name):
    os.makedirs(fp_generated+'Figures/SVG', exist_ok=True)
    os.makedirs(fp_generated+'Figures/PNG', exist_ok=True)
    
    fig.savefig(fp_generated+'Figures/SVG/%s.svg' % name, bbox_inches='tight')
    fig.savefig(fp_generated+'Figures/PNG/%s.png' % name, bbox_inches='tight')

# %% [code]
meta = pd.concat([
    # read metadata
    pd.concat([pd.read_csv(file, sep="\t", dtype=str, na_values=na_values).set_index('sample_name') for file in fps['meta'].values()], axis=0, sort=False),
    # read prep data
    pd.concat([pd.read_csv(file, sep="\t", dtype=str, na_values=na_values).set_index('sample_name') for file in fps['prep'].values()], axis=0, sort=False).rename(columns={'description': 'description_prep'})
], sort=False, axis=1)

# normalize date formats
for c in ['birth_timestamp',
          'collection_timestamp',
          ]:
    meta[c] = pd.to_datetime(meta[c])

# normalize float data
for c in ['elevation',
          'host_age',
          'host_weight',
          'latitude',
          'longitude',
          ]:
    meta[c] = meta[c].astype(float)
    
meta['timepoint'] = meta['generation'].apply(lambda x: {'RM': 0,
                                    'P1': 1,
                                    'F1': 2,
                                    'F2': 3,
                                    'F3': 4,
                                    'F4': 5,
                                    'F5': 6,
                                   }.get(x, x))
meta['i_generation'] = meta['generation'].apply(lambda x: {'RM': '0: RM',
                                    'P1': '1: P1',
                                    'F1': '2: F1',
                                    'F2': '3: F2',
                                    'F3': '4: F3',
                                    'F4': '5: F4',
                                    'F5': '6: F5',
                                   }.get(x, x))

# %% [code]
# load deblur feature data
counts_deblur = pd.concat([biom2pandas(file) for file in fps['deblur_150'].values()], axis=1, sort=False).fillna(0)

# load closedref feature data
counts_closedref = pd.concat([biom2pandas(file) for file in fps['closedref_150'].values()], axis=1, sort=False).fillna(0)

if not os.path.exists(fps['workdir']['combined_bioms_deblur']):
    pandas2biom(fps['workdir']['combined_bioms_deblur'], counts_deblur)
if not os.path.exists(fps['workdir']['combined_bioms_closedref']):
    pandas2biom(fps['workdir']['combined_bioms_closedref'], counts_closedref)

# create one shared insertion tree
res_sepp = sepp(counts_deblur, chunksize=3000, ppn=10, dry=False, wait=False)
if not os.path.exists(fps['workdir']['insertion_tree']):
    with open(fps['workdir']['insertion_tree'], 'w') as f:
        f.write(res_sepp['results']['tree'].replace('k__Bacteria)', 'k__Bacteria:0.0)')+"\n")
%time insertion_tree = TreeNode.read(fps['workdir']['insertion_tree'])

# %% [code]
# hier ein erster Aufruf eines meiner Q2 Wrapper. Schau doch mal, ob Du den ausführen kannst. 
# Es wird noch nichts gerechnet, sondern nur das Kommando ausgespruckt, dass den Job als Cluster-Job abschicken würde.
taxonomy_RDP(counts_deblur.iloc[:10,:], '/homes/sjanssen/MicrobiomeAnalyses/References/Q2-Naive_Bayes_classifiers/gg-13-8-99-nb-classifier_2021.11.qza', dry=True)

# %% [code]
# wenn obiges klappt, kannst Du "dry" auf False setzen und gucken, ob der Job nun wirklich im grid ausgeführt wird
taxonomy_RDP(counts_deblur.iloc[:11,:], '/homes/sjanssen/MicrobiomeAnalyses/References/Q2-Naive_Bayes_classifiers/gg-13-8-99-nb-classifier_2021.11.qza', dry=False)

# %% [markdown]
# Data Processing

# %% [code]
# sollten die beiden oberen taxonomy_RDP Aufrufe klappen, kannst Du Dich an das Monster process_study wagen und gucken, ob auch das etwas rechnet

# %% [code]
res_all = process_study(
    metadata=meta,
    control_samples=set(meta[meta['sample_type'].isin(['sterile water'])].index),
    fp_deblur_biom=fps['workdir']['combined_bioms_deblur'],
    fp_insertiontree=fps['workdir']['insertion_tree'],
    fp_closedref_biom=None, #fps['workdir']['combined_bioms_closedref'],
    rarefaction_depth=6000,
    fp_taxonomy_trained_classifier='/homes/sjanssen/MicrobiomeAnalyses/References/Q2-Naive_Bayes_classifiers/gg-13-8-99-nb-classifier_2021.11.qza',
    tree_insert=insertion_tree,
    is_v4_region=False,
    fix_zero_len_branches=False,
    emperor_fp=fp_generated+'/Emperor/',
    emperor_infix='_all_',
    dry=False,
    use_grid=True,
    #deblur_remove_features_lessthanXreads=10, # as this is now the default
)

print('Overall, the study recovered %s sOTUs (from %s samples)' % res_all['rarefaction']['results'].shape)

# %% [code]
emperor(meta, res_all['beta_diversity']['results'], fp_generated+'/Emperor2/', run_tsne_umap=True, dry=False, nocache=True)

# %% [code]
ganttChart(meta, col_birth='birth_timestamp', col_entities='host_subject_id', col_events='collection_timestamp', col_death='collection_timestamp')[0]

# %% [markdown]
## Metadata correlation

# %% [code]
mouse_meta = meta[meta['host_common_name'] == 'mouse']

cols_unique = {c for c in mouse_meta.columns if mouse_meta[c].dropna().unique().shape[0] == 1}
cols_alldiff = {c for c in mouse_meta.columns if mouse_meta[c].dropna().unique().shape[0] == mouse_meta[c].dropna().shape[0]}

cols_categorial = {'i7_index_id', 'sex', 'barcode', 'description', 'parental_cage', 'i7_index', 'plate_column', 'plate_number', 'genotype', 'plate_row', 'iacuc_protocol_id',
                   'i5_index', 'i5_index_id', 'tube_id', 'cage_id'}
cols_ordinal = {'father_subject_id': ['P1', 'F1', 'F2', 'F3', 'F4', 'fatherF4'],
                'mother_subject_id': ['P4', 'P3', 'P2', 'P1', 'recipient', 'F1', 'F2', 'F3', 'F4'],
                'generation': ['P1', 'RM', 'F1', 'F2', 'F3', 'F4', 'F5'],
               }
cols_interval = ['host_weight', 'host_age']
cols_date = {'collection_timestamp': 'yyyy-mm-dd hh:mm:ss', 'birth_timestamp': 'yyyy-mm-dd hh:mm:ss'}


res_corr = correlation_diversity_metacolumns(
    mouse_meta, categorial=cols_categorial, 
    alpha_diversities=res_all['alpha_diversity']['results'], beta_diversities=res_all['beta_diversity']['results'], dry=False, wait=False)
cols_significant = {c for c,p in res_corr['results'].groupby('column')['p-value'].min().sort_values().iteritems() if p < 0.05}
cols_categorial &= cols_significant
cols_ordinal = {k:v for k,v in cols_ordinal.items() if k in cols_significant}
cols_interval = list(set(cols_interval) & cols_significant)
cols_date = {k:v for k,v in cols_date.items() if k in cols_significant}

corr = correlate_metadata(mouse_meta, categorials=list(cols_categorial), ordinals=cols_ordinal, intervals=cols_interval, dates=cols_date)
#mouse_meta[set(mouse_meta.columns) - cols_unique - cols_alldiff - cols_categorial - set(cols_intervals) - set(cols_date.keys())].T
#mouse_meta['generation'].value_counts()

# %% [code]


# %% [markdown]
# Time Course

# %% [code]
metric = 'PD_whole_tree'
fig, axes = plt.subplots(1,1, figsize=(10,5))
meta_data = plot_timecourse(meta[(meta['host_common_name'] == 'mouse') & meta['genotype'].isin(['C57BL6/J','BALB/c'])],
                data=res_all['alpha_diversity']['results'][metric],
                col_events='i_generation', col_entities=None,
                cols_groups=['host_common_name', 'genotype'],
                test_groups=[(('mouse', 'C57BL6/J'), ('mouse', 'BALB/c'))],
                            #[(('Pax5+/-', 'no'), ('Pax5+/-', 'pB-ALL'))]
                pthreshold=0.05, print_samplesizes=False,
                ax=axes)

# %% [code]
res_all['alpha_diversity']['results'][metric]

# %% [code]
for i, metric in enumerate(res_all['beta_diversity']['results'].keys()):
    fig, axes = plt.subplots(1, 1, figsize=(10,5))
    #metric = 'unweighted_unifrac'
    data = []
    for (i_generation, genotype), g in meta[(meta['host_common_name'] == 'mouse') & meta['genotype'].isin(['C57BL6/J','BALB/c'])].groupby(['i_generation', 'genotype']):
        dm = res_all['beta_diversity']['results'][metric].filter(g.index, strict=False)
        for a,b in combinations(dm.ids, 2):
            data.append({'i_generation': i_generation, 'genotype': genotype, metric: dm[a,b], 'host_common_name': g['host_common_name'].iloc[0]})
    
    for i_generation, g in meta[(meta['host_common_name'] == 'mouse') & meta['genotype'].isin(['C57BL6/J','BALB/c'])].groupby(['i_generation']):
        dm = res_all['beta_diversity']['results'][metric].filter(g.index, strict=False)
        for a in g[g['genotype'] == 'C57BL6/J'].index:
            for b in g[g['genotype'] != 'C57BL6/J'].index:
                if (a in dm.ids) and (b in dm.ids):
                    data.append({'i_generation': i_generation, 'genotype': 'inter', metric: dm[a,b], 'host_common_name': g['host_common_name'].iloc[0]})
    data = pd.DataFrame(data)

    meta_data = plot_timecourse(
        data[[c for c in data.columns if c != metric]],
        data=data[metric],
        col_events='i_generation', col_entities=None,
        cols_groups=['host_common_name', 'genotype'],
        print_samplesizes=True,
        ax=axes)
    plt.figure()

# %% [code]
meta = pd.read_csv('../Incoming/13422_20210909-070806.txt', sep="\t", index_col=0, dtype=str)
meta[meta['host_body_habitat'] == "UBERON:gastrointestinal system"].fillna('?').groupby(['generation']).size()

# %% [markdown]
# Metabolites

# %% [code]
### load GC-MS
fp_metabolites = '../Incoming/GC_results/220511__Data_21-0025.xlsx'
x = pd.read_excel(fp_metabolites, sheet_name="01.AllData").rename(columns={'Sample.label.on.tube': 'sample_name'})
x['sample_name'] = 'sample.' + x['sample_name'].astype(str)
data = x[['sample_name', "ISTD", "Resp", "ISTD.Resp", "Resp.Ratio", "UserAnno.Peak", "MI.Peak", "MZ", "MZ.Ex.Left", "MZ.Ex.Right", "MZ.Ex.Units", "RT.Compound", "RT.Peak", "MS.Replicates", "data.source", "ISTD.Resp.Ratio", "Acq.Method.Condensed", "MS.project", "Report.Name", "data.dir", "relativeResponse", "IQR_1.5_outlier", "Compound"]].set_index(['sample_name', 'Compound'])
counts_chems = pd.pivot_table(data=data, columns='sample_name', index="Compound", values="relativeResponse").fillna(0)
# remove compounds containing 'unknown' in its name
counts_chems = counts_chems.loc[[c for c in counts_chems.index if 'unknown' not in c], :]

# load metadata from qiita
meta_qiita = pd.read_csv('../FromQiita/13422_20230201-071657.txt', sep="\t", index_col=0, dtype=str).groupby(['host_subject_id']).head(1)
meta_qiita

# also load metadata from Excel sheet that Philipp Westhoff provided
m = pd.read_excel(fp_metabolites, sheet_name="01.AllData").rename(columns={'Sample.label.on.tube': 'sample_name'})
m['sample_name'] = 'sample.' + m['sample_name'].astype(str)
meta_chems = m[['sample_name', "date", "MS.Label", "Species", "Tissue.type", "Condition.1", "Generation", "Benga-Nr."]].set_index(['sample_name']).drop_duplicates().rename(columns={'Condition.1': 'genotype', 'Generation': 'generation'})
meta_chems['Benga-Nr.'] = meta_chems['Benga-Nr.'].astype(str)

# merge metadata from Philipp and Qiita
meta_chems = meta_chems.reset_index().merge(meta_qiita, left_on='Benga-Nr.', right_on='host_subject_id', how='left').set_index('sample_name')
meta_chems

# %% [code]
res_all_chems = {
    'alpha_diversity': alpha_diversity(counts_chems, metrics=['shannon'], dry=False, wait=False, rarefaction_depth=None),
    'beta_diversity': beta_diversity(counts_chems, metrics=['bray_curtis'], dry=False, wait=False)
}

dsFDR_experiment = dict()
# for dsFDR
pandas2biom('tmp_dsfrd_metabolome.biom', counts_chems)
meta_chems.to_csv('tmp_dsfdr_metabolome_meta.tsv', sep="\t", index=True)

dsFDR_experiment = ca.read_amplicon('tmp_dsfrd_metabolome.biom', 
                                    'tmp_dsfdr_metabolome_meta.tsv', min_reads=0, normalize=10000)

# %% [code]
emperor(meta_chems, res_all_chems["beta_diversity"]['results'], fp_results='Generated/Emperor/', infix='_metabolites', run_tsne_umap=True, dry=False, wait=True, dirty=True)

# %% [markdown]
## Serum Trigylceride

# %% [code]
m = pd.read_csv("../FromQiita/13422_20230704-030402.txt", sep="\t", dtype=str, na_values=['not applicable', 'not collected'], index_col=0)
t = pd.read_excel('manual_triglycerates.xlsx', index_col=0)
t.index = list(map(lambda x: x.replace('f', '').replace('e', ''), t.index))
mt = m.merge(t, left_on='host_subject_id', right_index=True, how='left')
m.shape, t.shape, mt.shape

#mt.to_excel("help.xlsx")

# %% [code]
m = pd.read_csv("../FromQiita/13422_20230704-030402.txt", sep="\t", dtype=str, index_col=0)
t = pd.read_excel('manual_triglycerates.xlsx', index_col=0)
t.index = list(map(lambda x: x.replace('f', '').replace('e', ''), t.index))
#m.merge(t[['serum_triglyceride','serum_triglyceride_unit']], left_on='host_subject_id', right_index=True, how='left').to_csv('new_meta.csv', sep="\t", index=True)


# %% [code]
prj_data['metadata']['host_scientific_name'].value_counts()

# %% [code]
# no monkeys, one value per host
sns.set_style("white")
data = prj_data['metadata'][prj_data['metadata']['host_scientific_name'] != 'callithrix jacchus'].groupby('host_subject_id').head(1)
fig, axes = plt.subplots(1,1,figsize=(5,3))
sns.boxplot(data=data, x='serum_triglyceride', y='genotype', orient='horizontal', palette=colors, ax=axes, order=['b6cf1','c57bl6/j','balb/c'])

annotator = Annotator(axes, 
    [('balb/c', 'c57bl6/j'), ('b6cf1', 'c57bl6/j')], 
    data=data, x='serum_triglyceride', y='genotype', orient='h', order=['b6cf1','c57bl6/j','balb/c'])
annotator.configure(test='Mann-Whitney', text_format='star', loc='outside', comparisons_correction="fdr_bh", correction_format="default")
_ = annotator.apply_and_annotate()

fig.savefig('Generated/triglycerate.svg')

# %% [markdown]
## genotype across generations

# %% [code]
x = plotNetworks('genotype_x', meta_chems, 
                 alpha=res_all_chems["alpha_diversity"]['results'], 
                 beta=res_all_chems["beta_diversity"]['results'], minnumalpha=1)

# %% [code]
field = 'genotype_x'
b = detect_distant_groups(
                    res_all_chems["beta_diversity"]['results']['bray_curtis'], 'bray_curtis', meta_chems[field],)
fig, axes = plt.subplots(1,1,figsize=(5,3))
g = plotGroup_permanovas(res_all_chems["beta_diversity"]['results']['bray_curtis'], meta_chems[field], **b, horizontal=True, ax=axes)
fig.savefig("Figures/Metabolites/beta_%s.svg" % field)

# %% [code]
field = 'generation_x'
b = detect_distant_groups(
                    res_all_chems["beta_diversity"]['results']['bray_curtis'], 'bray_curtis', meta_chems[field],)
fig, axes = plt.subplots(1,1,figsize=(5,3))
g = plotGroup_permanovas(res_all_chems["beta_diversity"]['results']['bray_curtis'], meta_chems[field], **b, horizontal=True, ax=axes)
fig.savefig("Figures/Metabolites/beta_%s.svg" % field)

# %% [markdown]
### genotype stratefied by generation

# %% [code]
for generation, g in meta_chems.groupby('generation_x'):
    _ = plotNetworks('genotype_x', g, 
                     alpha=res_all_chems["alpha_diversity"]['results'], 
                     beta=res_all_chems["beta_diversity"]['results'], minnumalpha=1, name=generation)

# %% [markdown]
#### genotype in F4

# %% [code]
field = 'genotype_x'
res_dsFDR_genotype = find_diff_taxa(dsFDR_experiment, meta_chems[meta_chems['generation_x'] == 'F4'], [field])

if len(res_dsFDR_genotype) > 0:
    p = plot_diff_taxa(counts_chems, meta_chems[meta_chems['generation_x'] == 'F4'][field], res_dsFDR_genotype,
                       min_mean_abundance=0.0,
#                       colors_boxplot={'Pax5+/-': 'red', 'WT': 'blue'}, 
                       color_barplot='gray'
                      )
    p[0].get_axes()[0].set_ylabel("compound")
    p[0].set_size_inches((12,3))
    p[0].savefig('Figures/Metabolites/dsFDR_genotype.svg')

# %% [markdown]
## generations across genotype

# %% [code]
_ = plotNetworks('generation_x', meta_chems, 
                 alpha=res_all_chems["alpha_diversity"]['results'], 
                 beta=res_all_chems["beta_diversity"]['results'], minnumalpha=1)

# %% [markdown]
### generations stratefied by genotype

# %% [code]
for genotype, g in meta_chems.groupby('genotype_x'):
    _ = plotNetworks('generation_x', g, 
                     alpha=res_all_chems["alpha_diversity"]['results'], 
                     beta=res_all_chems["beta_diversity"]['results'], minnumalpha=1, name=genotype)

# %% [markdown]
# Repeat from Scratch

# %% [markdown]
  - distance (weighted unifrac) of mice within the same cage is ~0.09
    - does not change over generations (outlyer in F4)

# %% [code]
prj_data = dict()

#import Prep- and Samplefile form Qiita
prep_file = pd.read_csv("../FromQiita/13422_prep_11649_20220103-015657.txt", sep="\t", dtype=str, na_values=['not applicable', 'not collected'], index_col=0)
sample_file = pd.read_csv("../FromQiita/13422_20230804-065447.txt", sep="\t", dtype=str, na_values=['not applicable', 'not collected'], index_col=0)
#sample_file =pd.read_csv("new_meta.tsv", sep="\t", dtype=str, na_values=['not applicable', 'not collected'], index_col=0)
#merge them to the metadata file
#metadata = pd.merge(sample_file, prep_file, right_index=True, left_index=True, how ='inner')
prj_data['metadata'] = pd.concat([sample_file, prep_file], axis=1)
assert pd.Series(prj_data['metadata'].columns).value_counts().max() == 1
#del prep_file, sample_file

gmap = {'f3':3, 'f4':4, 'f2':2, 'f5':5, 'f1':1, 'p1': 0}
prj_data['metadata']['timepoint'] = prj_data['metadata']['generation'].apply(lambda x: gmap.get(x, np.nan))

prj_data['metadata']['cl'] = prj_data['metadata']['cage_line'].apply(lambda x: x if type(x) == float else {'1': 'a', '4': 'a',
                                                                                '2': 'b', '6': 'b',
                                                                                '3': 'c', '5': 'c'}.get(x[-1], x) )
prj_data['metadata']['serum_triglyceride'] = prj_data['metadata']['serum_triglyceride'].astype(float)

prj_data['metadata'].head()

# %% [code]
prj_data['paths'] = {'deblur_table': '../FromQiita/128106_reference-hit.biom',
                     'insertion_tree': '../FromQiita/128106_insertion_tree.relabelled.tre',
                     'Generated/Emperor': 'Generated/Emperor'}
prj_data['insertion_tree'] = TreeNode.read(prj_data['paths']['insertion_tree'], format="newick")

# %% [code]
idx_control_samples = set(prj_data['metadata'][prj_data['metadata']['sample_type'].isin(['sterile water', 'control bedding', 'control food'])].index)
res_all = process_study(
    metadata=prj_data['metadata'],
    control_samples=idx_control_samples,
    fp_deblur_biom=prj_data['paths']['deblur_table'],
    fp_insertiontree=prj_data['paths']['insertion_tree'],
    rarefaction_depth=6000,
    fp_taxonomy_trained_classifier="/vol/jlab/MicrobiomeAnalyses/References/Q2-Naive_Bayes_classifiers/gg_2022_10_backbone_full_length.nb_2023.5.qza",
    tree_insert=prj_data['insertion_tree'],
    is_v4_region=False,
    emperor_fp=prj_data['paths']['Generated/Emperor'],
    emperor_infix='_deblur_',
    dry=False,
    #use_grid=True,
    use_grid=True, ppn=4,
    fix_zero_len_branches=False, # since this has already been corrected via the project_sepp call
)

# %% [code]
_ = rarefaction_curves(
    res_all['counts_plantsremoved'], 
    reference_tree=prj_data['paths']['insertion_tree'], 
    control_sample_names=idx_control_samples, 
    fix_zero_len_branches=False, # since this has already been corrected via the project_sepp call
    sample_grouping=prj_data["metadata"]['sample_type'], 
    dry=False)

# %% [code]
idx_control_samples = set(prj_data['metadata'][prj_data['metadata']['sample_type'].isin(['sterile water', 'control bedding', 'control food'])].index)
res_all_skin = process_study(
    metadata=prj_data['metadata'],
    control_samples=idx_control_samples,
    fp_deblur_biom=prj_data['paths']['deblur_table'],
    fp_insertiontree=prj_data['paths']['insertion_tree'],
    rarefaction_depth=1000,
    fp_taxonomy_trained_classifier="/vol/jlab/MicrobiomeAnalyses/References/Q2-Naive_Bayes_classifiers/gg_2022_10_backbone_full_length.nb_2023.5.qza",
    tree_insert=prj_data['insertion_tree'],
    is_v4_region=False,
    emperor_fp=prj_data['paths']['Generated/Emperor'],
    emperor_infix='_deblur_skin_',
    dry=False,
    #use_grid=True,
    use_grid=True, ppn=4,
    fix_zero_len_branches=False, # since this has already been corrected via the project_sepp call
)

# %% [markdown]
### beta dist within cage

# %% [code]
generation = 'f1'
metric = 'weighted_unifrac'
# dist per cage
dists_withincage = []
for cage, g in meta[meta['genotype'] != 'b6cf1'].groupby(['cage_id']):
    for (a,b) in combinations(g.index, 2):
        dists_withincage.append({'dist': res_all['beta_diversity']['results'][metric][a,b], 
                                 'genotype_a': g.loc[a,'genotype'],
                                 'genotype_b': g.loc[b,'genotype'],
                                 'cage_id': cage,
                                 'generation': g.loc[a, 'generation']})
        #print(cage, a,b)
dists_withincage = pd.DataFrame(dists_withincage)

#dists_withincage = dists_withincage[dists_withincage['genotype_a'] == 'c57bl6/j']
dists_withincage

# %% [code]
fix, axes = plt.subplots(1,1)
sns.boxplot(data=dists_withincage, y='dist', x='generation', order=['p1','f1','f2','f3','f4','f5'], ax=axes)
sns.swarmplot(data=dists_withincage, y='dist', x='generation', order=['p1','f1','f2','f3','f4','f5'], ax=axes, color='darkgray')

annotator = Annotator(axes, [
    ('p1','f1'),
    ('f1','f2'),
    ('f2','f3'),
    ('f3','f4'),
    ('f4','f5'),
], data=dists_withincage, y='dist', x='generation', order=['p1','f1','f2','f3','f4','f5'])
annotator.configure(test='Mann-Whitney', text_format='star', loc='outside', comparisons_correction="fdr_bh", correction_format="default")
annotator.apply_and_annotate()

axes.set_ylim((0, 0.3))

# %% [code]
dists_withincage['dist'].mean()

# %% [markdown]
### beta dist within cage groups

# %% [code]
generation = 'f1'
metric = 'unweighted_unifrac'
# dist per cage
dists_cg_within = []
dists_cg_between = []
for generation in meta[meta['genotype'] != 'b6cf1']['generation'].unique():
    for cagegrp, g in meta[(meta['generation'] == generation)].groupby(['cage_group']):
        for (a,b) in combinations(g.index, 2):
            dists_cg_within.append({'dist': res_all['beta_diversity']['results'][metric][a,b], 
                          'genotype_a': g.loc[a,'genotype'],
                          'genotype_b': g.loc[b,'genotype'],
                          'cage_group': cagegrp,
                          'generation': g.loc[a, 'generation']})
            
    for genotype in meta[(meta['generation'] == generation) & 
                         (meta['genotype'] != 'b6cf1')]['genotype'].unique():
        
        for (cg_a, cg_b) in combinations(meta[(meta['generation'] == generation) & 
                                              (meta['genotype'] == genotype)]['cage_group'].unique(), 2):
            idx_a = meta[(meta['generation'] == generation) & 
                         (meta['genotype'] == genotype) &
                         (meta['cage_group'] == cg_a)].index
            idx_b = meta[(meta['generation'] == generation) & 
                         (meta['genotype'] == genotype) &
                         (meta['cage_group'] == cg_b)].index
            for a in idx_a:
                for b in idx_b:
                    dists_cg_between.append({
                        'dist': res_all['beta_diversity']['results'][metric][a,b], 
                        'generation': generation,
                        'genotype': genotype,
                        'cg_cmp': '%s vs %s' % tuple(sorted([cg_a, cg_b])),
                    })
dists_cg_within = pd.DataFrame(dists_cg_within)
dists_cg_between = pd.DataFrame(dists_cg_between)

    #dists_withincage = dists_withincage[dists_withincage['genotype_a'] == 'c57bl6/j']
dists_cg_within

# %% [code]
fix, axes = plt.subplots(5,1, figsize=(5,5*5))
for (generation, ax) in zip(['f1','f2','f3','f4','f5'], axes):
    sns.violinplot(data=dists_cg_within[dists_cg_within['generation'] == generation], y='dist', x='cage_group', ax=ax)
    pairs = []
    for (a,b) in [
        ('b%s-1' % generation,'b%s-2' % generation),
        ('b%s-2' % generation,'b%s-3' % generation),
        ('b%s-1' % generation,'b%s-3' % generation),
        ('c%s-1' % generation,'c%s-2' % generation),
        ('c%s-2' % generation,'c%s-3' % generation),
        ('c%s-1' % generation,'c%s-3' % generation)]:
        if (a in meta['cage_group'].unique()) & (b in meta['cage_group'].unique()):
            pairs.append((a,b))
    
    annotator = Annotator(ax, pairs, data=dists_cg_within[dists_cg_within['generation'] == generation], y='dist', x='cage_group')
    annotator.configure(test='Mann-Whitney', text_format='star', loc='outside', comparisons_correction="fdr_bh", correction_format="default")
    annotator.apply_and_annotate()


# %% [code]
fix, axes = plt.subplots(1,1)
sns.boxplot(data=dists_cg_between[dists_cg_between['generation'] == 'f1'], y='dist', x='cg_cmp', ax=axes)


# %% [code]
plotNetworks('cage_group', meta[meta['cage_group'].isin(['bf1-1','bf1-2'])], None, res_all['beta_diversity']['results'], summarize=True)[0]

# %% [code]
fix, axes = plt.subplots(1,1)
sns.boxplot(data=dists_cg_between[dists_cg_between['generation'] == 'f2'], x='dist', y='cg_cmp', ax=axes)


# %% [code]
fix, axes = plt.subplots(1,1)
sns.boxplot(data=dists_cg_between[dists_cg_between['generation'] == 'f3'], x='dist', y='cg_cmp', ax=axes)


# %% [code]
fix, axes = plt.subplots(1,1)
sns.boxplot(data=dists_cg_between[dists_cg_between['generation'] == 'f4'], x='dist', y='cg_cmp', ax=axes)


# %% [code]
fix, axes = plt.subplots(1,1)
sns.boxplot(data=dists_cg_between[dists_cg_between['generation'] == 'f5'], x='dist', y='cg_cmp', ax=axes)


# %% [markdown]
# Eco Filter?

# %% [markdown]
## add host_subject_id, parents, cages to metadata (only run once!)

# %% [code]
if False:
    sample_file =pd.read_csv("../FromQiita/13422_20230201-071657.txt", sep="\t", dtype=str, index_col=0)

    res = dict()
    with open('map_hsid_nr.txt', 'r') as f:
        for i, line in enumerate(f.readlines()):
            (realID, microID) = line.strip().split()
            microID = microID.split('(')[-1].split(')')[0]
            if microID not in res.keys():
                res[microID] = realID
            else:
                #print('duplicate', i)
                if res[microID] != realID:
                    raise ValueError(line)
    res = pd.Series(res)
    res.name = 'host_subject_id_stefan'

    h = pd.read_excel('map_hsid_parent_cage.xlsx', dtype=str).set_index('Unnamed: 0')
    display(h.iloc[20:25,:])

    display(h['cage_line'].value_counts())

    assert list(h[pd.notnull(h['father_subject_id_stefan'])].merge(newMeta, left_on='father_subject_id_stefan', right_on='host_subject_id_stefan', how='left')['sex'].unique()) == ['male']

    ss = []
    for l in h['mothers_subject_ids_stefan'].dropna().apply(lambda x: x.split('|')).values:
        ss.extend(l)
    assert list(newMeta[newMeta['host_subject_id_stefan'].isin(list(set(ss)))]['sex'].unique())  == ['female']

    num_cl = 0
    while num_cl < 297:
        num_cl = h['cage_line'].dropna().shape[0]
        print(num_cl)
        for idx, row in h.iterrows():
            if pd.isnull(row['cage_line']) and pd.notnull(row['father_subject_id_stefan']):
                cl = h[h['host_subject_id_stefan'] == row['father_subject_id_stefan']]['cage_line'].unique()
                if len(cl) == 1:
                    h.loc[idx, 'cage_line'] = cl[0]

    display(h['cage_line'].value_counts())

    newMeta = sample_file.merge(h, left_on='host_subject_id', right_index=True, how='left')
    display(newMeta.head())
    newMeta.to_csv('new_meta.tsv', sep="\t")

    display(newMeta['cage_line'].value_counts())

    _ = emperor(meta, res_all['beta_diversity']['results'], fp_results='./Generated/Emperor', infix='_ecofilter_', dry=False)

# %% [markdown]
## Cage effect: are mice of same cage more similar than mice of different cages in same cage_line/generation/genotype? <br/> -->YES
Only consider c57bl6/j genotype, as ALL F>=1 mice of balb/c originate from the same male individual

# %% [code]
colors = {'intra': 'orange', 'inter': 'blue'}
order_generations = ['rm','p1','f1','f2','f3','f4','f5','control']
key_generations = lambda x: x.apply(lambda a: order_generations.index(a))
def map_geolocname(x):
    return {'France:Lanvier': 'Lanvier', 'Germany:Duesseldorf': 'Duesseldorf'}.get(x,x)
order_cagelines = ["C57BL6/J_line1", "C57BL6/J_line2", "C57BL6/J_line3", "BALB/c_line4", "BALB/c_line5", "BALB/c_line6"]

colors_animals_per_cage = {'few (1-3 mice)': '#7fc97f', 
                           'many (4-5 mice)': '#beaed4'}

colors = {
    'c57bl6/j': '#0000ff',
    "C57BL6/J_line1": "#deebf7", 
    "C57BL6/J_line2": "#9ecae1", 
    "C57BL6/J_line3": "#3182bd",
    
    "b6cf1": "#ff0000",
    
    'balb/c': '#f27304',
    "BALB/c_line4": "#fee8c8",
    "BALB/c_line5": "#fdbb84",
    "BALB/c_line6": "#e34a33",
    
    'p1': '#8e0152',
    'f1': '#d6589e',
    'f2': '#f5c4e1',
    'f3': '#c7e89f',
    'f4': '#6eae36',
    'f5': '#276419',
    
    'mJAX': '#ff0000',
    'mTAC': '#0000ff',
}

# %% [code]
meta = prj_data['metadata'][(prj_data['metadata']['sample_type'] == 'colon content') & 
                            (prj_data['metadata']['host_scientific_name'] != 'callithrix jacchus')]
meta = meta[(meta['genotype'].isin(["c57bl6/j","balb/c"])) & (meta['iacuc_institute'] == "LANUV")]
# drop data without sequences
meta = meta.loc[list(set(res_all['rarefaction']['results'].columns) & set(meta.index)), :]

meta['genotype'].value_counts()

# %% [code]
#meta[meta['genotype'] == 'c57bl6/j'].groupby(['genotype','generation','cage_line','cage_id']).size()
res = []
for metric in res_all['beta_diversity']['results'].keys():
    for (_, generation, cage_line), g in meta[meta['genotype'] == 'c57bl6/j'].groupby(['genotype','generation','cage_line']):
#    for (_, generation, cage_line), g in meta.groupby(['genotype','generation','cage_line']):
        if metric == list(res_all['beta_diversity']['results'].keys())[0]:
            print(generation, cage_line, g.shape[0])
        for (a,b) in combinations(g.index, 2):
            cmp = None
            if meta.loc[a, 'cage_id'] == meta.loc[b, 'cage_id']:
                cmp = 'intra'
            else:
                cmp = 'inter'
            dist = {'dist': res_all['beta_diversity']['results'][metric][a,b],
                    'metric': metric,
                    'cmp': cmp,
                    'generation': generation}
            res.append(dist)
res = pd.DataFrame(res)

# %% [code]
fig, axes = plt.subplots(1,1)
data = res
sns.boxplot(data=data, y='dist', x='metric', hue='cmp', ax=axes, palette=colors)
fig.suptitle("are mice of same cage more similar than mice of different cages in same cage_line/generation/C57BL6/J")
annotator = Annotator(axes, [
    ((m, 'inter'), (m, 'intra')) for m in res_all['beta_diversity']['results'].keys()
], data=data, y='dist', x='metric', hue='cmp')
annotator.configure(test='Mann-Whitney', text_format='star', loc='outside', comparisons_correction="fdr_bh", correction_format="default")
annotator.apply_and_annotate()

# %% [code]
fig, axes = plt.subplots(1,1)
data = res[(res['metric'] == 'bray_curtis') & (res['cmp'] == 'intra')]
sns.violinplot(data=data, y='dist', x='generation', order=order_generations, ax=axes)

annotator = Annotator(axes, [
    ('p1', 'f1'), ('f1', 'f2'), ('f2', 'f3'), ('f3', 'f4'), ('f4', 'f5'), 
], data=data, y='dist', x='generation', order=order_generations)
annotator.configure(test='Mann-Whitney', text_format='star', loc='outside', comparisons_correction="fdr_bh", correction_format="default")
annotator.apply_and_annotate()

# %% [markdown]
## Cage lineage effect: are mice of same cage_lineage more similar than mice of different cage_lines/generation/C57BL6/J? <br/> -->YES
cage lineage effect also persists if no comparisions within cage are considered

# %% [code]
#meta[meta['genotype'] == 'c57bl6/j'].groupby(['genotype','generation','cage_line','cage_id']).size()
res = []
for metric in res_all['beta_diversity']['results'].keys():
    for (_, generation), g in meta[meta['genotype'] == 'c57bl6/j'].groupby(['genotype','generation']):
        if metric == list(res_all['beta_diversity']['results'].keys())[0]:
            print(generation, g.shape[0])
        for (a,b) in combinations(g.index, 2):
            cmp = None
            if meta.loc[a, 'cage_id'] == meta.loc[b, 'cage_id']:
                # skip comparisions of two mice living in the same cage, as we already know that there is 
                # a cage effect (as always)
                continue
            if meta.loc[a, 'cage_line'] == meta.loc[b, 'cage_line']:
                cmp = 'intra'
            else:
                cmp = 'inter'
            dist = {'dist': res_all['beta_diversity']['results'][metric][a,b],
                    'metric': metric,
                    'cmp': cmp,
                    'generation': generation}
            res.append(dist)
res = pd.DataFrame(res)

# %% [code]
fig, axes = plt.subplots(1,1)
data = res
sns.boxplot(data=data, y='dist', x='metric', hue='cmp', ax=axes, palette=colors)

fig.suptitle("are mice of same cage_lineage more similar than mice of different cage_lines/generation/C57BL6/J?")
annotator = Annotator(axes, [
    ((m, 'inter'), (m, 'intra')) for m in res_all['beta_diversity']['results'].keys()
], data=data, y='dist', x='metric', hue='cmp')
annotator.configure(test='Mann-Whitney', text_format='star', loc='outside', comparisons_correction="fdr_bh", correction_format="default")
_ = annotator.apply_and_annotate()

# %% [markdown]
## do genotype microbiomes become more distant over time? <br/> --> Yes, but only up to f2
Furthermore: also distances within genotypes become larger, most probably because cage_lineage effects grow

# %% [code]
#meta[meta['genotype'] == 'c57bl6/j'].groupby(['genotype','generation','cage_line','cage_id']).size()
res = []
for metric in res_all['beta_diversity']['results'].keys():
    for generation, g in meta.groupby('generation'):
        if metric == list(res_all['beta_diversity']['results'].keys())[0]:
            print(generation, g.shape[0])
        for (a,b) in combinations(g.index, 2):
            cmp = None
            if meta.loc[a, 'genotype'] == meta.loc[b, 'genotype']:
                cmp = 'intra'
            else:
                cmp = 'inter'
            dist = {'dist': res_all['beta_diversity']['results'][metric][a,b],
                    'metric': metric,
                    'cmp': cmp,
                    'generation': generation}
            res.append(dist)
res = pd.DataFrame(res)

# %% [markdown]
### across genotype

# %% [code]
for metric in res['metric'].unique():
    fig, axes = plt.subplots(1,1)
    data = res[(res['metric'] == metric) & (res['cmp'] == 'inter')]
    sns.violinplot(data=data, y='dist', x='generation', order=order_generations, ax=axes)
    fig.suptitle(metric)
    annotator = Annotator(axes, [
        ('p1', 'f1'), ('f1', 'f2'), ('f2', 'f3'), ('f3', 'f4'), ('f4', 'f5'), 
        ('p1', 'f1'), ('p1', 'f2'), ('p1', 'f3'), ('p1', 'f4'), ('p1', 'f5'), 
    ], data=data, y='dist', x='generation', order=order_generations)
    #annotator.apply_test()
    annotator.configure(test='Mann-Whitney', text_format='star', loc='outside', comparisons_correction="fdr_bh", correction_format="default")
    annotator.apply_test(alternative='less')
    annotator.annotate()

# %% [markdown]
### within genotype

# %% [code]
for metric in res['metric'].unique():
    fig, axes = plt.subplots(1,1)
    data = res[(res['metric'] == metric) & (res['cmp'] == 'intra')]
    sns.violinplot(data=data, y='dist', x='generation', order=order_generations, ax=axes)
    fig.suptitle(metric)
    annotator = Annotator(axes, [
        ('p1', 'f1'), ('f1', 'f2'), ('f2', 'f3'), ('f3', 'f4'), ('f4', 'f5'), 
        ('p1', 'f1'), ('p1', 'f2'), ('p1', 'f3'), ('p1', 'f4'), ('p1', 'f5'), 
    ], data=data, y='dist', x='generation', order=order_generations)
    #annotator.apply_test()
    annotator.configure(test='Mann-Whitney', text_format='star', loc='outside', comparisons_correction="fdr_bh", correction_format="default")
    annotator.apply_test(alternative='less')
    annotator.annotate()

# %% [markdown]
## ASVs

# %% [code]
rank = 'Genus'
minreads = 1

taxa = dict()
featureTableAll = collapseCounts_objects(res_all['rarefaction']['results'], 
                                         rank, res_all['taxonomy']['RDP']['results']['Taxon'])[0]
m = meta[meta['cage_line'] == 'C57BL6/J_line2']
for g in m['generation'].unique():
    idx_samples = [s for s in m[m['generation'] == g].index if s in res_all['rarefaction']['results'].columns]
    
    # at ASV level
    #featureTable = res_all['rarefaction']['results']
    # at Genus Level
    featureTable = collapseCounts_objects(res_all['rarefaction']['results'], 
                                          rank, res_all['taxonomy']['RDP']['results']['Taxon'], out=None)[0]

    featureTable = featureTable[featureTable.loc[:, idx_samples].max(axis=1) >= minreads]
    
    taxa[g] = set(featureTable.index)
    print("%  i taxa with at least %i reads in 1 of %i samples for generation '%s'" % (len(taxa[g]), minreads, len(idx_samples), g))
    
taxaALL = None
for _, a in taxa.items():
    if taxaALL is None:
        taxaALL = a
    else:
        taxaALL |= a
print("total AVSs across generations: ", len(taxaALL))

# %% [code]
res = []
for s in taxaALL:
    for g in taxa.keys():
        idx_samples = m[m['generation'] == g].index
        res.append({'taxon': s, 'generation': g, 'present': featureTableAll.loc[s, idx_samples].sum() / (featureTableAll.loc[:, idx_samples].sum().sum())})
res = pd.pivot_table(pd.DataFrame(res), index='taxon', columns='generation', values='present')#.to_excel("test.xlsx")
res

# %% [code]
data = res[res.max(axis=1) >= 0.01]
data = (data.T / data.T.mean()).T
data.shape

# %% [code]
sns.lineplot(data=data.loc[:, order_generations].T, legend=False)

# %% [markdown]
### Venn

# %% [code]
meta_v = prj_data['metadata'][(prj_data['metadata']['sample_type'] == 'colon content') & 
                              (prj_data['metadata']['host_scientific_name'] != 'callithrix jacchus') & 
                              (prj_data['metadata']['iacuc_institute'] == 'LANUV')]
display(meta_v.fillna('?').groupby(['iacuc_institute','genotype','generation','cage_line']).size())

rank = 'raw'
featureTableAll = collapseCounts_objects(res_all['rarefaction']['results'], 
                                         rank, res_all['taxonomy']['RDP']['results']['Taxon'])[0]

# %% [code]
def some(values, ratio=1/3):
    res = values.value_counts()
    trues = 0
    if True in res.index:
        trues = res[True]
    return (trues / values.shape[0]) >= ratio
(ftab_geno.loc[:, idx_samples] >= minreads).apply(some, axis=1)

# %% [code]
minreads = 10
for genotype in ['balb/c', 'c57bl6/j']:
    ftab_geno = featureTableAll.loc[:, list(set(featureTableAll.columns) & set(meta_v[meta_v['genotype'].isin([genotype, 'b6cf1'])].index))]
    ftab_geno = ftab_geno[(ftab_geno >= minreads).any(axis=1)]
    
    print(genotype, ftab_geno.shape)
    
    # in foster mother
    idx_samples = list(set(meta_v[meta_v['generation'] == 'rm'].index) & set(ftab_geno.columns))
    #taxa_rm = ftab_geno[(ftab_geno.loc[:, idx_samples] >= minreads).any(axis=1)].loc[:, idx_samples]
    taxa_rm = ftab_geno[(ftab_geno.loc[:, idx_samples] >= minreads).apply(some, axis=1)].loc[:, idx_samples]
    
    # in f5
    idx_samples = list(set(meta_v[meta_v['generation'] == 'f5'].index) & set(ftab_geno.columns))
    #taxa_f5 = ftab_geno[(ftab_geno.loc[:, idx_samples] >= minreads).any(axis=1)].loc[:, idx_samples]
    taxa_f5 = ftab_geno[(ftab_geno.loc[:, idx_samples] >= minreads).apply(some, axis=1)].loc[:, idx_samples]

    # in p1-f4
    taxa_p1f4 = None
    idx_samples = []
    for generation in order_generations[1:-1]:
        idx_gen_samples = list(set(meta_v[meta_v['generation'] == generation].index) & set(ftab_geno.columns))
        idx_samples += idx_gen_samples
        #sel = set(ftab_geno[(ftab_geno.loc[:, idx_gen_samples] >= minreads).any(axis=1)].loc[:, idx_gen_samples].index)
        sel = set(ftab_geno[(ftab_geno.loc[:, idx_gen_samples] >= minreads).apply(some, axis=1)].loc[:, idx_gen_samples].index)
        if taxa_p1f4 is None:
            taxa_p1f4 = sel
        else:
            taxa_p1f4 &= sel
    taxa_p1f4 = ftab_geno.loc[list(taxa_p1f4), idx_samples]
    
    print(taxa_rm.shape, taxa_f5.shape, taxa_p1f4.shape)
    fig, axes = plt.subplots(1,1)
    venn3_unweighted([set(taxa_rm.index), 
           set(taxa_p1f4.index), 
           set(taxa_f5.index)], ('recipient mothers', 'p1,f1,f2,f3,f4', 'f5'), ax=axes)
    axes.set_title(genotype)
    
    fig, axes = plt.subplots(1,1)
    weights = {
        '100': (featureTableAll.loc[list(set(taxa_rm.index) - (set(taxa_p1f4.index) | set(taxa_f5.index))), :] / featureTableAll.iloc[:, 0].sum()).sum().mean(),
        '010': (featureTableAll.loc[list(set(taxa_p1f4.index) - (set(taxa_rm.index) | set(taxa_f5.index))), :] / featureTableAll.iloc[:, 0].sum()).sum().mean(),
        '001': (featureTableAll.loc[list(set(taxa_f5.index) - (set(taxa_rm.index) | set(taxa_p1f4.index))), :] / featureTableAll.iloc[:, 0].sum()).sum().mean(),

        '101': (featureTableAll.loc[list((set(taxa_rm.index) | set(taxa_f5.index)) - set(taxa_p1f4.index)), :] / featureTableAll.iloc[:, 0].sum()).sum().mean(),
        '110': (featureTableAll.loc[list((set(taxa_rm.index) | set(taxa_p1f4.index)) - set(taxa_f5.index)), :] / featureTableAll.iloc[:, 0].sum()).sum().mean(),
        '011': (featureTableAll.loc[list((set(taxa_f5.index) | set(taxa_p1f4.index)) - set(taxa_rm.index)), :] / featureTableAll.iloc[:, 0].sum()).sum().mean(),

        '111': (featureTableAll.loc[list((set(taxa_f5.index) & set(taxa_p1f4.index)) & set(taxa_rm.index)), :] / featureTableAll.iloc[:, 0].sum()).sum().mean()}
    venn3(weights, ('recipient mothers', 'p1,f1,f2,f3,f4', 'f5'), ax=axes, subset_label_formatter=lambda x: "%.2f" % x, normalize_to=sum(weights.values()))


# %% [markdown]
## what is the most prototypic generation/cage_line per genotype?

# %% [code]
meta.groupby('genotype').size()

# %% [code]
fig, axes = plt.subplots(3,1,figsize=(5,15))
for metric, ax in zip(res_all['beta_diversity']['results'].keys(), axes):
    res = []
    for genotype, g in meta.groupby('genotype'):
        for (generation, cage_line), g2 in g.groupby(['generation', 'cage_line']):
            dm_subset = res_all['beta_diversity']['results'][metric].to_data_frame().loc[g2.index, list(set(g.index) - set(g2.index))]
            res.append({'genotype': genotype, 
                        'generation': generation,
                        'cage_line': cage_line,
                        'avgdist': dm_subset.mean(axis=1).mean()})
            #print(genotype, generation, cage_line, g2.shape[0], g.shape[0], dm_subset.shape, dm_subset.sum(axis=1).mean())
    res = pd.DataFrame(res)
    sns.barplot(data=res, y='avgdist', x='generation', order=order_generations, hue='cage_line', ax=ax)
    ax.set_title(metric)
    ax.legend(bbox_to_anchor=(1.1, 1.05))
    winner = res.loc[res.groupby('cage_line')['avgdist'].idxmin(),:].set_index('cage_line')
    winner.index.name = metric
    display(winner)

# %% [code]
c_cat = ['genotype','generation','sex']
for c_cat in [['genotype','generation','sex'], 
              ['genotype','generation','cage_line'],
              ['genotype','generation','sex', 'cage_line'],
              ['genotype','generation','sex', 'cage_id'],
              ['genotype','generation','cage_id'],
              ['genotype','generation','cage_line', 'cage_id'],
              ['genotype','generation','sex', 'cage_line', 'cage_id'],
             ['cage_line','genotype']]:
    usegrid = True
    max_rows = max(len(res_all['alpha_diversity']['results'].keys()), len(res_all['beta_diversity']['results'].keys()))
    fig, axes = plt.subplots(max_rows, 2, figsize=(5*2, 5*max_rows))
    for i, metric in enumerate(res_all['alpha_diversity']['results'].keys()):
        redundancy_analysis_alpha(meta, res_all['alpha_diversity']['results'][metric],
                                  categorials=c_cat, intervals=[], dates={},
                                  ax=axes[i][0], title='%s' % metric,
                                  dry=False, wait=False, use_grid=True)
    for i, metric in enumerate(res_all['beta_diversity']['results'].keys()):
        redundancy_analysis_beta(meta, res_all['beta_diversity']['results'][metric], metric,
                                 categorials=c_cat, intervals=[], dates={},
                                 ax=axes[i][1], title='%s' % metric,
                                 dry=False, wait=False, use_grid=True)

# %% [markdown]
## RF to predict genotype <br/>--> 83%, i.e. "filtered microbiom" is not prototypic

# %% [code]
meta_rf = prj_data['metadata'][(prj_data['metadata']['sample_type'] == 'colon content') & 
                               (prj_data['metadata']['host_scientific_name'] != 'callithrix jacchus')]
meta_rf.groupby(['iacuc_institute', 'geo_loc_name', 'genotype']).size()

# %% [code]


# %% [code]


# %% [code]
for s in range(len(order_generations)):
    clf = RandomForestClassifier(n_estimators=1000, n_jobs=4)
    gs = order_generations[s:]
    idx_train = list(set(meta_rf[(meta_rf['iacuc_institute'] == 'LANUV') & 
                                 (meta_rf['genotype'].isin(['balb/c', 'c57bl6/j'])) &
                                 (meta_rf['generation'].isin(gs))].index) & set(res_all['rarefaction']['results'].columns))
    c_train = res_all['rarefaction']['results'].loc[:, idx_train].T
    #c_train = c_train[c_train.sum(axis=1) <= 0].T
    #print(c_train.shape)
    m_train = meta_rf.loc[idx_train, 'genotype']
    display(meta_rf.loc[idx_train,:].groupby(['genotype', 'generation']).size())
    clf = clf.fit(c_train, m_train)

    idx_test = list(set(meta_rf[(meta_rf['iacuc_institute'] == 'ZETT') & (meta_rf['genotype'].isin(['balb/c', 'c57bl6/j']))].index) & set(res_all['rarefaction']['results'].columns))
    c_test = res_all['rarefaction']['results'].loc[:, idx_test].T
    #c_test = c_test[c_test.sum(axis=1) <= 0].T
    #print(c_test.shape)
    m_test = meta_rf.loc[idx_test, 'genotype']

    print(clf.score(c_test, m_test))

# %% [code]
p = pd.DataFrame(clf.predict(c_test), columns=['prediction'], index=c_test.index)
p.merge(meta_rf[['genotype','geo_loc_name','iacuc_institute']], left_index=True, right_index=True, how="left")

# %% [code]
idx_test = list(set(meta_rf[(meta_rf['iacuc_institute'] == 'LANUV') & (meta_rf['genotype'].isin(['b6cf1']))].index) & set(res_all['rarefaction']['results'].columns))
c_test = res_all['rarefaction']['results'].loc[:, idx_test].T
#c_test = c_test[c_test.sum(axis=1) <= 0].T
print(c_test.shape)
m_test = meta_rf.loc[idx_test, 'genotype']

p = pd.DataFrame(clf.predict(c_test), columns=['prediction'], index=c_test.index)
p.merge(meta_rf[['genotype','geo_loc_name','iacuc_institute']], left_index=True, right_index=True, how="left")

# %% [code]
dists = []
for genotype, g in meta_rf[(meta_rf['iacuc_institute'] == 'ZETT') & (meta_rf['genotype'].isin(['balb/c', 'c57bl6/j']))].groupby('genotype'):
    for fm in meta_rf[meta_rf['generation'] == 'rm'].index:
        for cm in g.index:
            for metric in res_all['beta_diversity']['results'].keys():
                dists.append({'metric': metric,
                              'genotype': genotype,
                              'recipient mouse': meta_rf.loc[fm, 'host_subject_id'],
                              'dist': res_all['beta_diversity']['results'][metric][fm, cm]})
    print(genotype, g.shape)
    
g = sns.boxplot(data=pd.DataFrame(dists), y='dist', x='metric', hue='genotype')
g.set_title('dist foster mothers VS ZETT controls')
annotator = Annotator(g, [
        ((m, 'balb/c'), (m, 'c57bl6/j')) for m in res_all['beta_diversity']['results'].keys()
    ], data=pd.DataFrame(dists), y='dist', x='metric', hue='genotype')
annotator.configure(test='Mann-Whitney', text_format='star', loc='outside', comparisons_correction="fdr_bh", correction_format="default")
annotator.apply_and_annotate()

# %% [code]
refGeneration = 'p1'
for refGeneration in order_generations:
    fig, axes = plt.subplots(1,1)
    dists = []
    for genotype, g in meta_rf[(meta_rf['iacuc_institute'] == 'LANUV') & (meta_rf['genotype'].isin(['balb/c', 'c57bl6/j'])) & (meta_rf['generation'] == refGeneration)].groupby('genotype'):
        for fm in meta_rf[meta_rf['generation'] == 'rm'].index:
            for cm in g.index:
                for metric in res_all['beta_diversity']['results'].keys():
                    try:
                        dists.append({'metric': metric,
                                      'genotype': genotype,
                                      'recipient mouse': meta_rf.loc[fm, 'host_subject_id'],
                                      'dist': res_all['beta_diversity']['results'][metric][fm, cm]})
                    except MissingIDError:
                        pass
        print(genotype, g.shape)

    sns.boxplot(data=pd.DataFrame(dists), y='dist', x='metric', hue='genotype', ax=axes)
    axes.set_title('dist foster mothers VS %s' % refGeneration)
    annotator = Annotator(axes, [
            ((m, 'balb/c'), (m, 'c57bl6/j')) for m in res_all['beta_diversity']['results'].keys()
        ], data=pd.DataFrame(dists), y='dist', x='metric', hue='genotype')
    annotator.configure(test='Mann-Whitney', text_format='star', loc='outside', comparisons_correction="fdr_bh", correction_format="default")
    annotator.apply_and_annotate()

# %% [code]
dists = []
for generation in order_generations:
    for genotype, g in meta_rf[(meta_rf['iacuc_institute'] == 'LANUV') & 
                               (meta_rf['genotype'].isin(['balb/c', 'c57bl6/j'])) &
                               (meta_rf['generation'] == generation)
                              ].groupby('genotype'):
        for fm in meta_rf[(meta_rf['iacuc_institute'] == 'ZETT') &
                          (meta_rf['genotype'].isin(['balb/c', 'c57bl6/j']))].index:
            for cm in g.index:
                for metric in res_all['beta_diversity']['results'].keys():
                    try: 
                        dists.append({'metric': metric,
                                      'genotype': '%s <-> %s' % (genotype, meta_rf.loc[fm, 'genotype']),
                                      'generation': generation,
                                      'cmpGroup': 'ZETT controls',
                                      'dist': res_all['beta_diversity']['results'][metric][fm, cm]})
                    except MissingIDError:
                        pass
        for fm in meta_rf[meta_rf['generation'] == 'rm'].index:
            for cm in g.index:
                for metric in res_all['beta_diversity']['results'].keys():
                    try:
                        dists.append({'metric': metric,
                                      'genotype': '%s <-> %s' % (genotype, meta_rf.loc[fm, 'genotype']),
                                      'generation': generation,
                                      'cmpGroup': 'foster mothers',
                                      'dist': res_all['beta_diversity']['results'][metric][fm, cm]})
                    except MissingIDError:
                        pass
        
    fig, axes = plt.subplots(1,1)
    sns.boxplot(data=pd.DataFrame(dists), y='dist', x='metric', hue='genotype', ax=axes)
    axes.set_title('dist ZETTcontrols VS generation %s' % generation)
    annotator = Annotator(axes, [
            ((m, 'balb/c <-> balb/c'), (m, 'balb/c <-> c57bl6/j')) for m in res_all['beta_diversity']['results'].keys()
        ] + [
            ((m, 'c57bl6/j <-> c57bl6/j'), (m, 'c57bl6/j <-> balb/c')) for m in res_all['beta_diversity']['results'].keys()
        ]
        , data=pd.DataFrame(dists), y='dist', x='metric', hue='genotype')
    annotator.configure(test='Mann-Whitney', text_format='star', loc='outside', comparisons_correction="fdr_bh", correction_format="default")
    annotator.apply_and_annotate()

# %% [code]
d = pd.DataFrame(dists)
sns.boxplot(data=d[(d['cmpGroup'] == 'ZETT controls') & (d['genotype'].isin(['c57bl6/j <-> c57bl6/j',
                                                                             'balb/c <-> balb/c']))], y='dist', x='metric', hue_order=order_generations, hue='generation')

# %% [code]
sns.boxplot(data=pd.DataFrame(dists), y='dist', x='metric', hue='genotype', ax=axes)

# %% [markdown]
# Temporal Summary

# %% [code]
def comp_temporal(meta, beta, summarize_baseline=False):
    meta = meta[meta['host_scientific_name'] != 'callithrix jacchus']  # exclude monkeys
    
    meta = meta.loc[list(set(meta.index) & set(beta[list(beta.keys())[0]].ids))]  # exclude data without sequences
    
    # --------- compute distances to recipient mothers
    dists = []
    idx_reference = meta[meta['generation'] == 'rm'].index
    #print("%i references: rm mice" % len(idx_reference))
    for (genotype, generation), g in meta[meta['generation'] != 'rm'].groupby(['genotype','generation']):
        #print("%i cmp (%s, %s)" % (g.shape[0], genotype, generation))
        for metric in beta.keys():
            for idx_ref in idx_reference:
                for idx_cmp in g.index:
                    dists.append({'reference': idx_ref,
                                  'cmp': idx_cmp,
                                  'metric': metric,
                                  'generation': generation,
                                  'genotype': genotype,
                                  'sex': meta.loc[idx_cmp, 'sex'],
                                  'cage_line': meta.loc[idx_cmp, 'cage_line'],
                                  'cl': meta.loc[idx_cmp, 'cl'],
                                  'dist': beta[metric][idx_ref,idx_cmp]})
    dists = pd.DataFrame(dists)

    # --------- compute intra cage distances
    cage_variance = []
    for cage_id, g in meta.groupby('cage_id'):
        assert g['genotype'].unique().shape[0] == 1
        assert g['generation'].unique().shape[0] == 1
        for metric in beta.keys():
            for (a,b) in combinations(g.index, 2):
                assert meta.loc[[a,b],'generation'].unique().shape[0] == 1
                cage_variance.append({'dist': beta[metric][a,b],
                                      'cage_id': cage_id,
                                      'generation': meta.loc[[a,b],'generation'].unique()[0],
                                      'metric': metric})
    cage_variance = pd.DataFrame(cage_variance)
    if summarize_baseline:
        cage_variance = cage_variance.groupby(['metric','cage_id']).mean().reset_index().groupby('metric')['dist'].mean()
    else:
        cage_variance = cage_variance.set_index('metric')
    # inject dist np.nan rows for missing metric/generation, necessary for correct plot order
    if not summarize_baseline:
        inject = []
        for metric, g in cage_variance.groupby('metric'):
            for generation in order_generations:
                if generation not in g['generation'].unique():
                    print(generation)
                    inject.append({'metric': metric,
                                   'dist': np.nan,
                                   'cage_id': 'fake-%i' % (len(inject)),
                                   'generation': generation})
        if len(inject) > 0:
            cage_variance = pd.concat([cage_variance.reset_index(), pd.DataFrame(inject)]).reset_index().set_index('metric')

    # --------- compute intra cage_line distances (exclude comparisons within same cage_id)
    cageline_variance = []
    for (genotype, generation, cage_line), g in meta.groupby(['genotype','generation','cage_line']):
        assert g['genotype'].unique().shape[0] == 1
        assert g['generation'].unique().shape[0] == 1
        assert g['cage_id'].unique().shape[0] >= 1
        for metric in beta.keys():
            for (a,b) in combinations(g.index, 2):
                if (meta.loc[a, 'cage_id'] != meta.loc[b, 'cage_id']):
                    cageline_variance.append({'dist': beta[metric][a,b],
                                              'cage_line': cage_line,
                                              'cl': g.loc[a,'cl'],
                                              'genotype': genotype,
                                              'generation': generation,
                                              'metric': metric})
    cageline_variance = pd.DataFrame(cageline_variance)
    num_cageline = cageline_variance.groupby(['metric','genotype','generation','cage_line']).size()
    if summarize_baseline:
        cageline_variance = cageline_variance.groupby(['metric','genotype','generation','cage_line'])['dist'].mean().reset_index().groupby('metric')['dist'].mean()
    else:
        cageline_variance = cageline_variance.set_index('metric')

    # --------- compute distance between both genotypes
    dists_genotype = []
    for generation, g in meta[meta['generation'] != 'rm'].groupby('generation'):
        for a in g[g['genotype'] == 'balb/c'].index:
            for b in g[g['genotype'] == 'c57bl6/j'].index:
                for metric in beta.keys():
                    dists_genotype.append({'dist': beta[metric][a,b],
                                           'metric': metric,
                                           'generation': generation})
    dists_genotype = pd.DataFrame(dists_genotype)
    
    # --------- compute distance between genotypes and 4*6 ZETT controls
    ctrl_dists = []
    for metric in beta.keys():
        dm = beta[metric].to_data_frame()
        for (generation, genotype_focus), g_focus in meta.groupby(['generation', 'genotype']):
            for (geo_loc_name, genotype_control), g_control in meta[(meta['iacuc_institute'] == 'ZETT') & 
                             meta['genotype'].isin(['balb/c','c57bl6/j'])
                            ].groupby(['geo_loc_name','genotype']):
                if (genotype_focus == genotype_control) | (generation == 'rm'):
                    d = dm.loc[list(set(g_focus.index) & set(dm.index)),list(set(g_control.index) & set(dm.index))].stack()
                    d = d.reset_index().rename(columns={'level_0': 'sample_focus',
                                                        'level_1': 'sample_control',
                                                        0: 'dist'})
                    d['metric'] = metric
                    d['generation'] = generation
                    d['genotype'] = genotype_control
                    #d['cage_line'] = 
                    d['geo_loc_name'] = geo_loc_name
                    ctrl_dists.append(d)
    ctrl_dists = pd.concat(ctrl_dists)     
    ctrl_dists = ctrl_dists.merge(meta[['cage_line','cl']], left_on='sample_focus', right_index=True, how='left')

    return {
        'meta': meta,
        'dists': dists,
        'cage_variance': cage_variance,
        'cageline_variance': cageline_variance,
        'num_cageline': num_cageline,
        'dists_genotype': dists_genotype,
        'summarize_baseline': summarize_baseline,
        'ctrl_dists': ctrl_dists
    }

# %% [code]
res_gut = comp_temporal(prj_data['metadata'][(prj_data['metadata']['sample_type'] == 'colon content')],
                        {m: dm for (m,dm) in res_all['beta_diversity']['results'].items() if m == 'weighted_unifrac'}, 
                        False)
res_skin = comp_temporal(prj_data['metadata'][(prj_data['metadata']['sample_type'] == 'skin of external ear') & (prj_data['metadata']['batch'] != "2")],
                        {m: dm for (m,dm) in res_all_skin['beta_diversity']['results'].items() if m == 'unweighted_unifrac'}, 
                        True)

# %% [code]
def plot_temporal(meta, dists, cage_variance, cageline_variance, num_cageline, dists_genotype, summarize_baseline, ctrl_dists, title="", cage_line=True, alpha=None):
    sns.set_style("white")
    #fig, axes = plt.subplots(dists['metric'].unique().shape[0], 1, figsize=(12, 4*dists['metric'].unique().shape[0]))
    #if dists['metric'].unique().shape[0] == 1:
    #    axes = [axes]
    #for ((metric, g), ax) in zip(dists.groupby('metric'), axes):
    i_ax = 0
    for (metric, g) in dists.groupby('metric'):
        num_axes = 2
        if cage_line:
            num_axes = 3
        fig, axes = plt.subplots(num_axes+1+1, 1, figsize=(12, 4+3*(num_axes-1)+4+4), sharex=False, sharey=False)#, gridspec_kw={'height_ratios': [3] + ([3] * (num_axes-1)) + [3]})
        
        i_ax = 0
        ax = axes[i_ax]
        if summarize_baseline:
            ax.axhline(y=cage_variance[metric], linestyle='--', label='avg. intra cage', color=sns.color_palette()[7])
            ax.axhline(y=cageline_variance[metric], linestyle='--', label='avg. intra cage_line', color=sns.color_palette()[5])
        else:
            sns.lineplot(data=cage_variance.loc[metric,:].reset_index().sort_values(by='generation', key=key_generations), y='dist', x='generation', ax=ax, sort=False, label='intra cage', color=sns.color_palette()[7],  linestyle='--', errorbar=None, markers=True, marker="o")
            sns.lineplot(data=cageline_variance.loc[metric,:].sort_values(by='generation', key=key_generations), y='dist', x='generation', ax=ax, sort=False, label='intra cage_line', color=sns.color_palette()[5],  linestyle='--', errorbar=None, markers=True, marker="o")
            
        style = None
        if cage_line:
            style = 'cl' #'cage_line'
        sns.lineplot(data=g.sort_values(by='generation', key=key_generations), y='dist', x='generation', hue='genotype', ax=ax, sort=False, style=style, palette={'c57bl6/j': colors['c57bl6/j'],
                                                                                                                                                                  'balb/c':   colors['balb/c']}, marker="o")

        sns.lineplot(data=dists_genotype[dists_genotype['metric'] == metric].sort_values(by='generation', key=key_generations), y='dist', x='generation', ax=ax, sort=False, label='Δ(balb/c, c57bl6/j)', marker="o")

        #sns.violinplot(data=g, order=order_generations, y='dist', x='generation', hue='genotype', ax=ax)
        ax.set_title(metric)

        # make space for rm generation
        if summarize_baseline:
            rm_offset = 1
            ax.set_xlim((ax.get_xlim()[0]-rm_offset, ax.get_xlim()[1]))
            ax.set_xticks([-1*rm_offset] + ax.get_xticks())
            ax.set_xticklabels(order_generations[:-1])

        # top X axis labels
        top_x = ax.twiny()
        top_x.set_xticks(ax.get_xticks())
        top_x.set_xlim(ax.get_xlim())
        num_mice = meta.groupby(['generation', 'genotype']).size()
        num_cages = meta.groupby(['generation', 'genotype'])['cage_id'].unique().apply(lambda x: len(x))
        num_cagelines = meta.groupby(['generation', 'genotype'])['cage_line'].unique().apply(lambda x: len(x))
        labels = []
        for generation in order_generations[:-1]:
            label = ""
            for genotype in reversed(meta['genotype'].unique()):
                if generation == 'rm':
                    label += "%s: " % genotype
                try:
                    label += "%ii (%ic, %il)" % (num_mice.loc[generation, genotype], num_cages.loc[generation, genotype], num_cagelines.loc[generation, genotype])
                except KeyError as e:
                    if generation == 'rm':
                        label += "%ii (%ic, %il)" % (0, 0, 0)
                label += "\n"
            #labels.append('\n'.join(['%s: %ii (%ic)' % (geno, num, num_cages.loc[gen.get_text(), geno]) for (geno, num) in num_mice.loc[gen.get_text(), :].items()]))
            labels.append(label)
        if summarize_baseline:
            top_x.set_xticklabels(labels)

        # right Y axis labels
        if summarize_baseline:
            right_y = ax.twinx()
            right_y.set_ylim(ax.get_ylim())
            right_y.set_yticks([cage_variance[metric], cageline_variance[metric]])
            ncage = meta.groupby(['cage_id']).size()
            num_cages_perline = meta.groupby(['genotype','generation','cage_line'])['cage_id'].unique().apply(len)
            right_y.set_yticklabels(['%ic (Σ %ii, Ø %.1fi)' % (ncage[ncage > 1].shape[0], ncage[ncage > 1].sum(), ncage[ncage > 1].mean()), 
                                     'Σ %ic, Ø %.1fc\nΣ %ii-i, Ø %.1fi-i' % (num_cages_perline.sum(), num_cages_perline.mean(), num_cageline.loc[metric, :].sum(), num_cageline.loc[metric, :].mean())])

        # make Y axis label more speaking
        betalabel = 'β diversity: %s' % {'bray_curtis': 'Bray-Curtis',
                                           'unweighted_unifrac': 'unweighted UniFrac',
                                           'weighted_unifrac': 'weighted UniFrac'}.get(metric)
        ax.set_ylabel(betalabel)
        ax.set_title(title)

        L = ax.legend()
        for l in L.get_texts():
            if l.get_text() == 'c57bl6/j':
                l.set_text('Δ(rm, c57bl6/j)')
            elif l.get_text() == 'balb/c':
                l.set_text('Δ(rm, balb/c)')
        ax.set_xlabel("")
        if cage_line:
            L.set_bbox_to_anchor((1.2,1.05))
        
        geos = [['France:Janvier', 'Germany:Duesseldorf']]
        style = 'geo_loc_name'
        if cage_line:
            geos = [['France:Janvier'], ['Germany:Duesseldorf']]
            style = 'cl' #'cage_line'
        for loc in geos:
            i_ax += 1
            ax = axes[i_ax]
            sns.lineplot(data=ctrl_dists[(ctrl_dists['metric'] == metric) & (ctrl_dists['geo_loc_name'].isin(loc))].sort_values(by='generation', key=key_generations), y='dist', x='generation', hue='genotype', style=style, sort=False, ax=ax, palette={'c57bl6/j': colors['c57bl6/j'], 'balb/c':   colors['balb/c']}, marker="o")
            ax.set_ylabel(betalabel)
            L = ax.legend()
            for l in L.get_texts():
                if l.get_text() == 'c57bl6/j':
                    l.set_text('Δ(control, c57bl6/j)')
                elif l.get_text() == 'balb/c':
                    l.set_text('Δ(control, balb/c)')
            L.set_bbox_to_anchor((1.1,1.05))
            ax.set_title('  |  '.join(["%s %s: %ii" % (row['geo_loc_name'], row['genotype'], row[0]) 
                                       for _, row 
                                       in meta.loc[ctrl_dists[ctrl_dists['geo_loc_name'].isin(loc)]['sample_control'].unique(),:].groupby(['geo_loc_name','genotype']).size().reset_index().applymap(lambda x: {'France:Janvier': 'Janvier', 'Germany:Duesseldorf': 'Duesseldorf'}.get(x,x)).iterrows()]))

        if False:
            for ax in axes[:-2]:
                ax.set_ylim(
                    min(map(lambda x: x.get_ylim()[0], axes[:-2])),
                    max(map(lambda x: x.get_ylim()[1], axes[:-2])))
        #ax.set_xticks(axes[0].get_xticks())
        #print(axes[0].get_xlim(), axes[1].get_xlim())
        #print(axes[0].get_xticks(), axes[1].get_xticks())
        #break
        
        
    ### ALPHA
    ax = axes[i_ax+1]
    metaalpha = meta.merge(
                 alpha, left_index=True, right_index=True, how='left')
    sns.lineplot(metaalpha[(metaalpha['iacuc_institute'] == 'LANUV')].sort_values(by='generation', key=key_generations), y=alpha.iloc[:,0].name, x='generation', hue='genotype', sort=False,
             marker="o", palette={'c57bl6/j': colors['c57bl6/j'], 'balb/c':   colors['balb/c'], 'b6cf1': colors['b6cf1']}, ax=ax,
            )
    ax.set_ylabel("α diversity: %s" % ({"PD_whole_tree": "Faith's PD"}.get(alpha.iloc[:,0].name, alpha.iloc[:,0].name)))
    
    #fig, axes = plt.subplots(1,1,figsize=(4,3))
    ax = axes[i_ax+2]
    metaalpha['cat'] = metaalpha['generation'].fillna('?') + metaalpha['geo_loc_name'].fillna('?') + metaalpha['genotype'].fillna('?')
    sns.boxplot(data=metaalpha[(metaalpha['iacuc_institute'] == 'ZETT') | (metaalpha['generation'] == 'rm')], y=alpha.iloc[:,0].name, x='cat', color='lightgray', ax=ax,
                width=0.1
               )
    sns.swarmplot(data=metaalpha[(metaalpha['iacuc_institute'] == 'ZETT') | (metaalpha['generation'] == 'rm')], y=alpha.iloc[:,0].name, x='cat', ax=ax,
                  size=8)
    for ax in axes[-2:]:
        ax.set_ylim(
            min(map(lambda x: x.get_ylim()[0], axes[-2:])),
            max(map(lambda x: x.get_ylim()[1], axes[-2:]))*(0.94 if meta['sample_type'].unique()[0] == 'colon content' else 1.0))
        
    return fig

f = plot_temporal(**res_gut, title='all fecal data', cage_line=False, alpha=res_all['alpha_diversity']['results'][['PD_whole_tree']])
#f.savefig('Generated/temporal_gut.svg')

# %% [code]
f = plot_temporal(**res_skin, title='all skin data', cage_line=False, alpha=res_all_skin['alpha_diversity']['results'][['PD_whole_tree']])
#f.savefig('Generated/temporal_skin.svg')

# %% [code]
metric = 'weighted_unifrac'
sub_meta = prj_data['metadata'][(prj_data['metadata']['sample_type'] == 'colon content') &
                              (prj_data['metadata']['host_scientific_name'] != 'callithrix jacchus')
                              ].copy()

redundancy_analysis_beta(sub_meta, res_all['beta_diversity']['results'][metric], metric,
                         categorials=['genotype','generation','sex'], intervals=[], dates={},
                         title='%s' % metric,
                         dry=False, wait=True, use_grid=True)['results']

# %% [code]
metric = 'weighted_unifrac'
sub_meta = prj_data['metadata'][(prj_data['metadata']['sample_type'] == 'colon content') &
                              (prj_data['metadata']['host_scientific_name'] != 'callithrix jacchus') &
                              ((prj_data['metadata']['iacuc_institute'] == 'ZETT') 
                              # | (prj_data['metadata']['generation'] == 'f2')
                              )].copy()
sub_meta['cl'] = sub_meta['cl'].fillna("nanana")

redundancy_analysis_beta(sub_meta, res_all['beta_diversity']['results'][metric], metric,
                         categorials=['genotype','geo_loc_name','cl'], intervals=[], dates={},
                         title='%s' % metric,
                         dry=False, wait=True, use_grid=True)['results']

# %% [code]
res = []
dm = res_all['beta_diversity']['results']['weighted_unifrac'].to_data_frame()
for generation, g in prj_data['metadata'][(prj_data['metadata']['sample_type'] == 'colon content') & 
                                          (prj_data['metadata']['host_scientific_name'] != 'callithrix jacchus')].groupby('generation'):
    g = g.loc[list(set(g.index) & set(dm.index))]
    #print(generation, g.shape[0])
    for a in list(g['cage_line'].unique()):
        for b in list(g['cage_line'].unique()):
            #print("  ", a, b)
            res.append({'generation': generation,
                        'a': a,
                        'b': b,
                        'dist': dm.loc[g[g['cage_line'] == a].index, g[g['cage_line'] == b].index].stack().mean()
                       })
res = pd.DataFrame(res)

# %% [code]
la = ['C57BL6/J_line1', 'C57BL6/J_line2', 'C57BL6/J_line3']
lb = ['BALB/c_line4', 'BALB/c_line5', 'BALB/c_line6']

dmavg = pd.pivot_table(res, index='a', columns='b', values='dist').iloc[3:6,0:3]
triples = []
for t in combinations([(a,b) for a in la for b in lb], 3):
    if len(set(list(map(lambda x: x[0], list(t))) + list(map(lambda x: x[1], list(t))))) == 6:
        s = 0
        for (a,b) in list(t):
            s += dmavg.loc[a,b]
        triples.append((t, s))
# how we should pair cage lines to minimize distances across generations
pd.DataFrame(triples).set_index(0)[1].sort_values().head(1).index[0]

# %% [code]
for generation in order_generations:
    if generation == 'rm':
        continue
    sns.clustermap(pd.pivot_table(res[res['generation'] == generation], index='a', columns='b', values='dist'))
    plt.title(generation)

# %% [code]
sns.barplot(data=res[res['generation'] == 'f2'], y='dist', x='a', hue='b')
plt.legend(bbox_to_anchor=(1.1,0.5))

# %% [code]
metric = 'weighted_unifrac'
sub_meta = prj_data['metadata'][(prj_data['metadata']['sample_type'] == 'colon content') &
                                (prj_data['metadata']['host_scientific_name'] != 'callithrix jacchus') &
                                ((prj_data['metadata']['iacuc_institute'] == 'LANUV') 
                              # | (prj_data['metadata']['generation'] == 'f2')
                              )]
redundancy_analysis_beta(sub_meta, res_all['beta_diversity']['results'][metric], metric,
                         categorials=['genotype','cl'], intervals=[], dates={},
                         title='%s' % (metric),
                         dry=False, wait=False, use_grid=True)['results']

# %% [code]
metric = 'weighted_unifrac'
for generation, sub_meta in prj_data['metadata'][(prj_data['metadata']['sample_type'] == 'colon content') &
                                (prj_data['metadata']['host_scientific_name'] != 'callithrix jacchus') &
                                ((prj_data['metadata']['iacuc_institute'] == 'LANUV') 
                              # | (prj_data['metadata']['generation'] == 'f2')
                              )].groupby('generation'):
    print(generation, sub_meta.shape[0])
    redundancy_analysis_beta(sub_meta, res_all['beta_diversity']['results'][metric], metric,
                             categorials=['genotype','cl'], intervals=[], dates={},
                             title='%s %s' % (metric, generation),
                             dry=False, wait=False, use_grid=True)['results']

# %% [code]
def plot_taxonomy_emperor(prj_data, res_all, subset='gut'):
    m = None
    if subset == 'gut':
        m = prj_data['metadata'][prj_data['metadata']['sample_type'] == 'colon content'].copy()
    elif subset == 'skin':
        m = prj_data['metadata'][(prj_data['metadata']['sample_type'] == 'skin of external ear') & (prj_data['metadata']['batch'] == "3")].copy()
    m['g_cage_line'] = m['cage_line']

    # 6 French controls c57bl6/j
    idx = m[(m['genotype'] == 'c57bl6/j') & 
            (m['iacuc_institute'] == 'ZETT') &
            (m['geo_loc_name'] == 'France:Janvier')].index
    i1 = 1
    i2 = 1
    m.loc[idx, 'g1'] = str(i1) + ': ' + m.loc[idx, 'genotype'] + '\ncontrol ' + m.loc[idx, 'geo_loc_name']
    m.loc[idx, 'g2'] = str(i2) + ': none'
    m.loc[idx, 'g_cage_line'] = m.loc[idx, 'g1']
    m.loc[idx, 'g_generation'] = m.loc[idx, 'g1']

    # 6 German controls c57bl6/j
    idx = m[(m['genotype'] == 'c57bl6/j') & 
            (m['iacuc_institute'] == 'ZETT') &
            (m['geo_loc_name'] == 'Germany:Duesseldorf')].index
    i1 += 1
    i2 += 1
    m.loc[idx, 'g1'] = str(i1) + ': ' + m.loc[idx, 'genotype'] + '\ncontrol' + m.loc[idx, 'geo_loc_name']
    m.loc[idx, 'g2'] = str(i2) + ': none'
    m.loc[idx, 'g_cage_line'] = m.loc[idx, 'g1']
    m.loc[idx, 'g_generation'] = m.loc[idx, 'g1']

    # f5 to p1 c57bl6/j
    idx = m[(m['genotype'] == 'c57bl6/j') & 
            (m['iacuc_institute'] == 'LANUV')].index
    i1 += 1
    m.loc[idx, 'g1'] = str(i1) + ': ' + m.loc[idx, 'genotype']
    for generation in reversed(order_generations):
        i2 += 1
        idx_gen = list(set(idx) & set(m[m['generation'] == generation].index))
        m.loc[idx_gen, 'g2'] = str(i2) + ": " + generation
        m.loc[idx, 'g_cage_line'] = m.loc[idx, 'cage_line']
        m.loc[idx, 'g_generation'] = m.loc[idx, 'generation']


    # 6 recipient mice
    idx = m[(m['genotype'] == 'b6cf1')].index
    i1 += 1
    i2 += 1
    m.loc[idx, 'g1'] = str(i1) + ': ' + m.loc[idx, 'genotype'] + "\nrecipient mice"
    m.loc[idx, 'g2'] = str(i2) + ': none'
    m.loc[idx, 'g_cage_line'] = m.loc[idx, 'g1']
    m.loc[idx, 'g_generation'] = m.loc[idx, 'g1']


    # f5 to p1 balb/c
    idx = m[(m['genotype'] == 'balb/c') & 
            (m['iacuc_institute'] == 'LANUV')].index
    i1 += 1
    m.loc[idx, 'g1'] = str(i1) + ': ' + m.loc[idx, 'genotype']
    for generation in order_generations:
        i2 += 1
        idx_gen = list(set(idx) & set(m[m['generation'] == generation].index))
        m.loc[idx_gen, 'g2'] = str(i2) + ": " + generation
        m.loc[idx, 'g_cage_line'] = m.loc[idx, 'cage_line']
        m.loc[idx, 'g_generation'] = m.loc[idx, 'generation']

    # 6 German controls balb/c
    idx = m[(m['genotype'] == 'balb/c') & 
            (m['iacuc_institute'] == 'ZETT') &
            (m['geo_loc_name'] == 'Germany:Duesseldorf')].index
    i1 += 1
    i2 += 1
    m.loc[idx, 'g1'] = str(i1) + ': ' + m.loc[idx, 'genotype'] + '\ncontrol' + m.loc[idx, 'geo_loc_name']
    m.loc[idx, 'g2'] = str(i2) + ': none'
    m.loc[idx, 'g_cage_line'] = m.loc[idx, 'g1']
    m.loc[idx, 'g_generation'] = m.loc[idx, 'g1']

    # 6 French controls balb/c
    idx = m[(m['genotype'] == 'balb/c') & 
            (m['iacuc_institute'] == 'ZETT') &
            (m['geo_loc_name'] == 'France:Janvier')].index
    i1 += 1
    i2 += 1
    m.loc[idx, 'g1'] = str(i1) + ': ' + m.loc[idx, 'genotype'] + '\ncontrol ' + m.loc[idx, 'geo_loc_name']
    m.loc[idx, 'g2'] = str(i2) + ': none'
    m.loc[idx, 'g_cage_line'] = m.loc[idx, 'g1']
    m.loc[idx, 'g_generation'] = m.loc[idx, 'g1']

    minread = 60*309*0.5
    if subset == 'skin':
        minread = 10*309*0.5
    p_orig = plotTaxonomy(res_all['rarefaction']['results'], m.sort_values(by='cage_line'), file_taxonomy=res_all['taxonomy']['RDP']['results']['Taxon'],
                          group_l1='g1', group_l2='g2', print_meanrelabunances=True, grayscale=True, minreadnr=minread,
                          rank='Genus', 
                          min_abundance_grayscale=0.01,
                          sample_label_column="cage_line",
                          print_sample_labels=True,
                          #fct_aggregate=np.mean
                         )
    p_orig[0].get_axes()[0].set_title("all %s samples" % subset)
    p_orig[0].set_size_inches((40, 5))

    p_orig[0].savefig('Generated/tax_barplot_%s.svg' % subset)
    _ = emperor(m[m['host_scientific_name'] != 'callithrix jacchus'], res_all['beta_diversity']['results'], fp_results="./Generated/Emperor/", infix='_333%s_' % subset, dry=False, wait=True)

# %% [code]
plot_taxonomy_emperor(prj_data, res_all, subset='gut')

# %% [markdown]
The very sampe Düsseldorf control skin samples have been sequences both in batch 2 and 3. Although avg. read count in batch 2 is higher (7056.416667 vs 1716.166667) I decide to only go with batch 3 reads, as all other skin samples also have been exclusively sequenced in batch 3!

# %% [code]
prj_data['metadata']['iacuc_institute'].value_counts()

# %% [code]
idx = prj_data['metadata'][(prj_data['metadata']['sample_type'] == 'skin of external ear') & 
                           (prj_data['metadata']['batch'] != '2') & 
                           (prj_data['metadata']['iacuc_institute'] == 'ZETT')].index
prj_data['metadata'].loc[list(set(idx) & set(res_all_skin['rarefaction']['results'].columns))].groupby(['genotype','cage_id']).size()#,'genotype', 'cage_id']).size()

# %% [code]
x = plot_taxonomy_emperor(prj_data, res_all_skin, subset='skin')

# %% [code]
plot_temporal(**comp_temporal(meta[meta['sex'] == 'female'], {m: dm for (m,dm) in res_all['beta_diversity']['results'].items() if m == 'weighted_unifrac'}, True),
              title='only female mice')

# %% [markdown]
## rarefie mice

# %% [code]
def oppSex(sex):
    if sex == 'male':
        return 'female'
    else:
        return 'male'

depth = 8  # as all but p1 balb/c has at least 9 cages and we target for even sex distribution, i.e. 4 male, 4 female
idx_mice = dict()
for (genotype, generation), g in meta.groupby(['genotype', 'generation']):
    if generation == 'rm':
        idx_mice[(genotype, generation)] = set(g['host_subject_id'].unique())
    else:
        print(genotype, generation)
        target_no_sex = dict()
        target_no_sex['male'] = min(g['sex'].value_counts()['male'], int(depth/2))
        target_no_sex['female'] = min(g['sex'].value_counts()['female'], int(depth/2))
        for sex in ['male', 'female']:
            if target_no_sex[sex] < int(depth/2):
                target_no_sex[oppSex(sex)] = depth - target_no_sex['male']
                if target_no_sex[oppSex(sex)] > g['sex'].value_counts()[oppSex(sex)]:
                    print("Warning: insufficient number of mice in %s %s" % (genotype, generation), file=sys.stderr)
                    display(g['sex'].value_counts())
        
        for sex, g_sex in g.groupby('sex'):
            choices = pd.Series(combinations(g_sex['host_subject_id_stefan'].unique(), min(g_sex['host_subject_id_stefan'].unique().shape[0], target_no_sex[sex]))).to_frame()
            choices['div_cage_lines'] = choices[0].apply(lambda x: meta[meta['host_subject_id_stefan'].isin(x)]['cage_line'].unique().shape[0])
            #print("    ", choices.shape[0])
            max_cage_lines = choices['div_cage_lines'].max()
            choices = choices[choices['div_cage_lines'] == max_cage_lines]
            #print("    ", choices.shape[0])
            choices['div_cage_ids'] = choices[0].apply(lambda x: meta[meta['host_subject_id_stefan'].isin(x)]['cage_id'].unique().shape[0])
            max_cage_ids = choices['div_cage_ids'].max()
            choices = choices[choices['div_cage_ids'] == max_cage_ids]
            #print("    ", choices.shape[0])
            
            choice = choices[0].sample(1, random_state=42)
            if (genotype, generation) not in idx_mice:
                idx_mice[(genotype, generation)] = set(choice.iloc[0])
            else:
                idx_mice[(genotype, generation)] |= set(choice.iloc[0])
            #print("  ", sex, choices.shape)
            
        print("  ", target_no_sex['male'], target_no_sex['female'])

# %% [code]
ids = []
for l in idx_mice.values():
    ids.extend(l)
# add controls
ids.extend(list(prj_data['metadata'][(prj_data['metadata']['sample_type'] == 'colon content') &
                     (prj_data['metadata']['host_scientific_name'] != 'callithrix jacchus') &
                     (prj_data['metadata']['iacuc_institute'] == 'ZETT')]['host_subject_id_stefan'].values))
m = prj_data['metadata'][prj_data['metadata']['host_subject_id_stefan'].isin(ids)]
m = m[m['sample_type'] == 'colon content']
display(m.groupby(['iacuc_institute']).size())
res_rare = comp_temporal(m, 
                         {m: dm for (m,dm) in res_all['beta_diversity']['results'].items() if m == 'weighted_unifrac'}, True)
plot_temporal(**res_rare, title='rarefied to 8 mice')

# %% [code]
metaSkin = prj_data['metadata'][(prj_data['metadata']['sample_type'] == 'skin of external ear') & 
                                (prj_data['metadata']['host_scientific_name'] != 'callithrix jacchus')]
#exclude data without sequences
metaSkin = metaSkin.loc[list(set(metaSkin.index) & set(res_all_skin['rarefaction']['results'].columns)),:]
metaSkin.fillna('?').groupby(['genotype','generation','iacuc_institute',]).size()

# %% [code]
x = comp_temporal(metaSkin, res_all_skin['beta_diversity']['results'], True)

# %% [code]
plot_temporal(**x)

# %% [markdown]
# Quantify Stress
## animals per cage --> makes a difference!
### gut

# %% [code]
stress = prj_data['metadata'][(prj_data['metadata']['sample_type'] == 'colon content') & 
                              (prj_data['metadata']['host_scientific_name'] != 'callithrix jacchus') & 
                              (prj_data['metadata']['iacuc_institute'] == 'LANUV')].groupby(['genotype', 'generation', 'cage_line', 'cage_id', 'sex']).size().reset_index()
stress = stress.rename(columns={0: 'animals_per_cage'})
stress['animals_per_cage_cat'] = stress['animals_per_cage'].apply(lambda x: {
    1: 'few (1-3 mice)',
    2: 'few (1-3 mice)',
    3: 'few (1-3 mice)',
    4: 'many (4-5 mice)',
    5: 'many (4-5 mice)'}.get(x,x))    
if 'animals_per_cage' not in prj_data['metadata']:
    prj_data['metadata'] = prj_data['metadata'].reset_index().merge(stress, left_on=['genotype','generation','cage_line','cage_id','sex'], right_on=['genotype','generation','cage_line','cage_id','sex'], how='left').set_index('sample_name')
print(stress.shape[0])
stress.head()

# %% [code]
dists = []
for metric in res_all['beta_diversity']['results'].keys():
    dm = res_all['beta_diversity']['results'][metric].to_data_frame()
    for cage_id, g in prj_data['metadata'][(prj_data['metadata']['sample_type'] == 'colon content') & 
                                  (prj_data['metadata']['host_scientific_name'] != 'callithrix jacchus') & 
                                  (prj_data['metadata']['iacuc_institute'] == 'LANUV') & 
                                  (prj_data['metadata']['generation'] != 'rm')].groupby('cage_id'):
        num_animals = len(g['host_subject_id_stefan'].unique())
        assert g.shape[0] == num_animals
        idx = list(set(dm.index) & set(g.index))
        dm_sub = dm.loc[idx, idx]
        if True:
            for i in range(dm_sub.shape[0]):
                for j in range(dm_sub.shape[1]):
                    if i <= j:
                        dm_sub.iloc[i,j] = np.nan
        res = dm_sub.stack().dropna().reset_index().rename(columns={0: 'dist'})
        res['animals_per_cage'] = str(num_animals)
        res['metric'] = metric
        for field in ['genotype', 'cage_line', 'sex', 'generation', 'cage_id', 'animals_per_cage_cat']:
            res[field] = g[field].unique()[0]
        dists.append(res)
dists = pd.concat(dists)
print(dists.shape)
dists.head()

# %% [code]
data = prj_data['metadata'][(prj_data['metadata']['sample_type'] == 'colon content') & 
                            (prj_data['metadata']['host_scientific_name'] != 'callithrix jacchus') & 
                            (prj_data['metadata']['iacuc_institute'] == 'LANUV') & 
                            (prj_data['metadata']['generation'] != 'rm')]
data = data.loc[list(set(data.index) & set(res_all['rarefaction']['results'].columns))]

# %% [code]
sns.set_style("white")
fig, axes = plt.subplots(1,3, figsize=(6+3,5))
for i, genotype in enumerate(['c57bl6/j', 'balb/c']):
#for i, (genotype, g) in enumerate(data.sort_values(by='cage_line', ascending=True).groupby('genotype')):
    g = data[data['genotype'] == genotype]
    ax = axes[i]
    ax.axhspan(0.8, 3.2, facecolor=colors_animals_per_cage['few (1-3 mice)'], alpha=0.3, zorder=0)
    ax.axhspan(3.8, 5.2, facecolor=colors_animals_per_cage['many (4-5 mice)'], alpha=0.3, zorder=0)
    
    order_cl = [l for l in order_cagelines if l in g['cage_line'].unique()]
    sns.stripplot(data=g, y='animals_per_cage', x='cage_line', ax=ax, color="black", jitter=0.3, size=10, order=order_cl, palette=colors, edgecolor='white', linewidth=1)
    sns.boxplot(data=g, y='animals_per_cage', x='cage_line', ax=ax, order=order_cl, palette={n:alpha_blending(c,0.4) for n,c in colors.items()})

    annotator = None
    test = 'Mann-Whitney'
    if genotype == 'balb/c':
        annotator = Annotator(ax, [
            ("BALB/c_line4","BALB/c_line5"),
            ("BALB/c_line6","BALB/c_line5"),
            ("BALB/c_line4","BALB/c_line6"),
        ], data=g, y='animals_per_cage', x='cage_line', order=order_cl)
        test = 'Mann-Whitney-ls'
    elif genotype == 'c57bl6/j':
        annotator = Annotator(ax, [
            ("C57BL6/J_line1","C57BL6/J_line2"),
            ("C57BL6/J_line2","C57BL6/J_line3"),
            ("C57BL6/J_line1","C57BL6/J_line3"),
        #    ("BALB/c_line6","BALB/c_line5"),
        ], data=g, y='animals_per_cage', x='cage_line', order=order_cl)
        test = 'Mann-Whitney-gt'
    if annotator is not None:
        annotator.configure(test=test, text_format='star', loc='outside', comparisons_correction="fdr_bh", correction_format="default")
        annotator.apply_and_annotate()

    ax.set_xticklabels(['%s\n(%ii,%ic)' % ({"C57BL6/J_line1":"B1",
                                            "C57BL6/J_line2":"B2",
                                            "C57BL6/J_line3":"B3",
                                            "BALB/c_line4": "C1",
                                            "BALB/c_line5": "C2",
                                            "BALB/c_line6": "C3"}.get(cl.get_text(), cl.get_text()), g.groupby('cage_line').size().loc[cl.get_text()], g[g['cage_line'] == cl.get_text()]['cage_id'].value_counts().shape[0]) for cl in ax.get_xticklabels()])
    ax.set_xlabel("")
    ax.set_ylabel("mice per cage")

    
ax = axes[i+1]
if False:
    sns.violinplot(data=dists[dists['metric'] == 'weighted_unifrac'], y='dist', hue='animals_per_cage_cat', x='genotype', ax=ax, palette=colors_animals_per_cage)
    annotator = Annotator(ax, [
        (("balb/c", list(colors_animals_per_cage.keys())[0],("balb/c", list(colors_animals_per_cage.keys())[-1]))),
        (("c57bl6/j", list(colors_animals_per_cage.keys())[0]),("c57bl6/j", list(colors_animals_per_cage.keys())[-1])),
    ], data=dists[dists['metric'] == 'weighted_unifrac'], y='dist', hue='animals_per_cage_cat', x='genotype')
    annotator.configure(test='Mann-Whitney', text_format='star', loc='outside', comparisons_correction="fdr_bh", correction_format="default")
else:
    sns.boxplot(data=dists[dists['metric'] == 'weighted_unifrac'], y='dist', x='animals_per_cage_cat', ax=ax, palette=colors_animals_per_cage)
    sns.stripplot(data=dists[dists['metric'] == 'weighted_unifrac'], y='dist', x='animals_per_cage_cat', ax=ax, palette=colors_animals_per_cage, edgecolor='white', linewidth=1)
    annotator = Annotator(ax, [
        (list(colors_animals_per_cage.keys())[0],list(colors_animals_per_cage.keys())[-1]),
    ], data=dists[dists['metric'] == 'weighted_unifrac'], y='dist', x='animals_per_cage_cat')
    annotator.configure(test='Mann-Whitney', text_format='star', loc='outside', comparisons_correction="fdr_bh", correction_format="default")
ax.set_xticklabels(['%s\n%ii\n%ic' % (
    cat.get_text(), 
    data['animals_per_cage_cat'].value_counts().loc[cat.get_text()], 
    data[data['animals_per_cage_cat'] == cat.get_text()]['cage_id'].value_counts().shape[0]
) for cat in ax.get_xticklabels()])
ax.set_ylabel("weighted UniFrac")
ax.set_xlabel("mice per cage")
annotator.apply_and_annotate()
fig.suptitle("stress: mice per cage")

#fig.savefig('Generated/stress_gut.svg')

# %% [markdown]
### skin

# %% [code]
stress = prj_data['metadata'][(prj_data['metadata']['sample_type'] == 'skin of external ear') & 
                              (prj_data['metadata']['host_scientific_name'] != 'callithrix jacchus') & 
                              (prj_data['metadata']['iacuc_institute'] == 'LANUV')].groupby(['genotype', 'generation', 'cage_line', 'cage_id', 'sex']).size().reset_index()
stress = stress.rename(columns={0: 'animals_per_cage'})
stress['animals_per_cage_cat'] = stress['animals_per_cage'].apply(lambda x: {
    1: 'few (1-3 mice)',
    2: 'few (1-3 mice)',
    3: 'few (1-3 mice)',
    4: 'many (4-5 mice)',
    5: 'many (4-5 mice)'}.get(x,x))    
if 'animals_per_cage' not in prj_data['metadata']:
    prj_data['metadata'] = prj_data['metadata'].reset_index().merge(stress, left_on=['genotype','generation','cage_line','cage_id','sex'], right_on=['genotype','generation','cage_line','cage_id','sex'], how='left').set_index('sample_name')
print(stress.shape[0])
stress.head()

# %% [code]
dists = []
for metric in res_all_skin['beta_diversity']['results'].keys():
    dm = res_all_skin['beta_diversity']['results'][metric].to_data_frame()
    for cage_id, g in prj_data['metadata'][(prj_data['metadata']['sample_type'] == 'skin of external ear') & 
                                  (prj_data['metadata']['host_scientific_name'] != 'callithrix jacchus') & 
                                  (prj_data['metadata']['iacuc_institute'] == 'LANUV') & 
                                  (prj_data['metadata']['generation'] != 'rm')].groupby('cage_id'):
        num_animals = len(g['host_subject_id_stefan'].unique())
        assert g.shape[0] == num_animals
        idx = list(set(dm.index) & set(g.index))
        dm_sub = dm.loc[idx, idx]
        if True:
            for i in range(dm_sub.shape[0]):
                for j in range(dm_sub.shape[1]):
                    if i <= j:
                        dm_sub.iloc[i,j] = np.nan
        res = dm_sub.stack().dropna().reset_index().rename(columns={0: 'dist'})
        res['animals_per_cage'] = str(num_animals)
        res['metric'] = metric
        for field in ['genotype', 'cage_line', 'sex', 'generation', 'cage_id', 'animals_per_cage_cat']:
            res[field] = g[field].unique()[0]
        dists.append(res)
dists = pd.concat(dists)
print(dists.shape)
dists.head()

data = prj_data['metadata'][(prj_data['metadata']['sample_type'] == 'skin of external ear') & 
                            (prj_data['metadata']['host_scientific_name'] != 'callithrix jacchus') & 
                            (prj_data['metadata']['iacuc_institute'] == 'LANUV') & 
                            (prj_data['metadata']['generation'] != 'rm')]
data = data.loc[list(set(data.index) & set(res_all_skin['rarefaction']['results'].columns))]

# %% [code]
sns.set_style("white")
fig, axes = plt.subplots(1,3, figsize=(6+3,5))
for i, genotype in enumerate(['c57bl6/j', 'balb/c']):
#for i, (genotype, g) in enumerate(data.sort_values(by='cage_line', ascending=True).groupby('genotype')):
    g = data[data['genotype'] == genotype]
    ax = axes[i]
    ax.axhspan(0.8, 3.2, facecolor=colors_animals_per_cage['few (1-3 mice)'], alpha=0.3, zorder=0)
    ax.axhspan(3.8, 5.2, facecolor=colors_animals_per_cage['many (4-5 mice)'], alpha=0.3, zorder=0)
    
    order_cl = [l for l in order_cagelines if l in g['cage_line'].unique()]
    sns.stripplot(data=g, y='animals_per_cage', x='cage_line', ax=ax, color="black", jitter=0.3, size=10, order=order_cl, palette=colors, edgecolor='white', linewidth=1)
    sns.boxplot(data=g, y='animals_per_cage', x='cage_line', ax=ax, order=order_cl, palette={n:alpha_blending(c,0.4) for n,c in colors.items()})

    annotator = None
    test = 'Mann-Whitney'
    if genotype == 'balb/c':
        annotator = Annotator(ax, [
            ("BALB/c_line4","BALB/c_line5"),
            ("BALB/c_line6","BALB/c_line5"),
            ("BALB/c_line4","BALB/c_line6"),
        ], data=g, y='animals_per_cage', x='cage_line', order=order_cl)
        test = 'Mann-Whitney-ls'
    elif genotype == 'c57bl6/j':
        annotator = Annotator(ax, [
            ("C57BL6/J_line1","C57BL6/J_line2"),
            ("C57BL6/J_line2","C57BL6/J_line3"),
            ("C57BL6/J_line1","C57BL6/J_line3"),
        #    ("BALB/c_line6","BALB/c_line5"),
        ], data=g, y='animals_per_cage', x='cage_line', order=order_cl)
        test = 'Mann-Whitney-gt'
    if annotator is not None:
        annotator.configure(test=test, text_format='star', loc='outside', comparisons_correction="fdr_bh", correction_format="default")
        annotator.apply_and_annotate()

    ax.set_xticklabels(['%s\n(%ii,%ic)' % ({"C57BL6/J_line1":"B1",
                                            "C57BL6/J_line2":"B2",
                                            "C57BL6/J_line3":"B3",
                                            "BALB/c_line4": "C1",
                                            "BALB/c_line5": "C2",
                                            "BALB/c_line6": "C3"}.get(cl.get_text(), cl.get_text()), g.groupby('cage_line').size().loc[cl.get_text()], g[g['cage_line'] == cl.get_text()]['cage_id'].value_counts().shape[0]) for cl in ax.get_xticklabels()])
    ax.set_xlabel("")
    ax.set_ylabel("mice per cage")

    
ax = axes[i+1]
if False:
    sns.violinplot(data=dists[dists['metric'] == 'weighted_unifrac'], y='dist', hue='animals_per_cage_cat', x='genotype', ax=ax, palette=colors_animals_per_cage)
    annotator = Annotator(ax, [
        (("balb/c", list(colors_animals_per_cage.keys())[0],("balb/c", list(colors_animals_per_cage.keys())[-1]))),
        (("c57bl6/j", list(colors_animals_per_cage.keys())[0]),("c57bl6/j", list(colors_animals_per_cage.keys())[-1])),
    ], data=dists[dists['metric'] == 'weighted_unifrac'], y='dist', hue='animals_per_cage_cat', x='genotype')
    annotator.configure(test='Mann-Whitney', text_format='star', loc='outside', comparisons_correction="fdr_bh", correction_format="default")
else:
    sns.boxplot(data=dists[dists['metric'] == 'weighted_unifrac'], y='dist', x='animals_per_cage_cat', ax=ax, palette=colors_animals_per_cage)
    sns.stripplot(data=dists[dists['metric'] == 'weighted_unifrac'], y='dist', x='animals_per_cage_cat', ax=ax, palette=colors_animals_per_cage, edgecolor='white', linewidth=1)
    annotator = Annotator(ax, [
        (list(colors_animals_per_cage.keys())[0],list(colors_animals_per_cage.keys())[-1]),
    ], data=dists[dists['metric'] == 'weighted_unifrac'], y='dist', x='animals_per_cage_cat')
    annotator.configure(test='Mann-Whitney', text_format='star', loc='outside', comparisons_correction="fdr_bh", correction_format="default")
ax.set_xticklabels(['%s\n%ii\n%ic' % (
    cat.get_text(), 
    data['animals_per_cage_cat'].value_counts().loc[cat.get_text()], 
    data[data['animals_per_cage_cat'] == cat.get_text()]['cage_id'].value_counts().shape[0]
) for cat in ax.get_xticklabels()])
ax.set_ylabel("weighted UniFrac")
ax.set_xlabel("mice per cage")
annotator.apply_and_annotate()
fig.suptitle("stress: mice per cage")

fig.savefig('Generated/stress_skin.svg')

# %% [code]
fig, axes = plt.subplots(1,1)
sns.violinplot(data=dists[dists['metric'] == 'weighted_unifrac'], y='dist', x='animals_per_cage', ax=axes, order=["2","3","4","5"])

annotator = Annotator(axes, [
    ("2","3"),
    #('2','4'),
    #('2','5'),
    ('3','4'),
    #('3','5'),
    ('4','5'),
], data=dists[dists['metric'] == 'weighted_unifrac'], y='dist', x='animals_per_cage', order=["2","3","4","5"])
annotator.configure(test='Mann-Whitney', text_format='star', loc='outside', comparisons_correction="fdr_bh", correction_format="default")
annotator.apply_and_annotate()


# %% [markdown]
## birth offset

# %% [code]
x = prj_data['metadata'].groupby('cage_id')['birth_timestamp'].unique().apply(len)
x[x > 1]

# except rm1/rm2, a cage always houses mice born at the same day!

# %% [markdown]
### are cousins born on same day closer to each other than cousins born on different days? --> yes, but only in unweighted UniFrac

# %% [code]
dists = []
for msid, g in prj_data['metadata'][(prj_data['metadata']['sample_type'] == 'colon content') & 
                     (prj_data['metadata']['host_scientific_name'] != 'callithrix jacchus') & 
                     (prj_data['metadata']['iacuc_institute'] == 'LANUV')].groupby('mothers_subject_ids_stefan'):
    #print(msid, g.shape[0], g['cage_id'].value_counts().shape[0])
    
    for metric in res_all['beta_diversity']['results'].keys():
        dm = res_all['beta_diversity']['results'][metric].to_data_frame()
        g = g.loc[list(set(g.index) & set(dm.index))]
        for (cage_a, cage_b) in combinations(g['cage_id'].dropna().unique(), 2):
            for idx_a in g[g['cage_id'] == cage_a].index:
                for idx_b in g[g['cage_id'] == cage_b].index:
                    dists.append({'dist': dm.loc[idx_a,idx_b],
                                  'metric': metric,
                                  'same_birth': g.loc[idx_a, 'birth_timestamp'] == g.loc[idx_b, 'birth_timestamp'],
                                 })
dists=pd.DataFrame(dists)

# %% [code]
fig, axes = plt.subplots(1,1)

sns.barplot(data=dists, y='dist', x='metric', hue='same_birth', ax=axes)

annotator = Annotator(axes, [
    ((metric, False), (metric, True))
    for metric in dists['metric'].unique()
], data=dists, y='dist', x='metric', hue='same_birth')
annotator.configure(test='Mann-Whitney', text_format='star', loc='outside', comparisons_correction="fdr_bh", correction_format="default")
annotator.apply_and_annotate()

# %% [code]
birth_offset = prj_data['metadata'][(prj_data['metadata']['sample_type'] == 'colon content') & 
                                    (prj_data['metadata']['host_scientific_name'] != 'callithrix jacchus') & 
                                    (prj_data['metadata']['iacuc_institute'] == 'LANUV')].groupby(['genotype', 'generation', 'cage_line'])['birth_timestamp'].unique().apply(lambda x: sorted(list(map(lambda d: datetime.strptime(d, '%Y-%m-%d'), x)))).apply(lambda dates: (dates[-1] - dates[0]).days).reset_index()
birth_offset = birth_offset.rename(columns={'birth_timestamp': 'birth_offset_in_days'})
if 'birth_offset_in_days' not in prj_data['metadata'].columns:
    prj_data['metadata'] = prj_data['metadata'].reset_index().merge(birth_offset, left_on=['genotype','generation','cage_line'], right_on=['genotype','generation','cage_line'], how='left').set_index('sample_name')
birth_offset.head()

# %% [code]
data = prj_data['metadata'][(prj_data['metadata']['sample_type'] == 'colon content') & 
                            (prj_data['metadata']['host_scientific_name'] != 'callithrix jacchus') & 
                            (prj_data['metadata']['iacuc_institute'] == 'LANUV') & 
                            (prj_data['metadata']['generation'] != 'rm')]

# %% [code]
dists = []
metric = 'weighted_unifrac'
dm = res_all['beta_diversity']['results'][metric].to_data_frame()
for (generation, cage_line), g in data.groupby(['generation','cage_line']):
    for (a,b) in combinations(g.index, 2):
        if (a in dm.index) & (b in dm.index):
            dist = dm.loc[a,b]
            diff = abs((datetime.strptime(g.loc[a, 'birth_timestamp'], '%Y-%m-%d') - \
                        datetime.strptime(g.loc[b, 'birth_timestamp'], '%Y-%m-%d')).days)
            dists.append({'dist': dist,
                          'metric': metric,
                          'generation': generation,
                          'genotype': g.loc[a, 'genotype'],
                          'cage_line': cage_line,
                          'birth_offset_in_days': diff,
                          'cat': 'diff' if diff != 0 else 'same'})
dists = pd.DataFrame(dists)
dists

# %% [code]
fig, axes = plt.subplots(1,1)
sns.boxplot(data=dists, y='dist', hue='cat', x='genotype', ax=axes)

annotator = Annotator(axes, [
    (("balb/c","same"), ("balb/c", "diff")),
    (("c57bl6/j","same"), ("c57bl6/j", "diff"))
], data=dists, y='dist', x='genotype', hue='cat')
annotator.configure(test='Mann-Whitney', text_format='star', loc='outside', comparisons_correction="fdr_bh", correction_format="default")
annotator.apply_and_annotate()

# %% [code]
data.groupby(['genotype','generation','cage_line','father_subject_id_stefan'])['birth_timestamp'].value_counts()

# %% [code]
fig, axes = plt.subplots(1,3, figsize=(12+6,5))
for i, genotype in enumerate(['c57bl6/j', 'balb/c']):
    g = data[data['genotype'] == genotype]
    ax = axes[i]
    #ax.axhspan(0.8, 3.2, facecolor=colors_animals_per_cage['few (1-3 mice)'], alpha=0.3)
    #ax.axhspan(3.8, 5.2, facecolor=colors_animals_per_cage['many (4-5 mice)'], alpha=0.3)
    
    order_cl = [l for l in order_cagelines if l in g['cage_line'].unique()]
    sns.stripplot(data=g, y='birth_offset_in_days', x='cage_line', ax=ax, color="black", jitter=0.3, size=5, order=order_cl)
    sns.violinplot(data=g, y='birth_offset_in_days', x='cage_line', ax=ax, order=order_cl)

    #annotator = None
    #test = 'Mann-Whitney'
    if genotype == 'balb/c':
        annotator = Annotator(ax, [
            ("BALB/c_line4","BALB/c_line5"),
            ("BALB/c_line6","BALB/c_line5"),
            ("BALB/c_line4","BALB/c_line6"),
        ], data=g, y='birth_offset_in_days', x='cage_line', order=order_cl)
        test = 'Mann-Whitney'
    elif genotype == 'c57bl6/j':
        annotator = Annotator(ax, [
            ("C57BL6/J_line1","C57BL6/J_line2"),
            ("C57BL6/J_line2","C57BL6/J_line3"),
            ("C57BL6/J_line1","C57BL6/J_line3"),
        ], data=g, y='birth_offset_in_days', x='cage_line', order=order_cl)
        test = 'Mann-Whitney'
    if annotator is not None:
        annotator.configure(test=test, text_format='star', loc='outside', comparisons_correction="fdr_bh", correction_format="default")
        annotator.apply_and_annotate()

    ax.set_xticklabels(['%s\n%ii\n%ic' % (cl.get_text(), g.groupby('cage_line').size().loc[cl.get_text()], g[g['cage_line'] == cl.get_text()]['cage_id'].value_counts().shape[0]) for cl in ax.get_xticklabels()])
    ax.set_xlabel("")
    #ax.set_ylabel("mice per cage")

# %% [code]
_ = plotNetworks('birth_offset_in_days', prj_data['metadata'][(prj_data['metadata']['sample_type'] == 'colon content') 
                                                         # & (prj_data['metadata']['genotype'] != 'c57bl6/j') 
                                                         # (prj_data['metadata']['sex'] == 'female') &
                                                         # (prj_data['metadata']['generation'].isin(['f1','f2','f3','f4','f5']))
                                                         ], res_all['alpha_diversity']['results'], res_all['beta_diversity']['results'], permutations=9999)

# %% [code]
import skbio
r = detect_distant_groups(res_all['beta_diversity']['results']['weighted_unifrac'], 
                      'weighted_unifrac', 
                      prj_data['metadata'][(prj_data['metadata']['sample_type'] == 'colon content') &
                                                          (prj_data['metadata']['genotype'] == 'c57bl6/j') 
                                                         # (prj_data['metadata']['sex'] == 'female') &
                                                         # (prj_data['metadata']['generation'].isin(['f1','f2','f3','f4','f5']))
                                                         ]['birth_offset_in_days'],
                      fct_test=skbio.stats.distance.permanova,
                     )['network']

# %% [code]
r[4.0][18.0]

# %% [code]
x = prj_data['metadata'][(prj_data['metadata']['sample_type'] == 'colon content') &
                                                          (prj_data['metadata']['genotype'] == 'c57bl6/j') 
                                                         # (prj_data['metadata']['sex'] == 'female') &
                                                         # (prj_data['metadata']['generation'].isin(['f1','f2','f3','f4','f5']))
                                                         ]
x = x[x['birth_offset_in_days'].isin([4.0,18.0])]
dm = res_all['beta_diversity']['results']['weighted_unifrac'].filter(x.index)
m = x.loc[list(set(x.index) & set(dm.ids)),:]

# %% [code]
dm.shape

# %% [code]
m['birth_offset_in_days'].value_counts()

# %% [code]
skbio.stats.distance.permanova(dm, m['birth_offset_in_days'].loc[list(dm.ids)])

# %% [code]
dm.write("test.dm.csv")

# %% [code]
m['birth_offset_in_days'].loc[list(dm.ids)].apply(lambda x: 'bdo_%i' % x).to_csv('test.meta.csv', sep="\t")

# %% [markdown]
## mothers deliver in sync --> no difference

# %% [code]
delivery = prj_data['metadata'][(prj_data['metadata']['sample_type'] == 'colon content') & 
                                (prj_data['metadata']['host_scientific_name'] != 'callithrix jacchus') & 
                                (prj_data['metadata']['iacuc_institute'] == 'LANUV')]

dists = []
for metric in res_all['beta_diversity']['results'].keys():
    dm = res_all['beta_diversity']['results'][metric].to_data_frame()
    
    for (genotype,cage_line,generation,mothers_subject_ids_stefan), g in delivery.groupby(['genotype','cage_line','generation','mothers_subject_ids_stefan']):
        sync_delivery = len(g['birth_timestamp'].unique()) == 1
        idx_mothers = list(set(dm.index) & set(delivery[delivery['host_subject_id_stefan'].isin(mothers_subject_ids_stefan.split('|'))].index))
        if len(idx_mothers) == 2:
            dists.append({'dist': dm.loc[idx_mothers[0], idx_mothers[-1]],
                          'sync_delivery': sync_delivery,
                          'metric': metric})

dists = pd.DataFrame(dists)

# %% [code]
fig, axes = plt.subplots(1,1)

sns.barplot(data=dists, y='dist', x='metric', hue='sync_delivery', ax=axes)

annotator = Annotator(axes, [
    ((metric, False), (metric, True))
    for metric in dists['metric'].unique()
], data=dists, y='dist', x='metric', hue='sync_delivery')
annotator.configure(test='Mann-Whitney', text_format='star', loc='outside', comparisons_correction="fdr_bh", correction_format="default")
annotator.apply_and_annotate()

# %% [markdown]
# food / beeding as taxa sources

# %% [code]
_ = plotTaxonomy(res_all['counts_plantsStillIn'].loc[:, prj_data['metadata'][prj_data['metadata']['sample_type'].isin(['control bedding', 'control food'])].index], prj_data['metadata'], file_taxonomy=res_all['taxonomy']['RDP']['results']['Taxon'],
                 rank='Genus', group_l1='sample_type', print_meanrelabunances=True)

# %% [markdown]
# maternal legacy
## gut

# %% [code]
# collect subjects that gave birth (independent of genotype, generation, cage_line)
idx_mothers = {i for l in prj_data['metadata']['mothers_subject_ids_stefan'].dropna().apply(lambda x: x.split('|')).values for i in l}

#metric = 'bray_curtis'
metric = 'unweighted_unifrac'
dm = res_all['beta_diversity']['results'][metric].to_data_frame()
meta = prj_data['metadata'][(prj_data['metadata']['host_scientific_name'] != 'callithrix jacchus') & 
                            (prj_data['metadata']['sample_type'] == 'colon content')]
dists = []
for (generation, genotype), g in meta[pd.notnull(meta['mothers_subject_ids_stefan'])].groupby(['generation', 'genotype']):
    idx_prevGen_mothers = set(meta[meta['host_subject_id_stefan'].isin({i for l in g['mothers_subject_ids_stefan'].apply(lambda x: x.split('|')).values for i in l})].index)
    #idx_prevGen_mothers |= set(meta[meta['host_subject_id_stefan'].isin(list(g['father_subject_id_stefan'].unique()))].index)
    
    idx_prevGen_diffGenotype_mothers = set(meta[(meta['host_subject_id_stefan'].isin(idx_mothers)) & # only mothers
                                            (meta['generation'] == order_generations[order_generations.index(generation)-1]) & # previous generation
                                            (meta['genotype'] != genotype)].index) # the other genotype
    #print(generation, prevGen, genotype, g.shape[0], idx_prevGen_mothers)
    #display("HIR", genotype, .groupby(['generation','genotype']).size())
    
    d = dm.loc[list(set(dm.index) & idx_prevGen_mothers), list(set(g.index) & set(dm.index))].stack().reset_index().rename(columns={'level_0': 'idx_mother', 'level_1': 'idx_child', 0: metric})
    for f in ['cage_line', 'generation', 'genotype']:
        for gen in ['mother', 'child']:
            d = d.merge(meta[[f]], left_on='idx_%s' % gen, right_index=True, how='left').rename(columns={f: '%s_%s' % (f, gen)})
    d['maternal'] = d.apply(lambda x: 'true mother' if x['cage_line_mother'] == x['cage_line_child'] else 'mother other cage lineage', axis=1)
    dists.append(d)
    
    d = dm.loc[list(set(dm.index) & idx_prevGen_diffGenotype_mothers), list(set(g.index) & set(dm.index))].stack().reset_index().rename(columns={'level_0': 'idx_mother', 'level_1': 'idx_child', 0: metric})
    for f in ['cage_line', 'generation', 'genotype']:
        for gen in ['mother', 'child']:
            d = d.merge(meta[[f]], left_on='idx_%s' % gen, right_index=True, how='left').rename(columns={f: '%s_%s' % (f, gen)})
    d['maternal'] = 'mother other genotype'
    dists.append(d)
    
dists = pd.concat(dists)
dists['cat'] = dists['genotype_child'] + ": " + dists['generation_child']
dists = dists.sort_values(by='cat')
print(dists.shape)
nums = dists.groupby(['generation_child','genotype_child','maternal']).size()
display(nums)
dists.head()

# %% [code]
fig, axes = plt.subplots(1, 1, figsize=(1.9*len(dists['generation_child'].unique()),5))
ho = ['true mother', 'mother other cage lineage', 'mother other genotype']
o = ['c57bl6/j: f1', 'c57bl6/j: f2', 'c57bl6/j: f3', 'c57bl6/j: f4', 'c57bl6/j: f5', 
     'balb/c: f1', 'balb/c: f2', 'balb/c: f3', 'balb/c: f4', 'balb/c: f5']
sns.boxplot(data=dists, y=metric, x='cat', hue='maternal', ax=axes, hue_order=ho, order=o)

_ = axes.set_xticklabels(['%s\n%ii\n%ii\n%ii' % (t.get_text(), 
                                       nums.loc[t.get_text().split(': ')[-1], t.get_text().split(': ')[0], 'true mother'],
                                       nums.loc[t.get_text().split(': ')[-1], t.get_text().split(': ')[0], 'mother other cage lineage'],
                                       nums.loc[t.get_text().split(': ')[-1], t.get_text().split(': ')[0], 'mother other genotype'],
                                 ) 
                          for t in axes.get_xticklabels()])
axes.set_ylim((axes.get_ylim()[0], axes.get_ylim()[1]*1.1))

annotator = Annotator(axes, 
    [((cat, 'true mother'), (cat, 'mother other cage lineage')) for cat in dists['cat'].unique()] + \
    [((cat, 'mother other genotype'), (cat, 'mother other cage lineage')) for cat in dists['cat'].unique()], 
    data=dists, y=metric, x='cat', hue='maternal', hue_order=ho, order=o)
annotator.configure(test='Mann-Whitney', text_format='star', loc='outside', comparisons_correction="fdr_bh", correction_format="default")
_ = annotator.apply_and_annotate()

#fig.savefig('Generated/maternal_legacy_gut.svg')

# %% [code]
fig, axes = plt.subplots(1, 1, )

d = dists[(dists['maternal'] == 'mother other cage lineage') & (dists['maternal'] == 'mother other cage lineage')]
sns.barplot(data=d,
            y=metric, x='genotype_child', ax=axes)#['genotype_child'].value_counts()

annotator = Annotator(axes, 
    [('balb/c','c57bl6/j')], 
    data=d, y=metric, x='genotype_child')
annotator.configure(test='Mann-Whitney', text_format='star', loc='outside', comparisons_correction="fdr_bh", correction_format="default")
_ = annotator.apply_and_annotate()

#dists['maternal'].value_counts()

# %% [code]
metric = 'unweighted_unifrac'
dm = res_all['beta_diversity']['results'][metric].to_data_frame()
dists = []
for (genotype, generation), g in prj_data['metadata'][(prj_data['metadata']['host_scientific_name'] != 'callithrix jacchus') & 
                                                      (prj_data['metadata']['sample_type'] == 'colon content')].groupby(['genotype','generation']):
    if generation == 'rm':
        continue
    print(genotype, generation, g.shape[0])
    for (a,b) in combinations(list(set(g.index) & set(dm.index)), 2):
        if (g.loc[a,'cage_line'] != g.loc[b,'cage_line']):
            dists.append({metric: dm.loc[a,b],
                          'generation': generation,
                          'genotype': genotype,
                          'cls': ' vs. '.join(sorted(g.loc[[a,b],'cage_line'].values))})
dists = pd.DataFrame(dists)
dists.shape[0]

# %% [code]
fig, axes = plt.subplots(1,1)
sns.lineplot(data=dists.sort_values(by='generation', key=key_generations), y=metric, x='generation', sort=False, hue='cls', ax=axes, style='genotype')
#        sns.lineplot(data=g, y='dist', x='generation', hue='genotype', ax=ax, style=style, palette={'c57bl6/j': colors['c57bl6/j'],
axes.legend(bbox_to_anchor=(1.1, 1.05))

# %% [markdown]
## skin

# %% [code]
# collect subjects that gave birth (independent of genotype, generation, cage_line)
idx_mothers = {i for l in prj_data['metadata']['mothers_subject_ids_stefan'].dropna().apply(lambda x: x.split('|')).values for i in l}

#metric = 'bray_curtis'
metric = 'unweighted_unifrac'
dm = res_all_skin['beta_diversity']['results'][metric].to_data_frame()
meta = prj_data['metadata'][(prj_data['metadata']['host_scientific_name'] != 'callithrix jacchus') & 
                            (prj_data['metadata']['sample_type'] == 'skin of external ear')]
dists = []
for (generation, genotype), g in meta[pd.notnull(meta['mothers_subject_ids_stefan'])].groupby(['generation', 'genotype']):
    idx_prevGen_mothers = set(meta[meta['host_subject_id_stefan'].isin({i for l in g['mothers_subject_ids_stefan'].apply(lambda x: x.split('|')).values for i in l})].index)
    #idx_prevGen_mothers |= set(meta[meta['host_subject_id_stefan'].isin(list(g['father_subject_id_stefan'].unique()))].index)
    
    idx_prevGen_diffGenotype_mothers = set(meta[(meta['host_subject_id_stefan'].isin(idx_mothers)) & # only mothers
                                            (meta['generation'] == order_generations[order_generations.index(generation)-1]) & # previous generation
                                            (meta['genotype'] != genotype)].index) # the other genotype
    #print(generation, prevGen, genotype, g.shape[0], idx_prevGen_mothers)
    #display("HIR", genotype, .groupby(['generation','genotype']).size())
    
    d = dm.loc[list(set(dm.index) & idx_prevGen_mothers), list(set(g.index) & set(dm.index))].stack().reset_index().rename(columns={'level_0': 'idx_mother', 'level_1': 'idx_child', 0: metric})
    for f in ['cage_line', 'generation', 'genotype']:
        for gen in ['mother', 'child']:
            d = d.merge(meta[[f]], left_on='idx_%s' % gen, right_index=True, how='left').rename(columns={f: '%s_%s' % (f, gen)})
    d['maternal'] = d.apply(lambda x: 'true mother' if x['cage_line_mother'] == x['cage_line_child'] else 'mother other cage lineage', axis=1)
    dists.append(d)
    
    d = dm.loc[list(set(dm.index) & idx_prevGen_diffGenotype_mothers), list(set(g.index) & set(dm.index))].stack().reset_index().rename(columns={'level_0': 'idx_mother', 'level_1': 'idx_child', 0: metric})
    for f in ['cage_line', 'generation', 'genotype']:
        for gen in ['mother', 'child']:
            d = d.merge(meta[[f]], left_on='idx_%s' % gen, right_index=True, how='left').rename(columns={f: '%s_%s' % (f, gen)})
    d['maternal'] = 'mother other genotype'
    dists.append(d)
    
dists = pd.concat(dists)
dists['cat'] = dists['genotype_child'] + ": " + dists['generation_child']
dists = dists.sort_values(by='cat')
print(dists.shape)
nums = dists.groupby(['generation_child','genotype_child','maternal']).size()
display(nums)
dists.head()

# %% [code]
fig, axes = plt.subplots(1, 1, figsize=(1.9*len(dists['generation_child'].unique()),5))
ho = ['true mother', 'mother other cage lineage', 'mother other genotype']
o = ['c57bl6/j: f1', 'c57bl6/j: f2', 'c57bl6/j: f3', 'c57bl6/j: f4', 'c57bl6/j: f5', 
     'balb/c: f1', 'balb/c: f2', 'balb/c: f3', 'balb/c: f4', 'balb/c: f5']
sns.boxplot(data=dists, y=metric, x='cat', hue='maternal', ax=axes, hue_order=ho, order=o)

_ = axes.set_xticklabels(['%s\n%ii\n%ii\n%ii' % (t.get_text(), 
                                       nums.loc[t.get_text().split(': ')[-1], t.get_text().split(': ')[0], 'true mother'],
                                       nums.loc[t.get_text().split(': ')[-1], t.get_text().split(': ')[0], 'mother other cage lineage'],
                                       nums.loc[t.get_text().split(': ')[-1], t.get_text().split(': ')[0], 'mother other genotype'],
                                 ) 
                          for t in axes.get_xticklabels()])
axes.set_ylim((axes.get_ylim()[0], axes.get_ylim()[1]*1.1))

annotator = Annotator(axes, 
    [((cat, 'true mother'), (cat, 'mother other cage lineage')) for cat in dists['cat'].unique()] + \
    [((cat, 'mother other genotype'), (cat, 'mother other cage lineage')) for cat in dists['cat'].unique()], 
    data=dists, y=metric, x='cat', hue='maternal', hue_order=ho, order=o)
annotator.configure(test='Mann-Whitney', text_format='star', loc='outside', comparisons_correction="fdr_bh", correction_format="default")
_ = annotator.apply_and_annotate()

fig.savefig('Generated/maternal_legacy_skin.svg')

# %% [markdown]
# Main Text Statement

# %% [code]
def getStar(value):
    if value <= 0.0001:
        return "****"
    if 0.0001 < value <= 0.001:
        return "***"
    if 0.001 < value <= 0.01:
        return "**"
    if 0.01 < value <= 0.05:
        return "*"
    return "ns"

# %% [code]
bmetric = 'weighted_unifrac'
meta = prj_data['metadata'][(prj_data['metadata']['host_scientific_name'] != 'callithrix jacchus') & 
                            (prj_data['metadata']['sample_type'] == 'colon content')]
meta = meta.loc[list(set(meta.index) & set(res_all['rarefaction']['results'].columns))]
dm = res_all['beta_diversity']['results'][bmetric].to_data_frame()

# %% [markdown]
Wir sehen in den Daten sehr klar, dass der Genotyp bei gleichem maternalen Effekt einen signifikanten Unterschied im (Colon) Mikrobiom ausmacht - allerdings noch nicht in der P Generation

# %% [code]
for generation, g in meta.groupby('generation'):
    if generation == 'rm':
        continue
    print(generation)
    print("  alpha", getStar(kruskal(
        res_all['alpha_diversity']['results'].loc[g[g['genotype'] == 'c57bl6/j'].index, 'PD_whole_tree'],
        res_all['alpha_diversity']['results'].loc[g[g['genotype'] == 'balb/c'].index, 'PD_whole_tree']).pvalue))
    print("  beta", getStar(permanova(DistanceMatrix(dm.loc[g.index, g.index], ids=g.index), g['genotype'],)['p-value']))

# %% [markdown]
Die RM stammen von einem C57BL6/J Weibchen und einem BALB/c Männchen ab. Verglichen mit den 2*6 Düsseldorf/ZETT Kontrollen sind die RM Colon Mikrobiome den C57BL6/J ähnlicher (fast signifikant: Kruskal-Wallis H-test auf weighted UniFrac p=0.0534, beinahe konstante blaue Linie in Fig YB), was biologisch Sinn ergibt, da der Kontakt zur Mutter größer sein dürfte als zum Vater. 

# %% [code]
kruskal(
    dm.loc[meta[(meta['iacuc_institute'] == 'ZETT') & (meta['genotype'] == 'c57bl6/j') & (meta['geo_loc_name'] == 'Germany:Duesseldorf')].index, 
               meta[meta['generation'] == 'rm'].index].stack().values,
    dm.loc[meta[(meta['iacuc_institute'] == 'ZETT') & (meta['genotype'] == 'balb/c') & (meta['geo_loc_name'] == 'Germany:Duesseldorf')].index, 
               meta[meta['generation'] == 'rm'].index].stack().values)

# %% [code]
kruskal(
    dm.loc[meta[(meta['iacuc_institute'] == 'ZETT') & (meta['genotype'] == 'c57bl6/j') & (meta['geo_loc_name'] == 'France:Janvier')].index, 
               meta[meta['generation'] == 'rm'].index].stack().values,
    dm.loc[meta[(meta['iacuc_institute'] == 'ZETT') & (meta['genotype'] == 'balb/c') & (meta['geo_loc_name'] == 'France:Janvier')].index, 
               meta[meta['generation'] == 'rm'].index].stack().values)

# %% [code]
for geo_loc_name, g in meta[(meta['iacuc_institute'] == 'ZETT')].groupby('geo_loc_name'):
    print(geo_loc_name, kruskal(
        list(res_all['alpha_diversity']['results'].loc[g[g['genotype'] == 'c57bl6/j'].index, 'PD_whole_tree'].values), 
        list(res_all['alpha_diversity']['results'].loc[g[g['genotype'] == 'balb/c'].index, 'PD_whole_tree'].values)).pvalue)

# %% [code]
print("Janvier VS Duesseldorf", getStar(kruskal(
    list(res_all['alpha_diversity']['results'].loc[meta[(meta['iacuc_institute'] == 'ZETT') & (meta['geo_loc_name'] == 'France:Janvier')].index, 'PD_whole_tree'].values), 
    list(res_all['alpha_diversity']['results'].loc[meta[(meta['iacuc_institute'] == 'ZETT') & (meta['geo_loc_name'] == 'Germany:Duesseldorf')].index, 'PD_whole_tree'].values)).pvalue))

# %% [code]
# dist within RM individuals (living in same cage)
res_gut['cage_variance'][res_gut['cage_variance']['generation'] == 'rm']['dist'].mean()

# %% [code]
# p1 is sig diff from RM
idx_rm_p1 = meta[meta['generation'].isin(['rm','p1'])].index
print(permanova(DistanceMatrix(dm.loc[idx_rm_p1,idx_rm_p1], ids=idx_rm_p1), meta.loc[idx_rm_p1,'generation']))

kruskal(
    res_all['alpha_diversity']['results'].loc[meta[meta['generation'] == 'p1'].index, 'PD_whole_tree'], 
    res_all['alpha_diversity']['results'].loc[meta[meta['generation'] == 'rm'].index, 'PD_whole_tree'])

# %% [code]
6+16+5

# %% [markdown]
## for skin

# %% [code]
bmetric = 'unweighted_unifrac'
ametric = 'shannon'#'observed_features'#'shannon'#'PD_whole_tree'
meta = prj_data['metadata'][(prj_data['metadata']['host_scientific_name'] != 'callithrix jacchus') & 
                            (prj_data['metadata']['sample_type'] == 'skin of external ear')]
meta = meta.loc[list(set(meta.index) & set(res_all_skin['rarefaction']['results'].columns))]
dm = res_all_skin['beta_diversity']['results'][bmetric].to_data_frame()

# %% [code]
for generation, g in meta.groupby('generation'):
    if generation == 'rm':
        continue
    print(generation)
    print("  alpha", getStar(kruskal(
        res_all_skin['alpha_diversity']['results'].loc[g[g['genotype'] == 'c57bl6/j'].index, ametric],
        res_all_skin['alpha_diversity']['results'].loc[g[g['genotype'] == 'balb/c'].index, ametric]).pvalue))
    print("  beta", (permanova(DistanceMatrix(dm.loc[g.index, g.index], ids=g.index), g['genotype'],)['p-value']))

# %% [code]
for geo_loc_name, g in meta[(meta['iacuc_institute'] == 'ZETT')].groupby('geo_loc_name'):
    print(geo_loc_name, kruskal(
        list(res_all_skin['alpha_diversity']['results'].loc[g[g['genotype'] == 'c57bl6/j'].index, ametric].values), 
        list(res_all_skin['alpha_diversity']['results'].loc[g[g['genotype'] == 'balb/c'].index, ametric].values)).pvalue)

# %% [code]
print("Janvier VS Duesseldorf", getStar(kruskal(
    list(res_all_skin['alpha_diversity']['results'].loc[meta[(meta['iacuc_institute'] == 'ZETT') & (meta['geo_loc_name'] == 'France:Janvier')].index, ametric].values), 
    list(res_all_skin['alpha_diversity']['results'].loc[meta[(meta['iacuc_institute'] == 'ZETT') & (meta['geo_loc_name'] == 'Germany:Duesseldorf')].index, ametric].values)).pvalue))

# %% [code]
print("Controls VS breeding experiment", (kruskal(
    list(res_all_skin['alpha_diversity']['results'].loc[meta[(meta['iacuc_institute'] == 'ZETT')].index, ametric].values), 
    list(res_all_skin['alpha_diversity']['results'].loc[meta[(meta['iacuc_institute'] != 'ZETT')].index, ametric].values)).pvalue))

# %% [code]
# p1 is sig diff from RM
idx_rm_p1 = meta[meta['generation'].isin(['rm','p1'])].index
print(permanova(DistanceMatrix(dm.loc[idx_rm_p1,idx_rm_p1], ids=idx_rm_p1), meta.loc[idx_rm_p1,'generation']))

kruskal(
    res_all_skin['alpha_diversity']['results'].loc[meta[meta['generation'] == 'p1'].index, ametric], 
    res_all_skin['alpha_diversity']['results'].loc[meta[meta['generation'] == 'rm'].index, ametric])

# %% [markdown]
# Meta: Robertsen

# %% [code]
metaRob = pd.concat([prj_data['metadata'], pd.read_excel('../Robertsen/metadata_stefan.xlsx', dtype=str, index_col=0)])
metaRob['vendor'] = metaRob['vendor'].replace('Jackson Laboratory', 'mJAX').replace('Taconic Farms', 'mTAC')
metaRob

# %% [code]
# merge feature table
fp_counts_both = 'tmp_dir/counts_both.biom'
pandas2biom(fp_counts_both, pd.concat([
    biom2pandas(prj_data['paths']['deblur_table']),
    biom2pandas("../Robertsen/no_backup/deblur/deblur_res/reference-hit.biom")], axis=1).fillna(0))

# %% [code]
# construct shared insertion tree
seppres = project_sepp({'paths': {'deblur_table': fp_counts_both,
                                  'tmp_workdir': 'tmp_dir/'}})

# %% [code]
idx_control_samples = set(metaRob[metaRob['sample_type'].isin(['sterile water', 'control bedding', 'control food'])].index)
res_all_both = process_study(
    metadata=metaRob,
    control_samples=idx_control_samples,
    fp_deblur_biom=fp_counts_both,
    fp_insertiontree=seppres['paths']['insertion_tree'],
    rarefaction_depth=6000,
    fp_taxonomy_trained_classifier="/vol/jlab/MicrobiomeAnalyses/References/Q2-Naive_Bayes_classifiers/gg_2022_10_backbone_full_length.nb_2023.5.qza",
    tree_insert=seppres['insertion_tree'],
    is_v4_region=False,
    emperor_fp=prj_data['paths']['Generated/Emperor'],
    emperor_infix='_both_',
    dry=False,
    #use_grid=False, ppn=4,
    use_grid=True, 
    fix_zero_len_branches=False, # since this has already been corrected via the project_sepp call
    emperor_skip_tsne_umap=True,
)

# %% [code]
_ = rarefaction_curves(
    res_all_both['counts_plantsremoved'], 
    reference_tree=seppres['paths']['insertion_tree'], 
    control_sample_names=idx_control_samples, 
    fix_zero_len_branches=False, # since this has already been corrected via the project_sepp call
    sample_grouping=metaRob["qiita_study_id"], 
    dry=False)

# %% [code]
metaRob[(metaRob['host_scientific_name'] == "mus musculus") & 
        (metaRob['sample_type'] == "colon content") & 
        (metaRob['host_body_site'] == "uberon:colon")].fillna('?').groupby(['qiita_study_id','genotype','generation','host_body_site']).size()

# %% [code]
_ = emperor(metaRob[(metaRob['host_scientific_name'] == "mus musculus") & 
                    (metaRob['sample_type'] == "colon content") & 
                    (metaRob['host_body_site'] == "uberon:colon")], 
            res_all_both['beta_diversity']['results'], 
            fp_results=prj_data['paths']['Generated/Emperor'],
            infix='_bothColon_',
            dry=False, )

# %% [code]
p = plotTaxonomy(res_all_both['counts_plantsStillIn'], metaRob[metaRob['host_body_site'] == 'uberon:colon'].fillna('?'), file_taxonomy=res_all_both['taxonomy']['RDP']['results']['Taxon'],
                 group_l1='qiita_study_id', rank='Phylum', group_l2='generation',
                 print_sample_labels=False, minreadnr=1000)
p[0].set_size_inches((25, 5))

# %% [code]
dists = []
metric = 'unweighted_unifrac'
dm = res_all_both['beta_diversity']['results'][metric].to_data_frame()
for (genotype_benga, generation_benga), g_benga in metaRob[
                                       (metaRob['host_scientific_name'] == "mus musculus") & 
                                       (metaRob['sample_type'] == "colon content") & 
                                       (metaRob['host_body_site'] == "uberon:colon") & 
                                       (metaRob['qiita_study_id'] != 'Robertsen')].fillna({'generation': 'control'}).groupby(['genotype', 'generation']):
    if genotype_benga == "b6cf1":
        continue
    for (genotype_rob, generation_rob, vendor_rob, experiment_rob, sample_title_rob), g_rob in metaRob[
                                       (metaRob['host_scientific_name'] == "mus musculus") & 
                                       (metaRob['sample_type'] == "colon content") & 
                                       (metaRob['host_body_site'] == "uberon:colon") & 
                                       (metaRob['qiita_study_id'] == 'Robertsen') & 
                                       #(metaRob["experiment"].isin(["sentinels","littermates"])) & 
                                       (pd.isnull(metaRob["skipped_samples"]))].fillna({'sample_title':'?'}).groupby(['genotype', 'generation','vendor','experiment', 'sample_title']):
    #print()
        print(genotype_benga, generation_benga, g_benga.shape[0], genotype_rob, generation_rob, vendor_rob, experiment_rob, g_rob.shape[0])
        dm_sub = dm.loc[list(set(g_benga.index) & set(dm.index)),
                        list(set(g_rob.index)   & set(dm.index))]
        dm_sub = dm_sub.stack().reset_index()
        dm_sub['genotype_benga'] = genotype_benga
        dm_sub['generation_benga'] = generation_benga
        dm_sub['generation_rob'] = generation_rob
        dm_sub['vendor_rob'] = vendor_rob
        dm_sub['experiment_rob'] = experiment_rob
        dm_sub['sample_title'] = sample_title_rob
        dists.append(dm_sub)
dists = pd.concat(dists)

# %% [code]
nums = dict()
d1 = dists[dists['experiment_rob'].isin(['littermates','sentinels'])].copy()
d1['cat'] = 'genotype : ' + d1['genotype_benga']
#print(d1.shape[0], metaRob.loc[d1['level_0'].unique(),:].groupby('genotype').size(), metaRob.loc[d1['level_1'].unique(),:].groupby('vendor').size(), "\n--------")

d2 = dists[dists['experiment_rob'].isin(['littermates','sentinels']) &
           (dists['genotype_benga'] == 'c57bl6/j')].copy()
d2['cat'] = 'vendor : ' + d2['vendor_rob']
#print(d2.shape[0], metaRob.loc[d2['level_0'].unique(),:].groupby('genotype').size(), metaRob.loc[d2['level_1'].unique(),:].groupby('vendor').size(), "\n--------")

d3 = dists[dists['experiment_rob'].isin(['littermates','sentinels']) &
           (dists['genotype_benga'] == 'c57bl6/j') & 
           (dists['vendor_rob'] == 'mJAX')].copy()
d3['cat'] = 'generation : ' + d3['generation_rob']
#print(d3.shape[0], metaRob.loc[d3['level_0'].unique(),:].groupby('genotype').size(), metaRob.loc[d3['level_1'].unique(),:].groupby('vendor').size(), "\n--------")

d4 = dists[dists['experiment_rob'].isin(['littermates','sentinels']) &
           (dists['genotype_benga'] == 'c57bl6/j') & 
           (dists['vendor_rob'] == 'mJAX') & 
           (dists['generation_rob'] == 'f2')].copy()
d4['cat'] = 'breeding : ' + d4['generation_benga']
#print(d4.shape[0], metaRob.loc[d4['level_0'].unique(),:].groupby('genotype').size(), metaRob.loc[d4['level_1'].unique(),:].groupby('vendor').size(), "\n--------")

d5 = dists[(dists['experiment_rob'] == 'co-housing') & 
           (dists['genotype_benga'] == 'c57bl6/j')].copy()
d5['cat'] = 'co-housing : ' + d5['sample_title']

comb_dists = pd.concat([d1,d2,d3,d4,d5],axis=0)
comb_dists = comb_dists.rename(columns={0: metric})

nums = dict()
for c, g in comb_dists.groupby('cat'):
    nums[c] = pd.concat([metaRob.loc[g['level_0'].unique(),:].groupby('genotype').size(),
                         metaRob.loc[g['level_1'].unique(),:].groupby('vendor').size()])

comb_dists

# %% [code]
fig, axes = plt.subplots(1,1, figsize=(8/14*16,5)) #(8,5)

o_cat = [
    'genotype : c57bl6/j', 'genotype : balb/c',
    'vendor : mJAX', 'vendor : mTAC',
    'generation : p1', 'generation : f1', 'generation : f2',
    'breeding : p1', 'breeding : f1', 'breeding : f2', 'breeding : f3', 'breeding : f4', 'breeding : f5','breeding : control', 
    'co-housing : non-co-housed', 'co-housing : co-housed']
pal = {
    'genotype : c57bl6/j': colors['c57bl6/j'], 
    'genotype : balb/c': colors['balb/c'],
    'vendor : mJAX': colors['mJAX'],
    'vendor : mTAC': colors['mTAC'],
    'generation : p1': colors['mJAX'], 
    'generation : f1': '#fc0f0d', 
    'generation : f2': '#e16d1a',
    'breeding : p1': colors['p1'], 
    'breeding : f1': colors['f1'], 
    'breeding : f2': colors['f2'], 
    'breeding : f3': colors['f3'], 
    'breeding : f4': colors['f4'], 
    'breeding : f5': colors['f5'],
    'breeding : control': '#555555',
    'co-housing : co-housed': '#00ff00',
    'co-housing : non-co-housed': '#00ffff',
}
#sns.stripplot(data=comb_dists, y=metric, x='cat', ax=axes, order=o_cat, palette=pal, edgecolor='white', linewidth=1)
sns.boxplot(data=comb_dists, y=metric, x='cat', ax=axes, order=o_cat, palette=pal, zorder=10)

_ = axes.set_xticklabels(['%s\n%ii\n%ii\n%ii\n%ii' % (t.get_text(), 
                                       nums[t.get_text()].to_dict().get('c57bl6/j', 0),
                                       nums[t.get_text()].to_dict().get('balb/c', 0),
                                       nums[t.get_text()].to_dict().get('mJAX', 0),
                                       nums[t.get_text()].to_dict().get('mTAC', 0)) 
                          for t in axes.get_xticklabels()])

annotator = Annotator(axes, [
    ('genotype : c57bl6/j', 'genotype : balb/c'),
    
    ('vendor : mJAX', 'vendor : mTAC'),
    
    ('generation : p1', 'generation : f1'),
    ('generation : f1', 'generation : f2'),
    ('generation : p1', 'generation : f2'),
    
    ('breeding : p1', 'breeding : f1'),
    ('breeding : f1', 'breeding : f2'),
    ('breeding : f2', 'breeding : f3'),
    ('breeding : f3', 'breeding : f4'),
    
    ('co-housing : co-housed','co-housing : non-co-housed'),
    ('co-housing : co-housed','generation : f2'),
    
    #('breeding : f4', 'breeding : f5'),
], data=comb_dists, y=metric, x='cat', order=o_cat)
annotator.configure(test='Mann-Whitney', text_format='star', loc='outside', comparisons_correction="fdr_bh", correction_format="default")
annotator.apply_and_annotate()

axes.axhline(y=comb_dists.groupby('cat').mean(numeric_only=True).loc['generation : f2',metric], color='#e16d1a')

fig.savefig('Generated/robertsen.svg')

# %% [code]
fig, axes = plt.subplots(1,1)

sns.boxplot(data=dists, y=0, x='genotype_benga', ax=axes)

annotator = Annotator(axes, [
    ('balb/c', 'c57bl6/j')
], data=dists, y=0, x='genotype_benga')
annotator.configure(test='Mann-Whitney', text_format='star', loc='outside', comparisons_correction="fdr_bh", correction_format="default")
annotator.apply_and_annotate()

axes.text(-0.5, 0.6, 'Benga c57bl6/j are significantly more similar to Robertsen than Benga balb/c')

# %% [code]
fig, axes = plt.subplots(1,1)

sns.boxplot(data=dists.sort_values(by='generation_benga', key=key_generations), y=0, hue='genotype_benga', x='generation_benga', ax=axes)

annotator = Annotator(axes, [
    ((g, 'balb/c'), (g,'c57bl6/j')) for g in (dists['generation_benga'].unique())
] + [(('p1', 'c57bl6/j'), (g, 'c57bl6/j')) for g in (dists['generation_benga'].unique()) if g != 'p1'], data=dists.sort_values(by='generation_benga', key=key_generations), y=0, x='generation_benga', hue='genotype_benga')
annotator.configure(test='Mann-Whitney', text_format='star', loc='outside', comparisons_correction="fdr_bh", correction_format="default")
annotator.apply_and_annotate()

axes.legend(bbox_to_anchor=(1.1, 1.05))
axes.text(-0.3, 0.9, 'a) Benga genotypes are significantly different in all but p1, f5 generations (surprise: f2, f5 BALB are closer?!)\n'
                     'b) similarity grows over time (except f5)', fontsize=12)

# %% [code]
fig, axes = plt.subplots(1,1,figsize=(20, 6))

dists['g2'] = dists['genotype_benga'] + "_" + dists['vendor_rob']
hue_order = ['%s_%s' % (geno, time) for geno in ['c57bl6/j','balb/c'] for time in ['mJAX','mTAC']]
sns.boxplot(data=dists.sort_values(by='generation_benga', key=key_generations), y=0, hue='g2', x='generation_benga', ax=axes,hue_order=hue_order)
axes.legend(bbox_to_anchor=(1.1, 1.05))

annotator = Annotator(axes, [
    ((g, 'c57bl6/j_mJAX'), (g,'c57bl6/j_mTAC')) for g in dists['generation_benga'].unique()
] + [
    ((g, 'balb/c_mJAX'), (g,'balb/c_mTAC')) for g in dists['generation_benga'].unique()
], data=dists.sort_values(by='generation_benga', key=key_generations), y=0, x='generation_benga', hue='g2', hue_order=hue_order)
annotator.configure(test='Mann-Whitney', text_format='star', loc='outside', comparisons_correction="fdr_bh", correction_format="default")
annotator.apply_and_annotate()

axes.text(-0.3, 0.6, 'distance to mTAC is significantly higher than to mJAX in all generations', fontsize=12)

# %% [code]
fig, axes = plt.subplots(1,1,figsize=(20, 6))

dists['g1'] = dists['genotype_benga'] + "_" + dists['generation_rob']
d = dists[dists['vendor_rob'] == 'mJAX'].sort_values(by='generation_benga', key=key_generations)
sns.boxplot(data=d, y=0, hue='g1', x='generation_benga', ax=axes,
            hue_order=['%s_%s' % (geno, time) for geno in ['c57bl6/j','balb/c'] for time in ['p1','f1','f2']]
           )

axes.legend(bbox_to_anchor=(1.1, 1.05))
axes.set_title("only mJAX, but no mTAC")

# %% [code]
dists_rob = []
bodysite = "uberon:colon"
metric = 'weighted_unifrac'
st = "colon content"
if bodysite == 'uberon:feces':
    st = 'feces'
dm = res_all_both['beta_diversity']['results'][metric].to_data_frame()
num_samples = dict()#{'Littermates': {}, 'Co-housing': {}}
for generation_rob, g_rob in metaRob[(metaRob['host_scientific_name'] == "mus musculus") & 
                                   (metaRob['sample_type'] == st) & 
                                   (metaRob['host_body_site'] == bodysite) & 
                                   (metaRob['qiita_study_id'] == 'Robertsen') & 
                                   (metaRob["experiment"].isin(["sentinels","littermates"])) &
                                   (pd.isnull(metaRob["skipped_samples"]))].groupby('generation'):
    g_rob['vendor'] = g_rob['vendor'].replace('Jackson Laboratory', 'mJAX').replace('Taconic Farms', 'mTAC')
    #vendor_rob = vendor_rob.replace('Jackson Laboratory', 'mJAX').replace('Taconic Farms', 'mTAC')
    #print(genotype_rob, generation_rob, g_rob['vendor'].unique(), g_rob.shape[0])
    idx_jax = list(set(g_rob[g_rob['vendor'] == 'mJAX'].index) & set(dm.index))
    idx_tac = list(set(g_rob[g_rob['vendor'] == 'mTAC'].index) & set(dm.index))
    dm_sub = dm.loc[idx_jax, idx_tac]
    dm_sub = dm_sub.stack().reset_index().rename(columns={0: metric})    
    dm_sub['generation_rob'] = generation_rob
    dm_sub['cat'] = 'Littermates:\n' + generation_rob
    num_samples[('Littermates', generation_rob, 'mJAX')] = len(idx_jax)
    num_samples[('Littermates', generation_rob, 'mTAC')] = len(idx_tac)
    dists_rob.append(dm_sub)
dists_rob = pd.concat(dists_rob)

dists_rob_CH = []
for housing_rob, g_rob in metaRob[(metaRob['host_scientific_name'] == "mus musculus") & 
                                     (metaRob['sample_type'] == st) & 
                                     (metaRob['host_body_site'] == bodysite) & 
                                     (metaRob['qiita_study_id'] == 'Robertsen') & 
                                     (metaRob["experiment"].isin(["co-housing"]))].groupby('sample_title'):
    g_rob['vendor'] = g_rob['vendor']
    #vendor_rob = vendor_rob.replace('Jackson Laboratory', 'mJAX').replace('Taconic Farms', 'mTAC')
    #print(housing_rob, g_rob.shape[0], g_rob.groupby('vendor').size())
    idx_jax = list(set(g_rob[g_rob['vendor'] == 'mJAX'].index) & set(dm.index))
    idx_tac = list(set(g_rob[g_rob['vendor'] == 'mTAC'].index) & set(dm.index))
    dm_sub = dm.loc[idx_jax, idx_tac]
    dm_sub = dm_sub.stack().reset_index().rename(columns={0: metric})
    dm_sub['housing_rob'] = housing_rob
    dm_sub['cat'] = 'Co-housing:\n' + housing_rob
    num_samples[('Co-housing', housing_rob, 'mJAX')] = len(idx_jax)
    num_samples[('Co-housing', housing_rob, 'mTAC')] = len(idx_tac)
    dists_rob_CH.append(dm_sub)
dists_rob_CH = pd.concat(dists_rob_CH)

dists_rob_both = pd.concat([dists_rob, dists_rob_CH])

# %% [code]
fig, axes = plt.subplots(1,1, figsize=(4,5))
o = ['Co-housing:\nnon-co-housed', 'Co-housing:\nco-housed', 
     'Littermates:\np1', 'Littermates:\nf1', 'Littermates:\nf2']
sns.boxplot(data=dists_rob_both, y=metric, x='cat', order=o, ax=axes, palette={n: alpha_blending(c, 0.4) for (n,c) in zip(o, sns.color_palette())})
sns.stripplot(data=dists_rob_both, y=metric, x='cat', order=o, ax=axes, edgecolor='white', linewidth=1, palette={n: c for (n,c) in zip(o, sns.color_palette())})

axes.set_xticklabels(['%s\n%ii\n%ii\n%ii-i' % (
    t.get_text(), 
    num_samples[(t.get_text().split('\n')[0][:-1], t.get_text().split('\n')[1], 'mJAX')],
    num_samples[(t.get_text().split('\n')[0][:-1], t.get_text().split('\n')[1], 'mTAC')],
    dists_rob_both[dists_rob_both['cat'] == t.get_text()].shape[0],
    ) for t in axes.get_xticklabels()])

annotator = Annotator(axes, 
                      [('Co-housing:\nco-housed', 'Littermates:\nf2'),
                       ('Co-housing:\nco-housed', 'Co-housing:\nnon-co-housed'),
                       ('Littermates:\np1', 'Littermates:\nf1'),
                       ('Littermates:\np1', 'Littermates:\nf2'),
                       ('Littermates:\nf1', 'Littermates:\nf2')], 
                      data=dists_rob_both, y=metric, x='cat', order=o)
annotator.configure(test='Mann-Whitney', text_format='star', loc='outside', comparisons_correction="fdr_bh", correction_format="default")
annotator.apply_and_annotate()

fig.savefig('Generated/orig_robertsen.svg')

# %% [markdown]
In the pellet, the Bray-Curtis dis-
tances between the JAX-CH and the JAX
siblings and between the TAC-CH and the
TAC siblings were significantly lower (indi-
cating higher similarity) than comparison of the JAX-CH and
TAC-CH groups (Figure S2B). In the colon and ileum, the Bray-
Curtis distances were about equal between the co-housed
groups and their separately caged sibling controls.
![image-2.png](attachment:image-2.png)

I don't find any test on beta diversity distances in their paper that supports this statement: "In conclusion, our results establish that F2 littermates have
greater homogeneity in microbiota than co-housed mice"!!

# %% [markdown]
# Source tracking skin
using sources: 5 food samples, 5 bedding samples, all F2 C57BL6/J and BALB/c gut samples
sinks: all B57BL6/J and BALB/c skin samples

# %% [markdown]
## prototypic gut microbiome + food/bedding

# %% [code]
meta_st2 = prj_data['metadata'][(prj_data['metadata']['batch'] != '2') &
                                (prj_data['metadata']['host_scientific_name'] != 'callithrix jacchus') &
                                ((prj_data['metadata']['generation'].isin(['f2']) & 
                                 (prj_data['metadata']['sample_type'] == 'colon content')) | 
                                  prj_data['metadata']['sample_type'].isin(['control bedding','control food','skin of external ear']))
                               ].copy()
meta_st2.fillna('?').groupby(['host_scientific_name', 'sample_type','generation']).size()

# %% [code]
meta_st2['st2_type'] = 'Sink'
meta_st2.loc[meta_st2[meta_st2['sample_type'].isin(['control bedding', 'control food', 'colon content'])].index, 'st2_type'] = 'Source'
meta_st2['st2_env'] = meta_st2.apply(lambda r: r['genotype'] if pd.notnull(r['genotype']) else r['sample_type'], axis=1)
meta_st2.groupby(['st2_type', 'st2_env']).size()

# %% [code]
res_st2 = sourcetracker2(res_all['counts_plantsremoved'], meta_st2, col_envname='st2_env', col_type='st2_type', dry=False)

# %% [code]
p_st2 = plotTaxonomy(
    res_st2['results'], meta_st2, rank='raw', minreadnr=0, file_taxonomy=pd.Series(),
    group_l1='genotype', 
    fct_aggregate=np.mean,
    #print_sample_labels=True, sample_label_column='host_subject_id', no_sample_numbers=True,
    #colors={'Pax5+/-': colors_srtr['SPF@tp01@Pax5+/-'], 'WT': colors_srtr['SPF@tp01@WT'], 'Unknown': 'gray'}
)

# %% [code]
res_st2['results'].T.describe()

# %% [markdown]
## gut seeds skin?

# %% [code]
meta_st2_gs = prj_data['metadata'][#(prj_data['metadata']['batch'] != '2') &
                                (prj_data['metadata']['host_scientific_name'] != 'callithrix jacchus') &
                                 prj_data['metadata']['sample_type'].isin(['control bedding','control food','skin of external ear','colon content'])
                               ].copy().fillna({'generation': 'control'})
meta_st2_gs.groupby(['sample_type','generation']).size()

# %% [code]
cols = colors.copy()
cols.update({'control_bedding': 'magenta', 'control_food': 'cyan', 'Unknown': 'gray'})

def get_type(row):
    if row['Source'] == 'Unknown':
        return row['Source']
    elif row['Source'] == 'control_bedding':
        return "bedding"
    elif row['Source'] == 'control_food':
        return "food"
    elif row['Source'] == row['genotype']:
        return "same genotype"
    elif row['Source'] != row['genotype']:
        return "other genotype"

# %% [markdown]
### per generation, i.e. cross genotypes

# %% [code]
res_impact = []
res_st2_gs = dict()
for generation, g in meta_st2_gs[meta_st2_gs['sample_type'].isin(['colon content', 'skin of external ear'])].groupby('generation'):
    meta_gen = pd.concat([g, meta_st2_gs[meta_st2_gs['sample_type'].isin(['control bedding', 'control food'])]], axis=0)
    
    meta_gen['st2_type'] = 'Sink'
    meta_gen.loc[meta_gen[meta_gen['sample_type'].isin(['control bedding', 'control food', 'colon content'])].index, 'st2_type'] = 'Source'
    meta_gen['st2_env'] = meta_gen.apply(lambda r: r['genotype'] if pd.notnull(r['genotype']) else r['sample_type'], axis=1)
    print(generation, meta_gen.shape[0])
    res_st2_gs[generation] = sourcetracker2(res_all['counts_plantsremoved'], meta_gen, col_envname='st2_env', col_type='st2_type', dry=False, wait=False)
    
    if res_st2_gs[generation]['results'] is not None:
        p_st2 = plotTaxonomy(
            res_st2_gs[generation]['results'], meta_st2_gs, rank='raw', minreadnr=0, file_taxonomy=pd.Series(),
            group_l1='genotype', 
            fct_aggregate=np.mean, colors=cols,
            #print_sample_labels=True, sample_label_column='host_subject_id', no_sample_numbers=True,
            #colors={'Pax5+/-': colors_srtr['SPF@tp01@Pax5+/-'], 'WT': colors_srtr['SPF@tp01@WT'], 'Unknown': 'gray'}
        )
        p_st2[0].get_axes()[0].set_title('%s' % (generation))
        
        d = res_st2_gs[generation]['results'].stack().reset_index().rename(columns={0: 'impact'}).merge(prj_data['metadata'][['genotype','generation','geo_loc_name']], left_on='Sink', right_index=True, how='left')
        d['type'] = d.apply(get_type, axis=1)
        d['generation'] = generation
        
        res_impact.append(d)
res_impact = pd.concat(res_impact)

# %% [code]
res_impact.groupby('type').mean()

# %% [code]
fig, axes = plt.subplots(1,1,figsize=(15,5))
sns.boxplot(data=res_impact, y='impact', hue='type', x='generation', order=['rm', 'p1', 'f1', 'f2', 'f3', 'f4', 'f5', 'control'], ax=axes)
axes.legend(bbox_to_anchor=(1.1, 1.05))
axes.set_title("Source Tracker: does gut microbiome seed skin microbiome? Using matching gut generation data as source.")

# %% [markdown]
### per cage lineage

# %% [code]
res_st2_gs = dict()
for (generation, cage_line), g in meta_st2_gs[meta_st2_gs['sample_type'].isin(['colon content', 'skin of external ear'])].groupby(['generation','cage_line']):
    meta_gen = pd.concat([g, meta_st2_gs[meta_st2_gs['sample_type'].isin(['control bedding', 'control food'])]], axis=0)
    
    meta_gen['st2_type'] = 'Sink'
    meta_gen.loc[meta_gen[meta_gen['sample_type'].isin(['control bedding', 'control food', 'colon content'])].index, 'st2_type'] = 'Source'
    meta_gen['st2_env'] = meta_gen.apply(lambda r: r['genotype'] if pd.notnull(r['genotype']) else r['sample_type'], axis=1)
    print(generation, meta_gen.shape[0])
    res_st2_gs[(generation, cage_line)] = sourcetracker2(res_all['counts_plantsremoved'], meta_gen, col_envname='st2_env', col_type='st2_type', dry=False, wait=False)

# %% [code]
for (generation, cage_line) in res_st2_gs.keys():
    if res_st2_gs[(generation, cage_line)]['results'] is not None:
        print(generation, cage_line)
        p_st2 = plotTaxonomy(
            res_st2_gs[(generation, cage_line)]['results'], meta_st2_gs, rank='raw', minreadnr=0, file_taxonomy=pd.Series(),
            group_l1='genotype', 
            fct_aggregate=np.mean, colors=cols,
            #print_sample_labels=True, sample_label_column='host_subject_id', no_sample_numbers=True,
            #colors={'Pax5+/-': colors_srtr['SPF@tp01@Pax5+/-'], 'WT': colors_srtr['SPF@tp01@WT'], 'Unknown': 'gray'}
        )
        p_st2[0].get_axes()[0].set_title('%s %s' % (generation, cage_line))

# %% [code]
```
