## Additional File 2: Analysis repository for "The host genotype actively shapes its microbiome across generations in laboratory mice": README.md

## Computational Analysis for "The host genotype actively shapes its microbiome across generations."

Laurentiu Benga, Anna Rehm, Christina Gougoula, Philipp Westhoff, Thorsten Wachtmeister, W. Peter M. Benten, Eva Engelhardt, Andreas Weber, Karl Köhrer, Martin Sager and Stefan Janssen

#### Abstract
The microbiome greatly affects health and wellbeing. Evolutionarily, it seems unlikely that a host leaves microbial colonization of its offsprings to mere chance. Existing literature is inconclusive about two alternatively hypothesized mechanisms, active microbial shaping by host genetic factors or transmission of a microbial maternal legacy. To disentangle both factors we here obtained 2-cell stage embryos of two representative wildtypes, C57BL6/J and BALB/c, and transferred a mix of both genotype embryos into hybrid recipient mice to be inoculated by an identical microbiome at birth. Following the offspring along six generations clearly favors host genetic factors over maternal legacy in constant environments, like in murine laboratory experiments. Interestingly, only in the first offspring generation, maternal legacy dominated composition of the microbiome; reconciling the above dispute, as existing data in favor of maternal legacy did never extend beyond this first generation.

#### Instructions
You will find our analysis [jupyter notebook](https://jupyter.org/) in the file [benga_hostgenotype.ipynb](benga_hostgenotype.ipynb).

Raw sequence data can be retrieved from [Qiita](https://qiita.ucsd.edu/), study IDs 13422. For your convenience, the pre-processing results (i.e. feature-tables) are contained in `FromQiita` together with preparation and sample metadata information.
You might alternatively download sequences from ENA: ERPxxxxxx.

There is a bunch of additional software necessary to actually re-compute each and every analysis step. Most notabely
  - [qiime2](qiime2.org/) (version 2023.5)
  - [ggmap](https://github.com/sjanssen2/ggmap) (a collection of custom build helper scripts, use release [1.1.0](https://github.com/sjanssen2/ggmap/tree/1.1.0/ggmap))
Functions in ggmap will call several conda environments, whose build recipies are contained in ggmap sources.

Shoot your questions to

#### Figures
In principle, figures are generated via the notebook. However, some of the figures in the manuscript have manually been polished via Inkscape or other graphic software. Those manually touched versions are stored in the sub-directory [Figures](Figures) as \*.svg files.
