## Supplementary figures and images for "The host genotype actively shapes its microbiome across generations in laboratory mice"

### GA.jpg

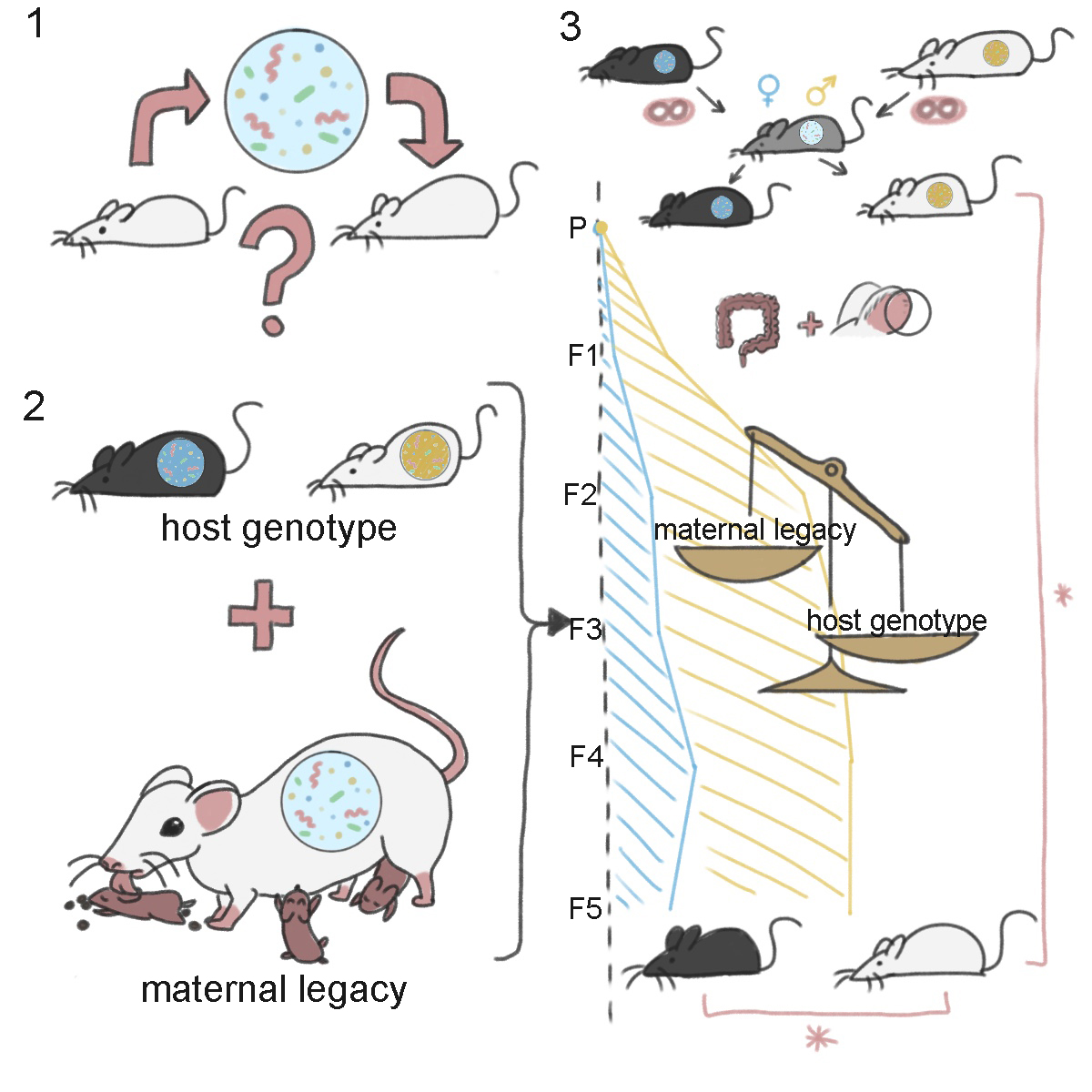
